# Commensal Yeast Promotes *Salmonella* Typhimurium Virulence

**DOI:** 10.1101/2024.08.08.606421

**Authors:** Kanchan Jaswal, Olivia A Todd, Roberto C Flores Audelo, William Santus, Saikat Paul, Manmeet Singh, Jian Miao, David M Underhill, Brian M Peters, Judith Behnsen

## Abstract

Enteric pathogens engage in complex interactions with the host and the resident microbiota to establish gut colonization. Although mechanistic interactions between enteric pathogens and bacterial commensals have been extensively studied, whether and how commensal fungi affect pathogenesis of enteric infections remains largely unknown. Here we show that colonization with the common human gut commensal fungus *Candida albicans* worsened infections with the enteric pathogen *Salmonella enterica* serovar Typhimurium. Presence of *C. albicans* in the mouse gut increased *Salmonella* cecum colonization and systemic dissemination. We investigated the underlying mechanism and found that *Salmonella* binds to *C. albicans* via Type 1 fimbriae and uses its Type 3 Secretion System (T3SS) to deliver effector proteins into *C. albicans*. A specific effector, SopB, was sufficient to manipulate *C. albicans* metabolism, triggering increased arginine biosynthesis in *C. albicans* and the release of millimolar amounts of arginine into the extracellular environment. The released arginine, in turn, induced T3SS expression in *Salmonella*, increasing its invasion of epithelial cells. *C. albicans* deficient in arginine production was unable to increase *Salmonella* virulence *in vitro* or *in vivo*. In addition to modulating pathogen invasion, arginine also directly influenced the host response to infection. Arginine-producing *C. albicans* dampened the inflammatory response during *Salmonella* infection, whereas *C. albicans* deficient in arginine production did not. Arginine supplementation in the absence of *C. albicans* increased the systemic spread of *Salmonella* and decreased the inflammatory response, phenocopying the presence of *C. albicans*. In summary, we identified *C. albicans* colonization as a susceptibility factor for disseminated *Salmonella* infection, and arginine as a central metabolite in the cross-kingdom interaction between fungi, bacteria, and host.

## Introduction

Decades of research have illuminated the central role of the gut microbiome for human health. Among a multitude of functions, gut microbes provide colonization resistance to pathogens ^1–6^, train the immune system ^7,8^, aid in digestion ^9,10^, and modulate distal organ functions via microbial products ^11–13^. Gut bacteria are the most abundant members of the gut microbiome and have thus been the focus of mechanistic research. Conversely, our knowledge on the roles of other members of the gut microbiome, such as viruses, fungi, and other microeukaryotes is still lacking. Their study has only recently received increased attention ^14–17^. New studies show that gut commensal fungi are a medically important part of the gut microbiota. Abundance and composition of the fungal microbiome is greatly altered in multiple gastrointestinal diseases ^18,19^, *e.g.* in inflammatory bowel disease (IBD) ^20^. However, it is largely unknown how fungi metabolically integrate into the gastrointestinal environment and interact with commensal and pathogenic bacteria ^14^, as these interactions have been only rarely studied in the gut environment ^21–26^.

To colonize the gut, enteric pathogens have been shown to exploit the bacterial microbiota ^2,27–30^. Some of the best studied pathogens in this regard are non-typhoidal *Salmonella* (NTS), which infect an estimated 100 million patients per year globally ^31^. In individuals with a healthy immune system, NTS like *Salmonella enterica* serovar Typhimurium (*Salmonella* or STm) cause a localized infection of the gastrointestinal tract, resulting in inflammatory diarrhea ^32,33^. In immunocompromised individuals, a patient group that encompasses children and the elderly, *Salmonella* can escape the gastrointestinal environment and disseminate to peripheral organs causing potentially fatal disease ^32,33^. To establish gut colonization, *Salmonella* must compete with resident microbes in the gut. Several studies now show that *Salmonella* uses a variety of commensal microbial products to boost gut colonization ^6,34–36^. However, these studies predominantly focus on gut resident bacteria while the potential contributions of the mycobiome have been largely excluded. Commensal fungi are found in all tested mammalian species ^37^, but their role during infection with enteric pathogens has been largely unexplored.

One of the most prominent fungal colonizers of human mucosal surfaces is *Candida albicans.* This yeast is a commensal of the human oral cavity, vagina, and gut, and represents the most frequently isolated fungus from human feces ^38^. Recent studies determined *C. albicans* to be present in more than 60% of healthy humans ^39,40^. While usually a commensal ^41,42^, *C. albicans* can become pathogenic, particularly in immunocompromised patients. *C. albicans* can escape the gastrointestinal tract to cause systemic candidiasis or expand in the vagina and oral mucosa to cause vulvovaginal candidiasis and oral thrush, respectively ^38,43,44^. An important virulence mechanism of *C. albicans* is the ability to switch morphology from rapidly growing yeast to epithelium-penetrating hyphae ^43^. *C. albicans* is associated with IBD, specifically Crohn’s Disease ^38,43,44^. While *C. albicans* cannot induce gut inflammation, it was shown to exacerbate it ^45,46^. *Salmonella* and *C. albicans* therefore both thrive under inflammatory conditions in the gut and both have high pathogenic potential. *C. albicans* represents an important human gut mycobiome member and is potentially present in the gut of a significant number of patients when they become infected with *Salmonella*. One clinical study underscores the potential importance of the inter-kingdom interaction of *Salmonella enterica* serovars and *C. albicans*.

Cross-sectional analysis of 2500 patients in Cameroon found a significantly increased recurrence of *S*. Typhi and *S*. Paratyphi infection when patients were colonized with *C. albicans*^47^. This study highlights the potential importance of *C. albicans* carriage in the modulation of *Salmonella* infection.

Our investigation on mechanistic interactions between *Salmonella* and *C. albicans* supports an important role for *C. albicans* carriage in the pathogenesis of *Salmonella* infection and delineates intricate fungal-bacterial-host crosstalk in the gut. Carriage of *C. albicans* might constitute a previously unappreciated susceptibility factor for disseminated *Salmonella* infections in susceptible populations.

## Results

### Presence of *C. albicans* increases *Salmonella* virulence

Inflammatory diseases, such as IBD, have often been associated with changes in abundance or composition of the fungal microbiome. We therefore wanted to test if infection with *Salmonella*, an enteric pathogen which causes acute gut inflammation, is associated with changes in the mycobiome. Using the streptomycin pre-treatment mouse model ^48^, we infected C57BL/6 mice with *Salmonella*, and sequenced the mycobiome of fecal samples collected before and after *Salmonella* infection (Fig. 1a). In feces from mice in two cages we observed a marked increase in the abundance of the order *Saccharomycetales* (Fig. 1b, red bars, Table S1), particularly in the genus *Candida* (Fig. S1a), after infection. *Candida* spp. were only present in two mice before infection but showed a high relative abundance in almost all cage mates after infection (Fig. 1b and S1a, red closed circles). Feces of mice from a third cage did not contain any *Candida* spp. before or after infection. Usage of an antibiotic likely opened a niche for *Candida* spp. that is usually occupied by bacteria. While expansion of *Candida* spp. and spreading to cage mates was therefore not unexpected, the serendipitous effect of this expansion on the outcome of *Salmonella* infection was. Mice in cages 2 and 3 with increased *Candida* spp. relative abundance showed markedly higher dissemination of *Salmonella* to the spleen and liver (Fig. 1c and S1b) than mice with no increase in *Candida* spp. in cage 1.

**Fig 1.**
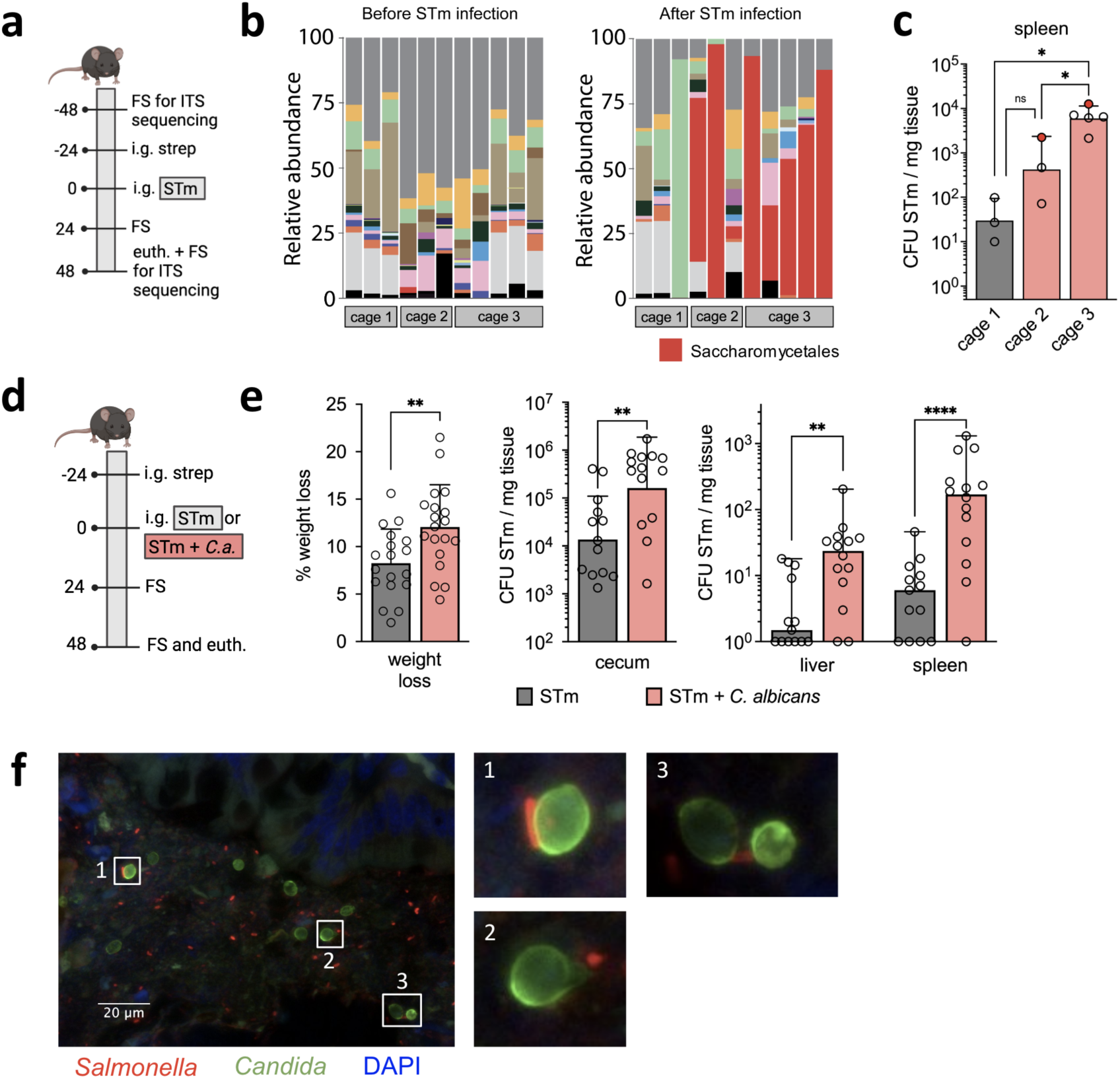
Candida albicans increases Salmonella colonization and dissemination. **a**, Schematic representation of experimental setup for sequencing analysis. FS, fecal sample; ITS, internally transcribed spacer; i.g., intragastrically; strep, streptomycin; STm, *Salmonella*; euth, euthanasia. **b**, Relative abundance of fungal genera identified with ITS sequencing in fecal samples of mouse before and after STm infection. n = 3 (Cage 1 and 2) and n = 5 (Cage 3). **c,** STm colonization in spleen 72h p.i. Mice with reads for *C. albicans* before infection are represented as red circles. Data are geometric mean ± s.d., n=3 (Cage 1 and 2) and n=5 (Cage 3). Ordinary one-way ANOVA for comparison. ns = not significant, * = p≤0.05. CFU, colony-forming units. **d**, Schematic representation of experimental setup for STm or STm and *C. albicans* ATCC infection. i.g., intragastrically; strep, streptomycin; FS, fecal sample; euth, euthanasia. **e,** Weight loss and STm colonization in C57BL/6 mice infected with STm or STm and *C. albicans* ATCC in the streptomycin pre-treatment model at 48h p.i. Data are geometric mean ± s.d. (for weight loss and cecum) and median (for liver and spleen) of 3 independent experiments, n ≥ 13. Significance determined by two-tailed Mann– Whitney test (for weight loss and cecum colonization) and mixed effect analysis with Šídák’s multiple comparisons test (for liver and spleen dissemination). ** = p ≤ 0.01, **** = p ≤ 0.0001. CFU, colony-forming units. **f,** Fluorescence image of lumen of colon during mouse infection with STm and *C. albicans* ATCC.

The species identified in mice was *Candida albicans*, which is a frequent commensal of humans but not a known commensal of mice. It is possible that the presence of *C. albicans* was an environmental contamination that bloomed in the presence of an antibiotic. Given that *C. albicans* is an important commensal in humans and seemed to be modulating *Salmonella* disease severity, we further analyzed this cross-domain interaction. We first determined whether the initial observation of higher *Salmonella* burden was indeed caused by presence of *C. albicans*. *Candida*-free mice received either *Salmonella* alone or *Salmonella* and *C. albicans* (10:1 ratio) (Fig. 1d). Colonization in the cecum and dissemination to peripheral sites was determined 48h p.i. (Fig. 1e and S1c,d). Co-infected mice lost 50% more weight and showed higher *Salmonella* colonization in the cecum than mice infected only with *Salmonella* at 48h post infection (Fig. 1e). Co-infected mice also had significantly increased *Salmonella* burden in the spleen and the liver compared to mice infected only with *Salmonella* (Fig. 1e). On the other hand, colonization of *C. albicans* was not significantly affected by presence of *Salmonella* and the yeast did not disseminate to peripheral organs (Fig. S1d). Imaging confirmed that *Salmonella* and *C. albicans* interact with each other in close proximity to cecum epithelial cells (Fig. 1f). In humans, *C. albicans* would be present in the gut as a commensal when they become infected with *Salmonella*. We therefore colonized CBA/J mice with the *C. albicans* strain 529L, which is known to colonize mice in the absence of antibiotics ^49^, and then infected the mice with *Salmonella*. Inflammation develops slower in this model, with peak inflammation reached at 9-10 days post infection. Also in this model, presence of *C. albicans* increased *Salmonella* colonization in the cecum. However, *Salmonella* dissemination to peripheral organs was not increased (Fig. S1e). Presence of two different *C. albicans* strains thus modulated *Salmonella* colonization or dissemination in two different mouse models.

### L-Arginine increases *Salmonella* virulence

*C. albicans* may modulate virulence of *Salmonella* through direct contact, secreted molecules, or altered immune response to *Salmonella* infection. We first tested whether presence of *C. albicans* led to higher gut epithelial cell invasion *in vitro*. We cultured *Salmonella* alone or with *C. albicans* for 2h and then infected two different colonic epithelial cell lines. Presence of *C. albicans* increased recovery of intracellular *Salmonella* to 180% compared to *Salmonella* alone (Fig. 2a and S2a,b). However, heat-killed *C. albicans* did not increase *Salmonella* invasion (Fig. 2a). Therefore, viable *C. albicans* is required to mediate increased *Salmonella* epithelial cell invasion. Consequently, purified cell wall components (curdlan, *i.e.* β-(1→3)-glucan) did not increase invasion of *Salmonella* (Fig. S2c). Imaging of the gut indicated that *C. albicans* and *Salmonella* directly interact with one another (Fig. 1f). To test if the two microbes indeed bind, we mixed *Salmonella* and *C. albicans* and determined sedimentation rates. As aggregates of yeast and bacteria sediment faster, increased sedimentation indicates binding. *Salmonella* bound to both live and heat-killed *C. albicans* yeast cells (Fig. 2b). Binding was additionally confirmed using fluorescence microscopy (Fig. 2c). We used a small library of *Salmonella* mutants to identify surface determinants that could mediate binding to *C. albicans* (Fig. S2d). A *Salmonella* mutant deficient in Type 1 Fimbriae (T1F), Δ*fim*, was unable to aggregate *C. albicans* (Fig. 2b,c and S2d). T1F binds to mannose residues ^50^, such as the ones present in the epithelial cell extracellular glycome or mannans present in the cell wall of yeast cells. As expected, binding of *Salmonella* to *C. albicans* was therefore inhibited by addition of excess mannose (Fig. 2b and S2e). To test if binding to *C. albicans* was required to increase *Salmonella* invasion, we measured invasion of *Salmonella* Δ*fim* into epithelial cells in the presence or absence of *C. albicans*. We also measured invasion of WT *Salmonella* in the presence of *C. albicans* and excess mannose. In either case we recovered equal numbers of *Salmonella*, indicating that *Salmonella* binding to *C. albicans* is required to increase *Salmonella* invasion into epithelial cells (Fig. 2d and S2f). Secreted factors could additionally modulate the interaction between *Salmonella* and *C. albicans*. We tested this by exposing *Salmonella* to cell-free supernatants of 2h single culture of *C. albicans* or co-cultures of *Salmonella* and *C. albicans*. In contrast to the mono-culture supernatant, co-culture supernatant was sufficient to increase *Salmonella* invasion to 160% (Fig. 2e), nearly the same level as exposure to live *C. albicans* (Fig. 2a). However, exposure to the supernatant of a co-culture of *C. albicans* and Δ*fim Salmonella* did not increase WT *Salmonella* invasion (Fig. 2e). Therefore, binding of *Salmonella* to *C. albicans* seemed to result in secreted molecules that alter *Salmonella* virulence. This hypothesis was supported by the observation that if *Salmonella* and *C. albicans* were directly added to epithelial cells without prior contact, *Salmonella* invasion was only marginally increased (Fig. S2g).

**Fig 2.**
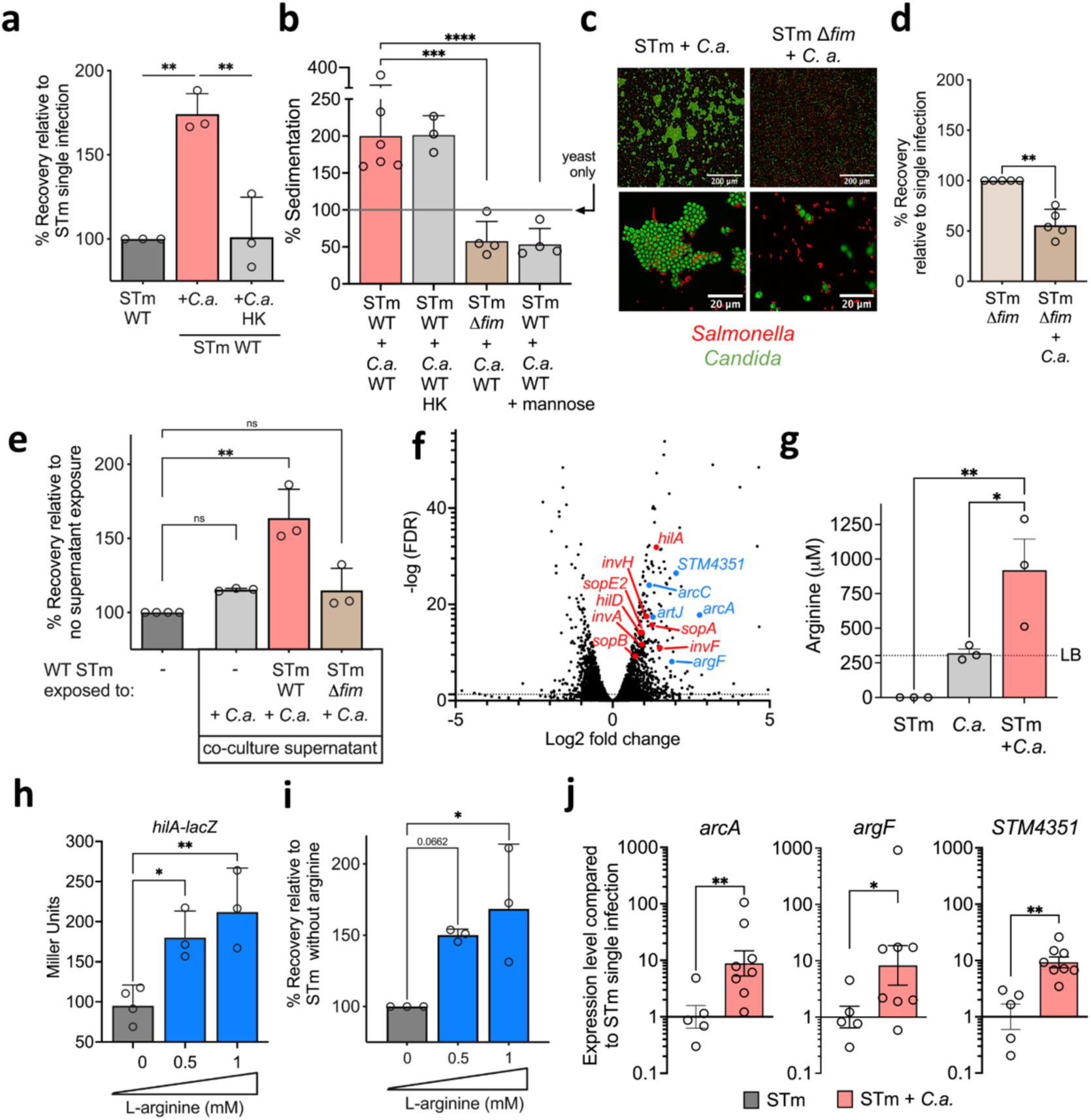
Binding to live Candida is required to increase Salmonella invasion. **a,** Invasion assay of STm infected (MOI:1) colonic epithelial cells (T84). STm either alone or with *C. albicans* ATCC in a 10:1 ratio were incubated for 2h prior to the assay. Data are geometric mean ± s.d., n = 3. Ordinary one-way ANOVA for comparison. ** = p ≤ 0.01. STm, *Salmonella*; HK, Heat-killed. **b,** Sedimentation assay of STm and *C. albicans* SC5314. Sedimentation of yeast single cultures is indicated as a line at 100%. Data are geometric mean ± s.d., n ≥ 3. Ordinary one-way ANOVA for comparison. *** = p ≤ 0.001, **** = p ≤ 0.0001. **c,** Fluorescence image of STm and *C. albicans* SC5314 *in vitro*. **d,** Invasion assay of STm infected (MOI:1) colonic epithelial cells (Caco2). STm either alone or with *C. albicans* ATCC in 10:1 ratio were incubated for 2h prior to the assay. Data are geometric mean ± s.d., n = 5. two-tailed Mann–Whitney test was used for comparison. ** = p ≤ 0.01. **e,** Invasion assay of STm infected (MOI:1) colonic epithelial cells (Caco2). STm either alone or with the supernatant from *C. albicans* ATCC containing cultures were incubated for 2h prior to the assay. Data are geometric mean ± s.d., n = 3. Ordinary one-way ANOVA for comparison. ns = not significant, ** = p ≤ 0.01. **f,** Volcano plot of RNAseq data showing STm genes differentially regulated in co-culture with *C. albicans* ATCC compared to STm alone. In red: genes involved in invasion and that are part of SPI-I. In blue: genes involved in arginine transport and downstream metabolism. **g,** Arginine levels measured in cell-free supernatants of STm, *C. albicans* SC5314 or STm and *C. albicans* SC5314 cultures incubated for 2h. Data are mean ± SEM., n = 3. Ordinary one-way ANOVA for comparison. * = p ≤ 0.05, ** = p ≤ 0.01. Dashed line indicates levels in LB. **h,** ß-gal assay performed on STm carrying chromosomal fusion of *lacZ* with the promoter of *hilA* after incubation with L-arginine for 2h. Data are geometric mean ± s.d., n ≥ 3. Ordinary one-way ANOVA for comparison. * = p ≤ 0.05, ** = p ≤ 0.01. **i,** Invasion assay of STm infected (MOI:1) colonic epithelial cells (Caco2). Data are geometric mean ± s.d., n = 3. Ordinary one-way ANOVA for comparison. * = p ≤ 0.05. **j,** qRT-PCR analysis of genes encoding for STm arginine import and metabolism from the cecal content of STm or STm and *C. albicans* ATCC infected mice 48h p.i. Data are mean ± SEM. of 2 independent exp, n ≥ 5. two-tailed Welch’s t-test for comparison. * = p ≤ 0.05, ** = p ≤ 0.01.

To determine *Salmonella* genes responsive to *C. albicans*, we compared transcriptomes of mono- and co-cultures (Table S2). KEGG analysis revealed significant regulation of multiple pathways in the presence of *C. albicans*, including *Salmonella* infection (Fig. S2h and Table S3). Incubation with *C. albicans* resulted in upregulation of *Salmonella* pathogenicity island (SPI)-1 regulator HilA and Type 3 Secretion System (T3SS) structural proteins and effectors (Fig. 2f and Table S2). We confirmed upregulation of *hilA* and *invA*, which encodes a SPI-1 T3SS structural protein (Fig. S2j,k). As SPI-1 is important for epithelial cell invasion, this finding could explain the increased invasion of *Salmonella* into colonic epithelial cells (Fig. 2a and S2a,b). Binding of *Salmonella* to *C. albicans* might directly result in increased SPI-1 gene expression. However, further analysis of RNAseq data suggested a more complex regulatory network. Genes involved in arginine uptake and metabolism represented some of the most highly upregulated genes in the *Salmonella* genome in the presence of *C. albicans* (Fig. 2f). In our KEGG analysis, arginine and proline metabolism was not significantly regulated (Fig. S2h and Table S2). However, the upregulated genes presented a specific subset of genes that coded for proteins involved in catabolism of arginine via the arginine deiminase pathway (ADI) (Fig. S2i). *arcA*, encoding the *Salmonella* arginine deiminase (and not the oxygen flux sensor with the same identifier) was upregulated 6.8-fold. We confirmed upregulation of *arcA, argF* (also annotated as *arcB*) and the arginine transporter subunit *STM4351* ^51^ by qRT-PCR using two different *Salmonella* and *C. albicans* strains (Fig. S2j,k). As arginine was previously shown to modulate virulence of pathogens, such as *Citrobacter rodentium* ^52^, we further investigated the role of arginine in the interaction of *Salmonella* and *C. albicans*. Expression of ADI pathway genes is known to be regulated by arginine availability ^53–55^. Metabolomic analysis of *in vitro* cultures indeed revealed significant differences in arginine concentrations. *Salmonella* depleted the arginine available in Luria Broth within 2h, whereas growth of *C. albicans* did not change the arginine concentration. During co-culture of *Salmonella* and *C. albicans*, arginine levels increased to 500-1300 μM, 2 to 4-fold the concentration of arginine in LB (Fig. 2g). Importantly, concentration of no other amino acid increased during co-culture (Fig. S3). We next tested if arginine can regulate SPI-1 expression in *Salmonella* in the absence of *C. albicans*. Indeed, gene expression of the SPI-1 regulator *hilA* increased in a dose dependent manner by addition of 0.5 mM and 1 mM L-arginine (Fig. 2h and S2l) and *invA* showed a trend for higher expression (Fig. S2l). To test if L-arginine increases invasion of *Salmonella* into epithelial cells, we incubated *Salmonella* with L-arginine for 2h and washed the bacteria before their addition to epithelial cells. Transient exposure to L-arginine increased invasion of *Salmonella* into epithelial cells in a dose-dependent manner (Fig. 2i and S2m). We next tested if the presence of *C. albicans* also increases arginine concentrations during *Salmonella* infections of mice. In our batch culture *in vitro*, *Salmonella* was the only consumer of arginine. In the gut, metabolites are in constant flux as the microbiome and the host are also metabolizing this amino acid. As observed for other metabolites utilized by *Salmonella in vivo*^56,57^, metabolomic analysis showed no significant change in arginine concentrations in cecum content (Fig. S4). We therefore assessed whether *Salmonella* was metabolizing arginine during co-infection and found that *Salmonella* expression of *arcA*, *argF*, and *STM4351* was 10-fold higher in the cecum of mice when *C. albicans* was present, compared to single infection without *C. albicans* (Fig. 2j). This indicates that *Salmonella* is metabolizing arginine in the presence of *C. albicans in vivo*.

### *C. albicans* produces arginine in the presence of *Salmonella*

We next examined the source of the increased arginine concentrations during *Salmonella* and *C. albicans* co-cultures *in vitro*. As *Salmonella* was catabolizing arginine in single cultures (Fig. 2g and S3), *C. albicans* remained the only possible producer. We therefore tested if the presence of *Salmonella* increased expression of genes required for arginine biosynthesis (Fig. S5a) in *C. albicans*. Indeed, expression of *ARG1* and *ARG4* was highly increased in the presence of *Salmonella* (Fig. 3a). We hypothesized that a *C. albicans* mutant deficient in the production of arginine would not increase *Salmonella* expression of SPI-1 genes and its invasion into epithelial cells. A *C. albicans arg4*Δ/Δ strain indeed failed to increase the expression of SPI-1 genes (Fig. S5b). The strain was still able to bind to *Salmonella* (Fig. S5c) but did not increase *Salmonella* invasion into epithelial cells, whereas the revertant strain expressing *ARG4* increased *Salmonella* invasion to the same extent as WT *C. albicans* (Fig. 3b). Arginine produced by *C. albicans* was thus required to increase *Salmonella* invasion into epithelial cells *in vitro*. We also tested whether the *C. albicans arg4*Δ/Δ strain would increase *Salmonella* virulence *in vivo*. To this end, we infected mice with *Salmonella* alone, with *Salmonella* and WT *C. albicans*, or with *Salmonella* and *C. albicans arg4*Δ/Δ (Fig. 3c). Cecum colonization and dissemination was determined at 48h p.i. (Fig. 3d,e and S5d,e). Both *C. albicans* strains colonized the small intestine, cecum, and colon/feces equally well (Fig. S5d). As previously observed (Fig. 1e), WT *C. albicans* increased *Salmonella* colonization in the cecum (Fig. 3d). However, *Salmonella* cecum colonization was not significantly increased in the presence of *arg4*Δ/Δ *C. albicans* (Fig. 3d). Similarly, WT *C. albicans* showed a trend toward higher *Salmonella* dissemination to the spleen (p=0.067), whereas *arg4*Δ/Δ *C. albicans* did not (Fig. 3e). Similar to co-infection with WT *C. albicans* and *Salmonella*, arginine concentrations *in vivo* did not change (Fig. S5f).

**Fig 3.**
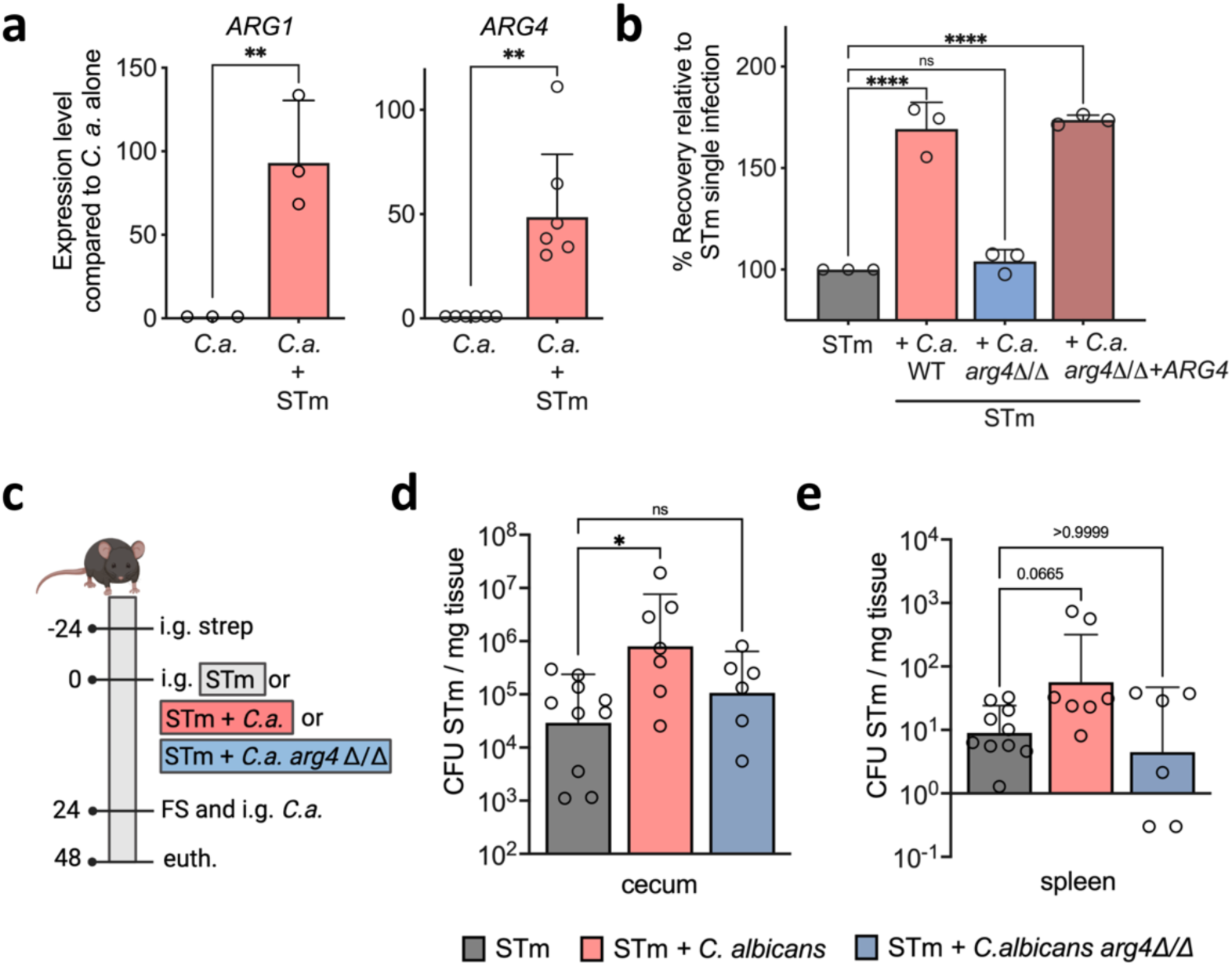
Role of arginine production in C. albicans for Salmonella virulence. **a,** qRT-PCR analysis of genes encoding for *C. albicans* arginine biosynthesis from the *C. albicans* SC5314 or STm and *C. albicans* SC5314 cultures incubated for 2h. Data are geometric mean ± s.d., n ≥ 3. Two-tailed Unpaired t-test for comparison. ** = p ≤ 0.01. STm, *Salmonella.* **b,** Invasion assay of STm infected (MOI:1) colonic epithelial cells (Caco2). STm either alone or with *C. albicans* SC5314 in a 10:1 ratio were incubated for 2h prior to the assay. Data are geometric mean ± s.d., n = 3. Ordinary one-way ANOVA for comparison. ns = not significant, **** = p≤0.0001. **c,** Schematic representation of experimental setup for STm or STm and *C. albicans* infection. i.g., intra-gastrically; strep, streptomycin; FS, fecal sample; euth, euthanasia. **d,** STm colonization in C57BL/6 mice infected with STm or STm and *C. albicans* SC5314 in the streptomycin pre-treatment model for 48h p.i. Data are geometric mean ± s.d. of 2 independent experiments, n ≥ 6. Kruskal-Wallis test for comparison. ns = not significant, * = p≤0.05. CFU, colony-forming units.

### *Salmonella* uses T3SS-1 effector SopB to trigger arginine production

*C. albicans* production of arginine was essential to increase *Salmonella* invasion into epithelial cells *in vitro* and *in vivo*. However, it was unclear why *C. albicans* would biosynthesize and release the amino acid when in contact with *Salmonella*. We hypothesized that *Salmonella* directly triggered this response in *C. albicans*. As a eukaryote, *C. albicans* shares many similarities with the mammalian cells *Salmonella* has evolved to interact with, and *Salmonella* T3SS effectors are functional when exogenously expressed in yeast (*e.g.*, *Saccharomyces cerevisiae*) ^58^. We therefore tested whether the T3SS-1 that is required for *Salmonella* invasion into epithelial cells was also required to induce the expression of arginine biosynthesis genes in *C. albicans*. We incubated a *Salmonella* Δ*invA* strain, deficient in the assembly of the T3SS-1 needle, with *C. albicans* and tested whether the cell-free supernatant would increase WT *Salmonella* invasion into epithelial cells. This supernatant did not increase invasion of *Salmonella* (Fig. 4a). We therefore measured the concentration of arginine and found that only co-cultures of *C. albicans* with WT *Salmonella,* but not with Δ*invA Salmonella,* had increased levels of arginine (Fig. 4b). Arginine biosynthesis genes *ARG1* (Fig. 4c) and *ARG4* (Fig. S6a) showed high expression in the presence of WT *Salmonella*, modest expression with the *Salmonella* Δ*fim* mutant that cannot aggregate with *C. albicans,* and no increase in the presence of *Salmonella* Δ*invA,* which was able to aggregate with *C. albicans* (Fig S2d). Therefore, the *Salmonella* T3SS was required to increase arginine biosynthesis in *C. albicans*. The *Salmonella* T3SS-1 can secrete many different effector proteins. To determine which effector might elicit the increase in arginine biosynthesis, we tested a mutant deficient in many effectors of T3SS-1 (*sipA, sopA, sopB, sopD,* and *sopE2*) and it failed to increase arginine biosynthesis in *C. albicans* (Fig. 4d and S6b). Further tests with *Salmonella* mutants pinpointed that deletion of a single effector, *sopB*, was sufficient to abrogate induction of *C. albicans* arginine biosynthesis (Fig. 4d and S6b). *Salmonella* Δ*sopB* shows no defect in binding to *C. albicans*, but the co-culture supernatant did not increase invasion of *Salmonella* and did not contain measurable arginine levels (Fig. 4e,f and S6c). *C. albicans* also did not increase invasion of a *Salmonella* strain deficient in SopB into epithelial cells (Fig. 4g). All our experiments were performed with the *Salmonella* strain IR715 (Nal^R^ derivative of ATCC 14028), but we also used the *Salmonella* SL1344 strain to confirm these findings (Fig. S6d-f). All tested *Salmonella* SL1344 strains showed similar binding to *C. albicans* (Fig. S6f). However, the function of SopB was redundant in SL1344, as the *Salmonella* SL1344 Δ*sopB* strain induced the same level of expression of *ARG1* and *ARG4* in *C. albicans* compared to the WT *Salmonella*. When we deleted both *sopB* and *sopE*, which was shown in mammalian systems to have overlapping functions with SopB ^59^, the strain failed to induce arginine biosynthesis (Fig. S6d). Overexpression of *sopB* in the complemented strain resulted in 3x higher expression of *ARG1* and *ARG4 in C. albicans* compared to WT (Fig. S6e). A functional *Salmonella* T3SS is therefore required to increase arginine biosynthesis in *C. albicans* and SopB (and SopE) are the translocated effectors that can trigger this response. We also expressed SopB under control of the doxycycline-repressible *tetO* promoter in *C. albicans* (Fig. S6g). Upon removal of doxycycline (Fig. S6g), *C. albicans ARG1* and *ARG4* genes were induced (Fig. 4h and S6h), confirming that presence of SopB in *C. albicans* results in upregulation of arginine biosynthesis.

**Fig 4.**
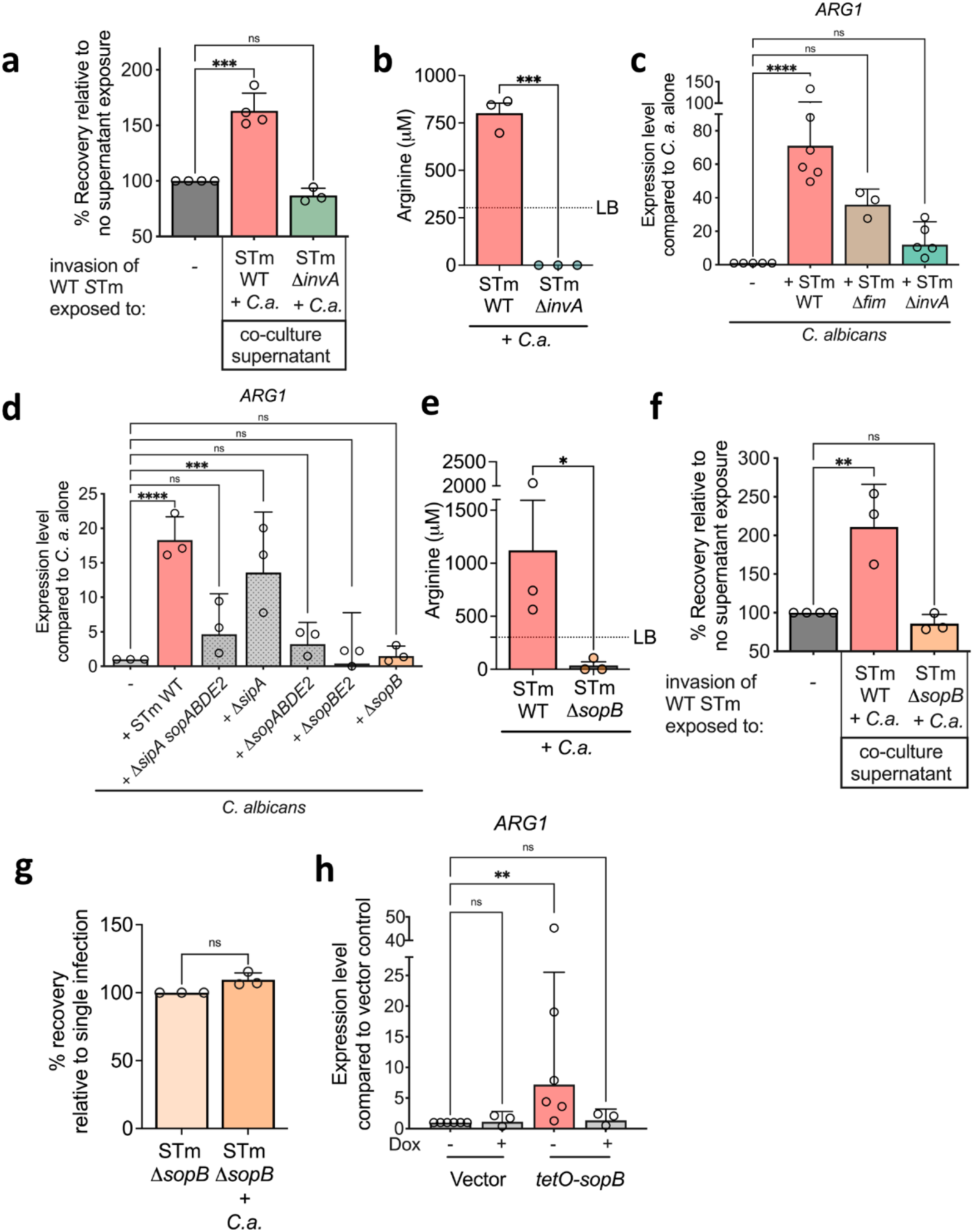
Salmonella uses T3SS-1 effector SopB to trigger arginine production in C. albicans. **a,** Invasion assay of STm infected (MOI:1) colonic epithelial cells (Caco2). STm either alone or with the supernatant from *C. albicans* ATCC containing cultures were incubated for 2h prior to the assay. Data are geometric mean ± s.d., n ≥ 3. Ordinary one-way ANOVA for comparison. ns = not significant, *** = p≤0.001. STm, *Salmonella.* **b,** Arginine levels measured in cell-free supernatants of STm and *C. albicans* SC5314 cultures incubated for 2h. Data are mean ± SEM., n = 3. Two-tailed unpaired t-test for comparison. *** = p≤0.001. Dashed line indicates levels in LB. **c & d,** qRT-PCR analysis of genes encoding for *C. albicans* arginine biosynthesis from the *C. albicans* SC5314 or STm and *C. albicans* SC5314 cultures incubated for 2h. Data are geometric mean ± s.d., n ≥ 3. Kruskal-Wallis test (for **c**) and Ordinary one-way ANOVA (for **d**) for comparison. ns = not significant, *** = p≤0.001, **** = p≤0.0001. **e,** Arginine levels measured in cell-free supernatants of STm and *C. albicans* SC5314 cultures incubated for 2h. Data are mean ± SEM., n = 3. One-tailed unpaired t-test for comparison. * = p≤0.05. Dashed line indicates levels in LB. **f,** Invasion assay of STm infected (MOI:1) colonic epithelial cells (Caco2). STm either alone or with the supernatant from *C. albicans* SC5314 containing cultures were incubated for 2h prior to the assay. Data are geometric mean ± s.d., n ≥ 3. Ordinary one-way ANOVA for comparison. ns = not significant, ** = p≤0.01. **g,** Invasion assay of STm infected (MOI:1) colonic epithelial cells (Caco2). STm either alone or with *C. albicans* SC5314 in a 10:1 ratio were incubated for 2h prior to the assay. Data are geometric mean ± s.d., n = 5. Two-tailed paired t-test test for comparison. ns = not significant. **h,** qRT-PCR analysis of genes encoding for *C. albicans* arginine biosynthesis from *C. albicans* SC5314 cultures expressing vector or *tetO*-*sopB* incubated for 6h with or without doxycycline. Data are geometric mean ± s.d., n ≥ 3. Kruskal-Wallis test for comparison. ns = not significant, ** = p ≤ 0.01.

### Blunted immune response to *Salmonella* and *C. albicans* co-infection

In addition to direct microbial interactions between *C. albicans* and *Salmonella*, we also found evidence that presence of *C. albicans* modulates the host immune response to *Salmonella* infection. In the gastroenteritis mouse model (Fig. 5a), infection with *Salmonella* resulted in a strong inflammatory response in the cecum (Fig. 5b and S7a,b). We expected an opportunistic pathogen like *C. albicans*, which is known to induce IL-17 during candidiasis ^60–62^, to further increase the inflammatory response. However, presence of *C. albicans* during *Salmonella* infection resulted in a blunted inflammatory response (Fig. 5b and S7a,b). The decreased response was particularly pronounced early after infection at 24h (Fig. 5b and S7a) when *Salmonella* has not yet disseminated to the liver and the spleen, and colonization of the cecum was equal between groups (Fig. S7c). However, expression of genes such as *Tnfa*, *Il17*, *Nos2,* and *Cxcl1* was 5 to 10-fold lower after infection with *Salmonella* and *C. albicans* compared to infection with *Salmonella* alone (Fig. 5b), with an increase in Arginase II (*Arg2*) expression (Fig. S7d). Reduced expression of *Cxcl1* at 24h post co-infection likely resulted in significantly reduced neutrophil infiltration at 48h post co-infection (Fig. S7e) and slightly reduced pathology score (Fig. S7f).

**Fig 5.**
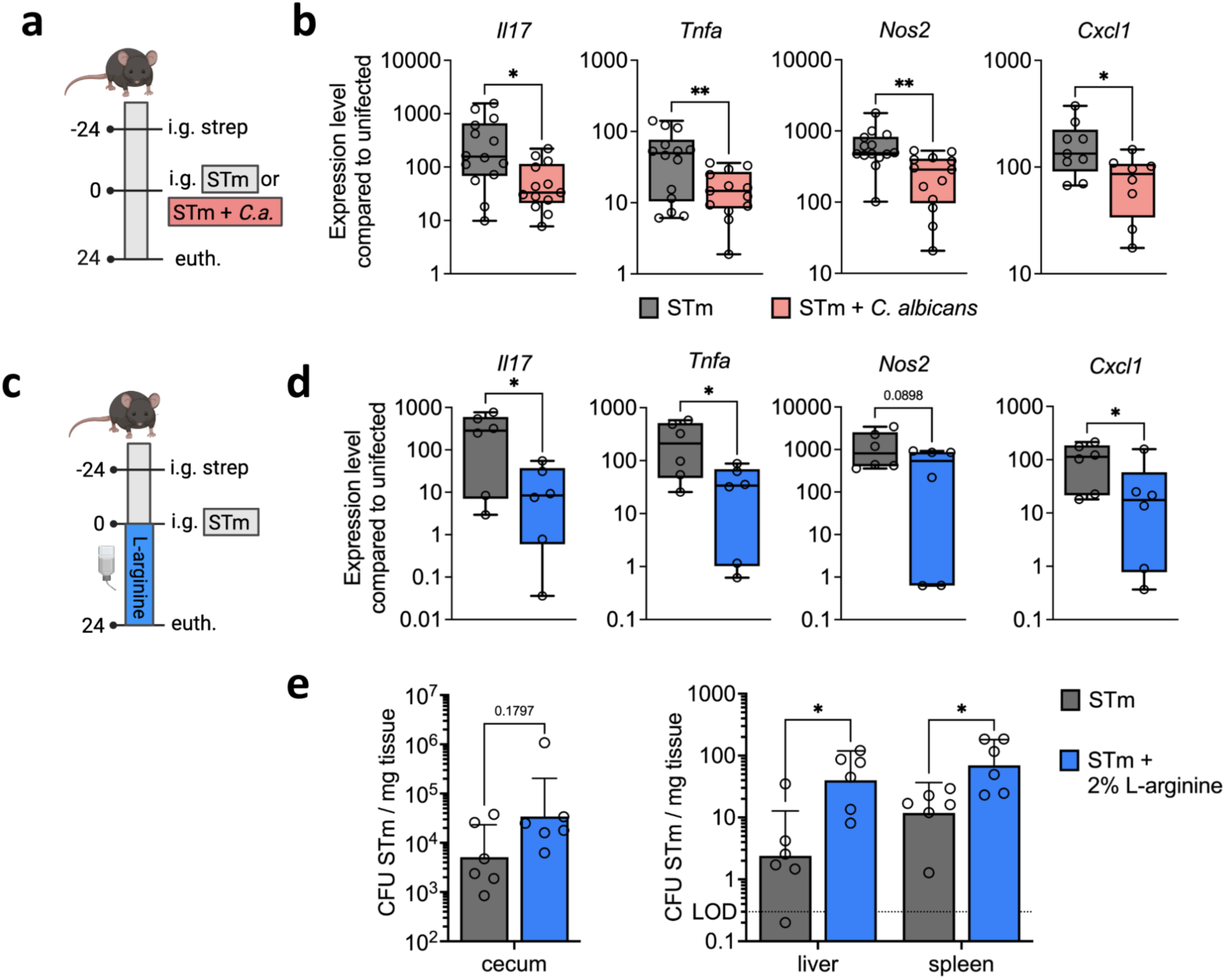
Immune response to Salmonella and C. albicans co-infection. **a,** Schematic representation of experimental setup for STm or STm and *C. albicans* infection. i.g., intra-gastrically; strep, streptomycin; STm, *Salmonella*; euth, euthanasia. **b**, qRT-PCR analysis of genes encoding for host inflammatory response from the cecum tissue of STm or STm and *C. albicans* ATCC infected mice 24h p.i. Data is from 3 independent experiments, n ≥ 8. Data are presented as box and whiskers plot from 25th to 75th percentile, median, minimum and maximum. Significance determined by two-tailed Unpaired t-test. * = p≤0.05, ** = p≤0.01. **c,** Schematic representation of experimental setup for STm infection. 2% L-arginine was supplemented in the drinking water. i.g., intra-gastrically; strep, streptomycin; euth, euthanasia. **d,** qRT-PCR analysis of genes encoding for host inflammatory response from the cecum tissue of STm or STm with L-arginine treated mice 24h p.i. Data is from 2 independent experiments, n = 6. Data are presented as box and whiskers plot from 25th to 75th percentile, median, minimum and maximum. Significance determined by two-tailed unpaired t-test. * = p≤0.05. **e,** STm colonization in C57BL/6 mice infected with STm or STm treated with 2% L-arginine in the streptomycin pre-treatment model for 48h p.i. Data are geometric mean ± s.d. of 2 independent experiments, n ≥ 6. Mann-Whitney test for comparison. ns = not significant, * = p≤0.05. CFU, colony-forming units.

Results of our *in vitro* studies allowed us to mechanistically link the consequence of contact between *Salmonella* and *C. albicans* to this initially unexpected reduced inflammatory response. *In vitro*, *C. albicans* produced arginine in the presence of *Salmonella*. While *in vivo* metabolomic analysis of cecum content did not indicate an overall increased availability of arginine, *Salmonella* gene expression indicated that there are likely locally increased levels of arginine (Fig. 2j and S4). Arginine was previously shown to exert anti-inflammatory effects ^63–66^ and supplementation of arginine resulted in reduced inflammation after *Salmonella* infection in broiler chickens ^67^. We therefore tested if supplementation of L-arginine in the gastrointestinal environment (Fig. 5c) would phenocopy the presence of *C. albicans*. Indeed, addition of 2% L-arginine to the drinking water (pH-adjusted to 7) resulted in a reduced inflammatory response (Fig. 5d, S8a,b), a trend toward higher *Salmonella* cecum colonization, and significantly increased *Salmonella* dissemination to spleen and liver (Fig. 5e). With L-arginine supplementation, induction of *Tnfa*, *Il17*, *Nos2*, and *Cxcl1* was significantly lower at 24h post infection with *Salmonella* (Fig 5d). L-arginine in the drinking water did not change immune gene expression in uninfected mice (Table S4). Exogenous supply of arginine therefore reduced the inflammatory response to *Salmonella* infection and increased *Salmonella* systemic dissemination similar to coinfection with *C. albicans*. We next tested if the reduced inflammatory response during infection with *Salmonella* and *C. albicans* was dependent on the ability of *C. albicans* to produce arginine. Indeed, in mice infected with *Salmonella* and *arg4*Δ/Δ *C. albicans* (Fig. 3c), inflammatory genes were expressed significantly higher compared to mice infected with *Salmonella* and WT *C. albicans* (Fig. S7b)*. Ifng* and *Il22* expression during co-infection with *arg4*Δ/Δ *C. albicans* was higher than with WT *C. albicans* and similar or higher compared to single infection with *Salmonella*. Many other genes (*e.g.*, *Il17* and *Tnfa*) also showed a trend toward higher expression, similar to infection with *Salmonella* alone. However, expression of *Cxcl1* and *Nos2* was not dependent on the ability of *C. albicans* to produce arginine (Fig. S7b). While clearly not the only factor relevant to this cross-kingdom interaction, arginine plays a crucial role to increase the virulence of *Salmonella* in the presence of *C. albicans*.

## Discussion

Our study delineates intricate molecular interactions that govern an inter-kingdom interaction in the gut between *Salmonella*, *C. albicans*, and the host (Fig. S9). The body of literature elucidating the role of fungi in disease has been steadily increasing but remains largely correlative. Molecular details are known only for a few interactions, such as *Staphylococcus aureus* and *C. albicans* in extraintestinal sites^22,68^, or in the oral mucosa, where *Enterococcus faecalis* inhibits *C. albicans* morphogenesis by secreting the peptide EntV ^69–71^. Our present study combines analysis of metabolic cross-feeding, T3SS usage against a non-mammalian organism, and host immune response to uncover the complex interactions between fungi, bacteria, and the host.

The amino acid arginine is emerging as a key metabolite in the gut. Its role as a substrate for the mammalian inducible nitric oxide synthase (iNOS) and arginases has been well established. However, an increasing body of literature indicates that arginine availability and metabolism also directly influence colonization resistance to pathogens and microbial pathogenesis. *C. rodentium* upregulates virulence genes in the gut in the presence of arginine ^52^. On the other hand, lack of arginine due to *Enterococcus faecalis* catabolism serves as a metabolic cue for *Clostridium difficile* to increase virulence gene expression ^72^. In our study, *Salmonella* used its T3SS to trigger arginine production by *C. albicans* in the gut, which in turn increased *Salmonella* virulence. Contrary to *E. faecalis*, *Salmonella* did not metabolize arginine to ornithine and export it, as indicated by a lack of ornithine in the culture medium. However, *Salmonella* might convert arginine to polyamines as the expression of spermidine biosynthesis genes was increased (Table S2). Arginine availability in the host is rate limiting for the generation of reactive nitrogen species and some pathogens, such as *Helicobacter pylori*, *Mycobacterium tuberculosis, Streptococcus pneumonia,* and *Pseudomonas* spp. deplete arginine to reduce its availability to iNOS ^73–76^. During *Salmonella* infection, *C. albicans* arginine production or exogenously supplied L-arginine were anti-inflammatory and detrimental to an adequate host response, resulting in more severe disease.

In prior decades, *Salmonella* effector functions were studied in the yeast *Saccharomyces cerevisiae* and the functions elucidated in this model were confirmed in mammalian cells ^77–79^. However, our study shows that yeast cells respond to SopB differently as compared to mammalian cells. SopB increased arginine biosynthesis, a phenotype absent in mammalian cells. Divergent evolution of a key protein with both enzymatic and scaffolding activity, Arg82p, likely explains this differing phenotype ^80,81^. SopB is an inositol phosphatase with catalytic activity for inositol 1,3,4,5,6-pentakisphosphate ^59,82,83^. Arg82p is a kinase that catalyzes the reverse reaction. Arg82p has a secondary function as an essential scaffolding protein of the arginine repressor complex that represses arginine biosynthetic genes ^84,85^. The mechanism by which Arg82p responds and switches between these activities has been extensively studied but is still not completely understood. The common substrate between Arg82 and SopB makes it likely that SopB is impacting arginine biosynthesis either by feedback regulation or making Arg82p physically unavailable to stabilize the arginine repressor complex. Detailed studies to address this hypothesis are ongoing in our laboratories.

Our study shows effector activity of SopB in the cytoplasm of *C. albicans*. Native translocation of SopB into *C. albicans* using the T3SS has also been shown once before ^86^. Considering the thickness of the *C. albicans* cell wall and the length of the T3SS needle, such activity should be physically impossible. However, other microbes are known to secrete fungal cell wall degrading enzymes ^87–90^. *Salmonella* secretes chitinases ^91–93^ that might result in local changes of the fungal cell wall that enable T3SS deployment.

Currently, the prevalence of *C. albicans* during *Salmonella* Typhimurium infections of human patients is unknown. *C. albicans* is not part of diagnostic panels used to differentiate the etiology of gastroenteritis. However, particularly in at-risk patients receiving antibiotic therapy that would allow fungal expansion, presence of *C. albicans* in the gut could be an important unappreciated risk factor. While infection rates with *Salmonella* Typhimurium in the United States remain high, most cases remain undiagnosed, rendering clinical studies unfeasible. A study in Cameroon found that the recurrence of *Salmonella enterica* serovar Typhi and Paratyphi infections increased 4-fold when patients were colonized with *Candida* spp ^47^. While this indicated an importance of *Candida* spp. carriage for *Salmonella* disease susceptibility, further studies are warranted to examine correlation of *Candida* carriage and severity of disease and the role of *C. albicans* during *Salmonella* Typhimurium infection. In vulnerable populations, administration of an antimycotic might represent a novel approach to limiting sepsis risk after *Salmonella* infection.

Our study identifies *C. albicans* as a susceptibility factor for *Salmonella* infection and arginine as a key metabolic cue in the interaction of *Salmonella*, *C. albicans*, and the host.

## Acknowledgements

This work was supported by grants from the National Institute of Health to J.B. (R01AI143641 and R01AI175223), to B.M.P (R01AI177615, R01DK113788, and R01AI134796), and by startup funds from the Department of Microbiology and Immunology at the University of Illinois Chicago. Olivia Todd was supported by the Training Program in Lung Biology and Pathobiology T32HL007829. We want to express our gratitude to the following scientists who generously shared their microbial strains: Andreas Baumler provided the *Salmonella* Δ*fim* mutant and T3SS-1 effector deletions, Leigh Knodler provided the plasmid pFPV-*mCherry*, *Salmonella* mutants and complemented strains in background strain SL1344, and James Slaugh provided the *hilA-lacZ* fusion *Salmonella* strain. Vector CIpSATtetTrans was a kind gift from Dr. Steve Saville (Univ. Texas at San Antonio). RNA-seq was performed by the UIC Genomic Research Core. Bioinformatic analysis was performed by the UIC Research Informatics Core, supported in part by NCATS through Grant UL1TR002003. Amino acid analysis was performed by the Microbial Culture & Metabolomics Core of the PennCHOP Microbiome Program and the Center for Molecular Studies in Digestive and Liver Diseases (NIH P30DK050306). Histopathology was performed by the UIC Research Histology core. We want to thank all lab members for critical reading of the manuscript. Figures were created with BioRender.com.

## Author Contributions

J.B. conceived and administered the study and was the primary supervisor. K.J. additionally conceived and supervised parts of the study and was responsible for execution and analysis of the majority of the experimental work. O.A.T. performed additional *in vitro* experiments, aided in mouse experiments, and performed extended colonization mouse experiments. R.F.A. established binding assays and performed *in vitro* microscopy. W.S. performed initial mouse experiments and *in vivo* imaging with gnotobiotic mice. K.J., O.A.T., W.S., R.F.A., and J.B. were involved in the husbandry of germ-free and gnotobiotic mice. D.M.U was responsible for microbiome sequencing and analysis. S.P. generated all *C. albicans* mutant strains for this study, except for SC5314 GFPy, which was generated by J.M. B.M.P. provided conceptual insight and supervised *C. albicans* mutant generation. J.B. and K.J. wrote the original draft of the manuscript. J.B. and K.J. revised the draft with input from all authors.

## Materials & Correspondence

Correspondence and material requests should be addressed to Judith Behnsen. All strains and materials used in this study are available upon request

## Data availability

Raw sequence reads have been deposited at NCBI Sequence Read Archive under project PRJNA1143068, and the UNITE (v.8.2) database^94^ was used for all microbiome analysis.

## Code availability

No custom code has been used.

## Materials and Methods

### Bacterial and fungal culture conditions

Fungal strains, bacterial strains, and plasmids used in this study are listed in Supplementary Table S5 and S6. *Salmonella enterica* serovar Typhimurium strains were grown at 37°C in LB broth (per liter: 10 g tryptone, 5 g yeast extract and 10 g sodium chloride). *Candida albicans* strains were grown at 37°C in YPD media (per liter: 10 g yeast extract, 20 g Peptone and 20 g Dextrose). Antibiotics and antifungals were added at the following concentrations (mg l^−1^) to LB-agar or YPD-agar, as needed: carbenicillin/ampicillin, 100; chloramphenicol, 30; doxycycline, 1 or 10; amphotericin, 2.5; nourseothricin (NAT), 200 or 25; hygromycin B (HygB), 600 or 75. All strains were cultured aerobically shaking at 200 rpm overnight except for *in vitro* experiments where *Salmonella* strains were grown overnight in liquid LB without shaking at 37°C. Co-incubation of *S.* Typhimurium strains and *C. albicans* strains were done by quantifying cell number by measuring the OD_600_ of the cultures. 1x10^9^ cells of *Salmonella* strains and 1x10^8^ cells of *C. albicans* strains were centrifuged and resuspended in 1 ml LB and co-incubated for 2h in 37°C without shaking, unless indicated otherwise. The cell-free supernatant from the 2h co-incubations was filter-sterilized and used to co-incubate with *Salmonella* to analyze the effect of secreted factors. When indicated, 1x10^8^ cells of *C. albicans* were heat-killed (HK) at 100°C for 1h, before co-incubation with *Salmonella*.

### Animal models

*SPF mice:* All animal experiments were reviewed and approved by the Institutional Animal Care and Use Committee at the University of Illinois Chicago in protocols 17-045, 20-016 and 22-192 and were in agreement with ethical regulations. Eight-to-nine-week-old C57BL/6J female WT mice (strain number 000664) and CBAJ female WT mice (strain number 000656) were obtained from Jackson Laboratories, maximum barrier, and were 9–10 weeks of age by the start of every experiment. Mice were housed in the Biologic Resources Laboratory facility at the University of Illinois Chicago in individual cages with filter tops (maximum occupancy, five mice) containing corn cob bedding and paper nesting material. The room the mice were housed in was kept under 14 h:10 h light/dark cycles, 70–76 °F and 30– 70% humidity. Mice had access to food and autoclaved water *ad libitum*. Mice were fed chow LM485. For infections with C57BL/6J mice, mice were pre-treated with streptomycin (0.1 ml of a 200 mg ml^-1^ solution in sterile water) intra-gastrically 24 h before inoculation with *Salmonella* or *C. albicans* (WT) or *Salmonella* and *C. albicans* (WT or *arg4*Δ/Δ). For initial experiments, 1×10^9^ colony-forming units (CFU) ml^-1^ of *Salmonella* and 1×10^8^ CFU ml^-1^ of *C. albicans* were used (Fig 1a-e, S1, 5a-b, S7a, S7c-f). The same dose, however, yielded significantly higher colonization and dissemination of *Salmonella* in single infected mice in subsequent experiments. We therefore maintained the same ratio of *Salmonella* to *C. albicans* but decreased the overall dose to 1×10^7^ CFU ml^-1^ of *Salmonella* and 1×10^6^ CFU ml^-1^ of *C. albicans.* This dose yielded *Salmonella* cecum colonization and dissemination levels similar to previous experiments (Fig 2j, S4, 3c-e, S5d,e, 5d, 5e, S7b, S8). We also observed that some mice were losing *C. albicans* in the cecum, so we re-colonized the mice with *C. albicans* at 24h p.i. (Fig. 3c,d and S5d,e) and excluded the mice if *C. albicans* didn’t colonize the cecum. When required, mice were treated with 2% L-arginine (pH adjusted to ∼7) in their drinking water *ad libitum*. For infections with CBA/J mice, mice were colonized i.g. with 1×10^8^ CFU ml^-1^ of *C. albicans* 529L or PBS control 3 and 1 days prior to and, to maintain equal colonization levels across mice, 4 days post i.g. infection with 1×10^9^ CFU ml^-1^ of *Salmonella*. Weights were monitored daily for both mouse models. Fecal samples were collected daily, and serial tenfold dilutions were plated for enumerating bacterial CFU on LB agar plates supplemented with carbenicillin and amphotericin or YPD agar plates with chloramphenicol. Mice were sacrificed at time points indicated. Cecum colonization and dissemination was assessed by homogenization and plating of cecum, spleen, liver, Peyer’s patches and mesenteric lymph nodes, respectively. Colon content was collected to enumerate *Salmonella* colonization and was flash frozen for microbiota analysis. Cecum luminal content was also collected, and flash frozen to either isolate RNA or perform metabolomic analysis. Cecum tissue was collected by flash freezing to analyze murine inflammatory gene expression and by fixing it with formalin for histological examination. No statistical methods were used to pre-determine sample sizes, but our sample sizes are similar to those reported in previous publications ^95–97^.

*Gnotobiotic mice:* Swiss Webster (Tac:SW) germ-free WT mice were purchased from Taconic and bred at the BRL facility at the University of Illinois Chicago in a room with 14 h:10 h light/dark cycles and 70–76 °F and 30–70% humidity. Mice were kept in isolators purchased from Park Bioservices LLC. Mice were fed autoclaved 5L79 chow and autoclaved super Q water *ad libitum*. Germ-free condition was tested at least once a month with aerobic liquid cultures (brain heart infusion, BHI), solid cultures (blood agar plates, Thermo Sci Remel, R01200), fungal cultures (Sabouraud slants), anaerobic liquid (BHI) and solid cultures (Brucella agar, Thermo Sci Remel, R01254) from isolators’ swabs, fecal samples and fungal traps placed inside the isolators. Fecal samples were also tested with Gram staining and qPCR to detect bacterial DNA.

For ASF experiments, mice were stably colonized with ASF purchased from Taconic Biosciences. Stable colonization was assessed via species-specific PCR ^98^. For *Salmonella* infection experiments, 10–15-week-old ASF-colonized Swiss Webster female mice were housed in a biosafety cabinet. Mice were then orally infected with 0.1 ml of 1×10^7^ CFU ml^-1^ of *Salmonella* or 1×10^7^ CFU ml^-1^ of *Salmonella* and 1×10^6^ CFU ml^-1^ of *C. albicans*. Fecal samples were collected, and serial tenfold dilutions were plated for enumerating bacterial CFU on LB agar plates supplemented with carbenicillin and amphotericin or YPD agar plates with chloramphenicol. Mice were euthanized at 6h and 24h and colon samples were collected for immunofluorescence.

### Microbiota and mycobiota sequencing and analysis

ITS1 sequencing of mouse fecal samples was performed largely as previously described ^19,24^. Briefly, for DNA isolation, one to two mouse fecal pellets were resuspended in 0.5 ml lyticase buffer (50 mM Tris, 1 mM EDTA and 0.2% 2-mercaptoethanol), homogenized briefly and treated with 200 U lyticase (Sigma-Aldrich) for 30 min at 30 °C. Material was pelleted and resuspended in 0.4 ml Stool DNA Stabilizer (B Bridge International), mixed with 0.1 ml 0.1 mm silica beads (Biospec) and 0.3 ml 0.5 mm beads (Biospec), heated at 95 °C for 5 min and subjected to bead beating twice on high (VWR) for 1 min. DNA was then further purified using the QIAmp DNA mini kit (Qiagen) according to the manufacturer’s instructions.

Primers used in this study are detailed in Table S7. Fungal ITS1 amplicons were generated using primers with the following recombinant DNA targeting sequences: ITS1 F (5′- CTTGGTCATTTAGAGGAAGTAA), ITS2 R (5′-GCTGCGTTCTTCATCGATGC). ITS1 amplicons were generated with 35 cycles using Invitrogen AccuPrime PCR reagents at an annealing temperature of 48 °C. Amplicons were then used in the second PCR reaction, using Illumina Nextera XT v2 barcoded primers to uniquely index each sample, and 2 × 300 paired-end sequencing was performed on the Illumina MiSeq.

ITS1 sequences were trimmed using ITSxpress (v1.7.4) ^99^ in QIIME 2 (v2019.7). Using the DADA2 package (v1.10.1) in R (v3.5.2), reads underwent further quality filtering as error rates were calculated and removed from the dereplicated reads. Where forward and reverse reads could be merged, they were, and where they could not, largely owing to ITS1 sequences sometimes being longer than sequence coverage, they were concatenated. An initial sequence table was constructed before chimaeras were identified using the removeBimeraDenovo function. Finally, taxonomy was assigned using DADA2’s native naïve Bayesian classifier against the UNITE (v.8.2) database ^94^.

Statistical analyses were performed with R 4.0.2 ^100^. Figures were produced using the packages ggplot2 ^101^, dplyr ^102^, ape ^103^ and RColorBrewer ^104^. Microbial communities were further analyzed using the microbiome ^105^ and phyloseq ^106^ packages.

### Fluorescence microscopy

*In vivo*: Tissue samples were fixed in 10% formalin for 24h, stored in 70% ethanol and finally embedded in paraffin and sectioned at 7μm with a microtome.

Deparaffinization was performed by immersing the sections in decreasing concentration of EtOH (100%, 90%, 70%). For antigen retrieval, the deparaffinized slides were immersed in 10mM Na-Citrate Buffer and heated in a microwave for 20 minutes. Blocking was performed by flooding the slides with PBS-10% FBS for 30 minutes at 4°C. *Salmonella* was stained with *Salmonella* antisera O-4 (Hardy Diagnostics, 1:100 in PBS) for 1h at room temperature.

Following washing with PBS-0.3% Tween20, slides were incubated for 30’ at room temperature with Alexa Fluor 594 goat anti-rabbit antibody (Invitrogen, 1:1000 in PBS). Slides were washed with PBS-0.3% Tween20 and *C. albicans* was stained using a FITC-labeled rabbit anti-fungal antibody (Meridian Life Science, 1:250 in PBS-10% FBS) for 1h at 37°C. Slides were washed with PBS-0.3% Tween20, mounted using ProLong Diamond Antifade Mounting with DAPI (Invitrogen) and images were taken using the BZ-X710 All-in-One Fluorescence Microscope at 60x.

*In vitro*: OD600 of overnight *Salmonella-*mCherry and *C. albicans-*GFP was measured. 4x10^9^ STm and 7.6x10^7^ *C. albicans* were centrifuged and resuspended in 1 and 2 ml of PBS, respectively. On a microscope slide, 20 µl of *Salmonella* and 20 µl of *C. albicans* were added to the slide and mixed by swirling the slide for 1 minute. 5 µl of the mixture was added to an agarose pad (0.75 g of agarose in 50 ml of H2O). Images were taken using the BZ-X710 All-in- One Fluorescence Microscope at 4x-40x.

### Invasion (Gentamicin protection) assay

The T84 colonic epithelial cell line (ATCC Cat# CCL- 248; RRID:CVCL_0555) or C2BBe1 colonic epithelial cell line (Caco2; ATCC Cat# CRL-2102; RRID:CVCL_1096) were seeded onto a 24-well tissue-culture treated plate at a density of 5x10^5^ cells/well with media lacking antibiotics/antimycotics and incubated overnight (for T84) and for 5 days (for Caco-2) at 37°C. *Salmonella* strains alone or co-incubated with either *C. albicans* strains or the cell-free supernatant were centrifuged, resuspended in DMEM/F12, and serially diluted. Epithelial cells were infected with 5x10^5^ cells of *Salmonella* (multiplicity of infection (MOI) = 1) and 5x10^4^ cells of *C. albicans* (MOI=0.1). Infected cells were incubated at 37°C for 1 h. The inoculum was serially diluted and plated on LB agar with amphotericin to confirm bacterial numbers. After infection, the media was removed via vacuum, and wells were washed 3 times with 500 μl phosphate-buffered saline (PBS). 500 μl of DMEM/F12 + 10% FBS + 0.1 mg ml^-1^ gentamicin was added to the wells and incubated at 37°C for 1 h to kill extracellular bacteria. After incubation, the wells were washed with PBS and lysed by incubation with 1% Triton X-100 for 5 mins. Cells were disrupted and harvested by scraping wells and pipetting, were serially diluted, and plated on LB agar with amphotericin to quantify bacterial cells that invaded. The percentage of cells recovered relative to the inoculum was calculated.

### Sedimentation assay

OD_600_ of overnight *Salmonella* and *C. albicans* cultures was measured. 4x10^9^ *Salmonella* and 7.6x10^7^ *C. albicans* were centrifuged and resuspended in 1 and 2 ml of PBS, respectively. In a 15 ml conical tube, 1 ml of *Salmonella* and 2 ml *C. albicans* were added and vortexed for 15 seconds before taking 200 μl from the top of the mixture, which was used to measure OD_600_ (1:5 dilution) for the initial starting value. For a baseline control, 2 ml *C. albicans* was mixed with 1 ml of PBS and OD_600_ was measured. 15 ml conical tubes were then incubated at 37°C for 20 minutes without shaking and after that time, OD_600_ was measured the same way as before. To calculate % sedimentation, the equation ((OD_0min_ – OD_20min_))/OD_0min_) x 100 was used. The value of *Salmonella* and *C. albicans* was divided over the *C. albicans* alone to measure sedimentation.

### RNA extraction

*In vivo*: Cecum tissue collected from mice 24 hpi and 48 hpi were homogenized by mortar and pestle, using liquid nitrogen. Because mice were treated with streptomycin 24 h prior to infection, we collected cecum from uninfected mice after streptomycin treatment as a control. The homogenate was transferred to 1 ml of Tri-Reagent (Molecular Research Center) for RNA extraction. RNA was extracted with 0.1 ml of bromo-3-chloropropane, centrifuged, and the upper phase was precipitated with 0.5 ml isopropanol. After centrifugation, pellets were washed twice with 1 ml of 75% ethanol in RNase-free water. The RNA pellet was then resuspended in RNase-free water. For cecum content, RNA was extracted from snap-frozen luminal cecal content samples collected during mouse experiments. RNA extraction was performed using the Qiagen RNeasy Power Microbiome kit. RNA was treated with DNase using the Turbo DNA-free kit (Invitrogen). *In vitro*: RNA extraction was performed on the cell pellet from mono and co-cultures of *Salmonella* and *C. albicans* after 2h of co-incubation, except to assess the expression of *sopB* in *C. albicans* in the *C. albicans tetO*-*sopB* strain. Here, transformants containing CIptet-SopB or CIpSATtetTranstADH1 (empty vector) were grown overnight in liquid YPD media containing 10 μg ml^-1^ doxycycline. Cells (1x10^5^ ml^-1^) were transferred into 5 ml YPD with or without 10 μg ml^-1^ doxycycline and incubated for 8 h at 30°C with continuous shaking. In all cases, RNA extraction was performed using the hot-phenol method as described previously ^107,108^ followed by DNase treatment using the Turbo DNA-free kit (Thermo Fisher).

### RNA sequencing and analysis

*Basic processing of RNAseq data*: Raw reads were aligned to the reference assembly (GCF_000022165.1) using BWA-MEM v 0.7.17. ^109^. Expression levels of gene features, i.e. CDS regions from the reference assembly, were quantitated using FeatureCounts v2.0.3 as raw read counts of the stranded libraries ^110^. *Differential analysis*: Differential analysis of quantitated gene features as compared with treatment was performed using the software package edgeR on raw sequence counts ^111^. Prior to analysis, the data were subsampled to a maximum depth of 1,750,000 counts per sample and filtered to remove any features that had less than 100 total counts summed across all samples. Data were normalized as counts per million and an additional normalization factor was computed using the TMM algorithm. Statistical tests were performed using the “exactTest” function in edgeR. [GC1] Adjusted p values (q values) were calculated using the Benjamini-Hochberg false discovery rate (FDR) correction ^112^. Significant gene features were determined based on an FDR threshold of 5% (0.05). *Enrichment analysis*: The enrichment or overrepresentation of differentially expressed gene features in the various gene groups, i.e. pathways, modules and BRITE categories, listed for the KEGG organism “seo” (KEGG genome T01714) was determined using Fisher’s Exact test in R. Briefly, a list of differentially expressed gene features was obtained from the results of the differential analysis based on a q value, i.e. FDR corrected p value, less than 0.05. The enrichment of significantly different gene features as compared with all genes listed in KEGG for the organism “seo” were then tested for the KEGG organism pathways, modules and BRITE level 1, 2, and 3 categories. Adjusted p values (q values) were calculated using the Benjamini-Hochberg false discovery rate (FDR) correction ^112^. Significant enrichment of gene groups was determined based on an FDR threshold of 5% (0.05).

### Quantitative Real-Time PCR (qRT-PCR)

Reverse transcription was performed with the High-Capacity cDNA Reverse Transcription Kit (Applied Biosystems). For *Salmonella* RNA, reactions were also performed without the addition of reverse transcriptase to confirm that there was no amplification of DNA in qRT-PCR reactions. 500 ng of RNA was used for the reverse transcription reaction. The reverse transcription cycle consisted of 10 minutes at 25°C followed by 120 minutes at 37°C and 5 minutes at 85°C. qRT-PCR was performed using the Fast SYBR Green Master Mix (Applied Biosystems) on the Viia7 Real-time PCR system at the Genome Research core at the University of Illinois at Chicago. The qRT-PCR reaction cycle consisted of 20 seconds at 95°C followed by 40 cycles of 3 seconds at 95°C and 30 seconds at 60°C. Reactions were performed in duplicate.

For transcript levels of *sopB* in doxycycline-repressible strains, samples were normalized to 1 µg RNA and treated with Turbo DNase (Thermo Fisher). First-strand cDNA synthesis was performed using the RevertAid RT kit following the manufacturer’s protocol. Amplification of ∼20 ng cDNA was performed using 2X Maxima SYBR Green/ROX qPCR Master Mix (Thermo Fisher) and gene-specific primers for *sopB* (CaSopBDETF + CaSopBDETR). qRT-PCR reactions were monitored and analyzed with a Bio-Rad CFX96 Real-Time System and software.

Relative expression was calculated based on the ΔCT values obtained by subtracting the CT value of the house-keeping gene with the gene of interest. *gapA*, *ActB*, and *ACT1* were used as house-keeping genes to normalize *Salmonella*, murine immune, and *C. albicans* gene expression, respectively.

### Metabolomics analyses

Amino acids were quantified using a Waters Acquity uPLC System with an AccQ-Tag Ultra C18 1.7 μm 2.1x100mm column and a Photodiode Detector Array.

Fecal samples were homogenized in methanol (5 μl/mg stool) and centrifuged twice at 13,000g for 5 minutes. Intestinal flushes were vortexed for 1 minute, and centrifuged twice at 13,000g for 5 minutes. Amino acids in the supernatant were derivatized using the Waters AccQ-Tag Ultra Amino Acid Derivatization Kit (Waters Corporation, Milford, MA) and analyzed using the UPLC AAA H-Class Application Kit (Waters Corporation, Milford, MA) according to manufacturer’s instructions. Blanks and standards were run every eight samples. All chemicals and reagents used were mass spectrometry grade.

### β-galactosidase assay

β-gal assays were performed as detailed in ^113^. Promoter activity was measured by monitoring β-gal expression from chromosomal transcriptional reporter fusions, as described ^114^.

### Statistical analyses

Statistical analyses were performed using GraphPad Version 10.2.2. All in vitro experiments were conducted at least in triplicate. In all graphs, each symbol represents an independent sample (n), and individual sample size per experiment is also indicated in the figure legends. Statistical test used in each experiment is described in the figure legends.

## Supplementary Materials

### *C. albicans* strain construction

*Construction of a C. albicans constitutive GFP reporter:* Strain SC5314 was transformed with SfiI-linearized vector pDUP3-GFPy via the lithium acetate method as described ^115^.

Transformants were selected on YPD plates containing NAT (200 mg/l). Integration of the cassette was confirmed with primer pairs NEUT5LAMPF + NAT1INTF and NEUT5LAMPR + tADH1AMPR-SpeI (Table S7). Positive fluorescence signal was confirmed by epifluorescence microscopy using a 488 nm laser and corresponding filters.

*Construction of arg4Δ/Δ strain using CRISPR-Cas9 mediated gene editing system*: The *arg4*Δ/Δ strain was constructed by amplifying linearized *SAT1*-flipper and *CaHygB*-flipper plasmids with ARG4CC9KO-F + ARG4CC9KO-R primers to make disruption repair templates bearing nourseothricin (NAT) and hygromycin B (HygB) resistance cassettes, respectively (Table S5).

Before the transformation, *C. albicans* competent cells were prepared by treating freshly grown cells with 1X Tris-EDTA and 0.1 M lithium acetate (pH 7.5) for 1 h at 37°C followed by 25 mM dithiothreitol for an additional 30 min. Cells were washed with sterile water, followed by washing with 1 M sorbitol and resuspended in residual sorbitol. Cells were mixed with *in vitro* assembled ribonucleoprotein (RNP) complexes containing universal tracrRNA, Cas9 protein (2 mg), *ARG4* gene-specific guide RNAs [crARG4up (5′-GATTTATAGGAATCACTTTT), and crARG4down (5′- AACAGCAGCTTTCAAGAATA) (Table S7)], 1 mg of each PCR-generated repair template. The transformation was performed with a single pulse at 1.8 kV using the Gene Pulser Xcell Electroporation System (Bio-Rad). Cells were immediately incubated at 30°C for 4-6 hours in YPD medium and plated on YPD agar containing NAT (200 mg/l) and HygB (600mg/L) (GoldBio). Selected colonies were transferred to a yeast-peptone medium with 2% maltose (YPM) for overnight culture to induce cassette excision. Cells were then selected in YPD agar containing NAT (25 mg/l) and HygB (75 mg/l). Cassette excision and expected auxotrophy was confirmed by growing transformants on YPD, YPD containing NAT (200 mg/l), and YPD containing HygB (600 mg/l) or yeast nitrogen base (YNB) without amino acids and YNB without amino acids containing 100 µg ml^-1^ arginine, respectively. Targeted integration and *ARG4* deletion was confirmed by PCR via amplification of genomic DNA using the primers ARG4DETF + ARG4DETR, ARG4INTF + FLPINTR, and ARG4INTR + FLPINTF as described (Table S7)115,116.

*Construction of arg4Δ/Δ+ARG4 (ARG4 revertant) strain:* The *arg4*Δ/Δ+*ARG4* strain was constructed using overlap extension PCR as described ^116^. The integration cassette consisted of PCR-amplified *5’* (Neut5homology-pDIS3F + Nt5ADH1upUNIVOL-R) and 3’ (ADH1t-UNIVOL-F+ NEUT5homology-pDIS3R) flanking *NEUT5* homology arms using linearized plasmids pDIS3- tADH1. The 3’ fragment also contained the Ca*ADH1* terminator and NAT1 encoding for NAT resistance. The *ARG4* promoter and ORF were amplified from *C. albicans* gDNA using PrARG4OL-F + tARG4OL-R (Table S7). Overlap extension of the amplified products was performed as described ^116^. The overlap product was used as the template in a subsequent PCR reaction using NEUT5homology-pDIS3F + NEUT5homology-pDIS3R primers to make the repair template. The transformation was performed using the CRISPR-Cas9 protocol above, except that crNEUT5pDISup (5′-GCTCGGAGGAGGCTCCCCAA) crRNA was used during RNP assembly and introduced into *arg4*Δ/Δ. Phenotypic confirmation of the transformed cells was confirmed by restoration of growth on YNB without amino acids. The genotype was confirmed by using primers NAT1INTF + NEUT5LAMPR and ARG4DETF + ARG4DETR (Table S7) ^115,116^.

*Construction of SopB expressing C. albicans:* Vector CIpSATtetTrans contains the tetracycline-regulatable (TR) promoter *tetO* and TR transactivator ^117^. To construct a doxycycline-repressible SopB strain, the CIpSATtetTrans vector and the ADH1 terminator synthetic DNA preceded by a new multiple cloning site (MCS-tADH1) were restriction digested with HindIII and MluI, followed by ligation and transformation into *E. coli* DH5α. Transformants were screened on LB agar plates containing ampicillin. Furthermore, the transformed plasmid named CIpSATtetTransMCStADH1 was verified for integrity using restriction digestion and sequencing (Plasmidsaurus). *C. albicans*-optimized STm *sopB* synthetic DNA (CaOptSTSopB) was PCR-amplified using SynthGene-F + SynthGene-Rv2 primers. This PCR product and plasmid CIpSATtetTransMCStADH1 were digested with HindIII and EagI, subsequently ligated, and transformed into *E. coli* DH5α to create plasmid CIpSATtetTransMCStADH1-SopB. Plasmid sequence was verified as described above. The lithium acetate protocol was used as described previously with some modifications to create *C. albicans* strains containing doxycycline-repressible *sopB* or empty vector at the *RPS1* (*i.e., RP10*) locus^118,119^. In brief, 25 μl of overnight culture was mixed with 5 ml YPD and incubated at 30°C with continuous shaking for 6 hours.

After washing with sterile water, cells were resuspended in 1 ml 1X TELiAc buffer (100 mM LiAc, 1X TE). After centrifugation, the cells were resuspended in 50 μl 1X TELiAc, and 1 µg of StuI-linearized CIpSATtetTransMCStADH1-SopB or CIpSATtetTransMCStADH1 was added. Additionally, 5 μl of single-stranded salmon sperm carrier DNA and 300 μl of 40% PEG were added to the suspension. The transformation mix was incubated for 30 minutes at 30°C with gentle agitation every 10 minutes. Cells were then heat-shocked at 42°C for 15 minutes and allowed to recover in YPD at 30°C for 4 h with continuous shaking. Cells were transferred to YPD agar containing 200 mg/l NAT and 10 mg/l doxycycline for selection and transgene repression, respectively. Resulting colonies were further confirmed genotypically by PCR using primers CaSopBDETF + CaSopBDETR and ADH1term-F-ClaI + RP10AMPR (Table S7).

### Histopathology

Tissue sections were fixed with formalin and embedded in paraffin wax. Embedding was performed by the Research Histology core at the University of Illinois Chicago. The tissue was then sectioned via microtome and transferred to slides. Before staining, deparaffinization was performed. The tissue slides were immersed in xylene for 10 min, 100% ethanol for 10 min, 90% ethanol for 2 min, 70% ethanol for 2 min, and PBS for 5 min. The tissue slides were then stained with hematoxylin for 30 seconds, washed with tap water, then stained with eosin for 10 min. Slides were dehydrated by immersion in serial increases of ethanol concentrations (50%-100%), then immersed in xylene. Coverslips were then mounted to the tissue slides and allowed to dry.

Tissue sections were scored for pathology by a board-certified pathologist in a blinded fashion following an approach established by Barthel and colleagues^48^ as summarized below.

Submucosal edema was scored as follows: 0 = no pathological changes; 1 = mild edema (submucosa accounts for <50% of the diameter of the entire intestinal wall [tunica muscularis to epithelium]); 2 = moderate edema; the submucosa accounts for 50 to 80% of the diameter of the entire intestinal wall; and 3 = profound edema (the submucosa accounts for >80% of the diameter of the entire intestinal wall).

Polymorphonuclear granulocytes (PMN) in the lamina propria were enumerated in 10 high-power fields (x400 magnification), and the average number of PMN/high-power fields was calculated. The scores were defined as follows: 0 = <5 PMN/high-power field; 1 = 5 to 20 PMN/high-power field; 2 = 21 to 60/high-power field; 3 = 61 to 100/high-power field; and 4 = >100/high-power field. Transmigration of PMN into the intestinal lumen was consistently observed when the number of PMN was >60 PMN/high-power field.

The average number of goblet cells per high-power field (magnification, x400) was determined from 10 different regions of the cecal epithelium. Scoring was as follows: 0 = >28 goblet cells/high-power field (magnification, x400); 1 = 11 to 28 goblet cells/high-power field; 2 = 1 to 10 goblet cells/high-power field; and 3 = <1 goblet cell/high-power field.

Epithelial integrity was scored as follows: 0 = no pathological changes detectable in 10 high power fields (x400 magnification); 1 = epithelial desquamation; 2 = erosion of the epithelial surface (gaps of 1 to 10 epithelial cells/lesion); and 3 = epithelial ulceration (gaps of >10 epithelial cells/lesion).

Two independent scores for submucosal edema, PMN infiltration, goblet cells, and epithelial integrity were averaged for each tissue sample. The combined pathological score for each tissue sample was determined as the sum of these averaged scores. It ranges between 0 and 13 arbitrary units and covers the following levels of inflammation: 0 intestine intact without any signs of inflammation; 1 to 2 minimal signs of inflammation; 3 to 4 slight inflammation; 5 to moderate inflammation; and 9 to 13 profound inflammation.

## Supplementary Figures

**Fig S1.**
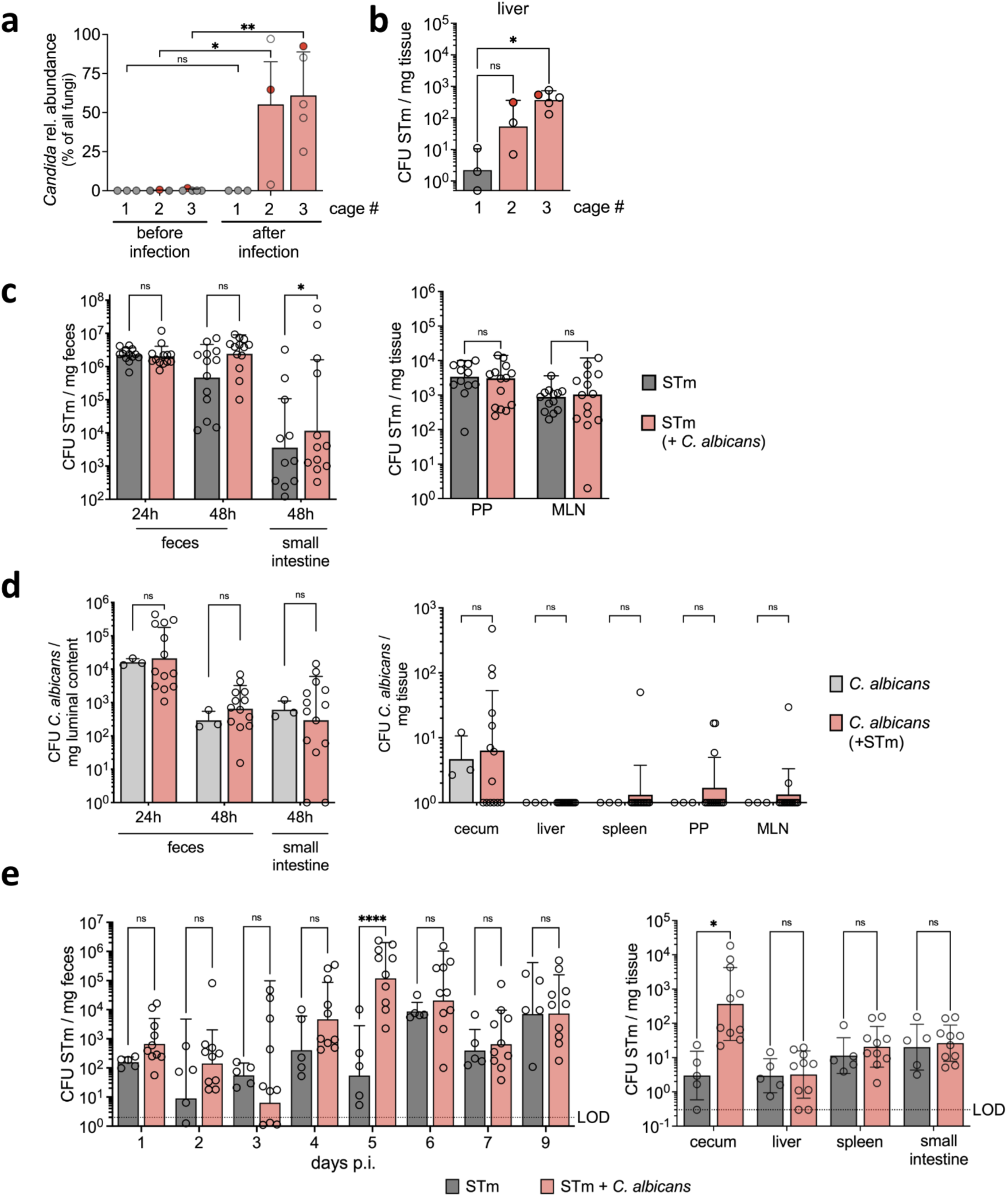
C. albicans increases Salmonella colonization and dissemination. **a,** Relative abundance of *C. albicans* identified with ITS sequencing in fecal samples of mouse before and after STm infection. Mice with reads for *C. albicans* before infection are represented as red circles. Data are mean ± SEM, n=3 (Cage 1 and 2) and n=5 (Cage 3). Two-way ANOVA for comparison. ns = not significant, * = p≤0.05, ** = p≤0.01. CFU, colony-forming units; STm, *Salmonella*. **b,** STm colonization in liver 72h p.i. Mice with reads for *C. albicans* before infection are represented as red circles. Data are geometric mean ± s.d., n=3 (Cage 1 and 2) and n=5 (Cage 3). Ordinary one-way ANOVA for comparison. ns = not significant, * = p≤0.05. CFU, colony-forming units. **c,** STm colonization in C57BL/6 mice infected with STm or STm and *C. albicans* ATCC in the streptomycin pre-treatment model for 48h p.i. Data are geometric mean ± s.d. (for feces and small intestine) and median with range (for PP and MLN), of 3 independent experiments, n ≥ 12. Two-way ANOVA for comparison. ns = not significant, * = p≤0.05. CFU, colony-forming units. **d,** *C. albicans* colonization in C57BL/6 mice infected with *C. albicans* ATCC or STm and *C. albicans* ATCC in the streptomycin pre-treatment model for 48h p.i. Data are geometric mean ± s.d., of 3 independent experiments, n ≥ 3. Two-way ANOVA for comparison. ns = not significant. CFU, colony-forming units. **e,** STm colonization in CBA/J mice infected with STm or STm and *C. albicans* 529L for 9d p.i. Data are geometric mean ± s.d. Two-way ANOVA for comparison. ns = not significant, * = p≤0.05, **** = p≤0.0001. CFU, colony-forming units.

**Fig S2.**
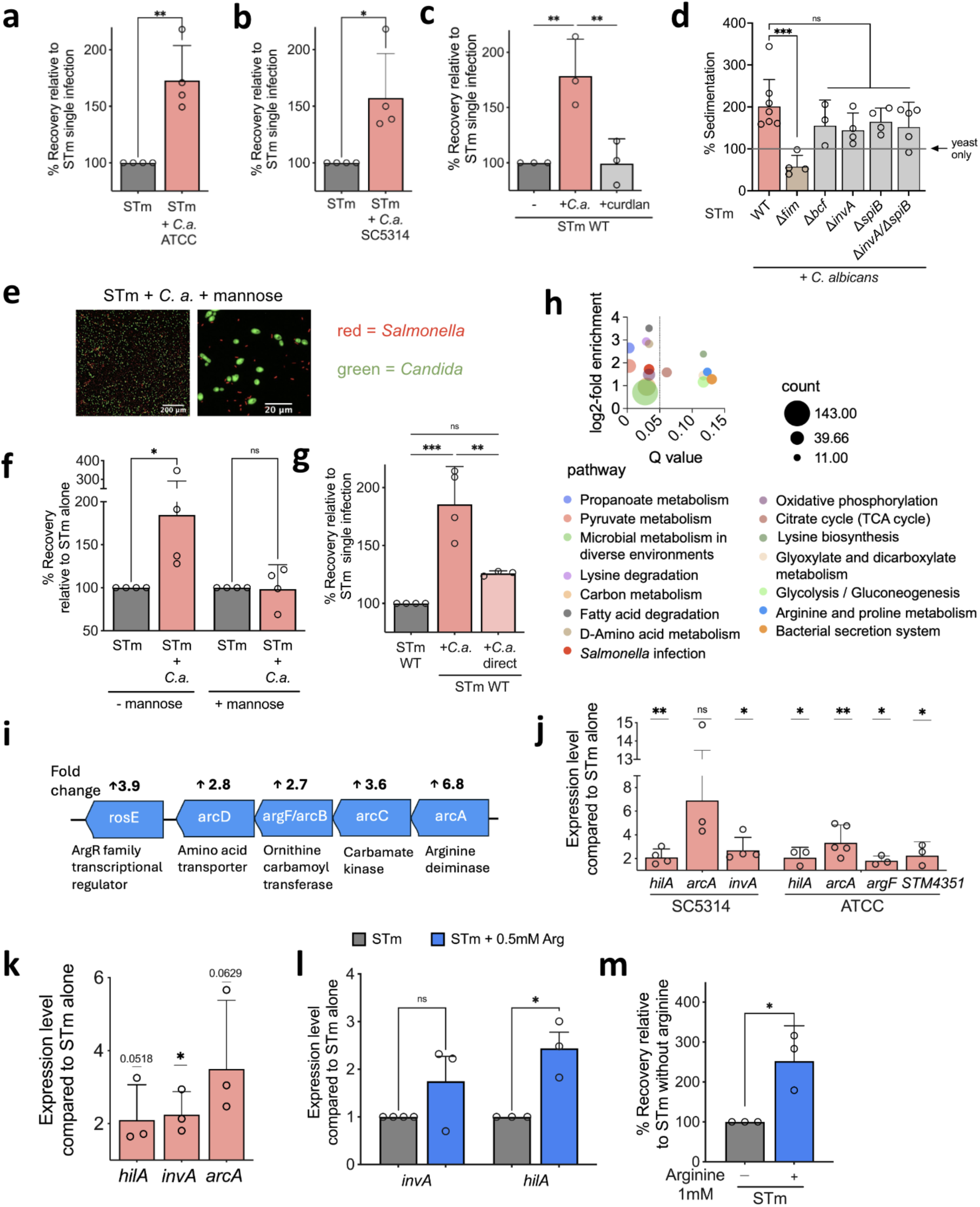
Salmonella invasion and transcriptomic changes in the presence of C. albicans. **a & b**, Invasion assay of STm infected (MOI:1) colonic epithelial cells (Caco2). STm either alone or with (in **a**) *C. albicans* ATCC (in **b**) *C. albicans* SC5314 in a 10:1 ratio were incubated for 2h prior to the assay. Data are geometric mean ± s.d., n = 4. Unpaired t-test for comparison. * = p≤0.05, ** = p≤0.01. STm, *Salmonella.* **c,** Invasion assay of STm infected (MOI:1) colonic epithelial cells (T84). STm either alone or with *C. albicans* ATCC in a 10:1 ratio or curdlan (ß-1,3-glucan) 100μg/ml were incubated for 2h prior to the assay. Data are geometric mean ± s.d., n = 3. Ordinary one-way ANOVA for comparison. ** = p≤0.01. **d,** Sedimentation assay of STm strains and *C. albicans* SC5314. Data are geometric mean ± s.d., n ≥ 3. Ordinary one-way ANOVA for comparison. ns = not significant, *** = p ≤ 0.001. For comparison, STm WT and Δ*fim* values from Fig. 2b were also plotted in this graph. **e,** Fluorescence image of STm (red) and *C. albicans* SC5314 (green) *in vitro* in presence of 5% mannose. **f**, Invasion assay of STm infected (MOI:1) colonic epithelial cells (Caco2). STm either alone or with *C. albicans* ATCC in a 10:1 ratio with or without mannose were incubated for 2h prior to the assay. Data are geometric mean ± s.d., n = 4. Two-way ANOVA for comparison. ns = not significant, * = p≤0.05. **g,** Invasion assay of STm infected (MOI:1) colonic epithelial cells (T84). STm either alone or with *C. albicans* ATCC in a 10:1 ratio incubated for 2h prior to the assay. For direct, both microbes were added to host cells without pre- incubation. Data are geometric mean ± s.d., n = 4. Kruskal-Wallis test for comparison. ns = not significant, ** = p≤0.01, *** = p≤0.001. **h,** Enrichment analysis showing upregulated KEGG pathways in STm in the presence of *C. albicans* ATCC. **i,** Genome organization of Arginine deiminase pathway and the corresponding fold increase in RNAseq analysis. **j,** qRT-PCR analysis of STm genes from the STm or STm with either *C. albicans* ATCC or *C. albicans* SC5314 cultures incubated for 2h. Data are geometric mean ± s.d., n ≥ 3. One sample t-test for comparison. ns = not significant, * = p≤0.05, ** = p≤0.01. **k,** qRT-PCR analysis of STm genes from the STm SL1344 or STm SL1344 with *C. albicans* SC5314 cultures incubated for 2h. Data are geometric mean ± s.d, n = 3. One sample t-test for comparison. * = p≤0.05. **l,** qRT-PCR analysis of STm genes from STm alone or in the presence of L-arginine cultures incubated for 2h. Data are mean ± SEM, n = 3. Two-way ANOVA for comparison. ns = not significant, * = p≤0.05. **m,** Invasion assay of STm infected (MOI:1) colonic epithelial cells (T84). STm either alone or with L-arginine were incubated for 2h prior to the assay. Data are geometric mean ± s.d., n = 3. Unpaired t-test for comparison. * = p≤0.05.

**Fig S3.**
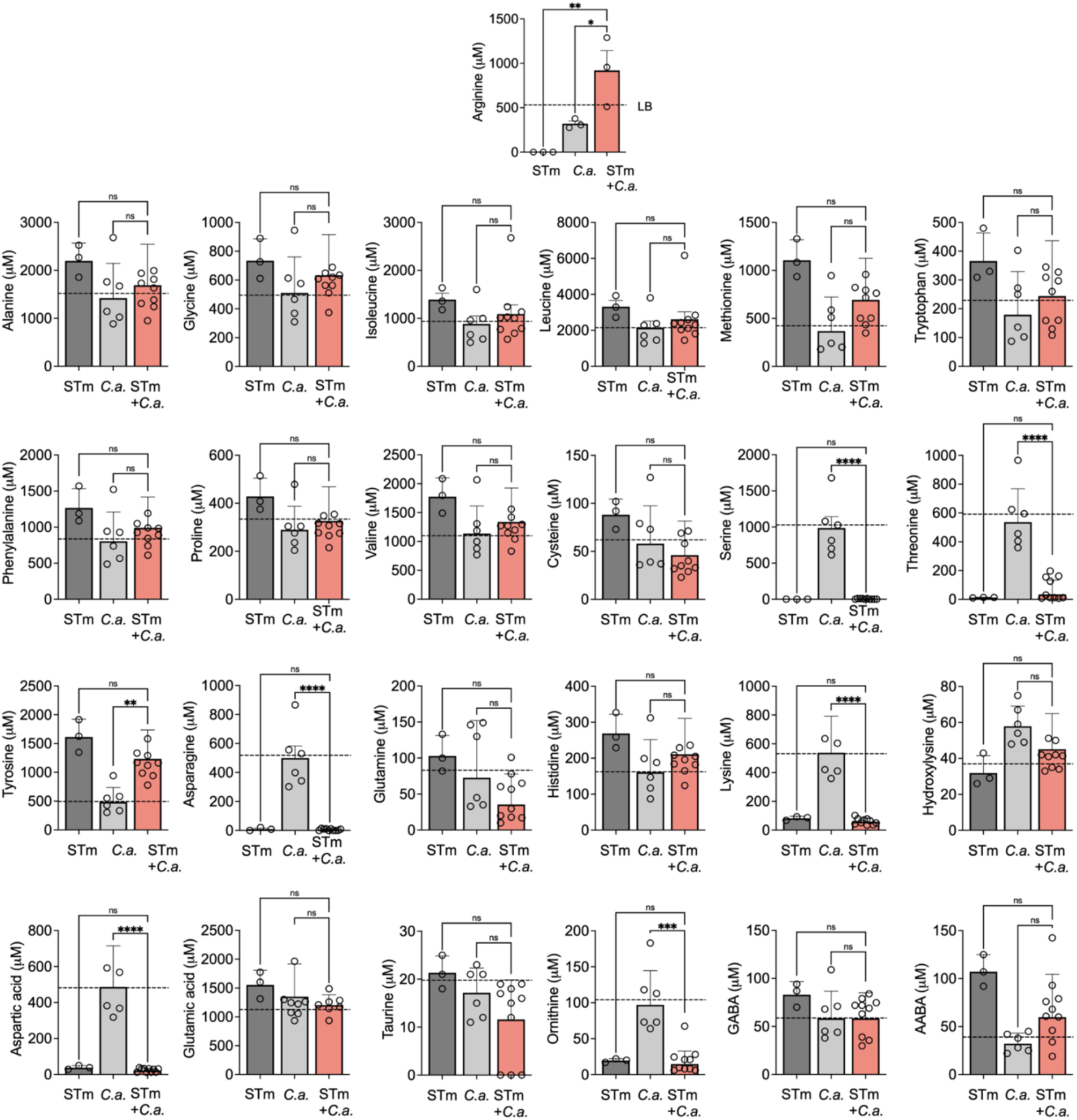
Amino acid levels in the supernatant of Salmonella and C. albicans cultures. Amino acid levels measured in cell-free supernatants of STm, *C. albicans* SC5314, or STm and *C. albicans* SC5314 cultures incubated for 2h. Data are mean ± SEM., n = 3. Ordinary one-way ANOVA for comparison. * = p≤0.05, ** = p≤0.01, *** = p≤0.001, **** = p≤0.0001. Dashed line indicates levels in LB. Arginine level (shown as upper panel for comparison) is the same as is presented in Fig 2g. STm, *Salmonella*.

**Fig S4.**
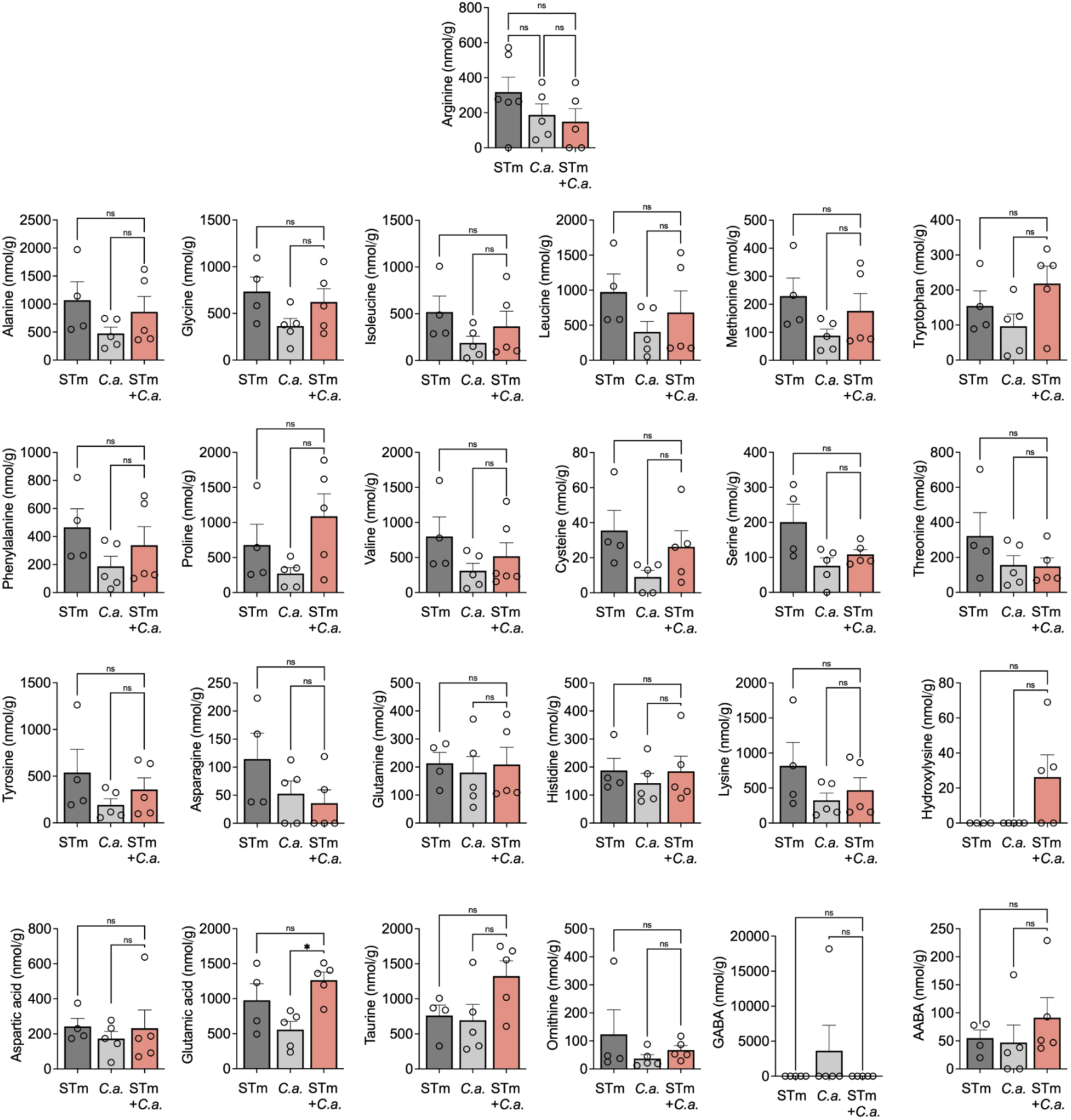
Amino acid levels in the cecum content of mice infected with Salmonella, C. albicans, or co-infected mice. Metabolomics analysis to detect amino acid levels in the cecum content of STm, *C. albicans* SC5314, or STm and *C. albicans* SC5314 infected mice 48h p.i. Data are mean ± SEM., n ≥ 4. Ordinary one-way ANOVA for comparison. ns = not significant, * = p≤0.05. STm, *Salmonella*.

**Fig S5.**
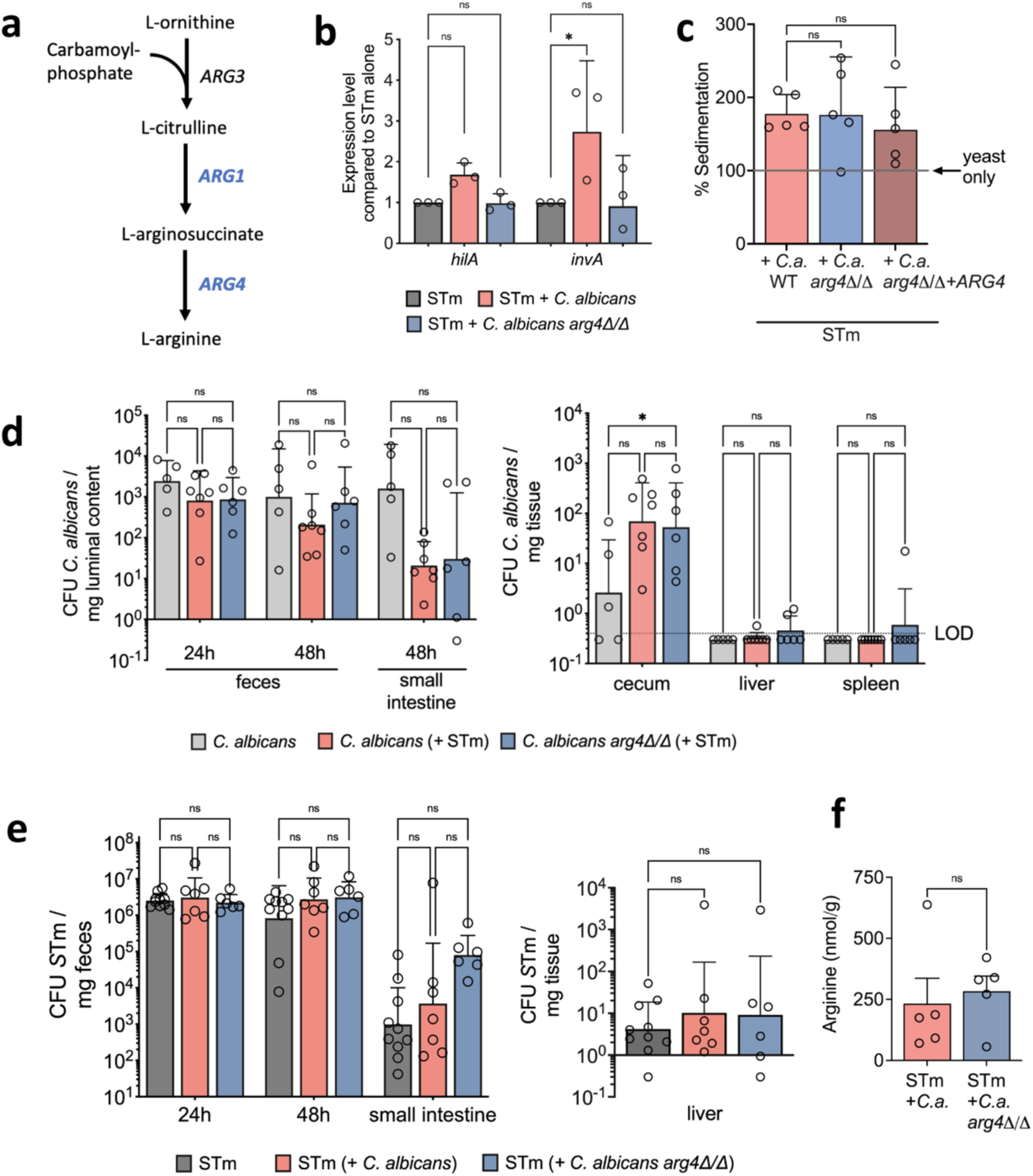
Role of C. albicans arginine biosynthesis for Salmonella virulence. **a,** Schematic of arginine biosynthesis in *C. albicans*. **b,** qRT-PCR analysis of STm genes from the STm or STm with *C. albicans* SC5314 cultures incubated for 2h. Data are geometric mean ± s.d, n = 3. Two-way ANOVA for comparison. ns = not significant, * = p≤0.05. STm, *Salmonella*. **c,** Sedimentation assay of STm and *C. albicans* SC5314. Data are geometric mean ± s.d., n = 5. Ordinary one-way ANOVA for comparison. ns = not significant. **d,** *C. albicans* colonization in C57BL/6 mice infected with *C. albicans* SC5314 alone or STm and *C. albicans* SC5314 in the streptomycin pre-treatment model for 48h p.i. Data are geometric mean ± s.d. of 2 independent experiments. Two-way ANOVA for comparison. ns = not significant, * = p≤0.05. CFU, colony-forming units. **e,** STm colonization in C57BL/6 mice infected with STm or STm and *C. albicans* SC5314 in the streptomycin pre-treatment model for 48h p.i. Data are geometric mean ± s.d. of 2 independent experiments. Ordinary one-way ANOVA (for liver) and Mixed-effects analysis (for fecal sample) for comparison. ns = not significant. CFU, colony-forming units. **f,** Arginine levels measured in the cecum content of STm and *C. albicans* SC5314 infected mice 48h p.i. Data are mean ± SEM., n = 5. Unpaired t-test for comparison. ns = not significant.

**Fig S6.**
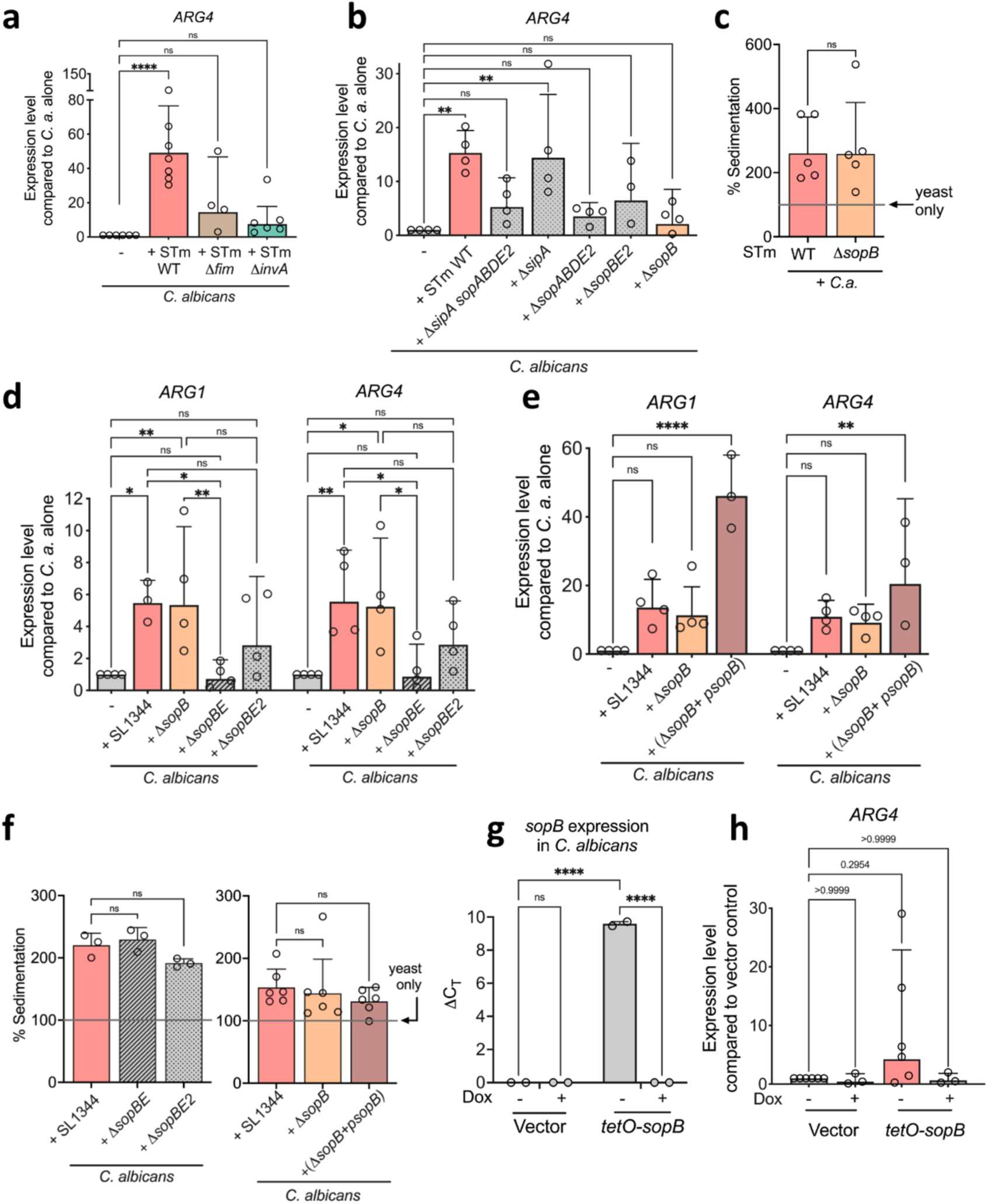
Salmonella uses T3SS-1 effector(s) to trigger arginine production in C. albicans. **a & b**, qRT-PCR analysis of genes encoding for *C. albicans* arginine biosynthesis from the *C. albicans* SC5314 or STm and *C. albicans* SC5314 cultures incubated for 2h. Data are geometric mean ± s.d., n ≥ 4. Kruskal-Wallis test (for **a**) and Ordinary one-way ANOVA (for **b**) for comparison. ns = not significant, ** = p≤0.01, **** = p≤0.0001. STm, *Salmonella*. **c,** Sedimentation assay of STm and *C. albicans* SC5314. Data are geometric mean ± s.d., n = 5. Two-tailed unpaired t-test for comparison. ns = not significant. **d & e,** qRT-PCR analysis of genes encoding for *C. albicans* arginine biosynthesis from the *C. albicans* SC5314 or STm SL1344 and *C. albicans* SC5314 cultures incubated for 2h. Data are geometric mean ± s.d., n ≥ 3. Mixed-effect analysis (for **a**) and Two-way ANOVA (for **b**) for comparison. ns = not significant, * = p≤0.05, ** = p≤0.01, **** = p≤0.0001. **f,** Sedimentation assay of STm SL1344 and *C. albicans* SC5314. Data are geometric mean ± s.d., n = 3 (for left panel) and n = 6 (for right panel). Ordinary one-way ANOVA for comparison. ns = not significant. **g,** qRT-PCR analysis to confirm *sopB* expression in *C. albicans* SC5314 cultures expressing vector or *tetO*-*sopB*. Data are mean ± SEM., n = 2. Two-way ANOVA for comparison. ns = not significant, **** = p≤0.0001. **h,** qRT-PCR analysis of genes encoding for *C. albicans* arginine biosynthesis from the *C. albicans* SC5314 cultures expressing vector or *tetO*-*sopB* incubated for 6h with or without doxycycline. Data are geometric mean ± s.d., n ≥ 3. Kruskal-Wallis test for comparison. ns = not significant.

**Fig S7.**
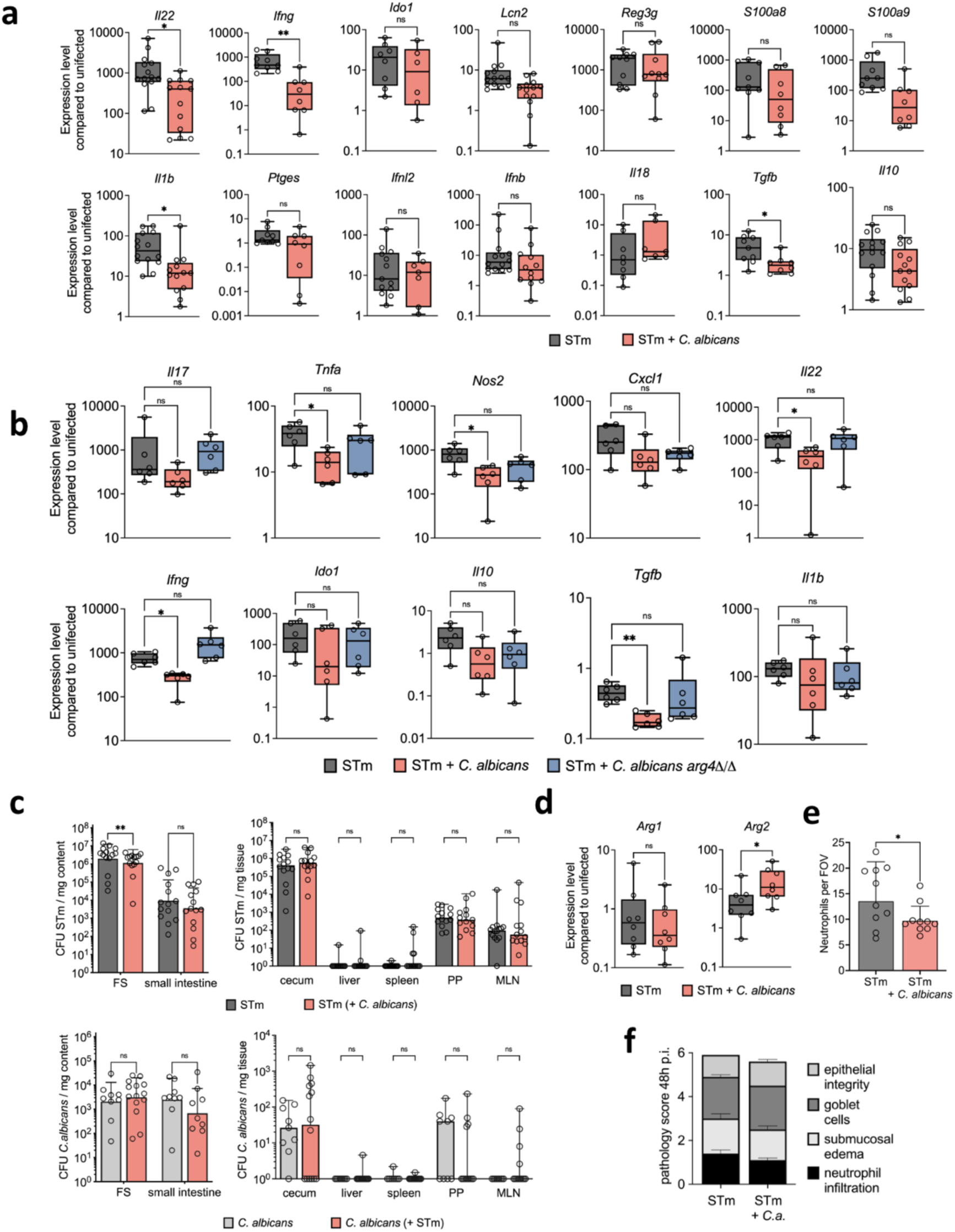
Immune response to Salmonella and C. albicans co-infection. **a & b,** qRT-PCR analysis of genes encoding for host inflammatory response from the cecum tissue of (for **a**) STm or STm and *C. albicans* ATCC infected mice 24h p.i. and (for **b**) STm or STm and *C. albicans* SC5314 infected mice 48h p.i. Data is from 3 (for **a**) and 2 (for **b)** independent experiments, n ≥ 6. Data are presented as box and whiskers plot from 25th to 75th percentile, median, minimum and maximum. Significance determined by two-tailed unpaired t-test (for **a**) and Kruskal-Wallis test (for **b**). ns = not significant, * = p≤0.05, ** = p≤0.01. STm, *Salmonella*. **c,** STm and *C. albicans* colonization in C57BL/6 mice infected with STm, *C. albicans* ATCC, or STm and *C. albicans* ATCC in the streptomycin pre-treatment model for 24 p.i. Data are geometric mean ± s.d. (for fecal samples and SI content), Median with range (for organ colonization) of 3 independent experiments, n ≥ 9. Two-way ANOVA (for FS) and mixed-effects analysis (for organs) for comparison. ns = not significant, ** = p≤0.01. CFU, colony-forming units. **d,** qRT-PCR analysis of genes encoding for host arginase genes from the cecum tissue of STm or STm and *C. albicans* ATCC infected mice 24h p.i. Data is from 2 independent experiments, n = 8. Significance determined by two-tailed Mann–Whitney test. ns = not significant, * = p≤0.05. **e,** Neutrophils counts for cecum 48h p.i. from STm or STm and *C. albicans* ATCC infected mice. Data are geometric mean ± s.d., n = 10. Welch’s t-test for comparison. * = p≤0.05. FOV: field of view. **f**, Pathological scores in cecum of STm or STm and *C. albicans* ATCC infected mice 48h p.i. Data are mean ± SEM., n = 10.

**Fig S8.**
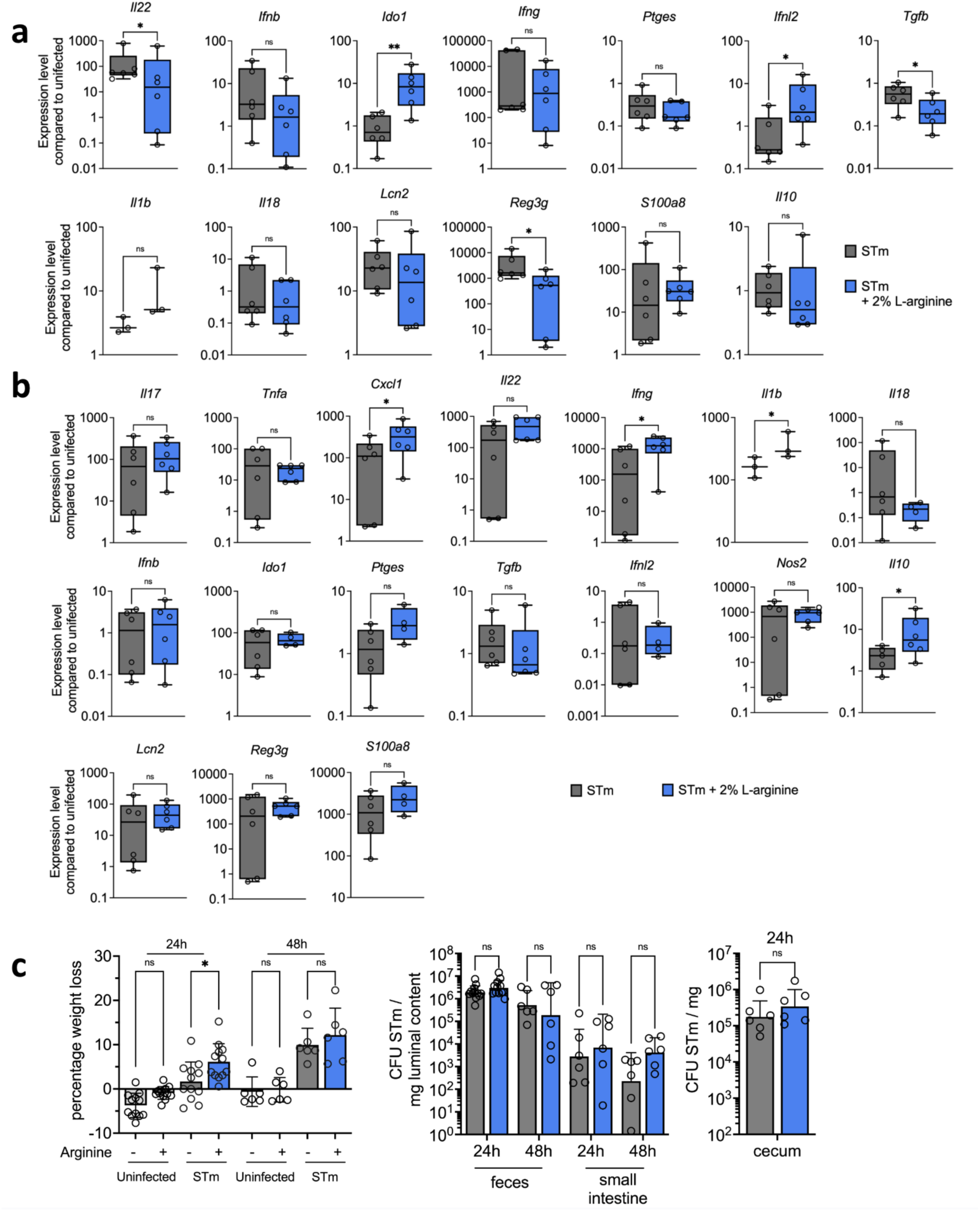
Effect of arginine supplementation during Salmonella infection. **a & b,** qRT-PCR analysis of genes encoding for host inflammatory response from the cecum tissue of STm infected mice with or without L-arginine supplementation 24h (for **a**) and 48h (for **b**) p.i. Data is from 2 independent experiments, n ≥ 3. Data are presented as box and whiskers plot from 25th to 75th percentile, median, minimum and maximum. Significance determined by two-tailed Mann–Whitney test. ns = not significant, * = p≤0.05, ** = p≤0.01. STm, *Salmonella*. **c,** STm colonization in STm or STm with L-arginine treated C57BL/6 mice in the streptomycin pre-treatment model for 24h and 48h p.i. Data are mean ± s.d. (for weight loss) and geometric mean ± s.d. (for fecal samples, SI content and cecum) of 2 independent experiments, n = 6. Mixed-effects analysis (for weight loss, fecal samples and SI content) and two-tailed Mann– Whitney test (for cecum) for comparison. ns = not significant, * = p≤0.05. CFU, colony-forming units.

**Fig S9.**
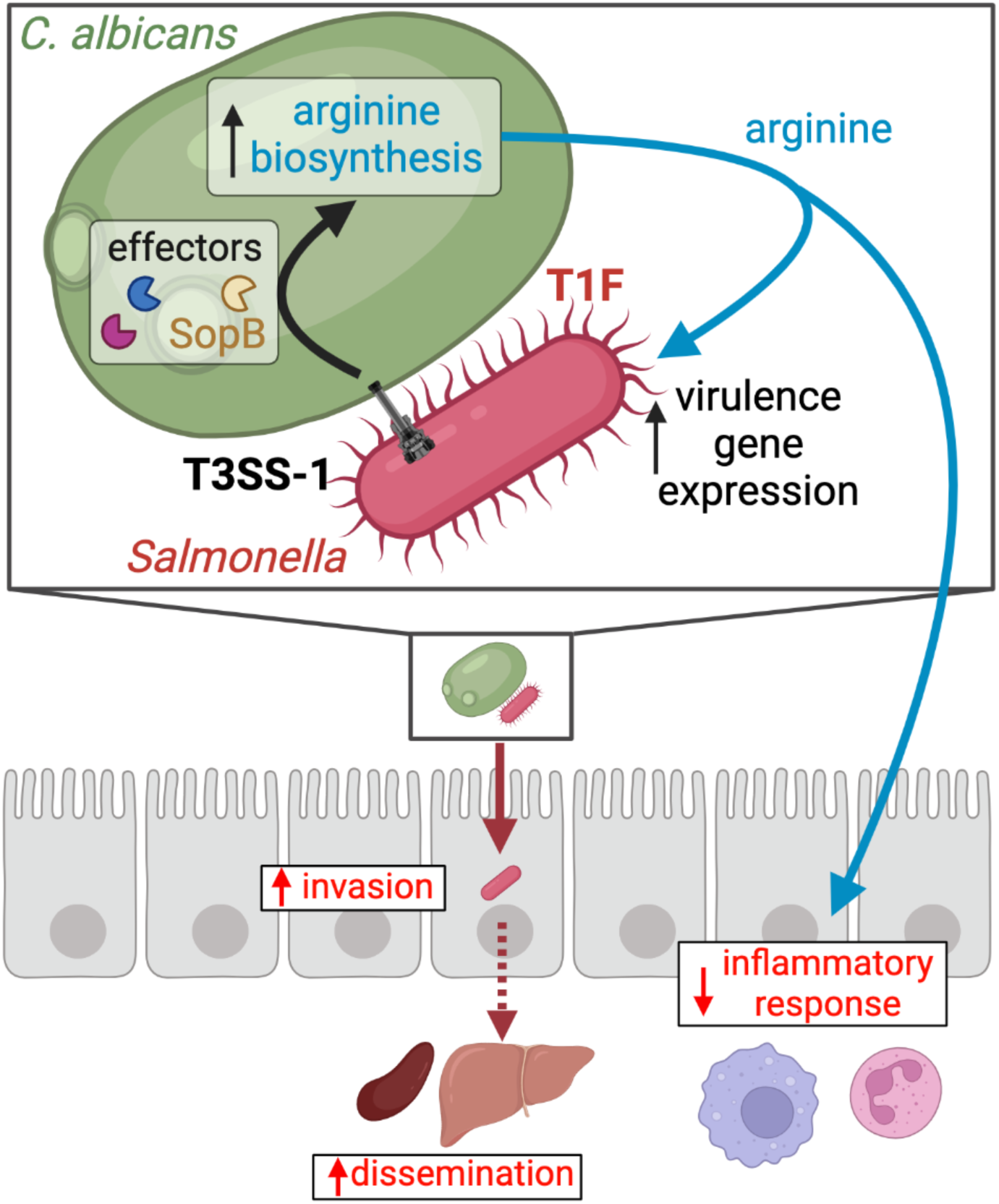
Proposed model of interactions between Salmonella, C. albicans, and the host during infection. *Salmonella* binds to *C. albicans* via Type 1 fimbriae (T1F) and uses its Type 3 Secretion System (T3SS)-1 to deliver effector molecules, including SopB, into *C. albicans*. The effector SopB increases arginine biosynthesis in *C. albicans*, which is exported into the extracellular environment. *Salmonella* senses the released arginine and increases virulence gene expression, which results in increased invasion of epithelial cells. Arginine also decreases the inflammatory response to the infection, further facilitating *Salmonella* pathogenicity and dissemination to peripheral organs.

**Table S5.** Strains used in this study.

| Strain name (as used in this study) | Relevant genotype/feature | Source |
| --- | --- | --- |
| *Salmonella enterica* serovar Typhimurium strains |  |  |
| STm | IR715; ATCC 14028 with spontaneous Nal<sup>R</sup> derivative | ATCC |
| STm (Mouse experiments) | IR715 + pHP45Ω | Vladimir E. Diaz-Ochoa |
| $\Delta fim$ | EHW2; IR715 $\Delta fimAICDHF$ | Weening et al 2005<sup>120</sup> |
| $\Delta bcf$ | EHW1; IR715 $bcfABCDEFG::kan$ | Weening et al 2005<sup>120</sup> |
| $\Delta invA$ | BL212; IR715 $\Delta invA$ | Devlin et al 2022<sup>91</sup> |
| $\Delta spiB$ | SPN450; IR715 $spiB::KSAC$ | Manuela Raffatellu et al 2009<sup>96</sup> |
| $\Delta invA/\Delta spiB$ | SPN452; IR715 $invA::tetRA spiB::KSAC$ | Manuela Raffatellu et al 2009<sup>96</sup> |
| STm (fluorescence microscopy) | IR715 + pFPV-*mCherry* | This study |
| *hilA-lacZ* | JS749; attλ::pDX1::*hilA'-lacZ* | Lin et al 2008<sup>121</sup> |
| $\Delta sipA \Delta sopABDE2$ | ZA21; IR715 $\Delta sipA \Delta sopABDE2$ | Zhang et al 2002<sup>122</sup> |
| $\Delta sipA$ | ZA10; IR715 $\Delta sipA$ | Zhang et al 2002<sup>122</sup> |
| $\Delta sopABDE2$ | ZA20; IR715 $\Delta sopABDE2$ | Zhang et al 2002<sup>122</sup> |
| $\Delta sopBE2$ | ZA16; IR715 $\Delta sopBE2$ | Zhang et al 2002<sup>122</sup> |
| $\Delta sopB$ | ZA15; IR715 $\Delta sopB$ | Zhang et al 2002<sup>122</sup> |
| SL1344 |  | Hoiseth S.K et al 1981<sup>123</sup> |
| SL1344 $\Delta sopB$ | SL1344 $\Delta sigDE$ | Knodler et al 2006<sup>124</sup> |
| SL1344 $\Delta sopB$ + *psopB* | SL1344 $\Delta sigDE$ pWSKDE (pWSK29+*sigDE*) | Knodler et al 2009<sup>125</sup> |
| SL1344 $\Delta sopBE$ | SL1344 $\Delta sopB \Delta sopE::aphT$ | Cooper et al 2011<sup>126</sup> |
| SL1344 $\Delta sopBE2$ | SL1344 $\Delta sopB \Delta sopE2::tet$ | Cooper et al 2011<sup>126</sup> |
| *Candida albicans* strains |  |  |
| *C. albicans* ATCC | *Candida albicans* (Robin) Berkhout, ATCC<sup>a</sup> #90028 | ATCC |
| *C. albicans* 529L |  | Rahman et al. 2007<sup>127</sup> |
| *C. albicans* SC5314 | wild-type | Gillium et al. 1984<sup>128</sup> |
| *C. albicans* (fluorescence microscopy) | SC5314 GFPy |  |
| SC5314 *arg4Δ/Δ* | *arg4Δ::FRT+/arg4Δ::FRT+* | This study |
| SC5314 *arg4Δ/Δ+ARG4* | *arg4Δ::FRT+/arg4Δ::FRT+ NEUT5L/neut5Δ::ARG4* | This study |
| SC5314 empty vector | *RPS1/RPS1::SAT1-tetTA-PrtetO-tADH1* | This study |
| SC5314 *tetO-sopB* | *RPS1/RPS1::SAT1-tetTA-PrtetO-sopB-tADH1* | This study |

**Table S6.** Plasmids used in this study.

| Plasmid name | Relevant feature | Source |
| --- | --- | --- |
| pFPV-*mCherry* | ColE1 *ori*, *PrpsM.mCherry* | Drecktrah et al. 2008<sup>129</sup> |
| *SAT1*-flipper | FRT-PrCaMAL2-FLP-*SAT1*-FRT | Reuss et al. 2004<sup>130</sup> |
| *CaHygB*-flipper | FRT-PrCaMAL2-FLP-*CaHygB*-FRT | Liu et al. 2022<sup>116</sup> |
| ClpSATtetTrans | *RPS1-tetR-SchAP4AD-PrtetO-SAT1* | Steve Saville |
| ClpSATtetTransMCStADH1 | MCS-tADH1 in ClpSATtetTrans | This study |
| ClpSATtetTransMCStADH1-SopB | *Salmonella sopB* in ClpSATtetTransMCStADH1 | This study |

**Table S7.**
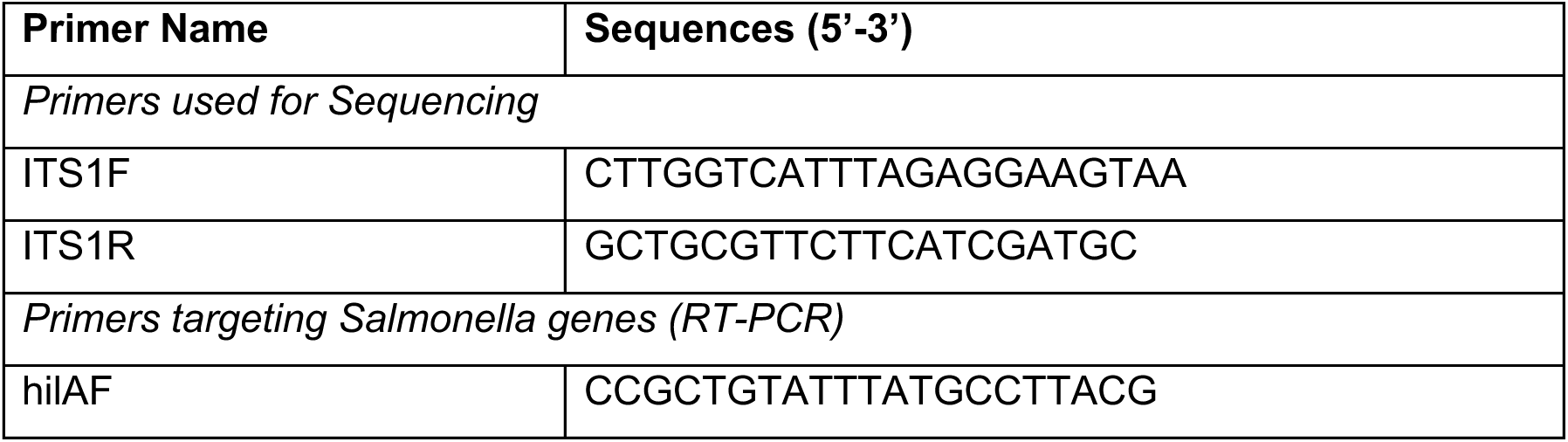
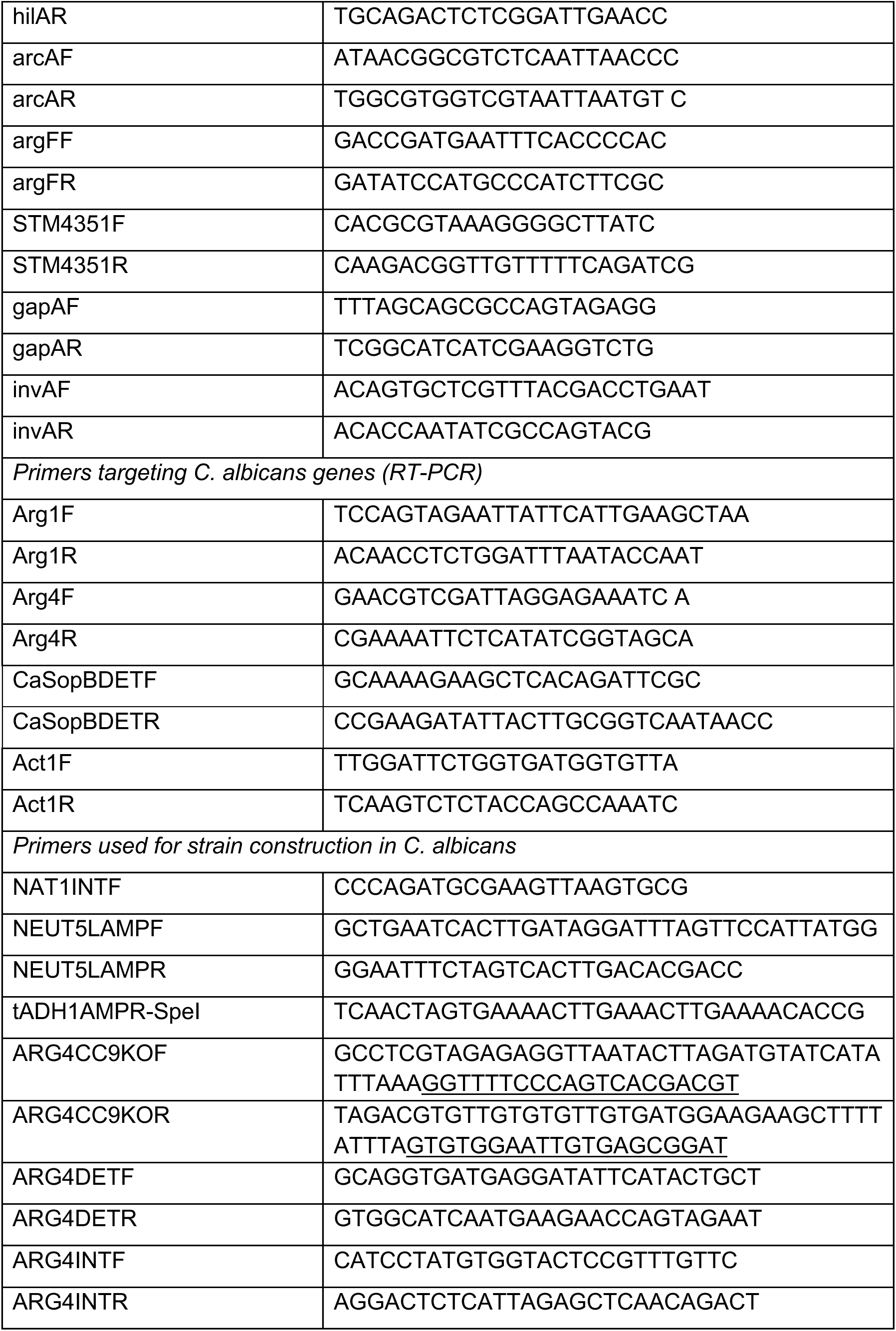
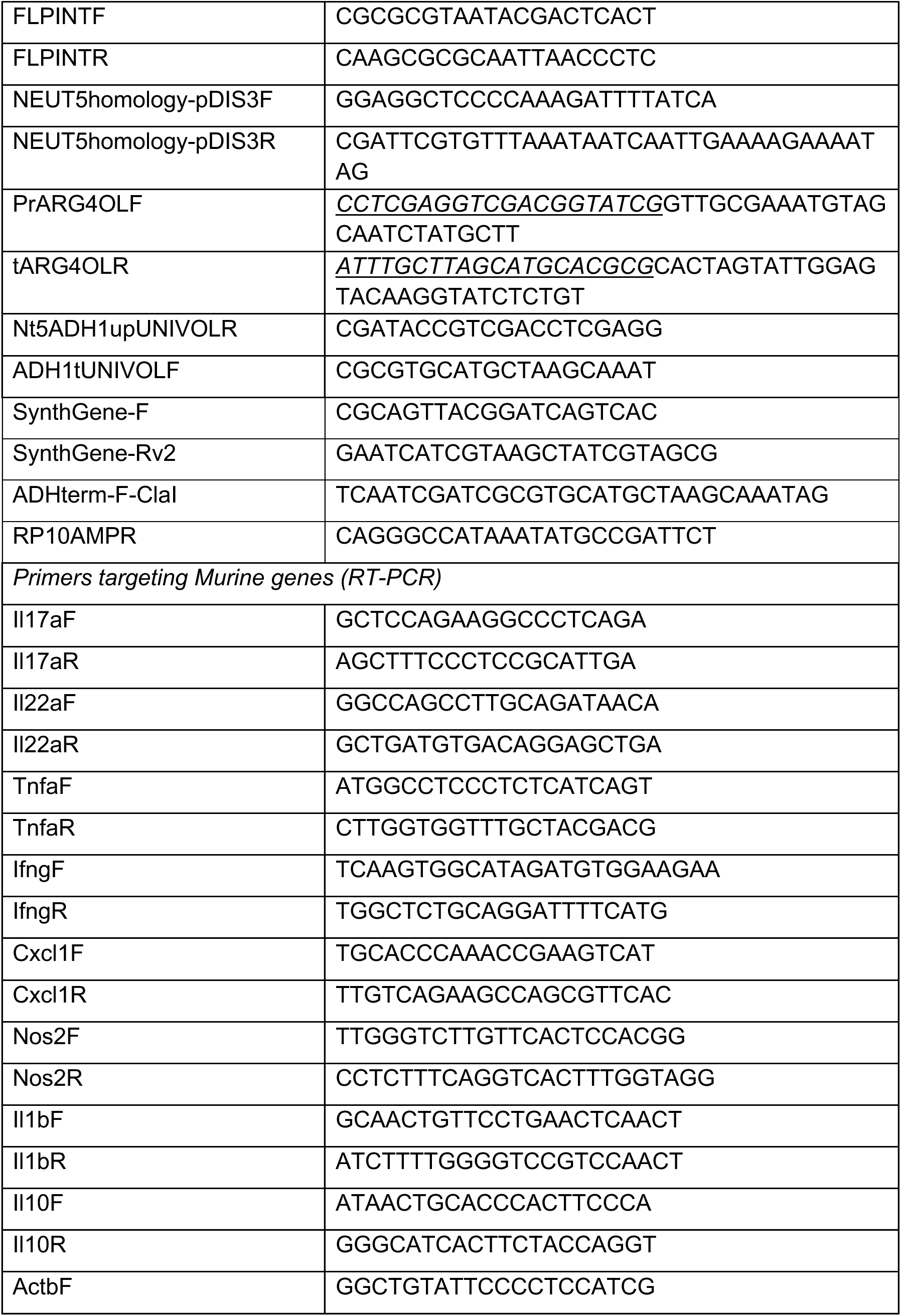
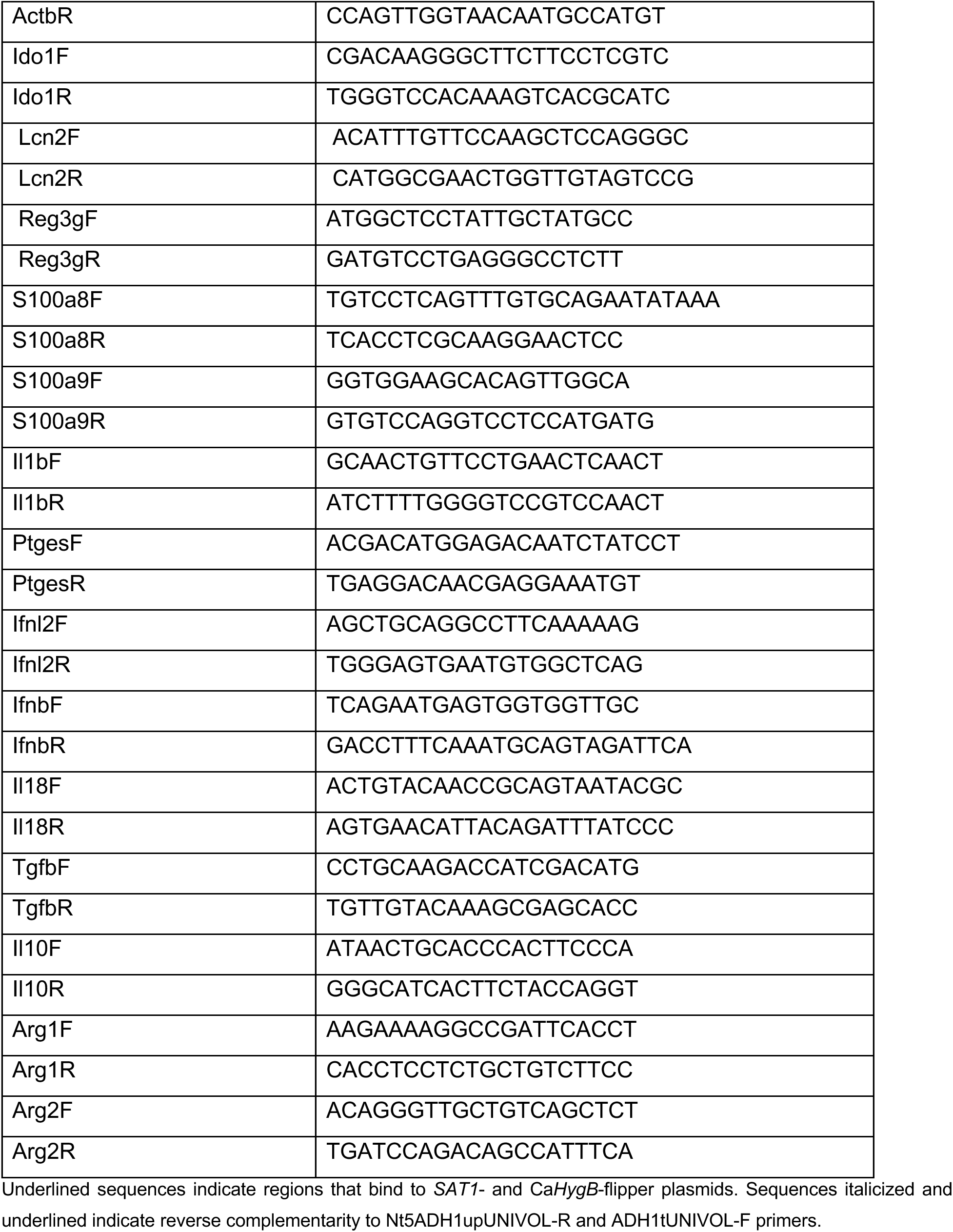
Primers used in this study.

