## Supplementary Table 1 for "Commensal Yeast Promotes *Salmonella* Typhimurium Virulence"

**Table S1: ITS sequencing results of fecal samples before and after STm infection.**

| ASV | Sample | Abundance | Infection | Kingdom | Phylum | Class | Order | Family |
|---|---|---|---|---|---|---|---|---|
| ASV2 | ITS-Behnsen-MY-1 | 0.26 | None | k__Fungi | p__Ascomycota | c__Eurotiomycetes | o__Eurotiales | f__Aspergillaceae |
| ASV5 | ITS-Behnsen-MY-1 | 0.22 | None | k__Fungi | p__Basidiomycota | c__Ustilaginomycetes | o__Ustilaginales | f__Ustilaginaceae |
| ASV3 | ITS-Behnsen-MY-1 | 0.20 | None | k__Fungi | p__Ascomycota | c__Dothideomycetes | o__Pleosporales | f__Didymellaceae |
| ASV4 | ITS-Behnsen-MY-1 | 0.11 | None | k__Fungi | p__Ascomycota | c__Dothideomycetes | o__Capnodiales | f__Cladosporiaceae |
| ASV14 | ITS-Behnsen-MY-1 | 0.06 | None | k__Fungi | p__Basidiomycota | c__Tremellomycetes | o__Tremellales | f__Bulleribasidiaceae |
| ASV16 | ITS-Behnsen-MY-1 | 0.03 | None | k__Fungi | p__Basidiomycota | c__Wallemiomycetes | o__Wallemiales | f__Wallemiaceae |
| ASV8 | ITS-Behnsen-MY-1 | 0.03 | None | k__Fungi | p__Ascomycota | c__Dothideomycetes | o__Capnodiales | f__Mycosphaerellaceae |
| ASV24 | ITS-Behnsen-MY-1 | 0.03 | None | k__Fungi | p__Basidiomycota | c__Microbotryomycetes | o__Sporidiobolales | f__Sporidiobolaceae |
| ASV11 | ITS-Behnsen-MY-1 | 0.02 | None | k__Fungi | p__Ascomycota | c__Eurotiomycetes | o__Eurotiales | f__Trichocomaceae |
| ASV7 | ITS-Behnsen-MY-1 | 0.02 | None | k__Fungi | p__Ascomycota | c__Dothideomycetes | o__Pleosporales | f__Pleosporaceae |
| ASV30 | ITS-Behnsen-MY-1 | 0.01 | None | k__Fungi | p__Ascomycota | c__Leotiomycetes | o__Helotiales | f__Myxotrichaceae |
| ASV12 | ITS-Behnsen-MY-1 | 0.01 | None | k__Fungi | p__Ascomycota | c__Sordariomycetes | o__Diaporthales | f__Diaporthaceae |
| ASV39 | ITS-Behnsen-MY-1 | 0.00 | None | k__Fungi | p__Ascomycota | c__Sordariomycetes | o__Hypocreales | f__Hypocreaceae |
| ASV159 | ITS-Behnsen-MY-1 | 0.00 | None | k__Fungi | p__Mucoromycota | c__Mucoromycetes | o__Mucorales | f__Lichtheimiaceae |
| ASV1 | ITS-Behnsen-MY-1 | 0.00 | None | k__Fungi | p__Ascomycota | c__Saccharomycetes | o__Saccharomycetales | f__Saccharomycetales_fam_Incertae_sedis |
| ASV44 | ITS-Behnsen-MY-1 | 0.00 | None | k__Fungi | p__Basidiomycota | c__Tremellomycetes | o__Filobasidiales | f__Filobasidiaceae |
| ASV28 | ITS-Behnsen-MY-1 | 0.00 | None | k__Fungi | p__Ascomycota | c__Sordariomycetes | o__Microascales | f__Microascaceae |
| ASV2 | ITS-Behnsen-MY-2 | 0.40 | None | k__Fungi | p__Ascomycota | c__Eurotiomycetes | o__Eurotiales | f__Aspergillaceae |
| ASV3 | ITS-Behnsen-MY-2 | 0.19 | None | k__Fungi | p__Ascomycota | c__Dothideomycetes | o__Pleosporales | f__Didymellaceae |
| ASV5 | ITS-Behnsen-MY-2 | 0.17 | None | k__Fungi | p__Basidiomycota | c__Ustilaginomycetes | o__Ustilaginales | f__Ustilaginaceae |
| ASV4 | ITS-Behnsen-MY-2 | 0.08 | None | k__Fungi | p__Ascomycota | c__Dothideomycetes | o__Capnodiales | f__Cladosporiaceae |
| ASV11 | ITS-Behnsen-MY-2 | 0.03 | None | k__Fungi | p__Ascomycota | c__Eurotiomycetes | o__Eurotiales | f__Trichocomaceae |
| ASV14 | ITS-Behnsen-MY-2 | 0.03 | None | k__Fungi | p__Basidiomycota | c__Tremellomycetes | o__Tremellales | f__Bulleribasidiaceae |
| ASV8 | ITS-Behnsen-MY-2 | 0.02 | None | k__Fungi | p__Ascomycota | c__Dothideomycetes | o__Capnodiales | f__Mycosphaerellaceae |
| ASV24 | ITS-Behnsen-MY-2 | 0.02 | None | k__Fungi | p__Basidiomycota | c__Microbotryomycetes | o__Sporidiobolales | f__Sporidiobolaceae |
| ASV16 | ITS-Behnsen-MY-2 | 0.02 | None | k__Fungi | p__Basidiomycota | c__Wallemiomycetes | o__Wallemiales | f__Wallemiaceae |
| ASV30 | ITS-Behnsen-MY-2 | 0.01 | None | k__Fungi | p__Ascomycota | c__Leotiomycetes | o__Helotiales | f__Myxotrichaceae |
| ASV7 | ITS-Behnsen-MY-2 | 0.01 | None | k__Fungi | p__Ascomycota | c__Dothideomycetes | o__Pleosporales | f__Pleosporaceae |
| ASV39 | ITS-Behnsen-MY-2 | 0.01 | None | k__Fungi | p__Ascomycota | c__Sordariomycetes | o__Hypocreales | f__Hypocreaceae |
| ASV44 | ITS-Behnsen-MY-2 | 0.00 | None | k__Fungi | p__Basidiomycota | c__Tremellomycetes | o__Filobasidiales | f__Filobasidiaceae |
| ASV159 | ITS-Behnsen-MY-2 | 0.00 | None | k__Fungi | p__Mucoromycota | c__Mucoromycetes | o__Mucorales | f__Lichtheimiaceae |
| ASV28 | ITS-Behnsen-MY-2 | 0.00 | None | k__Fungi | p__Ascomycota | c__Sordariomycetes | o__Microascales | f__Microascaceae |
| ASV1 | ITS-Behnsen-MY-2 | 0.00 | None | k__Fungi | p__Ascomycota | c__Saccharomycetes | o__Saccharomycetales | f__Saccharomycetales_fam_Incertae_sedis |
| ASV12 | ITS-Behnsen-MY-2 | 0.00 | None | k__Fungi | p__Ascomycota | c__Sordariomycetes | o__Diaporthales | f__Diaporthaceae |
| ASV3 | ITS-Behnsen-MY-3 | 0.42 | None | k__Fungi | p__Ascomycota | c__Dothideomycetes | o__Pleosporales | f__Didymellaceae |
| ASV2 | ITS-Behnsen-MY-3 | 0.21 | None | k__Fungi | p__Ascomycota | c__Eurotiomycetes | o__Eurotiales | f__Aspergillaceae |
| ASV5 | ITS-Behnsen-MY-3 | 0.15 | None | k__Fungi | p__Basidiomycota | c__Ustilaginomycetes | o__Ustilaginales | f__Ustilaginaceae |
| ASV4 | ITS-Behnsen-MY-3 | 0.09 | None | k__Fungi | p__Ascomycota | c__Dothideomycetes | o__Capnodiales | f__Cladosporiaceae |
| ASV11 | ITS-Behnsen-MY-3 | 0.03 | None | k__Fungi | p__Ascomycota | c__Eurotiomycetes | o__Eurotiales | f__Trichocomaceae |
| ASV14 | ITS-Behnsen-MY-3 | 0.03 | None | k__Fungi | p__Basidiomycota | c__Tremellomycetes | o__Tremellales | f__Bulleribasidiaceae |
| ASV8 | ITS-Behnsen-MY-3 | 0.01 | None | k__Fungi | p__Ascomycota | c__Dothideomycetes | o__Capnodiales | f__Mycosphaerellaceae |
| ASV16 | ITS-Behnsen-MY-3 | 0.01 | None | k__Fungi | p__Basidiomycota | c__Wallemiomycetes | o__Wallemiales | f__Wallemiaceae |
| ASV7 | ITS-Behnsen-MY-3 | 0.01 | None | k__Fungi | p__Ascomycota | c__Dothideomycetes | o__Pleosporales | f__Pleosporaceae |
| ASV28 | ITS-Behnsen-MY-3 | 0.01 | None | k__Fungi | p__Ascomycota | c__Sordariomycetes | o__Microascales | f__Microascaceae |
| ASV24 | ITS-Behnsen-MY-3 | 0.01 | None | k__Fungi | p__Basidiomycota | c__Microbotryomycetes | o__Sporidiobolales | f__Sporidiobolaceae |
| ASV39 | ITS-Behnsen-MY-3 | 0.00 | None | k__Fungi | p__Ascomycota | c__Sordariomycetes | o__Hypocreales | f__Hypocreaceae |
| ASV44 | ITS-Behnsen-MY-3 | 0.00 | None | k__Fungi | p__Basidiomycota | c__Tremellomycetes | o__Filobasidiales | f__Filobasidiaceae |
| ASV30 | ITS-Behnsen-MY-3 | 0.00 | None | k__Fungi | p__Ascomycota | c__Leotiomycetes | o__Helotiales | f__Myxotrichaceae |
| ASV1 | ITS-Behnsen-MY-3 | 0.00 | None | k__Fungi | p__Ascomycota | c__Saccharomycetes | o__Saccharomycetales | f__Saccharomycetales_fam_Incertae_sedis |
| ASV12 | ITS-Behnsen-MY-3 | 0.00 | None | k__Fungi | p__Ascomycota | c__Sordariomycetes | o__Diaporthales | f__Diaporthaceae |
| ASV159 | ITS-Behnsen-MY-3 | 0.00 | None | k__Fungi | p__Mucoromycota | c__Mucoromycetes | o__Mucorales | f__Lichtheimiaceae |
| ASV2 | ITS-Behnsen-MY-35 | 0.62 | None | k__Fungi | p__Ascomycota | c__Eurotiomycetes | o__Eurotiales | f__Aspergillaceae |
| ASV12 | ITS-Behnsen-MY-35 | 0.16 | None | k__Fungi | p__Ascomycota | c__Sordariomycetes | o__Diaporthales | f__Diaporthaceae |
| ASV7 | ITS-Behnsen-MY-35 | 0.06 | None | k__Fungi | p__Ascomycota | c__Dothideomycetes | o__Pleosporales | f__Pleosporaceae |
| ASV4 | ITS-Behnsen-MY-35 | 0.06 | None | k__Fungi | p__Ascomycota | c__Dothideomycetes | o__Capnodiales | f__Cladosporiaceae |
| ASV14 | ITS-Behnsen-MY-35 | 0.04 | None | k__Fungi | p__Basidiomycota | c__Tremellomycetes | o__Tremellales | f__Bulleribasidiaceae |
| ASV1 | ITS-Behnsen-MY-35 | 0.02 | None | k__Fungi | p__Ascomycota | c__Saccharomycetes | o__Saccharomycetales | f__Saccharomycetales_fam_Incertae_sedis |
| ASV8 | ITS-Behnsen-MY-35 | 0.02 | None | k__Fungi | p__Ascomycota | c__Dothideomycetes | o__Capnodiales | f__Mycosphaerellaceae |
| ASV16 | ITS-Behnsen-MY-35 | 0.01 | None | k__Fungi | p__Basidiomycota | c__Wallemiomycetes | o__Wallemiales | f__Wallemiaceae |
| ASV24 | ITS-Behnsen-MY-35 | 0.01 | None | k__Fungi | p__Basidiomycota | c__Microbotryomycetes | o__Sporidiobolales | f__Sporidiobolaceae |
| ASV44 | ITS-Behnsen-MY-35 | 0.01 | None | k__Fungi | p__Basidiomycota | c__Tremellomycetes | o__Filobasidiales | f__Filobasidiaceae |
| ASV159 | ITS-Behnsen-MY-35 | 0.00 | None | k__Fungi | p__Mucoromycota | c__Mucoromycetes | o__Mucorales | f__Lichtheimiaceae |
| ASV11 | ITS-Behnsen-MY-35 | 0.00 | None | k__Fungi | p__Ascomycota | c__Eurotiomycetes | o__Eurotiales | f__Trichocomaceae |
| ASV28 | ITS-Behnsen-MY-35 | 0.00 | None | k__Fungi | p__Ascomycota | c__Sordariomycetes | o__Microascales | f__Microascaceae |
| ASV3 | ITS-Behnsen-MY-35 | 0.00 | None | k__Fungi | p__Ascomycota | c__Dothideomycetes | o__Pleosporales | f__Didymellaceae |
| ASV30 | ITS-Behnsen-MY-35 | 0.00 | None | k__Fungi | p__Ascomycota | c__Leotiomycetes | o__Helotiales | f__Myxotrichaceae |
| ASV39 | ITS-Behnsen-MY-35 | 0.00 | None | k__Fungi | p__Ascomycota | c__Sordariomycetes | o__Hypocreales | f__Hypocreaceae |
| ASV5 | ITS-Behnsen-MY-35 | 0.00 | None | k__Fungi | p__Basidiomycota | c__Ustilaginomycetes | o__Ustilaginales | f__Ustilaginaceae |
| ASV2 | ITS-Behnsen-MY-36 | 0.52 | None | k__Fungi | p__Ascomycota | c__Eurotiomycetes | o__Eurotiales | f__Aspergillaceae |
| ASV7 | ITS-Behnsen-MY-36 | 0.14 | None | k__Fungi | p__Ascomycota | c__Dothideomycetes | o__Pleosporales | f__Pleosporaceae |
| ASV14 | ITS-Behnsen-MY-36 | 0.12 | None | k__Fungi | p__Basidiomycota | c__Tremellomycetes | o__Tremellales | f__Bulleribasidiaceae |
| ASV8 | ITS-Behnsen-MY-36 | 0.05 | None | k__Fungi | p__Ascomycota | c__Dothideomycetes | o__Capnodiales | f__Mycosphaerellaceae |
| ASV12 | ITS-Behnsen-MY-36 | 0.05 | None | k__Fungi | p__Ascomycota | c__Sordariomycetes | o__Diaporthales | f__Diaporthaceae |
| ASV3 | ITS-Behnsen-MY-36 | 0.04 | None | k__Fungi | p__Ascomycota | c__Dothideomycetes | o__Pleosporales | f__Didymellaceae |
| ASV44 | ITS-Behnsen-MY-36 | 0.02 | None | k__Fungi | p__Basidiomycota | c__Tremellomycetes | o__Filobasidiales | f__Filobasidiaceae |

| | | | | | | | | |
|---|---|---|---|---|---|---|---|---|
| ASV16 | ITS-Behnsen-MY-36 | 0.02 | None | k__Fungi | p__Basidiomycota | c__Wallemiomycetes | o__Wallemiales | f__Wallemiaceae |
| ASV4 | ITS-Behnsen-MY-36 | 0.02 | None | k__Fungi | p__Ascomycota | c__Dothideomycetes | o__Capnodiales | f__Cladosporiaceae |
| ASV30 | ITS-Behnsen-MY-36 | 0.01 | None | k__Fungi | p__Ascomycota | c__Leotiomycetes | o__Helotiales | f__Myxotrichaceae |
| ASV24 | ITS-Behnsen-MY-36 | 0.01 | None | k__Fungi | p__Basidiomycota | c__Microbotryomycetes | o__Sporidiobolales | f__Sporidiobolaceae |
| ASV5 | ITS-Behnsen-MY-36 | 0.00 | None | k__Fungi | p__Basidiomycota | c__Ustilaginomycetes | o__Ustilaginales | f__Ustilaginaceae |
| ASV1 | ITS-Behnsen-MY-36 | 0.00 | None | k__Fungi | p__Ascomycota | c__Saccharomycetes | o__Saccharomycetales | f__Saccharomycetales_fam_Incertae_sedis |
| ASV11 | ITS-Behnsen-MY-36 | 0.00 | None | k__Fungi | p__Ascomycota | c__Eurotiomycetes | o__Eurotiales | f__Trichocomaceae |
| ASV159 | ITS-Behnsen-MY-36 | 0.00 | None | k__Fungi | p__Mucoromycota | c__Mucoromycetes | o__Mucorales | f__Lichtheimiaceae |
| ASV28 | ITS-Behnsen-MY-36 | 0.00 | None | k__Fungi | p__Ascomycota | c__Sordariomycetes | o__Microascales | f__Microascaceae |
| ASV39 | ITS-Behnsen-MY-36 | 0.00 | None | k__Fungi | p__Ascomycota | c__Sordariomycetes | o__Hypocreales | f__Hypocreaceae |
| ASV2 | ITS-Behnsen-MY-37 | 0.58 | None | k__Fungi | p__Ascomycota | c__Eurotiomycetes | o__Eurotiales | f__Aspergillaceae |
| ASV16 | ITS-Behnsen-MY-37 | 0.17 | None | k__Fungi | p__Basidiomycota | c__Wallemiomycetes | o__Wallemiales | f__Wallemiaceae |
| ASV7 | ITS-Behnsen-MY-37 | 0.07 | None | k__Fungi | p__Ascomycota | c__Dothideomycetes | o__Pleosporales | f__Pleosporaceae |
| ASV14 | ITS-Behnsen-MY-37 | 0.05 | None | k__Fungi | p__Basidiomycota | c__Tremellomycetes | o__Tremellales | f__Bulleribasidiaceae |
| ASV4 | ITS-Behnsen-MY-37 | 0.05 | None | k__Fungi | p__Ascomycota | c__Dothideomycetes | o__Capnodiales | f__Cladosporiaceae |
| ASV39 | ITS-Behnsen-MY-37 | 0.02 | None | k__Fungi | p__Ascomycota | c__Sordariomycetes | o__Hypocreales | f__Hypocreaceae |
| ASV12 | ITS-Behnsen-MY-37 | 0.02 | None | k__Fungi | p__Ascomycota | c__Sordariomycetes | o__Diaporthales | f__Diaporthaceae |
| ASV11 | ITS-Behnsen-MY-37 | 0.01 | None | k__Fungi | p__Ascomycota | c__Eurotiomycetes | o__Eurotiales | f__Trichocomaceae |
| ASV8 | ITS-Behnsen-MY-37 | 0.01 | None | k__Fungi | p__Ascomycota | c__Dothideomycetes | o__Capnodiales | f__Mycosphaerellaceae |
| ASV24 | ITS-Behnsen-MY-37 | 0.01 | None | k__Fungi | p__Basidiomycota | c__Microbotryomycetes | o__Sporidiobolales | f__Sporidiobolaceae |
| ASV159 | ITS-Behnsen-MY-37 | 0.01 | None | k__Fungi | p__Mucoromycota | c__Mucoromycetes | o__Mucorales | f__Lichtheimiaceae |
| ASV44 | ITS-Behnsen-MY-37 | 0.00 | None | k__Fungi | p__Basidiomycota | c__Tremellomycetes | o__Filobasidiales | f__Filobasidiaceae |
| ASV3 | ITS-Behnsen-MY-37 | 0.00 | None | k__Fungi | p__Ascomycota | c__Dothideomycetes | o__Pleosporales | f__Didymellaceae |
| ASV1 | ITS-Behnsen-MY-37 | 0.00 | None | k__Fungi | p__Ascomycota | c__Saccharomycetes | o__Saccharomycetales | f__Saccharomycetales_fam_Incertae_sedis |
| ASV30 | ITS-Behnsen-MY-37 | 0.00 | None | k__Fungi | p__Ascomycota | c__Leotiomycetes | o__Helotiales | f__Myxotrichaceae |
| ASV5 | ITS-Behnsen-MY-37 | 0.00 | None | k__Fungi | p__Basidiomycota | c__Ustilaginomycetes | o__Ustilaginales | f__Ustilaginaceae |
| ASV28 | ITS-Behnsen-MY-37 | 0.00 | None | k__Fungi | p__Ascomycota | c__Sordariomycetes | o__Microascales | f__Microascaceae |
| ASV2 | ITS-Behnsen-MY-38 | 0.54 | None | k__Fungi | p__Ascomycota | c__Eurotiomycetes | o__Eurotiales | f__Aspergillaceae |
| ASV14 | ITS-Behnsen-MY-38 | 0.19 | None | k__Fungi | p__Basidiomycota | c__Tremellomycetes | o__Tremellales | f__Bulleribasidiaceae |
| ASV7 | ITS-Behnsen-MY-38 | 0.06 | None | k__Fungi | p__Ascomycota | c__Dothideomycetes | o__Pleosporales | f__Pleosporaceae |
| ASV4 | ITS-Behnsen-MY-38 | 0.04 | None | k__Fungi | p__Ascomycota | c__Dothideomycetes | o__Capnodiales | f__Cladosporiaceae |
| ASV3 | ITS-Behnsen-MY-38 | 0.04 | None | k__Fungi | p__Ascomycota | c__Dothideomycetes | o__Pleosporales | f__Didymellaceae |
| ASV12 | ITS-Behnsen-MY-38 | 0.03 | None | k__Fungi | p__Ascomycota | c__Sordariomycetes | o__Diaporthales | f__Diaporthaceae |
| ASV24 | ITS-Behnsen-MY-38 | 0.03 | None | k__Fungi | p__Basidiomycota | c__Microbotryomycetes | o__Sporidiobolales | f__Sporidiobolaceae |
| ASV16 | ITS-Behnsen-MY-38 | 0.02 | None | k__Fungi | p__Basidiomycota | c__Wallemiomycetes | o__Wallemiales | f__Wallemiaceae |
| ASV11 | ITS-Behnsen-MY-38 | 0.01 | None | k__Fungi | p__Ascomycota | c__Eurotiomycetes | o__Eurotiales | f__Trichocomaceae |
| ASV8 | ITS-Behnsen-MY-38 | 0.01 | None | k__Fungi | p__Ascomycota | c__Dothideomycetes | o__Capnodiales | f__Mycosphaerellaceae |
| ASV30 | ITS-Behnsen-MY-38 | 0.01 | None | k__Fungi | p__Ascomycota | c__Leotiomycetes | o__Helotiales | f__Myxotrichaceae |
| ASV44 | ITS-Behnsen-MY-38 | 0.01 | None | k__Fungi | p__Basidiomycota | c__Tremellomycetes | o__Filobasidiales | f__Filobasidiaceae |
| ASV159 | ITS-Behnsen-MY-38 | 0.01 | None | k__Fungi | p__Mucoromycota | c__Mucoromycetes | o__Mucorales | f__Lichtheimiaceae |
| ASV1 | ITS-Behnsen-MY-38 | 0.00 | None | k__Fungi | p__Ascomycota | c__Saccharomycetes | o__Saccharomycetales | f__Saccharomycetales_fam_Incertae_sedis |
| ASV5 | ITS-Behnsen-MY-38 | 0.00 | None | k__Fungi | p__Basidiomycota | c__Ustilaginomycetes | o__Ustilaginales | f__Ustilaginaceae |
| ASV28 | ITS-Behnsen-MY-38 | 0.00 | None | k__Fungi | p__Ascomycota | c__Sordariomycetes | o__Microascales | f__Microascaceae |
| ASV39 | ITS-Behnsen-MY-38 | 0.00 | None | k__Fungi | p__Ascomycota | c__Sordariomycetes | o__Hypocreales | f__Hypocreaceae |
| ASV2 | ITS-Behnsen-MY-39 | 0.51 | None | k__Fungi | p__Ascomycota | c__Eurotiomycetes | o__Eurotiales | f__Aspergillaceae |
| ASV7 | ITS-Behnsen-MY-39 | 0.12 | None | k__Fungi | p__Ascomycota | c__Dothideomycetes | o__Pleosporales | f__Pleosporaceae |
| ASV12 | ITS-Behnsen-MY-39 | 0.08 | None | k__Fungi | p__Ascomycota | c__Sordariomycetes | o__Diaporthales | f__Diaporthaceae |
| ASV8 | ITS-Behnsen-MY-39 | 0.08 | None | k__Fungi | p__Ascomycota | c__Dothideomycetes | o__Capnodiales | f__Mycosphaerellaceae |
| ASV30 | ITS-Behnsen-MY-39 | 0.07 | None | k__Fungi | p__Ascomycota | c__Leotiomycetes | o__Helotiales | f__Myxotrichaceae |
| ASV14 | ITS-Behnsen-MY-39 | 0.06 | None | k__Fungi | p__Basidiomycota | c__Tremellomycetes | o__Tremellales | f__Bulleribasidiaceae |
| ASV4 | ITS-Behnsen-MY-39 | 0.03 | None | k__Fungi | p__Ascomycota | c__Dothideomycetes | o__Capnodiales | f__Cladosporiaceae |
| ASV24 | ITS-Behnsen-MY-39 | 0.03 | None | k__Fungi | p__Basidiomycota | c__Microbotryomycetes | o__Sporidiobolales | f__Sporidiobolaceae |
| ASV3 | ITS-Behnsen-MY-39 | 0.02 | None | k__Fungi | p__Ascomycota | c__Dothideomycetes | o__Pleosporales | f__Didymellaceae |
| ASV1 | ITS-Behnsen-MY-39 | 0.00 | None | k__Fungi | p__Ascomycota | c__Saccharomycetes | o__Saccharomycetales | f__Saccharomycetales_fam_Incertae_sedis |
| ASV11 | ITS-Behnsen-MY-39 | 0.00 | None | k__Fungi | p__Ascomycota | c__Eurotiomycetes | o__Eurotiales | f__Trichocomaceae |
| ASV16 | ITS-Behnsen-MY-39 | 0.00 | None | k__Fungi | p__Basidiomycota | c__Wallemiomycetes | o__Wallemiales | f__Wallemiaceae |
| ASV44 | ITS-Behnsen-MY-39 | 0.00 | None | k__Fungi | p__Basidiomycota | c__Tremellomycetes | o__Filobasidiales | f__Filobasidiaceae |
| ASV5 | ITS-Behnsen-MY-39 | 0.00 | None | k__Fungi | p__Basidiomycota | c__Ustilaginomycetes | o__Ustilaginales | f__Ustilaginaceae |
| ASV159 | ITS-Behnsen-MY-39 | 0.00 | None | k__Fungi | p__Mucoromycota | c__Mucoromycetes | o__Mucorales | f__Lichtheimiaceae |
| ASV28 | ITS-Behnsen-MY-39 | 0.00 | None | k__Fungi | p__Ascomycota | c__Sordariomycetes | o__Microascales | f__Microascaceae |
| ASV39 | ITS-Behnsen-MY-39 | 0.00 | None | k__Fungi | p__Ascomycota | c__Sordariomycetes | o__Hypocreales | f__Hypocreaceae |
| ASV2 | ITS-Behnsen-MY-4 | 0.28 | None | k__Fungi | p__Ascomycota | c__Eurotiomycetes | o__Eurotiales | f__Aspergillaceae |
| ASV3 | ITS-Behnsen-MY-4 | 0.24 | None | k__Fungi | p__Ascomycota | c__Dothideomycetes | o__Pleosporales | f__Didymellaceae |
| ASV5 | ITS-Behnsen-MY-4 | 0.23 | None | k__Fungi | p__Basidiomycota | c__Ustilaginomycetes | o__Ustilaginales | f__Ustilaginaceae |
| ASV4 | ITS-Behnsen-MY-4 | 0.09 | None | k__Fungi | p__Ascomycota | c__Dothideomycetes | o__Capnodiales | f__Cladosporiaceae |
| ASV11 | ITS-Behnsen-MY-4 | 0.05 | None | k__Fungi | p__Ascomycota | c__Eurotiomycetes | o__Eurotiales | f__Trichocomaceae |
| ASV14 | ITS-Behnsen-MY-4 | 0.04 | None | k__Fungi | p__Basidiomycota | c__Tremellomycetes | o__Tremellales | f__Bulleribasidiaceae |
| ASV7 | ITS-Behnsen-MY-4 | 0.03 | None | k__Fungi | p__Ascomycota | c__Dothideomycetes | o__Pleosporales | f__Pleosporaceae |
| ASV8 | ITS-Behnsen-MY-4 | 0.02 | None | k__Fungi | p__Ascomycota | c__Dothideomycetes | o__Capnodiales | f__Mycosphaerellaceae |
| ASV16 | ITS-Behnsen-MY-4 | 0.02 | None | k__Fungi | p__Basidiomycota | c__Wallemiomycetes | o__Wallemiales | f__Wallemiaceae |
| ASV30 | ITS-Behnsen-MY-4 | 0.00 | None | k__Fungi | p__Ascomycota | c__Leotiomycetes | o__Helotiales | f__Myxotrichaceae |
| ASV24 | ITS-Behnsen-MY-4 | 0.00 | None | k__Fungi | p__Basidiomycota | c__Microbotryomycetes | o__Sporidiobolales | f__Sporidiobolaceae |
| ASV39 | ITS-Behnsen-MY-4 | 0.00 | None | k__Fungi | p__Ascomycota | c__Sordariomycetes | o__Hypocreales | f__Hypocreaceae |
| ASV44 | ITS-Behnsen-MY-4 | 0.00 | None | k__Fungi | p__Basidiomycota | c__Tremellomycetes | o__Filobasidiales | f__Filobasidiaceae |
| ASV28 | ITS-Behnsen-MY-4 | 0.00 | None | k__Fungi | p__Ascomycota | c__Sordariomycetes | o__Microascales | f__Microascaceae |
| ASV1 | ITS-Behnsen-MY-4 | 0.00 | None | k__Fungi | p__Ascomycota | c__Saccharomycetes | o__Saccharomycetales | f__Saccharomycetales_fam_Incertae_sedis |
| ASV12 | ITS-Behnsen-MY-4 | 0.00 | None | k__Fungi | p__Ascomycota | c__Sordariomycetes | o__Diaporthales | f__Diaporthaceae |
| ASV159 | ITS-Behnsen-MY-4 | 0.00 | None | k__Fungi | p__Mucoromycota | c__Mucoromycetes | o__Mucorales | f__Lichtheimiaceae |

| | | | | | | | | |
|---|---|---|---|---|---|---|---|---|
| ASV2 | ITS-Behnsen-MY-5 | 0.38 | None | k__Fungi | p__Ascomycota | c__Eurotiomycetes | o__Eurotiales | f__Aspergillaceae |
| ASV5 | ITS-Behnsen-MY-5 | 0.22 | None | k__Fungi | p__Basidiomycota | c__Ustilaginomycetes | o__Ustilaginales | f__Ustilaginaceae |
| ASV3 | ITS-Behnsen-MY-5 | 0.12 | None | k__Fungi | p__Ascomycota | c__Dothideomycetes | o__Pleosporales | f__Didymellaceae |
| ASV4 | ITS-Behnsen-MY-5 | 0.09 | None | k__Fungi | p__Ascomycota | c__Dothideomycetes | o__Capnodiales | f__Cladosporiaceae |
| ASV16 | ITS-Behnsen-MY-5 | 0.05 | None | k__Fungi | p__Basidiomycota | c__Wallemiomycetes | o__Wallemiales | f__Wallemiaceae |
| ASV14 | ITS-Behnsen-MY-5 | 0.05 | None | k__Fungi | p__Basidiomycota | c__Tremellomycetes | o__Tremellales | f__Bulleribasidiaceae |
| ASV7 | ITS-Behnsen-MY-5 | 0.03 | None | k__Fungi | p__Ascomycota | c__Dothideomycetes | o__Pleosporales | f__Pleosporaceae |
| ASV8 | ITS-Behnsen-MY-5 | 0.02 | None | k__Fungi | p__Ascomycota | c__Dothideomycetes | o__Capnodiales | f__Mycosphaerellaceae |
| ASV11 | ITS-Behnsen-MY-5 | 0.02 | None | k__Fungi | p__Ascomycota | c__Eurotiomycetes | o__Eurotiales | f__Trichocomaceae |
| ASV30 | ITS-Behnsen-MY-5 | 0.01 | None | k__Fungi | p__Ascomycota | c__Leotiomycetes | o__Helotiales | f__Myxotrichaceae |
| ASV24 | ITS-Behnsen-MY-5 | 0.00 | None | k__Fungi | p__Basidiomycota | c__Microbotryomycetes | o__Sporidiobolales | f__Sporidiobolaceae |
| ASV12 | ITS-Behnsen-MY-5 | 0.00 | None | k__Fungi | p__Ascomycota | c__Sordariomycetes | o__Diaporthales | f__Diaporthaceae |
| ASV159 | ITS-Behnsen-MY-5 | 0.00 | None | k__Fungi | p__Mucoromycota | c__Mucoromycetes | o__Mucorales | f__Lichtheimiaceae |
| ASV39 | ITS-Behnsen-MY-5 | 0.00 | None | k__Fungi | p__Ascomycota | c__Sordariomycetes | o__Hypocreales | f__Hypocreaceae |
| ASV1 | ITS-Behnsen-MY-5 | 0.00 | None | k__Fungi | p__Ascomycota | c__Saccharomycetes | o__Saccharomycetales | f__Saccharomycetales_fam_Incertae_sedis |
| ASV28 | ITS-Behnsen-MY-5 | 0.00 | None | k__Fungi | p__Ascomycota | c__Sordariomycetes | o__Microascales | f__Microascaceae |
| ASV44 | ITS-Behnsen-MY-5 | 0.00 | None | k__Fungi | p__Basidiomycota | c__Tremellomycetes | o__Filobasidiales | f__Filobasidiaceae |
| ASV2 | ITS-Behnsen-MY-6 | 0.32 | None | k__Fungi | p__Ascomycota | c__Eurotiomycetes | o__Eurotiales | f__Aspergillaceae |
| ASV3 | ITS-Behnsen-MY-6 | 0.24 | None | k__Fungi | p__Ascomycota | c__Dothideomycetes | o__Pleosporales | f__Didymellaceae |
| ASV5 | ITS-Behnsen-MY-6 | 0.15 | None | k__Fungi | p__Basidiomycota | c__Ustilaginomycetes | o__Ustilaginales | f__Ustilaginaceae |
| ASV4 | ITS-Behnsen-MY-6 | 0.08 | None | k__Fungi | p__Ascomycota | c__Dothideomycetes | o__Capnodiales | f__Cladosporiaceae |
| ASV11 | ITS-Behnsen-MY-6 | 0.06 | None | k__Fungi | p__Ascomycota | c__Eurotiomycetes | o__Eurotiales | f__Trichocomaceae |
| ASV12 | ITS-Behnsen-MY-6 | 0.04 | None | k__Fungi | p__Ascomycota | c__Sordariomycetes | o__Diaporthales | f__Diaporthaceae |
| ASV16 | ITS-Behnsen-MY-6 | 0.03 | None | k__Fungi | p__Basidiomycota | c__Wallemiomycetes | o__Wallemiales | f__Wallemiaceae |
| ASV14 | ITS-Behnsen-MY-6 | 0.03 | None | k__Fungi | p__Basidiomycota | c__Tremellomycetes | o__Tremellales | f__Bulleribasidiaceae |
| ASV8 | ITS-Behnsen-MY-6 | 0.02 | None | k__Fungi | p__Ascomycota | c__Dothideomycetes | o__Capnodiales | f__Mycosphaerellaceae |
| ASV7 | ITS-Behnsen-MY-6 | 0.01 | None | k__Fungi | p__Ascomycota | c__Dothideomycetes | o__Pleosporales | f__Pleosporaceae |
| ASV39 | ITS-Behnsen-MY-6 | 0.01 | None | k__Fungi | p__Ascomycota | c__Sordariomycetes | o__Hypocreales | f__Hypocreaceae |
| ASV24 | ITS-Behnsen-MY-6 | 0.00 | None | k__Fungi | p__Basidiomycota | c__Microbotryomycetes | o__Sporidiobolales | f__Sporidiobolaceae |
| ASV28 | ITS-Behnsen-MY-6 | 0.00 | None | k__Fungi | p__Ascomycota | c__Sordariomycetes | o__Microascales | f__Microascaceae |
| ASV30 | ITS-Behnsen-MY-6 | 0.00 | None | k__Fungi | p__Ascomycota | c__Leotiomycetes | o__Helotiales | f__Myxotrichaceae |
| ASV44 | ITS-Behnsen-MY-6 | 0.00 | None | k__Fungi | p__Basidiomycota | c__Tremellomycetes | o__Filobasidiales | f__Filobasidiaceae |
| ASV159 | ITS-Behnsen-MY-6 | 0.00 | None | k__Fungi | p__Mucoromycota | c__Mucoromycetes | o__Mucorales | f__Lichtheimiaceae |
| ASV1 | ITS-Behnsen-MY-6 | 0.00 | None | k__Fungi | p__Ascomycota | c__Saccharomycetes | o__Saccharomycetales | f__Saccharomycetales_fam_Incertae_sedis |
| ASV2 | ITS-Behnsen-MY-48 | 0.34 | Salmonella | k__Fungi | p__Ascomycota | c__Eurotiomycetes | o__Eurotiales | f__Aspergillaceae |
| ASV5 | ITS-Behnsen-MY-48 | 0.28 | Salmonella | k__Fungi | p__Basidiomycota | c__Ustilaginomycetes | o__Ustilaginales | f__Ustilaginaceae |
| ASV3 | ITS-Behnsen-MY-48 | 0.21 | Salmonella | k__Fungi | p__Ascomycota | c__Dothideomycetes | o__Pleosporales | f__Didymellaceae |
| ASV4 | ITS-Behnsen-MY-48 | 0.06 | Salmonella | k__Fungi | p__Ascomycota | c__Dothideomycetes | o__Capnodiales | f__Cladosporiaceae |
| ASV8 | ITS-Behnsen-MY-48 | 0.06 | Salmonella | k__Fungi | p__Ascomycota | c__Dothideomycetes | o__Capnodiales | f__Mycosphaerellaceae |
| ASV16 | ITS-Behnsen-MY-48 | 0.02 | Salmonella | k__Fungi | p__Basidiomycota | c__Wallemiomycetes | o__Wallemiales | f__Wallemiaceae |
| ASV7 | ITS-Behnsen-MY-48 | 0.01 | Salmonella | k__Fungi | p__Ascomycota | c__Dothideomycetes | o__Pleosporales | f__Pleosporaceae |
| ASV17 | ITS-Behnsen-MY-48 | 0.01 | Salmonella | k__Fungi | p__Basidiomycota | c__Tremellomycetes | o__Tremellales | f__Bulleribasidiaceae |
| ASV11 | ITS-Behnsen-MY-48 | 0.01 | Salmonella | k__Fungi | p__Ascomycota | c__Eurotiomycetes | o__Eurotiales | f__Trichocomaceae |
| ASV1 | ITS-Behnsen-MY-48 | 0.00 | Salmonella | k__Fungi | p__Ascomycota | c__Saccharomycetes | o__Saccharomycetales | f__Saccharomycetales_fam_Incertae_sedis |
| ASV44 | ITS-Behnsen-MY-48 | 0.00 | Salmonella | k__Fungi | p__Basidiomycota | c__Tremellomycetes | o__Filobasidiales | f__Filobasidiaceae |
| ASV159 | ITS-Behnsen-MY-48 | 0.00 | Salmonella | k__Fungi | p__Mucoromycota | c__Mucoromycetes | o__Mucorales | f__Lichtheimiaceae |
| ASV24 | ITS-Behnsen-MY-48 | 0.00 | Salmonella | k__Fungi | p__Basidiomycota | c__Microbotryomycetes | o__Sporidiobolales | f__Sporidiobolaceae |
| ASV28 | ITS-Behnsen-MY-48 | 0.00 | Salmonella | k__Fungi | p__Ascomycota | c__Sordariomycetes | o__Microascales | f__Microascaceae |
| ASV30 | ITS-Behnsen-MY-48 | 0.00 | Salmonella | k__Fungi | p__Ascomycota | c__Leotiomycetes | o__Helotiales | f__Myxotrichaceae |
| ASV39 | ITS-Behnsen-MY-48 | 0.00 | Salmonella | k__Fungi | p__Ascomycota | c__Sordariomycetes | o__Hypocreales | f__Hypocreaceae |
| ASV2 | ITS-Behnsen-MY-49 | 0.29 | Salmonella | k__Fungi | p__Ascomycota | c__Eurotiomycetes | o__Eurotiales | f__Aspergillaceae |
| ASV5 | ITS-Behnsen-MY-49 | 0.28 | Salmonella | k__Fungi | p__Basidiomycota | c__Ustilaginomycetes | o__Ustilaginales | f__Ustilaginaceae |
| ASV4 | ITS-Behnsen-MY-49 | 0.20 | Salmonella | k__Fungi | p__Ascomycota | c__Dothideomycetes | o__Capnodiales | f__Cladosporiaceae |
| ASV11 | ITS-Behnsen-MY-49 | 0.06 | Salmonella | k__Fungi | p__Ascomycota | c__Eurotiomycetes | o__Eurotiales | f__Trichocomaceae |
| ASV17 | ITS-Behnsen-MY-49 | 0.06 | Salmonella | k__Fungi | p__Basidiomycota | c__Tremellomycetes | o__Tremellales | f__Bulleribasidiaceae |
| ASV7 | ITS-Behnsen-MY-49 | 0.03 | Salmonella | k__Fungi | p__Ascomycota | c__Dothideomycetes | o__Pleosporales | f__Pleosporaceae |
| ASV3 | ITS-Behnsen-MY-49 | 0.02 | Salmonella | k__Fungi | p__Ascomycota | c__Dothideomycetes | o__Pleosporales | f__Didymellaceae |
| ASV30 | ITS-Behnsen-MY-49 | 0.02 | Salmonella | k__Fungi | p__Ascomycota | c__Leotiomycetes | o__Helotiales | f__Myxotrichaceae |
| ASV16 | ITS-Behnsen-MY-49 | 0.02 | Salmonella | k__Fungi | p__Basidiomycota | c__Wallemiomycetes | o__Wallemiales | f__Wallemiaceae |
| ASV8 | ITS-Behnsen-MY-49 | 0.02 | Salmonella | k__Fungi | p__Ascomycota | c__Dothideomycetes | o__Capnodiales | f__Mycosphaerellaceae |
| ASV24 | ITS-Behnsen-MY-49 | 0.01 | Salmonella | k__Fungi | p__Basidiomycota | c__Microbotryomycetes | o__Sporidiobolales | f__Sporidiobolaceae |
| ASV28 | ITS-Behnsen-MY-49 | 0.00 | Salmonella | k__Fungi | p__Ascomycota | c__Sordariomycetes | o__Microascales | f__Microascaceae |
| ASV1 | ITS-Behnsen-MY-49 | 0.00 | Salmonella | k__Fungi | p__Ascomycota | c__Saccharomycetes | o__Saccharomycetales | f__Saccharomycetales_fam_Incertae_sedis |
| ASV159 | ITS-Behnsen-MY-49 | 0.00 | Salmonella | k__Fungi | p__Mucoromycota | c__Mucoromycetes | o__Mucorales | f__Lichtheimiaceae |
| ASV39 | ITS-Behnsen-MY-49 | 0.00 | Salmonella | k__Fungi | p__Ascomycota | c__Sordariomycetes | o__Hypocreales | f__Hypocreaceae |
| ASV44 | ITS-Behnsen-MY-49 | 0.00 | Salmonella | k__Fungi | p__Basidiomycota | c__Tremellomycetes | o__Filobasidiales | f__Filobasidiaceae |
| ASV4 | ITS-Behnsen-MY-50 | 0.92 | Salmonella | k__Fungi | p__Ascomycota | c__Dothideomycetes | o__Capnodiales | f__Cladosporiaceae |
| ASV2 | ITS-Behnsen-MY-50 | 0.08 | Salmonella | k__Fungi | p__Ascomycota | c__Eurotiomycetes | o__Eurotiales | f__Aspergillaceae |
| ASV17 | ITS-Behnsen-MY-50 | 0.00 | Salmonella | k__Fungi | p__Basidiomycota | c__Tremellomycetes | o__Tremellales | f__Bulleribasidiaceae |
| ASV3 | ITS-Behnsen-MY-50 | 0.00 | Salmonella | k__Fungi | p__Ascomycota | c__Dothideomycetes | o__Pleosporales | f__Didymellaceae |
| ASV1 | ITS-Behnsen-MY-50 | 0.00 | Salmonella | k__Fungi | p__Ascomycota | c__Saccharomycetes | o__Saccharomycetales | f__Saccharomycetales_fam_Incertae_sedis |
| ASV11 | ITS-Behnsen-MY-50 | 0.00 | Salmonella | k__Fungi | p__Ascomycota | c__Eurotiomycetes | o__Eurotiales | f__Trichocomaceae |
| ASV5 | ITS-Behnsen-MY-50 | 0.00 | Salmonella | k__Fungi | p__Basidiomycota | c__Ustilaginomycetes | o__Ustilaginales | f__Ustilaginaceae |
| ASV159 | ITS-Behnsen-MY-50 | 0.00 | Salmonella | k__Fungi | p__Mucoromycota | c__Mucoromycetes | o__Mucorales | f__Lichtheimiaceae |
| ASV16 | ITS-Behnsen-MY-50 | 0.00 | Salmonella | k__Fungi | p__Basidiomycota | c__Wallemiomycetes | o__Wallemiales | f__Wallemiaceae |
| ASV24 | ITS-Behnsen-MY-50 | 0.00 | Salmonella | k__Fungi | p__Basidiomycota | c__Microbotryomycetes | o__Sporidiobolales | f__Sporidiobolaceae |
| ASV28 | ITS-Behnsen-MY-50 | 0.00 | Salmonella | k__Fungi | p__Ascomycota | c__Sordariomycetes | o__Microascales | f__Microascaceae |
| ASV30 | ITS-Behnsen-MY-50 | 0.00 | Salmonella | k__Fungi | p__Ascomycota | c__Leotiomycetes | o__Helotiales | f__Myxotrichaceae |

| | | | | | | | | |
|---|---|---|---|---|---|---|---|---|
| ASV39 | ITS-Behnsen-MY-50 | 0.00 | Salmonella | k__Fungi | p__Ascomycota | c__Sordariomycetes | o__Hypocreales | f__Hypocreaceae |
| ASV44 | ITS-Behnsen-MY-50 | 0.00 | Salmonella | k__Fungi | p__Basidiomycota | c__Tremellomycetes | o__Filobasidiales | f__Filobasidiaceae |
| ASV7 | ITS-Behnsen-MY-50 | 0.00 | Salmonella | k__Fungi | p__Ascomycota | c__Dothideomycetes | o__Pleosporales | f__Pleosporaceae |
| ASV8 | ITS-Behnsen-MY-50 | 0.00 | Salmonella | k__Fungi | p__Ascomycota | c__Dothideomycetes | o__Capnodiales | f__Mycosphaerellaceae |
| ASV1 | ITS-Behnsen-MY-51 | 0.63 | Salmonella | k__Fungi | p__Ascomycota | c__Saccharomycetes | o__Saccharomycetales | f__Saccharomycetales_fam_Incertae_sedis |
| ASV5 | ITS-Behnsen-MY-51 | 0.12 | Salmonella | k__Fungi | p__Basidiomycota | c__Ustilaginomycetes | o__Ustilaginales | f__Ustilaginaceae |
| ASV2 | ITS-Behnsen-MY-51 | 0.07 | Salmonella | k__Fungi | p__Ascomycota | c__Eurotiomycetes | o__Eurotiales | f__Aspergillaceae |
| ASV8 | ITS-Behnsen-MY-51 | 0.05 | Salmonella | k__Fungi | p__Ascomycota | c__Dothideomycetes | o__Capnodiales | f__Mycosphaerellaceae |
| ASV4 | ITS-Behnsen-MY-51 | 0.05 | Salmonella | k__Fungi | p__Ascomycota | c__Dothideomycetes | o__Capnodiales | f__Cladosporiaceae |
| ASV16 | ITS-Behnsen-MY-51 | 0.02 | Salmonella | k__Fungi | p__Basidiomycota | c__Wallemiomycetes | o__Wallemiales | f__Wallemiaceae |
| ASV3 | ITS-Behnsen-MY-51 | 0.02 | Salmonella | k__Fungi | p__Ascomycota | c__Dothideomycetes | o__Pleosporales | f__Didymellaceae |
| ASV7 | ITS-Behnsen-MY-51 | 0.02 | Salmonella | k__Fungi | p__Ascomycota | c__Dothideomycetes | o__Pleosporales | f__Pleosporaceae |
| ASV17 | ITS-Behnsen-MY-51 | 0.01 | Salmonella | k__Fungi | p__Basidiomycota | c__Tremellomycetes | o__Tremellales | f__Bulleribasidiaceae |
| ASV11 | ITS-Behnsen-MY-51 | 0.00 | Salmonella | k__Fungi | p__Ascomycota | c__Eurotiomycetes | o__Eurotiales | f__Trichocomaceae |
| ASV159 | ITS-Behnsen-MY-51 | 0.00 | Salmonella | k__Fungi | p__Mucoromycota | c__Mucoromycetes | o__Mucorales | f__Lichtheimiaceae |
| ASV24 | ITS-Behnsen-MY-51 | 0.00 | Salmonella | k__Fungi | p__Basidiomycota | c__Microbotryomycetes | o__Sporidiobolales | f__Sporidiobolaceae |
| ASV28 | ITS-Behnsen-MY-51 | 0.00 | Salmonella | k__Fungi | p__Ascomycota | c__Sordariomycetes | o__Microascales | f__Microascaceae |
| ASV30 | ITS-Behnsen-MY-51 | 0.00 | Salmonella | k__Fungi | p__Ascomycota | c__Leotiomycetes | o__Helotiales | f__Myxotrichaceae |
| ASV39 | ITS-Behnsen-MY-51 | 0.00 | Salmonella | k__Fungi | p__Ascomycota | c__Sordariomycetes | o__Hypocreales | f__Hypocreaceae |
| ASV44 | ITS-Behnsen-MY-51 | 0.00 | Salmonella | k__Fungi | p__Basidiomycota | c__Tremellomycetes | o__Filobasidiales | f__Filobasidiaceae |
| ASV1 | ITS-Behnsen-MY-52 | 0.98 | Salmonella | k__Fungi | p__Ascomycota | c__Saccharomycetes | o__Saccharomycetales | f__Saccharomycetales_fam_Incertae_sedis |
| ASV4 | ITS-Behnsen-MY-52 | 0.02 | Salmonella | k__Fungi | p__Ascomycota | c__Dothideomycetes | o__Capnodiales | f__Cladosporiaceae |
| ASV3 | ITS-Behnsen-MY-52 | 0.00 | Salmonella | k__Fungi | p__Ascomycota | c__Dothideomycetes | o__Pleosporales | f__Didymellaceae |
| ASV2 | ITS-Behnsen-MY-52 | 0.00 | Salmonella | k__Fungi | p__Ascomycota | c__Eurotiomycetes | o__Eurotiales | f__Aspergillaceae |
| ASV5 | ITS-Behnsen-MY-52 | 0.00 | Salmonella | k__Fungi | p__Basidiomycota | c__Ustilaginomycetes | o__Ustilaginales | f__Ustilaginaceae |
| ASV11 | ITS-Behnsen-MY-52 | 0.00 | Salmonella | k__Fungi | p__Ascomycota | c__Eurotiomycetes | o__Eurotiales | f__Trichocomaceae |
| ASV159 | ITS-Behnsen-MY-52 | 0.00 | Salmonella | k__Fungi | p__Mucoromycota | c__Mucoromycetes | o__Mucorales | f__Lichtheimiaceae |
| ASV16 | ITS-Behnsen-MY-52 | 0.00 | Salmonella | k__Fungi | p__Basidiomycota | c__Wallemiomycetes | o__Wallemiales | f__Wallemiaceae |
| ASV17 | ITS-Behnsen-MY-52 | 0.00 | Salmonella | k__Fungi | p__Basidiomycota | c__Tremellomycetes | o__Tremellales | f__Bulleribasidiaceae |
| ASV24 | ITS-Behnsen-MY-52 | 0.00 | Salmonella | k__Fungi | p__Basidiomycota | c__Microbotryomycetes | o__Sporidiobolales | f__Sporidiobolaceae |
| ASV28 | ITS-Behnsen-MY-52 | 0.00 | Salmonella | k__Fungi | p__Ascomycota | c__Sordariomycetes | o__Microascales | f__Microascaceae |
| ASV30 | ITS-Behnsen-MY-52 | 0.00 | Salmonella | k__Fungi | p__Ascomycota | c__Leotiomycetes | o__Helotiales | f__Myxotrichaceae |
| ASV39 | ITS-Behnsen-MY-52 | 0.00 | Salmonella | k__Fungi | p__Ascomycota | c__Sordariomycetes | o__Hypocreales | f__Hypocreaceae |
| ASV44 | ITS-Behnsen-MY-52 | 0.00 | Salmonella | k__Fungi | p__Basidiomycota | c__Tremellomycetes | o__Filobasidiales | f__Filobasidiaceae |
| ASV7 | ITS-Behnsen-MY-52 | 0.00 | Salmonella | k__Fungi | p__Ascomycota | c__Dothideomycetes | o__Pleosporales | f__Pleosporaceae |
| ASV8 | ITS-Behnsen-MY-52 | 0.00 | Salmonella | k__Fungi | p__Ascomycota | c__Dothideomycetes | o__Capnodiales | f__Mycosphaerellaceae |
| ASV2 | ITS-Behnsen-MY-53 | 0.27 | Salmonella | k__Fungi | p__Ascomycota | c__Eurotiomycetes | o__Eurotiales | f__Aspergillaceae |
| ASV17 | ITS-Behnsen-MY-53 | 0.15 | Salmonella | k__Fungi | p__Basidiomycota | c__Tremellomycetes | o__Tremellales | f__Bulleribasidiaceae |
| ASV4 | ITS-Behnsen-MY-53 | 0.12 | Salmonella | k__Fungi | p__Ascomycota | c__Dothideomycetes | o__Capnodiales | f__Cladosporiaceae |
| ASV5 | ITS-Behnsen-MY-53 | 0.11 | Salmonella | k__Fungi | p__Basidiomycota | c__Ustilaginomycetes | o__Ustilaginales | f__Ustilaginaceae |
| ASV16 | ITS-Behnsen-MY-53 | 0.10 | Salmonella | k__Fungi | p__Basidiomycota | c__Wallemiomycetes | o__Wallemiales | f__Wallemiaceae |
| ASV44 | ITS-Behnsen-MY-53 | 0.06 | Salmonella | k__Fungi | p__Basidiomycota | c__Tremellomycetes | o__Filobasidiales | f__Filobasidiaceae |
| ASV1 | ITS-Behnsen-MY-53 | 0.05 | Salmonella | k__Fungi | p__Ascomycota | c__Saccharomycetes | o__Saccharomycetales | f__Saccharomycetales_fam_Incertae_sedis |
| ASV3 | ITS-Behnsen-MY-53 | 0.04 | Salmonella | k__Fungi | p__Ascomycota | c__Dothideomycetes | o__Pleosporales | f__Didymellaceae |
| ASV8 | ITS-Behnsen-MY-53 | 0.03 | Salmonella | k__Fungi | p__Ascomycota | c__Dothideomycetes | o__Capnodiales | f__Mycosphaerellaceae |
| ASV30 | ITS-Behnsen-MY-53 | 0.03 | Salmonella | k__Fungi | p__Ascomycota | c__Leotiomycetes | o__Helotiales | f__Myxotrichaceae |
| ASV7 | ITS-Behnsen-MY-53 | 0.02 | Salmonella | k__Fungi | p__Ascomycota | c__Dothideomycetes | o__Pleosporales | f__Pleosporaceae |
| ASV11 | ITS-Behnsen-MY-53 | 0.01 | Salmonella | k__Fungi | p__Ascomycota | c__Eurotiomycetes | o__Eurotiales | f__Trichocomaceae |
| ASV28 | ITS-Behnsen-MY-53 | 0.00 | Salmonella | k__Fungi | p__Ascomycota | c__Sordariomycetes | o__Microascales | f__Microascaceae |
| ASV159 | ITS-Behnsen-MY-53 | 0.00 | Salmonella | k__Fungi | p__Mucoromycota | c__Mucoromycetes | o__Mucorales | f__Lichtheimiaceae |
| ASV24 | ITS-Behnsen-MY-53 | 0.00 | Salmonella | k__Fungi | p__Basidiomycota | c__Microbotryomycetes | o__Sporidiobolales | f__Sporidiobolaceae |
| ASV39 | ITS-Behnsen-MY-53 | 0.00 | Salmonella | k__Fungi | p__Ascomycota | c__Sordariomycetes | o__Hypocreales | f__Hypocreaceae |
| ASV1 | ITS-Behnsen-MY-65 | 0.93 | Salmonella | k__Fungi | p__Ascomycota | c__Saccharomycetes | o__Saccharomycetales | f__Saccharomycetales_fam_Incertae_sedis |
| ASV2 | ITS-Behnsen-MY-65 | 0.07 | Salmonella | k__Fungi | p__Ascomycota | c__Eurotiomycetes | o__Eurotiales | f__Aspergillaceae |
| ASV4 | ITS-Behnsen-MY-65 | 0.00 | Salmonella | k__Fungi | p__Ascomycota | c__Dothideomycetes | o__Capnodiales | f__Cladosporiaceae |
| ASV11 | ITS-Behnsen-MY-65 | 0.00 | Salmonella | k__Fungi | p__Ascomycota | c__Eurotiomycetes | o__Eurotiales | f__Trichocomaceae |
| ASV159 | ITS-Behnsen-MY-65 | 0.00 | Salmonella | k__Fungi | p__Mucoromycota | c__Mucoromycetes | o__Mucorales | f__Lichtheimiaceae |
| ASV16 | ITS-Behnsen-MY-65 | 0.00 | Salmonella | k__Fungi | p__Basidiomycota | c__Wallemiomycetes | o__Wallemiales | f__Wallemiaceae |
| ASV17 | ITS-Behnsen-MY-65 | 0.00 | Salmonella | k__Fungi | p__Basidiomycota | c__Tremellomycetes | o__Tremellales | f__Bulleribasidiaceae |
| ASV24 | ITS-Behnsen-MY-65 | 0.00 | Salmonella | k__Fungi | p__Basidiomycota | c__Microbotryomycetes | o__Sporidiobolales | f__Sporidiobolaceae |
| ASV28 | ITS-Behnsen-MY-65 | 0.00 | Salmonella | k__Fungi | p__Ascomycota | c__Sordariomycetes | o__Microascales | f__Microascaceae |
| ASV3 | ITS-Behnsen-MY-65 | 0.00 | Salmonella | k__Fungi | p__Ascomycota | c__Dothideomycetes | o__Pleosporales | f__Didymellaceae |
| ASV30 | ITS-Behnsen-MY-65 | 0.00 | Salmonella | k__Fungi | p__Ascomycota | c__Leotiomycetes | o__Helotiales | f__Myxotrichaceae |
| ASV39 | ITS-Behnsen-MY-65 | 0.00 | Salmonella | k__Fungi | p__Ascomycota | c__Sordariomycetes | o__Hypocreales | f__Hypocreaceae |
| ASV44 | ITS-Behnsen-MY-65 | 0.00 | Salmonella | k__Fungi | p__Basidiomycota | c__Tremellomycetes | o__Filobasidiales | f__Filobasidiaceae |
| ASV5 | ITS-Behnsen-MY-65 | 0.00 | Salmonella | k__Fungi | p__Basidiomycota | c__Ustilaginomycetes | o__Ustilaginales | f__Ustilaginaceae |
| ASV7 | ITS-Behnsen-MY-65 | 0.00 | Salmonella | k__Fungi | p__Ascomycota | c__Dothideomycetes | o__Pleosporales | f__Pleosporaceae |
| ASV8 | ITS-Behnsen-MY-65 | 0.00 | Salmonella | k__Fungi | p__Ascomycota | c__Dothideomycetes | o__Capnodiales | f__Mycosphaerellaceae |
| ASV1 | ITS-Behnsen-MY-66 | 0.29 | Salmonella | k__Fungi | p__Ascomycota | c__Saccharomycetes | o__Saccharomycetales | f__Saccharomycetales_fam_Incertae_sedis |
| ASV2 | ITS-Behnsen-MY-66 | 0.28 | Salmonella | k__Fungi | p__Ascomycota | c__Eurotiomycetes | o__Eurotiales | f__Aspergillaceae |
| ASV7 | ITS-Behnsen-MY-66 | 0.16 | Salmonella | k__Fungi | p__Ascomycota | c__Dothideomycetes | o__Pleosporales | f__Pleosporaceae |
| ASV3 | ITS-Behnsen-MY-66 | 0.10 | Salmonella | k__Fungi | p__Ascomycota | c__Dothideomycetes | o__Pleosporales | f__Didymellaceae |
| ASV16 | ITS-Behnsen-MY-66 | 0.07 | Salmonella | k__Fungi | p__Basidiomycota | c__Wallemiomycetes | o__Wallemiales | f__Wallemiaceae |
| ASV17 | ITS-Behnsen-MY-66 | 0.07 | Salmonella | k__Fungi | p__Basidiomycota | c__Tremellomycetes | o__Tremellales | f__Bulleribasidiaceae |
| ASV30 | ITS-Behnsen-MY-66 | 0.02 | Salmonella | k__Fungi | p__Ascomycota | c__Leotiomycetes | o__Helotiales | f__Myxotrichaceae |
| ASV4 | ITS-Behnsen-MY-66 | 0.02 | Salmonella | k__Fungi | p__Ascomycota | c__Dothideomycetes | o__Capnodiales | f__Cladosporiaceae |
| ASV24 | ITS-Behnsen-MY-66 | 0.00 | Salmonella | k__Fungi | p__Basidiomycota | c__Microbotryomycetes | o__Sporidiobolales | f__Sporidiobolaceae |
| ASV11 | ITS-Behnsen-MY-66 | 0.00 | Salmonella | k__Fungi | p__Ascomycota | c__Eurotiomycetes | o__Eurotiales | f__Trichocomaceae |

| | | | | | | | | |
|---|---|---|---|---|---|---|---|---|
| ASV159 | ITS-Behnsen-MY-66 | 0.00 | Salmonella | k__Fungi | p__Mucoromycota | c__Mucoromycetes | o__Mucorales | f__Lichtheimiaceae |
| ASV28 | ITS-Behnsen-MY-66 | 0.00 | Salmonella | k__Fungi | p__Ascomycota | c__Sordariomycetes | o__Microascales | f__Microascaceae |
| ASV39 | ITS-Behnsen-MY-66 | 0.00 | Salmonella | k__Fungi | p__Ascomycota | c__Sordariomycetes | o__Hypocreales | f__Hypocreaceae |
| ASV44 | ITS-Behnsen-MY-66 | 0.00 | Salmonella | k__Fungi | p__Basidiomycota | c__Tremellomycetes | o__Filobasidiales | f__Filobasidiaceae |
| ASV5 | ITS-Behnsen-MY-66 | 0.00 | Salmonella | k__Fungi | p__Basidiomycota | c__Ustilaginomycetes | o__Ustilaginales | f__Ustilaginaceae |
| ASV8 | ITS-Behnsen-MY-66 | 0.00 | Salmonella | k__Fungi | p__Ascomycota | c__Dothideomycetes | o__Capnodiales | f__Mycosphaerellaceae |
| ASV1 | ITS-Behnsen-MY-67 | 0.52 | Salmonella | k__Fungi | p__Ascomycota | c__Saccharomycetes | o__Saccharomycetales | f__Saccharomycetales_fam_Incertae_sedis |
| ASV2 | ITS-Behnsen-MY-67 | 0.26 | Salmonella | k__Fungi | p__Ascomycota | c__Eurotiomycetes | o__Eurotiales | f__Aspergillaceae |
| ASV30 | ITS-Behnsen-MY-67 | 0.07 | Salmonella | k__Fungi | p__Ascomycota | c__Leotiomycetes | o__Helotiales | f__Myxotrichaceae |
| ASV4 | ITS-Behnsen-MY-67 | 0.05 | Salmonella | k__Fungi | p__Ascomycota | c__Dothideomycetes | o__Capnodiales | f__Cladosporiaceae |
| ASV7 | ITS-Behnsen-MY-67 | 0.04 | Salmonella | k__Fungi | p__Ascomycota | c__Dothideomycetes | o__Pleosporales | f__Pleosporaceae |
| ASV3 | ITS-Behnsen-MY-67 | 0.03 | Salmonella | k__Fungi | p__Ascomycota | c__Dothideomycetes | o__Pleosporales | f__Didymellaceae |
| ASV28 | ITS-Behnsen-MY-67 | 0.02 | Salmonella | k__Fungi | p__Ascomycota | c__Sordariomycetes | o__Microascales | f__Microascaceae |
| ASV11 | ITS-Behnsen-MY-67 | 0.01 | Salmonella | k__Fungi | p__Ascomycota | c__Eurotiomycetes | o__Eurotiales | f__Trichocomaceae |
| ASV159 | ITS-Behnsen-MY-67 | 0.00 | Salmonella | k__Fungi | p__Mucoromycota | c__Mucoromycetes | o__Mucorales | f__Lichtheimiaceae |
| ASV16 | ITS-Behnsen-MY-67 | 0.00 | Salmonella | k__Fungi | p__Basidiomycota | c__Wallemiomycetes | o__Wallemiales | f__Wallemiaceae |
| ASV17 | ITS-Behnsen-MY-67 | 0.00 | Salmonella | k__Fungi | p__Basidiomycota | c__Tremellomycetes | o__Tremellales | f__Bulleribasidiaceae |
| ASV24 | ITS-Behnsen-MY-67 | 0.00 | Salmonella | k__Fungi | p__Basidiomycota | c__Microbotryomycetes | o__Sporidiobolales | f__Sporidiobolaceae |
| ASV39 | ITS-Behnsen-MY-67 | 0.00 | Salmonella | k__Fungi | p__Ascomycota | c__Sordariomycetes | o__Hypocreales | f__Hypocreaceae |
| ASV44 | ITS-Behnsen-MY-67 | 0.00 | Salmonella | k__Fungi | p__Basidiomycota | c__Tremellomycetes | o__Filobasidiales | f__Filobasidiaceae |
| ASV5 | ITS-Behnsen-MY-67 | 0.00 | Salmonella | k__Fungi | p__Basidiomycota | c__Ustilaginomycetes | o__Ustilaginales | f__Ustilaginaceae |
| ASV8 | ITS-Behnsen-MY-67 | 0.00 | Salmonella | k__Fungi | p__Ascomycota | c__Dothideomycetes | o__Capnodiales | f__Mycosphaerellaceae |
| ASV1 | ITS-Behnsen-MY-68 | 0.67 | Salmonella | k__Fungi | p__Ascomycota | c__Saccharomycetes | o__Saccharomycetales | f__Saccharomycetales_fam_Incertae_sedis |
| ASV2 | ITS-Behnsen-MY-68 | 0.23 | Salmonella | k__Fungi | p__Ascomycota | c__Eurotiomycetes | o__Eurotiales | f__Aspergillaceae |
| ASV17 | ITS-Behnsen-MY-68 | 0.05 | Salmonella | k__Fungi | p__Basidiomycota | c__Tremellomycetes | o__Tremellales | f__Bulleribasidiaceae |
| ASV4 | ITS-Behnsen-MY-68 | 0.03 | Salmonella | k__Fungi | p__Ascomycota | c__Dothideomycetes | o__Capnodiales | f__Cladosporiaceae |
| ASV30 | ITS-Behnsen-MY-68 | 0.01 | Salmonella | k__Fungi | p__Ascomycota | c__Leotiomycetes | o__Helotiales | f__Myxotrichaceae |
| ASV39 | ITS-Behnsen-MY-68 | 0.01 | Salmonella | k__Fungi | p__Ascomycota | c__Sordariomycetes | o__Hypocreales | f__Hypocreaceae |
| ASV7 | ITS-Behnsen-MY-68 | 0.01 | Salmonella | k__Fungi | p__Ascomycota | c__Dothideomycetes | o__Pleosporales | f__Pleosporaceae |
| ASV44 | ITS-Behnsen-MY-68 | 0.00 | Salmonella | k__Fungi | p__Basidiomycota | c__Tremellomycetes | o__Filobasidiales | f__Filobasidiaceae |
| ASV28 | ITS-Behnsen-MY-68 | 0.00 | Salmonella | k__Fungi | p__Ascomycota | c__Sordariomycetes | o__Microascales | f__Microascaceae |
| ASV159 | ITS-Behnsen-MY-68 | 0.00 | Salmonella | k__Fungi | p__Mucoromycota | c__Mucoromycetes | o__Mucorales | f__Lichtheimiaceae |
| ASV11 | ITS-Behnsen-MY-68 | 0.00 | Salmonella | k__Fungi | p__Ascomycota | c__Eurotiomycetes | o__Eurotiales | f__Trichocomaceae |
| ASV16 | ITS-Behnsen-MY-68 | 0.00 | Salmonella | k__Fungi | p__Basidiomycota | c__Wallemiomycetes | o__Wallemiales | f__Wallemiaceae |
| ASV24 | ITS-Behnsen-MY-68 | 0.00 | Salmonella | k__Fungi | p__Basidiomycota | c__Microbotryomycetes | o__Sporidiobolales | f__Sporidiobolaceae |
| ASV3 | ITS-Behnsen-MY-68 | 0.00 | Salmonella | k__Fungi | p__Ascomycota | c__Dothideomycetes | o__Pleosporales | f__Didymellaceae |
| ASV5 | ITS-Behnsen-MY-68 | 0.00 | Salmonella | k__Fungi | p__Basidiomycota | c__Ustilaginomycetes | o__Ustilaginales | f__Ustilaginaceae |
| ASV8 | ITS-Behnsen-MY-68 | 0.00 | Salmonella | k__Fungi | p__Ascomycota | c__Dothideomycetes | o__Capnodiales | f__Mycosphaerellaceae |
| ASV1 | ITS-Behnsen-MY-69 | 0.88 | Salmonella | k__Fungi | p__Ascomycota | c__Saccharomycetes | o__Saccharomycetales | f__Saccharomycetales_fam_Incertae_sedis |
| ASV2 | ITS-Behnsen-MY-69 | 0.12 | Salmonella | k__Fungi | p__Ascomycota | c__Eurotiomycetes | o__Eurotiales | f__Aspergillaceae |
| ASV17 | ITS-Behnsen-MY-69 | 0.00 | Salmonella | k__Fungi | p__Basidiomycota | c__Tremellomycetes | o__Tremellales | f__Bulleribasidiaceae |
| ASV11 | ITS-Behnsen-MY-69 | 0.00 | Salmonella | k__Fungi | p__Ascomycota | c__Eurotiomycetes | o__Eurotiales | f__Trichocomaceae |
| ASV159 | ITS-Behnsen-MY-69 | 0.00 | Salmonella | k__Fungi | p__Mucoromycota | c__Mucoromycetes | o__Mucorales | f__Lichtheimiaceae |
| ASV16 | ITS-Behnsen-MY-69 | 0.00 | Salmonella | k__Fungi | p__Basidiomycota | c__Wallemiomycetes | o__Wallemiales | f__Wallemiaceae |
| ASV24 | ITS-Behnsen-MY-69 | 0.00 | Salmonella | k__Fungi | p__Basidiomycota | c__Microbotryomycetes | o__Sporidiobolales | f__Sporidiobolaceae |
| ASV28 | ITS-Behnsen-MY-69 | 0.00 | Salmonella | k__Fungi | p__Ascomycota | c__Sordariomycetes | o__Microascales | f__Microascaceae |
| ASV3 | ITS-Behnsen-MY-69 | 0.00 | Salmonella | k__Fungi | p__Ascomycota | c__Dothideomycetes | o__Pleosporales | f__Didymellaceae |
| ASV30 | ITS-Behnsen-MY-69 | 0.00 | Salmonella | k__Fungi | p__Ascomycota | c__Leotiomycetes | o__Helotiales | f__Myxotrichaceae |
| ASV39 | ITS-Behnsen-MY-69 | 0.00 | Salmonella | k__Fungi | p__Ascomycota | c__Sordariomycetes | o__Hypocreales | f__Hypocreaceae |
| ASV4 | ITS-Behnsen-MY-69 | 0.00 | Salmonella | k__Fungi | p__Ascomycota | c__Dothideomycetes | o__Capnodiales | f__Cladosporiaceae |
| ASV44 | ITS-Behnsen-MY-69 | 0.00 | Salmonella | k__Fungi | p__Basidiomycota | c__Tremellomycetes | o__Filobasidiales | f__Filobasidiaceae |
| ASV5 | ITS-Behnsen-MY-69 | 0.00 | Salmonella | k__Fungi | p__Basidiomycota | c__Ustilaginomycetes | o__Ustilaginales | f__Ustilaginaceae |
| ASV7 | ITS-Behnsen-MY-69 | 0.00 | Salmonella | k__Fungi | p__Ascomycota | c__Dothideomycetes | o__Pleosporales | f__Pleosporaceae |
| ASV8 | ITS-Behnsen-MY-69 | 0.00 | Salmonella | k__Fungi | p__Ascomycota | c__Dothideomycetes | o__Capnodiales | f__Mycosphaerellaceae |
