## Supplementary Table 2 for "Commensal Yeast Promotes *Salmonella* Typhimurium Virulence"

**Table S2: RNAseq data showing differentially regulated STm genes in co-culture with *C. albicans* ATCC compared to STm alone.**

| Gene ID | gene | product | protein_id | fungus Candida albicans/Untreated : log2FC | fungus Candida albicans/Untreated : PValue | fungus Candida albicans/Untreated : QValue |
|---|---|---|---|---|---|---|
| STM14_RS16155 | | multidrug/biocide efflux PACE transporter | WP_001519547.1 | 5.979134173 | 2.889E-192 | 1.064E-188 |
| STM14_RS26775 | | tail fiber assembly protein | | 5.921577193 | 3.3349E-30 | 2.6694E-28 |
| STM14_RS11795 | | polysaccharide export protein | WP_000978077.1 | 4.673324566 | 6.4886E-52 | 2.1719E-49 |
| STM14_RS26375 | hisL | his operon leader peptide | WP_001364200.1 | 4.479886886 | 4.5018E-85 | 4.1439E-82 |
| STM14_RS00855 | citS | citrate/sodium symporter CitS | WP_000183602.1 | 4.320431755 | 2.8029E-33 | 2.7893E-31 |
| STM14_RS17335 | | tagatose bisphosphate family class II aldolase | WP_000469985.1 | 4.209747869 | 5.4796E-50 | 1.4411E-47 |
| STM14_RS09285 | mpaA | murein tripeptide amidase MpaA | WP_000219126.1 | 3.209412415 | 8.9384E-51 | 2.5316E-48 |
| STM14_RS23410 | arcA | arginine deiminase | WP_000410846.1 | 2.766606832 | 2.5401E-05 | 0.00012454 |
| STM14_RS21515 | menA | 1%2C4-dihydroxy-2-naphthoate polyprenyltransferase | WP_000139639.1 | 2.21694057 | 8.2861E-41 | 1.2204E-38 |
| STM14_RS03765 | tatE | twin-arginine translocase subunit TatE | WP_000503938.1 | 2.195204998 | 3.838E-119 | 7.065E-116 |
| STM14_RS26925 | | hypothetical protein | WP_001670836.1 | 2.125016363 | 5.6509E-13 | 9.5443E-12 |
| STM14_RS07950 | ssaR | SPI-2 type III secretion system export apparatus protein SsaR | WP_000056367.1 | 2.053708078 | 6.7923E-21 | 2.5009E-19 |
| STM14_RS07515 | | hypothetical protein | WP_011233063.1 | 1.991172499 | 3.175E-10 | 3.6533E-09 |
| STM14_RS23390 | | arginine repressor | WP_000666051.1 | 1.979178184 | 6.0945E-05 | 0.000272 |
| STM14_RS08870 | | M15 family metallopeptidase | WP_000237715.1 | 1.944259965 | 7.8648E-22 | 3.3672E-20 |
| STM14_RS26350 | mgtS | protein MgtS | WP_000901531.1 | 1.935738467 | 4.0843E-80 | 3.0077E-77 |
| STM14_RS13795 | ratB | DUF823/DUF824 repeat adhesin RatB | | 1.93150098 | 1.6005E-49 | 3.9288E-47 |
| STM14_RS26335 | | YnhF family membrane protein | WP_000102277.1 | 1.914824763 | 1.6554E-30 | 1.3852E-28 |
| STM14_RS26410 | ypfM | protein YpfM | WP_001386977.1 | 1.858766427 | 6.8399E-78 | 4.1974E-75 |
| STM14_RS10545 | uspC | universal stress protein UspC | WP_000122606.1 | 1.847310894 | 1.4303E-25 | 8.494E-24 |
| STM14_RS23405 | arcC | carbamate kinase | WP_000428751.1 | 1.840984155 | 0.0020123 | 0.00597524 |
| STM14_RS01410 | speD | adenosylmethionine decarboxylase | WP_000734276.1 | 1.834246787 | 2.0566E-14 | 4.2541E-13 |
| STM14_RS11140 | | hypothetical protein | WP_000008351.1 | 1.809350617 | 1.3788E-12 | 2.1695E-11 |
| STM14_RS02015 | tssM | type VI secretion system membrane subunit TssM | WP_001168961.1 | 1.794859255 | 4.7352E-18 | 1.3109E-16 |
| STM14_RS24325 | traY | conjugal transfer relaxosome protein TraY | WP_001676655.1 | 1.768178067 | 2.1405E-18 | 6.2057E-17 |
| STM14_RS06305 | bssS | biofilm formation regulator BssS | WP_000414441.1 | 1.761745031 | 1.0299E-35 | 1.264E-33 |
| STM14_RS10245 | | lytic enzyme | WP_000789530.1 | 1.735208581 | 2.1874E-25 | 1.2784E-23 |
| STM14_RS15535 | invF | type III secretion system transcriptional activator InvF | WP_001674874.1 | 1.699035409 | 5.8129E-51 | 1.7836E-48 |
| STM14_RS09380 | steC | SPI-2 type III secretion system effector kinase SteC | WP_001116926.1 | 1.696865145 | 8.3353E-34 | 9.3002E-32 |
| STM14_RS00090 | pefA | major pilin PefA | WP_000748205.1 | 1.690993356 | 1.7746E-15 | 4.0584E-14 |
| STM14_RS07735 | | major outer membrane lipoprotein | WP_001082332.1 | 1.674524154 | 2.8324E-25 | 1.6044E-23 |
| STM14_RS21065 | | hypothetical protein | WP_001541209.1 | 1.650634974 | 2.9327E-87 | 3.5994E-84 |
| STM14_RS24685 | | PagK family vesicle-borne virulence factor | WP_014344029.1 | 1.617862741 | 2.8965E-52 | 1.0665E-49 |
| STM14_RS14020 | cadA | lysine decarboxylase CadA | WP_001100652.1 | 1.602434407 | 3.5874E-35 | 4.2609E-33 |
| STM14_RS23170 | iolR | myo-inositol utilization transcriptional regulator IolR | WP_000033840.1 | 1.590315118 | 2.308E-21 | 8.8522E-20 |
| STM14_RS25255 | | hypothetical protein | WP_014343883.1 | 1.586918816 | 3.3443E-16 | 8.1548E-15 |
| STM14_RS14345 | | DUF1367 family protein | WP_000929791.1 | 1.579356981 | 1.2982E-06 | 8.143E-06 |
| STM14_RS14205 | | phage tail tape measure protein | WP_000372072.1 | 1.572912284 | 5.6639E-15 | 1.2488E-13 |
| STM14_RS15505 | spaM | SPI-1 type III secretion system protein SpaM | WP_001520714.1 | 1.566014099 | 3.6942E-28 | 2.5665E-26 |
| STM14_RS14025 | dtpD | dipeptide permease DtpD | WP_000856793.1 | 1.545779099 | 1.4155E-06 | 8.8184E-06 |
| STM14_RS04845 | | Cof-type HAD-IIB family hydrolase | WP_000217910.1 | 1.541957909 | 1.6857E-21 | 6.603E-20 |
| STM14_RS13125 | | sulfite exporter TauE/SafE family protein | WP_000368599.1 | 1.530320032 | 7.9153E-12 | 1.1611E-10 |
| STM14_RS20365 | pstS | phosphate ABC transporter substrate-binding protein PstS | WP_000867130.1 | 1.503260631 | 6.6852E-22 | 2.8959E-20 |
| STM14_RS02965 | ykgO | type B 50S ribosomal protein L36 | WP_001197749.1 | 1.500377503 | 6.6088E-15 | 1.4314E-13 |
| STM14_RS04525 | | AcrZ family multidrug efflux pump-associated protein | WP_000891513.1 | 1.499011191 | 1.7592E-15 | 4.0483E-14 |
| STM14_RS00570 | thrL | thr operon leader peptide | WP_001575544.1 | 1.495832844 | 1.2815E-33 | 1.3482E-31 |
| STM14_RS23395 | | YfcC family protein | WP_000514425.1 | 1.480057964 | 0.00891296 | 0.02164745 |
| STM14_RS16975 | | putative quinol monooxygenase | WP_000958587.1 | 1.479760959 | 3.4345E-07 | 2.4555E-06 |
| STM14_RS05685 | | phage tail tape measure protein | WP_010989010.1 | 1.476903499 | 4.1644E-20 | 1.4328E-18 |
| STM14_RS21045 | | acyltransferase | WP_000196890.1 | 1.476393786 | 1.5917E-16 | 4.0141E-15 |
| STM14_RS05095 | | lipoprotein | WP_001270724.1 | 1.473240836 | 1.0992E-23 | 5.3252E-22 |
| STM14_RS18465 | | YhfL family protein | WP_000472323.1 | 1.470749249 | 1.6069E-11 | 2.2757E-10 |
| STM14_RS05985 | hpaB | 4-hydroxyphenylacetate 3-monooxygenase%2C oxygenase component | WP_000801496.1 | 1.449503265 | 8.8223E-05 | 0.00037467 |
| STM14_RS23400 | argF | ornithine carbamoyltransferase | WP_000237031.1 | 1.436461459 | 0.01666812 | 0.03659631 |
| STM14_RS15420 | hilA | transcriptional regulator HilA | WP_001120085.1 | 1.434977902 | 4.7461E-37 | 6.4723E-35 |
| STM14_RS23040 | priB | primosomal replication protein N | WP_001519453.1 | 1.42969723 | 3.0745E-05 | 0.00014759 |
| STM14_RS22550 | | AraC family transcriptional regulator | WP_001217074.1 | 1.411444927 | 4.3772E-25 | 2.4055E-23 |
| STM14_RS14575 | raiA | ribosome-associated translation inhibitor RaiA | WP_000178449.1 | 1.410609531 | 1.0696E-53 | 4.9227E-51 |
| STM14_RS06950 | | phage tail tape measure protein | WP_000372072.1 | 1.405411235 | 1.6141E-13 | 2.9568E-12 |
| STM14_RS25330 | | hypothetical protein | WP_001626563.1 | 1.403724801 | 3.2811E-10 | 3.7635E-09 |
| STM14_RS10720 | fliT | flagella biosynthesis regulatory protein FliT | WP_000204899.1 | 1.396482688 | 1.2303E-22 | 5.6624E-21 |
| STM14_RS17760 | nanR | transcriptional regulator NanR | WP_000382926.1 | 1.386404528 | 5.1583E-20 | 1.7266E-18 |
| STM14_RS01190 | ftsI | peptidoglycan glycosyltransferase FtsI | WP_000642235.1 | 1.381932237 | 4.7978E-44 | 8.0297E-42 |
| STM14_RS10710 | fliD | flagellar filament capping protein FliD | WP_000146802.1 | 1.377625665 | 5.5733E-46 | 1.14E-43 |
| STM14_RS14485 | grcA | autonomous glycyl radical cofactor GrcA | WP_000627811.1 | 1.368255119 | 1.4563E-45 | 2.8221E-43 |
| STM14_RS11535 | sopA | SPI-1 type III secretion system effector HECT-type E3 ubiquitin transfera | WP_000703998.1 | 1.367289082 | 1.1548E-21 | 4.6216E-20 |
| STM14_RS12440 | pagL | lipid A 3-O-deacylase PagL | WP_000281877.1 | 1.357078014 | 1.2889E-10 | 1.6142E-09 |
| STM14_RS03630 | | phosphoadenosine phosphosulfate reductase | WP_000118144.1 | 1.354341719 | 1.0034E-06 | 6.4588E-06 |
| STM14_RS26425 | rseD | rpoE leader peptide RseD | WP_014344135.1 | 1.354180428 | 2.3648E-33 | 2.4187E-31 |
| STM14_RS22835 | | arginine ABC transporter substrate-binding protein | WP_000727896.1 | 1.352581406 | 5.1168E-16 | 1.2234E-14 |

| | | | | | | |
|---|---|---|---|---|---|---|
| STM14_RS10165 | | membrane protein | WP_001134856.1 | 1.344518057 | 6.1094E-30 | 4.6864E-28 |
| STM14_RS02890 | fadM | long-chain acyl-CoA thioesterase FadM | WP_001194543.1 | 1.328587946 | 9.0941E-08 | 7.2951E-07 |
| STM14_RS09250 | | YgdI/YgdR family lipoprotein | WP_001168019.1 | 1.315832298 | 4.3793E-09 | 4.3345E-08 |
| STM14_RS09650 | hns | DNA-binding transcriptional regulator H-NS | WP_001287383.1 | 1.284718203 | 2.2251E-44 | 4.0964E-42 |
| STM14_RS03475 | | PTS system mannose/fructose/sorbose family transporter subunit IID | WP_000005803.1 | 1.27537212 | 1.1328E-16 | 2.8764E-15 |
| STM14_RS05565 | | DUF1367 family protein | WP_000929805.1 | 1.274781767 | 0.00015957 | 0.00063245 |
| STM14_RS07035 | msgA | virulence protein MsgA | WP_000497451.1 | 1.272359471 | 5.323E-11 | 7.0249E-10 |
| STM14_RS08800 | | YgdI/YgdR family lipoprotein | WP_000732362.1 | 1.268985425 | 3.7221E-05 | 0.00017615 |
| STM14_RS23360 | mgtA | magnesium-translocating P-type ATPase | WP_001738655.1 | 1.260351051 | 1.6983E-24 | 8.9333E-23 |
| STM14_RS06625 | sifA | SPI-2 type III secretion system effector SifA | WP_001520468.1 | 1.259536733 | 1.6537E-18 | 4.8713E-17 |
| STM14_RS04615 | uvrB | excinuclease ABC subunit UvrB | WP_000042502.1 | 1.256248245 | 1.1403E-32 | 1.0496E-30 |
| STM14_RS08710 | fdnI | formate dehydrogenase-N subunit gamma | WP_000045657.1 | 1.252134241 | 3.0784E-20 | 1.0795E-18 |
| STM14_RS05070 | artJ | ABC transporter substrate-binding protein ArtJ | WP_000737538.1 | 1.249083656 | 0.0315812 | 0.06278724 |
| STM14_RS26490 | | integrase arm-type DNA-binding domain-containing protein | | 1.24707583 | 9.142E-07 | 5.9367E-06 |
| STM14_RS25905 | | hypothetical protein | WP_010989001.1 | 1.242498717 | 5.1018E-16 | 1.2234E-14 |
| STM14_RS12895 | ackA | acetate kinase | WP_000095689.1 | 1.236803785 | 8.2425E-26 | 5.0582E-24 |
| STM14_RS15475 | spaS | SPI-1 type III secretion system export apparatus protein SpaS | WP_000097719.1 | 1.225166341 | 2.6329E-16 | 6.5063E-15 |
| STM14_RS05930 | pipB | SPI-2 type III secretion system effector PipB | WP_001123045.1 | 1.217954751 | 1.6748E-10 | 2.0219E-09 |
| STM14_RS08780 | | PhzF family isomerase | WP_000804282.1 | 1.210789208 | 1.1E-06 | 7.0562E-06 |
| STM14_RS09200 | | mechanosensitive ion channel family protein | WP_000493665.1 | 1.209117362 | 4.7562E-07 | 3.2733E-06 |
| STM14_RS15425 | iagB | SPI-1 type III secretion system invasion protein IagB | WP_000558928.1 | 1.205354043 | 3.8577E-19 | 1.1837E-17 |
| STM14_RS01125 | | glucose uptake inhibitor SgrT | WP_001248755.1 | 1.197473918 | 8.9263E-33 | 8.4273E-31 |
| STM14_RS07220 | | hypothetical protein | WP_000762776.1 | 1.197048146 | 0.00016638 | 0.00065519 |
| STM14_RS10175 | | DUF1398 domain-containing protein | WP_000422882.1 | 1.192841383 | 1.4025E-06 | 8.7527E-06 |
| STM14_RS22635 | | hypothetical protein | WP_010989096.1 | 1.186407967 | 7.9766E-07 | 5.2634E-06 |
| STM14_RS22590 | | hypothetical protein | WP_010989095.1 | 1.18490241 | 0.00077584 | 0.0025782 |
| STM14_RS04910 | | electron transfer flavoprotein subunit beta/FixA family protein | WP_000442433.1 | 1.18278878 | 3.9078E-05 | 0.00018376 |
| STM14_RS18620 | nfuA | Fe-S biogenesis protein NfuA | WP_000619387.1 | 1.180126266 | 2.3332E-08 | 2.0701E-07 |
| STM14_RS09820 | | hydrogenase 1 small subunit | WP_001058312.1 | 1.176824877 | 2.1418E-08 | 1.9049E-07 |
| STM14_RS17290 | tdcC | threonine/serine transporter TdcC | WP_000108129.1 | 1.174338856 | 2.2929E-10 | 2.7059E-09 |
| STM14_RS06485 | acpP | acyl carrier protein | WP_000103754.1 | 1.167617745 | 4.4149E-41 | 6.7732E-39 |
| STM14_RS24010 | yjjX | inosine/xanthosine triphosphatase | WP_000554311.1 | 1.16653371 | 8.5667E-07 | 5.6126E-06 |
| STM14_RS15540 | invH | SPI-1 type III secretion system invasion lipoprotein InvH | WP_000715092.1 | 1.165443491 | 3.632E-24 | 1.8319E-22 |
| STM14_RS03480 | | PTS mannose/fructose/sorbose/N-acetylgalactosamine transporter sub | WP_000927523.1 | 1.153571764 | 5.6369E-07 | 3.8223E-06 |
| STM14_RS20570 | ilvL | ilv operon leader peptide | WP_001311244.1 | 1.152543261 | 4.0075E-09 | 3.9772E-08 |
| STM14_RS05150 | | VirK/YbjX family protein | WP_001519667.1 | 1.149530658 | 2.8273E-34 | 3.2532E-32 |
| STM14_RS26045 | | tyrosine-type recombinase/integrase | | 1.149245393 | 0.00026276 | 0.00098521 |
| STM14_RS02510 | | DUF2754 family protein | WP_001239243.1 | 1.148785326 | 1.3316E-10 | 1.662E-09 |
| STM14_RS21205 | | type II toxin-antitoxin system HigB family toxin | WP_000558166.1 | 1.146761081 | 3.6537E-09 | 3.636E-08 |
| STM14_RS22195 | psiE | phosphate-starvation-inducible protein PsiE | WP_000982749.1 | 1.142150247 | 0.00025166 | 0.00094746 |
| STM14_RS11385 | pocR | regulatory protein PocR | WP_000622326.1 | 1.140548889 | 1.3851E-09 | 1.4488E-08 |
| STM14_RS03490 | | PTS sugar transporter subunit IIA | WP_001216659.1 | 1.137707655 | 8.0649E-10 | 8.5807E-09 |
| STM14_RS17295 | tdcB | bifunctional threonine ammonia-lyase/L-serine ammonia-lyase TdcB | WP_000548370.1 | 1.134091093 | 6.3219E-14 | 1.2123E-12 |
| STM14_RS22760 | ecnB | lipoprotein toxin entericidin B | WP_000239598.1 | 1.133820215 | 9.186E-34 | 9.9479E-32 |
| STM14_RS03355 | fimZ | fimbria biosynthesis transcriptional regulator FimZ | WP_000801273.1 | 1.12088791 | 5.0406E-17 | 1.3163E-15 |
| STM14_RS07210 | yeaR | DUF1971 domain-containing protein YeaR | WP_001248993.1 | 1.120231657 | 4.6984E-05 | 0.00021733 |
| STM14_RS05085 | artJ | arginine ABC transporter substrate-binding protein | WP_000756586.1 | 1.113056252 | 3.9543E-18 | 1.12E-16 |
| STM14_RS07855 | sseC | SPI-2 type III secretion system translocon protein SseC | WP_001079760.1 | 1.112788412 | 1.3332E-26 | 8.7659E-25 |
| STM14_RS19110 | pdeH | cyclic-guanylate-specific phosphodiesterase | WP_000595626.1 | 1.112545248 | 6.2145E-19 | 1.8756E-17 |
| STM14_RS07105 | | Hsp20 family protein | WP_001218118.1 | 1.111933058 | 2.7081E-22 | 1.216E-20 |
| STM14_RS10525 | motA | flagellar motor stator protein MotA | WP_000906312.1 | 1.10842872 | 3.2273E-29 | 2.3766E-27 |
| STM14_RS19880 | mgtA | magnesium-translocating P-type ATPase | WP_000131288.1 | 1.105529906 | 1.7139E-17 | 4.6063E-16 |
| STM14_RS02270 | | alpha/beta hydrolase | WP_000105593.1 | 1.104268711 | 0.00339144 | 0.00940308 |
| STM14_RS07065 | pagC | virulence membrane protein PagC | WP_000789471.1 | 1.100257817 | 7.6475E-36 | 9.7097E-34 |
| STM14_RS19885 | | MgtC family protein | WP_000392694.1 | 1.097897766 | 1.0353E-08 | 9.7493E-08 |
| STM14_RS05220 | | membrane protein | WP_001539595.1 | 1.091800741 | 2.5186E-08 | 2.2239E-07 |
| STM14_RS15565 | | GNAT family N-acetyltransferase | WP_000971655.1 | 1.08957303 | 1.9633E-10 | 2.3547E-09 |
| STM14_RS07910 | ssaK | SPI-2 type III secretion system apparatus protein SsaK | WP_000011201.1 | 1.087566417 | 3.7843E-14 | 7.5318E-13 |
| STM14_RS26290 | speFL | leader peptide SpeFL | WP_015632840.1 | 1.083726357 | 5.088E-15 | 1.1354E-13 |
| STM14_RS05550 | | hypothetical protein | WP_014343823.1 | 1.082706661 | 6.3917E-24 | 3.1803E-22 |
| STM14_RS07430 | chbA | PTS N%2CN'-diacetylchitobiose transporter subunit IIA | WP_000459883.1 | 1.077593737 | 2.1309E-09 | 2.1674E-08 |
| STM14_RS26550 | | hypothetical protein | WP_012218875.1 | 1.077507955 | 2.8371E-26 | 1.8327E-24 |
| STM14_RS21555 | | aryl-sulfate sulfotransferase | WP_000899636.1 | 1.076897597 | 1.3508E-07 | 1.0471E-06 |
| STM14_RS16845 | | hypothetical protein | WP_000936444.1 | 1.076378416 | 2.4372E-09 | 2.4585E-08 |
| STM14_RS02945 | | YbaY family lipoprotein | WP_000779803.1 | 1.072248701 | 1.5356E-21 | 6.0797E-20 |
| STM14_RS04115 | speF | ornithine decarboxylase SpeF | WP_001292400.1 | 1.063114957 | 2.0869E-10 | 2.4867E-09 |
| STM14_RS14010 | cadC | lysine decarboxylation/transport transcriptional activator CadC | WP_001187150.1 | 1.050617327 | 1.9499E-13 | 3.5367E-12 |
| STM14_RS26230 | | hypothetical protein | WP_001595136.1 | 1.043978638 | 2.8096E-05 | 0.00013604 |
| STM14_RS08420 | | general stress protein | WP_000807642.1 | 1.042740266 | 1.6573E-20 | 5.8674E-19 |
| STM14_RS22675 | | MerR family transcriptional regulator | WP_001165471.1 | 1.04239017 | 1.49E-07 | 1.1403E-06 |
| STM14_RS09245 | | hypothetical protein | WP_000275493.1 | 1.041704283 | 2.8134E-30 | 2.3019E-28 |
| STM14_RS19510 | | hypothetical protein | WP_001062558.1 | 1.034201847 | 7.6152E-09 | 7.3209E-08 |
| STM14_RS07595 | aroH | 3-deoxy-7-phosphoheptulonate synthase AroH | WP_001082196.1 | 1.034025291 | 4.3561E-10 | 4.89E-09 |
| STM14_RS12370 | | acyltransferase family protein | WP_000639473.1 | 1.032201914 | 3.0884E-23 | 1.4579E-21 |
| STM14_RS21605 | | DUF1287 domain-containing protein | WP_000666689.1 | 1.031430211 | 7.1011E-05 | 0.00030942 |
| STM14_RS07905 | | type III secretion system domain-containing protein | WP_001526945.1 | 1.031305743 | 1.1315E-08 | 1.0521E-07 |
| STM14_RS15470 | sicA | SycD/LcrH family type III secretion system chaperone SicA | WP_000386309.1 | 1.030758871 | 2.9796E-26 | 1.8915E-24 |
| STM14_RS08165 | blr | division septum protein Blr | WP_001708800.1 | 1.030632344 | 5.3347E-05 | 0.0002428 |
| STM14_RS09825 | hyaB | Ni/Fe-hydrogenase large subunit | WP_000070978.1 | 1.027206139 | 5.1501E-07 | 3.5051E-06 |
| STM14_RS02140 | pagN | adhesin/invasin protein PagN | WP_000787603.1 | 1.024578224 | 5.9855E-16 | 1.4127E-14 |
| STM14_RS18650 | | type II toxin-antitoxin system RelB/DinJ family antitoxin | WP_000729713.1 | 1.022856108 | 3.9985E-17 | 1.0592E-15 |
| STM14_RS17210 | mzrA | EnvZ/OmpR regulon moderator MzrA | WP_000917516.1 | 1.01708437 | 2.4057E-05 | 0.00011858 |

| | | | | | | |
|---|---|---|---|---|---|---|
| STM14_RS15545 | | membrane protein | WP_000381610.1 | 1.015322981 | 4.0024E-14 | 7.8982E-13 |
| STM14_RS14875 | | FliC/FljB family flagellin | WP_000079794.1 | 1.015143682 | 1.438E-28 | 1.0182E-26 |
| STM14_RS15415 | hilD | transcriptional regulator HilD | WP_000432699.1 | 1.012350767 | 5.2938E-14 | 1.0313E-12 |
| STM14_RS21750 | birA | bifunctional biotin--[acetyl-CoA-carboxylase] ligase/biotin operon repre | WP_000655752.1 | 1.011452729 | 3.2456E-13 | 5.6907E-12 |
| STM14_RS10705 | | FliC/FljB family flagellin | WP_000079805.1 | 1.00965052 | 4.157E-28 | 2.8344E-26 |
| STM14_RS04625 | slrP | SPI-1 type III secretion system effector E3 ubiquitin transferase SlrP | WP_000481997.1 | 1.009503228 | 3.1421E-20 | 1.0914E-18 |
| STM14_RS11220 | | SEL1-like repeat protein | WP_010989035.1 | 1.008231091 | 0.00035598 | 0.00128755 |
| STM14_RS11895 | steD | type III secretion system effector SteD | WP_001542166.1 | 1.006787307 | 2.4364E-14 | 4.9562E-13 |
| STM14_RS00240 | | hypothetical protein | WP_001135407.1 | 1.005643041 | 1.7522E-07 | 1.3244E-06 |
| STM14_RS17870 | yhcN-B | DUF1471 family stress response protein YhcN-B | WP_000853277.1 | 1.003885078 | 9.9431E-07 | 6.4229E-06 |
| STM14_RS23480 | | LacI family DNA-binding transcriptional regulator | WP_001244065.1 | 1.003716165 | 2.2297E-06 | 1.3503E-05 |
| STM14_RS00510 | | conjugal transfer protein TraH | WP_001161719.1 | 1.002808734 | 0.0045055 | 0.01192614 |
| STM14_RS17155 | | methyl-accepting chemotaxis protein | WP_000478472.1 | 1.001256457 | 1.817E-29 | 1.3654E-27 |
| STM14_RS13565 | pta | phosphate acetyltransferase | WP_000582931.1 | 0.99566175 | 0.00074576 | 0.00249172 |
| STM14_RS07960 | ssaT | SPI-2 type III secretion system apparatus protein SsaT | WP_000068764.1 | 0.995138027 | 4.1439E-05 | 0.00019412 |
| STM14_RS15525 | invE | type III secretion system gatekeeper InvE | WP_000612171.1 | 0.99475256 | 7.2698E-14 | 1.3727E-12 |
| STM14_RS19950 | | type II toxin-antitoxin system RelE/ParE family toxin | WP_001258724.1 | 0.994210538 | 1.9947E-08 | 1.7827E-07 |
| STM14_RS14920 | | VirK family antimicrobial peptide resistance protein | WP_000178733.1 | 0.989720053 | 6.7947E-23 | 3.1669E-21 |
| STM14_RS14915 | pipB2 | SPI-2 type III secretion system effector PipB2 | WP_001738474.1 | 0.987619649 | 1.3907E-19 | 4.4917E-18 |
| STM14_RS08790 | steA | type III secretion system effector SteA | WP_001147134.1 | 0.986144409 | 6.0796E-30 | 4.6864E-28 |
| STM14_RS08885 | sifB | SPI-2 type III secretion system effector SifB | WP_001122926.1 | 0.984970925 | 2.4702E-08 | 2.1863E-07 |
| STM14_RS15375 | hilC | invasion transcriptional regulator HilC | WP_000243999.1 | 0.984194135 | 1.1225E-21 | 4.5891E-20 |
| STM14_RS15530 | invG | type III secretion system outer membrane ring protein InvG | WP_000848113.1 | 0.983556407 | 1.351E-17 | 3.6847E-16 |
| STM14_RS25790 | | hypothetical protein | WP_001687482.1 | 0.982539805 | 3.1793E-06 | 1.882E-05 |
| STM14_RS00265 | | Rpn family recombination-promoting nuclease/putative transposase | WP_000728917.1 | 0.981346293 | 5.795E-29 | 4.1838E-27 |
| STM14_RS22630 | | YjiK family protein | WP_001268200.1 | 0.978511425 | 4.1526E-11 | 5.6007E-10 |
| STM14_RS10845 | | YodC family protein | WP_000232161.1 | 0.976173147 | 2.6158E-24 | 1.3377E-22 |
| STM14_RS13610 | | hypothetical protein | WP_000700635.1 | 0.976000115 | 2.2155E-12 | 3.3849E-11 |
| STM14_RS07510 | | DNA/RNA non-specific endonuclease | WP_001044098.1 | 0.973132134 | 5.637E-14 | 1.0867E-12 |
| STM14_RS07865 | sseE | LcrR family type III secretion system chaperone SseE | WP_000249606.1 | 0.969211338 | 4.6509E-09 | 4.5788E-08 |
| STM14_RS20630 | rep | DNA helicase Rep | WP_001238842.1 | 0.967061507 | 6.3402E-15 | 1.3813E-13 |
| STM14_RS10715 | fliS | flagellar export chaperone FliS | WP_000287764.1 | 0.963326882 | 9.0866E-15 | 1.9452E-13 |
| STM14_RS07885 | ssaG | type III secretion system needle filament protein SsaG | WP_000350226.1 | 0.959194203 | 1.4782E-08 | 1.3539E-07 |
| STM14_RS14415 | | PD-(D/E)XK nuclease-like domain-containing protein | WP_000017127.1 | 0.958709024 | 0.00033169 | 0.00121157 |
| STM14_RS05500 | | PD-(D/E)XK nuclease-like domain-containing protein | WP_000017133.1 | 0.957999067 | 0.00210199 | 0.00619657 |
| STM14_RS03470 | | SIS domain-containing protein | WP_010988979.1 | 0.951732816 | 1.0119E-11 | 1.4611E-10 |
| STM14_RS12860 | lrhA | transcriptional regulator LrhA | WP_000606288.1 | 0.945485948 | 8.6226E-14 | 1.6035E-12 |
| STM14_RS06850 | | DUF2441 domain-containing protein | WP_000029438.1 | 0.944795593 | 5.3528E-18 | 1.4708E-16 |
| STM14_RS11175 | | tyrosine-type recombinase/integrase | WP_000532847.1 | 0.944700173 | 1.3459E-08 | 1.2451E-07 |
| STM14_RS07925 | ssaV | SPI-2 type III secretion system apparatus protein SsaV | WP_001258231.1 | 0.944377451 | 5.9428E-11 | 7.7869E-10 |
| STM14_RS26835 | | hypothetical protein | WP_001519391.1 | 0.942631587 | 0.0002811 | 0.00104127 |
| STM14_RS07900 | ssaJ | SPI-2 type III secretion system apparatus lipoprotein SsaJ | WP_000862874.1 | 0.940849653 | 2.1503E-13 | 3.8622E-12 |
| STM14_RS12105 | | LysR substrate-binding domain-containing protein | WP_000024658.1 | 0.938054067 | 0.00019743 | 0.00076521 |
| STM14_RS11250 | | hypothetical protein | | 0.937239952 | 1.0228E-06 | 6.5724E-06 |
| STM14_RS02940 | tesB | acyl-CoA thioesterase II | WP_000084180.1 | 0.936791036 | 2.876E-05 | 0.00013861 |
| STM14_RS12385 | | hypothetical protein | WP_000554739.1 | 0.935594241 | 0.00220363 | 0.00643288 |
| STM14_RS00730 | | aryl-sulfate sulfotransferase | WP_001036514.1 | 0.93302878 | 4.8567E-08 | 4.0827E-07 |
| STM14_RS20460 | asnC | transcriptional regulator AsnC | WP_000433018.1 | 0.932335496 | 0.00043073 | 0.00153381 |
| STM14_RS06125 | | MurR/RpiR family transcriptional regulator | WP_000433674.1 | 0.931745205 | 4.1392E-11 | 5.6007E-10 |
| STM14_RS07040 | | RICIN domain-containing protein | WP_001520581.1 | 0.930455874 | 1.9886E-13 | 3.5892E-12 |
| STM14_RS10170 | sopE2 | SPI-1 type III secretion system guanine nucleotide exchange factor SopE | WP_000182072.1 | 0.929413598 | 2.1446E-17 | 5.7219E-16 |
| STM14_RS22765 | | LuxR C-terminal-related transcriptional regulator | WP_000912957.1 | 0.929157496 | 3.834E-07 | 2.7095E-06 |
| STM14_RS22260 | | hypothetical protein | WP_010989093.1 | 0.928563967 | 2.1211E-23 | 1.0143E-21 |
| STM14_RS15440 | | hypothetical protein | WP_000692254.1 | 0.926288296 | 5.191E-17 | 1.3366E-15 |
| STM14_RS09320 | pspE | thiosulfate sulfurtransferase PspE | WP_000913428.1 | 0.923955458 | 2.465E-19 | 7.7573E-18 |
| STM14_RS18120 | | alternative ribosome-rescue factor A | WP_001682446.1 | 0.921004557 | 5.1218E-10 | 5.6974E-09 |
| STM14_RS09935 | | addiction module toxin%2C GnsA/GnsB family | WP_001083582.1 | 0.919173053 | 2.6093E-24 | 1.3377E-22 |
| STM14_RS14750 | | helix-turn-helix transcriptional regulator | WP_010989065.1 | 0.917971734 | 4.4814E-05 | 0.00020834 |
| STM14_RS07020 | sopF | type III secretion system effector SopF | WP_000917257.1 | 0.917904039 | 5.9975E-10 | 6.5918E-09 |
| STM14_RS09480 | | YciN family protein | WP_000548616.1 | 0.916829364 | 2.5898E-13 | 4.6066E-12 |
| STM14_RS19255 | lpfD | long polar fimbrial protein LpfD | | 0.915175841 | 0.0033657 | 0.00933874 |
| STM14_RS16835 | | TIGR00645 family protein | WP_000439335.1 | 0.913700319 | 0.00015459 | 0.00061467 |
| STM14_RS06415 | flgK | flagellar hook-associated protein FlgK | WP_000096425.1 | 0.913635913 | 6.9642E-20 | 2.3101E-18 |
| STM14_RS10780 | fliK | flagellar hook length control protein FliK | WP_000631669.1 | 0.912461753 | 1.5524E-06 | 9.5907E-06 |
| STM14_RS09295 | tpx | thiol peroxidase | WP_000084401.1 | 0.911013091 | 1.5817E-20 | 5.7097E-19 |
| STM14_RS10700 | fliB | flagellin lysine-N-methylase | WP_000659236.1 | 0.910215124 | 8.3066E-12 | 1.2089E-10 |
| STM14_RS18565 | pckA | phosphoenolpyruvate carboxykinase (ATP) | WP_001265689.1 | 0.909182358 | 1.1387E-25 | 6.8731E-24 |
| STM14_RS09155 | ttcA | tRNA 2-thiocytidine(32) synthetase TtcA | WP_001169566.1 | 0.909173543 | 1.7993E-08 | 1.6237E-07 |
| STM14_RS11245 | | DUF1493 family protein | WP_000228158.1 | 0.908598948 | 1.7768E-06 | 1.0922E-05 |
| STM14_RS09815 | | DUF4427 domain-containing protein | WP_000794388.1 | 0.90796846 | 4.6866E-11 | 6.2749E-10 |
| STM14_RS07080 | | c-type lysozyme inhibitor | WP_000977725.1 | 0.907221432 | 2.6905E-25 | 1.5479E-23 |
| STM14_RS16665 | ripC | itaconate degradation C-C-lyase RipC | WP_001112897.1 | 0.905092443 | 0.00271926 | 0.00775917 |
| STM14_RS23175 | iolT1 | myo-inositol import MFS transporter IolT1 | WP_000107709.1 | 0.903233584 | 0.00127994 | 0.00399723 |
| STM14_RS19955 | | DNA-binding transcriptional regulator | WP_001274847.1 | 0.902826689 | 0.00437606 | 0.01164208 |
| STM14_RS08110 | gstA | glutathione transferase GstA | WP_000765727.1 | 0.901897976 | 2.0858E-07 | 1.5515E-06 |
| STM14_RS09880 | ycgR | flagellar brake protein YcgR | WP_000017421.1 | 0.901703479 | 2.2805E-10 | 2.6999E-09 |
| STM14_RS07500 | | lipid A deacylase LpxR family protein | WP_001046438.1 | 0.900121825 | 1.7928E-19 | 5.7399E-18 |
| STM14_RS06245 | mdoC | glucans biosynthesis protein MdoC | WP_000100055.1 | 0.896817963 | 0.00141001 | 0.00434812 |
| STM14_RS10580 | | YecR-like lipofamily protein | WP_000803229.1 | 0.895509451 | 0.00055202 | 0.00190712 |
| STM14_RS09190 | uspE | universal stress protein UspE | WP_001262138.1 | 0.895161573 | 3.1186E-21 | 1.1838E-19 |
| STM14_RS17040 | | thiol:disulfide interchange protein DsbA/DsbL | WP_000096230.1 | 0.893873044 | 0.00044297 | 0.00157284 |
| STM14_RS21905 | hupA | DNA-binding protein HU-alpha | WP_001044509.1 | 0.892642811 | 5.8082E-26 | 3.6247E-24 |

| | | | | | | |
|---|---|---|---|---|---|---|
| STM14_RS05925 | pipA | type III secretion system effector protease PipA | WP_000938191.1 | 0.891391502 | 6.888E-05 | 0.00030121 |
| STM14_RS18395 | crp | cAMP-activated global transcriptional regulator CRP | WP_000242758.1 | 0.890698263 | 4.2026E-20 | 1.4328E-18 |
| STM14_RS12640 | glpC | anaerobic glycerol-3-phosphate dehydrogenase subunit GlpC | WP_001285726.1 | 0.88986542 | 3.694E-06 | 2.1555E-05 |
| STM14_RS10785 | fliL | flagellar basal body-associated protein FliL | WP_000132169.1 | 0.888872911 | 0.00123795 | 0.00387597 |
| STM14_RS12525 | eco | serine protease inhibitor ecotin | WP_000781589.1 | 0.888436001 | 4.7268E-05 | 0.00021837 |
| STM14_RS01000 | | sulfatase-like hydrolase/transferase | WP_000066317.1 | 0.887942797 | 0.00057742 | 0.00198328 |
| STM14_RS15370 | sprB | transcriptional regulator SprB | WP_001246659.1 | 0.886586596 | 2.9377E-14 | 5.9106E-13 |
| STM14_RS08820 | | NADP-dependent oxidoreductase | WP_001517827.1 | 0.885962703 | 1.946E-11 | 2.7418E-10 |
| STM14_RS00035 | repA | incFII family plasmid replication initiator RepA | WP_001738692.1 | 0.885771636 | 7.2838E-08 | 5.9465E-07 |
| STM14_RS06360 | flgM | flagellar biosynthesis anti-sigma factor FlgM | WP_000020893.1 | 0.885024877 | 9.807E-22 | 4.1461E-20 |
| STM14_RS05600 | | hypothetical protein | WP_000798705.1 | 0.884974336 | 1.5258E-07 | 1.1607E-06 |
| STM14_RS12635 | glpB | glycerol-3-phosphate dehydrogenase subunit GlpB | WP_000667146.1 | 0.883703663 | 0.0002731 | 0.00101573 |
| STM14_RS03775 | | YbeF family transcriptional regulator | WP_001282817.1 | 0.883553868 | 5.5966E-08 | 4.6411E-07 |
| STM14_RS14355 | | hypothetical protein | WP_000877757.1 | 0.881111998 | 3.8219E-21 | 1.4215E-19 |
| STM14_RS22820 | asd | archaetidylserine decarboxylase | WP_000934949.1 | 0.880317254 | 5.5807E-08 | 4.6384E-07 |
| STM14_RS21255 | | YiiG family protein | WP_000710966.1 | 0.880116615 | 5.8849E-06 | 3.2934E-05 |
| STM14_RS10585 | ftnA | non-heme ferritin | WP_000920612.1 | 0.879751255 | 7.1909E-14 | 1.3648E-12 |
| STM14_RS15410 | prgH | type III secretion system inner membrane ring protein PrgH | WP_000450192.1 | 0.878328881 | 2.04E-10 | 2.4387E-09 |
| STM14_RS17285 | tdcD | propionate kinase | WP_001001853.1 | 0.878217166 | 0.00034181 | 0.00124732 |
| STM14_RS02530 | iraP | anti-adapter protein IraP | WP_001518423.1 | 0.875899679 | 1.3759E-22 | 6.2546E-21 |
| STM14_RS15480 | spaR | SPI-1 type III secretion system export apparatus protein SpaR | WP_000498965.1 | 0.871550753 | 9.026E-05 | 0.00038069 |
| STM14_RS17860 | argR | transcriptional regulator ArgR | WP_001257852.1 | 0.871418362 | 8.3288E-11 | 1.0685E-09 |
| STM14_RS21480 | | DUF805 domain-containing protein | WP_000155237.1 | 0.871260331 | 0.00067595 | 0.00227565 |
| STM14_RS23375 | pyrI | aspartate carbamoyltransferase regulatory subunit | WP_000148570.1 | 0.870479007 | 0.00261505 | 0.00749893 |
| STM14_RS25970 | | hypothetical protein | WP_000480529.1 | 0.86974552 | 3.6565E-06 | 2.1392E-05 |
| STM14_RS10075 | ftsI | peptidoglycan glycosyltransferase FtsI | WP_000730322.1 | 0.863322731 | 2.5776E-05 | 0.00012587 |
| STM14_RS26860 | | rho operon leader peptide | WP_134820045.1 | 0.858865424 | 1.7199E-16 | 4.308E-15 |
| STM14_RS04195 | | tyrosine-type DNA invertase | WP_000824704.1 | 0.85842518 | 0.00212337 | 0.00624959 |
| STM14_RS25045 | | small membrane protein YohP | WP_000698460.1 | 0.856677191 | 2.3319E-05 | 0.00011525 |
| STM14_RS06705 | | site-specific integrase | WP_000741325.1 | 0.856342444 | 0.00012632 | 0.00051565 |
| STM14_RS22700 | dcuA | anaerobic C4-dicarboxylate transporter DcuA | WP_000637595.1 | 0.854257983 | 3.2549E-18 | 9.2904E-17 |
| STM14_RS05820 | rmf | ribosome modulation factor | WP_001537784.1 | 0.854042241 | 8.5222E-20 | 2.8017E-18 |
| STM14_RS26630 | | PD-(D/E)XK nuclease-like domain-containing protein | | 0.853570233 | 2.5967E-06 | 1.5597E-05 |
| STM14_RS21185 | | virulence factor BrkB family protein | WP_000921427.1 | 0.853526595 | 6.8215E-06 | 3.7713E-05 |
| STM14_RS26950 | | hypothetical protein | WP_000469259.1 | 0.852150128 | 1.3052E-09 | 1.3692E-08 |
| STM14_RS08880 | | hypothetical protein | WP_000747896.1 | 0.850394066 | 5.4382E-13 | 9.2701E-12 |
| STM14_RS05745 | sseI | SPI-2 type III secretion system effector SseI | WP_001526469.1 | 0.850149468 | 0.00042645 | 0.00152149 |
| STM14_RS15385 | orgB | type III secretion system linker protein OrgB | WP_000916654.1 | 0.850146256 | 2.7116E-13 | 4.777E-12 |
| STM14_RS22560 | adiA | arginine decarboxylase | WP_001542850.1 | 0.849110441 | 4.3451E-07 | 3.0243E-06 |
| STM14_RS14470 | srmB | ATP-dependent RNA helicase SrmB | WP_000219174.1 | 0.849048519 | 2.6394E-08 | 2.3194E-07 |
| STM14_RS05940 | sigE | type III secretion system chaperone SigE | WP_000444724.1 | 0.846317405 | 1.6719E-12 | 2.5974E-11 |
| STM14_RS08930 | rimL | 50S ribosomal protein L7/L12-serine acetyltransferase | WP_000229304.1 | 0.844222561 | 1.762E-07 | 1.3267E-06 |
| STM14_RS12585 | | GntR family transcriptional regulator | WP_000944714.1 | 0.841700882 | 0.00013619 | 0.00055227 |
| STM14_RS07690 | | YdiH family protein | WP_001537749.1 | 0.841047447 | 7.3622E-15 | 1.5852E-13 |
| STM14_RS07830 | ssaD | SctD family type III secretion system inner membrane ring subunit SsaD | WP_000323786.1 | 0.840322907 | 8.7566E-09 | 8.3745E-08 |
| STM14_RS07845 | sseB | SPI-2 type III secretion system translocon protein SseB | WP_000094422.1 | 0.838737103 | 1.9781E-14 | 4.115E-13 |
| STM14_RS01640 | | DUF3461 family protein | WP_000272193.1 | 0.838681791 | 5.0489E-19 | 1.5364E-17 |
| STM14_RS17055 | ubiK | ubiquinone biosynthesis accessory factor UbiK | WP_000566825.1 | 0.838216659 | 6.3879E-10 | 6.9587E-09 |
| STM14_RS11890 | | CesT family type III secretion system chaperone | WP_001669236.1 | 0.837197732 | 0.00024751 | 0.00093668 |
| STM14_RS02955 | | EAL domain-containing protein | WP_000213139.1 | 0.836926981 | 3.4058E-06 | 2.0064E-05 |
| STM14_RS12780 | | chemotaxis protein | WP_000368558.1 | 0.834844615 | 4.253E-18 | 1.1863E-16 |
| STM14_RS05955 | | hypothetical protein | WP_000174813.1 | 0.83335154 | 0.0025473 | 0.00731032 |
| STM14_RS02245 | | DUF1889 family protein | WP_001539227.1 | 0.832428856 | 1.603E-20 | 5.7304E-19 |
| STM14_RS10445 | cutC | copper homeostasis protein CutC | WP_001185770.1 | 0.831006075 | 2.2358E-07 | 1.6531E-06 |
| STM14_RS10695 | tnpA | IS200/IS605-like element IS200F family transposase | WP_000502119.1 | 0.830441555 | 3.5205E-07 | 2.5024E-06 |
| STM14_RS16850 | | hypothetical protein | WP_000665657.1 | 0.829863544 | 4.4567E-07 | 3.0903E-06 |
| STM14_RS07935 | ssaO | SPI-2 type III secretion system apparatus protein SsaO | WP_000449089.1 | 0.829363951 | 0.00198694 | 0.0059047 |
| STM14_RS12175 | fghA | S-formylglutathione hydrolase | WP_000425488.1 | 0.828487561 | 2.17E-09 | 2.2011E-08 |
| STM14_RS15520 | invA | type III secretion system export apparatus protein InvA | WP_000927219.1 | 0.828369599 | 2.4621E-11 | 3.4209E-10 |
| STM14_RS10225 | | PagK family vesicle-borne virulence factor | WP_010989029.1 | 0.826439099 | 4.1528E-18 | 1.1672E-16 |
| STM14_RS15490 | spaP | SPI-1 type III secretion system export apparatus protein SpaP | WP_000526016.1 | 0.82598067 | 3.5721E-10 | 4.0846E-09 |
| STM14_RS06645 | tnpA | IS200/IS605-like element IS200F family transposase | WP_000502119.1 | 0.825687121 | 9.5811E-08 | 7.6524E-07 |
| STM14_RS13215 | pgtP | phosphoglycerate transporter PgtP | WP_000955748.1 | 0.825096274 | 0.00869862 | 0.02122487 |
| STM14_RS15080 | proV | glycine betaine/L-proline ABC transporter ATP-binding protein ProV | WP_000985529.1 | 0.819144584 | 4.9779E-13 | 8.5249E-12 |
| STM14_RS17300 | tdcA | transcriptional regulator TdcA | WP_001094910.1 | 0.818847238 | 4.366E-08 | 3.6956E-07 |
| STM14_RS10520 | motB | flagellar motor protein MotB | WP_000795653.1 | 0.818224863 | 1.9326E-14 | 4.0431E-13 |
| STM14_RS07610 | fadK | medium-chain fatty-acid--CoA ligase | WP_000117497.1 | 0.818116555 | 7.5639E-06 | 4.1259E-05 |
| STM14_RS14015 | cadB | cadaverine/lysine antiporter | WP_000100008.1 | 0.817293664 | 6.5612E-12 | 9.7413E-11 |
| STM14_RS18460 | tnpA | IS200/IS605-like element IS200F family transposase | WP_000502119.1 | 0.817034454 | 2.7888E-07 | 2.0174E-06 |
| STM14_RS12615 | | LysR substrate-binding domain-containing protein | WP_000176719.1 | 0.81646821 | 0.00691788 | 0.01739865 |
| STM14_RS17035 | | aryl-sulfate sulfotransferase | WP_000460259.1 | 0.815880495 | 0.00414338 | 0.01116802 |
| STM14_RS08680 | hdeB | acid-activated periplasmic chaperone HdeB | WP_001032161.1 | 0.815253122 | 1.6175E-15 | 3.7457E-14 |
| STM14_RS08575 | | PhoPQ-activated pathogenicity-related family protein | WP_000476912.1 | 0.814706604 | 2.8669E-06 | 1.7136E-05 |
| STM14_RS15390 | orgA | oxygen-regulated invasion protein OrgA | WP_001574876.1 | 0.809004797 | 4.0113E-14 | 7.8982E-13 |
| STM14_RS26830 | | hypothetical protein | | 0.80851433 | 0.01095284 | 0.02575245 |
| STM14_RS22375 | siiF | SPI-4 type I secretion system protein SiiF | WP_000358566.1 | 0.808431694 | 1.0362E-12 | 1.6588E-11 |
| STM14_RS03000 | acrR | multidrug efflux transporter transcriptional repressor AcrR | WP_000101755.1 | 0.808363205 | 8.6001E-06 | 4.6092E-05 |
| STM14_RS19990 | | GntR family transcriptional regulator | WP_000854956.1 | 0.806854671 | 5.8854E-15 | 1.2899E-13 |
| STM14_RS18285 | bfd | bacterioferritin-associated ferredoxin | WP_000289082.1 | 0.806455176 | 4.8721E-07 | 3.3406E-06 |
| STM14_RS15320 | hypC | hydrogenase 3 maturation protein HypC | WP_000334871.1 | 0.804596948 | 0.00011792 | 0.00048349 |
| STM14_RS15455 | sipD | SPI-1 type III secretion system needle tip complex protein SipD | WP_000932246.1 | 0.804288813 | 7.5912E-13 | 1.2423E-11 |
| STM14_RS05000 | | GrxA family glutaredoxin | WP_000495513.1 | 0.803148904 | 2.5963E-15 | 5.901E-14 |

| | | | | | | |
|---|---|---|---|---|---|---|
| STM14_RS16205 | | YfdX family protein | WP_000835265.1 | 0.803101669 | 1.203E-07 | 9.4448E-07 |
| STM14_RS10755 | fliF | flagellar basal-body MS-ring/collar protein FliF | WP_001276834.1 | 0.798581248 | 8.1571E-08 | 6.6301E-07 |
| STM14_RS02505 | | YaiY family protein | WP_000763139.1 | 0.796681697 | 0.00191178 | 0.00571826 |
| STM14_RS23765 | | PTS sugar transporter subunit IIA | WP_000820693.1 | 0.796481219 | 0.01270408 | 0.029217 |
| STM14_RS23600 | | hypothetical protein | WP_000556777.1 | 0.796297834 | 4.4401E-05 | 0.00020694 |
| STM14_RS13220 | | YfdY family protein | WP_000639889.1 | 0.796289512 | 6.8818E-13 | 1.1414E-11 |
| STM14_RS17665 | ptsN | PTS IIA-like nitrogen regulatory protein PtsN | WP_000609332.1 | 0.796112973 | 3.8375E-08 | 3.2707E-07 |
| STM14_RS15395 | prgK | type III secretion system inner membrane ring lipoprotein PrgK | WP_000621238.1 | 0.795577376 | 1.9494E-09 | 1.9882E-08 |
| STM14_RS00020 | repA | replication regulatory protein RepA | WP_010999936.1 | 0.794620787 | 1.4455E-13 | 2.6612E-12 |
| STM14_RS07825 | spiA | SPI-2 type III secretion system protein SpiA | WP_000261284.1 | 0.7928022 | 1.1087E-08 | 1.0335E-07 |
| STM14_RS07820 | spiC | SPI-2 type III secretion system protein SpiC | WP_001738217.1 | 0.792377445 | 5.944E-05 | 0.00026593 |
| STM14_RS24665 | | hypothetical protein | WP_000069465.1 | 0.78870369 | 0.00322154 | 0.00899977 |
| STM14_RS12220 | nfo | deoxyribonuclease IV | WP_000873909.1 | 0.78799311 | 0.00150696 | 0.00461616 |
| STM14_RS07835 | ssaE | YscE family type III secretion system co-chaperone SsaE | WP_000210202.1 | 0.783244124 | 0.00021288 | 0.0008182 |
| STM14_RS07915 | ssaL | SPI-2 type III secretion system apparatus protein SsaL | WP_000238877.1 | 0.782815149 | 1.0621E-11 | 1.5276E-10 |
| STM14_RS07230 | | GlsB/YeaQ/YmgE family stress response membrane protein | WP_000512149.1 | 0.782502425 | 1.3593E-18 | 4.0689E-17 |
| STM14_RS10685 | fliZ | flagella biosynthesis regulatory protein FliZ | WP_000218080.1 | 0.780635825 | 1.3994E-13 | 2.5892E-12 |
| STM14_RS09105 | ldhA | D-lactate dehydrogenase | WP_000762206.1 | 0.779748089 | 8.1967E-10 | 8.6725E-09 |
| STM14_RS05180 | tnpA | IS200/IS605-like element IS200F family transposase | WP_000502119.1 | 0.778382269 | 2.0664E-06 | 1.2555E-05 |
| STM14_RS07815 | | two component system sensor kinase | WP_001050769.1 | 0.778090511 | 2.1025E-11 | 2.9324E-10 |
| STM14_RS12290 | | microcin C ABC transporter permease YejB | WP_000501623.1 | 0.777156146 | 0.01000541 | 0.0238755 |
| STM14_RS22370 | siiE | non-fimbrial adhesin SiiE | WP_000527216.1 | 0.775989168 | 1.8229E-12 | 2.8201E-11 |
| STM14_RS20050 | ivbL | ilvB operon leader peptide IvbL | WP_001541152.1 | 0.77552106 | 0.00282446 | 0.00802444 |
| STM14_RS13275 | | sensor domain-containing phosphodiesterase | WP_001112946.1 | 0.775057203 | 0.00014777 | 0.00059271 |
| STM14_RS15365 | avrA | type III secretion system YopJ family effector AvrA | | 0.774173064 | 3.9261E-10 | 4.448E-09 |
| STM14_RS04930 | | electron transfer flavoprotein-ubiquinone oxidoreductase | WP_001000303.1 | 0.773715588 | 0.00426058 | 0.01140077 |
| STM14_RS26615 | | ryhB-regulated fur leader peptide | | 0.772441083 | 1.1531E-06 | 7.3455E-06 |
| STM14_RS12285 | | extracellular solute-binding protein | WP_000561748.1 | 0.771677969 | 0.00030688 | 0.00113105 |
| STM14_RS17060 | glgS | cell surface composition regulator GlgS | WP_000928927.1 | 0.771334954 | 1.8042E-18 | 5.2723E-17 |
| STM14_RS14265 | | phage portal protein | WP_000831821.1 | 0.767167914 | 0.00040843 | 0.00146002 |
| STM14_RS04800 | ompX | outer membrane protein OmpX | WP_000716763.1 | 0.764864054 | 3.0422E-16 | 7.4676E-15 |
| STM14_RS09215 | | hypothetical protein | | 0.764735524 | 0.01662528 | 0.03654583 |
| STM14_RS23025 | yjfY | DUF1471 family protein YjfY | WP_001521305.1 | 0.763797241 | 0.00022242 | 0.0008513 |
| STM14_RS10095 | mgrB | PhoP/PhoQ regulator MgrB | WP_000714547.1 | 0.763630301 | 1.8066E-08 | 1.6264E-07 |
| STM14_RS09020 | | membrane protein | WP_000997195.1 | 0.761343505 | 0.00109467 | 0.00346566 |
| STM14_RS20600 | | type II toxin-antitoxin system RelE/ParE family toxin | WP_000535852.1 | 0.758626767 | 0.00084613 | 0.00277918 |
| STM14_RS08875 | | hypothetical protein | WP_001597849.1 | 0.756669715 | 2.0874E-05 | 0.0001047 |
| STM14_RS13625 | pbp4b | penicillin binding protein PBP4B | WP_000673300.1 | 0.755457806 | 0.00025126 | 0.00094691 |
| STM14_RS10605 | | glucose-6-phosphate dehydrogenase | WP_000240837.1 | 0.755235991 | 7.4385E-10 | 7.985E-09 |
| STM14_RS08890 | | DUF2554 family protein | WP_000956438.1 | 0.753113575 | 0.0001778 | 0.00069572 |
| STM14_RS03465 | | SIS domain-containing protein | WP_000109360.1 | 0.751770755 | 6.4292E-10 | 6.983E-09 |
| STM14_RS11020 | | terminase large subunit | WP_000088182.1 | 0.751376764 | 0.00032751 | 0.00119988 |
| STM14_RS09980 | tnpA | IS200/IS605-like element IS200F family transposase | WP_000502119.1 | 0.749790442 | 2.5755E-06 | 1.5495E-05 |
| STM14_RS14230 | | RNA-guided endonuclease TnpB family protein | WP_000817264.1 | 0.749754523 | 0.00246383 | 0.00713167 |
| STM14_RS05945 | sopB | SPI-1 type III secretion system effector inositol phosphate phosphatase | WP_001166946.1 | 0.749608579 | 2.4804E-10 | 2.8902E-09 |
| STM14_RS22085 | | glycosyltransferase family 2 protein | WP_000703632.1 | 0.748264169 | 0.00340005 | 0.00941986 |
| STM14_RS14780 | | DUF3883 domain-containing protein | WP_000889010.1 | 0.747808554 | 0.00105047 | 0.00334006 |
| STM14_RS10530 | flhC | flagellar transcriptional regulator FlhC | WP_000605987.1 | 0.747097151 | 9.0566E-11 | 1.1539E-09 |
| STM14_RS06925 | | RNA-guided endonuclease TnpB family protein | WP_000817264.1 | 0.745679147 | 0.00163541 | 0.0049683 |
| STM14_RS14715 | | hypothetical protein | WP_010989063.1 | 0.745314199 | 0.00421101 | 0.01129275 |
| STM14_RS14410 | | hypothetical protein | WP_001539619.1 | 0.743224505 | 0.00213665 | 0.00628366 |
| STM14_RS17050 | ribB | 3%2C4-dihydroxy-2-butanone-4-phosphate synthase | WP_001076978.1 | 0.742736827 | 3.4704E-11 | 4.7326E-10 |
| STM14_RS22350 | siiA | SPI-4 type I secretion system auxiliary protein SiiA | WP_000389233.1 | 0.742005448 | 1.2908E-05 | 6.647E-05 |
| STM14_RS02315 | | membrane protein | WP_001006010.1 | 0.741970326 | 0.00294387 | 0.00831979 |
| STM14_RS09640 | adhE | bifunctional acetaldehyde-CoA/alcohol dehydrogenase | WP_000301678.1 | 0.741037134 | 1.5221E-17 | 4.1209E-16 |
| STM14_RS08090 | mliC | C-type lysozyme inhibitor | WP_001132086.1 | 0.740469605 | 3.7289E-05 | 0.00017625 |
| STM14_RS02805 | | SEL1-like repeat protein | WP_000403151.1 | 0.73475081 | 0.00099846 | 0.00320516 |
| STM14_RS10915 | | hypothetical protein | WP_000554737.1 | 0.734653604 | 3.279E-05 | 0.00015659 |
| STM14_RS10790 | fliM | flagellar motor switch protein FliM | WP_000502811.1 | 0.733583862 | 2.5428E-06 | 1.5323E-05 |
| STM14_RS04320 | cydA | cytochrome ubiquinol oxidase subunit I | WP_000884369.1 | 0.732901538 | 3.6532E-16 | 8.8495E-15 |
| STM14_RS20120 | | YceK/YidQ family lipoprotein | WP_001581211.1 | 0.732773793 | 5.6854E-10 | 6.2675E-09 |
| STM14_RS05975 | hiuH | hydroxyisourate hydrolase | WP_000833188.1 | 0.731721431 | 8.8696E-07 | 5.8007E-06 |
| STM14_RS10775 | fliJ | flagellar export protein FliJ | WP_000046981.1 | 0.731687209 | 0.00366326 | 0.01002833 |
| STM14_RS00075 | | membrane protein | WP_010999940.1 | 0.73054729 | 0.01175501 | 0.02734173 |
| STM14_RS07840 | sseA | SPI-2 type III secretion system chaperone SseA | WP_001738219.1 | 0.730454752 | 2.2186E-09 | 2.2442E-08 |
| STM14_RS08160 | ydgT | transcription modulator YdgT | WP_000217946.1 | 0.729558486 | 1.2393E-06 | 7.827E-06 |
| STM14_RS24670 | | hypothetical protein | WP_014343878.1 | 0.729502363 | 0.00095759 | 0.00310375 |
| STM14_RS10765 | fliH | flagellar assembly protein FliH | WP_000064163.1 | 0.72817378 | 0.00013262 | 0.00053957 |
| STM14_RS21215 | | cytotoxic translational repressor of toxin-antitoxin stability system | WP_001159630.1 | 0.728157467 | 8.1639E-06 | 4.414E-05 |
| STM14_RS04275 | sdhC | succinate dehydrogenase cytochrome b556 subunit | WP_000263340.1 | 0.727796427 | 3.6683E-11 | 4.984E-10 |
| STM14_RS08285 | | MFS transporter | WP_000091796.1 | 0.727266519 | 1.9883E-08 | 1.7812E-07 |
| STM14_RS23585 | | YjhX family toxin | WP_001054380.1 | 0.726479475 | 0.00098782 | 0.00317774 |
| STM14_RS06410 | flgJ | flagellar assembly peptidoglycan hydrolase FlgJ | WP_000578692.1 | 0.724469672 | 0.00011516 | 0.00047388 |
| STM14_RS06025 | hpaX | 4-hydroxyphenylacetate permease | WP_001285800.1 | 0.72419811 | 0.00756539 | 0.0187834 |
| STM14_RS22080 | | DUF6056 family protein | WP_001095011.1 | 0.723873801 | 0.00055318 | 0.00190712 |
| STM14_RS01120 | sgrR | HTH-type transcriptional regulator SgrR | WP_001139040.1 | 0.723467938 | 0.00326571 | 0.00910935 |
| STM14_RS03785 | ybeD | DUF493 family protein YbeD | WP_000850547.1 | 0.720872438 | 1.4638E-09 | 1.514E-08 |
| STM14_RS22270 | zur | zinc uptake transcriptional repressor Zur | WP_000416272.1 | 0.720498713 | 7.6516E-06 | 4.1615E-05 |
| STM14_RS06315 | pyrC | dihydroorotase | WP_000126580.1 | 0.718948326 | 7.8442E-05 | 0.00033741 |
| STM14_RS15460 | sipC | SPI-1 type III secretion system needle tip complex protein SipC | WP_000909019.1 | 0.716399729 | 5.8599E-13 | 9.8521E-12 |
| STM14_RS26010 | steB | type III secretion system effector SteB | WP_001540100.1 | 0.715651805 | 1.8442E-05 | 9.3147E-05 |
| STM14_RS05315 | pflB | formate C-acetyltransferase | WP_001292799.1 | 0.714869411 | 4.3733E-15 | 9.8187E-14 |

| | | | | | | |
|---|---|---|---|---|---|---|
| STM14_RS23035 | rpsF | 30S ribosomal protein S6 | WP_001216673.1 | 0.713737796 | 0.00059805 | 0.00204649 |
| STM14_RS04430 | | FumA C-terminus/TtdB family hydratase beta subunit | WP_000970373.1 | 0.71353405 | 0.00082357 | 0.0027172 |
| STM14_RS16700 | | cupin domain-containing protein | WP_001147546.1 | 0.713164321 | 0.02663848 | 0.0545511 |
| STM14_RS19190 | | aromatic amino acid transport family protein | WP_001150163.1 | 0.711825426 | 0.02781001 | 0.05651019 |
| STM14_RS07050 | pagD | virulence protein PagD | WP_001535993.1 | 0.710785336 | 9.2228E-07 | 5.9786E-06 |
| STM14_RS15405 | prgI | type III secretion system needle filament protein PrgI | WP_000235228.1 | 0.710528881 | 1.7747E-07 | 1.3336E-06 |
| STM14_RS00175 | | helix-turn-helix domain-containing protein | WP_001541541.1 | 0.710071305 | 3.7447E-14 | 7.4935E-13 |
| STM14_RS11150 | | DUF262 domain-containing protein | WP_000071070.1 | 0.709147002 | 0.00475694 | 0.01249291 |
| STM14_RS02130 | | LysR family transcriptional regulator | WP_000970302.1 | 0.708144158 | 1.0833E-05 | 5.6577E-05 |
| STM14_RS15970 | rarD | EamA family transporter RarD | WP_014343890.1 | 0.707758886 | 3.1663E-06 | 1.8773E-05 |
| STM14_RS15450 | sipA | SPI-1 type III secretion system effector SipA | WP_000258812.1 | 0.706007769 | 5.1585E-11 | 6.8569E-10 |
| STM14_RS14350 | | DinI family protein | WP_001217669.1 | 0.705988415 | 0.00047025 | 0.0016594 |
| STM14_RS00540 | vapB | toxin-antitoxin system antitoxin VapB | WP_000450265.1 | 0.702858345 | 2.0737E-07 | 1.5456E-06 |
| STM14_RS26720 | | hypothetical protein | WP_212742690.1 | 0.70128553 | 4.1542E-08 | 3.5243E-07 |
| STM14_RS18330 | fkpA | FKBP-type peptidyl-prolyl cis-trans isomerase | WP_000838227.1 | 0.700860061 | 2.6683E-10 | 3.0993E-09 |
| STM14_RS14805 | | SgrR family transcriptional regulator | WP_000021514.1 | 0.700351076 | 0.01430183 | 0.03222386 |
| STM14_RS05585 | | antitermination protein | WP_001047631.1 | 0.699644011 | 0.00412846 | 0.01114443 |
| STM14_RS23490 | idnO | gluconate 5-dehydrogenase | WP_000998688.1 | 0.699467098 | 0.01316953 | 0.03009945 |
| STM14_RS26890 | | hypothetical protein | WP_212724070.1 | 0.698586895 | 0.00394185 | 0.01069557 |
| STM14_RS05695 | | Ail/Lom family outer membrane beta-barrel protein | WP_000725267.1 | 0.69715871 | 7.2672E-05 | 0.00031554 |
| STM14_RS02330 | | transcriptional regulator | WP_000081378.1 | 0.696804337 | 0.00102887 | 0.0032913 |
| STM14_RS00910 | tnpA | IS200/IS605-like element IS200F family transposase | WP_000502119.1 | 0.695441193 | 4.9211E-06 | 2.8049E-05 |
| STM14_RS16750 | | fructuronate reductase | WP_000437132.1 | 0.693924619 | 6.616E-07 | 4.4453E-06 |
| STM14_RS10535 | flhD | flagellar transcriptional regulator FlhD | WP_001518146.1 | 0.693543601 | 5.7045E-11 | 7.5014E-10 |
| STM14_RS16160 | | amino acid permease | WP_000036825.1 | 0.692410585 | 0.01830183 | 0.03956978 |
| STM14_RS23820 | | threonine/serine exporter ThrE family protein | WP_001674846.1 | 0.692111169 | 0.006128 | 0.01565809 |
| STM14_RS14825 | | membrane protein | WP_010989066.1 | 0.6916267 | 0.00050263 | 0.00176087 |
| STM14_RS26730 | | DUF1983 domain-containing protein | WP_001541993.1 | 0.688688929 | 0.04022217 | 0.0772551 |
| STM14_RS15560 | | DUF1778 domain-containing protein | WP_000855692.1 | 0.687331368 | 0.00522949 | 0.01358856 |
| STM14_RS06805 | | DinI family protein | WP_001217669.1 | 0.68568516 | 0.00016939 | 0.00066633 |
| STM14_RS21220 | | DNA-binding transcriptional regulator | WP_000362050.1 | 0.685420343 | 0.00019969 | 0.00077152 |
| STM14_RS00385 | traM | relaxosome protein TraM | WP_001151546.1 | 0.684118269 | 0.00829286 | 0.02035621 |
| STM14_RS17525 | tnpA | IS200/IS605-like element IS200F family transposase | WP_000502119.1 | 0.683759957 | 2.1989E-05 | 0.00010941 |
| STM14_RS11155 | | ead/Ea22-like family protein | WP_000065095.1 | 0.681712075 | 0.00113998 | 0.00359984 |
| STM14_RS08345 | | dimethyl sulfoxide reductase subunit A | WP_000705294.1 | 0.681562929 | 0.00010139 | 0.00042185 |
| STM14_RS12600 | nrdB | ribonucleotide-diphosphate reductase subunit beta | WP_000332026.1 | 0.680931048 | 4.415E-09 | 4.3582E-08 |
| STM14_RS12630 | glpA | anaerobic glycerol-3-phosphate dehydrogenase subunit A | WP_000857294.1 | 0.680865829 | 0.00185814 | 0.00557595 |
| STM14_RS08220 | ompC | porin OmpC | WP_000824309.1 | 0.679373436 | 0.00067614 | 0.00227565 |
| STM14_RS08530 | hypC | HypC/HybG/HupF family hydrogenase formation chaperone | WP_000334904.1 | 0.67932256 | 0.01219451 | 0.02823322 |
| STM14_RS08555 | | linear amide C-N hydrolase | WP_000976498.1 | 0.678408689 | 2.7169E-09 | 2.7333E-08 |
| STM14_RS15020 | | hypothetical protein | WP_000494942.1 | 0.67735308 | 3.5671E-08 | 3.0687E-07 |
| STM14_RS16400 | fbaA | class II fructose-bisphosphate aldolase | WP_000034386.1 | 0.676051816 | 9.3449E-13 | 1.5091E-11 |
| STM14_RS24090 | | tRNA/rRNA methyltransferase | WP_001266769.1 | 0.675233541 | 0.0041463 | 0.01116802 |
| STM14_RS11540 | sbcB | exodeoxyribonuclease I | WP_000216711.1 | 0.673273573 | 1.1515E-06 | 7.3455E-06 |
| STM14_RS10210 | | DMT family transporter | WP_001233446.1 | 0.673193423 | 0.00037461 | 0.00135093 |
| STM14_RS23250 | iolI2 | 2-keto-myo-inositol isomerase IolI2 | WP_001019386.1 | 0.670530401 | 0.00709532 | 0.01778419 |
| STM14_RS08075 | slyA | transcriptional regulator SlyA | WP_000764553.1 | 0.669648348 | 2.3678E-13 | 4.2322E-12 |
| STM14_RS07895 | ssaI | SctI family type III secretion system inner rod subunit SsaI | WP_001574409.1 | 0.667811919 | 7.8864E-05 | 0.00033883 |
| STM14_RS15400 | prgJ | type III secretion system inner rod protein PrgJ | WP_000020431.1 | 0.667263928 | 2.1165E-07 | 1.5712E-06 |
| STM14_RS21500 | | aquaporin | WP_000084285.1 | 0.666701048 | 4.8118E-06 | 2.7554E-05 |
| STM14_RS22310 | aphA | acid phosphatase AphA | WP_000724435.1 | 0.666657893 | 9.2714E-06 | 4.926E-05 |
| STM14_RS05855 | yccS | YccS family putative transporter | WP_000950880.1 | 0.665520737 | 0.00010085 | 0.00042054 |
| STM14_RS18150 | rpsK | 30S ribosomal protein S11 | WP_001029758.1 | 0.665386028 | 7.994E-12 | 1.168E-10 |
| STM14_RS08080 | slyB | outer membrane lipoprotein SlyB | WP_000597179.1 | 0.665270434 | 1.1355E-12 | 1.8099E-11 |
| STM14_RS08695 | sra | stationary-phase-induced ribosome-associated protein | WP_000841559.1 | 0.6639099 | 1.951E-11 | 2.7418E-10 |
| STM14_RS09025 | sseJ | SPI-2 type III secretion system effector SseJ | WP_001131430.1 | 0.662519357 | 0.00014869 | 0.00059574 |
| STM14_RS02395 | | MFS transporter | WP_000288496.1 | 0.66064975 | 0.00060062 | 0.00205146 |
| STM14_RS05270 | dmsA | dimethylsulfoxide reductase subunit A | WP_000850250.1 | 0.659961656 | 6.9499E-14 | 1.3259E-12 |
| STM14_RS09850 | appC | cytochrome bd-II oxidase subunit 1 | WP_000263611.1 | 0.659275501 | 0.0417833 | 0.07979569 |
| STM14_RS08610 | | type II toxin-antitoxin system Phd/YefM family antitoxin | WP_000885424.1 | 0.656371743 | 0.00019136 | 0.00074402 |
| STM14_RS10460 | mrdA | penicillin-binding protein 2 | WP_000142877.1 | 0.654849762 | 0.04906984 | 0.09157383 |
| STM14_RS22505 | phnO | aminoalkylphosphonate N-acetyltransferase | WP_001526214.1 | 0.654765584 | 0.00010879 | 0.00045059 |
| STM14_RS16760 | | methyl-accepting chemotaxis protein | WP_000475246.1 | 0.653070738 | 1.4569E-09 | 1.511E-08 |
| STM14_RS17205 | yqjA | DedA family general envelope maintenance protein YqjA | WP_000422143.1 | 0.652401926 | 4.1385E-07 | 2.8969E-06 |
| STM14_RS07930 | ssaN | SctN family type III secretion system ATPase SsaN | WP_000787213.1 | 0.652399171 | 2.7984E-06 | 1.6772E-05 |
| STM14_RS21505 | zapB | septal ring assembly protein ZapB | WP_000051370.1 | 0.651991604 | 7.6325E-13 | 1.2435E-11 |
| STM14_RS08740 | | GFA family protein | WP_014343841.1 | 0.651097492 | 0.00027633 | 0.00102564 |
| STM14_RS00985 | | hypothetical protein | WP_000802644.1 | 0.649527655 | 8.5007E-08 | 6.8489E-07 |
| STM14_RS10090 | | YebO family protein | WP_001000660.1 | 0.648507812 | 1.5293E-07 | 1.161E-06 |
| STM14_RS08970 | aac(6') | cryptic aminoglycoside N-acetyltransferase AAC(6')-Iy/Iaa | WP_000354867.1 | 0.647433129 | 0.02903444 | 0.05857798 |
| STM14_RS05870 | mgsA | methylglyoxal synthase | WP_000424187.1 | 0.647132558 | 3.7753E-06 | 2.196E-05 |
| STM14_RS18025 | | membrane protein | WP_000825645.1 | 0.644190307 | 0.00063712 | 0.00216237 |
| STM14_RS12650 | sseL | SPI-2 type III secretion system effector deubiquitinase SseL | WP_001017721.1 | 0.64371493 | 0.0001904 | 0.00074108 |
| STM14_RS10505 | tar | methyl-accepting chemotaxis protein II | WP_000483274.1 | 0.642940349 | 8.0056E-14 | 1.5039E-12 |
| STM14_RS20500 | rbsB | ribose ABC transporter substrate-binding protein RbsB | WP_001056260.1 | 0.642396469 | 2.6581E-14 | 5.3776E-13 |
| STM14_RS02970 | | YlaC family protein | WP_000136181.1 | 0.64233029 | 0.00932349 | 0.02246667 |
| STM14_RS04925 | | acyl-CoA dehydrogenase family protein | WP_000835387.1 | 0.642278172 | 0.00896712 | 0.02170739 |
| STM14_RS09140 | | SMR family transporter | WP_000159242.1 | 0.637613049 | 0.00120393 | 0.00377586 |
| STM14_RS21880 | rsd | sigma D regulator | WP_000934315.1 | 0.635682739 | 0.000128 | 0.00052193 |
| STM14_RS22540 | eptA | phosphoethanolamine transferase EptA | WP_000919289.1 | 0.635341757 | 4.9483E-07 | 3.3865E-06 |
| STM14_RS26360 | | YciY family protein | WP_001531561.1 | 0.634826832 | 7.8387E-06 | 4.2569E-05 |
| STM14_RS24535 | mntS | manganase accumulation protein MntS | WP_001001755.1 | 0.634363455 | 5.1702E-08 | 4.3167E-07 |

| | | | | | | |
|---|---|---|---|---|---|---|
| STM14_RS10610 | | SH3 domain-containing C40 family peptidase | WP_010989033.1 | 0.633631403 | 1.9602E-06 | 1.1969E-05 |
| STM14_RS06130 | | sodium:solute symporter | WP_000628082.1 | 0.633631252 | 4.6271E-07 | 3.1905E-06 |
| STM14_RS07765 | | PfkB family carbohydrate kinase | WP_000471807.1 | 0.632442641 | 0.03593103 | 0.07007311 |
| STM14_RS18795 | | glutamine amidotransferase | WP_001066509.1 | 0.629875787 | 0.04491383 | 0.08506827 |
| STM14_RS13080 | | hypothetical protein | WP_001051266.1 | 0.629542739 | 0.05167656 | 0.09595215 |
| STM14_RS06830 | | DUF1133 family protein | WP_000639984.1 | 0.629327338 | 0.0338681 | 0.06640167 |
| STM14_RS23075 | ytfE | iron-sulfur cluster repair protein YtfE | WP_000331468.1 | 0.629051176 | 0.03844211 | 0.0742233 |
| STM14_RS07215 | | DUF1869 domain-containing protein | WP_000513733.1 | 0.628879147 | 0.00559448 | 0.01447568 |
| STM14_RS18400 | | YccS/YhfK family putative transporter | WP_000269311.1 | 0.628500167 | 2.4091E-06 | 1.4565E-05 |
| STM14_RS23425 | rraB | ribonuclease E inhibitor RraB | WP_000002940.1 | 0.627920495 | 9.7224E-13 | 1.5632E-11 |
| STM14_RS03935 | | amino acid ABC transporter substrate-binding protein | WP_001526121.1 | 0.626243184 | 2.5386E-11 | 3.5139E-10 |
| STM14_RS05700 | sodC1 | superoxide dismutase [Cu-Zn] SodC1 | WP_000877926.1 | 0.626145616 | 2.3691E-12 | 3.6046E-11 |
| STM14_RS17865 | yhcN | peroxide/acid stress response protein YhcN | WP_000695693.1 | 0.625777576 | 0.0009493 | 0.00307957 |
| STM14_RS01480 | | polysaccharide deacetylase family protein | WP_000245304.1 | 0.625266864 | 8.1892E-06 | 4.4212E-05 |
| STM14_RS10085 | | DUF2627 domain-containing protein | WP_001537930.1 | 0.625166673 | 3.8132E-13 | 6.6227E-12 |
| STM14_RS12260 | yeiP | elongation factor P-like protein YeiP | WP_001136822.1 | 0.62507367 | 6.3685E-05 | 0.00028218 |
| STM14_RS09465 | | YmiA family putative membrane protein | WP_213083417.1 | 0.624130205 | 3.9719E-07 | 2.7963E-06 |
| STM14_RS03170 | sfbB | virulence-associated ABC transporter ATP-binding protein SfbB | WP_000569660.1 | 0.623733236 | 0.04897956 | 0.09148781 |
| STM14_RS15430 | sptP | SPI-1 type III secretion system effector GTPase-activating protein SptP | WP_000922300.1 | 0.621857079 | 2.0046E-05 | 0.0001007 |
| STM14_RS18125 | zntR | Zn(2+)-responsive transcriptional regulator | WP_000285585.1 | 0.621018178 | 0.00348053 | 0.00959947 |
| STM14_RS06880 | | phage terminase large subunit family protein | WP_000623088.1 | 0.620733215 | 0.02751001 | 0.05605526 |
| STM14_RS14530 | | DUF4385 domain-containing protein | WP_000985653.1 | 0.620200336 | 1.6908E-05 | 8.5871E-05 |
| STM14_RS04710 | | ABC transporter permease | WP_000469050.1 | 0.618957 | 2.2163E-05 | 0.00011013 |
| STM14_RS08495 | | bestrophin family protein | WP_001670736.1 | 0.617621159 | 0.02689228 | 0.05495122 |
| STM14_RS09830 | hyaC | Ni/Fe-hydrogenase b-type cytochrome subunit | WP_000147036.1 | 0.616945121 | 0.02404541 | 0.04971096 |
| STM14_RS04345 | ybgC | tol-pal system-associated acyl-CoA thioesterase | WP_001046240.1 | 0.616515211 | 6.8996E-06 | 3.8088E-05 |
| STM14_RS14145 | gogB | type III secretion effector GogB | WP_010989047.1 | 0.61609686 | 4.4061E-05 | 0.00020562 |
| STM14_RS06745 | | hypothetical protein | WP_000373340.1 | 0.615970595 | 0.03192208 | 0.06326002 |
| STM14_RS09910 | | SpoVR family protein | WP_001240769.1 | 0.615079801 | 4.672E-10 | 5.2287E-09 |
| STM14_RS23495 | idnD | L-idonate 5-dehydrogenase | WP_000453340.1 | 0.614327848 | 0.00623391 | 0.01590663 |
| STM14_RS10010 | | YoaH family protein | WP_000457328.1 | 0.611028977 | 4.6318E-05 | 0.00021479 |
| STM14_RS10905 | sseK | type III secretion system effector arginine glycosyltransferase | WP_000492926.1 | 0.610863883 | 0.01977071 | 0.04224942 |
| STM14_RS07965 | ssaU | SPI-2 type III secretion system apparatus protein SsaU | WP_001291716.1 | 0.609263072 | 0.00370111 | 0.01009442 |
| STM14_RS11120 | | helix-turn-helix transcriptional regulator | WP_001020636.1 | 0.608780324 | 1.694E-09 | 1.7472E-08 |
| STM14_RS24975 | | hypothetical protein | WP_014343859.1 | 0.606833513 | 1.2963E-07 | 1.0069E-06 |
| STM14_RS07655 | ydiB | quinate/shikimate dehydrogenase | WP_000383490.1 | 0.605711016 | 0.02534676 | 0.05212528 |
| STM14_RS15515 | spaK | SPI-1 type III secretion system chaperone SpaK | WP_001164066.1 | 0.605335058 | 1.4429E-07 | 1.1091E-06 |
| STM14_RS10750 | fliE | flagellar hook-basal body complex protein FliE | WP_000719036.1 | 0.604278423 | 0.03043281 | 0.06093529 |
| STM14_RS08290 | | LysR family transcriptional regulator | WP_001019577.1 | 0.604176825 | 0.03344963 | 0.06579142 |
| STM14_RS06155 | | Gfo/Idh/MocA family oxidoreductase | WP_000988585.1 | 0.603391475 | 3.999E-07 | 2.81E-06 |
| STM14_RS11645 | | MATE family efflux transporter | WP_001531712.1 | 0.600564385 | 0.0003955 | 0.00141931 |
| STM14_RS22530 | pmrB | two-component system sensor histidine kinase PmrB | WP_001212189.1 | 0.599474944 | 2.6957E-05 | 0.00013077 |
| STM14_RS12620 | glpQ | glycerophosphodiester phosphodiesterase | WP_000858952.1 | 0.598685471 | 3.0842E-09 | 3.0858E-08 |
| STM14_RS02850 | bolA | transcriptional regulator BolA | WP_000973433.1 | 0.598366924 | 2.4255E-10 | 2.8352E-09 |
| STM14_RS14285 | | PPC domain-containing protein | WP_000669690.1 | 0.598070074 | 0.00016564 | 0.00065298 |
| STM14_RS00750 | | anaerobic sulfatase maturase | WP_000971148.1 | 0.596884978 | 0.03203541 | 0.06338227 |
| STM14_RS08625 | | hypothetical protein | WP_010989018.1 | 0.59554979 | 0.00016513 | 0.00065165 |
| STM14_RS23740 | yjiA | GTPase | WP_000187839.1 | 0.594596073 | 0.00139594 | 0.00430834 |
| STM14_RS11225 | | hypothetical protein | WP_000669109.1 | 0.59451495 | 0.00568809 | 0.01465609 |
| STM14_RS07810 | | two component system response regulator | WP_000666335.1 | 0.592778615 | 2.0132E-06 | 1.2252E-05 |
| STM14_RS03440 | | DnaJ family molecular chaperone | WP_001033457.1 | 0.592033313 | 1.4324E-06 | 8.909E-06 |
| STM14_RS08845 | | virulence factor SrfB | WP_000960814.1 | 0.591699209 | 0.00038157 | 0.00137471 |
| STM14_RS13705 | bepA | beta-barrel assembly-enhancing protease | WP_000489630.1 | 0.591670622 | 1.6023E-05 | 8.16E-05 |
| STM14_RS15310 | hypA | hydrogenase maturation nickel metallochaperone HypA | WP_000544461.1 | 0.591072277 | 1.0957E-05 | 5.7142E-05 |
| STM14_RS15500 | spaN | SPI-1 type III secretion system protein SpaN | WP_000503098.1 | 0.590257336 | 7.0851E-06 | 3.8936E-05 |
| STM14_RS02490 | ampH | D-alanyl-D-alanine-carboxypeptidase/endopeptidase AmpH | WP_000830784.1 | 0.59024005 | 0.01130602 | 0.0264814 |
| STM14_RS21755 | coaA | type I pantothenate kinase | WP_000023069.1 | 0.585389783 | 6.2798E-05 | 0.00027892 |
| STM14_RS16285 | sdhE | FAD assembly factor SdhE | WP_000351186.1 | 0.585040968 | 2.272E-05 | 0.00011274 |
| STM14_RS17165 | ygjG | putrescine aminotransferase | WP_001520281.1 | 0.583782919 | 2.5085E-06 | 1.5141E-05 |
| STM14_RS22645 | | AraC family transcriptional regulator | WP_000921674.1 | 0.583684857 | 0.10463442 | 0.17535909 |
| STM14_RS10690 | | RNA polymerase sigma factor FliA | WP_001087453.1 | 0.583439956 | 2.7818E-08 | 2.4329E-07 |
| STM14_RS07870 | sscB | SycD/LcrH family type III secretion system chaperone SscB | WP_000979963.1 | 0.582973235 | 0.01003259 | 0.02392488 |
| STM14_RS16995 | | DUF1249 family protein | WP_000833405.1 | 0.582727133 | 1.411E-05 | 7.2459E-05 |
| STM14_RS10830 | yodD | YodD family peroxide/acid resistance protein | WP_000844798.1 | 0.581578444 | 9.319E-09 | 8.8663E-08 |
| STM14_RS23370 | ridA | 2-iminobutanoate/2-iminopropanoate deaminase | WP_000047544.1 | 0.581307073 | 6.1938E-09 | 6.0332E-08 |
| STM14_RS20230 | yidA | sugar-phosphatase | WP_000985572.1 | 0.580922899 | 2.6518E-05 | 0.00012915 |
| STM14_RS03365 | | diguanylate cyclase | WP_000226489.1 | 0.580637621 | 0.01786265 | 0.0387568 |
| STM14_RS21585 | | hypothetical protein | WP_001134807.1 | 0.579460155 | 0.00461148 | 0.01216294 |
| STM14_RS13750 | | sensor domain-containing phosphodiesterase | WP_000773647.1 | 0.577829954 | 0.00695267 | 0.0174623 |
| STM14_RS11515 | dacD | serine-type D-Ala-D-Ala carboxypeptidase DacD | WP_000925042.1 | 0.577434389 | 0.00882793 | 0.02148342 |
| STM14_RS13590 | tnpA | IS200/IS605-like element IS200F family transposase | WP_000502119.1 | 0.576935348 | 0.00012485 | 0.00051077 |
| STM14_RS10285 | yobA | CopC domain-containing protein YobA | WP_000168393.1 | 0.576667822 | 0.00079192 | 0.00262688 |
| STM14_RS18645 | | type II toxin-antitoxin system YafQ family toxin | WP_001526148.1 | 0.576368311 | 0.00086038 | 0.00281342 |
| STM14_RS12900 | pta | phosphate acetyltransferase | WP_000086692.1 | 0.576355565 | 6.9498E-06 | 3.8307E-05 |
| STM14_RS18000 | | DUF2556 family protein | WP_000620015.1 | 0.576090365 | 0.00825056 | 0.02028241 |
| STM14_RS16165 | rcnR | Ni(II)/Co(II)-binding transcriptional repressor RcnR | WP_000019953.1 | 0.575788864 | 0.00575463 | 0.01480682 |
| STM14_RS10615 | | membrane protein | WP_001682427.1 | 0.575737555 | 0.00239082 | 0.00694789 |
| STM14_RS11955 | | fimbrial biogenesis outer membrane usher protein | WP_000945364.1 | 0.575146242 | 0.02079923 | 0.0441145 |
| STM14_RS22545 | adiC | arginine/agmatine antiporter | WP_000093130.1 | 0.5747989 | 0.02328339 | 0.04843472 |
| STM14_RS19310 | | HTH-type transcriptional regulator | WP_000455790.1 | 0.57384186 | 1.2736E-10 | 1.6059E-09 |
| STM14_RS03750 | cspE | transcription antiterminator/RNA stability regulator CspE | WP_000034826.1 | 0.572669423 | 2.3252E-10 | 2.7353E-09 |
| STM14_RS12075 | dusC | tRNA dihydrouridine(16) synthase DusC | WP_001264832.1 | 0.572402028 | 0.08511006 | 0.14733203 |

| | | | | | | |
|---|---|---|---|---|---|---|
| STM14_RS19675 | rfaP | lipopolysaccharide core heptose(I) kinase RfaP | WP_000229825.1 | 0.572005971 | 0.00448422 | 0.01187834 |
| STM14_RS16220 | vapB | toxin-antitoxin system antitoxin VapB | WP_000557549.1 | 0.571649698 | 1.6156E-08 | 1.4652E-07 |
| STM14_RS17500 | | DUF1963 domain-containing protein | WP_000051504.1 | 0.570343115 | 0.00164253 | 0.00498582 |
| STM14_RS18130 | | DUF1992 domain-containing protein | WP_000266515.1 | 0.568623585 | 0.01558187 | 0.03462429 |
| STM14_RS04980 | | MFS transporter | WP_000217433.1 | 0.568376261 | 0.00045178 | 0.00159793 |
| STM14_RS02560 | yaiA | protein YaiA | WP_001143165.1 | 0.567370627 | 2.1298E-07 | 1.5778E-06 |
| STM14_RS08425 | | 2-oxo-tetronate isomerase | WP_001235357.1 | 0.567332502 | 0.00063468 | 0.00215779 |
| STM14_RS00270 | | AAA family ATPase | WP_000427676.1 | 0.566633265 | 7.3293E-06 | 4.0159E-05 |
| STM14_RS07360 | | YnjH family protein | WP_001254681.1 | 0.566038369 | 1.4994E-05 | 7.6783E-05 |
| STM14_RS14310 | | hypothetical protein | WP_000657897.1 | 0.564955063 | 0.00013572 | 0.00055097 |
| STM14_RS15445 | | acyl carrier protein | WP_001055577.1 | 0.560077591 | 3.1292E-05 | 0.00015002 |
| STM14_RS19315 | cspA | RNA chaperone/antiterminator CspA | WP_000014594.1 | 0.559525327 | 8.496E-08 | 6.8489E-07 |
| STM14_RS19975 | | PTS galactitol transporter subunit IIC | WP_000997824.1 | 0.558972381 | 0.00305811 | 0.00858228 |
| STM14_RS01340 | pdhR | pyruvate dehydrogenase complex transcriptional repressor PdhR | WP_000331771.1 | 0.558629135 | 1.1838E-05 | 6.1216E-05 |
| STM14_RS23540 | | DUF4435 domain-containing protein | WP_010989100.1 | 0.55822335 | 0.00297393 | 0.00837796 |
| STM14_RS05885 | hspQ | heat shock protein HspQ | WP_000561983.1 | 0.556954603 | 8.9192E-05 | 0.00037739 |
| STM14_RS05815 | pqiC | membrane integrity-associated transporter subunit PqiC | WP_000759136.1 | 0.555987246 | 0.00415999 | 0.01119669 |
| STM14_RS07945 | ssaQ | SPI-2 type III secretion system cytoplasmic ring protein SsaQ | WP_000944192.1 | 0.555254784 | 0.00020907 | 0.00080438 |
| STM14_RS05960 | | C69 family dipeptidase | WP_000760228.1 | 0.555069737 | 6.3396E-10 | 6.9266E-09 |
| STM14_RS07520 | | O-antigen polymerase | WP_000905563.1 | 0.554460689 | 0.0003296 | 0.00120634 |
| STM14_RS23530 | brxL | protease Lon-related BREX system protein BrxL | WP_001193068.1 | 0.55185579 | 3.545E-07 | 2.515E-06 |
| STM14_RS07435 | chbR | transcriptional regulator ChbR | WP_000983393.1 | 0.551031209 | 0.00340727 | 0.00943275 |
| STM14_RS16600 | | YggL family protein | WP_001107574.1 | 0.550952111 | 3.3156E-07 | 2.3751E-06 |
| STM14_RS14665 | bamE | outer membrane protein assembly factor BamE | WP_001203445.1 | 0.550852523 | 0.0011721 | 0.00368546 |
| STM14_RS16955 | | YgiW/YdeI family stress tolerance OB fold protein | WP_000731552.1 | 0.549987643 | 1.0179E-05 | 5.3391E-05 |
| STM14_RS13640 | acrD | multidrug efflux RND transporter permease AcrD | WP_001263072.1 | 0.549569332 | 0.00444516 | 0.01180033 |
| STM14_RS18245 | rpsS | 30S ribosomal protein S19 | WP_001138115.1 | 0.549150048 | 2.0395E-08 | 1.8182E-07 |
| STM14_RS17585 | sfsB | DNA-binding transcriptional regulator SfsB | WP_000444168.1 | 0.549059788 | 0.08391768 | 0.14581637 |
| STM14_RS04805 | | phosphoethanolamine transferase | WP_001089314.1 | 0.548901941 | 0.01709059 | 0.03730145 |
| STM14_RS02875 | hupB | DNA-binding protein HU-beta | WP_254888479.1 | 0.547462539 | 7.1602E-11 | 9.2182E-10 |
| STM14_RS04105 | | pyridoxamine 5'-phosphate oxidase family protein | WP_000070056.1 | 0.547389832 | 0.00358131 | 0.00982591 |
| STM14_RS05915 | yccA | FtsH protease modulator YccA | WP_000374050.1 | 0.547368192 | 4.8766E-05 | 0.00022444 |
| STM14_RS04265 | | citrate synthase | WP_000785811.1 | 0.54683825 | 9.9065E-08 | 7.8782E-07 |
| STM14_RS00070 | | hypothetical protein | WP_014343944.1 | 0.545982445 | 4.9427E-08 | 4.1455E-07 |
| STM14_RS04985 | | TetR/AcrR family transcriptional regulator | WP_000661875.1 | 0.54571853 | 0.0033119 | 0.00921028 |
| STM14_RS04670 | tnpA | IS200/IS605-like element IS200F family transposase | WP_000502119.1 | 0.544819861 | 0.00018763 | 0.00073106 |
| STM14_RS17255 | | pirin family protein | WP_000633547.1 | 0.544657553 | 0.06563986 | 0.11806837 |
| STM14_RS10195 | | hypothetical protein | WP_001617922.1 | 0.544627335 | 2.2956E-05 | 0.00011376 |
| STM14_RS24055 | | fimbrial outer membrane usher protein | WP_001160706.1 | 0.543947105 | 0.03222624 | 0.06372558 |
| STM14_RS00830 | | response regulator | WP_000377406.1 | 0.542622269 | 0.02499014 | 0.05143303 |
| STM14_RS20555 | hdfR | HTH-type transcriptional regulator HdfR | WP_001541181.1 | 0.541624208 | 8.8165E-06 | 4.7047E-05 |
| STM14_RS23330 | nrdG | anaerobic ribonucleoside-triphosphate reductase-activating protein | WP_001268859.1 | 0.540959423 | 0.02658957 | 0.05448124 |
| STM14_RS06015 | hpaH | 2-oxo-hepta-3-ene-1%2C7-dioic acid hydratase | WP_000884795.1 | 0.540713046 | 0.03763234 | 0.07292675 |
| STM14_RS09390 | | hypothetical protein | WP_010989025.1 | 0.539514889 | 0.03355723 | 0.06586482 |
| STM14_RS08595 | | S-adenosylmethionine:tRNA ribosyltransferase-isomerase | WP_000954688.1 | 0.539354936 | 0.00016254 | 0.00064281 |
| STM14_RS07740 | lpp | murein lipoprotein Lpp | WP_001082307.1 | 0.538054108 | 3.9957E-10 | 4.5129E-09 |
| STM14_RS07125 | | TRL-like family protein | WP_000182479.1 | 0.536905616 | 1.8505E-05 | 9.3334E-05 |
| STM14_RS06575 | bhsA | multiple stress resistance protein BhsA | WP_000800159.1 | 0.536461332 | 7.9421E-05 | 0.00034082 |
| STM14_RS05385 | ycaR | protein YcaR | WP_000350061.1 | 0.536165872 | 0.0041963 | 0.01126149 |
| STM14_RS02545 | proC | pyrroline-5-carboxylate reductase | WP_000412197.1 | 0.535258782 | 0.00055293 | 0.00190712 |
| STM14_RS22720 | yjeH | L-methionine/branched-chain amino acid transporter | WP_001670701.1 | 0.534766292 | 0.00069873 | 0.00234309 |
| STM14_RS05895 | | YbhB/YbcL family Raf kinase inhibitor-like protein | WP_000859419.1 | 0.534241721 | 0.02127135 | 0.04503801 |
| STM14_RS07455 | cedA | cell division activator CedA | WP_000977510.1 | 0.532886724 | 0.02155101 | 0.04549931 |
| STM14_RS11630 | rfbN | O antigen biosynthesis rhamnosyltransferase RfbN | WP_000705151.1 | 0.532070044 | 3.5332E-05 | 0.00016786 |
| STM14_RS08440 | ydeE | efflux MFS transporter YdeE | WP_001068050.1 | 0.530401391 | 0.06249726 | 0.11330129 |
| STM14_RS09845 | | hydrogenase expression/formation protein | WP_000063377.1 | 0.529628153 | 0.08961512 | 0.15404429 |
| STM14_RS03140 | | SDR family oxidoreductase | WP_001172802.1 | 0.528189156 | 0.01526547 | 0.03410648 |
| STM14_RS19720 | coaBC | bifunctional phosphopantothenoylcysteine decarboxylase/phosphopan | WP_000050168.1 | 0.526888851 | 0.00064392 | 0.00217915 |
| STM14_RS01900 | rnhA | ribonuclease HI | WP_000917872.1 | 0.526751544 | 0.00818971 | 0.02018374 |
| STM14_RS15800 | eno | phosphopyruvate hydratase | WP_000036734.1 | 0.526133559 | 5.0707E-07 | 3.4639E-06 |
| STM14_RS05275 | dmsB | dimethylsulfoxide reductase subunit B | WP_000213069.1 | 0.525024193 | 1.0699E-05 | 5.5958E-05 |
| STM14_RS11600 | wzzB | LPS O-antigen chain length determinant protein WzzB | WP_000215261.1 | 0.523631197 | 1.5061E-07 | 1.1481E-06 |
| STM14_RS18230 | rplP | 50S ribosomal protein L16 | WP_000941208.1 | 0.522051811 | 2.8566E-08 | 2.4865E-07 |
| STM14_RS06580 | | L%2CD-transpeptidase family protein | WP_000482967.1 | 0.521571585 | 0.01550523 | 0.03453736 |
| STM14_RS18335 | | SlyX family protein | WP_001152701.1 | 0.521423094 | 0.00023114 | 0.0008792 |
| STM14_RS15955 | | YgdI/YgdR family lipoprotein | WP_000750393.1 | 0.52088981 | 3.4794E-09 | 3.4719E-08 |
| STM14_RS15435 | sicP | chaperone SicP | WP_001147852.1 | 0.520275347 | 0.00025584 | 0.00096035 |
| STM14_RS09340 | pspA | phage shock protein PspA | WP_000510996.1 | 0.519899257 | 6.5019E-07 | 4.3846E-06 |
| STM14_RS10350 | pyk | pyruvate kinase | WP_000091156.1 | 0.519652799 | 9.664E-06 | 5.1125E-05 |
| STM14_RS09100 | | YdbH family protein | WP_000677012.1 | 0.516633901 | 0.00139256 | 0.00430502 |
| STM14_RS08915 | tehB | tellurite resistance methyltransferase TehB | WP_000218280.1 | 0.516561892 | 0.00024358 | 0.00092364 |
| STM14_RS18155 | rpsM | 30S ribosomal protein S13 | WP_000090780.1 | 0.51537033 | 8.4485E-08 | 6.8368E-07 |
| STM14_RS09720 | narX | nitrate/nitrite two-component system sensor histidine kinase NarX | WP_000476244.1 | 0.514873249 | 0.00246575 | 0.00713167 |
| STM14_RS11500 | | DUF496 family protein | WP_000450405.1 | 0.51466436 | 3.8579E-07 | 2.7212E-06 |
| STM14_RS18345 | | YheV family putative zinc ribbon protein | WP_001007735.1 | 0.514180227 | 0.00264246 | 0.00757164 |
| STM14_RS18940 | | methyl-accepting chemotaxis citrate transducer | WP_000789683.1 | 0.512654254 | 6.8757E-09 | 6.6447E-08 |
| STM14_RS10635 | pgsA | CDP-diacylglycerol--glycerol-3-phosphate 3-phosphatidyltransferase | WP_001160192.1 | 0.512224031 | 0.00017761 | 0.0006957 |
| STM14_RS20335 | adeP | adenine permease AdeP | WP_001519317.1 | 0.511784288 | 0.00161697 | 0.00492447 |
| STM14_RS08605 | | type II toxin-antitoxin system RelE/ParE family toxin | WP_000221343.1 | 0.511459184 | 0.00840322 | 0.02058593 |
| STM14_RS11690 | rfbB | dTDP-glucose 4%2C6-dehydratase | WP_000697840.1 | 0.510114512 | 7.5759E-07 | 5.0625E-06 |
| STM14_RS17510 | secG | preprotein translocase subunit SecG | WP_000272680.1 | 0.509679258 | 1.9975E-06 | 1.2177E-05 |
| STM14_RS06650 | | hypothetical protein | WP_001113672.1 | 0.509056529 | 0.00907195 | 0.02191792 |

| | | | | | | |
|---|---|---|---|---|---|---|
| STM14_RS15115 | | carbohydrate porin | WP_000645139.1 | 0.508453627 | 0.04120246 | 0.07880906 |
| STM14_RS18435 | nirB | NADPH-nitrite reductase large subunit | WP_000049249.1 | 0.507257689 | 0.0335585 | 0.06586482 |
| STM14_RS06420 | flgL | flagellar hook-associated protein FlgL | WP_001223033.1 | 0.506968124 | 3.4869E-07 | 2.4833E-06 |
| STM14_RS03485 | | PTS system mannose/fructose/N-acetylgalactosamine-transporter subu | WP_000594177.1 | 0.506789598 | 0.02393867 | 0.04951807 |
| STM14_RS06665 | phoP | two-component system response regulator PhoP | WP_001537766.1 | 0.506784452 | 1.3654E-08 | 1.26E-07 |
| STM14_RS15495 | spaO | SPI-1 type III secretion system protein SpaO | WP_000058734.1 | 0.505663273 | 0.00011166 | 0.00046142 |
| STM14_RS05310 | sopD2 | SPI-2 type III secretion system effector SopD2 | WP_001145561.1 | 0.50535748 | 0.15363221 | 0.23979389 |
| STM14_RS06145 | | oligogalacturonate-specific porin KdgM family protein | WP_000700610.1 | 0.505351626 | 0.01411706 | 0.0318537 |
| STM14_RS15100 | mprA | transcriptional repressor MprA | WP_000378431.1 | 0.505119035 | 6.1528E-06 | 3.4257E-05 |
| STM14_RS23320 | | type II toxin-antitoxin system RelB/DinJ family antitoxin | WP_000212724.1 | 0.504652975 | 0.01559468 | 0.03463186 |
| STM14_RS15465 | sipB | SPI-1 type III secretion system needle tip complex protein SipB | WP_000245788.1 | 0.504561111 | 4.2408E-05 | 0.00019841 |
| STM14_RS07190 | | biofilm/acid-resistance regulator YmgB/AriR | WP_000208089.1 | 0.504422206 | 4.6999E-08 | 3.969E-07 |
| STM14_RS16920 | | TRAP transporter substrate-binding protein | WP_000799367.1 | 0.503933682 | 0.00024795 | 0.00093734 |
| STM14_RS09585 | tonB | TonB system transport protein TonB | WP_001520573.1 | 0.503386732 | 0.01943704 | 0.04168153 |
| STM14_RS12730 | pmrD | signal transduction protein PmrD | WP_000455132.1 | 0.503345435 | 1.2809E-07 | 9.992E-07 |
| STM14_RS09420 | | DNA-binding transcriptional regulator YciT | WP_001088593.1 | 0.503259736 | 0.00195377 | 0.00582492 |
| STM14_RS07890 | | EscG/YscG/SsaH family type III secretion system needle protein co-cha | WP_000444713.1 | 0.503165427 | 0.01118012 | 0.02623659 |
| STM14_RS14925 | | antimicrobial resistance protein Mig-14 | WP_000701506.1 | 0.503126493 | 3.6162E-08 | 3.0965E-07 |
| STM14_RS18720 | | dihydrodipicolinate synthase family protein | WP_000192193.1 | 0.502893268 | 0.03953837 | 0.07614031 |
| STM14_RS20595 | ilvA | threonine ammonia-lyase%2C biosynthetic | WP_001518457.1 | 0.502707753 | 0.00083271 | 0.0027449 |
| STM14_RS16315 | | transporter | WP_000819369.1 | 0.502483714 | 0.0953076 | 0.16208895 |
| STM14_RS01865 | yafC | DNA-binding transcriptional regulator YafC | WP_000648534.1 | 0.502230511 | 0.02047386 | 0.04354983 |
| STM14_RS07200 | | chorismate mutase | WP_000617985.1 | 0.501893436 | 0.00029751 | 0.00109873 |
| STM14_RS23050 | rplI | 50S ribosomal protein L9 | WP_001196059.1 | 0.501592159 | 0.00975203 | 0.02337695 |
| STM14_RS08640 | | hypothetical protein | WP_000987828.1 | 0.501427822 | 0.00144509 | 0.00444885 |
| STM14_RS24195 | yncL | stress response membrane protein YncL | WP_001103198.1 | 0.500930837 | 0.00091751 | 0.00298696 |
| STM14_RS19130 | hmsP | biofilm formation regulator HmsP | WP_001266339.1 | 0.50040755 | 0.0017657 | 0.00533331 |
| STM14_RS09700 | narJ | nitrate reductase molybdenum cofactor assembly chaperone | WP_000571667.1 | 0.500062534 | 0.10481423 | 0.17550069 |
| STM14_RS05995 | hpaG | 4-hydroxyphenylacetate degradation bifunctional isomerase/decarboxy | WP_000679216.1 | 0.500049273 | 0.00438458 | 0.01165635 |
| STM14_RS03880 | | tetratricopeptide repeat protein | WP_001016626.1 | 0.49941477 | 0.00231434 | 0.00673629 |
| STM14_RS22915 | nsrR | nitric oxide-sensing transcriptional repressor NsrR | WP_001177632.1 | 0.499062765 | 6.5727E-05 | 0.00028914 |
| STM14_RS22225 | malM | maltose operon protein MalM | WP_000782504.1 | 0.49731287 | 0.04097174 | 0.07844927 |
| STM14_RS02055 | | polymorphic toxin type 47 domain-containing protein | | 0.496372829 | 0.01673574 | 0.03669849 |
| STM14_RS02845 | | lipoprotein | WP_001537832.1 | 0.49599985 | 0.00310287 | 0.00869465 |
| STM14_RS15575 | pphB | protein-serine/threonine phosphatase | WP_000420452.1 | 0.495745125 | 0.07860982 | 0.1379606 |
| STM14_RS02475 | hemB | porphobilinogen synthase | WP_000130724.1 | 0.494610139 | 0.00011519 | 0.00047388 |
| STM14_RS21430 | lsrR | transcriptional regulator LsrR | WP_001283048.1 | 0.494413202 | 5.5341E-06 | 3.1127E-05 |
| STM14_RS15760 | | CRISPR-associated helicase/endonuclease Cas3 | WP_000029738.1 | 0.494062203 | 0.00039264 | 0.00141183 |
| STM14_RS13140 | prmB | 50S ribosomal protein L3 N(5)-glutamine methyltransferase | WP_001542329.1 | 0.493612626 | 0.00373515 | 0.01017859 |
| STM14_RS15000 | lysM | peptidoglycan-binding protein LysM | WP_000522406.1 | 0.492775026 | 1.7773E-09 | 1.8279E-08 |
| STM14_RS00195 | spvB | SPI-2 type III secretion system effector NAD(+)--protein-arginine ADP-ri | WP_001676648.1 | 0.492530753 | 0.00704348 | 0.01767833 |
| STM14_RS05780 | pyrD | quinone-dependent dihydroorotate dehydrogenase | WP_000291723.1 | 0.492497975 | 0.00205694 | 0.00609305 |
| STM14_RS03620 | | pyridoxal phosphate-dependent aminotransferase | WP_001245592.1 | 0.492264523 | 0.01347942 | 0.03067443 |
| STM14_RS12000 | btsR | two-component system response regulator BtsR | WP_000598637.1 | 0.491856453 | 0.01163962 | 0.02710757 |
| STM14_RS21875 | thiC | phosphomethylpyrimidine synthase ThiC | WP_000108411.1 | 0.490543569 | 0.04780697 | 0.08967156 |
| STM14_RS05765 | gtgE | virulence effector protease GtgE | WP_001525490.1 | 0.489904796 | 0.00565675 | 0.01460599 |
| STM14_RS08150 | rsxA | electron transport complex subunit RsxA | WP_000133179.1 | 0.488814867 | 0.08532991 | 0.14764319 |
| STM14_RS11315 | | energy-coupling factor ABC transporter substrate-binding protein | WP_000753212.1 | 0.488621702 | 0.1495179 | 0.23506614 |
| STM14_RS22665 | | non-specific acid phosphatase | WP_000842434.1 | 0.487976235 | 2.09E-05 | 0.0001047 |
| STM14_RS15950 | gcvA | glycine cleavage system transcriptional regulator GcvA | WP_000044409.1 | 0.487906733 | 0.00026993 | 0.00100596 |
| STM14_RS09280 | | peptide ABC transporter substrate-binding protein | WP_001216554.1 | 0.487746585 | 4.3211E-07 | 3.0133E-06 |
| STM14_RS21745 | murB | UDP-N-acetylmuramate dehydrogenase | WP_000149795.1 | 0.487632641 | 0.00101326 | 0.00324702 |
| STM14_RS08670 | treZ | malto-oligosyltrehalose trehalohydrolase | WP_000095534.1 | 0.487591812 | 0.00022605 | 0.00086338 |
| STM14_RS11035 | | hypothetical protein | WP_001292890.1 | 0.484478699 | 0.01758264 | 0.03826198 |
| STM14_RS04810 | mntR | manganese-binding transcriptional regulator MntR | WP_000533542.1 | 0.484106149 | 0.00034536 | 0.00125653 |
| STM14_RS10825 | dsrB | protein DsrB | WP_000867218.1 | 0.484080178 | 9.4049E-05 | 0.00039441 |
| STM14_RS12685 | nudI | nucleoside triphosphatase NudI | WP_001249900.1 | 0.483306898 | 0.00084743 | 0.00278096 |
| STM14_RS02555 | aroL | shikimate kinase AroL | WP_000983569.1 | 0.482748653 | 0.01234059 | 0.0284747 |
| STM14_RS16730 | | carbon-nitrogen hydrolase family protein | | 0.481417114 | 0.00031301 | 0.00115134 |
| STM14_RS03010 | | Rpn family recombination-promoting nuclease/putative transposase | WP_000440523.1 | 0.481380187 | 0.00033074 | 0.00120931 |
| STM14_RS05170 | clpS | ATP-dependent Clp protease adapter ClpS | WP_000520789.1 | 0.480219839 | 2.5199E-07 | 1.8446E-06 |
| STM14_RS05850 | | TfoX/Sxy family DNA transformation protein | WP_001202375.1 | 0.479721842 | 0.03980771 | 0.0765389 |
| STM14_RS21035 | | serine/threonine protein kinase | WP_000999249.1 | 0.47873581 | 2.6822E-05 | 0.00013029 |
| STM14_RS00275 | | ParB family protein | WP_000064274.1 | 0.478240149 | 7.2984E-07 | 4.8859E-06 |
| STM14_RS05845 | sulA | cell division inhibitor SulA | WP_000288732.1 | 0.478023568 | 0.00146972 | 0.00451327 |
| STM14_RS06140 | | N-acetylneuraminate epimerase | WP_000525751.1 | 0.477962278 | 0.01979346 | 0.0422735 |
| STM14_RS04750 | ybiJ | DUF1471 family protein YbiJ | WP_000849089.1 | 0.477829796 | 0.11455727 | 0.18880925 |
| STM14_RS04940 | | GntP family permease | WP_000287113.1 | 0.476476033 | 0.00856726 | 0.0209599 |
| STM14_RS19645 | waaZ | 3-deoxy-D-manno-oct-2-ulosonate III transferase WaaZ | WP_000536012.1 | 0.476462641 | 0.00164595 | 0.00499207 |
| STM14_RS22640 | rtsB | transcriptional regulator RtsB | WP_001269916.1 | 0.474899566 | 0.00568649 | 0.01465609 |
| STM14_RS19360 | | YiaA/YiaB family protein | WP_000340935.1 | 0.474653319 | 0.13430751 | 0.21472872 |
| STM14_RS23380 | pyrB | aspartate carbamoyltransferase | WP_000013055.1 | 0.474366847 | 0.04343453 | 0.08260638 |
| STM14_RS06370 | flgB | flagellar basal body rod protein FlgB | WP_000887043.1 | 0.473617504 | 0.00743808 | 0.01855488 |
| STM14_RS12115 | | CidB/LrgB family autolysis modulator | WP_000989296.1 | 0.472953555 | 0.07022932 | 0.12516184 |
| STM14_RS06005 | hpaD | 3%2C4-dihydroxyphenylacetate 2%2C3-dioxygenase | WP_000516978.1 | 0.472552088 | 0.00671241 | 0.01693974 |
| STM14_RS17160 | | PAS domain-containing methyl-accepting chemotaxis protein | WP_000094651.1 | 0.470787332 | 2.4799E-07 | 1.8226E-06 |
| STM14_RS06750 | | ParA family protein | WP_000368620.1 | 0.470575186 | 5.4807E-07 | 3.7232E-06 |
| STM14_RS13015 | | alanine racemase | WP_000132305.1 | 0.470312429 | 0.01903109 | 0.04093019 |
| STM14_RS21550 | | TIGR02117 family protein | WP_000792788.1 | 0.470005668 | 0.12039657 | 0.19615052 |
| STM14_RS14275 | | phage terminase large subunit family protein | WP_000623091.1 | 0.469959148 | 0.09235463 | 0.15786896 |
| STM14_RS08505 | ompC | porin OmpC | WP_000758335.1 | 0.469632304 | 0.11972146 | 0.19522339 |
| STM14_RS09300 | tnpA | IS200/IS605-like element IS200F family transposase | WP_000502119.1 | 0.469108741 | 0.00194029 | 0.00579413 |

| | | | | | | |
|---|---|---|---|---|---|---|
| STM14_RS24075 | | hypothetical protein | WP_000749515.1 | 0.468929903 | 0.09213712 | 0.15764353 |
| STM14_RS06355 | flgN | flagella biosynthesis chaperone FlgN | WP_000197547.1 | 0.467993136 | 6.1591E-06 | 3.4257E-05 |
| STM14_RS10660 | sdiA | transcriptional regulator SdiA | WP_001157166.1 | 0.46733631 | 0.00012573 | 0.00051382 |
| STM14_RS19335 | | GNAT family N-acetyltransferase | WP_000702452.1 | 0.467181742 | 0.04963329 | 0.09248471 |
| STM14_RS01615 | btuF | vitamin B12 ABC transporter substrate-binding protein BtuF | WP_001118843.1 | 0.465341487 | 0.13310987 | 0.21318422 |
| STM14_RS17875 | | hypothetical protein | WP_001103887.1 | 0.465031513 | 4.5282E-06 | 2.6011E-05 |
| STM14_RS11625 | | mannose-1-phosphate guanylyltransferase/mannose-6-phosphate isom | WP_000009017.1 | 0.464765017 | 8.8102E-05 | 0.00037467 |
| STM14_RS21100 | | Cof-type HAD-IIB family hydrolase | WP_001270592.1 | 0.464751487 | 0.14734912 | 0.23225149 |
| STM14_RS23520 | | AAA domain-containing protein | WP_000342544.1 | 0.464728464 | 2.1706E-05 | 0.00010815 |
| STM14_RS01165 | ilvN | acetolactate synthase small subunit | WP_001563086.1 | 0.463312599 | 0.00133603 | 0.0041583 |
| STM14_RS21210 | | alpha/beta hydrolase | WP_001127703.1 | 0.462820655 | 0.07751689 | 0.13643269 |
| STM14_RS10910 | | tail fiber assembly protein | WP_015701331.1 | 0.462713496 | 0.02030453 | 0.0432396 |
| STM14_RS18975 | nikR | nickel-responsive transcriptional regulator NikR | WP_001190057.1 | 0.461621509 | 0.05262007 | 0.09760559 |
| STM14_RS07525 | thrS | threonine--tRNA ligase | WP_001144223.1 | 0.461115862 | 1.0141E-07 | 8.0469E-07 |
| STM14_RS20925 | tatA | Sec-independent protein translocase subunit TatA | WP_000508972.1 | 0.460335518 | 7.1828E-06 | 3.9415E-05 |
| STM14_RS18485 | dam | adenine-specific DNA-methyltransferase | WP_000742132.1 | 0.459354354 | 0.00146413 | 0.00449995 |
| STM14_RS07310 | gapA | glyceraldehyde-3-phosphate dehydrogenase | WP_000153505.1 | 0.459328946 | 1.2612E-07 | 9.8591E-07 |
| STM14_RS08865 | ycdR | GntR family transcriptional regulator YdcR | WP_000760692.1 | 0.458801853 | 0.00097862 | 0.00315522 |
| STM14_RS06455 | yceD | 23S rRNA accumulation protein YceD | WP_001174492.1 | 0.457261748 | 4.1131E-06 | 2.3849E-05 |
| STM14_RS06440 | | DUF3053 domain-containing protein | WP_010989014.1 | 0.45643047 | 0.00491602 | 0.01284655 |
| STM14_RS18225 | rpmC | 50S ribosomal protein L29 | WP_000644742.1 | 0.455954184 | 4.3675E-06 | 2.5205E-05 |
| STM14_RS07600 | ppsR | posphoenolpyruvate synthetase regulatory kinase/phosphorylase PpsR | WP_000370992.1 | 0.453865794 | 0.03919767 | 0.07556325 |
| STM14_RS09470 | cysB | HTH-type transcriptional regulator CysB | WP_000776233.1 | 0.453293057 | 2.4362E-05 | 0.00011976 |
| STM14_RS00040 | | AraC family transcriptional regulator | WP_000417898.1 | 0.452681592 | 0.01494315 | 0.03350833 |
| STM14_RS05245 | lrp | leucine-responsive transcriptional regulator Lrp | WP_000228469.1 | 0.451195907 | 4.772E-07 | 3.2781E-06 |
| STM14_RS06835 | | ORF6N domain-containing protein | WP_000211401.1 | 0.448093323 | 0.16193501 | 0.25010264 |
| STM14_RS10885 | | cold-shock protein | WP_000208509.1 | 0.446993121 | 0.00248675 | 0.00717036 |
| STM14_RS18700 | | PstS family phosphate ABC transporter substrate-binding protein | WP_010989078.1 | 0.445626401 | 0.0083404 | 0.02045927 |
| STM14_RS20960 | | aryl-sulfate sulfotransferase | WP_000669905.1 | 0.445485582 | 0.01335046 | 0.03045624 |
| STM14_RS14225 | | phage minor tail U family protein | WP_000033885.1 | 0.445291238 | 0.11689239 | 0.19170229 |
| STM14_RS11060 | | hypothetical protein | WP_000900722.1 | 0.445152468 | 0.05291032 | 0.09804519 |
| STM14_RS23605 | | DUF302 domain-containing protein | WP_001654330.1 | 0.444240094 | 0.00268297 | 0.0076698 |
| STM14_RS15765 | sopD | SPI-1 type III secretion system effector SopD | WP_001145543.1 | 0.443852612 | 0.00048071 | 0.00169213 |
| STM14_RS21630 | | PTS fructose-like transporter subunit IIB | WP_000161258.1 | 0.443772063 | 0.00097776 | 0.00315522 |
| STM14_RS19355 | | acyltransferase | WP_001182588.1 | 0.442718595 | 0.14227648 | 0.22502663 |
| STM14_RS00775 | | glycoside hydrolase family 31 protein | WP_001116457.1 | 0.44183868 | 0.10728402 | 0.17857802 |
| STM14_RS26515 | pmrR | LpxT activity modulator PmrR | WP_000844399.1 | 0.44075673 | 0.04370351 | 0.08307502 |
| STM14_RS07205 | leuE | leucine efflux protein LeuE | WP_000457190.1 | 0.440299712 | 0.00134342 | 0.00417777 |
| STM14_RS15915 | fucP | L-fucose:H+ symporter permease | WP_000528627.1 | 0.439314291 | 0.1054202 | 0.17619481 |
| STM14_RS06445 | | hypothetical protein | WP_000528430.1 | 0.438871832 | 0.01374769 | 0.03118854 |
| STM14_RS02810 | | tetratricopeptide repeat protein | WP_000595924.1 | 0.438537121 | 0.02065847 | 0.0438918 |
| STM14_RS13895 | | DUF5066 family protein | WP_000355342.1 | 0.438425137 | 0.11170047 | 0.18467944 |
| STM14_RS04440 | | LysR family transcriptional regulator | WP_000405925.1 | 0.438086854 | 0.02353337 | 0.04887189 |
| STM14_RS07695 | | L-cystine transporter | WP_000261866.1 | 0.437981933 | 0.01074091 | 0.02538383 |
| STM14_RS10475 | flhA | flagellar biosynthesis protein FlhA | WP_000002386.1 | 0.437728717 | 0.02451436 | 0.05062361 |
| STM14_RS17775 | | DUF695 domain-containing protein | WP_001739021.1 | 0.437249445 | 0.00029522 | 0.00109246 |
| STM14_RS08155 | | DUF2569 domain-containing protein | WP_000214061.1 | 0.436809498 | 0.01362692 | 0.03096584 |
| STM14_RS22740 | | YjeJ family protein | WP_001229227.1 | 0.436452339 | 0.10005633 | 0.16876198 |
| STM14_RS06380 | flgD | flagellar hook assembly protein FlgD | WP_000020450.1 | 0.436412905 | 7.1926E-05 | 0.00031294 |
| STM14_RS11130 | | hypothetical protein | WP_000997190.1 | 0.436301189 | 5.3209E-06 | 3.0003E-05 |
| STM14_RS16395 | | small-conductance mechanosensitive channel MscS | WP_000389833.1 | 0.435619081 | 0.00132304 | 0.00412482 |
| STM14_RS12280 | | cyclic di-GMP phosphodiesterase | WP_000957746.1 | 0.435512074 | 0.02206833 | 0.04632587 |
| STM14_RS15035 | | DUF2002 family protein | WP_000281305.1 | 0.435482432 | 0.09655309 | 0.16360262 |
| STM14_RS12590 | | bifunctional 3-demethylubiquinone 3-O-methyltransferase/2-octapreny | WP_001091009.1 | 0.434163633 | 0.00068607 | 0.00230486 |
| STM14_RS16245 | idi | isopentenyl-diphosphate Delta-isomerase | WP_000133994.1 | 0.433745925 | 0.00601717 | 0.01541768 |
| STM14_RS18145 | rpsD | 30S ribosomal protein S4 | WP_000135226.1 | 0.431234767 | 1.8888E-06 | 1.1572E-05 |
| STM14_RS06870 | | PPC domain-containing protein | WP_000669689.1 | 0.430879033 | 0.00210128 | 0.00619657 |
| STM14_RS15485 | spaQ | SPI-1 type III secretion system export apparatus protein SpaQ | WP_000342503.1 | 0.43033459 | 0.11448876 | 0.18878084 |
| STM14_RS07565 | | glutathione peroxidase | WP_001181565.1 | 0.429430215 | 0.03767156 | 0.07292675 |
| STM14_RS17280 | pflB | formate C-acetyltransferase | WP_000867323.1 | 0.428465198 | 0.00303723 | 0.00853019 |
| STM14_RS00900 | carA | glutamine-hydrolyzing carbamoyl-phosphate synthase small subunit | WP_000597287.1 | 0.42736107 | 0.11132956 | 0.18431449 |
| STM14_RS12960 | yfcG | GSH-dependent disulfide bond oxidoreductase | WP_000566563.1 | 0.426963673 | 0.07720227 | 0.13607408 |
| STM14_RS19845 | | virulence RhuM family protein | WP_000774139.1 | 0.426546844 | 0.01261699 | 0.029053 |
| STM14_RS13110 | mnmC | bifunctional tRNA (5-methylaminomethyl-2-thiouridine)(34)-methyltran | WP_000816091.1 | 0.425954085 | 0.00106126 | 0.00336568 |
| STM14_RS05560 | | hypothetical protein | WP_000807548.1 | 0.425637134 | 0.01702557 | 0.03718159 |
| STM14_RS18405 | | aspartate aminotransferase family protein | WP_000190029.1 | 0.425436144 | 0.03807095 | 0.07366119 |
| STM14_RS19065 | | GntR family transcriptional regulator | WP_000572568.1 | 0.425200473 | 0.01615612 | 0.03570639 |
| STM14_RS12275 | mepS | bifunctional murein DD-endopeptidase/murein LD-carboxypeptidase | WP_000241015.1 | 0.424227629 | 5.2251E-06 | 2.9598E-05 |
| STM14_RS07625 | | electron transfer flavoprotein subunit alpha | WP_001015871.1 | 0.4227595 | 0.11917725 | 0.19468553 |
| STM14_RS17070 | | flotillin family protein | WP_000342878.1 | 0.421877562 | 0.03070669 | 0.06140199 |
| STM14_RS12335 | | hypothetical protein | WP_000033440.1 | 0.419809877 | 0.0010378 | 0.0033099 |
| STM14_RS17660 | hpf | ribosome hibernation promoting factor | WP_001176589.1 | 0.41967267 | 2.1258E-06 | 1.2895E-05 |
| STM14_RS12120 | cdd | cytidine deaminase | WP_000553526.1 | 0.419465777 | 0.07780127 | 0.13679006 |
| STM14_RS13190 | pgtE | omptin family outer membrane protease PgtE | WP_000716007.1 | 0.419015671 | 0.00017 | 0.00066802 |
| STM14_RS21200 | | type II toxin-antitoxin system HigA family antitoxin | WP_001259011.1 | 0.418647947 | 0.00849933 | 0.02080755 |
| STM14_RS03390 | | DUF6056 family protein | WP_010988978.1 | 0.417641604 | 0.19274695 | 0.28697706 |
| STM14_RS22580 | fumA | class I fumarate hydratase FumA | WP_000037194.1 | 0.417570701 | 0.01389006 | 0.03147273 |
| STM14_RS06375 | flgC | flagellar basal body rod protein FlgC | WP_001196448.1 | 0.417410193 | 0.03822213 | 0.07391485 |
| STM14_RS05320 | focA | formate transporter FocA | WP_000642539.1 | 0.417341774 | 0.00010838 | 0.0004494 |
| STM14_RS23055 | | DMT family transporter | WP_000381075.1 | 0.41673363 | 0.09927697 | 0.16760101 |
| STM14_RS20655 | wecA | UDP-N-acetylglucosamine--undecaprenyl-phosphate N-acetylglucosar | WP_000771937.1 | 0.415840191 | 0.0004589 | 0.00162156 |
| STM14_RS23975 | | YtjB family periplasmic protein | WP_000090067.1 | 0.415235213 | 0.12878975 | 0.20698553 |

| | | | | | | |
|---|---|---|---|---|---|---|
| STM14_RS17090 | | TIGR04211 family SH3 domain-containing protein | WP_001125344.1 | 0.414957286 | 0.00019782 | 0.00076591 |
| STM14_RS09545 | | ferritin-like domain-containing protein | WP_001022822.1 | 0.413253981 | 0.00044297 | 0.00157284 |
| STM14_RS23445 | | YjgN family protein | WP_001066984.1 | 0.41257743 | 0.0639004 | 0.11544715 |
| STM14_RS22410 | | Na+/H+ antiporter | WP_001536604.1 | 0.412281223 | 0.00302713 | 0.00850831 |
| STM14_RS21615 | fsa | fructose-6-phosphate aldolase | WP_000424866.1 | 0.412009428 | 0.00217813 | 0.00638671 |
| STM14_RS09410 | pdeR | cyclic di-GMP phosphodiesterase | WP_001623919.1 | 0.411965279 | 0.01360981 | 0.03095203 |
| STM14_RS09090 | | YdbL family protein | WP_000760345.1 | 0.411635189 | 0.03301817 | 0.0649775 |
| STM14_RS07850 | sscA | SycD/LcrH family type III secretion system chaperone SscA | WP_000711029.1 | 0.411543457 | 0.00143313 | 0.0044157 |
| STM14_RS14305 | gogA | type III secretion system effector protease GogA | WP_001110783.1 | 0.410460372 | 0.03012597 | 0.06044895 |
| STM14_RS16720 | | Hcp family type VI secretion system effector | WP_000338754.1 | 0.410400431 | 0.0032783 | 0.00913065 |
| STM14_RS17575 | rplU | 50S ribosomal protein L21 | WP_000271396.1 | 0.410346896 | 1.2114E-06 | 7.6772E-06 |
| STM14_RS09375 | | EAL domain-containing protein | WP_001268403.1 | 0.410199707 | 0.00800245 | 0.01977519 |
| STM14_RS23795 | opgB | phosphatidylglycerol--membrane-oligosaccharide glycerophosphotrans | WP_001292726.1 | 0.410146068 | 5.1161E-05 | 0.00023371 |
| STM14_RS13345 | | outer membrane protein | WP_000722926.1 | 0.409706943 | 0.20523905 | 0.30227607 |
| STM14_RS10185 | | GNAT family N-acetyltransferase | WP_000354408.1 | 0.4087311 | 0.00245577 | 0.00711516 |
| STM14_RS23535 | pglZ | BREX-1 system phosphatase PglZ type A | WP_001738657.1 | 0.40835063 | 0.00070559 | 0.00236181 |
| STM14_RS26435 | | IS3 family transposase | | 0.408048204 | 0.12075688 | 0.19665052 |
| STM14_RS16685 | | LuxR family transcriptional regulator | WP_001275362.1 | 0.407928931 | 0.00368941 | 0.01007744 |
| STM14_RS01825 | gmhB | D-glycero-beta-D-manno-heptose 1%2C7-bisphosphate 7-phosphatase | WP_001051726.1 | 0.406260706 | 0.00222366 | 0.0064826 |
| STM14_RS00820 | | nitrite reductase | WP_000516322.1 | 0.406015538 | 0.0319893 | 0.06335066 |
| STM14_RS16895 | dkgA | 2%2C5-didehydrogluconate reductase DkgA | WP_000013100.1 | 0.406000508 | 4.2894E-05 | 0.00020043 |
| STM14_RS00735 | | bifunctional UDP-sugar hydrolase/5'-nucleotidase | WP_000262866.1 | 0.405015292 | 0.02463552 | 0.0508378 |
| STM14_RS02095 | | Saf-pilin pilus formation protein SafA | WP_010988970.1 | 0.403039985 | 0.22241989 | 0.32242128 |
| STM14_RS08735 | ompD | porin OmpD | WP_000769035.1 | 0.403024579 | 3.3629E-06 | 1.9844E-05 |
| STM14_RS21030 | | YihD family protein | WP_000649858.1 | 0.401319541 | 3.6655E-05 | 0.0001737 |
| STM14_RS13295 | | putative DNA-binding transcriptional regulator | WP_000826514.1 | 0.400840137 | 0.00309147 | 0.0086693 |
| STM14_RS15720 | | aminopeptidase | WP_000490481.1 | 0.400218378 | 0.00688191 | 0.01732002 |
| STM14_RS10495 | cheB | protein-glutamate methylesterase/protein glutamine deamidase | WP_000036392.1 | 0.399369061 | 5.8246E-05 | 0.00026154 |
| STM14_RS11615 | | undecaprenyl-phosphate galactose phosphotransferase | WP_000368707.1 | 0.399321085 | 0.0001642 | 0.0006487 |
| STM14_RS23255 | iolH | sugar phosphate isomerase/epimerase IolH | WP_000683955.1 | 0.39929221 | 0.11652191 | 0.19127672 |
| STM14_RS06635 | potA | spermidine/putrescine ABC transporter ATP-binding protein PotA | WP_000531607.1 | 0.398468133 | 0.00272056 | 0.00775917 |
| STM14_RS19605 | | glycosyltransferase | WP_000798206.1 | 0.39768828 | 0.26476843 | 0.37053491 |
| STM14_RS04415 | zitB | CDF family zinc transporter ZitB | WP_000951251.1 | 0.396801315 | 0.03830886 | 0.07400484 |
| STM14_RS21840 | | type III secretion system effector arginine glycosyltransferase SseK1 | WP_000400600.1 | 0.396265548 | 0.10704996 | 0.17827135 |
| STM14_RS01515 | dksA | RNA polymerase-binding protein DksA | WP_001155232.1 | 0.395053262 | 1.6695E-06 | 1.0279E-05 |
| STM14_RS02605 | phoB | phosphate response regulator transcription factor PhoB | WP_000113921.1 | 0.394751212 | 0.00516714 | 0.01346455 |
| STM14_RS00595 | | sodium:alanine symporter family protein | WP_001113820.1 | 0.394059304 | 0.03976371 | 0.07649425 |
| STM14_RS08265 | mdtJ | multidrug/spermidine efflux SMR transporter subunit MdtJ | WP_000500278.1 | 0.392988684 | 0.00314986 | 0.0088196 |
| STM14_RS01880 | | class I SAM-dependent methyltransferase | WP_000207224.1 | 0.392747516 | 0.05554431 | 0.10225708 |
| STM14_RS18815 | | DUF2756 family protein | WP_000826041.1 | 0.392187806 | 3.0378E-06 | 1.8069E-05 |
| STM14_RS11685 | rfbD | dTDP-4-dehydrorhamnose reductase | WP_001023662.1 | 0.391901471 | 0.00207397 | 0.00613855 |
| STM14_RS16905 | | HIT family protein | WP_000128242.1 | 0.391356639 | 0.00935735 | 0.02253353 |
| STM14_RS12110 | | CidA/LrgA family protein | WP_000045719.1 | 0.391209003 | 0.13321043 | 0.21325253 |
| STM14_RS06930 | | phage minor tail U family protein | WP_000033885.1 | 0.390625535 | 0.19636471 | 0.29153825 |
| STM14_RS22355 | | MotA/TolQ/ExbB proton channel family protein | WP_000875195.1 | 0.390576149 | 0.17541766 | 0.26612601 |
| STM14_RS14400 | | hypothetical protein | WP_000950426.1 | 0.390203745 | 0.00346552 | 0.00957242 |
| STM14_RS15890 | sdaC | HAAAP family serine/threonine permease SdaC | WP_000450462.1 | 0.389693265 | 1.9609E-05 | 9.8634E-05 |
| STM14_RS20360 | pstC | phosphate ABC transporter permease PstC | WP_000741630.1 | 0.389622949 | 0.13573177 | 0.21653569 |
| STM14_RS10515 | cheA | chemotaxis protein CheA | WP_000061302.1 | 0.388617602 | 4.7506E-05 | 0.00021919 |
| STM14_RS16360 | | 5-formyltetrahydrofolate cyclo-ligase | WP_001520956.1 | 0.387325701 | 0.00820763 | 0.02020086 |
| STM14_RS22170 | pgi | glucose-6-phosphate isomerase | WP_000790037.1 | 0.386883806 | 7.671E-05 | 0.00033112 |
| STM14_RS19985 | | PTS sugar transporter subunit IIA | WP_001152234.1 | 0.386501097 | 0.01672738 | 0.03669849 |
| STM14_RS01775 | | YaeP family protein | WP_001518678.1 | 0.386105271 | 0.00024192 | 0.00091832 |
| STM14_RS09060 | | class I SAM-dependent methyltransferase | WP_001520112.1 | 0.385698686 | 0.00026913 | 0.001004 |
| STM14_RS20170 | | cytochrome-c peroxidase | WP_000724476.1 | 0.385261032 | 0.0092099 | 0.02222205 |
| STM14_RS09510 | rnm | RNase AM | WP_000500442.1 | 0.385073015 | 0.00244325 | 0.00709467 |
| STM14_RS04935 | | LysR family transcriptional regulator | WP_000545327.1 | 0.384859385 | 0.00521119 | 0.01355057 |
| STM14_RS10510 | cheW | chemotaxis protein CheW | WP_000147295.1 | 0.384563995 | 0.00026785 | 0.00100084 |
| STM14_RS09005 | | methyl-accepting chemotaxis protein | WP_000528484.1 | 0.383305264 | 0.00064183 | 0.00217406 |
| STM14_RS09600 | | HI1450 family dsDNA-mimic protein | WP_000425069.1 | 0.382134766 | 0.00215917 | 0.00634321 |
| STM14_RS06760 | | YdaS family helix-turn-helix protein | WP_000145711.1 | 0.381807428 | 0.23301736 | 0.33499417 |
| STM14_RS23755 | tsr | methyl-accepting chemotaxis protein | WP_000919519.1 | 0.381085487 | 0.00018224 | 0.00071157 |
| STM14_RS15355 | sitC | iron/manganese ABC transporter permease subunit SitC | WP_001104336.1 | 0.381033712 | 0.0958271 | 0.16282204 |
| STM14_RS16085 | | YgdI/YgdR family lipoprotein | WP_000758662.1 | 0.380676301 | 0.00020858 | 0.00080332 |
| STM14_RS13270 | nupC | nucleoside permease NupC | WP_000376347.1 | 0.380419713 | 0.0078144 | 0.01933645 |
| STM14_RS09645 | tdk | thymidine kinase | WP_000068097.1 | 0.380105692 | 0.00560573 | 0.0144946 |
| STM14_RS14650 | grpE | nucleotide exchange factor GrpE | WP_001518875.1 | 0.379789724 | 4.7774E-05 | 0.00022016 |
| STM14_RS21520 | hslU | HslU--HslV peptidase ATPase subunit | WP_001293360.1 | 0.379743484 | 0.00052655 | 0.00183533 |
| STM14_RS19515 | | YibL family ribosome-associated protein | WP_000665687.1 | 0.378476978 | 0.00648179 | 0.01643662 |
| STM14_RS17540 | yhbY | ribosome assembly RNA-binding protein YhbY | WP_001592995.1 | 0.378323933 | 0.01056285 | 0.0250112 |
| STM14_RS06935 | | phage tail tube protein | WP_000971954.1 | 0.378016216 | 0.04002668 | 0.07691974 |
| STM14_RS10235 | | DUF1441 family protein | | 0.377744763 | 0.28387358 | 0.39132254 |
| STM14_RS23750 | btsT | pyruvate/proton symporter BtsT | WP_000379927.1 | 0.377688274 | 0.20217132 | 0.29895373 |
| STM14_RS21370 | cpxP | cell-envelope stress modulator CpxP | WP_001233463.1 | 0.377191832 | 0.00050578 | 0.00177025 |
| STM14_RS13115 | | YfcL family protein | WP_000559749.1 | 0.376389119 | 0.04641466 | 0.08750577 |
| STM14_RS02480 | | autotransporter outer membrane beta-barrel domain-containing protein | WP_000556518.1 | 0.375766556 | 0.00326907 | 0.00911182 |
| STM14_RS12445 | narP | nitrate/nitrite response regulator protein NarP | WP_001115498.1 | 0.375632167 | 0.0041444 | 0.01116802 |
| STM14_RS00235 | | carbonic anhydrase family protein | WP_001541566.1 | 0.37550262 | 9.8261E-06 | 5.1908E-05 |
| STM14_RS15805 | pyrG | CTP synthase (glutamine hydrolyzing) | WP_000210863.1 | 0.374800573 | 0.00019234 | 0.00074702 |
| STM14_RS07495 | | YdiY family protein | WP_000770981.1 | 0.374674574 | 0.06546433 | 0.1178678 |
| STM14_RS07920 | ssaM | SPI-2 type III secretion system apparatus protein SsaM | WP_000383702.1 | 0.374038676 | 0.14868066 | 0.23384973 |
| STM14_RS03510 | | RamA family antibiotic efflux transcriptional regulator | WP_000155631.1 | 0.373853873 | 0.04747234 | 0.0892257 |

| | | | | | | |
|---|---|---|---|---|---|---|
| STM14_RS20395 | | Bax inhibitor-1 family protein | WP_000009904.1 | 0.373748827 | 0.31203896 | 0.42193443 |
| STM14_RS10005 | | RidA family protein | WP_000029550.1 | 0.373341756 | 0.00705015 | 0.01768302 |
| STM14_RS10135 | yebS | membrane integrity lipid transport subunit YebS | WP_001207294.1 | 0.372154404 | 0.01598606 | 0.03539428 |
| STM14_RS06660 | phoQ | two-component system sensor histidine kinase PhoQ | WP_001031687.1 | 0.371973798 | 0.00026801 | 0.00100084 |
| STM14_RS08260 | | AI-2E family transporter | WP_001118268.1 | 0.371238259 | 0.04645232 | 0.08753196 |
| STM14_RS15875 | syd | SecY-interacting protein | WP_000343990.1 | 0.370921893 | 0.02599299 | 0.05337768 |
| STM14_RS11965 | yehD | fimbrial protein YehD | WP_000830686.1 | 0.370818292 | 0.01104169 | 0.02592825 |
| STM14_RS10800 | fliO | flagellar biosynthetic protein FliO | WP_000978276.1 | 0.370259846 | 0.15606148 | 0.24296761 |
| STM14_RS23560 | | DUF1788 domain-containing protein | WP_000566905.1 | 0.370235823 | 0.04145987 | 0.07921912 |
| STM14_RS15815 | | type II toxin-antitoxin system RelE/ParE family toxin | WP_001532579.1 | 0.370040172 | 0.2142321 | 0.31225532 |
| STM14_RS12345 | | YejL family protein | WP_001135904.1 | 0.369961693 | 0.00964448 | 0.02313418 |
| STM14_RS16150 | | LysR family transcriptional regulator | WP_001270766.1 | 0.369191476 | 0.0374814 | 0.07267325 |
| STM14_RS05515 | | helix-turn-helix domain-containing protein | WP_001038966.1 | 0.368785557 | 0.00605635 | 0.01550728 |
| STM14_RS06250 | mdoG | glucans biosynthesis protein MdoG | WP_001738192.1 | 0.368471383 | 0.00023042 | 0.00087735 |
| STM14_RS13985 | | DUF1007 family protein | WP_000805728.1 | 0.368343676 | 0.08432284 | 0.14624432 |
| STM14_RS12785 | | VWA domain-containing protein | WP_000931746.1 | 0.368295276 | 0.16763367 | 0.25664331 |
| STM14_RS01715 | lpxD | UDP-3-O-(3-hydroxymyristoyl)glucosamine N-acyltransferase | WP_001139265.1 | 0.367772269 | 0.00250335 | 0.00720669 |
| STM14_RS19015 | dtpB | dipeptide/tripeptide permease DtpB | WP_001098275.1 | 0.367215058 | 0.00218661 | 0.00640501 |
| STM14_RS18350 | kefB | glutathione-regulated potassium-efflux system protein KefB | WP_000398133.1 | 0.366935683 | 0.00587338 | 0.01509127 |
| STM14_RS15510 | invC | SctN family type III secretion system ATPase InvC | WP_000856766.1 | 0.366211534 | 0.0146499 | 0.03291087 |
| STM14_RS05495 | | recombinase RecT | WP_001539618.1 | 0.365541479 | 0.30390235 | 0.41320844 |
| STM14_RS26150 | | hypothetical protein | | 0.36485815 | 0.18992188 | 0.28391895 |
| STM14_RS20935 | tatC | Sec-independent protein translocase subunit TatC | WP_000257554.1 | 0.364529892 | 0.00682775 | 0.01719548 |
| STM14_RS19895 | | Rpn family recombination-promoting nuclease/putative transposase | WP_000749540.1 | 0.363960061 | 0.01325535 | 0.03025803 |
| STM14_RS03190 | allA | ureidoglycolate lyase | WP_000764664.1 | 0.363364756 | 0.08634302 | 0.14916107 |
| STM14_RS17190 | | Gfo/Idh/MocA family oxidoreductase | WP_000617687.1 | 0.363146599 | 0.01234264 | 0.0284747 |
| STM14_RS04110 | potE | putrescine-ornithine antiporter | WP_000042658.1 | 0.362311504 | 0.08980351 | 0.15422412 |
| STM14_RS20030 | rbsK | ribokinase | WP_000350272.1 | 0.361716211 | 0.00494642 | 0.01291682 |
| STM14_RS00120 | | hypothetical protein | WP_010999942.1 | 0.36118515 | 0.0009395 | 0.00305046 |
| STM14_RS22215 | malK | maltose/maltodextrin ABC transporter ATP-binding protein MalK | WP_000179176.1 | 0.360561109 | 0.13023048 | 0.20911847 |
| STM14_RS10080 | cspE | transcription antiterminator/RNA stability regulator CspE | WP_001062678.1 | 0.359720102 | 7.4329E-05 | 0.00032198 |
| STM14_RS00935 | caiB | L-carnitine CoA-transferase | WP_001016212.1 | 0.358487686 | 0.08755875 | 0.15093226 |
| STM14_RS09110 | hslJ | heat shock protein HslJ | WP_000725242.1 | 0.357986768 | 0.06695062 | 0.12001567 |
| STM14_RS07235 | | DUF333 domain-containing protein | WP_000673492.1 | 0.356739191 | 0.01187039 | 0.02755786 |
| STM14_RS08785 | nhoA | N-hydroxyarylamine O-acetyltransferase | WP_000200329.1 | 0.356454109 | 0.05760144 | 0.10551666 |
| STM14_RS22510 | yjdN | VOC family metalloprotein YjdN | WP_001131299.1 | 0.3563525 | 0.07022855 | 0.12516184 |
| STM14_RS13500 | | DUF1963 domain-containing protein | WP_000344439.1 | 0.355816952 | 0.04756425 | 0.08930728 |
| STM14_RS18255 | rplW | 50S ribosomal protein L23 | WP_000617546.1 | 0.355749452 | 0.00026705 | 0.00099928 |
| STM14_RS16050 | thyA | thymidylate synthase | WP_000816247.1 | 0.35538117 | 0.0902663 | 0.15480229 |
| STM14_RS04990 | | aspartate:alanine antiporter | WP_001024853.1 | 0.35528126 | 0.0166665 | 0.03659631 |
| STM14_RS20605 | | helix-turn-helix domain-containing protein | WP_000560824.1 | 0.355166933 | 0.05861549 | 0.10721422 |
| STM14_RS11115 | | helix-turn-helix transcriptional regulator | WP_001191666.1 | 0.355049262 | 0.00753722 | 0.01872608 |
| STM14_RS07180 | | helix-turn-helix transcriptional regulator | WP_001738208.1 | 0.354764257 | 0.11693735 | 0.19170229 |
| STM14_RS16535 | gshB | glutathione synthase | WP_000593248.1 | 0.35474805 | 0.0131937 | 0.03013599 |
| STM14_RS09705 | narH | nitrate reductase subunit beta | WP_000702625.1 | 0.354605119 | 0.13856887 | 0.22027424 |
| STM14_RS04280 | sdhD | succinate dehydrogenase membrane anchor subunit | WP_000254356.1 | 0.354435024 | 0.00202918 | 0.00602051 |
| STM14_RS01385 | yacL | protein YacL | WP_000384308.1 | 0.354416891 | 0.0001841 | 0.00071808 |
| STM14_RS21530 | ftsN | cell division protein FtsN | WP_000068811.1 | 0.354210406 | 0.0034209 | 0.00946338 |
| STM14_RS14605 | dgcN | diguanylate cyclase DgcN | WP_001030985.1 | 0.353686151 | 0.09736119 | 0.16466876 |
| STM14_RS19640 | | lipopolysaccharide N-acetylglucosaminyltransferase | WP_000591818.1 | 0.35259449 | 0.01231706 | 0.02846918 |
| STM14_RS14220 | | phage tail tube protein | WP_000971954.1 | 0.351904573 | 0.04749891 | 0.08923009 |
| STM14_RS09540 | | general stress protein | WP_000807657.1 | 0.351548676 | 0.00177773 | 0.00536524 |
| STM14_RS02610 | phoR | phosphate regulon sensor histidine kinase PhoR | WP_000893645.1 | 0.351132636 | 0.02253384 | 0.04719546 |
| STM14_RS04790 | dps | DNA starvation/stationary phase protection protein Dps | WP_000100805.1 | 0.350786044 | 1.7878E-05 | 9.0424E-05 |
| STM14_RS20435 | atpB | F0F1 ATP synthase subunit A | WP_000135632.1 | 0.35042338 | 0.00162583 | 0.00494735 |
| STM14_RS01285 | coaE | dephospho-CoA kinase | WP_001270913.1 | 0.350388318 | 0.1576602 | 0.24504215 |
| STM14_RS26605 | | DUF932 domain-containing protein | | 0.350072342 | 0.16926118 | 0.25849011 |
| STM14_RS15315 | hypB | hydrogenase nickel incorporation protein HypB | WP_000337636.1 | 0.349456943 | 0.00220486 | 0.00643288 |
| STM14_RS09630 | oppA | oligopeptide ABC transporter substrate-binding protein OppA | WP_001521098.1 | 0.348237834 | 0.00031966 | 0.00117463 |
| STM14_RS00915 | caiF | carnitine metabolism transcriptional regulator CaiF | WP_000333324.1 | 0.348112151 | 0.00061582 | 0.00209947 |
| STM14_RS04015 | nagD | ribonucleotide monophosphatase NagD | WP_000153143.1 | 0.34802988 | 0.06559803 | 0.1180508 |
| STM14_RS21465 | | epimerase | WP_001088049.1 | 0.347961626 | 0.01695183 | 0.03704251 |
| STM14_RS01070 | | YdcF family protein | WP_000877512.1 | 0.347915756 | 0.18075709 | 0.27265367 |
| STM14_RS11640 | rfbV | abequosyltransferase RfbV | WP_000908622.1 | 0.347708973 | 0.00606292 | 0.01551333 |
| STM14_RS24965 | | helix-turn-helix domain-containing protein | | 0.347604396 | 0.01452288 | 0.03264546 |
| STM14_RS14105 | | YfhL family 4Fe-4S dicluster ferredoxin | WP_001196290.1 | 0.34710051 | 0.19397302 | 0.28856916 |
| STM14_RS09570 | | YciC family protein | WP_000028507.1 | 0.34661768 | 0.02007571 | 0.04282663 |
| STM14_RS23670 | | DUF445 domain-containing protein | WP_001033021.1 | 0.34643361 | 0.06190678 | 0.11234142 |
| STM14_RS11040 | | KilA-N domain-containing protein | | 0.346234846 | 0.1601604 | 0.2481947 |
| STM14_RS06150 | | MFS transporter | WP_000560761.1 | 0.345858507 | 0.2236071 | 0.32388724 |
| STM14_RS08295 | | ROK family transcriptional regulator | WP_000225096.1 | 0.345437446 | 0.00574046 | 0.01478068 |
| STM14_RS23150 | ppa | inorganic diphosphatase | WP_000055079.1 | 0.345238816 | 0.00048502 | 0.00170246 |
| STM14_RS08840 | srfA | virulence-associated effector SrfA | WP_001129948.1 | 0.344921042 | 0.11934348 | 0.19470492 |
| STM14_RS01255 | secM | secA translation cis-regulator SecM | WP_000014333.1 | 0.344039013 | 0.01233539 | 0.0284747 |
| STM14_RS08520 | | hydrogenase expression/formation C-terminal domain-containing prote | WP_001016743.1 | 0.343179219 | 0.0115202 | 0.02691458 |
| STM14_RS18240 | rplV | 50S ribosomal protein L22 | WP_000447529.1 | 0.342996153 | 0.00056915 | 0.0019585 |
| STM14_RS24695 | | IS481 family transposase | | 0.342248306 | 0.09294577 | 0.15865847 |
| STM14_RS23545 | | ATP-binding protein | WP_001526178.1 | 0.340130945 | 0.00666138 | 0.01685719 |
| STM14_RS16230 | | DUF1460 domain-containing protein | WP_010989071.1 | 0.339877065 | 0.01158587 | 0.02701658 |
| STM14_RS15195 | srlA | PTS glucitol/sorbitol transporter subunit IIC | WP_000573333.1 | 0.339770503 | 0.32830705 | 0.44005335 |
| STM14_RS02985 | tomB | Hha toxicity modulator TomB | WP_000344807.1 | 0.338963259 | 0.00104008 | 0.0033099 |
| STM14_RS06230 | | type 1 fimbrial protein | WP_000111254.1 | 0.338940527 | 0.17651994 | 0.26713786 |

| | | | | | | |
|---|---|---|---|---|---|---|
| STM14_RS23095 | | bifunctional 2'%2C3'-cyclic-nucleotide 2'-phosphodiesterase/3'-nucleo | WP_000589395.1 | 0.338300709 | 0.01680334 | 0.03676168 |
| STM14_RS00940 | caiA | crotonobetainyl-CoA dehydrogenase | WP_000347134.1 | 0.338261854 | 0.24151203 | 0.34362383 |
| STM14_RS22705 | aspA | aspartate ammonia-lyase | WP_000069440.1 | 0.338167005 | 6.4207E-05 | 0.00028386 |
| STM14_RS04325 | cydB | cytochrome d ubiquinol oxidase subunit II | WP_000568258.1 | 0.337729502 | 0.00098819 | 0.00317774 |
| STM14_RS12595 | nrdA | ribonucleoside-diphosphate reductase subunit alpha | WP_001076487.1 | 0.337727426 | 0.00062524 | 0.00212765 |
| STM14_RS01185 | ftsL | cell division protein FtsL | WP_000625651.1 | 0.337402459 | 0.01399499 | 0.03167152 |
| STM14_RS10365 | znuA | zinc ABC transporter substrate-binding protein ZnuA | WP_000939597.1 | 0.337344798 | 0.01820507 | 0.03938371 |
| STM14_RS12330 | rplY | 50S ribosomal protein L25 | WP_000494192.1 | 0.336680533 | 0.00044567 | 0.00157938 |
| STM14_RS05590 | | hypothetical protein | WP_010989004.1 | 0.336363361 | 0.01142618 | 0.02672885 |
| STM14_RS21560 | metJ | met regulon transcriptional regulator MetJ | WP_000852810.1 | 0.336259285 | 0.00207829 | 0.00614639 |
| STM14_RS17275 | tdcG | L-serine ammonia-lyase | WP_000622081.1 | 0.336088186 | 0.02242327 | 0.04701735 |
| STM14_RS21375 | fieF | CDF family cation-efflux transporter FieF | WP_001077324.1 | 0.335518007 | 0.00159976 | 0.00487608 |
| STM14_RS03445 | pheP | phenylalanine transporter | WP_000786288.1 | 0.335342498 | 0.09587284 | 0.16282462 |
| STM14_RS17845 | | GntR family transcriptional regulator | WP_001171579.1 | 0.335292249 | 0.10085408 | 0.16981864 |
| STM14_RS10370 | znuC | zinc ABC transporter ATP-binding protein ZnuC | WP_000203014.1 | 0.335280905 | 0.0664557 | 0.11918651 |
| STM14_RS03865 | | 2-keto-3-deoxygluconate permease | WP_000694477.1 | 0.33380051 | 0.02886084 | 0.05825966 |
| STM14_RS00530 | traD | type IV conjugative transfer system coupling protein TraD | WP_000009318.1 | 0.332404498 | 0.11523515 | 0.18984154 |
| STM14_RS15725 | cas2e | type I-E CRISPR-associated endoribonuclease Cas2e | WP_001518648.1 | 0.331829055 | 0.16472526 | 0.25334937 |
| STM14_RS23200 | reiD | myo-inositol utilization transcriptional regulator ReiD | WP_000332219.1 | 0.331277986 | 0.04441877 | 0.08421726 |
| STM14_RS14725 | | hypothetical protein | WP_001161781.1 | 0.331232382 | 0.01894932 | 0.04077815 |
| STM14_RS24080 | arcA | two-component system response regulator ArcA | WP_001194359.1 | 0.327333052 | 9.2255E-05 | 0.00038821 |
| STM14_RS03675 | uspG | universal stress protein UspG | WP_000278499.1 | 0.327287273 | 8.9964E-05 | 0.00037987 |
| STM14_RS12065 | | SDR family oxidoreductase | WP_000169608.1 | 0.327104427 | 0.05969554 | 0.10897322 |
| STM14_RS20455 | mioC | FMN-binding protein MioC | WP_000763722.1 | 0.326423358 | 0.01613562 | 0.0356825 |
| STM14_RS12670 | | IclR family transcriptional regulator | WP_000894170.1 | 0.326283714 | 0.04103526 | 0.07853006 |
| STM14_RS12715 | arnT | lipid IV(A) 4-amino-4-deoxy-L-arabinosyltransferase | WP_000978038.1 | 0.326151811 | 0.08419311 | 0.14615702 |
| STM14_RS13930 | iscA | iron-sulfur cluster assembly protein IscA | WP_000028952.1 | 0.324807887 | 0.00113876 | 0.00359906 |
| STM14_RS12235 | fruB | fused PTS fructose transporter subunit IIA/HPr protein | WP_000487287.1 | 0.323820252 | 0.1261295 | 0.20350955 |
| STM14_RS00110 | | RepB family plasmid replication initiator protein | WP_001527061.1 | 0.323665061 | 0.00085013 | 0.00278487 |
| STM14_RS23550 | pglX | BREX-1 system adenine-specific DNA-methyltransferase PglX | WP_001095591.1 | 0.322877686 | 0.00365798 | 0.01002135 |
| STM14_RS12875 | | SLC13 family permease | WP_001012861.1 | 0.322389269 | 0.04033751 | 0.07743624 |
| STM14_RS14475 | | LysR family transcriptional regulator | WP_001526875.1 | 0.322021014 | 0.05426652 | 0.10030588 |
| STM14_RS05890 | rlmI | 23S rRNA (cytosine(1962)-C(5))-methyltransferase RlmI | WP_000140480.1 | 0.321939422 | 0.19538755 | 0.29032161 |
| STM14_RS07650 | aroD | type I 3-dehydroquinate dehydratase | WP_000860225.1 | 0.321900843 | 0.04321379 | 0.08227156 |
| STM14_RS16485 | | FadR/GntR family transcriptional regulator | WP_001576357.1 | 0.32094083 | 0.00085683 | 0.00280432 |
| STM14_RS16355 | zapA | cell division protein ZapA | WP_001276011.1 | 0.320825466 | 0.00055172 | 0.00190712 |
| STM14_RS12775 | rbn | ribonuclease BN | WP_024132429.1 | 0.320751346 | 0.05529261 | 0.10188215 |
| STM14_RS16785 | hybG | hydrogenase maturation factor HybG | WP_000334887.1 | 0.319968633 | 0.01386705 | 0.03143994 |
| STM14_RS06670 | purB | adenylosuccinate lyase | WP_000423763.1 | 0.319341419 | 0.03262643 | 0.06441314 |
| STM14_RS16445 | | M48 family metallopeptidase | WP_000701830.1 | 0.319334172 | 0.06185455 | 0.11230199 |
| STM14_RS04010 | asnB | asparagine synthase B | WP_000337041.1 | 0.319180172 | 0.01502042 | 0.03366109 |
| STM14_RS23950 | | fimbrial biogenesis outer membrane usher protein | WP_000876250.1 | 0.316833101 | 0.177289 | 0.26786135 |
| STM14_RS05950 | | YlcI/YnfO family protein | WP_001111006.1 | 0.316238 | 0.04246912 | 0.08097945 |
| STM14_RS14830 | | integrase arm-type DNA-binding domain-containing protein | WP_001738699.1 | 0.3155558 | 0.02168689 | 0.04567254 |
| STM14_RS09150 | | tyrosine-type recombinase/integrase | | 0.314385484 | 0.10497078 | 0.17568291 |
| STM14_RS19875 | cigR | anti-virulence regulatory inner membrane protein CigR | WP_001065324.1 | 0.313897911 | 0.00717761 | 0.01797821 |
| STM14_RS18160 | rpmJ | 50S ribosomal protein L36 | WP_000868187.1 | 0.312700261 | 0.00473096 | 0.01244242 |
| STM14_RS23155 | fbp | class 1 fructose-bisphosphatase | WP_000853764.1 | 0.312610384 | 0.00296775 | 0.00837338 |
| STM14_RS05835 | matP | macrodomain Ter protein MatP | WP_000877172.1 | 0.312082829 | 0.00616789 | 0.01574907 |
| STM14_RS08690 | bdm | biofilm-dependent modulation protein | WP_000495700.1 | 0.311810733 | 0.02143838 | 0.04533654 |
| STM14_RS12690 | pmrG | lipopolysaccharide core heptose(II)-phosphate phosphatase PmrG | WP_000879248.1 | 0.310807566 | 0.01967809 | 0.04212483 |
| STM14_RS09710 | | nitrate reductase subunit alpha | WP_000032881.1 | 0.310479634 | 0.06452609 | 0.11652039 |
| STM14_RS21965 | | acetyltransferase | WP_000973242.1 | 0.310432588 | 0.00497468 | 0.0129814 |
| STM14_RS21460 | lsrG | (4S)-4-hydroxy-5-phosphonooxypentane-2%2C3-dione isomerase | WP_001738619.1 | 0.3101841 | 0.02372241 | 0.04918125 |
| STM14_RS14170 | | prophage tail fiber N-terminal domain-containing protein | WP_001144691.1 | 0.309467914 | 0.29288326 | 0.40074179 |
| STM14_RS03495 | nfsB | oxygen-insensitive NAD(P)H nitroreductase | WP_000355879.1 | 0.308446593 | 0.03016437 | 0.06049303 |
| STM14_RS01905 | dnaQ | DNA polymerase III subunit epsilon | WP_001670675.1 | 0.308300377 | 0.01651651 | 0.03632842 |
| STM14_RS08405 | | GntR family transcriptional regulator | WP_000215563.1 | 0.308295949 | 0.02974574 | 0.05984908 |
| STM14_RS16680 | | anaerobic sulfatase maturase | WP_000933432.1 | 0.307593993 | 0.22885342 | 0.33018742 |
| STM14_RS05050 | rlmC | 23S rRNA (uracil(747)-C(5))-methyltransferase RlmC | WP_001149782.1 | 0.3058389 | 0.23960958 | 0.34129303 |
| STM14_RS00990 | srfN | DUF1471 family virulence protein SrfN | WP_000745961.1 | 0.305300908 | 0.03183535 | 0.06313114 |
| STM14_RS06285 | | YceI family protein | WP_000739886.1 | 0.305140264 | 0.31966756 | 0.43051059 |
| STM14_RS02905 | cof | HMP-PP phosphatase | WP_000113030.1 | 0.304936165 | 0.10344595 | 0.1736481 |
| STM14_RS02675 | yajD | HNH nuclease YajD | WP_000974818.1 | 0.304857988 | 0.00393943 | 0.01069557 |
| STM14_RS15215 | gutM | transcriptional regulator GutM | WP_000255198.1 | 0.304556278 | 0.25684957 | 0.36137566 |
| STM14_RS18300 | rpsG | 30S ribosomal protein S7 | WP_001138042.1 | 0.304551857 | 0.00247645 | 0.0071516 |
| STM14_RS15380 | | type III secretion system effector protein OrgC | WP_000612175.1 | 0.303450917 | 0.01125816 | 0.02638609 |
| STM14_RS11820 | | MASE1 domain-containing protein | WP_000042956.1 | 0.303297668 | 0.12885758 | 0.2070042 |
| STM14_RS21590 | | bifunctional UDP-sugar hydrolase/5'-nucleotidase | WP_000865249.1 | 0.30326264 | 0.20764843 | 0.30509238 |
| STM14_RS03165 | | MetQ/NlpA family ABC transporter substrate-binding protein | WP_000524317.1 | 0.303084675 | 0.1315394 | 0.21094428 |
| STM14_RS01485 | panD | aspartate 1-decarboxylase | WP_000621526.1 | 0.302826871 | 0.09109632 | 0.15608034 |
| STM14_RS08835 | | YdcY family protein | WP_000018720.1 | 0.301872341 | 0.02169167 | 0.04567254 |
| STM14_RS11620 | rfbK | O-antigen biosynthesis phosphomannomutase RfbK | WP_001103643.1 | 0.301536975 | 0.00723782 | 0.01811669 |
| STM14_RS26310 | | host specificity protein J | | 0.298654975 | 0.19723542 | 0.29259501 |
| STM14_RS01275 | yacG | DNA gyrase inhibitor YacG | WP_001286419.1 | 0.298379716 | 0.27655714 | 0.38295728 |
| STM14_RS06755 | | helix-turn-helix domain-containing protein | WP_000169863.1 | 0.298358483 | 0.00643532 | 0.01634126 |
| STM14_RS23115 | | hemolysin family protein | WP_000934974.1 | 0.29723209 | 0.0119647 | 0.02774556 |
| STM14_RS03875 | | zinc ribbon-containing protein | WP_001044884.1 | 0.297189103 | 0.00069235 | 0.00232383 |
| STM14_RS19770 | gmk | guanylate kinase | WP_000046969.1 | 0.296015363 | 0.00628962 | 0.01602656 |
| STM14_RS18780 | | oxidoreductase | WP_000172397.1 | 0.295698113 | 0.14068798 | 0.22299318 |
| STM14_RS16655 | ripA | itaconate CoA-transferase RipA/Ict | WP_000982337.1 | 0.295427667 | 0.18262149 | 0.2751278 |
| STM14_RS19890 | | EamA family transporter | WP_001682335.1 | 0.295044835 | 0.15925579 | 0.24720903 |

| | | | | | | |
|---|---|---|---|---|---|---|
| STM14_RS06890 | | phage portal protein | WP_000831821.1 | 0.294124196 | 0.1679701 | 0.25694471 |
| STM14_RS05295 | | amino acid permease | WP_001134268.1 | 0.293165523 | 0.07681586 | 0.13545785 |
| STM14_RS18455 | bigA | autotransporter adhesin BigA | WP_014344197.1 | 0.29229804 | 0.11597329 | 0.19063109 |
| STM14_RS15325 | hypD | hydrogenase formation protein HypD | WP_001212958.1 | 0.29226191 | 0.03168243 | 0.06292056 |
| STM14_RS09265 | | aldo/keto reductase | WP_000456774.1 | 0.292090176 | 0.03348757 | 0.06583088 |
| STM14_RS02960 | | type B 50S ribosomal protein L31 | WP_000801415.1 | 0.291273857 | 0.24786915 | 0.35129107 |
| STM14_RS08175 | add | adenosine deaminase | WP_000565569.1 | 0.290620603 | 0.06370186 | 0.11520149 |
| STM14_RS08185 | | YdgA family protein | WP_000753327.1 | 0.28998303 | 0.00993009 | 0.02374195 |
| STM14_RS08560 | | FadR/GntR family transcriptional regulator | WP_001518759.1 | 0.289201522 | 0.02442124 | 0.0504596 |
| STM14_RS03185 | allS | HTH-type transcriptional activator AllS | WP_000460169.1 | 0.288768168 | 0.4193694 | 0.52971462 |
| STM14_RS05255 | lolA | outer membrane lipoprotein chaperone LolA | WP_001519746.1 | 0.288730362 | 0.00791303 | 0.01956734 |
| STM14_RS13620 | nudK | GDP-mannose pyrophosphatase NudK | WP_000084037.1 | 0.288377893 | 0.04913794 | 0.09165445 |
| STM14_RS22600 | | sensor histidine kinase | WP_000682911.1 | 0.288284224 | 0.06792743 | 0.121471 |
| STM14_RS11495 | pduX | L-threonine kinase PduX | WP_001526166.1 | 0.286808108 | 0.34550883 | 0.45586429 |
| STM14_RS17655 | rpoN | RNA polymerase factor sigma-54 | WP_000809020.1 | 0.2866511 | 0.00254035 | 0.00729608 |
| STM14_RS00680 | bcfC | fimbrial usher BcfC | WP_001245854.1 | 0.28638301 | 0.32448876 | 0.43588749 |
| STM14_RS02060 | sciX | protein SciX | WP_000935097.1 | 0.286338359 | 0.38170334 | 0.49278812 |
| STM14_RS03145 | tesA | multifunctional acyl-CoA thioesterase I/protease I/lysophospholipase L | WP_001010574.1 | 0.286256776 | 0.27986094 | 0.38695005 |
| STM14_RS08315 | osmW | osmoprotectant ABC transporter permease OsmW | WP_000379677.1 | 0.285737753 | 0.21040983 | 0.30816587 |
| STM14_RS08390 | | MFS transporter | | 0.285650856 | 0.3829258 | 0.4940199 |
| STM14_RS16840 | | RNA helicase | WP_000433046.1 | 0.285557229 | 0.0286356 | 0.05787499 |
| STM14_RS10065 | mntP | manganese efflux pump MntP | WP_001518359.1 | 0.285537967 | 0.3400187 | 0.45147813 |
| STM14_RS18310 | tusB | sulfurtransferase complex subunit TusB | WP_000903400.1 | 0.284295914 | 0.26521572 | 0.37101987 |
| STM14_RS06815 | | DUF1367 family protein | WP_000929785.1 | 0.283751604 | 0.43802399 | 0.54831827 |
| STM14_RS18105 | rsmB | 16S rRNA (cytosine(967)-C(5))-methyltransferase RsmB | WP_000744604.1 | 0.28233934 | 0.16807672 | 0.25700103 |
| STM14_RS11530 | phsA | thiosulfate reductase PhsA | WP_000028228.1 | 0.281922099 | 0.00539894 | 0.01398938 |
| STM14_RS13740 | ppk1 | polyphosphate kinase 1 | WP_000529555.1 | 0.281830385 | 0.00568419 | 0.01465609 |
| STM14_RS16280 | | protein YgfX | WP_000244756.1 | 0.2817311 | 0.03682244 | 0.07162188 |
| STM14_RS13145 | smrB | endonuclease SmrB | WP_000730794.1 | 0.281056603 | 0.09610324 | 0.16314067 |
| STM14_RS08230 | rstA | two-component system response regulator RstA | WP_001080048.1 | 0.2810373 | 0.0466211 | 0.08780506 |
| STM14_RS15005 | | YqaE/Pmp3 family membrane protein | WP_000508175.1 | 0.28073542 | 0.10673467 | 0.17790723 |
| STM14_RS19125 | dctA | C4-dicarboxylate transporter DctC | WP_000858228.1 | 0.280098353 | 0.00418917 | 0.01125878 |
| STM14_RS05090 | artP | arginine ABC transporter ATP-binding protein ArtP | WP_000027186.1 | 0.279450652 | 0.0236228 | 0.04900233 |
| STM14_RS17215 | | DUF1090 domain-containing protein | WP_000877297.1 | 0.27933104 | 0.00081667 | 0.00270168 |
| STM14_RS10655 | | DUF2594 family protein | WP_000106483.1 | 0.279077825 | 0.11877628 | 0.19432572 |
| STM14_RS07535 | rpmI | 50S ribosomal protein L35 | WP_001124225.1 | 0.278830765 | 0.00103991 | 0.0033099 |
| STM14_RS02335 | | hypothetical protein | WP_000545421.1 | 0.278816699 | 0.41005805 | 0.52110646 |
| STM14_RS18280 | bfr | bacterioferritin | WP_000675530.1 | 0.27871271 | 0.00097349 | 0.0031442 |
| STM14_RS11545 | plaP | putrescine/proton symporter PlaP | WP_001519005.1 | 0.278109907 | 0.03766316 | 0.07292675 |
| STM14_RS09160 | dbpA | ATP-dependent RNA helicase DbpA | WP_000206674.1 | 0.276784284 | 0.17178356 | 0.26158274 |
| STM14_RS02170 | dpaA | peptidoglycan meso-diaminopimelic acid protein amidase | WP_001225658.1 | 0.275811678 | 0.0360364 | 0.07024141 |
| STM14_RS10680 | tcyJ | cystine ABC transporter substrate-binding protein | WP_000761635.1 | 0.275780584 | 0.00466539 | 0.01229633 |
| STM14_RS22230 | ubiC | chorismate lyase | WP_000019219.1 | 0.274934918 | 0.02689356 | 0.05495122 |
| STM14_RS03015 | rsmS | pleiotropic regulatory protein RsmS | WP_000051161.1 | 0.274606095 | 0.30215507 | 0.41145889 |
| STM14_RS03125 | fetA | iron ABC transporter ATP-binding protein FetA | WP_000140195.1 | 0.27321771 | 0.19741094 | 0.29273745 |
| STM14_RS06080 | | YccJ family protein | WP_001143125.1 | 0.272641059 | 0.00105137 | 0.00334006 |
| STM14_RS05480 | | site-specific integrase | WP_001262308.1 | 0.272349805 | 0.01034009 | 0.02455911 |
| STM14_RS21425 | lsrK | autoinducer-2 kinase | WP_000113085.1 | 0.271518978 | 0.01531289 | 0.03419167 |
| STM14_RS03050 | htpG | molecular chaperone HtpG | WP_001520210.1 | 0.271126965 | 0.01006681 | 0.02397542 |
| STM14_RS14395 | | hypothetical protein | WP_000356948.1 | 0.271114481 | 0.13440717 | 0.21479479 |
| STM14_RS21695 | fabR | HTH-type transcriptional repressor FabR | WP_000971958.1 | 0.270920587 | 0.02199383 | 0.04619582 |
| STM14_RS02145 | | Spy/CpxP family protein refolding chaperone | WP_000788200.1 | 0.270360315 | 0.33334547 | 0.44518609 |
| STM14_RS11875 | | DUF4034 domain-containing protein | WP_000434241.1 | 0.269140245 | 0.38490668 | 0.49605404 |
| STM14_RS09195 | | DUF2534 family protein | WP_000605645.1 | 0.268249341 | 0.00668455 | 0.0168926 |
| STM14_RS15025 | stpA | DNA-binding protein StpA | WP_001051100.1 | 0.267927643 | 0.00351997 | 0.00966483 |
| STM14_RS11055 | | phage holin%2C lambda family | WP_001527046.1 | 0.267228989 | 0.35980138 | 0.46994986 |
| STM14_RS20495 | rbsC | ribose ABC transporter permease | WP_000211345.1 | 0.266337809 | 0.1554714 | 0.24256174 |
| STM14_RS08470 | | sugar transporter | WP_000154625.1 | 0.266006399 | 0.42139581 | 0.53165354 |
| STM14_RS05140 | | lysine exporter LysO family protein | WP_000491118.1 | 0.265966245 | 0.13101004 | 0.21018692 |
| STM14_RS12140 | preA | NAD-dependent dihydropyrimidine dehydrogenase subunit PreA | WP_000956095.1 | 0.265573368 | 0.00904911 | 0.0218771 |
| STM14_RS17270 | | amino acid permease | WP_000443187.1 | 0.264872924 | 0.09386304 | 0.15985371 |
| STM14_RS19560 | | MFS transporter | WP_001092104.1 | 0.264422856 | 0.4153406 | 0.52606952 |
| STM14_RS08655 | | pyridoxal phosphate-dependent aminotransferase | WP_000347159.1 | 0.264262754 | 0.25820908 | 0.36287246 |
| STM14_RS01475 | | PTS sugar transporter subunit IIA | WP_000901960.1 | 0.263850298 | 0.02386366 | 0.04939068 |
| STM14_RS18500 | aroK | shikimate kinase AroK | WP_000818621.1 | 0.263738525 | 0.00641197 | 0.01630448 |
| STM14_RS16255 | prfB | peptide chain release factor 2 | WP_010989230.1 | 0.263566808 | 0.00873008 | 0.02125936 |
| STM14_RS26900 | timP | small toxic inner membrane protein TimP | WP_242617172.1 | 0.263209967 | 0.03703218 | 0.07195382 |
| STM14_RS07530 | infC | translation initiation factor IF-3 | WP_001574431.1 | 0.263046172 | 0.00116076 | 0.00365293 |
| STM14_RS16585 | | DUF1202 domain-containing protein | WP_000252197.1 | 0.262600261 | 0.25169477 | 0.35548146 |
| STM14_RS08685 | | OsmC family protein | WP_000152297.1 | 0.262220875 | 0.00599869 | 0.01538104 |
| STM14_RS06265 | | MysB family protein | WP_000180050.1 | 0.261182887 | 0.00476547 | 0.0125064 |
| STM14_RS12555 | rcsD | phosphotransferase RcsD | WP_000083201.1 | 0.260935446 | 0.01619421 | 0.03574766 |
| STM14_RS10490 | cheY | chemotaxis response regulator CheY | WP_000763861.1 | 0.260907774 | 0.00999163 | 0.02385809 |
| STM14_RS07120 | | cytochrome b | WP_000929982.1 | 0.260161215 | 0.06596099 | 0.11847237 |
| STM14_RS00080 | pefD | fimbrial chaperone PefD | WP_001038510.1 | 0.260132922 | 0.44858549 | 0.55863166 |
| STM14_RS14720 | | SEC-C domain-containing protein | WP_000248794.1 | 0.259811241 | 0.06335774 | 0.11468204 |
| STM14_RS06405 | | flagellar basal body P-ring protein FlgI | WP_001518955.1 | 0.259387646 | 0.1741612 | 0.26465602 |
| STM14_RS00605 | mog | molybdopterin adenylyltransferase | WP_000380372.1 | 0.259282622 | 0.08713863 | 0.15027842 |
| STM14_RS14640 | | HlyC/CorC family transporter | WP_127172650.1 | 0.259131573 | 0.09040298 | 0.15496452 |
| STM14_RS18945 | tusA | sulfurtransferase TusA | WP_001541054.1 | 0.258953574 | 0.0863692 | 0.14916107 |
| STM14_RS23500 | idnK | gluconokinase | WP_000896756.1 | 0.258870271 | 0.21438931 | 0.31225532 |
| STM14_RS23110 | | DUF1107 domain-containing protein | WP_000689229.1 | 0.258387554 | 0.0087211 | 0.02125936 |

| | | | | | | |
|---|---|---|---|---|---|---|
| STM14_RS00980 | | YgdI/YgdR family lipoprotein | WP_001172089.1 | 0.258259613 | 0.2353548 | 0.3373205 |
| STM14_RS21090 | glnA | glutamate--ammonia ligase | WP_001271699.1 | 0.257797402 | 0.00945481 | 0.02273847 |
| STM14_RS10105 | kdgR | DNA-binding transcriptional regulator KdgR | WP_001262195.1 | 0.25758031 | 0.00681751 | 0.01718675 |
| STM14_RS18535 | nudE | ADP compounds hydrolase NudE | WP_000045725.1 | 0.257374895 | 0.11113064 | 0.18406794 |
| STM14_RS18715 | | dihydroxy-acid dehydratase | WP_000078494.1 | 0.25667505 | 0.27631778 | 0.38276978 |
| STM14_RS01175 | mraZ | division/cell wall cluster transcriptional repressor MraZ | WP_000488294.1 | 0.255663896 | 0.0176835 | 0.03841337 |
| STM14_RS08925 | ydcK | YdcK family protein | WP_000170589.1 | 0.255470591 | 0.23045948 | 0.33211421 |
| STM14_RS17950 | msrP | protein-methionine-sulfoxide reductase catalytic subunit MsrP | WP_000723876.1 | 0.255046272 | 0.32060004 | 0.43129314 |
| STM14_RS13485 | amiA | N-acetylmuramoyl-L-alanine amidase AmiA | WP_000102881.1 | 0.254832036 | 0.07051719 | 0.12561407 |
| STM14_RS00835 | | sensor histidine kinase | WP_001737977.1 | 0.25476616 | 0.27689712 | 0.38328391 |
| STM14_RS25930 | | transposase | | 0.254436431 | 0.12861503 | 0.206868 |
| STM14_RS18200 | rpsN | 30S ribosomal protein S14 | WP_001118932.1 | 0.254070439 | 0.00626187 | 0.0159669 |
| STM14_RS10595 | tyrP | tyrosine transporter TyrP | WP_001562965.1 | 0.253351142 | 0.40661629 | 0.51751165 |
| STM14_RS15820 | | type II toxin-antitoxin system ParD family antitoxin | WP_000480218.1 | 0.251206131 | 0.08096855 | 0.14122511 |
| STM14_RS21540 | priA | primosomal protein N' | WP_000109379.1 | 0.250544964 | 0.05983692 | 0.10906908 |
| STM14_RS10015 | pabB | aminodeoxychorismate synthase component 1 | WP_000978464.1 | 0.250404739 | 0.12100754 | 0.19688457 |
| STM14_RS09550 | | ferritin-like domain-containing protein | WP_001109977.1 | 0.249254413 | 0.02079736 | 0.0441145 |
| STM14_RS06135 | | N-acetylmannosamine-6-phosphate 2-epimerase | WP_000054243.1 | 0.24920484 | 0.33039679 | 0.44188921 |
| STM14_RS19860 | | membrane protein | WP_001653911.1 | 0.248948028 | 0.39655796 | 0.50698833 |
| STM14_RS26795 | | baseplate J/gp47 family protein | | 0.24859606 | 0.3701102 | 0.48068633 |
| STM14_RS22810 | | YjeO family protein | WP_000074657.1 | 0.24837409 | 0.50108051 | 0.60870288 |
| STM14_RS14705 | | integrase domain-containing protein | | 0.248239264 | 0.08654037 | 0.14938661 |
| STM14_RS16705 | | FAD-binding oxidoreductase | WP_001092814.1 | 0.247764494 | 0.32432131 | 0.43582156 |
| STM14_RS05645 | | Clp protease ClpP | WP_010989008.1 | 0.247722115 | 0.23818243 | 0.34018142 |
| STM14_RS14645 | | hypothetical protein | WP_001287923.1 | 0.247528865 | 0.26077047 | 0.36577405 |
| STM14_RS23655 | | nucleoside recognition domain-containing protein | WP_000068750.1 | 0.247075792 | 0.04544848 | 0.08577207 |
| STM14_RS19760 | | trimeric intracellular cation channel family protein | WP_000924334.1 | 0.246677742 | 0.44871361 | 0.55863166 |
| STM14_RS07270 | | YbaK/prolyl-tRNA synthetase associated domain-containing protein | WP_010989016.1 | 0.246484579 | 0.29300485 | 0.40074581 |
| STM14_RS02035 | | type VI secretion system Vgr family protein | WP_000011535.1 | 0.246330535 | 0.51410892 | 0.62047696 |
| STM14_RS08400 | ydfZ | putative selenium delivery protein YdfZ | WP_000989267.1 | 0.245477184 | 0.0156439 | 0.03472021 |
| STM14_RS02980 | | HHA domain-containing protein | WP_001280991.1 | 0.244166866 | 0.03049803 | 0.06102921 |
| STM14_RS10385 | ruvA | Holliday junction branch migration protein RuvA | WP_000580335.1 | 0.243732782 | 0.12336041 | 0.20009385 |
| STM14_RS15880 | queF | NADPH-dependent 7-cyano-7-deazaguanine reductase QueF | WP_000100474.1 | 0.243726123 | 0.10079295 | 0.16981864 |
| STM14_RS05555 | | DinI family protein | WP_001217665.1 | 0.241811475 | 0.23067381 | 0.33216307 |
| STM14_RS09520 | trpD | bifunctional anthranilate synthase glutamate amidotransferase compor | WP_000763492.1 | 0.241751354 | 0.43764461 | 0.54809778 |
| STM14_RS16175 | | transcriptional regulator | WP_000134791.1 | 0.241603903 | 0.36409961 | 0.4745539 |
| STM14_RS22365 | siiD | SPI-4 type I secretion system protein SiiD | WP_001081935.1 | 0.2415231 | 0.11834137 | 0.19374518 |
| STM14_RS20560 | maoP | macrodomain Ori organization protein MaoP | WP_000840996.1 | 0.241478671 | 0.03633951 | 0.07075731 |
| STM14_RS19030 | | phosphatase PAP2 family protein | WP_000766151.1 | 0.241311456 | 0.11087199 | 0.18372217 |
| STM14_RS23845 | | GGDEF domain-containing protein | WP_000211207.1 | 0.240487325 | 0.26728842 | 0.37306898 |
| STM14_RS22020 | | DUF2058 domain-containing protein | WP_001096724.1 | 0.240212162 | 0.07467704 | 0.13193899 |
| STM14_RS07010 | sspH1 | type III secretion system effector E3 ubiquitin transferase SspH1 | WP_000481981.1 | 0.239923016 | 0.05993502 | 0.10919383 |
| STM14_RS21455 | lsrF | 3-hydroxy-5-phosphonooxypentane-2%2C4-dione thiolase | WP_000774147.1 | 0.239534315 | 0.12129392 | 0.19717625 |
| STM14_RS05370 | msbA | lipid A ABC transporter ATP-binding protein/permease MsbA | WP_000551246.1 | 0.239177728 | 0.06623955 | 0.11891468 |
| STM14_RS19060 | fraB | 6-phosphofructose-aspartate deglycase | WP_010989080.1 | 0.23846459 | 0.02382714 | 0.04937058 |
| STM14_RS00785 | rpsT | 30S ribosomal protein S20 | WP_001518655.1 | 0.237833367 | 0.02062176 | 0.04383909 |
| STM14_RS23805 | dnaC | DNA replication protein DnaC | WP_000799924.1 | 0.237184511 | 0.06313886 | 0.11435183 |
| STM14_RS12005 | | sensor histidine kinase | WP_000272845.1 | 0.235824445 | 0.11807613 | 0.19339694 |
| STM14_RS21445 | lsrD | autoinducer 2 ABC transporter permease LsrD | WP_000981826.1 | 0.235611445 | 0.0850113 | 0.14723029 |
| STM14_RS01450 | | ABC transporter permease | WP_000983414.1 | 0.235424877 | 0.47067239 | 0.57959561 |
| STM14_RS09565 | | YkgJ family cysteine cluster protein | WP_000114955.1 | 0.235359851 | 0.25131211 | 0.35507721 |
| STM14_RS08570 | | MFS transporter | WP_000824240.1 | 0.235337519 | 0.37436877 | 0.48553217 |
| STM14_RS23335 | nrdD | anaerobic ribonucleoside-triphosphate reductase | WP_000187818.1 | 0.235157059 | 0.02637034 | 0.05409225 |
| STM14_RS21915 | zraP | zinc resistance sensor/chaperone ZraP | WP_000828129.1 | 0.235082835 | 0.0615838 | 0.11186559 |
| STM14_RS09745 | chaA | sodium-potassium/proton antiporter ChaA | WP_000148418.1 | 0.234694793 | 0.28316497 | 0.39049191 |
| STM14_RS08045 | | DUF1289 domain-containing protein | WP_001519323.1 | 0.233974192 | 0.39488977 | 0.50555777 |
| STM14_RS09180 | ogt | methylated-DNA--[protein]-cysteine S-methyltransferase | WP_000945036.1 | 0.23338644 | 0.0284156 | 0.05752072 |
| STM14_RS20265 | dnaA | chromosomal replication initiator protein DnaA | WP_000059093.1 | 0.23315927 | 0.04807369 | 0.09008007 |
| STM14_RS09835 | | hydrogenase 1 maturation protease | WP_000993803.1 | 0.232511956 | 0.43811851 | 0.54831827 |
| STM14_RS20100 | | YidH family protein | WP_000703964.1 | 0.232238881 | 0.34866356 | 0.45898435 |
| STM14_RS12865 | alaA | alanine transaminase AlaA | WP_000074540.1 | 0.231597352 | 0.34555058 | 0.45586429 |
| STM14_RS23920 | | YjjI family glycine radical enzyme | WP_001111691.1 | 0.230675945 | 0.08002761 | 0.13984891 |
| STM14_RS14865 | | IS3 family transposase | WP_085983317.1 | 0.230667867 | 0.28416996 | 0.39143801 |
| STM14_RS07745 | pykF | pyruvate kinase PykF | WP_000751447.1 | 0.230235475 | 0.02726519 | 0.05561797 |
| STM14_RS22500 | lpxO | lipid A hydroxylase LpxO | WP_000457024.1 | 0.229655318 | 0.07647184 | 0.13491581 |
| STM14_RS02585 | araJ | MFS transporter AraJ | WP_000755620.1 | 0.22964497 | 0.45945639 | 0.56883605 |
| STM14_RS21625 | | PTS fructose transporter subunit EIIC | WP_000661597.1 | 0.229586803 | 0.22562209 | 0.32654895 |
| STM14_RS15165 | csrA | carbon storage regulator CsrA | WP_000906486.1 | 0.228457622 | 0.0162418 | 0.03583123 |
| STM14_RS13290 | | putative DNA-binding transcriptional regulator | WP_000472034.1 | 0.22833439 | 0.04667841 | 0.08786806 |
| STM14_RS00335 | | DNA polymerase III subunit theta | WP_000015983.1 | 0.227799159 | 0.46733169 | 0.57626098 |
| STM14_RS13940 | iscS | cysteine desulfurase | WP_000775263.1 | 0.226394001 | 0.03108842 | 0.06197486 |
| STM14_RS18090 | dprA | DNA-protecting protein DprA | WP_000124529.1 | 0.226327824 | 0.12414909 | 0.20101888 |
| STM14_RS08015 | | C40 family peptidase | WP_001520344.1 | 0.226160001 | 0.11562145 | 0.19015834 |
| STM14_RS23580 | | class I SAM-dependent methyltransferase | WP_000979786.1 | 0.225414768 | 0.23830133 | 0.34021927 |
| STM14_RS17880 | aaeB | p-hydroxybenzoic acid efflux pump subunit AaeB | WP_000510913.1 | 0.224904201 | 0.45923612 | 0.56880407 |
| STM14_RS22745 | epmB | EF-P beta-lysylation protein EpmB | WP_000940484.1 | 0.22489995 | 0.2242808 | 0.32473531 |
| STM14_RS00380 | | transglycosylase SLT domain-containing protein | WP_001526856.1 | 0.224147132 | 0.26658621 | 0.37237117 |
| STM14_RS04900 | bssR | biofilm formation regulator BssR | WP_000259651.1 | 0.222961455 | 0.00820595 | 0.02020086 |
| STM14_RS06120 | phoH | phosphate starvation-inducible protein PhoH | WP_000533527.1 | 0.222048612 | 0.01739283 | 0.03789373 |
| STM14_RS21450 | lsrB | autoinducer 2 ABC transporter substrate-binding protein LsrB | WP_000090737.1 | 0.221196718 | 0.14677064 | 0.23153791 |
| STM14_RS18085 | smg | DUF494 family protein Smg | WP_000460663.1 | 0.221134558 | 0.01599579 | 0.03539454 |
| STM14_RS13165 | | YfcZ/YiiS family protein | WP_000030904.1 | 0.220712456 | 0.01347732 | 0.03067443 |

| | | | | | | |
|---|---|---|---|---|---|---|
| STM14_RS14420 | | recombinase RecT | WP_077248255.1 | 0.220549546 | 0.44190543 | 0.55249433 |
| STM14_RS07255 | | helix-turn-helix transcriptional regulator | WP_000579793.1 | 0.220203142 | 0.43407747 | 0.54511366 |
| STM14_RS10945 | | hypothetical protein | WP_001066636.1 | 0.219757512 | 0.36548094 | 0.47601727 |
| STM14_RS10500 | cheR | protein-glutamate O-methyltransferase CheR | WP_000204362.1 | 0.217274649 | 0.08217982 | 0.14306672 |
| STM14_RS03905 | | molecular chaperone HscC | WP_000368034.1 | 0.216920381 | 0.13851928 | 0.22027424 |
| STM14_RS03915 | rihA | pyrimidine-specific ribonucleoside hydrolase RihA | WP_001207419.1 | 0.216887373 | 0.20007171 | 0.29620588 |
| STM14_RS18370 | | monooxygenase | WP_000047690.1 | 0.216784234 | 0.02771357 | 0.05637644 |
| STM14_RS12905 | yfcC | putative basic amino acid antiporter YfcC | WP_000117932.1 | 0.216734114 | 0.49855339 | 0.6064333 |
| STM14_RS19685 | rfaQ | lipopolysaccharide core heptosyltransferase RfaQ | WP_001526672.1 | 0.216348298 | 0.07730963 | 0.13615078 |
| STM14_RS10465 | | glycoside hydrolase family 105 protein | WP_000989908.1 | 0.216268542 | 0.32698064 | 0.43875463 |
| STM14_RS26805 | | hypothetical protein | | 0.215725484 | 0.12225981 | 0.19854088 |
| STM14_RS20200 | torS | TMAO reductase system sensor histidine kinase/response regulator Tor | WP_000106953.1 | 0.215548819 | 0.29882031 | 0.40765335 |
| STM14_RS17850 | | GntR family transcriptional regulator | WP_000723776.1 | 0.215463305 | 0.11164508 | 0.18467079 |
| STM14_RS00600 | tal | transaldolase | WP_000130175.1 | 0.215308433 | 0.05373252 | 0.09951869 |
| STM14_RS10045 | manX | PTS mannose transporter subunit IIAB | WP_000150521.1 | 0.215297495 | 0.02641926 | 0.05416243 |
| STM14_RS06770 | | replication protein | WP_000024048.1 | 0.214926994 | 0.42013251 | 0.53049654 |
| STM14_RS11920 | | ADP-ribosylglycohydrolase family protein | WP_000642793.1 | 0.214825307 | 0.26896564 | 0.3748416 |
| STM14_RS09530 | trpB | tryptophan synthase subunit beta | WP_000209485.1 | 0.213923278 | 0.41014843 | 0.52110646 |
| STM14_RS12575 | | mandelate racemase/muconate lactonizing enzyme family protein | WP_000705579.1 | 0.213692419 | 0.42673156 | 0.53735486 |
| STM14_RS06390 | | flagellar basal body rod protein FlgF | WP_000349278.1 | 0.213589798 | 0.1212901 | 0.19717625 |
| STM14_RS02355 | | DUF4156 domain-containing protein | WP_000722978.1 | 0.213560534 | 0.53085553 | 0.63481977 |
| STM14_RS22825 | rsgA | small ribosomal subunit biogenesis GTPase RsgA | WP_000041945.1 | 0.21335649 | 0.10528905 | 0.17613552 |
| STM14_RS11610 | gndA | NADP-dependent phosphogluconate dehydrogenase | WP_000043530.1 | 0.212784354 | 0.01831873 | 0.03958308 |
| STM14_RS05910 | tusE | sulfurtransferase TusE | WP_000904446.1 | 0.212766841 | 0.20460173 | 0.30169946 |
| STM14_RS19855 | misL | intestinal colonization autotransporter adhesin MisL | WP_001143053.1 | 0.212655928 | 0.40386465 | 0.51472123 |
| STM14_RS08010 | sodB | superoxide dismutase [Fe] | WP_000007301.1 | 0.211395601 | 0.01082055 | 0.02554142 |
| STM14_RS01565 | | fimbrial biogenesis outer membrane usher protein | WP_000951687.1 | 0.211114572 | 0.51982839 | 0.62488023 |
| STM14_RS09960 | minC | septum site-determining protein MinC | WP_000072527.1 | 0.211005659 | 0.06882504 | 0.12295672 |
| STM14_RS11945 | rcnB | Ni(II)/Co(II) efflux transporter accessory subunit RcnB | WP_000837534.1 | 0.210086932 | 0.23103264 | 0.33234948 |
| STM14_RS02790 | panE | 2-dehydropantoate 2-reductase | WP_000705816.1 | 0.20921507 | 0.11661886 | 0.19135055 |
| STM14_RS20305 | | retron St85 family effector protein | WP_000461381.1 | 0.208980378 | 0.2882985 | 0.39564483 |
| STM14_RS08730 | yddG | aromatic amino acid efflux DMT transporter YddG | WP_000203799.1 | 0.208693553 | 0.40533291 | 0.51641376 |
| STM14_RS19635 | rfaL | O-antigen ligase RfaL | WP_000958465.1 | 0.20863046 | 0.12169526 | 0.19774137 |
| STM14_RS05065 | | sulfatase | WP_000644017.1 | 0.208615394 | 0.37960084 | 0.49076204 |
| STM14_RS21020 | mobB | molybdopterin-guanine dinucleotide biosynthesis protein MobB | WP_010989085.1 | 0.20767286 | 0.219475 | 0.31840305 |
| STM14_RS16960 | qseB | two-component system response regulator QseB | WP_001221574.1 | 0.207090671 | 0.38722309 | 0.49816751 |
| STM14_RS07860 | sseD | SPI-2 type III secretion system translocon protein SseD | WP_000388323.1 | 0.206678249 | 0.22203099 | 0.32198428 |
| STM14_RS19320 | cueP | copper-binding periplasmic metallochaperone CueP | WP_000047149.1 | 0.206511323 | 0.257601 | 0.36215612 |
| STM14_RS08135 | rsxD | electron transport complex subunit RsxD | WP_000231891.1 | 0.205952428 | 0.50205797 | 0.60968913 |
| STM14_RS06320 | | lipoprotein | WP_000713057.1 | 0.205901094 | 0.03272272 | 0.06456863 |
| STM14_RS15635 | ygbI | DNA-binding transcriptional repressor YgbI | WP_000613185.1 | 0.205471703 | 0.25426914 | 0.3585672 |
| STM14_RS10480 | flhB | flagellar biosynthesis protein FlhB | WP_000797081.1 | 0.20543613 | 0.54783473 | 0.65027514 |
| STM14_RS22360 | siiC | SPI-4 type I secretion system protein SiiC | WP_001541306.1 | 0.205106098 | 0.55233894 | 0.65540186 |
| STM14_RS09580 | yciA | acyl-CoA thioester hydrolase YciA | WP_000210317.1 | 0.204913648 | 0.34359773 | 0.45459103 |
| STM14_RS07995 | punC | purine nucleoside transporter PunC | WP_001182309.1 | 0.204008226 | 0.5769011 | 0.6777294 |
| STM14_RS08325 | osmY | osmoprotectant ABC transporter permease OsmY | WP_000557304.1 | 0.203898944 | 0.26461045 | 0.37045463 |
| STM14_RS08370 | | DUF1283 family protein | WP_001066440.1 | 0.203859366 | 0.03618204 | 0.07048797 |
| STM14_RS12500 | napH | quinol dehydrogenase ferredoxin subunit NapH | WP_000013529.1 | 0.203356828 | 0.24152593 | 0.34362383 |
| STM14_RS06385 | flgE | flagellar hook protein FlgE | WP_000010567.1 | 0.2033299 | 0.06636058 | 0.11907391 |
| STM14_RS10645 | uvrY | UvrY/SirA/GacA family response regulator transcription factor | WP_000611323.1 | 0.203220265 | 0.05254992 | 0.09752459 |
| STM14_RS19330 | | DUF1778 domain-containing protein | WP_000965886.1 | 0.202946257 | 0.28506284 | 0.39237434 |
| STM14_RS01785 | arfB | alternative ribosome rescue aminoacyl-tRNA hydrolase ArfB | WP_000560527.1 | 0.201607092 | 0.2596804 | 0.3646618 |
| STM14_RS22265 | | CsbD family protein | WP_001030592.1 | 0.201241667 | 0.02027428 | 0.04320018 |
| STM14_RS20650 | rho | transcription termination factor Rho | WP_001054532.1 | 0.201118306 | 0.03362115 | 0.06595262 |
| STM14_RS05785 | zapC | cell division protein ZapC | WP_001220671.1 | 0.198969414 | 0.05389897 | 0.09972664 |
| STM14_RS05110 | | NAD(P)-dependent oxidoreductase | WP_000866904.1 | 0.198593653 | 0.234691 | 0.33663118 |
| STM14_RS16710 | | NAD-dependent phenylacetaldehyde dehydrogenase | WP_001286388.1 | 0.198260539 | 0.36234574 | 0.47277003 |
| STM14_RS12420 | sspH2 | SPI-2 type III secretion system effector E3 ubiquitin transferase SspH2 | WP_001115840.1 | 0.197958138 | 0.27830821 | 0.38509238 |
| STM14_RS09740 | chaB | putative cation transport regulator ChaB | WP_001526441.1 | 0.197713841 | 0.13592443 | 0.2166553 |
| STM14_RS20235 | | MFS transporter | WP_000061259.1 | 0.197687919 | 0.6298992 | 0.72545789 |
| STM14_RS20520 | | FadR/GntR family transcriptional regulator | WP_001131148.1 | 0.197650361 | 0.13606376 | 0.21678354 |
| STM14_RS16970 | | NAD(P)H-dependent oxidoreductase | WP_000065416.1 | 0.197553289 | 0.34210321 | 0.45359165 |
| STM14_RS06610 | cobB | NAD-dependent protein deacylase | WP_001191856.1 | 0.197444017 | 0.11341227 | 0.1872574 |
| STM14_RS16455 | | alpha/beta hydrolase | WP_014344408.1 | 0.195678807 | 0.37000341 | 0.48068633 |
| STM14_RS26665 | | hypothetical protein | | 0.195159473 | 0.27418951 | 0.38053741 |
| STM14_RS08650 | nhaC | Na+/H+ antiporter NhaC | WP_001276616.1 | 0.194886678 | 0.50978957 | 0.61718167 |
| STM14_RS01590 | stfH | fimbrial adhesin StfH | WP_000691870.1 | 0.194790167 | 0.27194946 | 0.37799846 |
| STM14_RS02115 | | polysaccharide deacetylase family protein | WP_000119390.1 | 0.194590815 | 0.58310519 | 0.6833206 |
| STM14_RS20515 | mdtD | multidrug transporter subunit MdtD | WP_000202071.1 | 0.194418209 | 0.36581952 | 0.47628977 |
| STM14_RS22815 | mscM | miniconductance mechanosensitive channel MscM | WP_001236755.1 | 0.194365652 | 0.2135569 | 0.31153586 |
| STM14_RS21645 | | PTS fructose-like transporter subunit IIB | WP_000290522.1 | 0.194327514 | 0.3186444 | 0.42960405 |
| STM14_RS05075 | artM | arginine ABC transporter permease ArtM | WP_000895397.1 | 0.192799269 | 0.20271944 | 0.29951723 |
| STM14_RS05300 | pflA | pyruvate formate lyase 1-activating protein | WP_000067973.1 | 0.192763281 | 0.09270536 | 0.1583215 |
| STM14_RS15305 | hycA | formate hydrogenlyase regulator HycA | WP_000158074.1 | 0.192702845 | 0.31150099 | 0.42136173 |
| STM14_RS02045 | | DUF6531 domain-containing protein | WP_000272656.1 | 0.192548497 | 0.25541455 | 0.35986357 |
| STM14_RS21655 | | phosphoethanolamine transferase CptA | WP_001192092.1 | 0.191875276 | 0.11895951 | 0.19449774 |
| STM14_RS21475 | | YiiQ family protein | WP_000802242.1 | 0.191355909 | 0.43721273 | 0.54806886 |
| STM14_RS07425 | chbC | PTS N%2CN'-diacetylchitobiose transporter subunit IIC | WP_000073064.1 | 0.191269411 | 0.26894449 | 0.3748416 |
| STM14_RS09170 | | methyl-accepting chemotaxis protein | WP_000906725.1 | 0.189396187 | 0.47884103 | 0.58750172 |
| STM14_RS19175 | bcsG | cellulose biosynthesis protein BcsG | WP_000192026.1 | 0.189301018 | 0.2330948 | 0.33499417 |
| STM14_RS16020 | recC | exodeoxyribonuclease V subunit gamma | WP_000946827.1 | 0.188023507 | 0.17000031 | 0.25940371 |
| STM14_RS01235 | ftsQ | cell division protein FtsQ | WP_000075728.1 | 0.187640225 | 0.12866043 | 0.206868 |

| | | | | | | |
|---|---|---|---|---|---|---|
| STM14_RS02885 | | helix-hairpin-helix domain-containing protein | WP_000680322.1 | 0.187273116 | 0.31665647 | 0.42723676 |
| STM14_RS14090 | | oxidoreductase | WP_000684022.1 | 0.185859506 | 0.502264 | 0.60973823 |
| STM14_RS07295 | | anaerobic sulfatase maturase | WP_000029246.1 | 0.184770482 | 0.59589391 | 0.6948327 |
| STM14_RS12560 | rcsB | response regulator transcription factor RcsB | WP_001061919.1 | 0.184662695 | 0.07407611 | 0.13106594 |
| STM14_RS02995 | acrA | multidrug efflux RND transporter periplasmic adaptor subunit AcrA | WP_001039202.1 | 0.183977132 | 0.06716646 | 0.12028547 |
| STM14_RS07185 | | MerR family transcriptional regulator | WP_000190263.1 | 0.182308892 | 0.32886057 | 0.44047457 |
| STM14_RS07445 | chbG | chitin disaccharide deacetylase | WP_000442738.1 | 0.1822605 | 0.39594848 | 0.50656092 |
| STM14_RS09750 | kdsA | 3-deoxy-8-phosphooctulonate synthase | WP_000811046.1 | 0.182230731 | 0.2456542 | 0.34842017 |
| STM14_RS03795 | rlpA | endolytic peptidoglycan transglycosylase RlpA | WP_001231300.1 | 0.181772018 | 0.28092125 | 0.38812459 |
| STM14_RS12430 | | tail fiber protein | WP_000624225.1 | 0.181642493 | 0.56290368 | 0.6653648 |
| STM14_RS07275 | dgcJ | diguanylate cyclase DgcJ | WP_001048646.1 | 0.180308733 | 0.23782074 | 0.33986505 |
| STM14_RS15790 | | MBL fold metallo-hydrolase | WP_001208013.1 | 0.180212909 | 0.28619646 | 0.39334654 |
| STM14_RS06550 | nagZ | beta-N-acetylhexosaminidase | WP_000529340.1 | 0.179935866 | 0.11718581 | 0.1920241 |
| STM14_RS08350 | | dimethyl sulfoxide reductase subunit A | WP_000982628.1 | 0.179828281 | 0.18654007 | 0.27943065 |
| STM14_RS16310 | bglA | 6-phospho-beta-glucosidase BglA | WP_001230141.1 | 0.179717697 | 0.16029747 | 0.2482736 |
| STM14_RS02325 | | EAL domain-containing protein | WP_000868285.1 | 0.179623804 | 0.39376195 | 0.50486732 |
| STM14_RS18195 | rpsH | 30S ribosomal protein S8 | WP_000062611.1 | 0.179305821 | 0.05619895 | 0.1033073 |
| STM14_RS22295 | dnaB | replicative DNA helicase | WP_000918353.1 | 0.178725856 | 0.3222531 | 0.43320041 |
| STM14_RS10855 | yedA | drug/metabolite exporter YedA | WP_001212264.1 | 0.178127098 | 0.57027899 | 0.67257118 |
| STM14_RS19350 | | YsaB family lipoprotein | WP_000605590.1 | 0.177771514 | 0.23768665 | 0.33986505 |
| STM14_RS04435 | | fumarate hydratase | WP_000086829.1 | 0.176374291 | 0.41033306 | 0.52116121 |
| STM14_RS15340 | | nitrous oxide-stimulated promoter family protein | WP_001117947.1 | 0.175699288 | 0.64667081 | 0.73854901 |
| STM14_RS18915 | rsmD | 16S rRNA (guanine(966)-N(2))-methyltransferase | WP_000743277.1 | 0.174992461 | 0.54028176 | 0.6444177 |
| STM14_RS11650 | | CDP-abequose synthase | WP_000143399.1 | 0.174461363 | 0.3399654 | 0.45147813 |
| STM14_RS11680 | rfbA | glucose-1-phosphate thymidylyltransferase RfbA | WP_000857529.1 | 0.174274143 | 0.08970048 | 0.15411907 |
| STM14_RS12520 | napF | ferredoxin-type protein NapF | WP_000228070.1 | 0.174067656 | 0.58984736 | 0.68924722 |
| STM14_RS04080 | fldA | flavodoxin FldA | WP_001018621.1 | 0.173373968 | 0.08383942 | 0.14574917 |
| STM14_RS13260 | ypeC | DUF2502 domain-containing protein YpeC | WP_000490091.1 | 0.173276343 | 0.24564483 | 0.34842017 |
| STM14_RS10050 | manY | PTS mannose transporter subunit IIC | WP_000406918.1 | 0.173059476 | 0.10537319 | 0.17619481 |
| STM14_RS17005 | tolC | outer membrane channel protein TolC | WP_014343899.1 | 0.172715347 | 0.06274187 | 0.11368875 |
| STM14_RS19650 | rfaY | lipopolysaccharide core heptose(II) kinase RfaY | WP_000582809.1 | 0.172491586 | 0.33625226 | 0.44790069 |
| STM14_RS14595 | aroF | 3-deoxy-7-phosphoheptulonate synthase AroF | WP_001168062.1 | 0.172002209 | 0.21006907 | 0.30788443 |
| STM14_RS13030 | cvpA | colicin V production protein | WP_000262100.1 | 0.17171532 | 0.29152706 | 0.39918283 |
| STM14_RS18480 | rpe | ribulose-phosphate 3-epimerase | WP_000816284.1 | 0.171414197 | 0.23450712 | 0.33649852 |
| STM14_RS11045 | | lysis protein | WP_001050825.1 | 0.171283925 | 0.63660385 | 0.72933499 |
| STM14_RS09755 | sirB1 | invasion regulator SirB1 | WP_001257065.1 | 0.171007481 | 0.21377227 | 0.31172653 |
| STM14_RS10725 | amyA | alpha-amylase | WP_000795487.1 | 0.1708464 | 0.24845006 | 0.35197889 |
| STM14_RS14385 | | helix-turn-helix domain-containing protein | WP_000660736.1 | 0.17076022 | 0.1418618 | 0.22456369 |
| STM14_RS17940 | csrD | RNase E specificity factor CsrD | WP_001241377.1 | 0.170748625 | 0.15794594 | 0.24538268 |
| STM14_RS18850 | | type II toxin-antitoxin system Phd/YefM family antitoxin | WP_001520924.1 | 0.170548076 | 0.51336513 | 0.62014778 |
| STM14_RS02665 | | VOC family protein | WP_000793035.1 | 0.170451096 | 0.38764794 | 0.49836583 |
| STM14_RS15810 | mazG | nucleoside triphosphate pyrophosphohydrolase | WP_000210449.1 | 0.169638739 | 0.1761236 | 0.26697698 |
| STM14_RS04045 | chiP | chitoporin | WP_001258803.1 | 0.16946432 | 0.18319059 | 0.2758723 |
| STM14_RS09555 | | manganese catalase family protein | WP_000488349.1 | 0.168979466 | 0.13408993 | 0.21447399 |
| STM14_RS19325 | | GNAT family N-acetyltransferase | WP_000533909.1 | 0.168772913 | 0.40598816 | 0.51706967 |
| STM14_RS21650 | | AraC family transcriptional regulator | WP_000274596.1 | 0.168006007 | 0.4604695 | 0.56970722 |
| STM14_RS05790 | | YcbX family protein | WP_000224069.1 | 0.167921234 | 0.66760222 | 0.75448477 |
| STM14_RS23365 | | transposase | WP_000252550.1 | 0.167378787 | 0.24276705 | 0.34498968 |
| STM14_RS21535 | cytR | DNA-binding transcriptional regulator CytR | WP_000841333.1 | 0.167287136 | 0.29849199 | 0.40735638 |
| STM14_RS19525 | sadA | trimeric autotransporter adhesin SadA | WP_001079584.1 | 0.16709414 | 0.25712315 | 0.3616224 |
| STM14_RS02635 | acpH | ACP phosphodiesterase | WP_001009858.1 | 0.166747258 | 0.07777862 | 0.13679006 |
| STM14_RS02765 | phnS | 2-aminoethylphosphonate ABC transporter substrate-binding protein | WP_000778012.1 | 0.166209702 | 0.64468727 | 0.73741489 |
| STM14_RS23730 | | BrnT family toxin | WP_001131963.1 | 0.166047088 | 0.17530689 | 0.26606759 |
| STM14_RS09915 | fadR | fatty acid metabolism transcriptional regulator FadR | WP_000234826.1 | 0.165908758 | 0.12403189 | 0.20091747 |
| STM14_RS16810 | hybB | Ni/Fe-hydrogenase cytochrome b subunit | WP_000017678.1 | 0.165332831 | 0.20800151 | 0.30548926 |
| STM14_RS18140 | rpoA | DNA-directed RNA polymerase subunit alpha | WP_001162094.1 | 0.165275355 | 0.07363732 | 0.1305405 |
| STM14_RS17125 | dnaG | DNA primase | WP_000918865.1 | 0.164971754 | 0.16038059 | 0.2482736 |
| STM14_RS20565 | | YifB family Mg chelatase-like AAA ATPase | WP_000050215.1 | 0.164405702 | 0.58466927 | 0.68471764 |
| STM14_RS03950 | ybeY | rRNA maturation RNase YbeY | WP_000084476.1 | 0.164210684 | 0.12485775 | 0.20181134 |
| STM14_RS13645 | | ArsC family reductase | WP_000631921.1 | 0.16416321 | 0.2934483 | 0.4010678 |
| STM14_RS19970 | | FGGY family carbohydrate kinase | WP_001113558.1 | 0.163839017 | 0.53315599 | 0.63695014 |
| STM14_RS02840 | ampG | muropeptide MFS transporter AmpG | WP_000098481.1 | 0.163792623 | 0.62694627 | 0.72341466 |
| STM14_RS22795 | frdA | fumarate reductase (quinol) flavoprotein subunit | WP_001192954.1 | 0.1637402 | 0.06462134 | 0.11663518 |
| STM14_RS05440 | aspC | aspartate transaminase | WP_000462653.1 | 0.16351711 | 0.13584213 | 0.2166179 |
| STM14_RS05345 | | M48 family metallopeptidase | WP_000792312.1 | 0.162731688 | 0.54102929 | 0.64496606 |
| STM14_RS11510 | sbmC | DNA gyrase inhibitor SbmC | WP_000384326.1 | 0.161241113 | 0.0833542 | 0.14504261 |
| STM14_RS16250 | lysS | lysine--tRNA ligase | WP_000003339.1 | 0.161034844 | 0.09680748 | 0.16388283 |
| STM14_RS08540 | cybH | Ni/Fe-hydrogenase%2C b-type cytochrome subunit | WP_000924456.1 | 0.160936103 | 0.3874984 | 0.49834758 |
| STM14_RS07420 | chbB | PTS N%2CN'-diacetylchitobiose transporter subunit IIB | WP_000412161.1 | 0.160676004 | 0.16657855 | 0.25545281 |
| STM14_RS14390 | | AAA family ATPase | WP_001111772.1 | 0.160284107 | 0.51493756 | 0.62068199 |
| STM14_RS05135 | hcp | hydroxylamine reductase | WP_000458775.1 | 0.160141004 | 0.33635128 | 0.44790069 |
| STM14_RS10795 | fliN | flagellar motor switch protein FliN | WP_001282115.1 | 0.160031086 | 0.52780622 | 0.6319943 |
| STM14_RS09335 | pspB | envelope stress response membrane protein PspB | WP_001274953.1 | 0.15938391 | 0.42768676 | 0.53818956 |
| STM14_RS03685 | rnk | nucleoside diphosphate kinase regulator | WP_000089748.1 | 0.159374979 | 0.39957473 | 0.51013667 |
| STM14_RS15980 | mltA | murein transglycosylase A | WP_000678626.1 | 0.158664067 | 0.23926188 | 0.34119374 |
| STM14_RS18725 | | IclR family transcriptional regulator | WP_000724553.1 | 0.15816458 | 0.21756738 | 0.31613382 |
| STM14_RS20350 | pstB | phosphate ABC transporter ATP-binding protein PstB | WP_000063118.1 | 0.158100349 | 0.342101 | 0.45359165 |
| STM14_RS09690 | | tetratricopeptide repeat protein | WP_001212712.1 | 0.157504993 | 0.20453764 | 0.30169946 |
| STM14_RS01820 | metN | methionine ABC transporter ATP-binding protein MetN | WP_000594021.1 | 0.157349047 | 0.30660839 | 0.41596614 |
| STM14_RS07315 | msrB | peptide-methionine (R)-S-oxide reductase MsrB | WP_001519539.1 | 0.155773381 | 0.10684371 | 0.1780084 |
| STM14_RS06695 | rluE | 23S rRNA pseudouridine(2457) synthase RluE | WP_001249412.1 | 0.15572992 | 0.35395263 | 0.46461804 |
| STM14_RS22575 | melB | melibiose:sodium transporter MelB | WP_000028075.1 | 0.155494218 | 0.63151492 | 0.72660017 |

| | | | | | | |
|---|---|---|---|---|---|---|
| STM14_RS10485 | cheZ | protein phosphatase CheZ | WP_000983586.1 | 0.15530907 | 0.12809196 | 0.20622414 |
| STM14_RS11675 | rfbC | dTDP-4-dehydrorhamnose 3%2C5-epimerase | WP_000973708.1 | 0.154324028 | 0.16875605 | 0.25793266 |
| STM14_RS21230 | fdoI | formate dehydrogenase cytochrome b556 subunit | WP_000829025.1 | 0.1541916 | 0.18474466 | 0.27723073 |
| STM14_RS08040 | | TetR/AcrR family transcriptional regulator | WP_001041821.1 | 0.153892913 | 0.30835004 | 0.41786708 |
| STM14_RS20090 | | radical SAM protein | WP_000147832.1 | 0.152945073 | 0.32992485 | 0.44141834 |
| STM14_RS20040 | ilvN | acetolactate synthase small subunit | WP_001171826.1 | 0.1519018 | 0.59594831 | 0.6948327 |
| STM14_RS23595 | | membrane protein | WP_001240360.1 | 0.151826202 | 0.2717236 | 0.37782715 |
| STM14_RS22750 | efp | elongation factor P | WP_000257282.1 | 0.151511787 | 0.12692073 | 0.20460689 |
| STM14_RS06975 | | host specificity protein J | WP_000514918.1 | 0.151469091 | 0.47071381 | 0.57959561 |
| STM14_RS05445 | ompF | porin OmpF | WP_000977713.1 | 0.150495735 | 0.21290761 | 0.31083788 |
| STM14_RS12270 | | phosphatase PAP2 family protein | WP_000178095.1 | 0.150471371 | 0.36133938 | 0.4716241 |
| STM14_RS02400 | | site-specific DNA-methyltransferase | WP_000910371.1 | 0.150263377 | 0.23421354 | 0.33620829 |
| STM14_RS05100 | | heavy metal-binding domain-containing protein | WP_001160725.1 | 0.148977465 | 0.15050976 | 0.23610687 |
| STM14_RS18260 | rplD | 50S ribosomal protein L4 | WP_000424395.1 | 0.148758204 | 0.17349103 | 0.26374648 |
| STM14_RS13205 | pgtC | phosphoglycerate transport regulator PgtC | WP_000467983.1 | 0.148596774 | 0.32869493 | 0.44041292 |
| STM14_RS22565 | melR | transcriptional regulator MelR | WP_000106992.1 | 0.146884044 | 0.28114995 | 0.38829487 |
| STM14_RS07805 | | MerR family transcriptional regulator | WP_000122327.1 | 0.145976079 | 0.53983553 | 0.64430289 |
| STM14_RS11850 | | MFS transporter | WP_000137816.1 | 0.145238183 | 0.72537403 | 0.80311665 |
| STM14_RS13395 | ptsI | phosphoenolpyruvate-protein phosphotransferase PtsI | WP_000623114.1 | 0.144884567 | 0.08183299 | 0.14259775 |
| STM14_RS19835 | | AAA family ATPase | WP_001541628.1 | 0.14475904 | 0.25212864 | 0.35595768 |
| STM14_RS20845 | pldA | phospholipase A | WP_001201692.1 | 0.144588965 | 0.18408447 | 0.27687869 |
| STM14_RS20780 | cyaY | iron donor protein CyaY | WP_000999933.1 | 0.143759992 | 0.31493669 | 0.42554014 |
| STM14_RS18490 | damX | cell division protein DamX | WP_000343156.1 | 0.143751833 | 0.19229753 | 0.28666418 |
| STM14_RS17325 | | sugar kinase | WP_001027807.1 | 0.143314591 | 0.28752627 | 0.39487942 |
| STM14_RS16865 | exbB | tol-pal system-associated acyl-CoA thioesterase | WP_000527859.1 | 0.143260748 | 0.16110064 | 0.24902291 |
| STM14_RS03790 | dacA | D-alanyl-D-alanine carboxypeptidase DacA | WP_000858711.1 | 0.142976289 | 0.31429247 | 0.42482558 |
| STM14_RS26505 | | YshB family small membrane protein | WP_001578131.1 | 0.142865584 | 0.42692994 | 0.53742086 |
| STM14_RS09270 | | LysR family transcriptional regulator | WP_000359201.1 | 0.142547606 | 0.54748379 | 0.65026946 |
| STM14_RS23555 | brxC | BREX system P-loop protein BrxC | WP_001019652.1 | 0.142489575 | 0.17821819 | 0.26904444 |
| STM14_RS16515 | galP | galactose/proton symporter | WP_001113171.1 | 0.142307935 | 0.19171885 | 0.28602463 |
| STM14_RS06400 | flgH | flagellar basal body L-ring protein FlgH | WP_001174897.1 | 0.142110975 | 0.52276247 | 0.62758768 |
| STM14_RS14180 | | host specificity protein J | WP_000514917.1 | 0.141830334 | 0.53707017 | 0.64141822 |
| STM14_RS07985 | | riboflavin synthase | WP_000493971.1 | 0.141821413 | 0.2143557 | 0.31225532 |
| STM14_RS17450 | nlpI | lipoprotein NlpI | WP_000802069.1 | 0.141703071 | 0.09132551 | 0.15640024 |
| STM14_RS21690 | sthA | Si-specific NAD(P)(+) transhydrogenase | WP_001120789.1 | 0.141514809 | 0.13300574 | 0.21311016 |
| STM14_RS21970 | metA | homoserine O-succinyltransferase | WP_001122767.1 | 0.140798897 | 0.56462338 | 0.66696929 |
| STM14_RS22315 | | secondary thiamine-phosphate synthase enzyme YjbQ | WP_000270393.1 | 0.140399141 | 0.18727916 | 0.2803097 |
| STM14_RS12885 | | YfbU family protein | WP_000426135.1 | 0.140369741 | 0.19012887 | 0.28399777 |
| STM14_RS16745 | uxuA | mannonate dehydratase | WP_000815487.1 | 0.140150269 | 0.16530296 | 0.25391968 |
| STM14_RS09775 | hemA | glutamyl-tRNA reductase | WP_000173208.1 | 0.140141758 | 0.31088227 | 0.42083401 |
| STM14_RS22535 | pmrA | two-component system response regulator PmrA | WP_000697899.1 | 0.139751466 | 0.26238559 | 0.36775932 |
| STM14_RS11160 | | hypothetical protein | WP_000208068.1 | 0.139614069 | 0.57353901 | 0.67533439 |
| STM14_RS18605 | | Rpn family recombination-promoting nuclease/putative transposase | WP_000235945.1 | 0.13959391 | 0.42199886 | 0.53194105 |
| STM14_RS02175 | dinB | DNA polymerase IV | WP_001226198.1 | 0.137922169 | 0.57676582 | 0.6777294 |
| STM14_RS16880 | yghB | DedA family general envelope maintenance protein YghB | WP_000268441.1 | 0.137876543 | 0.39436313 | 0.50529341 |
| STM14_RS03450 | | mechanosensitive ion channel family protein | WP_001156978.1 | 0.137547243 | 0.31633633 | 0.42709471 |
| STM14_RS07415 | osmE | osmotically-inducible lipoprotein OsmE | WP_001039301.1 | 0.136821949 | 0.10576557 | 0.17661171 |
| STM14_RS19590 | gpmM | 2%2C3-bisphosphoglycerate-independent phosphoglycerate mutase | WP_000116576.1 | 0.135408557 | 0.24305091 | 0.34525982 |
| STM14_RS14435 | | site-specific integrase | WP_001007939.1 | 0.135087585 | 0.63721006 | 0.72976903 |
| STM14_RS05020 | | YbjN domain-containing protein | WP_000624814.1 | 0.134942906 | 0.56454284 | 0.66696929 |
| STM14_RS15835 | barA | two-component sensor histidine kinase BarA | WP_000186400.1 | 0.134520925 | 0.32943999 | 0.4410902 |
| STM14_RS23710 | | restriction endonuclease subunit S | WP_000014001.1 | 0.133920902 | 0.37490425 | 0.48588435 |
| STM14_RS11655 | rfbH | lipopolysaccharide biosynthesis protein RfbH | WP_000126349.1 | 0.133751237 | 0.13338498 | 0.21343915 |
| STM14_RS02705 | ribE | 6%2C7-dimethyl-8-ribityllumazine synthase | WP_001021372.1 | 0.133568856 | 0.20485619 | 0.30183293 |
| STM14_RS19345 | glyQ | glycine--tRNA ligase subunit alpha | WP_001168551.1 | 0.132473839 | 0.36696159 | 0.47743907 |
| STM14_RS01890 | gloB | hydroxyacylglutathione hydrolase | WP_001052775.1 | 0.132452703 | 0.51084988 | 0.61771732 |
| STM14_RS10435 | cmoA | carboxy-S-adenosyl-L-methionine synthase CmoA | WP_000019609.1 | 0.132247734 | 0.43890857 | 0.54893388 |
| STM14_RS13465 | yfeX | porphyrinogen peroxidase | WP_000084583.1 | 0.132038743 | 0.33966244 | 0.45133061 |
| STM14_RS10320 | yebG | DNA damage-inducible protein YebG | WP_000257723.1 | 0.132033133 | 0.42159002 | 0.53165354 |
| STM14_RS13875 | | dimethyl sulfoxide reductase subunit A | WP_000508279.1 | 0.131926274 | 0.44241791 | 0.5529473 |
| STM14_RS16140 | kduI | 5-dehydro-4-deoxy-D-glucuronate isomerase | WP_000383266.1 | 0.13153536 | 0.66408363 | 0.75189296 |
| STM14_RS01815 | | methionine ABC transporter permease MetI | WP_001287487.1 | 0.131398998 | 0.52629239 | 0.63079706 |
| STM14_RS16365 | serA | phosphoglycerate dehydrogenase | WP_001151627.1 | 0.13026216 | 0.22612215 | 0.32701562 |
| STM14_RS02445 | yahO | DUF1471 family periplasmic protein YahO | WP_000848026.1 | 0.129511341 | 0.16394504 | 0.25254224 |
| STM14_RS11570 | hisC | histidinol-phosphate transaminase | WP_000102713.1 | 0.129312825 | 0.57315901 | 0.67510284 |
| STM14_RS08940 | sgcB | PTS sugar transporter subunit IIB SgcB | WP_000976132.1 | 0.12897364 | 0.54784179 | 0.65027514 |
| STM14_RS18205 | rplE | 50S ribosomal protein L5 | WP_001096206.1 | 0.128690245 | 0.1476896 | 0.23258901 |
| STM14_RS12950 | yfcE | phosphodiesterase | WP_000772458.1 | 0.128586475 | 0.35222478 | 0.46267986 |
| STM14_RS02795 | | YajQ family cyclic di-GMP-binding protein | WP_001138913.1 | 0.128503716 | 0.24273437 | 0.34498968 |
| STM14_RS11815 | udk | uridine kinase | WP_000132082.1 | 0.128294858 | 0.47049524 | 0.57959561 |
| STM14_RS18235 | rpsC | 30S ribosomal protein S3 | WP_000529945.1 | 0.127894966 | 0.16278428 | 0.25099318 |
| STM14_RS26760 | ymiC | small membrane protein YmiC | WP_223148964.1 | 0.127406258 | 0.65849181 | 0.7473861 |
| STM14_RS12680 | | YfaZ family protein | WP_001670658.1 | 0.125145032 | 0.59547387 | 0.69471952 |
| STM14_RS23725 | | restriction endonuclease | WP_000290512.1 | 0.124488225 | 0.40354339 | 0.51466809 |
| STM14_RS19380 | | protein bax | WP_001575119.1 | 0.124261234 | 0.2528133 | 0.35678749 |
| STM14_RS06845 | | hypothetical protein | WP_223151767.1 | 0.123624631 | 0.58623329 | 0.68633099 |
| STM14_RS20640 | rhlB | ATP-dependent RNA helicase RhlB | WP_000047525.1 | 0.123383855 | 0.27116578 | 0.37719396 |
| STM14_RS07475 | | YniB family protein | WP_000222158.1 | 0.123162408 | 0.41272077 | 0.52329127 |
| STM14_RS10180 | | hypothetical protein | WP_000030934.1 | 0.123122478 | 0.44381188 | 0.55412524 |
| STM14_RS07485 | ghoS | type V toxin-antitoxin system endoribonuclease antitoxin GhoS | WP_000146127.1 | 0.122465625 | 0.39456339 | 0.50531562 |
| STM14_RS21125 | | STM4015 family protein | WP_001042093.1 | 0.121899343 | 0.74051719 | 0.81512236 |
| STM14_RS23325 | | type II toxin-antitoxin system RelE/ParE family toxin | WP_000220578.1 | 0.120926433 | 0.69921277 | 0.78297629 |

| | | | | | | |
|---|---|---|---|---|---|---|
| STM14_RS17375 | diaA | DnaA initiator-associating protein DiaA | WP_000893481.1 | 0.120850232 | 0.38393328 | 0.49514618 |
| STM14_RS17955 | msrQ | protein-methionine-sulfoxide reductase heme-binding subunit MsrQ | WP_001240053.1 | 0.120369316 | 0.66118564 | 0.74914329 |
| STM14_RS16500 | yqgB | acid stress response protein YqgB | WP_000729109.1 | 0.120257392 | 0.72680614 | 0.80361337 |
| STM14_RS06480 | fabG | 3-oxoacyl-ACP reductase FabG | WP_000007236.1 | 0.117677764 | 0.2418459 | 0.34394615 |
| STM14_RS20310 | | retron St85 family RNA-directed DNA polymerase | WP_000353754.1 | 0.11742525 | 0.69152879 | 0.77557386 |
| STM14_RS11110 | | YmfL family putative regulatory protein | WP_000509731.1 | 0.117041169 | 0.35127529 | 0.46187942 |
| STM14_RS23720 | hsdR | type I restriction-modification system endonuclease | WP_001043484.1 | 0.116972805 | 0.30333676 | 0.41285287 |
| STM14_RS10760 | fliG | flagellar motor switch protein FliG | WP_000067735.1 | 0.116557657 | 0.55368614 | 0.65657725 |
| STM14_RS09485 | sohB | protease SohB | WP_000422082.1 | 0.116403206 | 0.30503902 | 0.41437194 |
| STM14_RS14140 | lepA | translation elongation factor 4 | WP_000790154.1 | 0.116183318 | 0.26956475 | 0.37539236 |
| STM14_RS10880 | ompS1 | porin OmpS1 | WP_001080680.1 | 0.11600144 | 0.69912763 | 0.78297629 |
| STM14_RS23430 | miaE | tRNA isopentenyl-2-thiomethyl-A-37 hydroxylase MiaE | WP_000218459.1 | 0.115791419 | 0.6317008 | 0.72660017 |
| STM14_RS03680 | | glutathione-dependent formaldehyde dehydrogenase | WP_000646113.1 | 0.114859101 | 0.44878665 | 0.55863166 |
| STM14_RS01795 | proS | proline--tRNA ligase | WP_001260683.1 | 0.114377305 | 0.40998377 | 0.52110646 |
| STM14_RS20225 | dgoR | D-galactonate utilization transcriptional regulator DgoR | WP_000174317.1 | 0.114147933 | 0.56899001 | 0.67126602 |
| STM14_RS03755 | crcB | fluoride efflux transporter CrcB | WP_000939753.1 | 0.113855914 | 0.48965343 | 0.59797809 |
| STM14_RS12150 | mglA | galactose/methyl galactoside ABC transporter ATP-binding protein MglA | WP_000535907.1 | 0.113835872 | 0.43604954 | 0.54684415 |
| STM14_RS09455 | ribA | GTP cyclohydrolase II | WP_001176284.1 | 0.113820106 | 0.37230091 | 0.48319067 |
| STM14_RS07580 | ydiV | anti-FlhDC factor YdiV | WP_000562005.1 | 0.113796976 | 0.42790702 | 0.53828277 |
| STM14_RS14855 | | DEAD/DEAH box helicase | WP_000073810.1 | 0.113421958 | 0.69414781 | 0.77803721 |
| STM14_RS01710 | skp | molecular chaperone Skp | WP_000758966.1 | 0.112909141 | 0.31073012 | 0.42078275 |
| STM14_RS22460 | nrfC | cytochrome c nitrite reductase Fe-S protein | WP_001278030.1 | 0.112776345 | 0.72242019 | 0.80094885 |
| STM14_RS03900 | | J domain-containing protein | WP_001017750.1 | 0.112570265 | 0.72635542 | 0.80337659 |
| STM14_RS21705 | trmA | tRNA (uridine(54)-C5)-methyltransferase TrmA | WP_000186981.1 | 0.112092952 | 0.58692565 | 0.68670488 |
| STM14_RS11660 | rfbG | CDP-glucose 4%2C6-dehydratase | WP_000565905.1 | 0.111861714 | 0.22880326 | 0.33018742 |
| STM14_RS09440 | lapB | lipopolysaccharide assembly protein LapB | WP_000891378.1 | 0.111708908 | 0.4038137 | 0.51472123 |
| STM14_RS02565 | | AroM family protein | WP_001276437.1 | 0.111263433 | 0.43251449 | 0.54352163 |
| STM14_RS03940 | lnt | apolipoprotein N-acyltransferase | WP_001575804.1 | 0.111073474 | 0.53293286 | 0.63693081 |
| STM14_RS16305 | yqfB | N(4)-acetylcytidine aminohydrolase | WP_001182976.1 | 0.110810744 | 0.38914437 | 0.49994053 |
| STM14_RS14035 | glrR | two-component system response regulator GlrR | WP_000625591.1 | 0.110342341 | 0.39147958 | 0.50224595 |
| STM14_RS09785 | ispE | 4-(cytidine 5'-diphospho)-2-C-methyl-D-erythritol kinase | WP_000988246.1 | 0.108988195 | 0.36805391 | 0.4786911 |
| STM14_RS22280 | dusA | tRNA dihydrouridine(20/20a) synthase DusA | WP_001182237.1 | 0.108934809 | 0.36957332 | 0.48049751 |
| STM14_RS18495 | aroB | 3-dehydroquinate synthase | WP_000439828.1 | 0.108268319 | 0.34082732 | 0.45238868 |
| STM14_RS04485 | gpmA | 2%2C3-diphosphoglycerate-dependent phosphoglycerate mutase | WP_000301556.1 | 0.108118977 | 0.20398686 | 0.30103392 |
| STM14_RS09575 | | septation protein A | WP_000808682.1 | 0.108085341 | 0.61058407 | 0.70853153 |
| STM14_RS02250 | | MFS transporter | WP_012543325.1 | 0.107932387 | 0.86756563 | 0.91320087 |
| STM14_RS13935 | iscU | Fe-S cluster assembly scaffold IscU | WP_000331704.1 | 0.107618037 | 0.30473438 | 0.41418678 |
| STM14_RS06985 | | prophage tail fiber N-terminal domain-containing protein | WP_001144681.1 | 0.107606817 | 0.71128636 | 0.7907477 |
| STM14_RS13200 | pgtB | two-component system sensor histidine kinase PgtB | WP_000678496.1 | 0.106735791 | 0.43867669 | 0.5488303 |
| STM14_RS21820 | rpoB | DNA-directed RNA polymerase subunit beta | WP_000263106.1 | 0.106523493 | 0.20910355 | 0.30674074 |
| STM14_RS09360 | sapC | peptide ABC transporter permease SapC | WP_001146150.1 | 0.106047644 | 0.75036085 | 0.82178128 |
| STM14_RS20480 | kup | low affinity potassium transporter Kup | WP_000102338.1 | 0.105725189 | 0.46274017 | 0.57194001 |
| STM14_RS20485 | rbsD | D-ribose pyranase | WP_000715944.1 | 0.105168301 | 0.58156221 | 0.68194652 |
| STM14_RS18110 | trkA | Trk system potassium transporter TrkA | WP_000691374.1 | 0.104718651 | 0.52715152 | 0.63162118 |
| STM14_RS03625 | | ParB/RepB/Spo0J family partition protein | WP_001164756.1 | 0.104518902 | 0.51417075 | 0.62047696 |
| STM14_RS22025 | rluF | 23S rRNA pseudouridine(2604) synthase RluF | WP_000954611.1 | 0.103519381 | 0.44967863 | 0.55929762 |
| STM14_RS02540 | adrA | diguanylate cyclase AdrA | WP_000484086.1 | 0.103509752 | 0.51481911 | 0.62068199 |
| STM14_RS08490 | | DUF4186 domain-containing protein | WP_000992449.1 | 0.103441237 | 0.71034915 | 0.79042175 |
| STM14_RS08550 | | hydrogenase small subunit | WP_001194448.1 | 0.103222323 | 0.41835842 | 0.52898204 |
| STM14_RS22250 | lexA | transcriptional repressor LexA | WP_000646079.1 | 0.10274898 | 0.39622295 | 0.50673598 |
| STM14_RS16830 | | methyl-accepting chemotaxis protein | WP_001238434.1 | 0.102471004 | 0.47214132 | 0.58102418 |
| STM14_RS12665 | rhmD | L-rhamnonate dehydratase | WP_001526367.1 | 0.101983721 | 0.44552718 | 0.55570158 |
| STM14_RS08565 | | zinc-binding alcohol dehydrogenase family protein | WP_000864924.1 | 0.101904565 | 0.65538153 | 0.74501847 |
| STM14_RS11390 | pduF | propanediol diffusion facilitator PduF | WP_001000023.1 | 0.101435421 | 0.73434733 | 0.81028065 |
| STM14_RS23525 | | restriction endonuclease | WP_010989099.1 | 0.101252169 | 0.3358443 | 0.44770205 |
| STM14_RS17480 | nusA | transcription termination factor NusA | WP_001031038.1 | 0.101160315 | 0.28705369 | 0.39437749 |
| STM14_RS06085 | wrbA | NAD(P)H:quinone oxidoreductase | WP_001062899.1 | 0.101129559 | 0.20818504 | 0.30563689 |
| STM14_RS01885 | mltD | murein transglycosylase D | WP_000644706.1 | 0.100844302 | 0.27441827 | 0.38056801 |
| STM14_RS14615 | trmD | tRNA (guanosine(37)-N1)-methyltransferase TrmD | WP_000469804.1 | 0.100462789 | 0.29137169 | 0.39911852 |
| STM14_RS05080 | artQ | arginine ABC transporter permease ArtQ | WP_001001677.1 | 0.100406202 | 0.6019729 | 0.70074746 |
| STM14_RS10555 | otsB | trehalose-phosphatase | WP_000830111.1 | 0.100254616 | 0.38436857 | 0.49553399 |
| STM14_RS04730 | cecR | transcriptional regulator CecR | WP_001025254.1 | 0.100068044 | 0.393802 | 0.50486732 |
| STM14_RS14570 | bamD | outer membrane protein assembly factor BamD | WP_000197660.1 | 0.100066384 | 0.41133971 | 0.5222437 |
| STM14_RS04850 | | glycyl radical protein | WP_000192673.1 | 0.100057729 | 0.60367787 | 0.70228813 |
| STM14_RS01280 | zapD | cell division protein ZapD | WP_000557441.1 | 0.099867968 | 0.69648365 | 0.78041777 |
| STM14_RS23675 | mdtM | multidrug efflux MFS transporter MdtM | WP_001188918.1 | 0.098408857 | 0.63663298 | 0.72933499 |
| STM14_RS01170 | cra | catabolite repressor/activator | WP_000762406.1 | 0.09832673 | 0.35801964 | 0.46862009 |
| STM14_RS03160 | | porin | WP_000152869.1 | 0.097699013 | 0.59968794 | 0.69830835 |
| STM14_RS04095 | seqA | replication initiation negative regulator SeqA | WP_000848395.1 | 0.097196258 | 0.35733241 | 0.46788689 |
| STM14_RS23015 | ulaE | L-ribulose-5-phosphate 3-epimerase UlaE | WP_000949525.1 | 0.096745785 | 0.80937088 | 0.87018371 |
| STM14_RS03070 | gsk | inosine/guanosine kinase | WP_014343969.1 | 0.096702536 | 0.45709867 | 0.56687009 |
| STM14_RS18325 | | transcriptional regulator | WP_000091471.1 | 0.09626087 | 0.46027563 | 0.56965878 |
| STM14_RS13245 | | ion channel protein | WP_000468920.1 | 0.096079937 | 0.73779891 | 0.81310254 |
| STM14_RS09895 | | potassium/proton antiporter | WP_000338376.1 | 0.095291194 | 0.45895537 | 0.56880407 |
| STM14_RS03870 | | sigma-54-dependent Fis family transcriptional regulator | WP_001044182.1 | 0.094177082 | 0.43484016 | 0.54574815 |
| STM14_RS20945 | rfaH | transcription/translation regulatory transformer protein RfaH | WP_001192410.1 | 0.094038256 | 0.51233807 | 0.61931345 |
| STM14_RS07115 | | exo-alpha-sialidase | WP_000761746.1 | 0.093780715 | 0.66688579 | 0.75413804 |
| STM14_RS05455 | | Lrp/AsnC family transcriptional regulator | WP_000762342.1 | 0.09349584 | 0.65974331 | 0.74835948 |
| STM14_RS04975 | | Cof-type HAD-IIB family hydrolase | WP_000023547.1 | 0.093489067 | 0.58335246 | 0.68339286 |
| STM14_RS20430 | atpE | F0F1 ATP synthase subunit C | WP_000429386.1 | 0.092870567 | 0.36387499 | 0.47442908 |
| STM14_RS04995 | | inner membrane protein YbjM | WP_000680850.1 | 0.092616504 | 0.7860854 | 0.8517853 |
| STM14_RS06330 | mdtH | multidrug efflux MFS transporter MdtH | WP_000092178.1 | 0.092106133 | 0.75501593 | 0.82614224 |

| | | | | | | |
|---|---|---|---|---|---|---|
| STM14_RS19660 | waaO | lipopolysaccharide 3-alpha-galactosyltransferase | WP_000088479.1 | 0.092091536 | 0.51955262 | 0.62475269 |
| STM14_RS02975 | maa | maltose O-acetyltransferase | WP_001278792.1 | 0.091493994 | 0.49106979 | 0.59950894 |
| STM14_RS06700 | icd | NADP-dependent isocitrate dehydrogenase | WP_000444507.1 | 0.091486256 | 0.27429957 | 0.38054673 |
| STM14_RS22970 | yjfN | DUF1471 family protease activator YjfN | WP_001518964.1 | 0.091458148 | 0.40632712 | 0.51732242 |
| STM14_RS17020 | ygiD | 4%2C5-DOPA dioxygenase extradiol | WP_001520560.1 | 0.091006261 | 0.61633868 | 0.71363491 |
| STM14_RS22520 | proP | glycine betaine/L-proline transporter ProP | WP_000919710.1 | 0.090509614 | 0.41751931 | 0.5281024 |
| STM14_RS11010 | | phage portal protein | WP_000466254.1 | 0.09050889 | 0.67030192 | 0.75637501 |
| STM14_RS09665 | rssA | patatin-like phospholipase RssA | WP_001230572.1 | 0.09029364 | 0.43335943 | 0.54439762 |
| STM14_RS08120 | nth | endonuclease III | WP_001030318.1 | 0.089513941 | 0.73435811 | 0.81028065 |
| STM14_RS17625 | | calcium/sodium antiporter | WP_000922736.1 | 0.089487658 | 0.77928971 | 0.84566599 |
| STM14_RS05265 | serS | serine--tRNA ligase | WP_000886697.1 | 0.088573579 | 0.34860719 | 0.45898435 |
| STM14_RS06325 | grxB | glutaredoxin 2 | WP_000780893.1 | 0.088495839 | 0.40322101 | 0.51443512 |
| STM14_RS21390 | | CDP-diacylglycerol diphosphatase | WP_000750761.1 | 0.088088796 | 0.52071105 | 0.62553281 |
| STM14_RS07440 | | 6-phospho-beta-glucosidase | WP_000078785.1 | 0.087693319 | 0.6690463 | 0.75542118 |
| STM14_RS01315 | ampD | 1%2C6-anhydro-N-acetylmuramyl-L-alanine amidase AmpD | WP_000936329.1 | 0.087656032 | 0.59200028 | 0.69132415 |
| STM14_RS11605 | ugd | UDP-glucose 6-dehydrogenase | WP_000704831.1 | 0.087645573 | 0.50254674 | 0.60983531 |
| STM14_RS04555 | | putative acyl-CoA thioester hydrolase | WP_001094775.1 | 0.086898591 | 0.61998039 | 0.71694969 |
| STM14_RS21190 | dtd | D-aminoacyl-tRNA deacylase | WP_000560974.1 | 0.086836706 | 0.78252442 | 0.84867596 |
| STM14_RS12490 | napC | cytochrome c-type protein NapC | WP_000431778.1 | 0.086801156 | 0.62779566 | 0.7241678 |
| STM14_RS09135 | uspF | universal stress protein UspF | WP_001082296.1 | 0.086588975 | 0.33147717 | 0.44301232 |
| STM14_RS07175 | aadA | cryptic aminoglycoside nucleotidyltransferase ANT(3'')/ANT(9) | WP_000175803.1 | 0.086511889 | 0.80434438 | 0.86697775 |
| STM14_RS21250 | fdhD | formate dehydrogenase accessory sulfurtransferase FdhD | WP_001059744.1 | 0.086388104 | 0.7059163 | 0.78750765 |
| STM14_RS15830 | rlmD | 23S rRNA (uracil(1939)-C(5))-methyltransferase RlmD | WP_000046849.1 | 0.086091446 | 0.44489884 | 0.55529408 |
| STM14_RS05155 | macA | macrolide transporter subunit MacA | WP_001201751.1 | 0.085960918 | 0.44372235 | 0.55412524 |
| STM14_RS04630 | yvcK | uridine diphosphate-N-acetylglucosamine-binding protein YvcK | WP_001246033.1 | 0.085891513 | 0.55871044 | 0.66210874 |
| STM14_RS12350 | yejM | LPS biosynthesis-modulating metalloenzyme YejM | WP_000256150.1 | 0.085162729 | 0.54378873 | 0.64685996 |
| STM14_RS13765 | | DUF1493 family protein | WP_001738968.1 | 0.085105127 | 0.79078833 | 0.85612544 |
| STM14_RS01625 | dgt | dGTPase | WP_000146443.1 | 0.084913927 | 0.45665085 | 0.56650554 |
| STM14_RS04480 | | ABC transporter ATP-binding protein | WP_001214710.1 | 0.084399251 | 0.86948539 | 0.91461211 |
| STM14_RS11075 | | KilA-N domain-containing protein | WP_001061457.1 | 0.084326124 | 0.79905942 | 0.86254377 |
| STM14_RS00200 | spvA | virulence protein SpvA | WP_001526990.1 | 0.084104256 | 0.63344564 | 0.72791032 |
| STM14_RS00015 | | phospholipase D family protein | | 0.083355732 | 0.78018124 | 0.84638401 |
| STM14_RS05415 | mukE | chromosome partition protein MukE | WP_000060025.1 | 0.083239547 | 0.64564145 | 0.73802443 |
| STM14_RS16675 | | arylsulfatase | WP_000712459.1 | 0.082513909 | 0.5967308 | 0.69552478 |
| STM14_RS19680 | | glycosyltransferase family 4 protein | WP_001261577.1 | 0.082461376 | 0.59147557 | 0.69093054 |
| STM14_RS13025 | purF | amidophosphoribosyltransferase | WP_000334204.1 | 0.082416323 | 0.54391049 | 0.64685996 |
| STM14_RS03110 | cueR | Cu(I)-responsive transcriptional regulator | WP_001026760.1 | 0.081575135 | 0.4873341 | 0.59613427 |
| STM14_RS21380 | pfkA | 6-phosphofructokinase | WP_000591793.1 | 0.081293831 | 0.39584521 | 0.50656092 |
| STM14_RS14655 | nadK | NAD(+) kinase | WP_001059155.1 | 0.080249082 | 0.55535618 | 0.65834561 |
| STM14_RS12300 | yejF | microcin C ABC transporter ATP-binding protein YejF | WP_000203622.1 | 0.079796726 | 0.79848683 | 0.86217844 |
| STM14_RS17545 | greA | transcription elongation factor GreA | WP_001148008.1 | 0.079070347 | 0.47082305 | 0.57959561 |
| STM14_RS20715 | | amino acid permease | WP_000818378.1 | 0.079040425 | 0.54386241 | 0.64685996 |
| STM14_RS18955 | | DcrB family lipoprotein | WP_001245277.1 | 0.07895017 | 0.48568223 | 0.59431106 |
| STM14_RS19260 | lpfC | outer membrane usher protein LpfC | WP_000220312.1 | 0.078649573 | 0.92785749 | 0.95965485 |
| STM14_RS13965 | asrA | anaerobic sulfite reductase subunit AsrA | WP_000985204.1 | 0.078517523 | 0.67140899 | 0.75739212 |
| STM14_RS01695 | cdsA | phosphatidate cytidylyltransferase | WP_000922422.1 | 0.078468027 | 0.5809828 | 0.68148412 |
| STM14_RS09840 | hyaE | hydrogenase-1 operon protein HyaE | WP_001262581.1 | 0.077971357 | 0.86183024 | 0.90950092 |
| STM14_RS22790 | frdB | fumarate reductase iron-sulfur protein | WP_000829509.1 | 0.077740445 | 0.41654041 | 0.52704529 |
| STM14_RS19495 | mtlA | PTS mannitol transporter subunit IICBA | WP_000093287.1 | 0.077090046 | 0.47275909 | 0.58159003 |
| STM14_RS15045 | | PLP-dependent aminotransferase family protein | WP_001138459.1 | 0.076763645 | 0.56227049 | 0.66504335 |
| STM14_RS01045 | lptD | LPS assembly protein LptD | WP_000746126.1 | 0.076287667 | 0.41373731 | 0.52421913 |
| STM14_RS18690 | glpD | glycerol-3-phosphate dehydrogenase | WP_000448179.1 | 0.075997561 | 0.67308572 | 0.75902567 |
| STM14_RS01215 | ftsW | cell division protein FtsW | WP_001239803.1 | 0.075347217 | 0.59308842 | 0.6921558 |
| STM14_RS12495 | napB | nitrate reductase cytochrome c-type subunit | WP_000835182.1 | 0.07508562 | 0.70614296 | 0.78750765 |
| STM14_RS07800 | mlrB | transcriptional regulator MlrB | WP_001060729.1 | 0.074632344 | 0.48533564 | 0.59408438 |
| STM14_RS11970 | | YehE family protein | WP_000760404.1 | 0.074572651 | 0.87320629 | 0.91651811 |
| STM14_RS10145 | rsmF | 16S rRNA (cytosine(1407)-C(5))-methyltransferase RsmF | WP_001542138.1 | 0.074328074 | 0.87068946 | 0.91475714 |
| STM14_RS11280 | ldtA | L%2CD-transpeptidase | WP_001252820.1 | 0.07424909 | 0.52925283 | 0.63331456 |
| STM14_RS09965 | minD | septum site-determining protein MinD | WP_000101047.1 | 0.07372726 | 0.44976756 | 0.55929762 |
| STM14_RS24020 | robA | MDR efflux pump AcrAB transcriptional activator RobA | WP_000371685.1 | 0.07338196 | 0.4819307 | 0.59070201 |
| STM14_RS14510 | pat | protein lysine acetyltransferase | WP_000082639.1 | 0.073317684 | 0.45927607 | 0.56880407 |
| STM14_RS15940 | rlmM | 23S rRNA (cytidine(2498)-2'-O)-methyltransferase RlmM | WP_001045494.1 | 0.07320495 | 0.57383238 | 0.67546382 |
| STM14_RS02190 | pepD | cytosol nonspecific dipeptidase | WP_001292977.1 | 0.072709542 | 0.48512625 | 0.59402556 |
| STM14_RS06270 | mdtG | multidrug efflux MFS transporter MdtG | WP_000075048.1 | 0.072690319 | 0.72466664 | 0.80271437 |
| STM14_RS03760 | | deaminated glutathione amidase | WP_000495780.1 | 0.072464471 | 0.73931325 | 0.81417252 |
| STM14_RS14445 | rseB | sigma-E factor regulatory protein RseB | WP_000812017.1 | 0.071957632 | 0.58663022 | 0.68657739 |
| STM14_RS08235 | | GlpM family protein | WP_000524877.1 | 0.071512493 | 0.86753054 | 0.91320087 |
| STM14_RS12125 | sanA | outer membrane permeability protein SanA | WP_000920081.1 | 0.071334571 | 0.70290203 | 0.78569681 |
| STM14_RS12040 | bglX | beta-glucosidase BglX | WP_000871560.1 | 0.071329104 | 0.62973122 | 0.72545789 |
| STM14_RS16605 | trmB | tRNA (guanosine(46)-N7)-methyltransferase TrmB | WP_000786895.1 | 0.070507785 | 0.57831207 | 0.67878389 |
| STM14_RS19055 | fraD | fructose-aspartate kinase | WP_000028429.1 | 0.070415927 | 0.66959392 | 0.75580773 |
| STM14_RS18220 | rpsQ | 30S ribosomal protein S17 | WP_000130101.1 | 0.069893133 | 0.57180629 | 0.6741565 |
| STM14_RS11635 | | glycosyltransferase family 1 protein | WP_001521643.1 | 0.069774988 | 0.7062331 | 0.78750765 |
| STM14_RS11980 | metG | methionine--tRNA ligase | WP_000195332.1 | 0.069348154 | 0.61433664 | 0.71198852 |
| STM14_RS08190 | manA | mannose-6-phosphate isomerase | WP_001170627.1 | 0.069283131 | 0.65105511 | 0.74170326 |
| STM14_RS06090 | | general stress protein | WP_001273663.1 | 0.069009738 | 0.46386391 | 0.57313654 |
| STM14_RS19750 | | LysR family transcriptional regulator | WP_001050870.1 | 0.068876529 | 0.82814393 | 0.88485953 |
| STM14_RS13760 | | YfgG family protein | WP_000075924.1 | 0.068800348 | 0.6187033 | 0.71592255 |
| STM14_RS03030 | apt | adenine phosphoribosyltransferase | WP_000127350.1 | 0.06855607 | 0.66378952 | 0.75179114 |
| STM14_RS00125 | ccdA | type II toxin-antitoxin system antitoxin CcdA | WP_000813641.1 | 0.068397286 | 0.62399931 | 0.72069179 |
| STM14_RS11670 | | CDP-6-deoxy-delta-3%2C4-glucoseen reductase | WP_000018226.1 | 0.068145724 | 0.50990403 | 0.61718167 |
| STM14_RS02550 | | YaiI/YqxD family protein | WP_000158137.1 | 0.067992312 | 0.85157369 | 0.90412178 |

| | | | | | | |
|---|---|---|---|---|---|---|
| STM14_RS18705 | | glycerol dehydrogenase | WP_000258838.1 | 0.067850637 | 0.82801419 | 0.88485953 |
| STM14_RS21420 | | cupin domain-containing protein | WP_001173080.1 | 0.067696135 | 0.64216798 | 0.7349899 |
| STM14_RS23415 | | YhcH/YjgK/YiaL family protein | WP_000583461.1 | 0.067546009 | 0.8251966 | 0.882735 |
| STM14_RS08715 | fdxH | formate dehydrogenase subunit beta | WP_000061599.1 | 0.067476073 | 0.88376768 | 0.92496663 |
| STM14_RS02645 | tgt | tRNA guanosine(34) transglycosylase Tgt | WP_000667305.1 | 0.066973789 | 0.59249338 | 0.69168061 |
| STM14_RS22885 | hfq | RNA chaperone Hfq | WP_001051875.1 | 0.066429586 | 0.44542463 | 0.55570158 |
| STM14_RS11165 | | 3'-5' exonuclease | WP_001061334.1 | 0.065774633 | 0.85494611 | 0.90593285 |
| STM14_RS16815 | hybA | hydrogenase 2 operon protein HybA | WP_001081721.1 | 0.065568563 | 0.53296682 | 0.63693081 |
| STM14_RS13375 | cysZ | sulfate transporter CysZ | WP_000255006.1 | 0.065037146 | 0.80535396 | 0.86755801 |
| STM14_RS05605 | gtgA | type III secretion system effector protease GtgA | WP_001141973.1 | 0.064851865 | 0.768843 | 0.83729072 |
| STM14_RS02860 | clpP | ATP-dependent Clp endopeptidase proteolytic subunit ClpP | WP_000122257.1 | 0.064417791 | 0.50663961 | 0.61403786 |
| STM14_RS01435 | hpt | hypoxanthine phosphoribosyltransferase | WP_000683342.1 | 0.063601486 | 0.65174194 | 0.74225605 |
| STM14_RS06730 | recT | recombination protein RecT | WP_001126029.1 | 0.063534534 | 0.94035805 | 0.96931645 |
| STM14_RS09205 | | helix-turn-helix domain-containing protein | WP_000163371.1 | 0.061965964 | 0.81521241 | 0.87389135 |
| STM14_RS21595 | metF | methylenetetrahydrofolate reductase | WP_000007508.1 | 0.061848249 | 0.74317883 | 0.81683118 |
| STM14_RS09175 | smrA | DNA endonuclease SmrA | WP_001046799.1 | 0.061496698 | 0.6996176 | 0.78297629 |
| STM14_RS08020 | grxD | monothiol glutaredoxin 4 | WP_000108176.1 | 0.060863647 | 0.52736216 | 0.63166801 |
| STM14_RS02730 | dxs | 1-deoxy-D-xylulose-5-phosphate synthase | WP_000006777.1 | 0.060799581 | 0.63187771 | 0.72660017 |
| STM14_RS08600 | | RidA family protein | WP_000194089.1 | 0.060739579 | 0.79204772 | 0.8566019 |
| STM14_RS17695 | arcB | aerobic respiration two-component sensor histidine kinase ArcB | WP_000809815.1 | 0.060702813 | 0.54549646 | 0.64811809 |
| STM14_RS18060 | | gamma carbonic anhydrase family protein | WP_001285640.1 | 0.06025519 | 0.68000636 | 0.76513384 |
| STM14_RS10455 | argS | arginine--tRNA ligase | WP_001025361.1 | 0.059850752 | 0.60873322 | 0.70727539 |
| STM14_RS05450 | asnS | asparagine--tRNA ligase | WP_000117870.1 | 0.059165792 | 0.55956824 | 0.66248562 |
| STM14_RS21145 | | alpha-glucosidase | WP_001087041.1 | 0.059058586 | 0.87204852 | 0.91556392 |
| STM14_RS14680 | smpB | SsrA-binding protein SmpB | WP_001518569.1 | 0.058685918 | 0.63057369 | 0.7260076 |
| STM14_RS19730 | slmA | nucleoid occlusion factor SlmA | WP_000818607.1 | 0.058118155 | 0.68688795 | 0.77130876 |
| STM14_RS05125 | poxB | ubiquinone-dependent pyruvate dehydrogenase | WP_000815313.1 | 0.058086314 | 0.66816149 | 0.75488512 |
| STM14_RS09735 | | DsrE/DsrF/TusD sulfur relay family protein | WP_001179794.1 | 0.05807142 | 0.69115508 | 0.77539092 |
| STM14_RS06490 | fabF | beta-ketoacyl-ACP synthase II | WP_000044664.1 | 0.057867148 | 0.58178608 | 0.68199183 |
| STM14_RS22910 | purA | adenylosuccinate synthase | WP_000527972.1 | 0.057725621 | 0.52458709 | 0.6295729 |
| STM14_RS17610 | mlaD | outer membrane lipid asymmetry maintenance protein MlaD | WP_001193591.1 | 0.057339696 | 0.64942556 | 0.74030493 |
| STM14_RS02685 | | nucleoside-specific channel-forming protein Tsx | WP_000752021.1 | 0.056632606 | 0.55895058 | 0.66218019 |
| STM14_RS19865 | | transcriptional regulator | WP_001278062.1 | 0.056383359 | 0.79462283 | 0.85825793 |
| STM14_RS21060 | yihA | ribosome biogenesis GTP-binding protein YihA/YsxC | WP_000183327.1 | 0.056246998 | 0.70507198 | 0.78692787 |
| STM14_RS09445 | | LapA family protein | WP_000876301.1 | 0.055634267 | 0.66524571 | 0.75251451 |
| STM14_RS20340 | yieH | 6-phosphogluconate phosphatase | WP_000078081.1 | 0.055462963 | 0.6903888 | 0.77476732 |
| STM14_RS13050 | | DedA family protein | WP_000364318.1 | 0.055267403 | 0.79105246 | 0.85615966 |
| STM14_RS15130 | | YqaA family protein | WP_001279499.1 | 0.054936848 | 0.67462155 | 0.76008462 |
| STM14_RS13840 | ispG | flavodoxin-dependent (E)-4-hydroxy-3-methylbut-2-enyl-diphosphate sy | WP_000551804.1 | 0.054670518 | 0.66110177 | 0.74914329 |
| STM14_RS02950 | | MGMT family protein | WP_000961285.1 | 0.054066694 | 0.82839999 | 0.88487634 |
| STM14_RS08355 | | DUF1161 domain-containing protein | WP_000975516.1 | 0.053804512 | 0.64504831 | 0.73759872 |
| STM14_RS18250 | rplB | 50S ribosomal protein L2 | WP_000301869.1 | 0.053276511 | 0.57471172 | 0.67628269 |
| STM14_RS19690 | waaA | lipid IV(A) 3-deoxy-D-manno-octulosonic acid transferase | WP_000891582.1 | 0.052951054 | 0.75854436 | 0.82937995 |
| STM14_RS08410 | ydfG | bifunctional NADP-dependent 3-hydroxy acid dehydrogenase/3-hydroxy | WP_000636564.1 | 0.052603255 | 0.73471344 | 0.81042987 |
| STM14_RS06545 | thiK | thiamine kinase | WP_001257342.1 | 0.051536651 | 0.72700601 | 0.80361337 |
| STM14_RS04965 | ybjG | undecaprenyl-diphosphate phosphatase | WP_001738110.1 | 0.051429159 | 0.88048948 | 0.92284722 |
| STM14_RS13020 | | sigma-54-dependent Fis family transcriptional regulator | WP_000168925.1 | 0.050801702 | 0.72604602 | 0.80327567 |
| STM14_RS09305 | tyrR | transcriptional regulator TyrR | WP_001235493.1 | 0.050703099 | 0.62934736 | 0.72527605 |
| STM14_RS06620 | potC | spermidine/putrescine ABC transporter permease PotC | WP_000580299.1 | 0.050694738 | 0.91019281 | 0.94510151 |
| STM14_RS06000 | hpaE | 5-carboxymethyl-2-hydroxymuconate semialdehyde dehydrogenase | WP_000723524.1 | 0.050025626 | 0.78287732 | 0.84880869 |
| STM14_RS08340 | dmsB | dimethylsulfoxide reductase subunit B | WP_000213064.1 | 0.049935793 | 0.88506418 | 0.92553431 |
| STM14_RS03910 | | hypothetical protein | WP_000498891.1 | 0.049855376 | 0.76423204 | 0.83343867 |
| STM14_RS00625 | dnaK | molecular chaperone DnaK | WP_000516125.1 | 0.049019592 | 0.58898166 | 0.68845412 |
| STM14_RS00285 | umuD | translesion error-prone DNA polymerase V autoproteolytic subunit | WP_000925627.1 | 0.048855732 | 0.94256665 | 0.97078504 |
| STM14_RS17245 | | DUF805 domain-containing protein | WP_000384139.1 | 0.047678243 | 0.70351575 | 0.78600564 |
| STM14_RS21195 | fabY | fatty acid biosynthesis protein FabY | WP_001518251.1 | 0.047196563 | 0.80615924 | 0.86789572 |
| STM14_RS05145 | | ATP-dependent endonuclease | WP_000599761.1 | 0.047083861 | 0.77442297 | 0.84179565 |
| STM14_RS18210 | rplX | 50S ribosomal protein L24 | WP_000729185.1 | 0.04597785 | 0.61990194 | 0.71694969 |
| STM14_RS18305 | rpsL | 30S ribosomal protein S12 | WP_000246815.1 | 0.045596119 | 0.6339273 | 0.72791032 |
| STM14_RS06520 | ptsG | PTS glucose transporter subunit IIBC | WP_000475705.1 | 0.045263647 | 0.64766122 | 0.73943833 |
| STM14_RS26455 | yrbN | protein YrbN | WP_010723222.1 | 0.044883062 | 0.87518267 | 0.91833075 |
| STM14_RS13230 | lpxP | kdo(2)-lipid IV(A) palmitoleoyltransferase | WP_000484423.1 | 0.044536665 | 0.86496255 | 0.91228648 |
| STM14_RS22725 | | co-chaperone GroES | WP_000027827.1 | 0.044500555 | 0.67677183 | 0.76180797 |
| STM14_RS12890 | yfbV | terminus macrodomain insulation protein YfbV | WP_000106617.1 | 0.044362038 | 0.76696606 | 0.83574106 |
| STM14_RS02625 | malZ | maltodextrin glucosidase | WP_000930727.1 | 0.044234083 | 0.84719295 | 0.90059412 |
| STM14_RS15750 | casB | type I-E CRISPR-associated protein Cse2/CasB | WP_000117946.1 | 0.044153093 | 0.85748863 | 0.9072624 |
| STM14_RS06280 | | rhodanese-related sulfurtransferase | WP_001144632.1 | 0.043799212 | 0.81032353 | 0.870365 |
| STM14_RS24705 | | isocitrate/isopropylmalate family dehydrogenase | | 0.043776431 | 1 | 1 |
| STM14_RS10360 | mepM | murein DD-endopeptidase MepM | WP_001184028.1 | 0.043543014 | 0.71485718 | 0.79351949 |
| STM14_RS05970 | hprR | response regulator transcription factor HprR | WP_000698205.1 | 0.04322981 | 0.84802547 | 0.90113414 |
| STM14_RS21410 | | GntR family transcriptional regulator | WP_001519915.1 | 0.043038116 | 0.83615984 | 0.89187154 |
| STM14_RS17725 | codB | cytosine permease | WP_000076266.1 | 0.042878986 | 0.9299297 | 0.96125804 |
| STM14_RS19725 | dut | dUTP diphosphatase | WP_000976078.1 | 0.042538841 | 0.91385187 | 0.94754675 |
| STM14_RS03745 | pagP | lipid IV(A) palmitoyltransferase PagP | WP_000289054.1 | 0.042336811 | 0.7842625 | 0.85006021 |
| STM14_RS13715 | | DnaA inactivator Hda | WP_061835882.1 | 0.042118011 | 0.80823043 | 0.86938488 |
| STM14_RS13040 | folC | bifunctional tetrahydrofolate synthase/dihydrofolate synthase | WP_000792275.1 | 0.041296659 | 0.76381077 | 0.83343867 |
| STM14_RS03295 | cysS | cysteine--tRNA ligase | WP_000912376.1 | 0.040732628 | 0.72451752 | 0.80271437 |
| STM14_RS17105 | folB | bifunctional dihydroneopterin aldolase/7%2C8-dihydroneopterin epime | WP_000355776.1 | 0.040427642 | 0.89959221 | 0.936736 |
| STM14_RS22830 | orn | oligoribonuclease | WP_001271546.1 | 0.040236607 | 0.74775564 | 0.82069471 |
| STM14_RS21070 | yihI | Der GTPase-activating protein YihI | WP_000743292.1 | 0.039211561 | 0.70823278 | 0.78878195 |
| STM14_RS02760 | phnT | 2-aminoethylphosphonate ABC transport system ATP-binding subunit P | WP_000928801.1 | 0.038685144 | 0.89633346 | 0.93387091 |
| STM14_RS12450 | | cytochrome c-type biogenesis protein CcmH | WP_001238333.1 | 0.038359083 | 0.92887048 | 0.96043277 |

| | | | | | | |
|---|---|---|---|---|---|---|
| STM14_RS26875 | ysgD | YsgD/CorL family protein | WP_212723238.1 | 0.038319161 | 1 | 1 |
| STM14_RS23640 | hypT | hypochlorite stress DNA-binding transcriptional regulator HypT | WP_000383509.1 | 0.038268889 | 0.80204946 | 0.86501058 |
| STM14_RS01750 | | chitinase | WP_000524168.1 | 0.038259342 | 0.86532178 | 0.91240401 |
| STM14_RS21405 | | ADP-ribosylglycohydrolase family protein | WP_000557889.1 | 0.037751945 | 0.80994747 | 0.87021494 |
| STM14_RS12340 | yejK | nucleoid-associated protein YejK | WP_000050806.1 | 0.03764981 | 0.74761854 | 0.82069471 |
| STM14_RS14455 | rpoE | RNA polymerase sigma factor RpoE | WP_000003307.1 | 0.037466817 | 0.70657455 | 0.78764986 |
| STM14_RS18170 | rplO | 50S ribosomal protein L15 | WP_001238917.1 | 0.036309884 | 0.6813303 | 0.76623646 |
| STM14_RS03960 | miaB | tRNA (N6-isopentenyl adenosine(37)-C2)-methylthiotransferase MiaB | WP_001519200.1 | 0.036158995 | 0.77247899 | 0.84047522 |
| STM14_RS14460 | nadB | L-aspartate oxidase | WP_000989177.1 | 0.036006402 | 1 | 1 |
| STM14_RS16410 | epd | erythrose-4-phosphate dehydrogenase | WP_000218338.1 | 0.035324263 | 0.75910115 | 0.82937995 |
| STM14_RS14610 | rplS | 50S ribosomal protein L19 | WP_000065257.1 | 0.035047733 | 0.717507 | 0.79622085 |
| STM14_RS19980 | | PTS sugar transporter subunit IIB | WP_000041396.1 | 0.034922929 | 0.94527441 | 0.97139279 |
| STM14_RS10960 | | phage tail assembly protein | | 0.034473426 | 0.9058772 | 0.94194856 |
| STM14_RS12705 | arnA | bifunctional UDP-4-amino-4-deoxy-L-arabinose formyltransferase/UDP | WP_000648776.1 | 0.03401138 | 0.77651298 | 0.84339846 |
| STM14_RS08430 | | YdeI family stress tolerance OB fold protein | WP_000776348.1 | 0.03391457 | 0.73943317 | 0.81417252 |
| STM14_RS20905 | rmuC | DNA recombination protein RmuC | WP_000355562.1 | 0.033146904 | 0.82807368 | 0.88485953 |
| STM14_RS15110 | emrB | multidrug efflux MFS transporter permease subunit EmrB | WP_001187300.1 | 0.033112821 | 0.95328645 | 0.97744381 |
| STM14_RS18295 | fusA | elongation factor G | WP_000124693.1 | 0.032861317 | 0.70457248 | 0.7868474 |
| STM14_RS04355 | tolR | colicin uptake protein TolR | WP_000124676.1 | 0.032780262 | 0.91149014 | 0.94591508 |
| STM14_RS13150 | sixA | phosphohistidine phosphatase SixA | WP_001195800.1 | 0.032405944 | 0.8063746 | 0.86789572 |
| STM14_RS08095 | pdxH | pyridoxamine 5'-phosphate oxidase | WP_001282334.1 | 0.032277605 | 0.85309637 | 0.90521638 |
| STM14_RS18660 | rtcB | RNA-splicing ligase RtcB | WP_001105521.1 | 0.031575102 | 0.81894048 | 0.87757242 |
| STM14_RS13995 | | 3-phenylpropionate MFS transporter | WP_001173729.1 | 0.031376234 | 1 | 1 |
| STM14_RS02695 | nrdR | transcriptional regulator NrdR | WP_000543533.1 | 0.030333124 | 0.85627614 | 0.90702208 |
| STM14_RS12135 | | NAD(P)-dependent oxidoreductase | WP_001136409.1 | 0.029720325 | 0.76982751 | 0.83786724 |
| STM14_RS24070 | | winged helix-turn-helix transcriptional regulator | WP_000811636.1 | 0.02891711 | 0.93929678 | 0.96849363 |
| STM14_RS11830 | yegD | molecular chaperone | WP_000469629.1 | 0.028782724 | 0.92759215 | 0.95964998 |
| STM14_RS05280 | | dimethyl sulfoxide reductase anchor subunit | WP_000534681.1 | 0.028718068 | 0.85122032 | 0.90400727 |
| STM14_RS04820 | ldtB | L%2CD-transpeptidase | WP_001056404.1 | 0.028114612 | 0.88261311 | 0.92428369 |
| STM14_RS23270 | cybC | cytochrome b562 | WP_001232231.1 | 0.027936549 | 0.83273925 | 0.88873795 |
| STM14_RS20115 | ibpA | heat shock chaperone IbpA | WP_001532742.1 | 0.027626719 | 0.82954358 | 0.88584091 |
| STM14_RS20920 | ubiB | ubiquinone biosynthesis regulatory protein kinase UbiB | WP_000187559.1 | 0.026906083 | 0.81158434 | 0.87121095 |
| STM14_RS14375 | | CII family transcriptional regulator | WP_010835408.1 | 0.026726296 | 1 | 1 |
| STM14_RS23125 | tamA | autotransporter assembly complex protein TamA | WP_001120231.1 | 0.026213245 | 0.85028483 | 0.90327431 |
| STM14_RS02745 | thiI | tRNA 4-thiouridine(8) synthase ThiI | WP_000668708.1 | 0.02571976 | 0.85187926 | 0.90418549 |
| STM14_RS14060 | mltF | membrane-bound lytic murein transglycosylase MltF | WP_000734258.1 | 0.025700674 | 0.95651693 | 0.97993749 |
| STM14_RS10335 | edd | phosphogluconate dehydratase | WP_001069113.1 | 0.024871524 | 0.87039346 | 0.91475714 |
| STM14_RS13010 | | APC family permease | WP_000072512.1 | 0.024763392 | 0.94726174 | 0.97234952 |
| STM14_RS03005 | mscK | mechanosensitive channel MscK | WP_000178249.1 | 0.02419351 | 0.87150762 | 0.91525701 |
| STM14_RS09450 | pgpB | phosphatidylglycerophosphatase B | WP_000948651.1 | 0.024011469 | 0.86989995 | 0.91461211 |
| STM14_RS22160 | | Mor transcription activator family protein | WP_000615248.1 | 0.02389208 | 0.84727269 | 0.90059412 |
| STM14_RS01705 | bamA | outer membrane protein assembly factor BamA | WP_001240935.1 | 0.023137258 | 0.7915357 | 0.85643093 |
| STM14_RS02615 | brnQ | branched-chain amino acid transporter carrier protein BrnQ | WP_000149625.1 | 0.023002547 | 0.90671777 | 0.94255642 |
| STM14_RS19755 | | HARLDQ motif MBL-fold protein | WP_001082981.1 | 0.02292494 | 1 | 1 |
| STM14_RS02495 | sbmA | peptide antibiotic transporter SbmA | WP_000477024.1 | 0.022893342 | 0.89801485 | 0.93535804 |
| STM14_RS23345 | treC | alpha%2Calpha-phosphotrehalase | WP_000155057.1 | 0.022139022 | 1 | 1 |
| STM14_RS13950 | trmJ | tRNA (cytosine(32)/uridine(32)-2'-O)-methyltransferase TrmJ | WP_000940032.1 | 0.021646648 | 0.88482995 | 0.92553431 |
| STM14_RS16170 | | nickel/cobalt efflux protein RcnA | WP_000503671.1 | 0.021516504 | 0.94445468 | 0.97109246 |
| STM14_RS05325 | | 30S ribosomal protein S12 methylthiotransferase accessory protein Yca | WP_000194823.1 | 0.020301714 | 0.90898707 | 0.94411577 |
| STM14_RS11525 | phsB | thiosulfate reductase electron transport protein PhsB | WP_001016236.1 | 0.019283639 | 0.89003723 | 0.92862484 |
| STM14_RS05160 | macB | macrolide ABC transporter ATP-binding protein/permease MacB | WP_000125877.1 | 0.017930026 | 0.88497707 | 0.92553431 |
| STM14_RS16650 | | VOC family protein | WP_000582675.1 | 0.01746748 | 0.93215728 | 0.96274982 |
| STM14_RS09805 | pth | aminoacyl-tRNA hydrolase | WP_000985595.1 | 0.017264161 | 0.95389737 | 0.97752578 |
| STM14_RS20385 | glmS | glutamine--fructose-6-phosphate transaminase (isomerizing) | WP_000334066.1 | 0.016946286 | 0.85966094 | 0.90888931 |
| STM14_RS19405 | yiaJ | IclR family transcriptional regulator YiaJ | WP_000081340.1 | 0.016445454 | 0.97858138 | 0.994243 |
| STM14_RS02050 | | hypothetical protein | WP_000932531.1 | 0.016366616 | 1 | 1 |
| STM14_RS07290 | | MipA/OmpV family protein | WP_000163762.1 | 0.015978276 | 0.86718115 | 0.91320087 |
| STM14_RS16000 | argA | amino-acid N-acetyltransferase | WP_000588964.1 | 0.015836458 | 0.97059658 | 0.98940659 |
| STM14_RS09185 | fnr | fumarate/nitrate reduction transcriptional regulator Fnr | WP_000611914.1 | 0.015762074 | 0.87078321 | 0.91475714 |
| STM14_RS01665 | rpsB | 30S ribosomal protein S2 | WP_000246886.1 | 0.01515725 | 0.8578374 | 0.90737067 |
| STM14_RS18095 | def | peptide deformylase | WP_000114987.1 | 0.015120222 | 0.91409182 | 0.94754675 |
| STM14_RS17495 | argG | argininosuccinate synthase | WP_000207654.1 | 0.014602225 | 0.97446075 | 0.99087669 |
| STM14_RS17990 | fis | DNA-binding transcriptional regulator Fis | WP_000462905.1 | 0.014515778 | 0.94089212 | 0.96959552 |
| STM14_RS08500 | | nuclear transport factor 2 family protein | WP_000513122.1 | 0.014380636 | 1 | 1 |
| STM14_RS19995 | | DUF1198 domain-containing protein | WP_001519640.1 | 0.014354287 | 1 | 1 |
| STM14_RS17330 | | DeoR/GlpR family DNA-binding transcription regulator | WP_001118363.1 | 0.014252769 | 0.94639563 | 0.97215496 |
| STM14_RS08115 | dtpA | dipeptide/tripeptide permease DtpA | WP_000100911.1 | 0.014101932 | 0.91873425 | 0.95155542 |
| STM14_RS16440 | tkt | transketolase | WP_000098655.1 | 0.013578773 | 0.88871536 | 0.92829786 |
| STM14_RS16885 | | AraC family transcriptional regulator | WP_001541751.1 | 0.012823509 | 0.94928612 | 0.97415594 |
| STM14_RS01335 | aroP | aromatic amino acid transporter AroP | WP_000969487.1 | 0.012797143 | 0.93025121 | 0.9613205 |
| STM14_RS21975 | aceB | malate synthase A | WP_001069372.1 | 0.012761589 | 0.93598746 | 0.96562226 |
| STM14_RS05355 | rpsA | 30S ribosomal protein S1 | WP_000140324.1 | 0.011724051 | 0.88941064 | 0.9283636 |
| STM14_RS01445 | | ABC transporter ATP-binding protein | WP_000150619.1 | 0.011635387 | 0.96775962 | 0.9870612 |
| STM14_RS21180 | yihX | glucose-1-phosphatase | WP_000965695.1 | 0.011617524 | 0.96391822 | 0.98505326 |
| STM14_RS15105 | emrA | multidrug efflux MFS transporter periplasmic adaptor subunit EmrA | WP_001275603.1 | 0.011212146 | 0.94495909 | 0.97133986 |
| STM14_RS16715 | iraD | anti-adapter protein IraD | WP_000988938.1 | 0.010927442 | 1 | 1 |
| STM14_RS13900 | sseB | enhanced serine sensitivity protein SseB | WP_001533245.1 | 0.010693159 | 0.97403076 | 0.99071306 |
| STM14_RS15695 | ftsB | cell division protein FtsB | WP_000517480.1 | 0.010647614 | 0.9538911 | 0.97752578 |
| STM14_RS20250 | gyrB | DNA topoisomerase (ATP-hydrolyzing) subunit B | WP_000072047.1 | 0.010537 | 0.91067241 | 0.9453329 |
| STM14_RS21545 | rpmE | 50S ribosomal protein L31 | WP_000715284.1 | 0.01031876 | 0.9161297 | 0.94939194 |
| STM14_RS06190 | | DUF1097 domain-containing protein | WP_001546652.1 | 0.009850552 | 0.96342916 | 0.98482681 |
| STM14_RS09415 | | hypothetical protein | WP_001279854.1 | 0.009847199 | 1 | 1 |

| | | | | | | |
|---|---|---|---|---|---|---|
| STM14_RS16265 | dsbC | bifunctional protein-disulfide isomerase/oxidoreductase DsbC | WP_000745625.1 | 0.00968543 | 0.94373983 | 0.97104225 |
| STM14_RS17355 | | DeoR/GlpR family DNA-binding transcription regulator | WP_001083896.1 | 0.009564095 | 0.94680817 | 0.97215496 |
| STM14_RS17670 | rapZ | RNase adapter RapZ | WP_000243749.1 | 0.009299953 | 0.96012337 | 0.98250426 |
| STM14_RS04050 | chiQ | ChiQ/YbfN family lipoprotein | WP_000722261.1 | 0.009204486 | 0.96690094 | 0.98700561 |
| STM14_RS24025 | creA | protein CreA | WP_000875811.1 | 0.008492016 | 0.94367384 | 0.97104225 |
| STM14_RS14745 | | DUF4435 domain-containing protein | WP_010989064.1 | 0.008404629 | 0.97969437 | 0.99482478 |
| STM14_RS10280 | copD | copper homeostasis membrane protein CopD | WP_000979702.1 | 0.008353381 | 1 | 1 |
| STM14_RS11185 | mtfA | DgsA anti-repressor MtfA | WP_000598920.1 | 0.008218491 | 0.96035656 | 0.98250426 |
| STM14_RS02720 | pgpA | phosphatidylglycerophosphatase A | WP_000154063.1 | 0.007987733 | 1 | 1 |
| STM14_RS08720 | fdnG | formate dehydrogenase-N subunit alpha | WP_010989019.1 | 0.007617252 | 0.97216295 | 0.99018091 |
| STM14_RS23760 | | sigma-54-dependent transcriptional regulator | WP_001224211.1 | 0.007354119 | 0.96305532 | 0.98471805 |
| STM14_RS15995 | amiC | N-acetylmuramoyl-L-alanine amidase AmiC | WP_000011919.1 | 0.006365322 | 1 | 1 |
| STM14_RS15900 | xni | flap endonuclease Xni | WP_001519360.1 | 0.006279899 | 1 | 1 |
| STM14_RS20270 | rpmH | 50S ribosomal protein L34 | WP_000831330.1 | 0.005252382 | 0.96519715 | 0.98581301 |
| STM14_RS17645 | lptA | lipopolysaccharide ABC transporter substrate-binding protein LptA | WP_000669770.1 | 0.004885561 | 0.97544019 | 0.99159878 |
| STM14_RS20445 | rsmG | 16S rRNA (guanine(527)-N(7))-methyltransferase RsmG | WP_001519938.1 | 0.004872624 | 1 | 1 |
| STM14_RS05825 | fabA | bifunctional 3-hydroxydecanoyl-ACP dehydratase/trans-2-decenoyl-AC | WP_000227928.1 | 0.003416075 | 0.98365465 | 0.99829559 |
| STM14_RS14580 | pheA | bifunctional chorismate mutase/prephenate dehydratase | WP_000200080.1 | 0.002945542 | 1 | 1 |
| STM14_RS21040 | dsbA | thiol:disulfide interchange protein DsbA | WP_000725364.1 | 0.002680419 | 0.98419756 | 0.99857134 |
| STM14_RS10570 | | non-heme ferritin-like protein | WP_000741707.1 | 0.002661362 | 0.97888207 | 0.99427414 |
| STM14_RS08920 | tehA | dicarboxylate transporter/tellurite-resistance protein TehA | WP_001244002.1 | 0.002598763 | 1 | 1 |
| STM14_RS17380 | dolP | division/outer membrane stress-associated lipid-binding lipoprotein | WP_000643280.1 | 0.001698921 | 1 | 1 |
| STM14_RS05260 | rarA | replication-associated recombination protein RarA | WP_000067785.1 | 0.001223489 | 1 | 1 |
| STM14_RS10405 | | YebC/PmpR family DNA-binding transcriptional regulator | WP_000907242.1 | 0.000215639 | 1 | 1 |
| STM14_RS19810 | | AsmA family protein | WP_000766669.1 | 0.000122958 | 1 | 1 |
| STM14_RS05350 | cmk | (d)CMP kinase | WP_000125006.1 | -0.000160466 | 1 | 1 |
| STM14_RS03130 | fetB | iron export ABC transporter permease subunit FetB | WP_001002130.1 | -0.000773037 | 1 | 1 |
| STM14_RS06840 | | hypothetical protein | WP_000658040.1 | -0.001040712 | 1 | 1 |
| STM14_RS21085 | glnL | nitrogen regulation protein NR(II) | WP_000146194.1 | -0.001126769 | 1 | 1 |
| STM14_RS16860 | exbD | TonB system transport protein ExbD | WP_001251476.1 | -0.002290709 | 1 | 1 |
| STM14_RS23315 | | BglG family transcription antiterminator | WP_001212137.1 | -0.002938297 | 1 | 1 |
| STM14_RS05360 | ihfB | integration host factor subunit beta | WP_000167332.1 | -0.003082161 | 0.97327758 | 0.9905883 |
| STM14_RS06335 | rimJ | ribosomal protein S5-alanine N-acetyltransferase | WP_000468204.1 | -0.003259219 | 0.98055319 | 0.99542234 |
| STM14_RS13135 | aroC | chorismate synthase | WP_000918456.1 | -0.003279747 | 1 | 1 |
| STM14_RS18215 | rplN | 50S ribosomal protein L14 | WP_000613954.1 | -0.003802693 | 0.96996961 | 0.98904129 |
| STM14_RS26945 | | hypothetical protein | WP_001751809.1 | -0.004607321 | 1 | 1 |
| STM14_RS18165 | secY | preprotein translocase subunit SecY | WP_001118864.1 | -0.004709707 | 0.95792009 | 0.98055651 |
| STM14_RS22035 | panS | ketopantoate/pantoate/pantothenate transporter PanS | | -0.004972698 | 1 | 1 |
| STM14_RS19930 | dagR | transcriptional regulator DagR | WP_001252662.1 | -0.005381884 | 1 | 1 |
| STM14_RS22220 | | maltoporin | WP_000973645.1 | -0.005511752 | 0.98501195 | 0.99912232 |
| STM14_RS23940 | deoD | purine-nucleoside phosphorylase | WP_000224870.1 | -0.005547284 | 0.96754591 | 0.9870612 |
| STM14_RS18265 | rplC | 50S ribosomal protein L3 | WP_000579838.1 | -0.005772624 | 0.9573546 | 0.98037323 |
| STM14_RS10890 | | Y-family DNA polymerase | WP_000457662.1 | -0.006074765 | 1 | 1 |
| STM14_RS22190 | | YjbH domain-containing protein | WP_000750804.1 | -0.006347989 | 1 | 1 |
| STM14_RS23645 | iadA | beta-aspartyl-peptidase | WP_001113069.1 | -0.006427174 | 0.9672194 | 0.98705705 |
| STM14_RS09210 | | B3/4 domain-containing protein | WP_000097253.1 | -0.006709004 | 1 | 1 |
| STM14_RS15730 | cas1e | type I-E CRISPR-associated endonuclease Cas1e | WP_000144831.1 | -0.007021004 | 1 | 1 |
| STM14_RS07750 | | APC family permease | WP_000756089.1 | -0.007165432 | 1 | 1 |
| STM14_RS00670 | | fimbrial protein BcfA | WP_000743357.1 | -0.007697287 | 1 | 1 |
| STM14_RS00745 | | sulfatase-like hydrolase/transferase | WP_000831880.1 | -0.008340265 | 0.97172568 | 0.99003257 |
| STM14_RS19010 | uspA | universal stress protein UspA | WP_000323565.1 | -0.008487618 | 0.92071792 | 0.95307376 |
| STM14_RS13710 | arsC | arsenate reductase (glutaredoxin) | WP_000132648.1 | -0.008842376 | 0.9769915 | 0.99290166 |
| STM14_RS19665 | waaB | lipopolysaccharide 1%2C6-galactosyltransferase | WP_000683977.1 | -0.009025622 | 1 | 1 |
| STM14_RS01240 | ftsA | cell division protein FtsA | WP_000588463.1 | -0.009133824 | 0.93373665 | 0.9638403 |
| STM14_RS14590 | tyrA | bifunctional chorismate mutase/prephenate dehydrogenase | WP_000225191.1 | -0.009179231 | 1 | 1 |
| STM14_RS20025 | fucP | L-fucose:H+ symporter permease | WP_000998359.1 | -0.009320958 | 0.97337003 | 0.9905883 |
| STM14_RS02380 | golS | Au(I) sensor transcriptional regulator GolS | WP_001020583.1 | -0.009787724 | 1 | 1 |
| STM14_RS15745 | cas7e | type I-E CRISPR-associated protein Cas7/Cse4/CasC | WP_000206415.1 | -0.011106376 | 0.94660831 | 0.97215496 |
| STM14_RS11995 | | DUF1456 family protein | WP_000950413.1 | -0.011215399 | 1 | 1 |
| STM14_RS22450 | nrfA | ammonia-forming nitrite reductase cytochrome c552 subunit | WP_000101771.1 | -0.011479396 | 0.91776898 | 0.95082313 |
| STM14_RS08280 | asr | acid resistance repetitive basic protein Asr | WP_000756314.1 | -0.013015755 | 1 | 1 |
| STM14_RS18190 | rplF | 50S ribosomal protein L6 | WP_000091939.1 | -0.013025178 | 0.89333512 | 0.93127404 |
| STM14_RS03855 | | UxaA family hydrolase | WP_001574029.1 | -0.013492183 | 1 | 1 |
| STM14_RS20390 | glmU | bifunctional UDP-N-acetylglucosamine diphosphorylase/glucosamine- | WP_000934854.1 | -0.013628918 | 0.89190068 | 0.93030547 |
| STM14_RS13000 | | amino acid permease | WP_000380357.1 | -0.013636347 | 0.97387138 | 0.99071306 |
| STM14_RS20245 | | LysR family transcriptional regulator | WP_001054589.1 | -0.013918659 | 0.90444058 | 0.9407204 |
| STM14_RS14955 | | tripartite tricarboxylate transporter substrate binding protein | WP_000744413.1 | -0.014146319 | 0.91921363 | 0.95178419 |
| STM14_RS10345 | | MurR/RpiR family transcriptional regulator | WP_001056720.1 | -0.014400686 | 0.95278617 | 0.97720297 |
| STM14_RS01510 | gluQRS | tRNA glutamyl-Q(34) synthetase GluQRS | WP_000132295.1 | -0.014931065 | 1 | 1 |
| STM14_RS17790 | zapE | cell division protein ZapE | WP_001191222.1 | -0.015103045 | 0.94414211 | 0.97104225 |
| STM14_RS06240 | | phospholipase D family protein | WP_000976323.1 | -0.015314882 | 0.93299978 | 0.96334974 |
| STM14_RS19280 | tag | DNA-3-methyladenine glycosylase I | WP_001188414.1 | -0.015929921 | 0.96519532 | 0.98581301 |
| STM14_RS08300 | bioD | dethiobiotin synthase | WP_000919233.1 | -0.016356641 | 0.86930975 | 0.91461211 |
| STM14_RS05040 | potI | putrescine ABC transporter permease PotI | WP_001061629.1 | -0.016805966 | 1 | 1 |
| STM14_RS20220 | | 2-dehydro-3-deoxygalactonokinase | WP_000127125.1 | -0.016869639 | 1 | 1 |
| STM14_RS23105 | | YtfJ family protein | WP_000175305.1 | -0.017502231 | 0.97174843 | 0.99003257 |
| STM14_RS16805 | hybC | hydrogenase 2 large subunit | WP_000083044.1 | -0.017731639 | 0.84508889 | 0.89879182 |
| STM14_RS14905 | | TonB-dependent siderophore receptor | WP_001521281.1 | -0.01830107 | 0.95747478 | 0.98037323 |
| STM14_RS16755 | uxaC | glucuronate isomerase | WP_000190182.1 | -0.019578933 | 0.86146572 | 0.90950092 |
| STM14_RS18020 | | efflux RND transporter permease subunit | WP_001273295.1 | -0.019944168 | 1 | 1 |
| STM14_RS09800 | ychH | stress-induced protein YchH | WP_000823878.1 | -0.020386333 | 0.82170925 | 0.88010418 |
| STM14_RS08275 | | serine protease | WP_000549642.1 | -0.020574966 | 0.8698006 | 0.91461211 |

| | | | | | | |
|---|---|---|---|---|---|---|
| STM14_RS22010 | pepE | dipeptidase PepE | WP_000421792.1 | -0.02094918 | 0.86195148 | 0.90950092 |
| STM14_RS13790 | shdA | fibronectin-binding autotransporter adhesin ShdA | | -0.021306662 | 0.92267619 | 0.95483241 |
| STM14_RS11105 | | Rha family phage regulatory protein | WP_226949723.1 | -0.021675691 | 1 | 1 |
| STM14_RS11380 | | cobyrinate a%2Cc-diamide synthase | WP_000741259.1 | -0.023304735 | 0.8861155 | 0.9263706 |
| STM14_RS03650 | ahpF | alkyl hydroperoxide reductase subunit F | WP_000887644.1 | -0.024057917 | 0.85745266 | 0.9072624 |
| STM14_RS16110 | | LacI family DNA-binding transcriptional regulator | WP_000968354.1 | -0.025219606 | 0.96556904 | 0.98591936 |
| STM14_RS23420 | argF | ornithine carbamoyltransferase | WP_000103041.1 | -0.025250544 | 0.97262755 | 0.99038016 |
| STM14_RS05290 | | MFS transporter | WP_000109271.1 | -0.025405073 | 0.8944007 | 0.93212097 |
| STM14_RS10115 | htpX | protease HtpX | WP_000984498.1 | -0.025455663 | 0.80990414 | 0.87021494 |
| STM14_RS10355 | lpxM | lauroyl-Kdo(2)-lipid IV(A) myristoyltransferase | WP_000448401.1 | -0.02662164 | 0.87939177 | 0.92195914 |
| STM14_RS18730 | glgP | glycogen phosphorylase | WP_000993428.1 | -0.026753493 | 0.76395354 | 0.83343867 |
| STM14_RS20930 | tatB | Sec-independent protein translocase protein TatB | WP_000459612.1 | -0.027415419 | 0.86075763 | 0.90939056 |
| STM14_RS18410 | pabA | aminodeoxychorismate synthase component 2 | WP_000601880.1 | -0.027441624 | 0.95109693 | 0.97574223 |
| STM14_RS08140 | rsxC | electron transport complex subunit RsxC | WP_000915668.1 | -0.02784743 | 0.9428374 | 0.97078504 |
| STM14_RS15050 | | carboxymuconolactone decarboxylase family protein | WP_000209813.1 | -0.027975678 | 0.93168505 | 0.96253209 |
| STM14_RS15795 | queE | 7-carboxy-7-deazaguanine synthase QueE | WP_001199961.1 | -0.028536728 | 0.83048513 | 0.88658923 |
| STM14_RS06515 | | metal-dependent hydrolase | WP_000480277.1 | -0.028787581 | 0.8271205 | 0.88453607 |
| STM14_RS17700 | | TIGR01212 family radical SAM protein | WP_001176887.1 | -0.02896731 | 0.94269619 | 0.97078504 |
| STM14_RS01415 | speE | polyamine aminopropyltransferase | WP_000829968.1 | -0.029360846 | 0.87852179 | 0.92130938 |
| STM14_RS22655 | | DUF1778 domain-containing protein | WP_001110452.1 | -0.029971939 | 0.88662791 | 0.92659143 |
| STM14_RS03600 | entH | proofreading thioesterase EntH | WP_000637965.1 | -0.030068809 | 0.93513362 | 0.96501177 |
| STM14_RS08815 | mcbR | colanic acid/biofilm transcriptional regulator McbR | WP_001119592.1 | -0.030271082 | 0.85500181 | 0.90593285 |
| STM14_RS06050 | cbpA | curved DNA-binding protein | WP_000420603.1 | -0.030371401 | 0.74890494 | 0.82106178 |
| STM14_RS05115 | | SDR family oxidoreductase | WP_000069000.1 | -0.03090746 | 0.96152889 | 0.98343038 |
| STM14_RS00220 | | ExeA family protein | WP_001576625.1 | -0.031329541 | 0.74739403 | 0.82069471 |
| STM14_RS16080 | | TerC family protein | WP_000895635.1 | -0.031786489 | 0.87550045 | 0.91840247 |
| STM14_RS02520 | | extensin family protein | WP_000480794.1 | -0.031919755 | 0.85654271 | 0.9070435 |
| STM14_RS16560 | | YggS family pyridoxal phosphate-dependent enzyme | WP_000997812.1 | -0.032491952 | 0.86629305 | 0.91264407 |
| STM14_RS14515 | pssA | CDP-diacylglycerol--serine O-phosphatidyltransferase | WP_000949286.1 | -0.032840066 | 0.7777009 | 0.84443961 |
| STM14_RS10120 | prc | carboxy terminal-processing peptidase | WP_001091237.1 | -0.033368579 | 0.73870418 | 0.81385661 |
| STM14_RS17475 | infB | translation initiation factor IF-2 | WP_000133064.1 | -0.03382173 | 0.70494841 | 0.78692787 |
| STM14_RS19775 | rpoZ | DNA-directed RNA polymerase subunit omega | WP_000135058.1 | -0.033843805 | 0.80628889 | 0.86789572 |
| STM14_RS01250 | lpxC | UDP-3-O-acyl-N-acetylglucosamine deacetylase | WP_000595487.1 | -0.034468224 | 0.70060751 | 0.7836078 |
| STM14_RS08900 | | XRE family transcriptional regulator | WP_000369555.1 | -0.034650228 | 0.90776861 | 0.94311626 |
| STM14_RS01180 | rsmH | 16S rRNA (cytosine(1402)-N(4))-methyltransferase RsmH | WP_000970444.1 | -0.034655168 | 0.7414093 | 0.81545934 |
| STM14_RS07320 | | YeaC family protein | WP_001046864.1 | -0.034710552 | 0.71289637 | 0.79158156 |
| STM14_RS14565 | rluD | 23S rRNA pseudouridine(1911/1915/1917) synthase RluD | WP_000079130.1 | -0.034770766 | 0.75987704 | 0.82998138 |
| STM14_RS17985 | dusB | tRNA dihydrouridine synthase DusB | WP_001219664.1 | -0.035357129 | 0.82199262 | 0.88010418 |
| STM14_RS18270 | rpsJ | 30S ribosomal protein S10 | WP_001181005.1 | -0.03545869 | 0.7235756 | 0.80198836 |
| STM14_RS15135 | yqaB | fructose-1-phosphate/6-phosphogluconate phosphatase | WP_000271296.1 | -0.035607292 | 0.81446134 | 0.87355357 |
| STM14_RS17905 | yhdP | AsmA2 domain-containing protein YhdP | WP_001253505.1 | -0.036135852 | 0.83972517 | 0.89356461 |
| STM14_RS20255 | recF | DNA replication/repair protein RecF | WP_000060085.1 | -0.036161304 | 0.83993132 | 0.89356461 |
| STM14_RS22335 | uvrA | excinuclease ABC subunit UvrA | WP_000357724.1 | -0.036199325 | 0.82202017 | 0.88010418 |
| STM14_RS21510 | rraA | ribonuclease E activity regulator RraA | WP_000872918.1 | -0.036587894 | 0.67621538 | 0.7616473 |
| STM14_RS09315 | | YcjX family protein | WP_000825847.1 | -0.037023669 | 0.8383009 | 0.89356461 |
| STM14_RS08810 | | TonB-dependent receptor | WP_000692042.1 | -0.03748849 | 0.83468898 | 0.89056066 |
| STM14_RS20035 | | DeoR family transcriptional regulator | WP_000448638.1 | -0.037505064 | 0.83977751 | 0.89356461 |
| STM14_RS14525 | | MFS transporter | WP_000807809.1 | -0.037959493 | 0.90134716 | 0.93803285 |
| STM14_RS22175 | yjbE | exopolysaccharide production protein YjbE | WP_001541297.1 | -0.038717057 | 0.9439018 | 0.97104225 |
| STM14_RS14075 | | PTS transporter subunit EIIC | WP_000361663.1 | -0.038760397 | 0.86207448 | 0.90950092 |
| STM14_RS09355 | sapB | peptide ABC transporter permease SapB | WP_000583298.1 | -0.038877115 | 1 | 1 |
| STM14_RS14235 | | phage tail protein | WP_000677089.1 | -0.038912371 | 1 | 1 |
| STM14_RS05840 | ompA | porin OmpA | WP_001674965.1 | -0.039486485 | 0.64982162 | 0.74052715 |
| STM14_RS14935 | | HoxN/HupN/NixA family nickel/cobalt transporter | WP_001059352.1 | -0.03949233 | 0.88953471 | 0.9283636 |
| STM14_RS06555 | ycfP | alpha/beta hydrolase YcfP | WP_000587943.1 | -0.039803509 | 0.69951464 | 0.78297629 |
| STM14_RS18115 | mscL | large-conductance mechanosensitive channel protein MscL | WP_000008119.1 | -0.040205398 | 0.65292993 | 0.74314931 |
| STM14_RS15040 | | DUF883 domain-containing protein | WP_001519557.1 | -0.040305765 | 0.64831928 | 0.7395557 |
| STM14_RS20355 | pstA | phosphate ABC transporter permease PstA | WP_000212197.1 | -0.040574347 | 0.90226031 | 0.93871785 |
| STM14_RS07305 | | D-hexose-6-phosphate mutarotase | WP_000608662.1 | -0.04065156 | 0.79033043 | 0.85588137 |
| STM14_RS07370 | xthA | exodeoxyribonuclease III | WP_000673954.1 | -0.040761704 | 0.85411657 | 0.90577685 |
| STM14_RS03800 | mrdB | peptidoglycan glycosyltransferase MrdB | WP_000131713.1 | -0.040868596 | 0.91272996 | 0.94666809 |
| STM14_RS01755 | ldcC | lysine decarboxylase LdcC | WP_001021054.1 | -0.040918293 | 0.80792639 | 0.8693118 |
| STM14_RS09370 | sapF | peptide ABC transporter ATP-binding protein SapF | WP_000230462.1 | -0.04139533 | 0.93876747 | 0.968219 |
| STM14_RS19480 | selB | selenocysteine-specific translation elongation factor | WP_000582394.1 | -0.04307229 | 0.7254664 | 0.80311665 |
| STM14_RS18185 | rplR | 50S ribosomal protein L18 | WP_000358956.1 | -0.043133955 | 0.65260768 | 0.74301221 |
| STM14_RS06465 | plsX | phosphate acyltransferase PlsX | WP_001518286.1 | -0.043576569 | 0.77268024 | 0.84047522 |
| STM14_RS21310 | rhaR | HTH-type transcriptional activator RhaR | WP_000013290.1 | -0.044007741 | 0.85440156 | 0.90581818 |
| STM14_RS13685 | | glycine cleavage system transcriptional repressor | WP_000189061.1 | -0.04457682 | 0.74148774 | 0.81545934 |
| STM14_RS00655 | | glycosyl hydrolase family 18 protein | WP_000235798.1 | -0.044897678 | 0.88915194 | 0.9283636 |
| STM14_RS01690 | ispU | (2E%2C6E)-farnesyl-diphosphate-specific ditrans%2Cpolycis-undecap | WP_000947413.1 | -0.045041484 | 0.74820339 | 0.82074127 |
| STM14_RS19245 | eptB | kdo(2)-lipid A phosphoethanolamine 7''-transferase | WP_001269259.1 | -0.045111477 | 0.7926243 | 0.8566019 |
| STM14_RS21670 | argC | N-acetyl-gamma-glutamyl-phosphate reductase | WP_000935340.1 | -0.04582356 | 0.91182762 | 0.94599868 |
| STM14_RS10410 | nudB | dihydroneopterin triphosphate diphosphatase | WP_000653693.1 | -0.046042274 | 0.76440043 | 0.83343867 |
| STM14_RS21435 | lsrA | autoinducer 2 ABC transporter ATP-binding protein LsrA | WP_001167250.1 | -0.047000596 | 0.65438317 | 0.74434317 |
| STM14_RS21790 | secE | preprotein translocase subunit SecE | WP_001275691.1 | -0.047045462 | 0.6602065 | 0.74865425 |
| STM14_RS02700 | ribD | bifunctional diaminohydroxyphosphoribosylaminopyrimidine deaminas | WP_001150216.1 | -0.047277212 | 0.72820161 | 0.80469338 |
| STM14_RS08000 | punR | DNA-binding transcriptional activator PunR | WP_000269523.1 | -0.047368491 | 0.90745162 | 0.94305303 |
| STM14_RS20065 | emrD | multidrug efflux MFS transporter EmrD | WP_000828740.1 | -0.04743276 | 0.77921476 | 0.84566599 |
| STM14_RS01735 | rnhB | ribonuclease HII | WP_000569412.1 | -0.048426747 | 0.85683483 | 0.90709196 |
| STM14_RS19780 | spoT | bifunctional GTP diphosphokinase/guanosine-3'%2C5'-bis pyrophospha | WP_000280481.1 | -0.049360578 | 0.6767057 | 0.76180797 |
| STM14_RS08535 | hybD | HyaD/HybD family hydrogenase maturation endopeptidase | WP_000852664.1 | -0.049778395 | 0.81386981 | 0.87340969 |
| STM14_RS19075 | | STY4199 family HEPN domain-containing protein | WP_000185023.1 | -0.049843497 | 0.65868221 | 0.7473861 |

| | | | | | | |
|---|---|---|---|---|---|---|
| STM14_RS03645 | ahpC | alkyl hydroperoxide reductase subunit C | WP_000052802.1 | -0.0504166 | 0.5887601 | 0.68841369 |
| STM14_RS02515 | ddlA | D-alanine--D-alanine ligase | WP_001096588.1 | -0.050530154 | 0.6879866 | 0.77230691 |
| STM14_RS18990 | | HlyD family efflux transporter periplasmic adaptor subunit | WP_000359836.1 | -0.051047587 | 0.75869079 | 0.82937995 |
| STM14_RS09760 | sirB2 | invasion regulator SirB2 | WP_000150667.1 | -0.051877172 | 0.88683004 | 0.92659143 |
| STM14_RS19375 | | XylR family transcriptional regulator | WP_000459826.1 | -0.051909955 | 0.81447838 | 0.87355357 |
| STM14_RS10100 | yobH | invasion protein YobH | WP_001236777.1 | -0.05262832 | 0.86098194 | 0.90939056 |
| STM14_RS14195 | | phage minor tail protein L | WP_001152689.1 | -0.052702481 | 0.89222838 | 0.93038372 |
| STM14_RS23985 | radA | DNA repair protein RadA | WP_001029709.1 | -0.052735298 | 0.74780846 | 0.82069471 |
| STM14_RS09505 | | L-threonylcarbamoyladenylate synthase | WP_000077250.1 | -0.053025219 | 0.71106919 | 0.7907477 |
| STM14_RS18475 | gph | phosphoglycolate phosphatase | WP_000359655.1 | -0.053503985 | 0.71115332 | 0.7907477 |
| STM14_RS21680 | argH | argininosuccinate lyase | WP_001230051.1 | -0.054194978 | 0.76775901 | 0.83635759 |
| STM14_RS22610 | dmsB | dimethylsulfoxide reductase subunit B | WP_000816143.1 | -0.054591216 | 0.69196031 | 0.77582152 |
| STM14_RS15640 | | MarR family winged helix-turn-helix transcriptional regulator | WP_000423344.1 | -0.056017057 | 0.77342703 | 0.84103908 |
| STM14_RS14250 | | major capsid protein | WP_000522566.1 | -0.056144634 | 0.90071374 | 0.93763868 |
| STM14_RS00535 | | type II toxin-antitoxin system VapC family toxin | WP_000911322.1 | -0.05661296 | 0.8002366 | 0.86330828 |
| STM14_RS21985 | aceK | bifunctional isocitrate dehydrogenase kinase/phosphatase | WP_001137266.1 | -0.056705348 | 0.7482967 | 0.82074127 |
| STM14_RS04740 | dinG | ATP-dependent DNA helicase DinG | WP_001218630.1 | -0.057634136 | 0.74997601 | 0.82178128 |
| STM14_RS20470 | viaA | ATPase RavA stimulator ViaA | WP_000956588.1 | -0.058125687 | 0.73054103 | 0.80703633 |
| STM14_RS06395 | flgG | flagellar basal-body rod protein FlgG | WP_000625851.1 | -0.058372021 | 0.70988677 | 0.79014603 |
| STM14_RS18560 | | DUF4153 domain-containing protein | WP_000446437.1 | -0.058606211 | 0.71962891 | 0.79833493 |
| STM14_RS21395 | | MFS transporter | WP_000777317.1 | -0.058653537 | 0.88168547 | 0.92357493 |
| STM14_RS07330 | ansA | asparaginase | WP_001170134.1 | -0.059463804 | 0.79241612 | 0.8566019 |
| STM14_RS17805 | degS | outer membrane-stress sensor serine endopeptidase DegS | WP_000497705.1 | -0.060257118 | 0.79218415 | 0.8566019 |
| STM14_RS22735 | | DUF4156 domain-containing protein | WP_000558210.1 | -0.060486788 | 0.52830622 | 0.63238735 |
| STM14_RS19705 | rpmG | 50S ribosomal protein L33 | WP_001051798.1 | -0.060562355 | 0.65799801 | 0.7470702 |
| STM14_RS00520 | | entry exclusion protein | WP_010999946.1 | -0.060767001 | 0.54496209 | 0.64790133 |
| STM14_RS06640 | pepT | peptidase T | WP_000359410.1 | -0.06079074 | 0.55919071 | 0.66225159 |
| STM14_RS04155 | | YbfA family protein | WP_000401437.1 | -0.061437105 | 0.60322166 | 0.70197919 |
| STM14_RS14045 | qseE | two component system sensor histidine kinase QseE/GlrK | WP_001670672.1 | -0.062184084 | 0.68688537 | 0.77130876 |
| STM14_RS18550 | hslR | ribosome-associated heat shock protein Hsp15 | WP_000660723.1 | -0.062379401 | 0.86012641 | 0.90900845 |
| STM14_RS01540 | fhuA | ferrichrome porin FhuA | WP_000113220.1 | -0.062428193 | 0.60626725 | 0.7050777 |
| STM14_RS07770 | ttrA | tetrathionate reductase subunit TtrA | WP_000002375.1 | -0.062677558 | 0.77458003 | 0.84179565 |
| STM14_RS15755 | casA | type I-E CRISPR-associated protein Cse1/CasA | WP_000368584.1 | -0.062836553 | 0.83848913 | 0.89356461 |
| STM14_RS12700 | arnC | undecaprenyl-phosphate 4-deoxy-4-formamido-L-arabinose transferase | WP_000458893.1 | -0.063516204 | 0.70088225 | 0.78367703 |
| STM14_RS13490 | hemF | oxygen-dependent coproporphyrinogen oxidase | WP_000801350.1 | -0.063844922 | 0.77621832 | 0.84332719 |
| STM14_RS10140 | | PqiB family protein | WP_001535256.1 | -0.063872805 | 0.63235093 | 0.72691731 |
| STM14_RS20985 | trkH | Trk system potassium transporter TrkH | WP_000545688.1 | -0.063943596 | 0.75905299 | 0.82937995 |
| STM14_RS03850 | leuS | leucine--tRNA ligase | WP_001157918.1 | -0.064092341 | 0.48062776 | 0.58930117 |
| STM14_RS07135 | | ABC transporter permease | WP_001520270.1 | -0.064333136 | 0.815268 | 0.87389135 |
| STM14_RS10160 | pphA | protein-serine/threonine phosphatase | WP_000986176.1 | -0.064759123 | 0.8049261 | 0.86735086 |
| STM14_RS03225 | allB | allantoinase AllB | WP_000006865.1 | -0.065558488 | 0.83906577 | 0.89356461 |
| STM14_RS18540 | igaA | intracellular growth attenuator protein IgaA | WP_000104094.1 | -0.065626689 | 0.63103433 | 0.72631084 |
| STM14_RS05120 | ltaE | low-specificity L-threonine aldolase | WP_000566344.1 | -0.065926307 | 0.73330744 | 0.8096066 |
| STM14_RS09730 | | YchO/YchP family invasin | WP_000008985.1 | -0.066538668 | 0.6466817 | 0.73854901 |
| STM14_RS20475 | ravA | ATPase RavA | WP_000940991.1 | -0.066760007 | 0.62473761 | 0.72131825 |
| STM14_RS09075 | hrpA | ATP-dependent RNA helicase HrpA | WP_000139702.1 | -0.066892171 | 0.52215325 | 0.62706076 |
| STM14_RS10450 | | VOC family protein | WP_000024794.1 | -0.067330597 | 0.88120513 | 0.92333446 |
| STM14_RS12530 | | multidrug ABC transporter permease/ATP-binding protein | WP_000421590.1 | -0.067883854 | 0.7516361 | 0.82293313 |
| STM14_RS02165 | | class II glutamine amidotransferase | WP_000333387.1 | -0.067972001 | 0.63382687 | 0.72791032 |
| STM14_RS21565 | metB | cystathionine gamma-synthase | WP_000197217.1 | -0.067988939 | 0.73597991 | 0.81158371 |
| STM14_RS07155 | | mechanosensitive ion channel family protein | WP_000456065.1 | -0.068619005 | 0.83947388 | 0.89356461 |
| STM14_RS16615 | | oxidative damage protection protein | WP_000091706.1 | -0.068635509 | 0.61573737 | 0.71316295 |
| STM14_RS26715 | | IS4 family transposase | | -0.068740096 | 0.76494809 | 0.83378889 |
| STM14_RS12305 | | YejG family protein | WP_001094639.1 | -0.068813313 | 0.54126701 | 0.64496606 |
| STM14_RS21240 | fdnG | formate dehydrogenase-N subunit alpha | WP_010989087.1 | -0.068896694 | 0.41257976 | 0.52329127 |
| STM14_RS07325 | pncA | bifunctional nicotinamidase/pyrazinamidase | WP_000184708.1 | -0.069022964 | 0.74903491 | 0.82106178 |
| STM14_RS13845 | rodZ | cytoskeleton protein RodZ | WP_001090890.1 | -0.069111694 | 0.50708042 | 0.6143699 |
| STM14_RS11575 | hisB | bifunctional histidinol-phosphatase/imidazoleglycerol-phosphate dehy | WP_000080044.1 | -0.069636818 | 0.66124815 | 0.74914329 |
| STM14_RS09350 | sapA | peptide ABC transporter substrate-binding protein SapA | WP_001241629.1 | -0.070416176 | 0.71257566 | 0.79146413 |
| STM14_RS15335 | flhA | formate hydrogenlyase transcriptional activator FlhA | WP_000122692.1 | -0.071041985 | 0.57756457 | 0.67812269 |
| STM14_RS01760 | | VOC family protein | WP_000901088.1 | -0.07115661 | 0.75742311 | 0.82852997 |
| STM14_RS19070 | | alpha%2Calpha-trehalase | WP_000934257.1 | -0.071236662 | 0.62053831 | 0.71735243 |
| STM14_RS15225 | srlR | glucitol operon DNA-binding transcriptional repressor SrlR | WP_000804536.1 | -0.071241351 | 0.57522658 | 0.67667229 |
| STM14_RS22605 | | molybdopterin-dependent oxidoreductase | WP_000403482.1 | -0.072231383 | 0.46254827 | 0.5718948 |
| STM14_RS06045 | cbpM | chaperone modulator CbpM | WP_001284251.1 | -0.072394406 | 0.51498737 | 0.62068199 |
| STM14_RS08100 | tyrS | tyrosine--tRNA ligase | WP_000168626.1 | -0.072654133 | 0.45409006 | 0.56370856 |
| STM14_RS06110 | putP | sodium/proline symporter PutP | WP_001018467.1 | -0.07268126 | 0.53748582 | 0.64170648 |
| STM14_RS00515 | traG | conjugal transfer mating pair stabilization protein TraG | WP_001007086.1 | -0.07289156 | 0.54520011 | 0.64797508 |
| STM14_RS06495 | pabC | aminodeoxychorismate lyase | WP_000478660.1 | -0.073022926 | 0.70817018 | 0.78878195 |
| STM14_RS21910 | | DUF1481 domain-containing protein | WP_001083925.1 | -0.073483405 | 0.71244348 | 0.79146413 |
| STM14_RS09885 | emtA | membrane-bound lytic murein transglycosylase EmtA | WP_000776974.1 | -0.07352883 | 0.75300936 | 0.82419157 |
| STM14_RS11005 | | HK97 family phage prohead protease | WP_001193639.1 | -0.073586575 | 0.83970918 | 0.89356461 |
| STM14_RS17095 | | multifunctional CCA addition/repair protein | WP_000708443.1 | -0.074346875 | 0.61422181 | 0.71198852 |
| STM14_RS03505 | | TetR/AcrR family transcriptional regulator | WP_000113609.1 | -0.075318964 | 0.76329364 | 0.83321885 |
| STM14_RS18375 | | hydrolase | WP_001214118.1 | -0.075421841 | 0.58714762 | 0.68674635 |
| STM14_RS08665 | treY | malto-oligosyltrehalose synthase | WP_000613145.1 | -0.075680888 | 0.60652123 | 0.70515035 |
| STM14_RS00280 | | Y-family DNA polymerase | WP_000457541.1 | -0.077037052 | 0.79027611 | 0.85588137 |
| STM14_RS19120 | | pitrilysin family protein | WP_001163182.1 | -0.077115316 | 0.55985248 | 0.66260908 |
| STM14_RS02300 | | fimbrial outer membrane usher protein | WP_000075343.1 | -0.077304356 | 0.7429994 | 0.81683118 |
| STM14_RS03635 | | LysR family transcriptional regulator | WP_000002497.1 | -0.077512474 | 0.56861877 | 0.67104305 |
| STM14_RS03425 | rclA | reactive chlorine resistance oxidoreductase RclA | WP_000195929.1 | -0.077633033 | 0.68324629 | 0.76815659 |
| STM14_RS02910 | cdsH | cysteine desulfhydrase | WP_000987846.1 | -0.078504209 | 0.65600709 | 0.74549942 |

| | | | | | | |
|---|---|---|---|---|---|---|
| STM14_RS06365 | flgA | flagellar basal body P-ring formation chaperone FlgA | WP_001194082.1 | -0.078952862 | 0.74701853 | 0.82069471 |
| STM14_RS10430 | | MAPEG family protein | WP_000252975.1 | -0.079522402 | 0.81149548 | 0.87121095 |
| STM14_RS07675 | ydiK | AI-2E family transporter YdiK | WP_000248615.1 | -0.080155919 | 0.65698866 | 0.74638453 |
| STM14_RS04530 | modA | molybdate ABC transporter substrate-binding protein | WP_000951419.1 | -0.081573798 | 0.43191402 | 0.54295235 |
| STM14_RS01605 | erpA | iron-sulfur cluster insertion protein ErpA | WP_001278668.1 | -0.081737191 | 0.5143117 | 0.62047696 |
| STM14_RS01790 | nlpE | envelope stress response activation lipoprotein NlpE | WP_000252573.1 | -0.081946483 | 0.54265411 | 0.64640972 |
| STM14_RS03325 | fimA | type 1 fimbrial protein subunit FimA | WP_000681030.1 | -0.082599484 | 0.32989276 | 0.44141834 |
| STM14_RS21025 | mobA | molybdenum cofactor guanylyltransferase MobA | WP_001046887.1 | -0.083528832 | 0.67451172 | 0.76008462 |
| STM14_RS12695 | arnB | UDP-4-amino-4-deoxy-L-arabinose aminotransferase | WP_001279284.1 | -0.083980732 | 0.48873233 | 0.59742591 |
| STM14_RS12165 | yeiB | DUF418 domain-containing protein YeiB | WP_001526153.1 | -0.084032835 | 0.76971286 | 0.83786724 |
| STM14_RS16015 | ptrA | pitrilysin | WP_001138254.1 | -0.084554618 | 0.55254582 | 0.65543611 |
| STM14_RS09605 | | ion transporter | WP_000059074.1 | -0.084794321 | 0.70057628 | 0.7836078 |
| STM14_RS15010 | | metalloregulator ArsR/SmtB family transcription factor | WP_000137432.1 | -0.084874059 | 0.80944574 | 0.87018371 |
| STM14_RS18775 | | pirin family protein | WP_000639789.1 | -0.085271972 | 0.74077783 | 0.81516557 |
| STM14_RS21495 | glpK | glycerol kinase GlpK | WP_000136809.1 | -0.08581221 | 0.44277332 | 0.55320372 |
| STM14_RS14560 | yfiH | purine nucleoside phosphorylase YfiH | WP_000992639.1 | -0.085911556 | 0.37824558 | 0.48938921 |
| STM14_RS05775 | pepN | aminopeptidase N | WP_000193784.1 | -0.086669961 | 0.35666737 | 0.46751487 |
| STM14_RS07260 | | DUF441 domain-containing protein | WP_000460698.1 | -0.087118546 | 0.8032684 | 0.86607152 |
| STM14_RS18005 | | bifunctional diguanylate cyclase/phosphodiesterase | WP_001145331.1 | -0.087498859 | 0.88312926 | 0.92456126 |
| STM14_RS10150 | | YebV family protein | WP_024131108.1 | -0.087762535 | 0.41929529 | 0.52971462 |
| STM14_RS13820 | der | ribosome biogenesis GTPase Der | WP_000249411.1 | -0.088036883 | 0.49611112 | 0.6041876 |
| STM14_RS02710 | nusB | transcription antitermination factor NusB | WP_000801129.1 | -0.088504502 | 0.45607192 | 0.56597802 |
| STM14_RS06300 | solA | N-methyl-L-tryptophan oxidase | WP_000872773.1 | -0.088565819 | 0.52478611 | 0.62960202 |
| STM14_RS16820 | hybO | hydrogenase 2 small subunit | WP_000145429.1 | -0.088628015 | 0.31068679 | 0.42078275 |
| STM14_RS01260 | secA | preprotein translocase subunit SecA | WP_000905756.1 | -0.088929018 | 0.34148425 | 0.4530973 |
| STM14_RS02735 | ispA | (2E%2C6E)-farnesyl diphosphate synthase | WP_000348074.1 | -0.089255413 | 0.57560114 | 0.67689664 |
| STM14_RS05810 | pqiB | intermembrane transport protein PqiB | WP_000433414.1 | -0.089608139 | 0.47300844 | 0.58170242 |
| STM14_RS21900 | | YjaG family protein | WP_000940092.1 | -0.090831468 | 0.48897635 | 0.59742591 |
| STM14_RS18385 | | phosphoribulokinase | WP_001274690.1 | -0.090849106 | 0.57899286 | 0.67936638 |
| STM14_RS06685 | | NUDIX hydrolase | WP_000476067.1 | -0.090999322 | 0.83769001 | 0.89324489 |
| STM14_RS11505 | | FUSC family protein | WP_001200854.1 | -0.09103015 | 0.48903902 | 0.59742591 |
| STM14_RS12660 | | MFS transporter | WP_000028931.1 | -0.091053598 | 0.62820876 | 0.72425362 |
| STM14_RS19715 | radC | DNA repair protein RadC | WP_000380129.1 | -0.091445079 | 0.66717548 | 0.75423399 |
| STM14_RS13405 | | hypothetical protein | WP_000844541.1 | -0.092701208 | 0.82370673 | 0.88165354 |
| STM14_RS20740 | hemY | protoheme IX biogenesis protein HemY | WP_001546072.1 | -0.092720374 | 0.496379 | 0.6041876 |
| STM14_RS04360 | tolA | cell envelope integrity protein TolA | WP_000030590.1 | -0.092854626 | 0.49629595 | 0.6041876 |
| STM14_RS23220 | iolI1 | 2-keto-myo-inositol isomerase IolI1 | WP_000976924.1 | -0.093085156 | 0.82493454 | 0.88271112 |
| STM14_RS22320 | | MmcQ/YjbR family DNA-binding protein | WP_000155666.1 | -0.093355622 | 0.47337029 | 0.58195306 |
| STM14_RS12850 | nuoA | NADH-quinone oxidoreductase subunit NuoA | WP_000062993.1 | -0.093548454 | 0.34817483 | 0.45883312 |
| STM14_RS18920 | | DUF1145 family protein | WP_000906757.1 | -0.093689513 | 0.86577595 | 0.91262154 |
| STM14_RS01675 | pyrH | UMP kinase | WP_000224567.1 | -0.094351655 | 0.37997986 | 0.49107962 |
| STM14_RS06105 | putA | trifunctional transcriptional regulator/proline dehydrogenase/L-glutama | WP_000537528.1 | -0.094415865 | 0.34321674 | 0.45457298 |
| STM14_RS22400 | | glutathione S-transferase | WP_000958575.1 | -0.094453718 | 0.51599954 | 0.62149503 |
| STM14_RS18555 | hslO | Hsp33 family molecular chaperone HslO | WP_001669339.1 | -0.094625181 | 0.51674836 | 0.62209698 |
| STM14_RS15300 | hycB | formate hydrogenlyase subunit HycB | WP_001079164.1 | -0.094867184 | 0.86605518 | 0.91264407 |
| STM14_RS07550 | pheT | phenylalanine--tRNA ligase subunit beta | WP_000672402.1 | -0.09523484 | 0.26271267 | 0.36807764 |
| STM14_RS08945 | sgcC | PTS sugar transporter subunit IIC SgcC | WP_000460835.1 | -0.09584108 | 0.50267522 | 0.60983531 |
| STM14_RS02895 | queC | 7-cyano-7-deazaguanine synthase QueC | WP_000817201.1 | -0.09651473 | 0.60839667 | 0.70710749 |
| STM14_RS20105 | | putative transporter | WP_001279793.1 | -0.096635556 | 0.48468528 | 0.59368303 |
| STM14_RS18380 | | YheU family protein | WP_000586568.1 | -0.09699971 | 0.75024978 | 0.82178128 |
| STM14_RS10565 | | DJ-1/PfpI family protein | WP_000754800.1 | -0.097531924 | 0.61027884 | 0.7084006 |
| STM14_RS07760 | | glutamine amidotransferase | WP_000697187.1 | -0.097809483 | 0.64394506 | 0.73679482 |
| STM14_RS09475 | topA | type I DNA topoisomerase | WP_000513593.1 | -0.097814156 | 0.33914156 | 0.45096397 |
| STM14_RS20710 | rffM | lipopolysaccharide N-acetylmannosaminouronosyltransferase | WP_000183621.1 | -0.097816847 | 0.73081447 | 0.80709625 |
| STM14_RS08750 | | TetR family transcriptional regulator | WP_000121043.1 | -0.097891836 | 0.63642882 | 0.72933499 |
| STM14_RS13180 | | formate/nitrite transporter family protein | WP_000377772.1 | -0.098202763 | 0.51392498 | 0.62047696 |
| STM14_RS08825 | mddA | L-methionine sulfoximine/L-methionine sulfone acetyltransferase | WP_000155373.1 | -0.099565482 | 0.65362312 | 0.74370838 |
| STM14_RS12870 | yfbR | 5'-deoxynucleotidase | WP_000813882.1 | -0.099922477 | 0.60895536 | 0.70731029 |
| STM14_RS11880 | yegQ | tRNA 5-hydroxyuridine modification protein YegQ | WP_001517981.1 | -0.100520956 | 0.63400554 | 0.72791032 |
| STM14_RS04285 | sdhA | succinate dehydrogenase flavoprotein subunit | WP_000775561.1 | -0.101091785 | 0.26594655 | 0.37161867 |
| STM14_RS17370 | | YraN family protein | WP_000057285.1 | -0.101201735 | 0.72080204 | 0.79939551 |
| STM14_RS18065 | | DUF1488 domain-containing protein | WP_001070571.1 | -0.101312618 | 0.64856745 | 0.7395557 |
| STM14_RS22585 | dcuB | anaerobic C4-dicarboxylate transporter DcuB | WP_000899149.1 | -0.101345493 | 0.60963133 | 0.70787213 |
| STM14_RS01800 | tsaA | tRNA (N6-threonylcarbamoyladenosine(37)-N6)-methyltransferase Trm | WP_000093986.1 | -0.101458996 | 0.8536108 | 0.90550128 |
| STM14_RS18530 | mrcA | peptidoglycan glycosyltransferase/peptidoglycan DD-transpeptidase M | WP_000673808.1 | -0.101586983 | 0.4521664 | 0.56150984 |
| STM14_RS19160 | bcsR | cellulose biosynthesis protein BcsR | WP_000282616.1 | -0.102148217 | 0.61078257 | 0.70853857 |
| STM14_RS02855 | tig | trigger factor | WP_001198406.1 | -0.102206247 | 0.2296335 | 0.33105346 |
| STM14_RS23090 | | helix-turn-helix domain-containing protein | WP_001201297.1 | -0.102426615 | 0.67326938 | 0.75902567 |
| STM14_RS15825 | relA | GTP diphosphokinase | WP_000226838.1 | -0.103153656 | 0.25568186 | 0.36000788 |
| STM14_RS10550 | otsA | alpha%2Calpha-trehalose-phosphate synthase | WP_000089042.1 | -0.103444236 | 0.37827314 | 0.48938921 |
| STM14_RS20290 | mnmE | tRNA uridine-5-carboxymethylaminomethyl(34) synthesis GTPase Mnm | WP_000019077.1 | -0.103708168 | 0.68014206 | 0.76513384 |
| STM14_RS09165 | zntB | zinc transporter ZntB | WP_000387373.1 | -0.10378702 | 0.45208989 | 0.56150984 |
| STM14_RS04715 | | ABC transporter permease | WP_000045765.1 | -0.103862247 | 0.57704554 | 0.6777294 |
| STM14_RS12295 | | ABC transporter permease | WP_001137764.1 | -0.104871366 | 0.85976683 | 0.90888931 |
| STM14_RS01420 | | YacC family pilotin-like protein | WP_000846613.1 | -0.105505461 | 0.62826346 | 0.72425362 |
| STM14_RS15030 | alaE | L-alanine exporter AlaE | WP_000492669.1 | -0.105699716 | 0.37387191 | 0.48505862 |
| STM14_RS05800 | | ABC transporter ATP-binding protein | WP_000053044.1 | -0.105719365 | 0.38982779 | 0.50064386 |
| STM14_RS23630 | | Hcp family type VI secretion system effector | WP_000907671.1 | -0.105734706 | 0.29275875 | 0.40072035 |
| STM14_RS08515 | | ATP/GTP-binding protein | WP_001189401.1 | -0.106356894 | 0.76161229 | 0.83163003 |
| STM14_RS23800 | | DUF2501 domain-containing protein | WP_001090314.1 | -0.106539001 | 0.56747143 | 0.66990376 |
| STM14_RS10600 | | YecA family protein | WP_000781520.1 | -0.106601848 | 0.20936782 | 0.30700609 |
| STM14_RS17570 | rpmA | 50S ribosomal protein L27 | WP_000940593.1 | -0.10732865 | 0.27115916 | 0.37719396 |

| | | | | | | |
|---|---|---|---|---|---|---|
| STM14_RS08590 | | MarR family winged helix-turn-helix transcriptional regulator | WP_001293638.1 | -0.107710526 | 0.66459952 | 0.75201458 |
| STM14_RS04165 | | MFS transporter | WP_001041123.1 | -0.107815822 | 0.64787616 | 0.73945444 |
| STM14_RS19000 | pitA | inorganic phosphate transporter PitA | WP_000902764.1 | -0.10800772 | 0.49446786 | 0.60305752 |
| STM14_RS02500 | yaiW | surface-exposed outer membrane lipoprotein YaiW | WP_001271421.1 | -0.108358396 | 0.56166904 | 0.66454544 |
| STM14_RS00230 | | recombinase family protein | WP_001240331.1 | -0.109362057 | 0.26931559 | 0.37518728 |
| STM14_RS13400 | crr | PTS glucose transporter subunit IIA | WP_000522253.1 | -0.109659641 | 0.21245006 | 0.31053637 |
| STM14_RS17015 | | glutathionylspermidine synthase family protein | WP_000442833.1 | -0.109989454 | 0.29709365 | 0.40574883 |
| STM14_RS24040 | creD | cell envelope integrity protein CreD | WP_000920374.1 | -0.110272199 | 0.5400204 | 0.64431468 |
| STM14_RS03730 | dpiB | sensor histidine kinase DpiB | WP_001277836.1 | -0.110979956 | 0.59936099 | 0.69814843 |
| STM14_RS05225 | aat | leucyl/phenylalanyl-tRNA--protein transferase | WP_001241650.1 | -0.111209803 | 0.72568416 | 0.80311665 |
| STM14_RS06435 | rluC | 23S rRNA pseudouridine(955/2504/2580) synthase RluC | WP_000846319.1 | -0.111221465 | 0.51312028 | 0.62005542 |
| STM14_RS23565 | | DUF1819 family protein | WP_001740084.1 | -0.111336783 | 0.41170214 | 0.5223595 |
| STM14_RS18100 | fmt | methionyl-tRNA formyltransferase | WP_001285165.1 | -0.111633925 | 0.38611607 | 0.4971761 |
| STM14_RS19585 | | rhodanese-like domain-containing protein | WP_001156179.1 | -0.111872734 | 0.26680606 | 0.37253694 |
| STM14_RS23990 | nadR | multifunctional transcriptional regulator/nicotinamide-nucleotide aden | WP_000093829.1 | -0.113576247 | 0.2357082 | 0.33769556 |
| STM14_RS18545 | yrfG | GMP/IMP nucleotidase | WP_001520508.1 | -0.114384397 | 0.49216037 | 0.60064119 |
| STM14_RS17640 | lptC | LPS export ABC transporter periplasmic protein LptC | WP_000047845.1 | -0.115017978 | 0.39063323 | 0.50150333 |
| STM14_RS01680 | frr | ribosome recycling factor | WP_000622423.1 | -0.115319735 | 0.23107405 | 0.33234948 |
| STM14_RS03020 | priC | primosomal replication protein N'' | WP_001738050.1 | -0.115658149 | 0.52997138 | 0.63396836 |
| STM14_RS11000 | | phage major capsid protein | WP_000257528.1 | -0.115834033 | 0.47430135 | 0.58270856 |
| STM14_RS02410 | ssaA | small effector SsaA | WP_000934834.1 | -0.115915522 | 0.3801936 | 0.49118345 |
| STM14_RS10055 | | PTS mannose transporter subunit IID | WP_001518647.1 | -0.117484317 | 0.28020075 | 0.38727446 |
| STM14_RS16135 | kduD | 2-dehydro-3-deoxy-D-gluconate 5-dehydrogenase KduD | WP_000602490.1 | -0.117591634 | 0.70360527 | 0.78600564 |
| STM14_RS12155 | mglB | galactose/glucose ABC transporter substrate-binding protein MglB | WP_001036943.1 | -0.117631325 | 0.33136848 | 0.44301232 |
| STM14_RS18640 | malT | HTH-type transcriptional regulator MalT | WP_000907029.1 | -0.117678699 | 0.24380368 | 0.34619558 |
| STM14_RS13915 | fdx | ISC system 2Fe-2S type ferredoxin | WP_001124466.1 | -0.117912102 | 0.41146904 | 0.5222437 |
| STM14_RS17650 | lptB | LPS export ABC transporter ATP-binding protein | WP_000224104.1 | -0.120879469 | 0.33957246 | 0.45133061 |
| STM14_RS13410 | pdxK | pyridoxine/pyridoxal/pyridoxamine kinase | WP_000529613.1 | -0.120895887 | 0.49521486 | 0.60359394 |
| STM14_RS11590 | hisF | imidazole glycerol phosphate synthase subunit HisF | WP_000880125.1 | -0.121769836 | 0.49662695 | 0.60428963 |
| STM14_RS12055 | | Yip1 family protein | WP_001017057.1 | -0.122835702 | 0.73655014 | 0.81196934 |
| STM14_RS20645 | trxA | thioredoxin TrxA | WP_001280776.1 | -0.122887816 | 0.17697151 | 0.26771121 |
| STM14_RS02900 | | SgrR family transcriptional regulator | WP_001238163.1 | -0.123007562 | 0.50986367 | 0.61718167 |
| STM14_RS20000 | uhpT | hexose-6-phosphate:phosphate antiporter | WP_000879181.1 | -0.123366707 | 0.63515099 | 0.7285439 |
| STM14_RS13120 | | elongation factor P hydroxylase | WP_001066345.1 | -0.123611926 | 0.65791415 | 0.7470702 |
| STM14_RS01545 | fhuC | Fe3+-hydroxamate ABC transporter ATP-binding protein FhuC | WP_001157204.1 | -0.124254414 | 0.7934785 | 0.85727343 |
| STM14_RS01645 | | MFS transporter | | -0.124580121 | 0.39440882 | 0.50529341 |
| STM14_RS20450 | mnmG | tRNA uridine-5-carboxymethylaminomethyl(34) synthesis enzyme Mnm | WP_000499872.1 | -0.124907634 | 0.29310388 | 0.40074581 |
| STM14_RS05475 | pncB | nicotinate phosphoribosyltransferase | WP_000191399.1 | -0.125388467 | 0.57312136 | 0.67510284 |
| STM14_RS00685 | | fimbrial protein BcfD | WP_000701247.1 | -0.125987991 | 0.79951289 | 0.86278032 |
| STM14_RS04695 | clsB | cardiolipin synthase ClsB | WP_000649648.1 | -0.126040971 | 0.65537368 | 0.74501847 |
| STM14_RS23615 | uxuR | Uxu operon transcriptional regulator | WP_000841654.1 | -0.126485326 | 0.26544131 | 0.37105349 |
| STM14_RS06680 | mnmA | tRNA 2-thiouridine(34) synthase MnmA | WP_000004541.1 | -0.126891444 | 0.34333717 | 0.45457298 |
| STM14_RS09015 | | metal/formaldehyde-sensitive transcriptional repressor | WP_001120770.1 | -0.126894463 | 0.5626314 | 0.66525652 |
| STM14_RS20900 | udp | uridine phosphorylase | WP_000045169.1 | -0.127168974 | 0.23936182 | 0.34120411 |
| STM14_RS15910 | fucA | L-fuculose-phosphate aldolase | WP_000440819.1 | -0.1271975 | 0.66438862 | 0.75200704 |
| STM14_RS19575 | secB | protein-export chaperone SecB | WP_000003370.1 | -0.129012263 | 0.16989815 | 0.2593553 |
| STM14_RS01600 | clcA | H(+)/Cl(-) exchange transporter ClcA | WP_000845425.1 | -0.129120238 | 0.44725566 | 0.55729114 |
| STM14_RS21920 | zraS | two-component system sensor histidine kinase ZraS | WP_001526924.1 | -0.129450635 | 0.34392517 | 0.45469748 |
| STM14_RS09670 | | YchJ family protein | WP_001682412.1 | -0.12948853 | 0.35932605 | 0.46952033 |
| STM14_RS03805 | mrdA | peptidoglycan DD-transpeptidase MrdA | WP_000828700.1 | -0.129914427 | 0.51726377 | 0.62240693 |
| STM14_RS23230 | iolC | 5-dehydro-2-deoxygluconokinase | WP_014343940.1 | -0.13031525 | 0.49233563 | 0.60065599 |
| STM14_RS16130 | araE | arabinose-proton symporter AraE | WP_000253650.1 | -0.130341773 | 0.68492381 | 0.76957262 |
| STM14_RS16595 | | DUF2884 domain-containing protein | WP_000984838.1 | -0.130704613 | 0.33730426 | 0.44884506 |
| STM14_RS05045 | | DUF2593 family protein | WP_000505788.1 | -0.13096414 | 0.62107429 | 0.71753861 |
| STM14_RS04720 | | ATP-binding cassette domain-containing protein | WP_001287616.1 | -0.130965647 | 0.35166304 | 0.46210682 |
| STM14_RS17920 | mreD | rod shape-determining protein MreD | WP_000179403.1 | -0.131241686 | 0.63477157 | 0.72833559 |
| STM14_RS05250 | ftsK | DNA translocase FtsK | WP_001526798.1 | -0.13236835 | 0.19230324 | 0.28666418 |
| STM14_RS05165 | cspD | cold shock-like protein CspD | WP_000447499.1 | -0.132494358 | 0.25547535 | 0.35986357 |
| STM14_RS03290 | ppiB | peptidylprolyl isomerase B | WP_000255991.1 | -0.132922633 | 0.15292314 | 0.2388897 |
| STM14_RS18180 | rpsE | 30S ribosomal protein S5 | WP_000940121.1 | -0.132959961 | 0.16442767 | 0.25304773 |
| STM14_RS15885 | ppnN | nucleotide 5'-monophosphate nucleosidase PpnN | WP_000627973.1 | -0.132994628 | 0.18065281 | 0.27260804 |
| STM14_RS21700 | | YijD family membrane protein | WP_000813834.1 | -0.132999914 | 0.46891444 | 0.57801908 |
| STM14_RS16405 | pgk | phosphoglycerate kinase | WP_000111274.1 | -0.13324908 | 0.14820935 | 0.23320805 |
| STM14_RS16055 | lgt | prolipoprotein diacylglyceryl transferase | WP_000204645.1 | -0.133573766 | 0.41319234 | 0.52370885 |
| STM14_RS00255 | | hypothetical protein | WP_000988946.1 | -0.133627085 | 0.43759764 | 0.54809778 |
| STM14_RS12625 | glpT | glycerol-3-phosphate transporter | WP_000948739.1 | -0.134081664 | 0.30784014 | 0.41732967 |
| STM14_RS17135 | mug | G/U mismatch-specific DNA glycosylase | WP_000237776.1 | -0.134146068 | 0.16209095 | 0.25019872 |
| STM14_RS23575 | | YebC/PmpR family DNA-binding transcriptional regulator | WP_000532937.1 | -0.134457146 | 0.23411382 | 0.33619621 |
| STM14_RS04255 | nei | endonuclease VIII | WP_001113970.1 | -0.134550371 | 0.34483322 | 0.45557083 |
| STM14_RS16765 | gss | bifunctional glutathionylspermidine amidase/synthase | WP_000033693.1 | -0.134888447 | 0.22573817 | 0.32658858 |
| STM14_RS20910 | ubiE | bifunctional demethylmenaquinone methyltransferase/2-methoxy-6-po | WP_000229009.1 | -0.135098871 | 0.2113747 | 0.30945592 |
| STM14_RS06765 | | CII family transcriptional regulator | WP_072143007.1 | -0.135589201 | 0.6355324 | 0.72875438 |
| STM14_RS22465 | nrfD | cytochrome c nitrite reductase subunit NrfD | WP_000199637.1 | -0.135686571 | 0.66900385 | 0.75542118 |
| STM14_RS09970 | minE | cell division topological specificity factor MinE | WP_001185666.1 | -0.135741778 | 0.29379128 | 0.40138757 |
| STM14_RS22240 | plsB | glycerol-3-phosphate 1-O-acyltransferase PlsB | WP_000017359.1 | -0.135806689 | 0.18677121 | 0.27966312 |
| STM14_RS07560 | btuC | vitamin B12 ABC transporter permease BtuC | WP_000954984.1 | -0.136136832 | 0.70957055 | 0.79003289 |
| STM14_RS21485 | fpr | ferredoxin--NADP(+) reductase | WP_000796300.1 | -0.137015665 | 0.24966938 | 0.35316276 |
| STM14_RS04425 | aroG | 3-deoxy-7-phosphoheptulonate synthase AroG | WP_001109234.1 | -0.137367382 | 0.15187069 | 0.23764891 |
| STM14_RS16350 | | YecA family protein | WP_000027236.1 | -0.137597013 | 0.18418865 | 0.2769065 |
| STM14_RS19115 | | sugar kinase | WP_000037583.1 | -0.138366264 | 0.33579403 | 0.44770205 |
| STM14_RS03955 | | PhoH family protein | WP_001018045.1 | -0.138413689 | 0.17575245 | 0.2665241 |
| STM14_RS02440 | | LysE family transporter | WP_000433131.1 | -0.139222228 | 0.36665782 | 0.47721248 |

| | | | | | | |
|---|---|---|---|---|---|---|
| STM14_RS22385 | | EAL domain-containing protein | WP_000083640.1 | -0.139598322 | 0.51526258 | 0.62081047 |
| STM14_RS16725 | | polysaccharide deacetylase | WP_000540889.1 | -0.139607166 | 0.37777889 | 0.48926552 |
| STM14_RS08240 | | amino acid permease | WP_000412353.1 | -0.139848719 | 0.70789843 | 0.78878195 |
| STM14_RS04075 | fur | ferric iron uptake transcriptional regulator | WP_000131699.1 | -0.140102548 | 0.17444393 | 0.2649763 |
| STM14_RS08445 | eamA | O-acetylserine/cysteine exporter | WP_000987751.1 | -0.140137609 | 0.61566015 | 0.71316295 |
| STM14_RS13370 | zipA | cell division protein ZipA | WP_000983110.1 | -0.140182313 | 0.17815619 | 0.26904444 |
| STM14_RS19840 | | DUF4942 domain-containing protein | | -0.140848877 | 0.52006235 | 0.62495744 |
| STM14_RS11805 | asmA | outer membrane assembly protein AsmA | WP_001252270.1 | -0.141239772 | 0.26144458 | 0.36657995 |
| STM14_RS19935 | | hypothetical protein | WP_000824217.1 | -0.14167256 | 0.17263863 | 0.26266754 |
| STM14_RS16270 | xerD | site-specific tyrosine recombinase XerD | WP_000434302.1 | -0.142023134 | 0.37490283 | 0.48588435 |
| STM14_RS13460 | ucpA | SDR family oxidoreductase UcpA | WP_000517460.1 | -0.142211004 | 0.09402155 | 0.16004963 |
| STM14_RS23305 | | DgaE family pyridoxal phosphate-dependent ammonia lyase | WP_001139187.1 | -0.14275639 | 0.61785793 | 0.7151691 |
| STM14_RS17635 | kdsC | 3-deoxy-manno-octulosonate-8-phosphatase KdsC | WP_000030021.1 | -0.143624995 | 0.3056452 | 0.41481224 |
| STM14_RS14125 | era | GTPase Era | WP_000102232.1 | -0.14370761 | 0.17526501 | 0.26606759 |
| STM14_RS02535 | psiF | phosphate starvation-inducible protein PsiF | WP_000705165.1 | -0.144281105 | 0.14980435 | 0.23523943 |
| STM14_RS20400 | | F0F1 ATP synthase subunit epsilon | WP_001251971.1 | -0.146643007 | 0.14526708 | 0.22936251 |
| STM14_RS22060 | | tail fiber protein | WP_000368196.1 | -0.146827225 | 0.61377612 | 0.71178699 |
| STM14_RS14450 | rseA | anti-sigma-E factor RseA | WP_001168374.1 | -0.147145829 | 0.11550574 | 0.19011718 |
| STM14_RS20410 | atpG | F0F1 ATP synthase subunit gamma | WP_000896506.1 | -0.147298832 | 0.15117586 | 0.23667145 |
| STM14_RS17620 | mlaF | phospholipid ABC transporter ATP-binding protein MlaF | WP_000531566.1 | -0.147379465 | 0.37885629 | 0.48997149 |
| STM14_RS17895 | aaeR | HTH-type transcriptional activator AaeR | WP_000440300.1 | -0.147522709 | 0.45194287 | 0.56150984 |
| STM14_RS16610 | mutY | A/G-specific adenine glycosylase | WP_001148956.1 | -0.147836292 | 0.33219379 | 0.44380897 |
| STM14_RS23120 | msrA | peptide-methionine (S)-S-oxide reductase MsrA | WP_000051472.1 | -0.148141567 | 0.2076195 | 0.30509238 |
| STM14_RS08910 | | DUF3313 domain-containing protein | WP_001259025.1 | -0.148234603 | 0.2721628 | 0.37815224 |
| STM14_RS22715 | fxsA | membrane protein FxsA | WP_001267291.1 | -0.148581891 | 0.40005668 | 0.51057494 |
| STM14_RS08850 | srfC | virulence-associated effector SrfC | WP_000165648.1 | -0.148940316 | 0.50598605 | 0.6134477 |
| STM14_RS16590 | ansB | L-asparaginase 2 | WP_000394189.1 | -0.14902465 | 0.13680773 | 0.21777189 |
| STM14_RS03815 | rsfS | ribosome silencing factor | WP_001161670.1 | -0.149546126 | 0.41558109 | 0.52619311 |
| STM14_RS14030 | glnB | nitrogen regulatory protein P-II | WP_000717694.1 | -0.149754317 | 0.43487916 | 0.54574815 |
| STM14_RS19005 | uspB | universal stress protein UspB | WP_000626193.1 | -0.151030362 | 0.4081173 | 0.51924253 |
| STM14_RS10590 | | YecH family protein | WP_000377238.1 | -0.15141285 | 0.52503156 | 0.62960202 |
| STM14_RS09660 | rssB | two-component system response regulator RssB | WP_000193432.1 | -0.151841801 | 0.28786297 | 0.39519443 |
| STM14_RS01065 | polB | DNA polymerase II | WP_000035723.1 | -0.151880879 | 0.34507365 | 0.45572496 |
| STM14_RS14130 | rnc | ribonuclease III | WP_001068341.1 | -0.15188396 | 0.16641258 | 0.25541105 |
| STM14_RS22030 | | DUF3811 domain-containing protein | WP_001207628.1 | -0.152121072 | 0.46449003 | 0.57355412 |
| STM14_RS16060 | ptsP | phosphoenolpyruvate--protein phosphotransferase | WP_000957887.1 | -0.152601516 | 0.1560282 | 0.24296761 |
| STM14_RS19365 | xylB | xylulokinase | WP_000275326.1 | -0.15332387 | 0.45914233 | 0.56880407 |
| STM14_RS08310 | osmV | osmoprotectant ABC transporter ATP-binding protein OsmV | WP_000593086.1 | -0.153573954 | 0.28401667 | 0.39137327 |
| STM14_RS20380 | | SgrR family transcriptional regulator | WP_000213163.1 | -0.154024359 | 0.63467596 | 0.72833559 |
| STM14_RS06570 | comR | TetR family copper-responsive transcriptional repressor ComR | WP_000205990.1 | -0.1548622 | 0.1420436 | 0.22475485 |
| STM14_RS11695 | galF | GalU regulator GalF | WP_000981469.1 | -0.155200401 | 0.14691771 | 0.23167067 |
| STM14_RS05240 | trxB | thioredoxin-disulfide reductase | WP_000537407.1 | -0.155514669 | 0.10475734 | 0.17548522 |
| STM14_RS17120 | rpsU | 30S ribosomal protein S21 | WP_001144069.1 | -0.155522908 | 0.16707832 | 0.25600598 |
| STM14_RS01670 | tsf | translation elongation factor Ts | WP_000808106.1 | -0.155860181 | 0.07378864 | 0.13068292 |
| STM14_RS11235 | | AMP nucleosidase | WP_000428191.1 | -0.156400037 | 0.17300646 | 0.26311846 |
| STM14_RS22245 | | diacylglycerol kinase | WP_000002902.1 | -0.15680672 | 0.44622678 | 0.55619736 |
| STM14_RS16620 | mltC | membrane-bound lytic murein transglycosylase MltC | WP_000976289.1 | -0.157743993 | 0.51683722 | 0.62209698 |
| STM14_RS01265 | mutT | 8-oxo-dGTP diphosphatase MutT | WP_000736056.1 | -0.157816628 | 0.50369326 | 0.6108691 |
| STM14_RS17425 | | LLM class flavin-dependent oxidoreductase | WP_000130437.1 | -0.158007131 | 0.49523555 | 0.60359394 |
| STM14_RS17555 | dacB | serine-type D-Ala-D-Ala carboxypeptidase | WP_001212680.1 | -0.158276916 | 0.36506093 | 0.47563848 |
| STM14_RS23935 | deoB | phosphopentomutase | WP_000816454.1 | -0.158488217 | 0.25548163 | 0.35986357 |
| STM14_RS22775 | | lipocalin family protein | WP_001238397.1 | -0.158631091 | 0.35041366 | 0.46095859 |
| STM14_RS22690 | | protein-disulfide reductase DsbD | WP_000068899.1 | -0.158646849 | 0.51039419 | 0.61736905 |
| STM14_RS13725 | upp | uracil phosphoribosyltransferase | WP_000706208.1 | -0.158666061 | 0.15070805 | 0.23623118 |
| STM14_RS01160 | ilvI | acetolactate synthase 3 large subunit | WP_000425604.1 | -0.159228369 | 0.49556811 | 0.60379939 |
| STM14_RS05230 | cydC | cysteine/glutathione ABC transporter ATP-binding protein/permease Cy | WP_001202267.1 | -0.159339697 | 0.31592757 | 0.42672241 |
| STM14_RS08125 | | electron transport complex subunit E | WP_001289628.1 | -0.159384899 | 0.5435772 | 0.64685996 |
| STM14_RS21600 | katG | catalase/peroxidase HPI | WP_000108103.1 | -0.159801706 | 0.07970432 | 0.1393501 |
| STM14_RS04025 | nagA | N-acetylglucosamine-6-phosphate deacetylase | WP_000271186.1 | -0.16004179 | 0.22699405 | 0.32801888 |
| STM14_RS16105 | galR | HTH-type transcriptional regulator GalR | WP_000201030.1 | -0.160347613 | 0.23656946 | 0.33879765 |
| STM14_RS18970 | acpT | 4'-phosphopantetheinyl transferase AcpT | WP_000285376.1 | -0.160351995 | 0.46491594 | 0.57373291 |
| STM14_RS04040 | glnS | glutamine--tRNA ligase | WP_001287181.1 | -0.16098654 | 0.09312153 | 0.15888484 |
| STM14_RS18390 | | OsmC family protein | WP_001148919.1 | -0.161477589 | 0.2129096 | 0.31083788 |
| STM14_RS06600 | lolE | lipoprotein-releasing ABC transporter permease subunit LolE | WP_000168091.1 | -0.161857798 | 0.25591943 | 0.36020465 |
| STM14_RS11930 | | GntR family transcriptional regulator | WP_000434062.1 | -0.161960579 | 0.30351186 | 0.41285287 |
| STM14_RS10000 | | ATP-dependent DNA helicase | WP_000128807.1 | -0.16254806 | 0.34637592 | 0.45668106 |
| STM14_RS16670 | ripR | itaconate degradation transcriptional regulator RipR | WP_000423170.1 | -0.1634861 | 0.63976197 | 0.7324638 |
| STM14_RS16510 | metK | methionine adenosyltransferase | WP_001062140.1 | -0.164531416 | 0.09231612 | 0.15786896 |
| STM14_RS16630 | | ornithine decarboxylase | WP_000839738.1 | -0.165047269 | 0.43594048 | 0.54684415 |
| STM14_RS01810 | metQ | methionine ABC transporter substrate-binding lipoprotein MetQ | WP_000874215.1 | -0.165351147 | 0.1690674 | 0.25830131 |
| STM14_RS13380 | cysK | cysteine synthase A | WP_000036904.1 | -0.166703058 | 0.08899528 | 0.15305027 |
| STM14_RS14630 | ffh | signal recognition particle protein | WP_000460052.1 | -0.166957605 | 0.09564622 | 0.16258974 |
| STM14_RS15125 | gshA | glutamate--cysteine ligase | WP_000611821.1 | -0.167302505 | 0.08797783 | 0.15151281 |
| STM14_RS12565 | rcsC | two-component system sensor histidine kinase RcsC | WP_000876078.1 | -0.167790976 | 0.19414477 | 0.28870802 |
| STM14_RS04030 | nagB | glucosamine-6-phosphate deaminase | WP_001237059.1 | -0.168054712 | 0.18949646 | 0.28339803 |
| STM14_RS18880 | panM | aspartate 1-decarboxylase autocleavage activator PanM | WP_000778873.1 | -0.16820715 | 0.62651014 | 0.72313804 |
| STM14_RS01490 | panC | pantoate--beta-alanine ligase | WP_000905344.1 | -0.169055345 | 0.35470426 | 0.46527292 |
| STM14_RS21800 | rplK | 50S ribosomal protein L11 | WP_001085926.1 | -0.169088684 | 0.07348541 | 0.13033395 |
| STM14_RS02880 | ppiD | peptidylprolyl isomerase | WP_000969434.1 | -0.169406319 | 0.09242456 | 0.15791519 |
| STM14_RS18860 | livG | high-affinity branched-chain amino acid ABC transporter ATP-binding pr | WP_000082083.1 | -0.169650572 | 0.56737195 | 0.66990376 |
| STM14_RS06615 | potD | spermidine/putrescine ABC transporter substrate-binding protein PotD | WP_000759329.1 | -0.169865179 | 0.13897226 | 0.22065366 |
| STM14_RS05905 | yccX | acylphosphatase | WP_000072884.1 | -0.169984599 | 0.52512433 | 0.62960202 |

| | | | | | | |
|---|---|---|---|---|---|---|
| STM14_RS04635 | moaA | GTP 3'%2C8-cyclase MoaA | WP_000168180.1 | -0.170217906 | 0.23877348 | 0.34076123 |
| STM14_RS09365 | sapD | peptide ABC transporter ATP-binding protein SapD | WP_001128868.1 | -0.170601016 | 0.42596539 | 0.53657358 |
| STM14_RS07240 | | hypothetical protein | WP_000487126.1 | -0.170646898 | 0.21451754 | 0.31231854 |
| STM14_RS10665 | yecC | L-cystine ABC transporter ATP-binding protein YecC | WP_001273026.1 | -0.170660956 | 0.49883098 | 0.60657056 |
| STM14_RS13005 | | alanine/ornithine racemase family PLP-dependent enzyme | WP_000281907.1 | -0.170963002 | 0.42162638 | 0.53165354 |
| STM14_RS01535 | mrcB | bifunctional glycosyl transferase/transpeptidase | WP_000918127.1 | -0.17134735 | 0.15056462 | 0.23610687 |
| STM14_RS20660 | wzzE | ECA polysaccharide chain length modulation protein | WP_000193413.1 | -0.171841729 | 0.15743714 | 0.2447988 |
| STM14_RS06040 | iraM | anti-adapter protein IraM | WP_001537781.1 | -0.172489074 | 0.68374172 | 0.76847895 |
| STM14_RS18610 | bioH | pimeloyl-ACP methyl ester esterase BioH | WP_000998146.1 | -0.173003291 | 0.36982224 | 0.48065143 |
| STM14_RS23715 | hsdM | type I restriction-modification system methyltransferase | WP_001063191.1 | -0.17306248 | 0.24508133 | 0.34787566 |
| STM14_RS20080 | dsdX | D-serine transporter DsdX | WP_000543314.1 | -0.173112144 | 0.71225331 | 0.79146413 |
| STM14_RS21235 | fdxH | formate dehydrogenase subunit beta | WP_000331361.1 | -0.173170759 | 0.14981979 | 0.23523943 |
| STM14_RS03460 | | sigma-54-dependent transcriptional regulator | WP_001031460.1 | -0.173872309 | 0.23786862 | 0.33986505 |
| STM14_RS12325 | | DEAD/DEAH box helicase | WP_000578121.1 | -0.173881212 | 0.39807255 | 0.50857152 |
| STM14_RS11265 | | EmmdR/YeeO family multidrug/toxin efflux MATE transporter | WP_001669456.1 | -0.174423383 | 0.38914334 | 0.49994053 |
| STM14_RS09330 | pspC | envelope stress response membrane protein PspC | WP_000508938.1 | -0.174579717 | 0.35881562 | 0.46916162 |
| STM14_RS02405 | | type III restriction-modification system endonuclease | WP_001651666.1 | -0.174703643 | 0.15652644 | 0.24348558 |
| STM14_RS21805 | rplA | 50S ribosomal protein L1 | WP_001096676.1 | -0.175208679 | 0.0432558 | 0.08230897 |
| STM14_RS20370 | | fructose-specific PTS transporter subunit EIIC | WP_000162545.1 | -0.175647558 | 0.51794894 | 0.62302777 |
| STM14_RS03830 | cobD | threonine-phosphate decarboxylase CobD | WP_001173255.1 | -0.176471126 | 0.33387241 | 0.44572815 |
| STM14_RS05395 | | YcbJ family phosphotransferase | WP_000436889.1 | -0.176776769 | 0.07923462 | 0.13872652 |
| STM14_RS17360 | rsmI | 16S rRNA (cytidine(1402)-2'-O)-methyltransferase | WP_000812288.1 | -0.176777707 | 0.34840605 | 0.45897355 |
| STM14_RS13955 | suhB | inositol-1-monophosphatase | WP_000553467.1 | -0.178103463 | 0.16092679 | 0.24885865 |
| STM14_RS03740 | dcuC | anaerobic C4-dicarboxylate transporter DcuC | WP_000958610.1 | -0.178292491 | 0.62071831 | 0.71735243 |
| STM14_RS01405 | | DeoR/GlpR family DNA-binding transcription regulator | WP_000678254.1 | -0.178442428 | 0.19077766 | 0.28485131 |
| STM14_RS20425 | atpF | F0F1 ATP synthase subunit B | WP_001052212.1 | -0.178465929 | 0.06075238 | 0.1105189 |
| STM14_RS09035 | | transporter substrate-binding domain-containing protein | WP_000949370.1 | -0.178623657 | 0.48796996 | 0.59671385 |
| STM14_RS00005 | | hypothetical protein | WP_000477658.1 | -0.179098338 | 0.30209996 | 0.41145889 |
| STM14_RS05420 | mukB | chromosome partition protein MukB | WP_000572724.1 | -0.179131558 | 0.0588305 | 0.10755408 |
| STM14_RS22005 | | Na/Pi cotransporter family protein | WP_000956811.1 | -0.179521289 | 0.20084899 | 0.29711772 |
| STM14_RS14500 | trxC | thioredoxin TrxC | WP_001098732.1 | -0.180158828 | 0.19271589 | 0.28697706 |
| STM14_RS20180 | torA | trimethylamine-N-oxide reductase TorA | WP_000790665.1 | -0.180591398 | 0.64582149 | 0.73802443 |
| STM14_RS14155 | | PagK family vesicle-borne virulence factor | WP_014344379.1 | -0.180594037 | 0.3205065 | 0.43129314 |
| STM14_RS07130 | | ABC transporter substrate-binding protein | WP_000946082.1 | -0.180649762 | 0.12449529 | 0.20149084 |
| STM14_RS16215 | vapC | tRNA(fMet)-specific endonuclease VapC | WP_000911336.1 | -0.180856263 | 0.27543742 | 0.38183757 |
| STM14_RS06960 | | phage minor tail protein L | WP_001152689.1 | -0.181133131 | 0.54284037 | 0.64642246 |
| STM14_RS04515 | modF | molybdate ABC transporter ATP-binding protein ModF | WP_000096928.1 | -0.181171824 | 0.27307721 | 0.37913661 |
| STM14_RS20465 | asnA | aspartate--ammonia ligase | WP_000845121.1 | -0.181556574 | 0.43732382 | 0.54806886 |
| STM14_RS13635 | narQ | nitrate/nitrite two-component system sensor histidine kinase NarQ | WP_000216821.1 | -0.181852613 | 0.2685547 | 0.37455242 |
| STM14_RS09975 | rnd | ribonuclease D | WP_001109130.1 | -0.182370616 | 0.35390817 | 0.46461804 |
| STM14_RS22455 | nrfB | cytochrome c nitrite reductase pentaheme subunit | WP_000115065.1 | -0.183202905 | 0.46512568 | 0.57373291 |
| STM14_RS17785 | rplM | 50S ribosomal protein L13 | WP_000847559.1 | -0.183285625 | 0.03660711 | 0.07124069 |
| STM14_RS03660 | | molybdopterin-dependent oxidoreductase | WP_000064328.1 | -0.183932135 | 0.44575964 | 0.55580325 |
| STM14_RS09525 | trpCF | bifunctional indole-3-glycerol-phosphate synthase TrpC/phosphoribosy | WP_001195296.1 | -0.183957437 | 0.57600605 | 0.67715654 |
| STM14_RS00760 | | aryl-sulfate sulfotransferase | WP_001094892.1 | -0.184464972 | 0.25913113 | 0.36402932 |
| STM14_RS10375 | znuB | zinc ABC transporter permease subunit ZnuB | WP_000571508.1 | -0.184495549 | 0.64851991 | 0.7395557 |
| STM14_RS02590 | sbcC | exonuclease subunit SbcC | WP_001526312.1 | -0.184564945 | 0.21013405 | 0.30788443 |
| STM14_RS17130 | rpoD | RNA polymerase sigma factor RpoD | WP_001120159.1 | -0.185246846 | 0.02946438 | 0.05938032 |
| STM14_RS22485 | yjcO | Sel1 family TPR-like repeat protein YjcO | WP_000771454.1 | -0.185489929 | 0.25101444 | 0.35479277 |
| STM14_RS19745 | | YicC family protein | WP_000621352.1 | -0.185523909 | 0.079006 | 0.13845793 |
| STM14_RS07285 | yeaG | protein kinase YeaG | WP_001518010.1 | -0.185525052 | 0.02302598 | 0.04803493 |
| STM14_RS22805 | yjeM | glutamate/gamma-aminobutyrate family transporter YjeM | WP_000149876.1 | -0.186079355 | 0.5990507 | 0.69800781 |
| STM14_RS05330 | | DUF421 domain-containing protein | WP_000642866.1 | -0.186235213 | 0.19118917 | 0.28535002 |
| STM14_RS11555 | | SDR family oxidoreductase | WP_000754712.1 | -0.186267308 | 0.06129971 | 0.1114045 |
| STM14_RS09795 | dauA | C4-dicarboxylic acid transporter DauA | WP_001037181.1 | -0.186268041 | 0.47954782 | 0.58817291 |
| STM14_RS14490 | ung | uracil-DNA glycosylase | WP_000179978.1 | -0.186414459 | 0.1450077 | 0.2290512 |
| STM14_RS18595 | feoB | Fe(2+) transporter permease subunit FeoB | WP_000736973.1 | -0.187009577 | 0.11548073 | 0.19011718 |
| STM14_RS06180 | | phosphatase | WP_000283643.1 | -0.187167344 | 0.14120504 | 0.22371642 |
| STM14_RS19710 | rpmB | 50S ribosomal protein L28 | WP_001519051.1 | -0.187179423 | 0.04091613 | 0.07838355 |
| STM14_RS03180 | mnmH | tRNA 2-selenouridine(34) synthase MnmH | WP_001154078.1 | -0.187464949 | 0.16485075 | 0.25343652 |
| STM14_RS01505 | pcnB | polynucleotide adenylyltransferase PcnB | WP_203422695.1 | -0.187651363 | 0.21226176 | 0.31038435 |
| STM14_RS03530 | | TonB-dependent siderophore receptor | WP_001034955.1 | -0.187921385 | 0.26798786 | 0.37390349 |
| STM14_RS21825 | rpoC | DNA-directed RNA polymerase subunit beta' | WP_000653965.1 | -0.187954703 | 0.02808124 | 0.05696702 |
| STM14_RS09310 | | YcjF family protein | WP_001294459.1 | -0.188084647 | 0.26446269 | 0.3703886 |
| STM14_RS10275 | | YebY family protein | WP_000722368.1 | -0.188911828 | 0.34528704 | 0.45584327 |
| STM14_RS22920 | rnr | ribonuclease R | WP_000076357.1 | -0.189307533 | 0.0308936 | 0.06165324 |
| STM14_RS10770 | fliI | flagellar protein export ATPase FliI | WP_000213257.1 | -0.18930823 | 0.47681055 | 0.58559588 |
| STM14_RS15170 | alaS | alanine--tRNA ligase | WP_000047224.1 | -0.190215422 | 0.044952 | 0.08509679 |
| STM14_RS21095 | typA | ribosome-dependent GTPase TypA | WP_000572067.1 | -0.190757468 | 0.04263007 | 0.08124426 |
| STM14_RS15235 | norR | nitric oxide reductase transcriptional regulator NorR | WP_000010814.1 | -0.191149502 | 0.32498987 | 0.43640142 |
| STM14_RS05405 | cmoM | tRNA uridine 5-oxyacetic acid(34) methyltransferase CmoM | WP_001154025.1 | -0.191364276 | 0.28554931 | 0.39278342 |
| STM14_RS22870 | amiB | N-acetylmuramoyl-L-alanine amidase AmiB | WP_000640362.1 | -0.191878892 | 0.150789 | 0.23625749 |
| STM14_RS12755 | menD | 2-succinyl-5-enolpyruvyl-6-hydroxy-3-cyclohexene-1-carboxylic-acid sy | WP_000116392.1 | -0.192166827 | 0.19008666 | 0.28399777 |
| STM14_RS17945 | | oxidoreductase | WP_001148246.1 | -0.192574389 | 0.14261574 | 0.22546636 |
| STM14_RS03930 | gltJ | glutamate/aspartate ABC transporter permease GltJ | WP_000020930.1 | -0.192698605 | 0.30545024 | 0.41470051 |
| STM14_RS19735 | pyrE | orotate phosphoribosyltransferase | WP_000806167.1 | -0.192931195 | 0.32773399 | 0.4396053 |
| STM14_RS16740 | | MFS transporter | WP_000784371.1 | -0.193090467 | 0.07876215 | 0.13814163 |
| STM14_RS17265 | | serine dehydratase subunit alpha family protein | WP_000463090.1 | -0.193927724 | 0.15627027 | 0.24318983 |
| STM14_RS13130 | mepA | penicillin-insensitive murein endopeptidase | WP_000750428.1 | -0.19439759 | 0.43186513 | 0.54295235 |
| STM14_RS04660 | | Bax inhibitor-1/YccA family protein | WP_000373611.1 | -0.195121016 | 0.04056304 | 0.07774759 |
| STM14_RS09810 | ychF | redox-regulated ATPase YchF | WP_000505857.1 | -0.195235905 | 0.074632 | 0.13192272 |
| STM14_RS03085 | ushA | bifunctional UDP-sugar hydrolase/5'-nucleotidase UshA | WP_000670396.1 | -0.195309021 | 0.04506534 | 0.08526752 |

| | | | | | | |
|---|---|---|---|---|---|---|
| STM14_RS03055 | adk | adenylate kinase | WP_001220237.1 | -0.195317138 | 0.04429379 | 0.08406687 |
| STM14_RS14675 | | type II toxin-antitoxin system RatA family toxin | WP_001537508.1 | -0.195336318 | 0.08213441 | 0.14305529 |
| STM14_RS09955 | | YcgL domain-containing protein | WP_001525604.1 | -0.195872081 | 0.16041361 | 0.2482736 |
| STM14_RS20020 | | aldose 1-epimerase family protein | WP_000213211.1 | -0.197101324 | 0.27249769 | 0.37847472 |
| STM14_RS14040 | qseG | two-component system QseEF-associated lipoprotein QseG | WP_001054236.1 | -0.197774571 | 0.2127122 | 0.31079616 |
| STM14_RS01765 | tilS | tRNA lysidine(34) synthetase TilS | WP_000210056.1 | -0.19897075 | 0.29040411 | 0.39808115 |
| STM14_RS10030 | | cyclic diguanylate phosphodiesterase | WP_001192513.1 | -0.19920821 | 0.359345 | 0.46952033 |
| STM14_RS14850 | | hypothetical protein | WP_000061088.1 | -0.199213188 | 0.28577841 | 0.39291863 |
| STM14_RS03035 | dnaX | DNA polymerase III subunit gamma/tau | WP_000121962.1 | -0.199317857 | 0.16166681 | 0.24979321 |
| STM14_RS20755 | hemC | hydroxymethylbilane synthase | WP_001521319.1 | -0.199833763 | 0.07918178 | 0.13869996 |
| STM14_RS23210 | iolG1 | inositol 2-dehydrogenase IolG1 | WP_000172704.1 | -0.199915301 | 0.46451342 | 0.57355412 |
| STM14_RS19045 | fraE | fructose-asparagine asparaginase | WP_000702755.1 | -0.201766673 | 0.24096408 | 0.34308962 |
| STM14_RS17885 | aaeA | p-hydroxybenzoic acid efflux pump subunit AaeA | WP_000855134.1 | -0.202533903 | 0.35464312 | 0.46527292 |
| STM14_RS13970 | asrB | anaerobic sulfite reductase subunit AsrB | WP_000017587.1 | -0.203618778 | 0.42592374 | 0.53657358 |
| STM14_RS21440 | lsrC | autoinducer 2 ABC transporter permease LsrC | WP_000911134.1 | -0.204233935 | 0.09640032 | 0.16341897 |
| STM14_RS20855 | rhtC | threonine export protein RhtC | WP_000980080.1 | -0.204461695 | 0.50028949 | 0.60814325 |
| STM14_RS06920 | | phage tail protein | WP_000677089.1 | -0.20489896 | 0.54109773 | 0.64496606 |
| STM14_RS11135 | | DUF2303 family protein | WP_000080415.1 | -0.204913066 | 0.1037507 | 0.17403346 |
| STM14_RS08895 | | U32 family peptidase | WP_001249753.1 | -0.205320256 | 0.20602197 | 0.30310495 |
| STM14_RS22235 | ubiA | 4-hydroxybenzoate octaprenyltransferase | WP_000455249.1 | -0.205329525 | 0.18536408 | 0.27800837 |
| STM14_RS05875 | | DUF2057 family protein | WP_000847719.1 | -0.206075677 | 0.51018856 | 0.61732313 |
| STM14_RS14635 | | inner membrane protein YpjD | WP_001537507.1 | -0.206319249 | 0.3194147 | 0.43032745 |
| STM14_RS09945 | | YcgN family cysteine cluster protein | WP_001285703.1 | -0.206674849 | 0.22654287 | 0.32749542 |
| STM14_RS06460 | rpmF | 50S ribosomal protein L32 | WP_000290727.1 | -0.207103402 | 0.03253501 | 0.06430161 |
| STM14_RS22595 | dcuR | two-component system response regulator DcuR | WP_000611307.1 | -0.207183675 | 0.39705382 | 0.50744609 |
| STM14_RS10850 | | DUF808 domain-containing protein | WP_085983315.1 | -0.207652933 | 0.25979045 | 0.36467725 |
| STM14_RS08700 | | NAD-dependent malic enzyme | WP_000450511.1 | -0.2079208 | 0.03355492 | 0.06586482 |
| STM14_RS06530 | hinT | purine nucleoside phosphoramidase | WP_001532068.1 | -0.208975029 | 0.05120049 | 0.09516416 |
| STM14_RS24005 | trpR | trp operon repressor | WP_000192003.1 | -0.209162642 | 0.28870656 | 0.39605722 |
| STM14_RS22620 | | molecular chaperone | WP_001112218.1 | -0.209182817 | 0.07783139 | 0.13679006 |
| STM14_RS02525 | | multidrug efflux MFS transporter | WP_000446768.1 | -0.209662442 | 0.47782465 | 0.58664567 |
| STM14_RS07540 | rplT | 50S ribosomal protein L20 | WP_000124850.1 | -0.209677293 | 0.01411875 | 0.0318537 |
| STM14_RS20405 | atpD | F0F1 ATP synthase subunit beta | WP_000190499.1 | -0.210153415 | 0.02615784 | 0.05368627 |
| STM14_RS18600 | feoC | [Fe-S]-dependent transcriptional repressor FeoC | WP_000157589.1 | -0.210695402 | 0.44977736 | 0.55929762 |
| STM14_RS23930 | deoA | thymidine phosphorylase | WP_000477838.1 | -0.211898732 | 0.09014777 | 0.15467106 |
| STM14_RS05380 | | winged helix-turn-helix domain-containing protein | WP_000056911.1 | -0.211944755 | 0.38054806 | 0.49146894 |
| STM14_RS20750 | hemD | uroporphyrinogen-III synthase | WP_000025498.1 | -0.212328658 | 0.10242196 | 0.17227851 |
| STM14_RS20620 | ppiC | peptidylprolyl isomerase PpiC | WP_001096806.1 | -0.212983361 | 0.17081997 | 0.26054645 |
| STM14_RS21055 | polA | DNA polymerase I | WP_000249972.1 | -0.213480948 | 0.02722557 | 0.05559875 |
| STM14_RS21470 | tpiA | triose-phosphate isomerase | WP_001216339.1 | -0.213723627 | 0.0185687 | 0.04005269 |
| STM14_RS09635 | | YchE family NAAT transporter | WP_000615840.1 | -0.214118854 | 0.48350684 | 0.59243667 |
| STM14_RS08645 | | LacI family DNA-binding transcriptional regulator | WP_000201080.1 | -0.214268326 | 0.47357862 | 0.58201484 |
| STM14_RS18175 | rpmD | 50S ribosomal protein L30 | WP_001140434.1 | -0.215008168 | 0.0780199 | 0.13705595 |
| STM14_RS02865 | clpX | ATP-dependent protease ATP-binding subunit ClpX | WP_000130314.1 | -0.215169465 | 0.01042454 | 0.0247155 |
| STM14_RS04825 | | ABC-F family ATPase | WP_000961479.1 | -0.215602683 | 0.04883681 | 0.0913241 |
| STM14_RS04350 | tolQ | Tol-Pal system protein TolQ | WP_000131318.1 | -0.217027771 | 0.18126648 | 0.27331007 |
| STM14_RS08035 | | alkene reductase | WP_000092932.1 | -0.21758725 | 0.05026353 | 0.0936117 |
| STM14_RS12930 | | PTS sugar transporter subunit IIA | WP_000901812.1 | -0.217648813 | 0.25413545 | 0.35851599 |
| STM14_RS14625 | rpsP | 30S ribosomal protein S16 | WP_000256453.1 | -0.217956305 | 0.02734335 | 0.05574652 |
| STM14_RS08195 | fumB | class I fumarate hydratase | WP_000066601.1 | -0.218688298 | 0.01096467 | 0.02576383 |
| STM14_RS17150 | | PadR family transcriptional regulator | WP_000983441.1 | -0.218735429 | 0.09703174 | 0.16418697 |
| STM14_RS15265 | hycI | hydrogenase maturation peptidase HycI | WP_000132984.1 | -0.219355321 | 0.18158592 | 0.27367965 |
| STM14_RS15090 | proX | glycine betaine/L-proline ABC transporter substrate-binding protein Pro | WP_001216622.1 | -0.220526552 | 0.05831929 | 0.10672547 |
| STM14_RS13980 | | nickel/cobalt transporter | WP_000111027.1 | -0.220971012 | 0.46173722 | 0.57108378 |
| STM14_RS02800 | | MFS transporter | WP_001000938.1 | -0.221089728 | 0.23372006 | 0.33576171 |
| STM14_RS10440 | cmoB | tRNA 5-methoxyuridine(34)/uridine 5-oxyacetic acid(34) synthase Cmo | WP_000569041.1 | -0.221089976 | 0.22724768 | 0.32812782 |
| STM14_RS04540 | modC | molybdenum ABC transporter ATP-binding protein ModC | WP_000891710.1 | -0.221233593 | 0.12387411 | 0.20083861 |
| STM14_RS08365 | speG | spermidine N1-acetyltransferase | WP_000199997.1 | -0.221663177 | 0.0489989 | 0.09148781 |
| STM14_RS17530 | ftsH | ATP-dependent zinc metalloprotease FtsH | WP_001107481.1 | -0.22245284 | 0.00817715 | 0.02016629 |
| STM14_RS07780 | ttrB | tetrathionate reductase subunit TtrB | WP_000269830.1 | -0.2225193 | 0.35720804 | 0.46788689 |
| STM14_RS19290 | | molybdopterin guanine dinucleotide-containing S/N-oxide reductase | WP_000148453.1 | -0.222965891 | 0.12305571 | 0.19968758 |
| STM14_RS22480 | gltP | glutamate/aspartate:proton symporter GltP | WP_000793278.1 | -0.223000074 | 0.05747074 | 0.1053821 |
| STM14_RS20790 | | lipoprotein | WP_000799903.1 | -0.223456504 | 0.14787266 | 0.23277774 |
| STM14_RS14505 | tapT | tRNA-uridine aminocarboxypropyltransferase | WP_000183639.1 | -0.223513717 | 0.46497772 | 0.57373291 |
| STM14_RS08805 | | YncE family protein | WP_000550636.1 | -0.223737562 | 0.10780323 | 0.17920158 |
| STM14_RS23915 | | YjjW family glycine radical enzyme activase | WP_001064938.1 | -0.224173563 | 0.28316471 | 0.39049191 |
| STM14_RS04505 | galE | UDP-glucose 4-epimerase GalE | WP_001265465.1 | -0.225133743 | 0.08668953 | 0.14957397 |
| STM14_RS05175 | clpA | ATP-dependent Clp protease ATP-binding subunit ClpA | WP_000934063.1 | -0.225720304 | 0.00746508 | 0.01857453 |
| STM14_RS20705 | wzyE | ECA oligosaccharide polymerase | WP_000055609.1 | -0.225965009 | 0.15864829 | 0.24636989 |
| STM14_RS18315 | tusC | sulfurtransferase complex subunit TusC | WP_000820704.1 | -0.226209765 | 0.38618242 | 0.4971761 |
| STM14_RS04020 | nagC | DNA-binding transcriptional regulator NagC | WP_000187577.1 | -0.226286446 | 0.11374967 | 0.18773029 |
| STM14_RS13545 | eutJ | ethanolamine utilization protein EutJ | WP_000929675.1 | -0.226556933 | 0.50083955 | 0.60861097 |
| STM14_RS12845 | nuoB | NADH-quinone oxidoreductase subunit NuoB | WP_000386728.1 | -0.226653389 | 0.03559289 | 0.0694873 |
| STM14_RS18965 | | AI-2E family transporter | WP_000179589.1 | -0.226892902 | 0.41911078 | 0.52971462 |
| STM14_RS17080 | glnE | bifunctional [glutamate--ammonia ligase]-adenylyl-L-tyrosine phosphor | WP_000188309.1 | -0.227186531 | 0.07152069 | 0.127217 |
| STM14_RS21980 | aceA | isocitrate lyase | WP_000857881.1 | -0.227754504 | 0.0572344 | 0.10505337 |
| STM14_RS17455 | pnp | polyribonucleotide nucleotidyltransferase | WP_001670767.1 | -0.22812571 | 0.01342296 | 0.03058376 |
| STM14_RS13925 | hscB | co-chaperone HscB | WP_000384396.1 | -0.228690223 | 0.12097078 | 0.19688457 |
| STM14_RS17595 | ibaG | BolA family iron metabolism protein IbaG | WP_001518349.1 | -0.229286412 | 0.30216862 | 0.41145889 |
| STM14_RS01700 | rseP | sigma E protease regulator RseP | WP_000949017.1 | -0.229516747 | 0.03747319 | 0.07267325 |
| STM14_RS09940 | | hypothetical protein | WP_001519337.1 | -0.230058057 | 0.11639416 | 0.19123753 |
| STM14_RS13850 | | bifunctional tRNA (adenosine(37)-C2)-methyltransferase TrmG/riboson | WP_000003211.1 | -0.230141248 | 0.06970605 | 0.12440992 |

| | | | | | | |
|---|---|---|---|---|---|---|
| STM14_RS01635 | cdaR | DNA-binding transcriptional regulator CdaR | WP_000929420.1 | -0.230922456 | 0.21836637 | 0.31716962 |
| STM14_RS16550 | | IclR family transcriptional regulator | WP_000098329.1 | -0.231332347 | 0.57303944 | 0.67510284 |
| STM14_RS22345 | ssb1 | single-stranded DNA-binding protein SSB1 | WP_000168322.1 | -0.231548067 | 0.02249183 | 0.04713428 |
| STM14_RS15735 | cas6e | type I-E CRISPR-associated protein Cas6/Cse3/CasE | WP_000281486.1 | -0.231690712 | 0.35820239 | 0.46869269 |
| STM14_RS22800 | epmA | elongation factor P--(R)-beta-lysine ligase | WP_000004794.1 | -0.232435314 | 0.15587102 | 0.24296761 |
| STM14_RS17485 | rimP | ribosome maturation factor RimP | WP_000105461.1 | -0.232446678 | 0.06474916 | 0.11680862 |
| STM14_RS21280 | rhaM | L-rhamnose mutarotase | WP_000619478.1 | -0.232760717 | 0.33571786 | 0.44770205 |
| STM14_RS20950 | ubiD | 4-hydroxy-3-polyprenylbenzoate decarboxylase | WP_000339764.1 | -0.233060684 | 0.04514564 | 0.08533174 |
| STM14_RS17560 | cgtA | Obg family GTPase CgtA | WP_000673556.1 | -0.233727425 | 0.04728923 | 0.08892694 |
| STM14_RS05365 | | ComEC family protein | WP_000705784.1 | -0.233819732 | 0.39148449 | 0.50224595 |
| STM14_RS02105 | | fimbrial biogenesis outer membrane usher protein | WP_000669634.1 | -0.234128676 | 0.15594377 | 0.24296761 |
| STM14_RS03075 | ybaL | YbaL family putative K(+) efflux transporter | WP_000546207.1 | -0.234595505 | 0.07510875 | 0.1325745 |
| STM14_RS13835 | hisS | histidine--tRNA ligase | WP_001107141.1 | -0.235260301 | 0.01937906 | 0.04158141 |
| STM14_RS02925 | | SmdB family multidrug efflux ABC transporter permease/ATP-binding pr | WP_001256068.1 | -0.235414088 | 0.28259984 | 0.39015096 |
| STM14_RS17960 | accB | acetyl-CoA carboxylase biotin carboxyl carrier protein | WP_000354626.1 | -0.235530931 | 0.01184377 | 0.02751342 |
| STM14_RS14585 | | SMP-30/gluconolactonase/LRE family protein | WP_000210995.1 | -0.235683532 | 0.08402546 | 0.14593479 |
| STM14_RS17855 | mdh | malate dehydrogenase | WP_000861586.1 | -0.235962133 | 0.00989308 | 0.02366882 |
| STM14_RS12980 | | histidine ABC transporter permease HisQ | WP_000965498.1 | -0.236198789 | 0.24931067 | 0.35292652 |
| STM14_RS15845 | gudD | glucarate dehydratase | WP_000107113.1 | -0.236440612 | 0.14366746 | 0.22703158 |
| STM14_RS08105 | pdxY | pyridoxal kinase PdxY | WP_000789732.1 | -0.236788428 | 0.153433 | 0.23958452 |
| STM14_RS07460 | tcyP | cystine/sulfocysteine:cation symporter | WP_001010662.1 | -0.237216444 | 0.25009538 | 0.35362949 |
| STM14_RS07300 | | aldo/keto reductase | WP_000843751.1 | -0.237262014 | 0.13861364 | 0.22027424 |
| STM14_RS19165 | bcsE | cellulose biosynthesis c-di-GMP-binding protein BcsE | WP_001205771.1 | -0.239033902 | 0.06247392 | 0.11330129 |
| STM14_RS11520 | | thiosulfate reductase cytochrome B subunit | WP_001092547.1 | -0.239198379 | 0.04234349 | 0.08078173 |
| STM14_RS22275 | | conjugal transfer protein TraF | WP_001039339.1 | -0.23934401 | 0.10048005 | 0.16939906 |
| STM14_RS10420 | | hydrolase | WP_000916144.1 | -0.239583338 | 0.16257839 | 0.25078074 |
| STM14_RS09430 | yciH | stress response translation initiation inhibitor YciH | WP_001285199.1 | -0.239818403 | 0.37811928 | 0.48938921 |
| STM14_RS23195 | iolA | malonate-semialdehyde dehydrogenase IolA | WP_000450729.1 | -0.240137753 | 0.12631813 | 0.20372463 |
| STM14_RS03690 | rna | ribonuclease I | WP_001092580.1 | -0.240163179 | 0.015656 | 0.03472615 |
| STM14_RS01035 | pdxA | 4-hydroxythreonine-4-phosphate dehydrogenase PdxA | WP_000093011.1 | -0.240253504 | 0.17124821 | 0.26087543 |
| STM14_RS13720 | uraA | uracil permease | WP_000198335.1 | -0.240808209 | 0.17098689 | 0.26069306 |
| STM14_RS06185 | | molecular chaperone | WP_001001883.1 | -0.240899837 | 0.06584947 | 0.11837348 |
| STM14_RS16875 | metC | cystathionine beta-lyase | WP_000130262.1 | -0.241170942 | 0.10625397 | 0.17726648 |
| STM14_RS01650 | dapD | 2%2C3%2C4%2C5-tetrahydropyridine-2%2C6-dicarboxylate N-succiny | WP_001186675.1 | -0.241688899 | 0.01948899 | 0.04176861 |
| STM14_RS06255 | mdoH | glucans biosynthesis glucosyltransferase MdoH | WP_001044610.1 | -0.241807714 | 0.03506372 | 0.06852687 |
| STM14_RS11665 | rfbF | glucose-1-phosphate cytidylyltransferase | WP_000648784.1 | -0.242175947 | 0.02164013 | 0.04563513 |
| STM14_RS10950 | | DNA circularization N-terminal domain-containing protein | WP_000863818.1 | -0.243067406 | 0.20323513 | 0.30004481 |
| STM14_RS26640 | | hypothetical protein | WP_000029029.1 | -0.243510586 | 0.1673277 | 0.25628145 |
| STM14_RS13390 | ptsH | phosphocarrier protein Hpr | WP_000487600.1 | -0.243768987 | 0.00745691 | 0.01857453 |
| STM14_RS02630 | | peroxiredoxin | WP_000244731.1 | -0.24377857 | 0.00287861 | 0.00814683 |
| STM14_RS01720 | fabZ | 3-hydroxyacyl-ACP dehydratase FabZ | WP_000210741.1 | -0.243904956 | 0.05623201 | 0.1033165 |
| STM14_RS09085 | | GFA family protein | WP_001521629.1 | -0.244518379 | 0.15271723 | 0.23866928 |
| STM14_RS23650 | | YjiG family protein | WP_000211979.1 | -0.245120002 | 0.11646922 | 0.19127549 |
| STM14_RS09395 | fabI | enoyl-ACP reductase FabI | WP_000506505.1 | -0.24653003 | 0.00909624 | 0.02196221 |
| STM14_RS22875 | mutL | DNA mismatch repair endonuclease MutL | WP_001122537.1 | -0.247541135 | 0.08497292 | 0.14723029 |
| STM14_RS10955 | | phage tail tape measure protein | WP_000785385.1 | -0.247762097 | 0.06701668 | 0.12007562 |
| STM14_RS17415 | | peptidase U32 family protein | WP_000421309.1 | -0.247854825 | 0.11404861 | 0.18813933 |
| STM14_RS03840 | holA | DNA polymerase III subunit delta | WP_000620491.1 | -0.247874559 | 0.10274738 | 0.17266811 |
| STM14_RS17175 | rlmG | 23S rRNA (guanine(1835)-N(2))-methyltransferase RlmG | WP_000019989.1 | -0.247996488 | 0.2060488 | 0.30310495 |
| STM14_RS14670 | | RnfH family protein | WP_001112990.1 | -0.248014956 | 0.07294104 | 0.1294932 |
| STM14_RS18820 | ugpQ | glycerophosphodiester phosphodiesterase | WP_000073548.1 | -0.248203987 | 0.31643519 | 0.42709471 |
| STM14_RS15740 | cas5e | type I-E CRISPR-associated protein Cas5/CasD | WP_000085114.1 | -0.24925187 | 0.35136455 | 0.46187942 |
| STM14_RS02690 | | DUF3251 domain-containing protein | WP_000198484.1 | -0.249356815 | 0.17633317 | 0.26711769 |
| STM14_RS12090 | | fumarylacetoacetate hydrolase family protein | WP_000170621.1 | -0.24937496 | 0.3550197 | 0.46552085 |
| STM14_RS01140 | leuB | 3-isopropylmalate dehydrogenase | WP_000042325.1 | -0.249433669 | 0.33595893 | 0.44770205 |
| STM14_RS19580 | grxC | glutaredoxin 3 | WP_001273795.1 | -0.249778165 | 0.01439221 | 0.03239127 |
| STM14_RS13195 | pgtA | two-component system response regulator PgtA | WP_000930843.1 | -0.249796083 | 0.09839076 | 0.16633368 |
| STM14_RS05900 | | helix-turn-helix domain-containing protein | WP_000548082.1 | -0.250020291 | 0.21209523 | 0.31026406 |
| STM14_RS04420 | | YbgS-like family protein | WP_000784380.1 | -0.250701792 | 0.00591431 | 0.01517525 |
| STM14_RS21885 | nudC | NAD(+) diphosphatase | WP_000373966.1 | -0.251087964 | 0.07932392 | 0.13881686 |
| STM14_RS20320 | yidZ | HTH-type transcriptional regulator YidZ | WP_000749366.1 | -0.252176954 | 0.34386134 | 0.45469748 |
| STM14_RS23735 | | BrnA antitoxin family protein | WP_000063142.1 | -0.252267917 | 0.17636152 | 0.26711769 |
| STM14_RS01390 | | 2-keto-3-deoxygluconate permease 1 | WP_001022089.1 | -0.252338385 | 0.36297551 | 0.47342395 |
| STM14_RS03545 | entF | enterobactin non-ribosomal peptide synthetase EntF | WP_000194139.1 | -0.252533456 | 0.28489734 | 0.3922932 |
| STM14_RS23840 | fhuF | siderophore-iron reductase FhuF | WP_000331539.1 | -0.252596582 | 0.34323662 | 0.45457298 |
| STM14_RS10640 | uvrC | excinuclease ABC subunit UvrC | WP_001289464.1 | -0.252770579 | 0.03177 | 0.06306045 |
| STM14_RS00245 | | DNA-binding protein | WP_001247118.1 | -0.252823649 | 0.0253548 | 0.05212528 |
| STM14_RS06565 | | glycine zipper 2TM domain-containing protein | WP_001043449.1 | -0.252857144 | 0.13461224 | 0.21502918 |
| STM14_RS01345 | aceE | pyruvate dehydrogenase (acetyl-transferring)%2C homodimeric type | WP_000003852.1 | -0.253105988 | 0.0154881 | 0.03452008 |
| STM14_RS01770 | rof | Rho-binding antiterminator | WP_000062330.1 | -0.253136337 | 0.09979551 | 0.1683992 |
| STM14_RS22685 | | transcriptional regulator | WP_001188509.1 | -0.253235939 | 0.03882969 | 0.07489309 |
| STM14_RS04375 | cpoB | cell division protein CpoB | WP_000097552.1 | -0.253483625 | 0.01363271 | 0.03096584 |
| STM14_RS06560 | ndh | NADH-quinone dehydrogenase | WP_000211053.1 | -0.25356155 | 0.01279981 | 0.02940044 |
| STM14_RS19305 | | DUF3053 domain-containing protein | WP_000190524.1 | -0.253656052 | 0.01971529 | 0.04215545 |
| STM14_RS03670 | | dimethyl sulfoxide reductase anchor subunit family protein | WP_000417097.1 | -0.253784904 | 0.35699266 | 0.46777472 |
| STM14_RS18895 | rpoH | RNA polymerase sigma factor RpoH | WP_000159621.1 | -0.253925548 | 0.00325522 | 0.00908698 |
| STM14_RS17085 | | inorganic triphosphatase | WP_000046230.1 | -0.254483666 | 0.0935414 | 0.15937966 |
| STM14_RS16260 | recJ | single-stranded-DNA-specific exonuclease RecJ | WP_000813394.1 | -0.254677344 | 0.02358735 | 0.04895639 |
| STM14_RS13990 | csiE | stationary phase inducible protein CsiE | WP_000982961.1 | -0.254987427 | 0.03719399 | 0.0722301 |
| STM14_RS15585 | mutS | DNA mismatch repair protein MutS | WP_001005807.1 | -0.255537364 | 0.03948633 | 0.0760799 |
| STM14_RS23020 | | L-ribulose-5-phosphate 4-epimerase | WP_001170771.1 | -0.256080647 | 0.39950494 | 0.51013667 |
| STM14_RS23875 | | DNA polymerase III subunit psi | WP_000203998.1 | -0.256352796 | 0.36013046 | 0.47021289 |

| | | | | | | |
|---|---|---|---|---|---|---|
| STM14_RS20110 | ibpB | heat shock chaperone IbpB | WP_001246919.1 | -0.256355342 | 0.31137714 | 0.421349 |
| STM14_RS07470 | hxpB | hexitol phosphatase HxpB | WP_000106848.1 | -0.256483293 | 0.09353099 | 0.15937966 |
| STM14_RS21925 | zraR | sigma-54-dependent response regulator transcription factor ZraR | WP_000617942.1 | -0.256536829 | 0.04938469 | 0.09206807 |
| STM14_RS13745 | ppx | exopolyphosphatase | WP_001123330.1 | -0.256692955 | 0.08541189 | 0.14771563 |
| STM14_RS05640 | | phage portal protein | WP_001009207.1 | -0.256822876 | 0.2992142 | 0.40803952 |
| STM14_RS07640 | | acyl-CoA dehydrogenase | WP_000347819.1 | -0.256909967 | 0.18791453 | 0.28114641 |
| STM14_RS23135 | | gamma-glutamylcyclotransferase | WP_001219166.1 | -0.257169584 | 0.12752474 | 0.20549062 |
| STM14_RS10865 | | DNA cytosine methyltransferase | WP_001157322.1 | -0.257583819 | 0.06378762 | 0.11529995 |
| STM14_RS08795 | ansP | L-asparagine permease | WP_000857101.1 | -0.257895438 | 0.19549809 | 0.29036869 |
| STM14_RS10070 | rlmA | 23S rRNA (guanine(745)-N(1))-methyltransferase | WP_000010923.1 | -0.258341261 | 0.23922623 | 0.34119374 |
| STM14_RS12170 | folE | GTP cyclohydrolase I FolE | WP_001139611.1 | -0.258821569 | 0.06520995 | 0.11746725 |
| STM14_RS02660 | secF | protein translocase subunit SecF | WP_000046629.1 | -0.2593327 | 0.05056929 | 0.09408596 |
| STM14_RS13155 | fadJ | fatty acid oxidation complex subunit alpha FadJ | WP_000214154.1 | -0.259613679 | 0.03074599 | 0.06142525 |
| STM14_RS02655 | secD | protein translocase subunit SecD | WP_000934811.1 | -0.260672775 | 0.01030353 | 0.02449167 |
| STM14_RS20415 | atpA | F0F1 ATP synthase subunit alpha | WP_001176751.1 | -0.260822611 | 0.00455557 | 0.01202408 |
| STM14_RS17535 | rlmE | 23S rRNA (uridine(2552)-2'-O)-methyltransferase RlmE | WP_000145974.1 | -0.26090864 | 0.01400767 | 0.03168074 |
| STM14_RS22210 | malE | maltose/maltodextrin ABC transporter substrate-binding protein MalE | WP_000695417.1 | -0.261288907 | 0.05602804 | 0.10304458 |
| STM14_RS02595 | sbcD | exonuclease subunit SbcD | WP_001221261.1 | -0.261361265 | 0.25996464 | 0.3647827 |
| STM14_RS02670 | | YafY family protein | WP_000198318.1 | -0.261519832 | 0.30739939 | 0.41688565 |
| STM14_RS17755 | nanA | N-acetylneuraminate lyase | WP_001029667.1 | -0.261555653 | 0.11140595 | 0.18435806 |
| STM14_RS22855 | queG | tRNA epoxyqueuosine(34) reductase QueG | WP_001670426.1 | -0.262117603 | 0.07429188 | 0.13138458 |
| STM14_RS12050 | pbpG | D-alanyl-D-alanine endopeptidase | WP_000911555.1 | -0.262212302 | 0.17643754 | 0.26712295 |
| STM14_RS18135 | rplQ | 50S ribosomal protein L17 | WP_001216370.1 | -0.262236151 | 0.01525336 | 0.03410011 |
| STM14_RS24015 | gpmB | 2%2C3-diphosphoglycerate-dependent phosphoglycerate mutase Gpm | WP_000942363.1 | -0.262430038 | 0.03147374 | 0.0626074 |
| STM14_RS17765 | sspB | ClpXP protease specificity-enhancing factor | WP_000366112.1 | -0.262899919 | 0.03973864 | 0.07648598 |
| STM14_RS19870 | | isochorismatase family protein | WP_000106461.1 | -0.262958047 | 0.03096594 | 0.06176413 |
| STM14_RS15895 | sdaB | L-serine ammonia-lyase II | WP_000626381.1 | -0.262971358 | 0.01910465 | 0.04104044 |
| STM14_RS16300 | | hemolysin III family protein | WP_000250289.1 | -0.263494873 | 0.03747605 | 0.07267325 |
| STM14_RS10305 | ptrB | oligopeptidase B | WP_000936999.1 | -0.263520236 | 0.00669541 | 0.01690843 |
| STM14_RS05965 | | heavy metal sensor histidine kinase | WP_000240046.1 | -0.264274131 | 0.37738162 | 0.48892298 |
| STM14_RS20325 | | 4'-phosphopantetheinyl transferase superfamily protein | WP_000190678.1 | -0.26455794 | 0.08608748 | 0.14881413 |
| STM14_RS04885 | gsiC | glutathione ABC transporter permease GsiC | WP_000936064.1 | -0.265211055 | 0.40890746 | 0.52006814 |
| STM14_RS10310 | | tellurite resistance TerB family protein | WP_000024706.1 | -0.267128994 | 0.06338347 | 0.11468204 |
| STM14_RS11800 | | TerC family protein | WP_000454713.1 | -0.267449907 | 0.01273468 | 0.02926909 |
| STM14_RS21175 | | DeoR/GlpR family DNA-binding transcription regulator | WP_000059693.1 | -0.267853614 | 0.01846671 | 0.03985606 |
| STM14_RS18785 | yhhY | N-acetyltransferase | WP_001290276.1 | -0.267890055 | 0.42162122 | 0.53165354 |
| STM14_RS02435 | foxA | ferrioxamine B receptor FoxA | WP_001127725.1 | -0.268114586 | 0.326438 | 0.43818618 |
| STM14_RS02180 | | RNA ligase RtcB family protein | | -0.268269024 | 0.34260783 | 0.45409721 |
| STM14_RS15710 | cysN | sulfate adenylyltransferase subunit CysN | WP_001092263.1 | -0.268711931 | 0.22897547 | 0.33023411 |
| STM14_RS23790 | | SIS domain-containing protein | WP_000900312.1 | -0.268834461 | 0.18477029 | 0.27723073 |
| STM14_RS04295 | sucA | 2-oxoglutarate dehydrogenase E1 component | WP_001181501.1 | -0.268894225 | 0.00103636 | 0.00330952 |
| STM14_RS18910 | ftsY | signal recognition particle-docking protein FtsY | WP_001118686.1 | -0.269183525 | 0.01146086 | 0.02679293 |
| STM14_RS03090 | ybaK | Cys-tRNA(Pro)/Cys-tRNA(Cys) deacylase YbaK | WP_000186618.1 | -0.269340662 | 0.14989518 | 0.23525748 |
| STM14_RS01060 | rapA | RNA polymerase-associated protein RapA | WP_001116976.1 | -0.269712211 | 0.05783087 | 0.10588427 |
| STM14_RS02460 | prpC | 2-methylcitrate synthase | WP_000132749.1 | -0.269879203 | 0.01649261 | 0.03630948 |
| STM14_RS03780 | lipB | lipoyl(octanoyl) transferase LipB | WP_000284009.1 | -0.27054303 | 0.20252736 | 0.29935998 |
| STM14_RS15255 | hydN | electron transport protein HydN | WP_001078791.1 | -0.271125659 | 0.34641776 | 0.45668106 |
| STM14_RS10415 | aspS | aspartate--tRNA ligase | WP_001258636.1 | -0.27208409 | 0.00693423 | 0.01742786 |
| STM14_RS03415 | | efflux RND transporter permease subunit | | -0.272148674 | 0.45087052 | 0.56046768 |
| STM14_RS21660 | ppc | phosphoenolpyruvate carboxylase | WP_001005548.1 | -0.272922398 | 0.0073709 | 0.01841225 |
| STM14_RS13480 | | GNAT family acetyltransferase | WP_000406008.1 | -0.273000405 | 0.05479526 | 0.10123239 |
| STM14_RS05025 | potF | spermidine/putrescine ABC transporter substrate-binding protein PotF | WP_000125769.1 | -0.273205362 | 0.06587378 | 0.11837348 |
| STM14_RS08380 | rspA | starvation-sensing protein RspA | WP_000706264.1 | -0.273296866 | 0.24855443 | 0.35199131 |
| STM14_RS05340 | aroA | 3-phosphoshikimate 1-carboxyvinyltransferase | WP_000445178.1 | -0.27431611 | 0.04773725 | 0.08958642 |
| STM14_RS01145 | leuA | 2-isopropylmalate synthase | WP_000082819.1 | -0.274641699 | 0.19267224 | 0.28697706 |
| STM14_RS19275 | | autotransporter outer membrane beta-barrel domain-containing protein | WP_000637070.1 | -0.27548811 | 0.0463689 | 0.08746428 |
| STM14_RS19600 | | divergent polysaccharide deacetylase family protein | WP_001135518.1 | -0.275547296 | 0.11927567 | 0.19470492 |
| STM14_RS16010 | recB | exodeoxyribonuclease V subunit beta | WP_001000527.1 | -0.275781962 | 0.03300853 | 0.0649775 |
| STM14_RS08395 | | mannitol dehydrogenase family protein | WP_001181258.1 | -0.276898299 | 0.16317968 | 0.25149752 |
| STM14_RS18580 | greB | transcription elongation factor GreB | WP_000856693.1 | -0.277348933 | 0.23713407 | 0.3394742 |
| STM14_RS10425 | | DUF72 domain-containing protein | WP_000639226.1 | -0.277440274 | 0.41595047 | 0.52647976 |
| STM14_RS20195 | torT | TMAO reductase system periplasmic protein TorT | WP_001202030.1 | -0.277523892 | 0.13909944 | 0.2206653 |
| STM14_RS20240 | | mandelate racemase/muconate lactonizing enzyme family protein | WP_001687378.1 | -0.277668201 | 0.16213312 | 0.25019872 |
| STM14_RS10025 | sdaA | L-serine ammonia-lyase | WP_001738311.1 | -0.277821829 | 0.05494652 | 0.10146092 |
| STM14_RS18070 | aroE | shikimate dehydrogenase | WP_000451193.1 | -0.277903724 | 0.0342891 | 0.06719131 |
| STM14_RS18290 | tuf | elongation factor Tu | WP_000031748.1 | -0.277941628 | 0.00102299 | 0.00327533 |
| STM14_RS09890 | ldcA | muramoyltetrapeptide carboxypeptidase | WP_000051601.1 | -0.277984318 | 0.11052896 | 0.18323622 |
| STM14_RS00525 | | complement resistance protein TraT | WP_000782438.1 | -0.278266419 | 0.00211091 | 0.0062179 |
| STM14_RS09070 | | YdcF family protein | WP_001098498.1 | -0.278354039 | 0.0272486 | 0.05561494 |
| STM14_RS21640 | | [formate-C-acetyltransferase]-activating enzyme | WP_000171724.1 | -0.278561089 | 0.04795771 | 0.08990851 |
| STM14_RS10870 | | phosphohydrolase | WP_000377041.1 | -0.278642937 | 0.20884833 | 0.30648846 |
| STM14_RS17910 | rng | ribonuclease G | WP_000123188.1 | -0.278794345 | 0.01641762 | 0.03617575 |
| STM14_RS19025 | prlC | oligopeptidase A | WP_000184178.1 | -0.278942945 | 0.02268482 | 0.04743073 |
| STM14_RS17395 | | YhbP family protein | WP_000380404.1 | -0.279042697 | 0.09875568 | 0.16687399 |
| STM14_RS14890 | iroC | salmochelin/enterobactin export ABC transporter IroC | WP_001111808.1 | -0.279077935 | 0.38574259 | 0.49695739 |
| STM14_RS11360 | | cobalt-precorrin-7 (C(5))-methyltransferase | WP_000958833.1 | -0.279155272 | 0.3193776 | 0.43032745 |
| STM14_RS22490 | | formate dehydrogenase H subunit alpha%2C selenocysteine-containing | WP_014344263.1 | -0.279768023 | 0.03692291 | 0.07177939 |
| STM14_RS06905 | | major capsid protein | WP_000522566.1 | -0.280584459 | 0.28557339 | 0.39278342 |
| STM14_RS04970 | | MFS transporter | WP_001181284.1 | -0.280657748 | 0.30509569 | 0.41437194 |
| STM14_RS08130 | rsxG | electron transport complex subunit RsxG | WP_000920822.1 | -0.282086234 | 0.23724014 | 0.33949405 |
| STM14_RS23190 | iolB | 5-deoxy-glucuronate isomerase | WP_000002010.1 | -0.282334625 | 0.1126263 | 0.18604308 |
| STM14_RS08995 | | VOC family protein | WP_000024095.1 | -0.282635351 | 0.00959977 | 0.02304194 |

| | | | | | | |
|---|---|---|---|---|---|---|
| STM14_RS12210 | yieE | DNA-binding transcriptional regulator YeiE | WP_000548308.1 | -0.283307658 | 0.01071795 | 0.02534585 |
| STM14_RS10330 | kdgA | bifunctional 4-hydroxy-2-oxoglutarate aldolase/2-dehydro-3-deoxy-pho | WP_000800497.1 | -0.284643676 | 0.03292594 | 0.06493481 |
| STM14_RS07705 | sufB | Fe-S cluster assembly protein SufB | WP_000089383.1 | -0.284875682 | 0.01090294 | 0.02566792 |
| STM14_RS12750 | menH | 2-succinyl-6-hydroxy-2%2C4-cyclohexadiene-1-carboxylate synthase | WP_000979146.1 | -0.285619108 | 0.10208594 | 0.17179178 |
| STM14_RS20975 | pepQ | Xaa-Pro dipeptidase | WP_000444529.1 | -0.285854781 | 0.01248116 | 0.02877622 |
| STM14_RS24000 | sltY | murein transglycosylase | WP_000373268.1 | -0.285905908 | 0.04225009 | 0.08064533 |
| STM14_RS12195 | | MFS transporter | WP_000023807.1 | -0.286010665 | 0.47824085 | 0.58696094 |
| STM14_RS17000 | nudF | ADP-ribose diphosphatase | WP_001231952.1 | -0.286265882 | 0.1305291 | 0.20950661 |
| STM14_RS14120 | recO | DNA repair protein RecO | WP_000399380.1 | -0.286386992 | 0.20478495 | 0.30183293 |
| STM14_RS04765 | rlmF | 23S rRNA (adenine(1618)-N(6))-methyltransferase RlmF | WP_001275962.1 | -0.286747754 | 0.10570071 | 0.17658349 |
| STM14_RS02195 | gpt | xanthine phosphoribosyltransferase | WP_001292018.1 | -0.287001723 | 0.04820888 | 0.09028744 |
| STM14_RS00250 | | hypothetical protein | WP_001541564.1 | -0.287130905 | 0.20728819 | 0.30480635 |
| STM14_RS03945 | corC | CNNM family magnesium/cobalt transport protein CorC | WP_001278615.1 | -0.287278578 | 0.01066715 | 0.02524192 |
| STM14_RS04960 | deoR | DNA-binding transcriptional repressor DeoR | WP_000450103.1 | -0.287899993 | 0.16500109 | 0.25356178 |
| STM14_RS15935 | fucR | L-fucose operon activator | WP_000642330.1 | -0.288394434 | 0.15206991 | 0.23785956 |
| STM14_RS07685 | menI | 1%2C4-dihydroxy-2-naphthoyl-CoA hydrolase | WP_000637930.1 | -0.288468987 | 0.21606894 | 0.31408047 |
| STM14_RS06500 | yceG | cell division protein YceG | WP_000736475.1 | -0.288609626 | 0.07223782 | 0.12836856 |
| STM14_RS13300 | gltX | glutamate--tRNA ligase | WP_000695623.1 | -0.288622948 | 0.00948877 | 0.02280526 |
| STM14_RS19700 | mutM | bifunctional DNA-formamidopyrimidine glycosylase/DNA-(apurinic or a | WP_001114508.1 | -0.289144592 | 0.19667422 | 0.29188009 |
| STM14_RS13700 | | AI-2E family transporter | WP_000892056.1 | -0.290892456 | 0.09632679 | 0.1633846 |
| STM14_RS22880 | miaA | tRNA (adenosine(37)-N6)-dimethylallyltransferase MiaA | WP_001000734.1 | -0.290908424 | 0.00145143 | 0.00446462 |
| STM14_RS23130 | tamB | autotransporter assembly complex protein TamB | WP_000060815.1 | -0.291430556 | 0.00473531 | 0.01244497 |
| STM14_RS07245 | | DUF488 domain-containing protein | WP_001222527.1 | -0.291483694 | 0.08431687 | 0.14624432 |
| STM14_RS22975 | bsmA | biofilm peroxide resistance protein BsmA | WP_000586836.1 | -0.291537039 | 0.0048501 | 0.0126923 |
| STM14_RS15855 | gudP | galactarate/glucarate/glycerate transporter GudP | WP_000097015.1 | -0.291735137 | 0.23950847 | 0.34128104 |
| STM14_RS02935 | amtB | ammonium transporter AmtB | WP_000684912.1 | -0.292100816 | 0.28275762 | 0.39022248 |
| STM14_RS10300 | exoX | exodeoxyribonuclease X | WP_000944282.1 | -0.29254866 | 0.10379616 | 0.17403346 |
| STM14_RS01100 | | DedA family protein | WP_001148368.1 | -0.292676008 | 0.23063007 | 0.33216307 |
| STM14_RS16565 | | YggT family protein | WP_001094848.1 | -0.293022963 | 0.15239259 | 0.23826307 |
| STM14_RS07555 | ihfA | integration host factor subunit alpha | WP_001229266.1 | -0.293437993 | 0.00089447 | 0.00291971 |
| STM14_RS20885 | metE | 5-methyltetrahydropteroyltriglutamate--homocysteine S-methyltransfer | WP_000154192.1 | -0.293843346 | 0.07731974 | 0.13615078 |
| STM14_RS04690 | | YbhN family protein | WP_001129814.1 | -0.294093593 | 0.16446041 | 0.25304773 |
| STM14_RS01040 | surA | peptidylprolyl isomerase SurA | WP_000800482.1 | -0.294297734 | 0.00219989 | 0.00642857 |
| STM14_RS12160 | galS | HTH-type transcriptional regulator GalS | WP_000628631.1 | -0.294580788 | 0.21549839 | 0.31337483 |
| STM14_RS20345 | phoU | phosphate signaling complex protein PhoU | WP_000377800.1 | -0.294717897 | 0.06801734 | 0.12157274 |
| STM14_RS04725 | hlyD | secretion protein HlyD | WP_000743436.1 | -0.294737756 | 0.01802182 | 0.03903314 |
| STM14_RS04125 | kdpE | two-component system response regulator KdpE | WP_000186054.1 | -0.29575559 | 0.08854672 | 0.15235001 |
| STM14_RS13880 | pbpC | peptidoglycan glycosyltransferase PbpC | WP_001014757.1 | -0.295977259 | 0.33796294 | 0.44955909 |
| STM14_RS03535 | fes | enterochelin esterase | WP_000656600.1 | -0.296546359 | 0.30352871 | 0.41285287 |
| STM14_RS12570 | gyrA | DNA topoisomerase (ATP-hydrolyzing) subunit A | WP_001281271.1 | -0.296714681 | 0.00180384 | 0.00543071 |
| STM14_RS15850 | | enolase C-terminal domain-like protein | WP_000211739.1 | -0.297104741 | 0.1989169 | 0.29473321 |
| STM14_RS11295 | cobU | bifunctional adenosylcobinamide kinase/adenosylcobinamide-phospha | WP_000973145.1 | -0.29744453 | 0.20279643 | 0.29951723 |
| STM14_RS05235 | cydD | cysteine/glutathione ABC transporter permease/ATP-binding protein Cy | WP_001044542.1 | -0.297819424 | 0.0298418 | 0.0599768 |
| STM14_RS17680 | yrbL | PhoP regulatory network protein YrbL | WP_000602196.1 | -0.29794019 | 0.00448389 | 0.01187834 |
| STM14_RS19535 | lldR | transcriptional regulator LldR | WP_000636696.1 | -0.298359 | 0.13909869 | 0.2206653 |
| STM14_RS07145 | | ATP-binding cassette domain-containing protein | WP_000950207.1 | -0.298365278 | 0.27930361 | 0.38632453 |
| STM14_RS17250 | yhaJ | DNA-binding transcriptional regulator YhaJ | WP_001041026.1 | -0.298481577 | 0.11880152 | 0.19432572 |
| STM14_RS04365 | tolB | Tol-Pal system beta propeller repeat protein TolB | WP_001562341.1 | -0.299159426 | 0.00182834 | 0.00549099 |
| STM14_RS11320 | cbiM | cobalt ECF transporter S component CbiM | WP_000764336.1 | -0.299194047 | 0.12402339 | 0.20091747 |
| STM14_RS03330 | fimI | type 1 fimbrial protein subunit FimI | WP_000619606.1 | -0.29997608 | 0.00181212 | 0.00545117 |
| STM14_RS20045 | ilvB | acetolactate synthase large subunit | WP_000168439.1 | -0.300000392 | 0.02986026 | 0.05998116 |
| STM14_RS16580 | hemW | radical SAM family heme chaperone HemW | WP_001096517.1 | -0.300450614 | 0.14757496 | 0.23250792 |
| STM14_RS08985 | | glucan biosynthesis protein | WP_001081943.1 | -0.300553274 | 0.02670845 | 0.05466399 |
| STM14_RS15360 | sitD | iron/manganese ABC transporter permease subunit SitD | WP_000480208.1 | -0.300702907 | 0.21474981 | 0.31253312 |
| STM14_RS23980 | serB | phosphoserine phosphatase | WP_001132983.1 | -0.300830094 | 0.07278526 | 0.12927898 |
| STM14_RS03080 | | MFS transporter | WP_001251593.1 | -0.300907217 | 0.23512548 | 0.33712305 |
| STM14_RS04895 | rimO | 30S ribosomal protein S12 methylthiotransferase RimO | WP_000073311.1 | -0.301521511 | 0.04275433 | 0.08143893 |
| STM14_RS00010 | | hypothetical protein | WP_001526822.1 | -0.302114067 | 0.16555135 | 0.2541952 |
| STM14_RS21850 | thiH | 2-iminoacetate synthase ThiH | WP_000847565.1 | -0.302301868 | 0.38715029 | 0.49816751 |
| STM14_RS21525 | hslV | ATP-dependent protease subunit HslV | WP_000208240.1 | -0.303257057 | 0.01761943 | 0.03831894 |
| STM14_RS22165 | lysC | lysine-sensitive aspartokinase 3 | WP_000136400.1 | -0.303468993 | 0.04140518 | 0.07915569 |
| STM14_RS12810 | nuoJ | NADH-quinone oxidoreductase subunit J | WP_000393522.1 | -0.304669817 | 0.01304197 | 0.02986353 |
| STM14_RS14620 | rimM | ribosome maturation factor RimM | WP_000043266.1 | -0.304672791 | 0.00084827 | 0.00278124 |
| STM14_RS12095 | gtdA | gentisate 1%2C2-dioxygenase | WP_001289857.1 | -0.305325227 | 0.18451114 | 0.2770677 |
| STM14_RS00925 | caiD | crotonobetainyl-CoA hydratase | WP_000004385.1 | -0.305464691 | 0.37079314 | 0.4814035 |
| STM14_RS22985 | ulaR | HTH-type transcriptional regulator UlaR | WP_000133618.1 | -0.30552132 | 0.10086729 | 0.16981864 |
| STM14_RS14135 | lepB | signal peptidase I | WP_000002559.1 | -0.305682056 | 0.02075405 | 0.04406945 |
| STM14_RS19100 | | MFS transporter | WP_001148983.1 | -0.305725132 | 0.38286213 | 0.4940199 |
| STM14_RS08030 | gloA | lactoylglutathione lyase | WP_001237787.1 | -0.305873129 | 0.00266742 | 0.00763127 |
| STM14_RS13775 | guaA | glutamine-hydrolyzing GMP synthase | WP_000138296.1 | -0.305930042 | 0.01534115 | 0.03423401 |
| STM14_RS02990 | acrB | efflux RND transporter permease AcrB | WP_001132508.1 | -0.306722128 | 0.00103426 | 0.00330568 |
| STM14_RS12840 | nuoC | NADH-quinone oxidoreductase subunit C/D | WP_000247855.1 | -0.30694933 | 0.00137 | 0.00424251 |
| STM14_RS22415 | | LysR family transcriptional regulator | WP_000354963.1 | -0.307260243 | 0.02340637 | 0.04866305 |
| STM14_RS13265 | | Nramp family divalent metal transporter | WP_000131735.1 | -0.307970116 | 0.01679802 | 0.03676168 |
| STM14_RS13055 | truA | tRNA pseudouridine(38-40) synthase TruA | WP_000016626.1 | -0.308067608 | 0.1089914 | 0.18101323 |
| STM14_RS20575 | ilvG | acetolactate synthase 2 catalytic subunit | WP_001012570.1 | -0.309480933 | 0.10950053 | 0.18169488 |
| STM14_RS15680 | truD | tRNA pseudouridine(13) synthase TruD | WP_000134246.1 | -0.309746714 | 0.00765481 | 0.01897981 |
| STM14_RS19370 | xylA | xylose isomerase | WP_001149561.1 | -0.31001445 | 0.04703709 | 0.088498 |
| STM14_RS03550 | wzz(fepE) | LPS O-antigen length regulator Wzz(fepE) | WP_001139583.1 | -0.310817434 | 0.01864452 | 0.04014569 |
| STM14_RS19740 | rph | ribonuclease PH | WP_001247078.1 | -0.312627895 | 0.02766006 | 0.0562987 |
| STM14_RS12185 | | L-serine ammonia-lyase | WP_000443208.1 | -0.3130668 | 0.18365606 | 0.27646019 |
| STM14_RS21620 | ptsP | phosphoenolpyruvate--protein phosphotransferase | WP_001128495.1 | -0.313710004 | 0.0267717 | 0.054763 |

| | | | | | | |
|---|---|---|---|---|---|---|
| STM14_RS14260 | | S49 family peptidase | WP_001189493.1 | -0.313897396 | 0.30192546 | 0.41145889 |
| STM14_RS17580 | ispB | octaprenyl diphosphate synthase | WP_001047354.1 | -0.314125116 | 0.00518785 | 0.01349942 |
| STM14_RS09500 | rluB | 23S rRNA pseudouridine(2605) synthase RluB | WP_001291233.1 | -0.314132307 | 0.02773817 | 0.05639534 |
| STM14_RS21710 | btuB | TonB-dependent vitamin B12 receptor BtuB | WP_000591398.1 | -0.314438332 | 0.01253186 | 0.02887503 |
| STM14_RS18770 | gntR | gluconate operon transcriptional repressor GntR | WP_000730239.1 | -0.315101753 | 0.02817168 | 0.05708758 |
| STM14_RS12550 | | porin OmpC | WP_000865526.1 | -0.316167007 | 0.00039049 | 0.00140545 |
| STM14_RS02770 | phnR | phosphonate utilization transcriptional regulator PhnR | WP_000836268.1 | -0.316240944 | 0.32815833 | 0.4400142 |
| STM14_RS12505 | napG | ferredoxin-type protein NapG | WP_000091673.1 | -0.316487425 | 0.16668927 | 0.25551619 |
| STM14_RS13350 | | LysR family transcriptional regulator | WP_001109831.1 | -0.317377017 | 0.32151759 | 0.43236953 |
| STM14_RS13690 | bcp | thioredoxin-dependent thiol peroxidase | WP_001068686.1 | -0.317659374 | 0.01281301 | 0.0294124 |
| STM14_RS21635 | | formate C-acetyltransferase | WP_000149714.1 | -0.317723451 | 0.00432484 | 0.01151414 |
| STM14_RS05830 | | Lon protease family protein | WP_000156453.1 | -0.318806889 | 0.02133722 | 0.04515153 |
| STM14_RS04405 | nadA | quinolinate synthase NadA | WP_000115347.1 | -0.318840878 | 0.34477551 | 0.45557083 |
| STM14_RS08320 | osmX | osmoprotectant ABC transporter substrate-binding protein OsmX | WP_001211193.1 | -0.319013506 | 0.09352645 | 0.15937966 |
| STM14_RS00930 | caiC | crotonobetaine/carnitine-CoA ligase | WP_000355806.1 | -0.31960629 | 0.35842076 | 0.46881181 |
| STM14_RS15180 | recA | recombinase RecA | WP_000963150.1 | -0.320367487 | 0.00054529 | 0.00189053 |
| STM14_RS17745 | | N-acetylmannosamine-6-phosphate 2-epimerase | WP_000054441.1 | -0.320476565 | 0.33687777 | 0.4484396 |
| STM14_RS13175 | mlaA | phospholipid-binding lipoprotein MlaA | WP_000776787.1 | -0.320592067 | 0.01196631 | 0.02774556 |
| STM14_RS02465 | | bifunctional 2-methylcitrate dehydratase/aconitate hydratase | WP_000107071.1 | -0.320966292 | 0.00468883 | 0.01234043 |
| STM14_RS19145 | bcsB | cellulose biosynthesis cyclic di-GMP-binding regulatory protein BcsB | WP_000823813.1 | -0.321634107 | 0.01521059 | 0.03402514 |
| STM14_RS22780 | frdD | fumarate reductase subunit FrdD | WP_000609649.1 | -0.321945401 | 0.00347625 | 0.00959488 |
| STM14_RS13975 | asrC | sulfite reductase subunit C | WP_000020685.1 | -0.321977293 | 0.03123264 | 0.06222867 |
| STM14_RS01225 | murC | UDP-N-acetylmuramate--L-alanine ligase | WP_001096070.1 | -0.322859236 | 0.00407774 | 0.01102367 |
| STM14_RS10860 | | very short patch repair endonuclease | WP_000785978.1 | -0.323122791 | 0.35009628 | 0.46070568 |
| STM14_RS24035 | creC | two-component system sensor histidine kinase CreC | WP_001219526.1 | -0.323487679 | 0.21519788 | 0.31306148 |
| STM14_RS09125 | nifJ | pyruvate:ferredoxin (flavodoxin) oxidoreductase | WP_001193265.1 | -0.323704718 | 0.00115581 | 0.00364046 |
| STM14_RS02785 | yajL | protein deglycase YajL | WP_001275806.1 | -0.323808327 | 0.21337541 | 0.31139448 |
| STM14_RS10380 | ruvB | Holliday junction branch migration DNA helicase RuvB | WP_000568508.1 | -0.32419675 | 0.01476535 | 0.03315 |
| STM14_RS05865 | helD | DNA helicase IV | WP_000420505.1 | -0.324287214 | 0.00428126 | 0.01143947 |
| STM14_RS03120 | | SPFH/Band 7/PHB domain protein | WP_000906146.1 | -0.325206416 | 0.00396856 | 0.01076013 |
| STM14_RS08660 | glgX | glycogen debranching protein GlgX | WP_000210601.1 | -0.325409912 | 0.00209644 | 0.00619511 |
| STM14_RS01495 | panB | 3-methyl-2-oxobutanoate hydroxymethyltransferase | WP_000805487.1 | -0.325496924 | 0.19325564 | 0.28761813 |
| STM14_RS23890 | prfC | peptide chain release factor 3 | WP_000175965.1 | -0.325525387 | 0.00838652 | 0.02055871 |
| STM14_RS04055 | tcuC | tricarballylate/proton symporter TcuC | WP_000057014.1 | -0.326111039 | 0.21912428 | 0.31814495 |
| STM14_RS22785 | frdC | fumarate reductase subunit FrdC | WP_000208749.1 | -0.326135472 | 0.00348908 | 0.0096015 |
| STM14_RS09625 | oppB | oligopeptide ABC transporter permease OppB | WP_000911097.1 | -0.326388153 | 0.03145518 | 0.0626043 |
| STM14_RS16335 | ubiI | FAD-dependent 2-octaprenylphenol hydroxylase | WP_001192144.1 | -0.326591209 | 0.00732494 | 0.01830986 |
| STM14_RS01520 | sfsA | DNA/RNA nuclease SfsA | WP_000899412.1 | -0.326619546 | 0.00812895 | 0.0200743 |
| STM14_RS19570 | gpsA | NAD(P)H-dependent glycerol-3-phosphate dehydrogenase | WP_001076596.1 | -0.326641248 | 0.00089657 | 0.00292398 |
| STM14_RS19395 | avtA | valine--pyruvate transaminase | WP_000144381.1 | -0.327046243 | 0.07377079 | 0.13068292 |
| STM14_RS06690 | | lysozyme inhibitor LprI family protein | WP_000825957.1 | -0.327150046 | 0.22724519 | 0.32812782 |
| STM14_RS18420 | | YhfG family protein | WP_001521109.1 | -0.327705273 | 0.09009488 | 0.15465237 |
| STM14_RS06425 | rne | ribonuclease E | WP_000827470.1 | -0.327877029 | 0.00158882 | 0.00485077 |
| STM14_RS03920 | gltL | glutamate/aspartate ABC transporter ATP-binding protein GltL | WP_000631369.1 | -0.327912665 | 0.03301086 | 0.0649775 |
| STM14_RS17320 | garD | galactarate dehydratase | WP_001273723.1 | -0.328015756 | 0.15118176 | 0.23667145 |
| STM14_RS17460 | rpsO | 30S ribosomal protein S15 | WP_000059465.1 | -0.328125486 | 0.00034261 | 0.00124902 |
| STM14_RS15190 | mltB | lytic murein transglycosylase B | WP_000476820.1 | -0.328453763 | 0.02317998 | 0.0483005 |
| STM14_RS23895 | osmY | molecular chaperone OsmY | WP_000178963.1 | -0.328609384 | 4.1235E-05 | 0.00019341 |
| STM14_RS03240 | allE | (S)-ureidoglycine aminohydrolase | WP_000540981.1 | -0.328781591 | 0.05029141 | 0.09361627 |
| STM14_RS18360 | | ABC transporter ATP-binding protein | WP_000634766.1 | -0.328817851 | 0.03071767 | 0.06140199 |
| STM14_RS00630 | dnaJ | molecular chaperone DnaJ | WP_001119009.1 | -0.328881479 | 0.00204338 | 0.00605775 |
| STM14_RS10975 | | hypothetical protein | WP_000779218.1 | -0.329106242 | 0.18620986 | 0.27916315 |
| STM14_RS23485 | idnT | gnt-II system L-idonate transporter | WP_001128356.1 | -0.329313166 | 0.01282334 | 0.02941779 |
| STM14_RS10840 | dgcQ | cellulose biosynthesis regulator diguanylate cyclase DgcQ | WP_001119821.1 | -0.329633447 | 0.01956348 | 0.04190386 |
| STM14_RS19550 | | LacI family DNA-binding transcriptional regulator | WP_010989081.1 | -0.329795222 | 0.17114963 | 0.26083317 |
| STM14_RS23970 | lplA | lipoate--protein ligase LplA | WP_000209756.1 | -0.32993057 | 0.06986204 | 0.12462791 |
| STM14_RS07450 | katE | catalase HPII | WP_000019119.1 | -0.330628592 | 0.00065361 | 0.00220777 |
| STM14_RS01375 | acnB | bifunctional aconitate hydratase 2/2-methylisocitrate dehydratase | WP_000888962.1 | -0.330646489 | 0.000281 | 0.00104127 |
| STM14_RS00660 | | chitinase | WP_000155792.1 | -0.331097832 | 0.2494688 | 0.35301465 |
| STM14_RS12180 | | helix-turn-helix transcriptional regulator | WP_000876817.1 | -0.331452752 | 0.3347733 | 0.44676887 |
| STM14_RS12545 | apbE | FAD:protein FMN transferase ApbE | WP_000784307.1 | -0.332037254 | 0.15965763 | 0.24762401 |
| STM14_RS22730 | groL | chaperonin GroEL | WP_000729126.1 | -0.33217464 | 0.00052222 | 0.00182257 |
| STM14_RS02835 | cyoA | cytochrome o ubiquinol oxidase subunit II | WP_001239449.1 | -0.332734986 | 0.00013896 | 0.00056164 |
| STM14_RS20190 | torR | two-component system response regulator TorR | WP_001562512.1 | -0.333462771 | 0.01308121 | 0.02992371 |
| STM14_RS06035 | | MBL fold metallo-hydrolase | WP_000976562.1 | -0.333868043 | 0.32040473 | 0.43129314 |
| STM14_RS06475 | fabD | ACP S-malonyltransferase | WP_000191343.1 | -0.334978859 | 0.00132657 | 0.00413234 |
| STM14_RS20615 | ilvC | ketol-acid reductoisomerase | WP_000024948.1 | -0.335162033 | 0.00587835 | 0.0150935 |
| STM14_RS13785 | xseA | exodeoxyribonuclease VII large subunit | WP_000953165.1 | -0.335924739 | 0.06100579 | 0.1109251 |
| STM14_RS23870 | rsmC | 16S rRNA (guanine(1207)-N(2))-methyltransferase RsmC | WP_001272276.1 | -0.33669322 | 0.29747097 | 0.40611351 |
| STM14_RS05805 | pqiA | membrane integrity-associated transporter subunit PqiA | WP_000333152.1 | -0.33688034 | 0.0404768 | 0.0776227 |
| STM14_RS20745 | hemX | uroporphyrinogen-III C-methyltransferase | WP_000138962.1 | -0.337072174 | 0.01054687 | 0.02498942 |
| STM14_RS07980 | mdtK | multidrug efflux MATE transporter MdtK | WP_001175077.1 | -0.337138761 | 0.02816973 | 0.05708758 |
| STM14_RS02275 | | LysR family transcriptional regulator | WP_000859023.1 | -0.337394496 | 0.1593664 | 0.24727648 |
| STM14_RS12320 | rsuA | 16S rRNA pseudouridine(516) synthase RsuA | WP_001234836.1 | -0.337608686 | 0.07344299 | 0.13032149 |
| STM14_RS12510 | napA | nitrate reductase catalytic subunit NapA | WP_000778094.1 | -0.337844 | 0.00422268 | 0.01131579 |
| STM14_RS16065 | rppH | RNA pyrophosphohydrolase | WP_000564481.1 | -0.338326909 | 0.00445384 | 0.01181487 |
| STM14_RS18320 | tusD | sulfurtransferase complex subunit TusD | WP_001268010.1 | -0.338551869 | 0.31708636 | 0.42766006 |
| STM14_RS17465 | truB | tRNA pseudouridine(55) synthase TruB | WP_000089674.1 | -0.338599033 | 0.01222855 | 0.02828236 |
| STM14_RS05105 | | N-acetylmuramoyl-L-alanine amidase | WP_000645852.1 | -0.339402865 | 0.02223865 | 0.04665682 |
| STM14_RS11290 | cobS | adenosylcobinamide-GDP ribazoletransferase | WP_000039997.1 | -0.340682029 | 0.0454146 | 0.08575209 |
| STM14_RS13355 | | bile acid:sodium symporter | WP_000765569.1 | -0.340750313 | 0.34349216 | 0.45459103 |
| STM14_RS18875 | livK | high-affinity branched-chain amino acid ABC transporter substrate-binc | WP_000822976.1 | -0.340826562 | 0.106571 | 0.17771486 |

| | | | | | | |
|---|---|---|---|---|---|---|
| STM14_RS01660 | map | type I methionyl aminopeptidase | WP_001018214.1 | -0.34104232 | 0.01091964 | 0.02569081 |
| STM14_RS03195 | allR | HTH-type transcriptional repressor AllR | WP_000141265.1 | -0.34130274 | 0.00753272 | 0.01872608 |
| STM14_RS18755 | asd | aspartate-semialdehyde dehydrogenase | WP_000799940.1 | -0.343150235 | 0.00189279 | 0.00566606 |
| STM14_RS01780 | | YaeQ family protein | WP_001185319.1 | -0.34414576 | 0.1993049 | 0.29518932 |
| STM14_RS01740 | dnaE | DNA polymerase III subunit alpha | WP_001294826.1 | -0.344178641 | 0.00189207 | 0.00566606 |
| STM14_RS13655 | | YpfN family protein | WP_000383839.1 | -0.34451751 | 0.01020367 | 0.02426998 |
| STM14_RS16390 | argO | arginine exporter ArgO | WP_000626863.1 | -0.345066082 | 0.11563404 | 0.19015834 |
| STM14_RS08525 | | hydrogenase | WP_000155978.1 | -0.345256784 | 0.23749166 | 0.33972195 |
| STM14_RS07015 | | SOS response-associated peptidase | WP_000334551.1 | -0.345343291 | 0.01160712 | 0.027049 |
| STM14_RS19485 | selA | L-seryl-tRNA(Sec) selenium transferase | WP_000200187.1 | -0.345783074 | 0.00253747 | 0.00729349 |
| STM14_RS16295 | ygfZ | tRNA-modifying protein YgfZ | WP_000874172.1 | -0.346471393 | 0.00294424 | 0.00831979 |
| STM14_RS09490 | | YciK family oxidoreductase | WP_000559310.1 | -0.347459212 | 0.00872485 | 0.02125936 |
| STM14_RS18505 | hofQ | DNA uptake porin HofQ | WP_000832802.1 | -0.347568296 | 0.07119812 | 0.12676571 |
| STM14_RS21075 | hemN | oxygen-independent coproporphyrinogen III oxidase | WP_000003520.1 | -0.348104417 | 0.00659516 | 0.01670855 |
| STM14_RS13515 | eutL | ethanolamine utilization microcompartment protein EutL | WP_001111056.1 | -0.348370045 | 0.07156059 | 0.1272265 |
| STM14_RS13035 | dedD | cell division protein DedD | WP_000157037.1 | -0.348393519 | 0.00746614 | 0.01857453 |
| STM14_RS22980 | yjfP | esterase | WP_000560303.1 | -0.349785537 | 0.01736618 | 0.03785807 |
| STM14_RS05335 | serC | 3-phosphoserine/phosphohydroxythreonine transaminase | WP_000079584.1 | -0.349975983 | 0.00128336 | 0.00400452 |
| STM14_RS06675 | hflD | high frequency lysogenization protein HflD | WP_001519653.1 | -0.350317436 | 0.12806461 | 0.20622414 |
| STM14_RS02455 | prpB | methylisocitrate lyase | WP_000052224.1 | -0.35045706 | 0.0039762 | 0.0107729 |
| STM14_RS08270 | mdtI | multidrug/spermidine efflux SMR transporter subunit MdtI | WP_001183821.1 | -0.351093474 | 0.1090842 | 0.18108567 |
| STM14_RS00225 | | IS481 family transposase | WP_000098783.1 | -0.351609767 | 0.00013742 | 0.00055662 |
| STM14_RS12710 | arnD | 4-deoxy-4-formamido-L-arabinose-phosphoundecaprenol deformylase | WP_000169761.1 | -0.351870051 | 0.04482346 | 0.08494079 |
| STM14_RS08085 | anmK | anhydro-N-acetylmuramic acid kinase | WP_000835011.1 | -0.352169606 | 0.03167637 | 0.06292056 |
| STM14_RS01115 | thiB | thiamine ABC transporter substrate binding subunit | WP_000915326.1 | -0.352716861 | 0.18432858 | 0.2769065 |
| STM14_RS02575 | rdgC | recombination-associated protein RdgC | WP_000964305.1 | -0.35294041 | 0.01745152 | 0.03799911 |
| STM14_RS03285 | lpxH | UDP-2%2C3-diacylglucosamine diphosphatase | WP_000212284.1 | -0.353292857 | 0.21930399 | 0.31828037 |
| STM14_RS12145 | mglC | galactose/methyl galactoside ABC transporter permease MglC | WP_001275101.1 | -0.354590689 | 0.05096085 | 0.09476659 |
| STM14_RS18080 | | type I DNA topoisomerase | WP_001129708.1 | -0.354652171 | 0.00827799 | 0.02033327 |
| STM14_RS07335 | sppA | signal peptide peptidase SppA | WP_001259876.1 | -0.355950536 | 0.01794339 | 0.03890905 |
| STM14_RS04535 | modB | molybdate ABC transporter permease subunit | WP_000604026.1 | -0.356426399 | 0.02843229 | 0.05752072 |
| STM14_RS22290 | | quinone oxidoreductase | WP_000235551.1 | -0.356631694 | 0.0036715 | 0.01004343 |
| STM14_RS02150 | | amidohydrolase | WP_000015789.1 | -0.356657671 | 0.02192286 | 0.04607303 |
| STM14_RS20990 | hemG | menaquinone-dependent protoporphyrinogen IX dehydrogenase | WP_000853952.1 | -0.357024328 | 0.01546452 | 0.0344884 |
| STM14_RS11330 | | sirohydrochlorin cobaltochelatase | WP_000709784.1 | -0.357255345 | 0.05515607 | 0.10174582 |
| STM14_RS04370 | pal | peptidoglycan-associated lipoprotein Pal | WP_001176621.1 | -0.357577352 | 5.8769E-05 | 0.00026357 |
| STM14_RS18960 | | MFS transporter | WP_001576500.1 | -0.358000511 | 0.09899953 | 0.16720929 |
| STM14_RS07700 | sufA | Fe-S cluster assembly scaffold SufA | WP_000418970.1 | -0.358798918 | 0.00889376 | 0.02161507 |
| STM14_RS11565 | hisD | histidinol dehydrogenase | WP_000009626.1 | -0.359463016 | 0.05416687 | 0.10017197 |
| STM14_RS16240 | actS | amidase activator ActS | WP_000244329.1 | -0.36029471 | 0.01308451 | 0.02992371 |
| STM14_RS02580 | mak | fructokinase | WP_001219277.1 | -0.360964712 | 0.00893646 | 0.02166164 |
| STM14_RS05390 | kdsB | 3-deoxy-manno-octulosonate cytidylyltransferase | WP_000011576.1 | -0.361044915 | 0.02148424 | 0.04538439 |
| STM14_RS17565 | | DMT family transporter | WP_000813052.1 | -0.361505086 | 0.01515652 | 0.0339248 |
| STM14_RS06605 | nagK | N-acetylglucosamine kinase | WP_000291338.1 | -0.361658519 | 0.00544957 | 0.01411063 |
| STM14_RS01005 | kefF | glutathione-regulated potassium-efflux system oxidoreductase KefF | WP_000600709.1 | -0.363348771 | 0.15597417 | 0.24296761 |
| STM14_RS10970 | | phage tail sheath subtilisin-like domain-containing protein | WP_001007991.1 | -0.363796188 | 0.05735214 | 0.10521704 |
| STM14_RS19620 | rfaD | ADP-glyceromanno-heptose 6-epimerase | WP_000587771.1 | -0.365913139 | 0.00010133 | 0.00042185 |
| STM14_RS02570 | ppnP | pyrimidine/purine nucleoside phosphorylase | WP_000941952.1 | -0.366228036 | 0.00681962 | 0.01718675 |
| STM14_RS20980 | | IMPACT family protein | WP_000378902.1 | -0.366346516 | 0.05506707 | 0.10163256 |
| STM14_RS16375 | argP | DNA-binding transcriptional regulator ArgP | WP_000828345.1 | -0.366473695 | 0.18408207 | 0.27687869 |
| STM14_RS23440 | | GNAT family N-acetyltransferase | WP_001519079.1 | -0.367371391 | 0.12468076 | 0.20164441 |
| STM14_RS17315 | garL | 2-dehydro-3-deoxyglucarate aldolase | WP_001057715.1 | -0.367491056 | 0.023168 | 0.0483005 |
| STM14_RS01230 | | D-alanine--D-alanine ligase | WP_000763905.1 | -0.367972783 | 0.00115134 | 0.00362948 |
| STM14_RS12080 | | 3-hydroxybenzoate 6-monooxygenase | WP_000150103.1 | -0.3679951 | 0.02990148 | 0.06003121 |
| STM14_RS02205 | crl | sigma factor-binding protein Crl | WP_000174696.1 | -0.368450656 | 0.00249844 | 0.00719816 |
| STM14_RS22890 | hflX | GTPase HflX | WP_000460338.1 | -0.369050818 | 7.0218E-05 | 0.00030633 |
| STM14_RS17770 | sspA | stringent starvation protein A | WP_000257298.1 | -0.369052195 | 0.00024944 | 0.000942 |
| STM14_RS22515 | | zinc ribbon domain-containing protein YjdM | WP_000056482.1 | -0.369610942 | 0.00528364 | 0.01371958 |
| STM14_RS02915 | decR | DNA-binding transcriptional regulator DecR | WP_000884593.1 | -0.369613679 | 0.055939 | 0.10293223 |
| STM14_RS19530 | lldP | L-lactate permease | WP_001055305.1 | -0.370245897 | 0.07496017 | 0.13237571 |
| STM14_RS12740 | menC | o-succinylbenzoate synthase | WP_001255561.1 | -0.371369968 | 0.01555717 | 0.03459028 |
| STM14_RS04130 | kdpD | two-component system sensor histidine kinase KdpD | WP_000997487.1 | -0.371406774 | 0.01861249 | 0.04010017 |
| STM14_RS17605 | mlaC | phospholipid-binding protein MlaC | WP_000476475.1 | -0.371630015 | 0.00096846 | 0.00313622 |
| STM14_RS08475 | | LysR family transcriptional regulator | WP_000366538.1 | -0.371940587 | 0.19881159 | 0.29469576 |
| STM14_RS03135 | cnoX | chaperedoxin | WP_000116058.1 | -0.372826749 | 0.00348432 | 0.0096015 |
| STM14_RS12795 | nuoM | NADH-quinone oxidoreductase subunit M | WP_000926431.1 | -0.373255597 | 0.00172132 | 0.00520355 |
| STM14_RS16495 | speA | biosynthetic arginine decarboxylase | WP_001278579.1 | -0.373697883 | 0.00023709 | 0.00090089 |
| STM14_RS06540 | lpoB | penicillin-binding protein activator LpoB | WP_000164359.1 | -0.374002483 | 0.0016842 | 0.00510388 |
| STM14_RS21360 | cpxA | envelope stress sensor histidine kinase CpxA | WP_000580402.1 | -0.374044754 | 0.00467868 | 0.01232254 |
| STM14_RS05630 | | phage terminase large subunit family protein | WP_000989241.1 | -0.375645614 | 0.10733111 | 0.17857802 |
| STM14_RS11310 | | energy-coupling factor ABC transporter transmembrane protein | WP_000147137.1 | -0.375973813 | 0.06767525 | 0.1211377 |
| STM14_RS15330 | hypE | hydrogenase expression/formation protein HypE | WP_001059953.1 | -0.376204936 | 0.00404736 | 0.01095764 |
| STM14_RS02650 | yajC | preprotein translocase subunit YajC | WP_000007628.1 | -0.376214388 | 0.00031178 | 0.00114797 |
| STM14_RS15185 | pncC | nicotinamide-nucleotide amidase | WP_000132239.1 | -0.376247805 | 0.00633891 | 0.01614098 |
| STM14_RS20870 | yigL | sugar/pyridoxal phosphate phosphatase YigL | WP_000285338.1 | -0.376310185 | 0.01674456 | 0.03669849 |
| STM14_RS06100 | | lysozyme inhibitor LprI family protein | WP_000821066.1 | -0.377043277 | 0.01430908 | 0.03222386 |
| STM14_RS23300 | | amidohydrolase/deacetylase family metallohydrolase | WP_000459930.1 | -0.378021276 | 0.26994139 | 0.37577475 |
| STM14_RS20440 | atpI | F0F1 ATP synthase subunit I | WP_000116685.1 | -0.378216829 | 0.00385654 | 0.01048729 |
| STM14_RS12205 | lysP | lysine-specific permease | WP_000535000.1 | -0.378243081 | 0.00417309 | 0.01122375 |
| STM14_RS07170 | zinT | metal-binding protein ZinT | WP_001574476.1 | -0.378416565 | 0.1865077 | 0.27943065 |
| STM14_RS04340 | ybgE | cyd operon protein YbgE | WP_000875305.1 | -0.37849215 | 0.05524622 | 0.10186108 |
| STM14_RS05130 | hcr | NADH oxidoreductase | WP_001669316.1 | -0.379117413 | 0.09633568 | 0.1633846 |

| | | | | | | |
|---|---|---|---|---|---|---|
| STM14_RS09495 | cobO | cob(I)yrinic acid a%2Cc-diamide adenosyltransferase | WP_001278856.1 | -0.379516511 | 0.00646079 | 0.01639465 |
| STM14_RS18675 | | DeoR/GlpR family transcriptional regulator | WP_000815085.1 | -0.379626455 | 0.00429855 | 0.01146684 |
| STM14_RS03060 | hemH | ferrochelatase | WP_001250078.1 | -0.379848458 | 0.01988596 | 0.04244643 |
| STM14_RS01730 | lpxB | lipid-A-disaccharide synthase | WP_000741212.1 | -0.381463953 | 0.05531299 | 0.10188215 |
| STM14_RS18930 | | lysoplasmalogenase | WP_000964735.1 | -0.381941212 | 0.01084483 | 0.02554745 |
| STM14_RS16795 | hybE | hydrogenase-2 assembly chaperone | WP_000004953.1 | -0.382520627 | 0.01720632 | 0.0375318 |
| STM14_RS22255 | dinF | MATE family efflux transporter DinF | WP_001128116.1 | -0.383003178 | 0.16657842 | 0.25545281 |
| STM14_RS04770 | ybiO | mechanosensitive channel protein | WP_001267705.1 | -0.383026559 | 0.14045593 | 0.22272124 |
| STM14_RS21675 | argB | acetylglutamate kinase | WP_001575262.1 | -0.383409155 | 0.12229514 | 0.19854088 |
| STM14_RS09610 | oppF | murein tripeptide/oligopeptide ABC transporter ATP-binding protein Op | WP_000994696.1 | -0.383431477 | 0.00270613 | 0.00772999 |
| STM14_RS20795 | dapF | diaminopimelate epimerase | WP_001160671.1 | -0.383895029 | 0.00051885 | 0.00181254 |
| STM14_RS17100 | bacA | undecaprenyl-diphosphate phosphatase | WP_001281933.1 | -0.38536518 | 0.05132883 | 0.09535458 |
| STM14_RS13475 | | DUF2919 domain-containing protein | WP_000842932.1 | -0.385422277 | 0.24738872 | 0.3507452 |
| STM14_RS04330 | cydX | cytochrome bd-I oxidase subunit CydX | WP_000270278.1 | -0.385964015 | 0.00431659 | 0.01150048 |
| STM14_RS06590 | lolC | lipoprotein-releasing ABC transporter permease subunit LolC | WP_001518787.1 | -0.386035892 | 0.00607467 | 0.01553258 |
| STM14_RS09290 | ycjG | L-Ala-D/L-Glu epimerase | WP_001258061.1 | -0.386069406 | 0.0064273 | 0.01633218 |
| STM14_RS16940 | plsC | 1-acylglycerol-3-phosphate O-acyltransferase | WP_000965730.1 | -0.386406814 | 0.04844285 | 0.09063342 |
| STM14_RS03580 | entC | isochorismate synthase EntC | WP_000367601.1 | -0.387516627 | 0.29050626 | 0.39808115 |
| STM14_RS21890 | hemE | uroporphyrinogen decarboxylase | WP_000137623.1 | -0.38764596 | 0.00349927 | 0.00962235 |
| STM14_RS19655 | rfaJ | lipopolysaccharide 1%2C2-glucosyltransferase RfaJ | WP_000376865.1 | -0.387819982 | 0.05388863 | 0.09972664 |
| STM14_RS09925 | dsbB | disulfide bond formation protein DsbB | WP_000943475.1 | -0.388127783 | 0.06895101 | 0.12312203 |
| STM14_RS02485 | iprA | hydrogen peroxide resistance inhibitor IprA | WP_000950242.1 | -0.388275364 | 0.28964573 | 0.39719761 |
| STM14_RS05670 | | phage tail protein | WP_000132756.1 | -0.388619198 | 0.20067969 | 0.29698659 |
| STM14_RS20685 | rffC | dTDP-4-amino-4%2C6-dideoxy-D-galactose acyltransferase | WP_001145156.1 | -0.388956355 | 0.0483659 | 0.09053546 |
| STM14_RS20130 | | cytochrome c-type biogenesis protein CcmH | WP_001238334.1 | -0.389104825 | 0.23105984 | 0.33234948 |
| STM14_RS17975 | panF | sodium/pantothenate symporter | WP_001175741.1 | -0.39013744 | 0.21188075 | 0.3100735 |
| STM14_RS18670 | rtcR | RNA repair transcriptional activator RtcR | WP_001232871.1 | -0.390144803 | 0.05664904 | 0.1040308 |
| STM14_RS17440 | mtr | tryptophan permease | WP_000224386.1 | -0.390411329 | 0.27586383 | 0.38228476 |
| STM14_RS14555 | clpB | ATP-dependent chaperone ClpB | WP_001235094.1 | -0.390734186 | 3.4642E-06 | 2.0376E-05 |
| STM14_RS11375 | cbiB | adenosylcobinamide-phosphate synthase CbiB | WP_000153648.1 | -0.391464146 | 0.1639947 | 0.25254224 |
| STM14_RS23925 | deoC | deoxyribose-phosphate aldolase | WP_001519713.1 | -0.39166769 | 0.00182219 | 0.005477 |
| STM14_RS12945 | yfcD | NUDIX hydrolase YfcD | WP_000230215.1 | -0.39208961 | 0.00106672 | 0.00338008 |
| STM14_RS08360 | | YnfC family lipoprotein | WP_000743122.1 | -0.392496312 | 0.00245007 | 0.00710888 |
| STM14_RS23265 | pmbA | metalloprotease PmbA | WP_000852994.1 | -0.392925397 | 0.00098374 | 0.00316897 |
| STM14_RS03245 | allC | allantoate deiminase | WP_000589769.1 | -0.393563056 | 0.10865206 | 0.18053109 |
| STM14_RS19050 | | anaerobic C4-dicarboxylate transporter family protein | WP_000498935.1 | -0.393807376 | 0.03857562 | 0.07444206 |
| STM14_RS13670 | | phosphoribosylaminoimidazolesuccinocarboxamide synthase | WP_001171630.1 | -0.393976577 | 0.00067842 | 0.00228121 |
| STM14_RS24030 | creB | two-component system response regulator CreB | WP_001187047.1 | -0.394550703 | 0.01489318 | 0.03341663 |
| STM14_RS02160 | lpcA | D-sedoheptulose 7-phosphate isomerase | WP_000284051.1 | -0.395035226 | 0.00081785 | 0.00270315 |
| STM14_RS13800 | ratA | DUF823/DUF824 repeat adhesin RatA | WP_000375791.1 | -0.395700341 | 0.00126298 | 0.00394762 |
| STM14_RS23810 | dnaT | primosomal protein DnaT | WP_000098578.1 | -0.395848682 | 0.02011715 | 0.04289018 |
| STM14_RS03610 | | YbdD/YjiX family protein | WP_000460619.1 | -0.395975744 | 0.00275578 | 0.00784142 |
| STM14_RS06450 | | Maf family protein | WP_001137605.1 | -0.396442605 | 0.02875277 | 0.05807335 |
| STM14_RS03860 | | UxaA family hydrolase | WP_001670689.1 | -0.396605403 | 0.00518779 | 0.01349942 |
| STM14_RS13680 | dapA | 4-hydroxy-tetrahydrodipicolinate synthase | WP_000494020.1 | -0.396615437 | 2.8899E-05 | 0.00013909 |
| STM14_RS05730 | | phage tail protein | WP_001144679.1 | -0.396734737 | 0.22355602 | 0.32388724 |
| STM14_RS01310 | nadC | carboxylating nicotinate-nucleotide diphosphorylase | WP_001135135.1 | -0.396773322 | 0.03574612 | 0.06974945 |
| STM14_RS09990 | | Slp family lipoprotein | WP_000290594.1 | -0.398180884 | 0.03441347 | 0.06739916 |
| STM14_RS04260 | | AbrB family transcriptional regulator | WP_001122057.1 | -0.399411315 | 0.12828172 | 0.20643938 |
| STM14_RS12035 | | ABC transporter substrate-binding protein | WP_000155869.1 | -0.399441061 | 0.00026424 | 0.00098978 |
| STM14_RS12820 | nuoH | NADH-quinone oxidoreductase subunit NuoH | WP_000118515.1 | -0.399606813 | 0.00052737 | 0.00183533 |
| STM14_RS15620 | | aldolase | WP_000153635.1 | -0.40021736 | 0.01505241 | 0.03371227 |
| STM14_RS15285 | hycE | formate hydrogenlyase subunit HycE | WP_001288155.1 | -0.400672131 | 0.05902375 | 0.10785381 |
| STM14_RS20865 | pldB | lysophospholipase L2 | WP_000487685.1 | -0.40165915 | 0.00151054 | 0.0046233 |
| STM14_RS09725 | narL | two-component system response regulator NarL | WP_001064598.1 | -0.401821686 | 0.04388755 | 0.08338181 |
| STM14_RS23885 | yjjG | pyrimidine 5'-nucleotidase | WP_000870686.1 | -0.402112041 | 0.08840158 | 0.1521714 |
| STM14_RS26450 | | fumarate hydratase | | -0.402434065 | 0.08457685 | 0.1466158 |
| STM14_RS01875 | | endonuclease/exonuclease/phosphatase family protein | WP_001230968.1 | -0.403945221 | 0.00455254 | 0.01202408 |
| STM14_RS10670 | tcyL | cystine ABC transporter permease | WP_001158238.1 | -0.40449919 | 0.04514341 | 0.08533174 |
| STM14_RS07405 | cho | excinuclease Cho | WP_001526281.1 | -0.405408127 | 0.02267337 | 0.04743073 |
| STM14_RS20820 | corA | magnesium/cobalt transporter CorA | WP_000947139.1 | -0.40587166 | 0.00188156 | 0.00564161 |
| STM14_RS08050 | | aldo/keto reductase family oxidoreductase | WP_000250717.1 | -0.407631754 | 0.03043455 | 0.06093529 |
| STM14_RS18450 | cysG | siroheme synthase CysG | WP_000349895.1 | -0.407800082 | 0.01934052 | 0.04152291 |
| STM14_RS01895 | | class I SAM-dependent methyltransferase | WP_000801238.1 | -0.407904349 | 0.16771071 | 0.25665455 |
| STM14_RS14465 | trmN | tRNA(1)(Val) (adenine(37)-N(6))-methyltransferase TrmN | WP_000083343.1 | -0.408002067 | 0.01833682 | 0.03959893 |
| STM14_RS18985 | rbbA | ribosome-associated ATPase/putative transporter RbbA | WP_000202926.1 | -0.408110956 | 0.00014652 | 0.00058833 |
| STM14_RS12400 | | KAP family P-loop domain protein | WP_000894640.1 | -0.408386637 | 0.19474549 | 0.28948441 |
| STM14_RS11975 | apbC | iron-sulfur cluster carrier protein ApbC | WP_001005424.1 | -0.409398296 | 5.4294E-05 | 0.0002462 |
| STM14_RS04495 | galK | galactokinase | WP_001049367.1 | -0.40964534 | 0.00101092 | 0.00324235 |
| STM14_RS22470 | nrfEF | heme lyase NrfEFG subunit NrfEF | WP_000551709.1 | -0.410107875 | 0.13686153 | 0.21777189 |
| STM14_RS08745 | smvA | methyl viologen efflux MFS transporter SmvA | WP_000489731.1 | -0.410216611 | 0.09165585 | 0.15689299 |
| STM14_RS15925 | fucK | L-fuculokinase | WP_000806780.1 | -0.410217986 | 0.08776335 | 0.15121415 |
| STM14_RS13235 | alaC | alanine transaminase | WP_000763118.1 | -0.410357031 | 7.5315E-05 | 0.00032548 |
| STM14_RS18980 | | ABC transporter permease | WP_001214750.1 | -0.411009679 | 0.01585738 | 0.03515163 |
| STM14_RS12835 | nuoE | NADH-quinone oxidoreductase subunit NuoE | WP_000545038.1 | -0.411875095 | 0.00077222 | 0.00256849 |
| STM14_RS08465 | | MarC family NAAT transporter | WP_000968968.1 | -0.41190368 | 0.00867993 | 0.02119331 |
| STM14_RS16275 | fldB | flavodoxin FldB | WP_001055885.1 | -0.412798824 | 0.00899162 | 0.02175239 |
| STM14_RS16825 | | DUF2623 family protein | WP_001059119.1 | -0.413311023 | 0.15979526 | 0.24773311 |
| STM14_RS22895 | hflK | FtsH protease activity modulator HflK | WP_000312505.1 | -0.413775956 | 5.3092E-06 | 2.9982E-05 |
| STM14_RS23060 | | OapA family protein | WP_001119448.1 | -0.413853598 | 0.00034706 | 0.00126149 |
| STM14_RS04865 | moeA | molybdopterin molybdotransferase MoeA | WP_000395790.1 | -0.414166539 | 0.14154801 | 0.22416335 |
| STM14_RS12655 | yfaU | 2-keto-3-deoxy-L-rhamnonate aldolase | WP_000992941.1 | -0.414394492 | 0.02467332 | 0.0508378 |

| | | | | | | |
|---|---|---|---|---|---|---|
| STM14_RS23905 | | patatin family protein | WP_000531533.1 | -0.41443009 | 0.00168688 | 0.00510779 |
| STM14_RS09765 | prmC | peptide chain release factor N(5)-glutamine methyltransferase | WP_000347310.1 | -0.415920855 | 0.09418303 | 0.16025042 |
| STM14_RS21080 | glnG | nitrogen regulation protein NR(I) | WP_001188783.1 | -0.416145291 | 0.01597073 | 0.03538161 |
| STM14_RS19910 | | PTS system mannose/fructose/sorbose family transporter subunit IID | WP_001526325.1 | -0.416352567 | 0.13191454 | 0.21145379 |
| STM14_RS04955 | dacC | serine-type D-Ala-D-Ala carboxypeptidase | WP_000195709.1 | -0.417132071 | 3.8043E-05 | 0.00017935 |
| STM14_RS02470 | prpE | propionate--CoA ligase | WP_000010229.1 | -0.417321029 | 0.00019635 | 0.00076183 |
| STM14_RS08335 | | dimethyl sulfoxide reductase anchor subunit | WP_000534700.1 | -0.417560875 | 0.04045021 | 0.07761213 |
| STM14_RS02870 | lon | endopeptidase La | WP_001067723.1 | -0.417657966 | 4.8739E-06 | 2.7866E-05 |
| STM14_RS14085 | | MurR/RpiR family transcriptional regulator | WP_000995703.1 | -0.418614929 | 0.17709148 | 0.26776354 |
| STM14_RS16890 | yqhD | alcohol dehydrogenase | WP_001058724.1 | -0.418771154 | 0.00126041 | 0.00394294 |
| STM14_RS13240 | glk | glucokinase | WP_000170380.1 | -0.419072605 | 0.00053277 | 0.00185184 |
| STM14_RS12030 | | ABC transporter permease | WP_001278110.1 | -0.419285173 | 0.00998273 | 0.02385231 |
| STM14_RS15775 | cysI | assimilatory sulfite reductase (NADPH) hemoprotein subunit | WP_001290660.1 | -0.41938153 | 0.12578486 | 0.20322065 |
| STM14_RS03250 | allD | ureidoglycolate dehydrogenase | WP_000703934.1 | -0.419675678 | 0.18427481 | 0.2769065 |
| STM14_RS01630 | degP | serine endoprotease DegP | WP_000753958.1 | -0.419939611 | 3.2472E-05 | 0.00015527 |
| STM14_RS22395 | soxR | redox-sensitive transcriptional activator SoxR | WP_000412686.1 | -0.420201559 | 0.09667999 | 0.16374229 |
| STM14_RS07410 | nadE | ammonia-dependent NAD(+) synthetase | WP_000174981.1 | -0.4202424 | 2.828E-05 | 0.00013665 |
| STM14_RS16695 | | APC family permease | WP_000475950.1 | -0.420580419 | 0.13621636 | 0.2169328 |
| STM14_RS13530 | eutA | ethanolamine ammonia-lyase reactivating factor EutA | WP_001097535.1 | -0.421179152 | 0.06790534 | 0.121471 |
| STM14_RS03105 | copA | copper-exporting P-type ATPase CopA | WP_000083899.1 | -0.421529538 | 7.7687E-05 | 0.00033455 |
| STM14_RS17630 | kdsD | arabinose-5-phosphate isomerase KdsD | WP_000018617.1 | -0.422076659 | 0.00428812 | 0.0114495 |
| STM14_RS09950 | | fumarylacetoacetate hydrolase family protein | WP_001519895.1 | -0.422215232 | 0.00195293 | 0.00582492 |
| STM14_RS22660 | | GNAT family N-acetyltransferase | WP_000626099.1 | -0.422743915 | 0.1026893 | 0.17264931 |
| STM14_RS20760 | cyaA | class I adenylate cyclase | WP_000281718.1 | -0.423029527 | 2.3509E-05 | 0.00011603 |
| STM14_RS00545 | traI | conjugative transfer relaxase/helicase TraI | WP_000986921.1 | -0.423633879 | 0.00077198 | 0.00256849 |
| STM14_RS08705 | adhP | alcohol dehydrogenase AdhP | WP_000642447.1 | -0.423667669 | 1.7983E-06 | 1.1035E-05 |
| STM14_RS16790 | hypA | hydrogenase maturation nickel metallochaperone HypA | WP_000544945.1 | -0.423875499 | 0.00774017 | 0.01916563 |
| STM14_RS15270 | | formate hydrogenlyase maturation HycH family protein | WP_000003726.1 | -0.423960729 | 0.12611106 | 0.20350955 |
| STM14_RS21285 | fucO | lactaldehyde reductase | WP_000009249.1 | -0.424117539 | 0.00047051 | 0.0016594 |
| STM14_RS20285 | yidC | membrane protein insertase YidC | WP_000378281.1 | -0.425492816 | 3.1988E-05 | 0.00015316 |
| STM14_RS04880 | gsiB | glutathione ABC transporter substrate-binding protein GsiB | WP_000191434.1 | -0.426992016 | 0.00148341 | 0.00454781 |
| STM14_RS01050 | djlA | co-chaperone DjlA | WP_001200595.1 | -0.427086016 | 0.00097331 | 0.0031442 |
| STM14_RS15920 | fucI | L-fucose isomerase | WP_000724126.1 | -0.427881976 | 0.01298052 | 0.02974131 |
| STM14_RS12200 | cirA | catecholate siderophore receptor CirA | WP_000488255.1 | -0.427999995 | 0.08139182 | 0.14189616 |
| STM14_RS01205 | mraY | phospho-N-acetylmuramoyl-pentapeptide-transferase | WP_000964141.1 | -0.428232162 | 0.00248684 | 0.00717036 |
| STM14_RS13665 | | neutral zinc metallopeptidase | WP_001267523.1 | -0.428241573 | 0.00023025 | 0.00087735 |
| STM14_RS14880 | hin | flagellar phase variation DNA invertase Hin | WP_000190912.1 | -0.429648931 | 0.02927318 | 0.05902731 |
| STM14_RS15015 | | rhodanese family protein | WP_000022009.1 | -0.429837642 | 0.00813589 | 0.02007798 |
| STM14_RS16990 | cpdA | 3'%2C5'-cyclic-AMP phosphodiesterase | WP_000444735.1 | -0.429965793 | 6.158E-05 | 0.0002745 |
| STM14_RS15690 | ispD | 2-C-methyl-D-erythritol 4-phosphate cytidylyltransferase | WP_000741650.1 | -0.430045014 | 0.00482579 | 0.01263768 |
| STM14_RS00555 | | fertility inhibition protein FinO | WP_000139307.1 | -0.430537023 | 0.00216034 | 0.00634321 |
| STM14_RS07605 | ppsA | phosphoenolpyruvate synthase | WP_000069340.1 | -0.43079418 | 3.1828E-07 | 2.2844E-06 |
| STM14_RS21350 | yiiM | 6-hydroxyaminopurine reductase | WP_000559232.1 | -0.430876363 | 0.03260929 | 0.06441314 |
| STM14_RS03095 | | TraB/GumN family protein | WP_000421859.1 | -0.431117346 | 0.07854576 | 0.13791392 |
| STM14_RS04170 | ybgI | radiation resistance protein YbgI | WP_000798844.1 | -0.431824569 | 0.00013932 | 0.00056248 |
| STM14_RS09425 | osmB | osmotically-inducible lipoprotein OsmB | WP_000481166.1 | -0.432162208 | 3.6601E-06 | 2.1392E-05 |
| STM14_RS10035 | yoaE | CNNM family cation transport protein YoaE | WP_000421815.1 | -0.4332589 | 0.02951928 | 0.0594259 |
| STM14_RS09405 | | exoribonuclease II | WP_000485055.1 | -0.433597295 | 0.0001323 | 0.00053884 |
| STM14_RS07710 | sufC | Fe-S cluster assembly ATPase SufC | WP_000948902.1 | -0.433662294 | 0.00089138 | 0.00291222 |
| STM14_RS18425 | ppiA | peptidylprolyl isomerase A | WP_000920478.1 | -0.433835827 | 0.00055764 | 0.0019207 |
| STM14_RS17800 | degQ | serine endoprotease DegQ | WP_000716693.1 | -0.433861515 | 9.3007E-05 | 0.00039093 |
| STM14_RS21995 | iclR | glyoxylate bypass operon transcriptional repressor IclR | WP_000226434.1 | -0.43445793 | 0.00246761 | 0.00713167 |
| STM14_RS13535 | eutH | ethanolamine utilization protein EutH | WP_000512359.1 | -0.434510931 | 0.04522995 | 0.08544725 |
| STM14_RS19150 | bcsA | UDP-forming cellulose synthase catalytic subunit | WP_001275340.1 | -0.434609967 | 0.00155557 | 0.00475321 |
| STM14_RS15615 | | HPr family phosphocarrier protein | WP_001136888.1 | -0.435471934 | 0.01552271 | 0.03455538 |
| STM14_RS16530 | rsmE | 16S rRNA (uracil(1498)-N(3))-methyltransferase | WP_001222489.1 | -0.435473914 | 0.01762962 | 0.03831894 |
| STM14_RS00575 | thrA | bifunctional aspartate kinase/homoserine dehydrogenase I | WP_001264681.1 | -0.435525591 | 0.00083687 | 0.00275613 |
| STM14_RS17180 | | M48 family metallopeptidase | WP_000202966.1 | -0.435595204 | 0.02265471 | 0.04742162 |
| STM14_RS04890 | gsiD | glutathione ABC transporter permease GsiD | WP_001236024.1 | -0.435686535 | 0.17253766 | 0.26262244 |
| STM14_RS23260 | yjgA | ribosome biogenesis factor YjgA | WP_000166287.1 | -0.436360836 | 0.00011443 | 0.00047181 |
| STM14_RS01430 | | pyrroloquinoline quinone-dependent dehydrogenase | WP_000829730.1 | -0.436711794 | 9.6353E-05 | 0.00040361 |
| STM14_RS18890 | livJ | branched chain amino acid ABC transporter substrate-binding protein L | WP_001521443.1 | -0.43697461 | 0.01801792 | 0.03903314 |
| STM14_RS17520 | folP | dihydropteroate synthase | WP_000764711.1 | -0.437414928 | 0.00369801 | 0.01009347 |
| STM14_RS17445 | deaD | ATP-dependent RNA helicase DeaD | WP_000807248.1 | -0.437857664 | 2.989E-05 | 0.00014367 |
| STM14_RS09435 | pyrF | orotidine-5'-phosphate decarboxylase | WP_001542125.1 | -0.438266608 | 0.05756583 | 0.10550393 |
| STM14_RS07400 | spy | ATP-independent periplasmic protein-refolding chaperone Spy | WP_001228998.1 | -0.438295144 | 0.00014218 | 0.00057153 |
| STM14_RS12060 | | DedA family protein | WP_001523476.1 | -0.438866033 | 0.04431842 | 0.08407028 |
| STM14_RS17365 | | penicillin-binding protein activator | WP_000249229.1 | -0.438952063 | 0.00039327 | 0.00141271 |
| STM14_RS06340 | | YceH family protein | WP_000873047.1 | -0.438959143 | 0.00015738 | 0.00062512 |
| STM14_RS09080 | azoR | FMN-dependent NADH-azoreductase | WP_000048924.1 | -0.440206879 | 0.10587353 | 0.17671185 |
| STM14_RS13825 | bamB | outer membrane protein assembly factor BamB | WP_001177074.1 | -0.440764188 | 1.5629E-05 | 7.9815E-05 |
| STM14_RS17010 | | DUF1190 family protein | WP_000831528.1 | -0.440825552 | 6.3002E-05 | 0.00027949 |
| STM14_RS07790 | ttrR | tetrathionate respiration response regulator TtrR | WP_000190927.1 | -0.441002252 | 0.03828453 | 0.07399666 |
| STM14_RS21355 | | nuclear transport factor 2 family protein | WP_000133444.1 | -0.441740209 | 0.06501241 | 0.11716872 |
| STM14_RS01350 | aceF | pyruvate dehydrogenase complex dihydrolipoyllysine-residue acetyltrar | WP_000963612.1 | -0.442042649 | 1.0147E-05 | 5.3374E-05 |
| STM14_RS15975 | tcdA | tRNA cyclic N6-threonylcarbamoyladenosine(37) synthase TcdA | WP_000117748.1 | -0.442332993 | 0.01269007 | 0.02920303 |
| STM14_RS00770 | nhaR | transcriptional activator NhaR | WP_000062871.1 | -0.442463351 | 0.00219285 | 0.00641307 |
| STM14_RS04545 | | pyridoxal phosphatase | WP_000127031.1 | -0.442963701 | 0.00159072 | 0.00485257 |
| STM14_RS02740 | xseB | exodeoxyribonuclease VII small subunit | WP_001124944.1 | -0.443425096 | 0.00197223 | 0.00587047 |
| STM14_RS21305 | rhaS | HTH-type transcriptional activator RhaS | WP_000217112.1 | -0.443495013 | 0.00949498 | 0.02280529 |
| STM14_RS20955 | fre | NAD(P)H-flavin reductase | WP_000209831.1 | -0.443979405 | 0.00024998 | 0.00094307 |
| STM14_RS08675 | | OBAP family protein | WP_000698598.1 | -0.444127006 | 0.03300303 | 0.0649775 |

| | | | | | | |
|---|---|---|---|---|---|---|
| STM14_RS15665 | nlpD | murein hydrolase activator NlpD | WP_223149274.1 | -0.445151716 | 4.7897E-08 | 4.0356E-07 |
| STM14_RS17025 | zupT | zinc transporter ZupT | WP_000115874.1 | -0.445594613 | 0.01082146 | 0.02554142 |
| STM14_RS09590 | | YciI family protein | WP_000967605.1 | -0.445947958 | 0.03183991 | 0.06313114 |
| STM14_RS04300 | odhB | 2-oxoglutarate dehydrogenase complex dihydrolipoyllysine-residue suc | WP_000099858.1 | -0.446075232 | 1.3989E-07 | 1.0798E-06 |
| STM14_RS03100 | | DUF1311 domain-containing protein | WP_000488548.1 | -0.446081188 | 0.0805381 | 0.14060754 |
| STM14_RS09260 | bdcA | SDR family oxidoreductase | WP_000171963.1 | -0.447348137 | 0.11935062 | 0.19470492 |
| STM14_RS20665 | wecB | UDP-N-acetylglucosamine 2-epimerase (non-hydrolyzing) | WP_000866685.1 | -0.449485877 | 0.00022551 | 0.00086223 |
| STM14_RS02215 | proB | glutamate 5-kinase | WP_001285275.1 | -0.450083538 | 0.0006997 | 0.0023442 |
| STM14_RS07280 | | YeaH/YhbH family protein | WP_000219714.1 | -0.45096306 | 7.8059E-07 | 5.188E-06 |
| STM14_RS23470 | lptF | LPS export ABC transporter permease LptF | WP_000584130.1 | -0.451101883 | 0.00059384 | 0.00203397 |
| STM14_RS13910 | iscX | Fe-S cluster assembly protein IscX | WP_000523621.1 | -0.45148743 | 0.02047024 | 0.04354983 |
| STM14_RS21225 | fdhE | formate dehydrogenase accessory protein FdhE | WP_000027730.1 | -0.452038813 | 0.00064147 | 0.00217406 |
| STM14_RS19500 | mtlD | mannitol-1-phosphate 5-dehydrogenase | WP_000645391.1 | -0.452226943 | 0.00027216 | 0.00101324 |
| STM14_RS09655 | galU | UTP--glucose-1-phosphate uridylyltransferase GalU | WP_000729450.1 | -0.452380234 | 1.152E-05 | 5.974E-05 |
| STM14_RS14820 | | membrane protein | WP_001521074.1 | -0.452729714 | 0.12606445 | 0.20350955 |
| STM14_RS19765 | ligB | NAD-dependent DNA ligase LigB | WP_001241834.1 | -0.454150688 | 0.14974319 | 0.23523943 |
| STM14_RS07585 | selO | protein adenylyltransferase SelO | WP_000175669.1 | -0.454710188 | 0.00024753 | 0.00093668 |
| STM14_RS16915 | | YgiQ family radical SAM protein | WP_001274458.1 | -0.45529414 | 0.01676234 | 0.03671561 |
| STM14_RS15675 | surE | 5'/3'-nucleotidase SurE | WP_001221538.1 | -0.455465817 | 0.00376155 | 0.01023654 |
| STM14_RS01685 | ispC | 1-deoxy-D-xylulose-5-phosphate reductoisomerase | WP_000811905.1 | -0.455621143 | 0.00119888 | 0.00376322 |
| STM14_RS00045 | rck | complement resistance protein Rck | WP_000725062.1 | -0.456101503 | 0.00582108 | 0.01496733 |
| STM14_RS06275 | | LpxL/LpxP family Kdo(2)-lipid IV(A) lauroyl/palmitoleoyl acyltransferase | WP_000163973.1 | -0.456384669 | 0.02279045 | 0.04759753 |
| STM14_RS23475 | lptG | LPS export ABC transporter permease LptG | WP_001182242.1 | -0.457300963 | 0.00178343 | 0.00537696 |
| STM14_RS17410 | | SCP2 domain-containing protein | WP_000884721.1 | -0.458134727 | 4.3138E-06 | 2.4935E-05 |
| STM14_RS12315 | | Bcr/CflA family multidrug efflux MFS transporter | WP_000213347.1 | -0.458634602 | 0.0796941 | 0.1393501 |
| STM14_RS12770 | | GNAT family N-acetyltransferase | WP_000568155.1 | -0.458695476 | 0.01858334 | 0.0400608 |
| STM14_RS12940 | | LacI family DNA-binding transcriptional regulator | WP_000026949.1 | -0.45929611 | 0.00147092 | 0.00451327 |
| STM14_RS01525 | thpR | RNA 2'%2C3'-cyclic phosphodiesterase | WP_001294178.1 | -0.46049598 | 0.1246997 | 0.20164441 |
| STM14_RS04070 | tcuR | tricarballylate utilization LysR family transcriptional regulator TcuR | WP_000423363.1 | -0.460868821 | 0.01404243 | 0.03172039 |
| STM14_RS20850 | recQ | ATP-dependent DNA helicase RecQ | WP_001526553.1 | -0.46178951 | 8.6835E-06 | 4.6405E-05 |
| STM14_RS13075 | | MFS transporter | WP_000127734.1 | -0.462108814 | 0.0168641 | 0.03687269 |
| STM14_RS03150 | | ABC transporter ATP-binding protein | WP_001110598.1 | -0.462911424 | 0.00563104 | 0.01454981 |
| STM14_RS05430 | | YcbK family protein | WP_000357052.1 | -0.463238281 | 1.7106E-05 | 8.6756E-05 |
| STM14_RS16005 | recD | exodeoxyribonuclease V subunit alpha | WP_000155129.1 | -0.463683577 | 0.00988 | 0.02365289 |
| STM14_RS18995 | | NAD(P)/FAD-dependent oxidoreductase | WP_000439219.1 | -0.463718761 | 0.00301342 | 0.00847624 |
| STM14_RS21810 | rplJ | 50S ribosomal protein L10 | WP_001207203.1 | -0.464547917 | 9.7959E-08 | 7.807E-07 |
| STM14_RS19595 | envC | murein hydrolase activator EnvC | WP_058814511.1 | -0.464719994 | 0.00136055 | 0.00421678 |
| STM14_RS05705 | | phage minor tail protein L | WP_001152415.1 | -0.465351666 | 0.10347201 | 0.1736481 |
| STM14_RS05375 | lpxK | tetraacyldisaccharide 4'-kinase | WP_000561681.1 | -0.465861145 | 0.06049988 | 0.11011397 |
| STM14_RS08385 | | Zn-dependent oxidoreductase | WP_000836487.1 | -0.465959445 | 0.11237684 | 0.18571433 |
| STM14_RS14065 | tadA | tRNA adenosine(34) deaminase TadA | WP_001134566.1 | -0.466126136 | 0.02978688 | 0.05989912 |
| STM14_RS07990 | cfa | cyclopropane fatty acyl phospholipid synthase | WP_000098879.1 | -0.466149865 | 5.0345E-06 | 2.8607E-05 |
| STM14_RS11560 | hisG | ATP phosphoribosyltransferase | WP_000886603.1 | -0.466456076 | 0.00825178 | 0.02028241 |
| STM14_RS16075 | mutH | DNA mismatch repair endonuclease MutH | WP_001274930.1 | -0.466544198 | 0.01401704 | 0.03168248 |
| STM14_RS01200 | murF | UDP-N-acetylmuramoyl-tripeptide--D-alanyl-D-alanine ligase | WP_000626630.1 | -0.466819202 | 0.00014955 | 0.00059723 |
| STM14_RS01860 | dkgB | 2%2C5-didehydrogluconate reductase DkgB | WP_000154871.1 | -0.467115877 | 0.14593431 | 0.23031725 |
| STM14_RS20125 | | DUF3748 domain-containing protein | WP_000809945.1 | -0.467591802 | 0.02169508 | 0.04567254 |
| STM14_RS13520 | eutC | ethanolamine ammonia-lyase subunit EutC | WP_000372335.1 | -0.468023606 | 0.01034524 | 0.02455911 |
| STM14_RS08330 | dmsD | Tat proofreading chaperone DmsD | WP_000206561.1 | -0.468500745 | 0.00419443 | 0.01126149 |
| STM14_RS04860 | moeB | molybdopterin-synthase adenylyltransferase MoeB | WP_000829280.1 | -0.468844685 | 0.26539297 | 0.37105349 |
| STM14_RS04290 | sdhB | succinate dehydrogenase iron-sulfur subunit SdhB | WP_001537859.1 | -0.468894191 | 2.7982E-07 | 2.0202E-06 |
| STM14_RS01290 | | GMP reductase | WP_001217365.1 | -0.469830439 | 1.5772E-05 | 8.0435E-05 |
| STM14_RS20490 | rbsA | ribose ABC transporter ATP-binding protein RbsA | WP_000339359.1 | -0.470618387 | 0.11918011 | 0.19468553 |
| STM14_RS19555 | | L-talarate/galactarate dehydratase | WP_001218249.1 | -0.471580587 | 0.08077843 | 0.14096027 |
| STM14_RS15660 | rpoS | RNA polymerase sigma factor RpoS | WP_000081498.1 | -0.471619859 | 1.5856E-08 | 1.4416E-07 |
| STM14_RS01055 | rluA | bifunctional tRNA pseudouridine(32) synthase/23S rRNA pseudouridine | WP_000525196.1 | -0.471658519 | 0.10328368 | 0.17349019 |
| STM14_RS06470 | fabH | beta-ketoacyl-ACP synthase III | WP_000288154.1 | -0.472124725 | 0.00084601 | 0.00277918 |
| STM14_RS08250 | pntA | Re/Si-specific NAD(P)(+) transhydrogenase subunit alpha | WP_001219362.1 | -0.473876643 | 1.2693E-06 | 8.0025E-06 |
| STM14_RS11325 | | cobalt-factor II C(20)-methyltransferase | WP_001013552.1 | -0.474418715 | 0.01156213 | 0.0269783 |
| STM14_RS22860 | nnr | bifunctional ADP-dependent NAD(P)H-hydrate dehydratase/NAD(P)H-h | WP_000968675.1 | -0.474551069 | 0.00027609 | 0.00102564 |
| STM14_RS08545 | | nickel-dependent hydrogenase large subunit | WP_000331199.1 | -0.475254518 | 0.00169333 | 0.00512312 |
| STM14_RS10130 | | GAF domain-containing protein | WP_001518229.1 | -0.475316639 | 5.7386E-05 | 0.00025862 |
| STM14_RS16800 | | HyaD/HybD family hydrogenase maturation endopeptidase | WP_001221960.1 | -0.475929348 | 0.0003527 | 0.00127817 |
| STM14_RS21990 | | ASCH domain-containing protein | WP_010989091.1 | -0.476016887 | 0.17715154 | 0.26776354 |
| STM14_RS16100 | aas | bifunctional acyl-ACP--phospholipid O-acyltransferase/long-chain-fatty | WP_000896102.1 | -0.476407094 | 0.00055246 | 0.00190712 |
| STM14_RS15350 | sitB | iron/manganese ABC transporter ATP-binding protein SitB | WP_000083153.1 | -0.477201898 | 0.00054035 | 0.00187518 |
| STM14_RS23635 | | aspartate/glutamate racemase family protein | WP_000683247.1 | -0.477697705 | 6.192E-06 | 3.4335E-05 |
| STM14_RS16345 | pepP | Xaa-Pro aminopeptidase | WP_000193679.1 | -0.477816194 | 2.8788E-06 | 1.7179E-05 |
| STM14_RS04500 | galT | galactose-1-phosphate uridylyltransferase | WP_000187473.1 | -0.478978377 | 0.00636355 | 0.01619252 |
| STM14_RS11580 | hisH | imidazole glycerol phosphate synthase subunit HisH | WP_001103591.1 | -0.479094459 | 0.02160215 | 0.04558115 |
| STM14_RS07480 | | fructosamine kinase family protein | WP_000267686.1 | -0.479693481 | 2.668E-07 | 1.9337E-06 |
| STM14_RS12880 | | sugar phosphatase | WP_014343872.1 | -0.480023652 | 0.00082335 | 0.0027172 |
| STM14_RS09870 | | alpha%2Calpha-trehalase | WP_000612821.1 | -0.480757518 | 9.6777E-05 | 0.00040446 |
| STM14_RS21365 | cpxR | envelope stress response regulator transcription factor CpxR | WP_001033731.1 | -0.481126671 | 6.2452E-05 | 0.00027771 |
| STM14_RS21490 | glpX | class II fructose-bisphosphatase | WP_001250625.1 | -0.481173092 | 0.00328091 | 0.00913099 |
| STM14_RS17965 | accC | acetyl-CoA carboxylase biotin carboxylase subunit | WP_000884627.1 | -0.481573308 | 3.0247E-07 | 2.1752E-06 |
| STM14_RS22900 | hflC | protease modulator HflC | WP_001232417.1 | -0.481604761 | 1.482E-06 | 9.1864E-06 |
| STM14_RS13505 | eutR | HTH-type transcriptional regulator EutR | WP_000753665.1 | -0.482011981 | 0.00386066 | 0.01049073 |
| STM14_RS16945 | parC | DNA topoisomerase IV subunit A | WP_001281908.1 | -0.482716961 | 0.00015373 | 0.00061194 |
| STM14_RS11050 | | glycoside hydrolase family 19 protein | WP_001005901.1 | -0.48276191 | 0.11003338 | 0.1824968 |
| STM14_RS08170 | | oxidoreductase | WP_000998242.1 | -0.483200926 | 0.00011666 | 0.00047887 |
| STM14_RS04570 | | histidine utilization repressor | WP_000287416.1 | -0.483389276 | 0.00197408 | 0.0058712 |

| | | | | | | |
|---|---|---|---|---|---|---|
| STM14_RS07795 | fumD | fumarate hydratase FumD | WP_000165779.1 | -0.483400944 | 0.02752795 | 0.05606079 |
| STM14_RS09595 | cls | cardiolipin synthase | WP_000206886.1 | -0.483745191 | 0.00351604 | 0.00966123 |
| STM14_RS23185 | iolT2 | myo-inositol import MFS transporter IolT2 | WP_000083110.1 | -0.484501911 | 0.078788 | 0.13814163 |
| STM14_RS11595 | hisIE | bifunctional phosphoribosyl-AMP cyclohydrolase/phosphoribosyl-ATP d | WP_000954848.1 | -0.485775082 | 0.02144928 | 0.04533654 |
| STM14_RS07680 | | FAD-binding and (Fe-S)-binding domain-containing protein | WP_000612921.1 | -0.486668567 | 7.5148E-05 | 0.00032514 |
| STM14_RS18665 | | TROVE domain-containing protein | WP_000013008.1 | -0.4869304 | 0.00114588 | 0.00361537 |
| STM14_RS08245 | ydgH | DUF1471 family protein YdgH | WP_000769294.1 | -0.488441754 | 5.308E-06 | 2.9982E-05 |
| STM14_RS08765 | narH | nitrate reductase subunit beta | WP_000702596.1 | -0.488500612 | 0.01814504 | 0.03927691 |
| STM14_RS16330 | gcvT | glycine cleavage system aminomethyltransferase GcvT | WP_000068738.1 | -0.488566403 | 5.7875E-08 | 4.7779E-07 |
| STM14_RS20940 | tatD | 3'-5' ssDNA/RNA exonuclease TatD | WP_000459635.1 | -0.48871309 | 0.13561717 | 0.21644665 |
| STM14_RS15230 | gutQ | arabinose-5-phosphate isomerase GutQ | WP_001278200.1 | -0.48975487 | 0.00135285 | 0.0042 |
| STM14_RS02620 | proY | proline-specific permease ProY | WP_001555879.1 | -0.489898131 | 0.02173235 | 0.04572486 |
| STM14_RS20505 | rbsK | ribokinase | WP_000844728.1 | -0.490355604 | 6.7772E-05 | 0.00029707 |
| STM14_RS13595 | maeB | NADP-dependent oxaloacetate-decarboxylating malate dehydrogenase | WP_000344297.1 | -0.490682907 | 8.182E-08 | 6.6357E-07 |
| STM14_RS14660 | recN | DNA repair protein RecN | WP_000880965.1 | -0.491034716 | 0.02841756 | 0.05752072 |
| STM14_RS06585 | mfd | transcription-repair coupling factor | WP_001114323.1 | -0.491538534 | 8.9273E-05 | 0.00037739 |
| STM14_RS00590 | yaaA | peroxide stress protein YaaA | WP_000906175.1 | -0.492055481 | 0.00030499 | 0.00112521 |
| STM14_RS15260 | | hypothetical protein | WP_000902280.1 | -0.493601537 | 0.01155793 | 0.0269783 |
| STM14_RS13600 | tal | transaldolase | WP_001072448.1 | -0.49390196 | 1.4601E-05 | 7.4877E-05 |
| STM14_RS06510 | holB | DNA polymerase III subunit delta' | WP_000872453.1 | -0.494236759 | 0.01213123 | 0.02811026 |
| STM14_RS18635 | malP | maltodextrin phosphorylase | WP_000082246.1 | -0.494352955 | 0.00015175 | 0.0006047 |
| STM14_RS13060 | | aspartate-semialdehyde dehydrogenase | WP_001289141.1 | -0.494358603 | 0.00080832 | 0.00267645 |
| STM14_RS06235 | ymdB | O-acetyl-ADP-ribose deacetylase | WP_000203944.1 | -0.496271292 | 0.00667792 | 0.01688742 |
| STM14_RS18470 | trpS | tryptophan--tRNA ligase | WP_000165562.1 | -0.496960676 | 8.598E-06 | 4.6092E-05 |
| STM14_RS15555 | | hypothetical protein | WP_000393302.1 | -0.497248507 | 0.03548522 | 0.06931384 |
| STM14_RS05205 | | pirin family protein | WP_000809973.1 | -0.497337677 | 0.03460303 | 0.06773437 |
| STM14_RS15840 | | glycerate kinase | WP_000706479.1 | -0.498015415 | 0.00319072 | 0.00892722 |
| STM14_RS20815 | uvrD | DNA helicase II | WP_000383441.1 | -0.498672122 | 3.3324E-05 | 0.00015894 |
| STM14_RS11910 | fbaB | class I fructose-bisphosphate aldolase | WP_000129590.1 | -0.499061956 | 9.3829E-05 | 0.00039393 |
| STM14_RS05400 | elyC | envelope biogenesis factor ElyC | WP_000899563.1 | -0.500087593 | 0.16091203 | 0.24885865 |
| STM14_RS02920 | | SmdA family multidrug ABC transporter permease/ATP-binding protein | WP_001235572.1 | -0.501109078 | 0.00767211 | 0.01900991 |
| STM14_RS09875 | | GlsB/YeaQ/YmgE family stress response membrane protein | WP_000511323.1 | -0.501267079 | 2.4527E-05 | 0.00012041 |
| STM14_RS07350 | topB | DNA topoisomerase III | WP_001235865.1 | -0.501755952 | 8.8622E-05 | 0.0003755 |
| STM14_RS21715 | murI | glutamate racemase | WP_000201805.1 | -0.502287314 | 0.0004396 | 0.00156386 |
| STM14_RS12225 | fruA | PTS fructose transporter subunit IIBC | WP_000854414.1 | -0.503244692 | 0.01420808 | 0.03203562 |
| STM14_RS15210 | srlD | sorbitol-6-phosphate dehydrogenase | WP_001077372.1 | -0.503413291 | 0.07148657 | 0.127217 |
| STM14_RS05435 | | MBL fold metallo-hydrolase | WP_001109471.1 | -0.503573983 | 0.00010808 | 0.00044867 |
| STM14_RS08485 | glsB | glutaminase B | WP_000090081.1 | -0.504191821 | 0.00335185 | 0.00931434 |
| STM14_RS04795 | rhtA | threonine/homoserine exporter RhtA | WP_001119554.1 | -0.504949547 | 0.20602634 | 0.30310495 |
| STM14_RS18825 | | sn-glycerol-3-phosphate import ATP-binding protein UgpC | WP_000907838.1 | -0.505509015 | 0.00532489 | 0.0138072 |
| STM14_RS04760 | | DksA/TraR family C4-type zinc finger protein | WP_000146330.1 | -0.505771453 | 8.6344E-05 | 0.00036796 |
| STM14_RS01210 | murD | UDP-N-acetylmuramoyl-L-alanine--D-glutamate ligase | WP_000796445.1 | -0.505856574 | 4.4559E-05 | 0.00020742 |
| STM14_RS00815 | ispH | 4-hydroxy-3-methylbut-2-enyl diphosphate reductase | WP_001166422.1 | -0.506900271 | 0.00058931 | 0.00202223 |
| STM14_RS18630 | malQ | 4-alpha-glucanotransferase | WP_000428721.1 | -0.507489148 | 0.00014148 | 0.00056933 |
| STM14_RS20260 | dnaN | DNA polymerase III subunit beta | WP_000673478.1 | -0.507984372 | 0.00011206 | 0.00046256 |
| STM14_RS04870 | iaaA | beta-aspartyl-peptidase | WP_001030503.1 | -0.509195719 | 0.00014917 | 0.00059637 |
| STM14_RS03640 | dsbG | thiol:disulfide interchange protein DsbG | WP_000597122.1 | -0.509326363 | 0.00060223 | 0.00205505 |
| STM14_RS14960 | | tripartite tricarboxylate transporter TctB family protein | WP_001574835.1 | -0.511203049 | 0.05278599 | 0.09786405 |
| STM14_RS05030 | potG | putrescine ABC transporter ATP-binding subunit PotG | WP_001000698.1 | -0.511298546 | 0.00765142 | 0.01897981 |
| STM14_RS19790 | recG | ATP-dependent DNA helicase RecG | WP_000016748.1 | -0.511577995 | 0.00075474 | 0.00251946 |
| STM14_RS18570 | envZ | two-component system sensor histidine kinase EnvZ | WP_001253818.1 | -0.511733669 | 0.00017326 | 0.00068012 |
| STM14_RS09675 | purU | formyltetrahydrofolate deformylase | WP_001191156.1 | -0.512244727 | 6.6067E-06 | 3.658E-05 |
| STM14_RS00155 | | CaiF/GrlA family transcriptional regulator | WP_001075507.1 | -0.512414892 | 0.13870716 | 0.22032776 |
| STM14_RS05795 | rlmKL | bifunctional 23S rRNA (guanine(2069)-N(7))-methyltransferase RlmK/23 | WP_001086485.1 | -0.512994709 | 0.00014088 | 0.00056752 |
| STM14_RS03040 | | YbaB/EbfC family nucleoid-associated protein | WP_000467098.1 | -0.513602321 | 9.676E-05 | 0.00040446 |
| STM14_RS18575 | ompR | two-component system response regulator OmpR | WP_001157751.1 | -0.514001585 | 1.916E-06 | 1.1719E-05 |
| STM14_RS00580 | thrB | homoserine kinase | WP_000241685.1 | -0.514630564 | 1.7418E-05 | 8.8216E-05 |
| STM14_RS03845 | lptE | LPS assembly lipoprotein LptE | WP_001269947.1 | -0.514720928 | 9.7051E-05 | 0.00040515 |
| STM14_RS05010 | nfsA | nitroreductase NfsA | WP_000075300.1 | -0.514939571 | 0.10750027 | 0.17877867 |
| STM14_RS16980 | parE | DNA topoisomerase IV subunit B | WP_000195318.1 | -0.51512603 | 1.3564E-06 | 8.4939E-06 |
| STM14_RS03155 | | ABC transporter permease | WP_000561776.1 | -0.515399831 | 0.00048503 | 0.00170246 |
| STM14_RS09790 | prs | ribose-phosphate diphosphokinase | WP_001518537.1 | -0.515744281 | 9.7755E-09 | 9.2528E-08 |
| STM14_RS09920 | nhaB | Na(+)/H(+) antiporter NhaB | WP_000406435.1 | -0.515977184 | 0.00053312 | 0.00185184 |
| STM14_RS18585 | | Tex family protein | WP_000980662.1 | -0.51654435 | 2.1532E-05 | 0.00010757 |
| STM14_RS00140 | | hypothetical protein | WP_000198608.1 | -0.516864775 | 0.0006372 | 0.00216237 |
| STM14_RS18900 | ftsX | permease-like cell division protein FtsX | WP_001081707.1 | -0.516871179 | 0.00052734 | 0.00183533 |
| STM14_RS20915 | ubiJ | ubiquinone biosynthesis protein UbiJ | WP_001116086.1 | -0.517129897 | 8.8476E-05 | 0.00037531 |
| STM14_RS17780 | rpsI | 30S ribosomal protein S9 | WP_000829815.1 | -0.517294185 | 4.5966E-07 | 3.1753E-06 |
| STM14_RS22000 | metH | methionine synthase | WP_000095958.1 | -0.517303882 | 5.7576E-05 | 0.00025916 |
| STM14_RS15085 | proW | glycine betaine/L-proline ABC transporter permease ProW | WP_000775022.1 | -0.517370447 | 0.00490949 | 0.0128386 |
| STM14_RS01440 | can | carbonate dehydratase | WP_000651591.1 | -0.517412708 | 9.2072E-06 | 4.9061E-05 |
| STM14_RS13470 | | RpoE-regulated lipoprotein | WP_000838976.1 | -0.517522616 | 0.00886453 | 0.02155826 |
| STM14_RS16935 | ftsP | cell division protein FtsP | WP_000010658.1 | -0.518169761 | 0.00035444 | 0.00128324 |
| STM14_RS12970 | hisP | histidine ABC transporter ATP-binding protein HisP | WP_001293591.1 | -0.518845029 | 0.00059067 | 0.002025 |
| STM14_RS20420 | atpH | F0F1 ATP synthase subunit delta | WP_001288957.1 | -0.519543559 | 7.0501E-07 | 4.7284E-06 |
| STM14_RS09065 | cybB | cytochrome b561 | WP_000527273.1 | -0.519578651 | 0.008625 | 0.02107316 |
| STM14_RS02725 | | aldo/keto reductase | WP_001199859.1 | -0.519870215 | 1.1897E-07 | 9.3597E-07 |
| STM14_RS08025 | rnt | ribonuclease T | WP_001282267.1 | -0.520478649 | 0.00035224 | 0.00127777 |
| STM14_RS13105 | fabB | beta-ketoacyl-ACP synthase I | WP_000817161.1 | -0.52112745 | 3.8165E-08 | 3.2604E-07 |
| STM14_RS07340 | | NAD(P)H nitroreductase | WP_000339313.1 | -0.522166403 | 3.7903E-05 | 0.00017892 |
| STM14_RS12790 | nuoN | NADH-quinone oxidoreductase subunit NuoN | WP_000156671.1 | -0.522338746 | 5.3832E-05 | 0.0002444 |
| STM14_RS12015 | | protein YohO | WP_001261696.1 | -0.523253656 | 0.00478489 | 0.0125484 |

| | | | | | | |
|---|---|---|---|---|---|---|
| STM14_RS20070 | | cellulase family glycosylhydrolase | WP_000238929.1 | -0.523674306 | 0.01180209 | 0.02743389 |
| STM14_RS12745 | menB | 1%2C4-dihydroxy-2-naphthoyl-CoA synthase | WP_000640013.1 | -0.525063665 | 8.0714E-05 | 0.00034557 |
| STM14_RS06505 | tmk | dTMP kinase | WP_000535399.1 | -0.525481613 | 0.00059944 | 0.00204935 |
| STM14_RS13045 | accD | acetyl-CoA carboxylase%2C carboxyltransferase subunit beta | WP_000118383.1 | -0.525925612 | 4.889E-05 | 0.00022473 |
| STM14_RS01620 | mtnN | 5'-methylthioadenosine/S-adenosylhomocysteine nucleosidase | WP_000689821.1 | -0.526574002 | 0.00139369 | 0.00430502 |
| STM14_RS00805 | lspA | signal peptidase II | WP_000042739.1 | -0.526975984 | 0.00044554 | 0.00157938 |
| STM14_RS13555 | eutN | ethanolamine utilization microcompartment protein EutN | WP_001522340.1 | -0.527021124 | 0.11981284 | 0.19528591 |
| STM14_RS13885 | | alpha-2-macroglobulin | WP_000681859.1 | -0.527284127 | 4.2697E-07 | 2.9831E-06 |
| STM14_RS06535 | | YcfL family protein | WP_000589563.1 | -0.52739988 | 9.0418E-05 | 0.00038091 |
| STM14_RS12675 | | nicotinamide mononucleotide deamidase-related protein YfaY | WP_000935674.1 | -0.528048466 | 6.2269E-05 | 0.00027723 |
| STM14_RS16125 | | aspartate/glutamate racemase | WP_000848625.1 | -0.528590985 | 0.00035762 | 0.00129219 |
| STM14_RS18655 | rtcA | RNA 3'-terminal phosphate cyclase | WP_000101027.1 | -0.530381232 | 0.0349166 | 0.06827559 |
| STM14_RS02715 | thiL | thiamine-phosphate kinase | WP_000752138.1 | -0.530409709 | 0.00050936 | 0.00178106 |
| STM14_RS25180 | | PagK family vesicle-borne virulence factor | WP_001542312.1 | -0.531354415 | 0.00511293 | 0.01333274 |
| STM14_RS17115 | tsaD | tRNA (adenosine(37)-N6)-threonylcarbamoyltransferase complex trans | WP_001264394.1 | -0.531887195 | 0.00153255 | 0.00468676 |
| STM14_RS01355 | lpdA | dihydrolipoyl dehydrogenase | WP_001519338.1 | -0.531967464 | 1.6525E-10 | 2.0081E-09 |
| STM14_RS15275 | hycG | formate hydrogenlyase subunit HycG | WP_000067419.1 | -0.532926293 | 0.0739498 | 0.13090537 |
| STM14_RS18810 | ggt | gamma-glutamyltransferase | WP_000805096.1 | -0.534013187 | 3.6453E-05 | 0.00017296 |
| STM14_RS17615 | mlaE | lipid asymmetry maintenance ABC transporter permease subunit MlaE | WP_000925786.1 | -0.534411655 | 0.00746072 | 0.01857453 |
| STM14_RS03320 | folD | bifunctional methylenetetrahydrofolate dehydrogenase/methenyltetrah | WP_000729165.1 | -0.536800475 | 0.00013508 | 0.00054898 |
| STM14_RS04410 | pnuC | nicotinamide riboside transporter PnuC | WP_000345368.1 | -0.53681538 | 0.00893121 | 0.02166164 |
| STM14_RS13830 | | YfgM family protein | WP_000409190.1 | -0.53684262 | 2.599E-05 | 0.00012675 |
| STM14_RS03335 | fimC | type 1 fimbria chaperone FimC | WP_000935025.1 | -0.536876994 | 1.3778E-07 | 1.0658E-06 |
| STM14_RS17795 | zapG | Z-ring associated protein ZapG | WP_001519074.1 | -0.537591861 | 2.261E-07 | 1.6683E-06 |
| STM14_RS26275 | | type I toxin-antitoxin system toxin Ldr family protein | WP_000176312.1 | -0.538032748 | 3.9613E-05 | 0.00018604 |
| STM14_RS11550 | | LysR family transcriptional regulator | WP_000803316.1 | -0.53810867 | 1.9568E-05 | 9.8564E-05 |
| STM14_RS05410 | mukF | chromosome partition protein MukF | WP_001288828.1 | -0.538550558 | 0.000344 | 0.00125281 |
| STM14_RS22615 | | dimethyl sulfoxide reductase anchor subunit | WP_000544929.1 | -0.539196333 | 0.00011587 | 0.00047614 |
| STM14_RS23065 | fklB | FKBP-type peptidyl-prolyl cis-trans isomerase | WP_000235161.1 | -0.539348276 | 5.9259E-05 | 0.00026544 |
| STM14_RS05210 | | LysR family transcriptional regulator | WP_000597928.1 | -0.539584203 | 0.00272542 | 0.00776702 |
| STM14_RS20635 | gppA | guanosine-5'-triphosphate%2C3'-diphosphate diphosphatase | WP_001089447.1 | -0.542138704 | 5.0739E-05 | 0.00023237 |
| STM14_RS11835 | mdtA | multidrug efflux RND transporter subunit MdtA | WP_000678822.1 | -0.542819545 | 0.02538403 | 0.05215624 |
| STM14_RS12265 | | GTP-binding protein | WP_000169350.1 | -0.544282351 | 0.00430084 | 0.01146684 |
| STM14_RS19340 | glyS | glycine--tRNA ligase subunit beta | WP_001291736.1 | -0.545526356 | 5.1226E-06 | 2.9062E-05 |
| STM14_RS12025 | | ABC transporter ATP-binding protein | WP_000569166.1 | -0.546277088 | 0.01084222 | 0.02554745 |
| STM14_RS05880 | | CoA-binding protein | WP_001537782.1 | -0.547156105 | 0.00040042 | 0.00143417 |
| STM14_RS25900 | | IS3-like element ISSen1 family transposase | WP_085983316.1 | -0.548468103 | 5.0022E-05 | 0.00022965 |
| STM14_RS09345 | pspF | phage shock protein operon transcriptional activator | WP_001738297.1 | -0.548610502 | 0.01640884 | 0.03617575 |
| STM14_RS15670 | | protein-L-isoaspartate(D-aspartate) O-methyltransferase | WP_000253545.1 | -0.549494722 | 0.00135543 | 0.00420445 |
| STM14_RS12815 | nuoI | NADH-quinone oxidoreductase subunit NuoI | WP_000172747.1 | -0.550027777 | 9.8627E-06 | 5.2027E-05 |
| STM14_RS12800 | nuoL | NADH-quinone oxidoreductase subunit L | WP_001056587.1 | -0.550263587 | 1.4928E-07 | 1.1403E-06 |
| STM14_RS04640 | moaB | molybdenum cofactor biosynthesis protein B | WP_000084604.1 | -0.550344784 | 8.8195E-05 | 0.00037467 |
| STM14_RS08415 | dcp | peptidyl-dipeptidase Dcp | WP_000106265.1 | -0.551622154 | 6.9531E-05 | 0.00030369 |
| STM14_RS08200 | fumC | class II fumarate hydratase | WP_000259121.1 | -0.553670391 | 7.2472E-10 | 7.8024E-09 |
| STM14_RS20835 | rarD | EamA family transporter RarD | WP_000339080.1 | -0.554154899 | 0.01907597 | 0.04100275 |
| STM14_RS19905 | dgaE | D-glucosaminate-6-phosphate ammonia lyase | WP_000188463.1 | -0.555353438 | 0.03028749 | 0.06070687 |
| STM14_RS12010 | mlrA | HTH-type transcriptional regulator MlrA | WP_001240418.1 | -0.556260825 | 7.728E-05 | 0.00033319 |
| STM14_RS23900 | | DUF1328 domain-containing protein | WP_000490276.1 | -0.556743993 | 1.511E-10 | 1.8545E-09 |
| STM14_RS17200 | sstT | serine/threonine transporter SstT | WP_000235363.1 | -0.557394348 | 1.1891E-06 | 7.5615E-06 |
| STM14_RS17145 | | siderophore-interacting protein | WP_000213760.1 | -0.558566111 | 0.02805266 | 0.0569404 |
| STM14_RS06595 | lolD | lipoprotein-releasing ABC transporter ATP-binding protein LolD | WP_001033714.1 | -0.559293755 | 0.02095756 | 0.04442472 |
| STM14_RS09115 | | DUF333 domain-containing protein | WP_001200680.1 | -0.559603679 | 0.00022778 | 0.00086911 |
| STM14_RS03455 | | autotransporter domain-containing esterase | WP_000192436.1 | -0.560120984 | 2.5653E-05 | 0.00012544 |
| STM14_RS20700 | | TDP-N-acetylfucosamine:lipid II N-acetylfucosaminyltransferase | WP_000217186.1 | -0.560879873 | 0.00348755 | 0.0096015 |
| STM14_RS22300 | alr | alanine racemase | WP_001147297.1 | -0.561820416 | 0.0098695 | 0.02364314 |
| STM14_RS18340 | slyD | peptidylprolyl isomerase | WP_000861341.1 | -0.561924501 | 1.4496E-09 | 1.5077E-08 |
| STM14_RS19040 | gorA | glutathione-disulfide reductase | WP_000160774.1 | -0.563242882 | 1.2431E-07 | 9.7383E-07 |
| STM14_RS11340 | | precorrin-3B C(17)-methyltransferase | WP_000953646.1 | -0.564395907 | 0.00479258 | 0.01255963 |
| STM14_RS17515 | glmM | phosphoglucosamine mutase | WP_000071169.1 | -0.564675822 | 3.8382E-06 | 2.2291E-05 |
| STM14_RS19400 | | 4Fe-4S dicluster domain-containing protein | WP_001078820.1 | -0.564764361 | 0.0028549 | 0.00809218 |
| STM14_RS01015 | folA | type 3 dihydrofolate reductase | WP_000624379.1 | -0.565377845 | 0.00412515 | 0.01114366 |
| STM14_RS02200 | frsA | esterase FrsA | WP_000189588.1 | -0.565490016 | 7.5634E-06 | 4.1259E-05 |
| STM14_RS16575 | | XTP/dITP diphosphatase | WP_001174769.1 | -0.565733942 | 0.00937182 | 0.02255362 |
| STM14_RS10325 | purT | formate-dependent phosphoribosylglycinamide formyltransferase | WP_000173431.1 | -0.567119122 | 0.05979779 | 0.10905174 |
| STM14_RS13065 | pdxB | 4-phosphoerythronate dehydrogenase PdxB | WP_000699183.1 | -0.567199312 | 2.8117E-05 | 0.00013604 |
| STM14_RS00795 | ribF | bifunctional riboflavin kinase/FAD synthetase | WP_000767311.1 | -0.567210728 | 8.1522E-06 | 4.414E-05 |
| STM14_RS16965 | qseC | two-component system sensor histidine kinase QseC | WP_000779328.1 | -0.567306833 | 0.05998783 | 0.10923601 |
| STM14_RS17420 | | U32 family peptidase | WP_014343907.1 | -0.567412859 | 0.00659811 | 0.01670855 |
| STM14_RS01380 | | HNH endonuclease | WP_000536990.1 | -0.568402032 | 0.02802905 | 0.05692385 |
| STM14_RS21610 | | glycerol dehydrogenase | WP_000374031.1 | -0.56854755 | 1.9433E-09 | 1.9876E-08 |
| STM14_RS09770 | prfA | peptide chain release factor 1 | WP_000804703.1 | -0.569361771 | 2.8655E-05 | 0.00013828 |
| STM14_RS20510 | rbsR | ribose operon transcriptional repressor RbsR | WP_000224491.1 | -0.570118983 | 0.00083981 | 0.00276335 |
| STM14_RS06780 | | ead/Ea22-like family protein | WP_000065092.1 | -0.570563602 | 0.08351045 | 0.14524585 |
| STM14_RS02830 | cyoB | cytochrome o ubiquinol oxidase subunit I | WP_000467158.1 | -0.57124762 | 2.3729E-10 | 2.7825E-09 |
| STM14_RS03605 | cstA | pyruvate/proton symporter CstA | WP_001043108.1 | -0.571313679 | 7.1139E-10 | 7.701E-09 |
| STM14_RS00145 | | site-specific integrase | WP_000082169.1 | -0.571817941 | 6.5864E-05 | 0.00028939 |
| STM14_RS01655 | glnD | bifunctional uridylyltransferase/uridylyl-removing protein GlnD | WP_001094519.1 | -0.571981282 | 5.2679E-05 | 0.00024006 |
| STM14_RS04090 | ybfF | esterase | WP_000773246.1 | -0.57370063 | 0.00442611 | 0.01175824 |
| STM14_RS14070 | yfhb | phosphatidylglycerophosphatase C | WP_000253558.1 | -0.574193417 | 1.2952E-06 | 8.1384E-06 |
| STM14_RS10965 | | phage tail tube protein | WP_000515952.1 | -0.574276402 | 0.02102837 | 0.04454918 |
| STM14_RS09615 | | ABC transporter ATP-binding protein | WP_000058857.1 | -0.574841294 | 7.6578E-07 | 5.108E-06 |
| STM14_RS10315 | yebF | protein YebF | WP_001042119.1 | -0.575406281 | 1.5683E-06 | 9.6726E-06 |

| | | | | | | |
|---|---|---|---|---|---|---|
| STM14_RS17305 | garK | glycerate 2-kinase | WP_000706459.1 | -0.575556864 | 0.00321148 | 0.00897848 |
| STM14_RS10400 | ruvC | crossover junction endodeoxyribonuclease RuvC | WP_000022509.1 | -0.575755871 | 0.03126777 | 0.06226497 |
| STM14_RS16325 | gcvH | glycine cleavage system protein GcvH | WP_000073211.1 | -0.575950016 | 6.3987E-09 | 6.2163E-08 |
| STM14_RS10155 | | YebW family protein | WP_000457836.1 | -0.576177234 | 0.06490625 | 0.11703467 |
| STM14_RS10340 | zwf | glucose-6-phosphate dehydrogenase | WP_000301711.1 | -0.577212469 | 2.5796E-07 | 1.8833E-06 |
| STM14_RS06350 | murJ | murein biosynthesis integral membrane protein MurJ | WP_001537775.1 | -0.577368347 | 0.00253388 | 0.00728885 |
| STM14_RS16775 | | molybdate ABC transporter substrate-binding protein | WP_001220944.1 | -0.578531901 | 0.02489637 | 0.0512687 |
| STM14_RS12965 | | TIGR01777 family oxidoreductase | WP_001166274.1 | -0.579759216 | 0.00021841 | 0.0008377 |
| STM14_RS04875 | gsiA | glutathione ABC transporter ATP-binding protein GsiA | WP_001120598.1 | -0.580306726 | 1.1272E-05 | 5.8618E-05 |
| STM14_RS06345 | | Gfo/Idh/MocA family oxidoreductase | WP_001259729.1 | -0.58045431 | 7.0087E-06 | 3.8574E-05 |
| STM14_RS00800 | ileS | isoleucine--tRNA ligase | WP_001670710.1 | -0.581265685 | 4.0614E-10 | 4.5732E-09 |
| STM14_RS11285 | cobT | nicotinate-nucleotide--dimethylbenzimidazole phosphoribosyltransfera | WP_001193983.1 | -0.581498028 | 0.00044963 | 0.00159185 |
| STM14_RS13735 | purN | phosphoribosylglycinamide formyltransferase | WP_001028593.1 | -0.58213261 | 0.00093662 | 0.00304379 |
| STM14_RS18355 | kefG | glutathione-regulated potassium-efflux system ancillary protein KefG | WP_000081820.1 | -0.582370956 | 0.01083107 | 0.02554745 |
| STM14_RS03965 | ubiF | 3-demethoxyubiquinol 3-hydroxylase | WP_000184633.1 | -0.584262111 | 8.4969E-07 | 5.5767E-06 |
| STM14_RS09995 | tsaB | tRNA (adenosine(37)-N6)-threonylcarbamoyltransferase complex dime | WP_001221014.1 | -0.584339386 | 0.00037266 | 0.00134522 |
| STM14_RS04705 | | YbhQ family protein | WP_000871970.1 | -0.584838858 | 3.5926E-08 | 3.0834E-07 |
| STM14_RS19610 | tdh | L-threonine 3-dehydrogenase | WP_000645990.1 | -0.585460196 | 6.9449E-08 | 5.6951E-07 |
| STM14_RS17915 | | Maf family protein | WP_000210306.1 | -0.586058827 | 0.00343384 | 0.00949203 |
| STM14_RS19465 | | glycoside hydrolase family 127 protein | WP_001101509.1 | -0.586181654 | 0.02270582 | 0.04744768 |
| STM14_RS21815 | rplL | 50S ribosomal protein L7/L12 | WP_000028882.1 | -0.586867614 | 2.9466E-11 | 4.0483E-10 |
| STM14_RS13365 | ligA | NAD-dependent DNA ligase LigA | WP_000433266.1 | -0.588613297 | 9.9311E-06 | 5.2312E-05 |
| STM14_RS14965 | | tripartite tricarboxylate transporter permease | WP_000382035.1 | -0.589856159 | 0.00025587 | 0.00096035 |
| STM14_RS15960 | csdA | cysteine desulfurase CsdA | WP_000991053.1 | -0.590434717 | 0.00064455 | 0.00217926 |
| STM14_RS02220 | proA | glutamate-5-semialdehyde dehydrogenase | WP_000893231.1 | -0.590520783 | 1.6569E-05 | 8.4263E-05 |
| STM14_RS05425 | ldtD | L%2CD-transpeptidase | WP_000925872.1 | -0.590940425 | 2.9384E-09 | 2.948E-08 |
| STM14_RS16855 | | SDR family oxidoreductase | WP_000019032.1 | -0.592622045 | 1.0937E-08 | 1.0234E-07 |
| STM14_RS20330 | | NADPH-dependent FMN reductase | WP_001527038.1 | -0.596077518 | 1.2187E-05 | 6.2937E-05 |
| STM14_RS08005 | purR | HTH-type transcriptional repressor PurR | WP_000190993.1 | -0.596316244 | 1.0497E-05 | 5.4981E-05 |
| STM14_RS17900 | tldD | metalloprotease TldD | WP_000055936.1 | -0.596649765 | 6.4654E-07 | 4.368E-06 |
| STM14_RS12985 | hisJ | histidine ABC transporter substrate-binding protein HisJ | WP_000731868.1 | -0.597031115 | 1.7822E-09 | 1.8279E-08 |
| STM14_RS08770 | narW | nitrate reductase molybdenum cofactor assembly chaperone | WP_001166296.1 | -0.598225727 | 0.05917726 | 0.10808069 |
| STM14_RS09620 | oppC | oligopeptide ABC transporter permease OppC | WP_000979653.1 | -0.598276764 | 0.00025526 | 0.00096002 |
| STM14_RS22570 | melA | alpha-galactosidase | WP_001520868.1 | -0.598498512 | 5.4383E-05 | 0.00024629 |
| STM14_RS17740 | nanK | N-acetylmannosamine kinase | WP_000208976.1 | -0.599066744 | 4.493E-05 | 0.00020861 |
| STM14_RS09560 | ompW | outer membrane protein OmpW | WP_000714799.1 | -0.600556655 | 1.4683E-11 | 2.0954E-10 |
| STM14_RS04700 | | endonuclease/exonuclease/phosphatase family protein | WP_000192039.1 | -0.60129082 | 0.01971139 | 0.04215545 |
| STM14_RS04815 | | anion transporter | WP_000050088.1 | -0.601401241 | 0.03474384 | 0.06797386 |
| STM14_RS17925 | mreC | rod shape-determining protein MreC | WP_000802538.1 | -0.602831329 | 0.0024561 | 0.00711516 |
| STM14_RS12215 | | YeiH family protein | WP_000137961.1 | -0.604585157 | 0.00097095 | 0.00314151 |
| STM14_RS07345 | selD | selenide%2C water dikinase SelD | WP_001294900.1 | -0.604801602 | 3.2802E-06 | 1.9387E-05 |
| STM14_RS23350 | treB | PTS trehalose transporter subunit IIBC | | -0.605966769 | 0.00010339 | 0.00042967 |
| STM14_RS19475 | | ROK family protein | WP_001188274.1 | -0.606300523 | 5.4693E-05 | 0.0002474 |
| STM14_RS17220 | | YqjD family protein | WP_000031219.1 | -0.606696705 | 1.6595E-12 | 2.5891E-11 |
| STM14_RS17710 | gltB | glutamate synthase large subunit | WP_000965023.1 | -0.606818284 | 3.8407E-05 | 0.00018084 |
| STM14_RS13560 | eutM | ethanolamine utilization microcompartment protein EutM | WP_000387716.1 | -0.606848464 | 0.01019738 | 0.02426998 |
| STM14_RS06175 | ghrA | glyoxylate/hydroxypyruvate reductase GhrA | WP_000402556.1 | -0.606950992 | 2.0946E-05 | 0.00010479 |
| STM14_RS01135 | leuC | 3-isopropylmalate dehydratase large subunit | WP_001140629.1 | -0.607098114 | 0.01340131 | 0.03055332 |
| STM14_RS20585 | ilvE | branched-chain-amino-acid transaminase | WP_000208528.1 | -0.607136253 | 1.561E-05 | 7.9815E-05 |
| STM14_RS04645 | moaC | cyclic pyranopterin monophosphate synthase MoaC | WP_000080894.1 | -0.608216608 | 0.00423118 | 0.01133034 |
| STM14_RS18750 | glgB | 1%2C4-alpha-glucan branching enzyme | WP_000098543.1 | -0.608732907 | 6.3142E-11 | 8.1863E-10 |
| STM14_RS18415 | | putative adenosine monophosphate-protein transferase Fic | WP_001280671.1 | -0.609510633 | 8.9036E-07 | 5.8126E-06 |
| STM14_RS03590 | | isochorismatase | WP_001007166.1 | -0.609629541 | 0.02319263 | 0.0483005 |
| STM14_RS14945 | tctD | transcriptional regulator TctD | WP_001237934.1 | -0.610227839 | 5.8856E-06 | 3.2934E-05 |
| STM14_RS00890 | dapB | 4-hydroxy-tetrahydrodipicolinate reductase | WP_000544031.1 | -0.610439291 | 4.546E-07 | 3.1463E-06 |
| STM14_RS19940 | | glycoside hydrolase family 1 protein | WP_001270489.1 | -0.610581256 | 1.1666E-07 | 9.2174E-07 |
| STM14_RS06655 | | cupin domain-containing protein | WP_000456516.1 | -0.611606138 | 8.987E-07 | 5.8463E-06 |
| STM14_RS12085 | maiA | maleylacetoacetate isomerase | WP_000781183.1 | -0.61166613 | 0.01441566 | 0.03242422 |
| STM14_RS19695 | coaD | pantetheine-phosphate adenylyltransferase | WP_001171890.1 | -0.613061666 | 0.0029811 | 0.00839175 |
| STM14_RS17930 | mreB | rod shape-determining protein MreB | WP_000913396.1 | -0.61329087 | 6.0909E-08 | 5.0171E-07 |
| STM14_RS19785 | trmH | tRNA (guanosine(18)-2'-O)-methyltransferase TrmH | WP_001068438.1 | -0.613911313 | 0.00286156 | 0.00810481 |
| STM14_RS12605 | yfaE | ferredoxin-like diferric-tyrosyl radical cofactor maintenance protein Yfal | WP_000533921.1 | -0.613956033 | 0.08029014 | 0.14024112 |
| STM14_RS18845 | | type II toxin-antitoxin system death-on-curing family toxin | WP_001521773.1 | -0.614740285 | 0.13519064 | 0.21585947 |
| STM14_RS17690 | elbB | isoprenoid biosynthesis glyoxalase ElbB | WP_000719822.1 | -0.616594949 | 1.291E-07 | 1.0049E-06 |
| STM14_RS03810 | rlmH | 23S rRNA (pseudouridine(1915)-N(3))-methyltransferase RlmH | WP_000776107.1 | -0.616601386 | 0.00740596 | 0.01848728 |
| STM14_RS07715 | sufD | Fe-S cluster assembly protein SufD | WP_000907925.1 | -0.616822868 | 1.391E-06 | 8.6956E-06 |
| STM14_RS21275 | | XRE family transcriptional regulator | WP_001517951.1 | -0.616971546 | 0.0019566 | 0.00582864 |
| STM14_RS11300 | | cobyric acid synthase | WP_000189662.1 | -0.617350723 | 5.0132E-05 | 0.00022987 |
| STM14_RS10675 | dcyD | D-cysteine desulfhydrase | WP_001128180.1 | -0.618137778 | 8.6705E-06 | 4.6402E-05 |
| STM14_RS19460 | | AraC family transcriptional regulator | WP_000891805.1 | -0.618211168 | 0.00894944 | 0.02167885 |
| STM14_RS21895 | nfi | deoxyribonuclease V | WP_000362359.1 | -0.618460866 | 0.00015096 | 0.00060222 |
| STM14_RS13440 | cysA | sulfate/thiosulfate ABC transporter ATP-binding protein CysA | WP_000021076.1 | -0.619634989 | 0.02385654 | 0.04939068 |
| STM14_RS12760 | menF | isochorismate synthase MenF | WP_000555674.1 | -0.619774326 | 6.4219E-05 | 0.00028386 |
| STM14_RS00765 | nhaA | Na+/H+ antiporter NhaA | WP_000681340.1 | -0.620239086 | 5.0937E-07 | 3.4731E-06 |
| STM14_RS20805 | xerC | tyrosine recombinase XerC | WP_000132874.1 | -0.620297739 | 0.00090792 | 0.00295836 |
| STM14_RS16370 | rpiA | ribose-5-phosphate isomerase RpiA | WP_000189741.1 | -0.620713645 | 1.4919E-10 | 1.8372E-09 |
| STM14_RS03925 | gltK | glutamate/aspartate ABC transporter permease GltK | WP_000272794.1 | -0.621937081 | 0.00055098 | 0.00190712 |
| STM14_RS11355 | | decarboxylating cobalt-precorrin-6B (C(15))-methyltransferase | WP_000650985.1 | -0.622140439 | 0.01219964 | 0.02823322 |
| STM14_RS01245 | ftsZ | cell division protein FtsZ | WP_000462785.1 | -0.622650693 | 5.4337E-12 | 8.1E-11 |
| STM14_RS21570 | metL | bifunctional aspartate kinase/homoserine dehydrogenase II | WP_000110806.1 | -0.622747224 | 0.00019898 | 0.00076959 |
| STM14_RS15860 | | flavodoxin | WP_000807739.1 | -0.62315461 | 1.1933E-06 | 7.5756E-06 |
| STM14_RS11935 | thiD | bifunctional hydroxymethylpyrimidine kinase/phosphomethylpyrimidin | WP_001188957.1 | -0.625038826 | 0.05976976 | 0.10905174 |

| | | | | | | |
|---|---|---|---|---|---|---|
| STM14_RS05200 | | hydrolase | WP_000097893.1 | -0.625964855 | 0.000122 | 0.00049969 |
| STM14_RS03615 | | oxidoreductase | WP_001016847.1 | -0.626054277 | 0.0037375 | 0.01017859 |
| STM14_RS12830 | nuoF | NADH-quinone oxidoreductase subunit NuoF | WP_000800048.1 | -0.626354106 | 2.9505E-08 | 2.5562E-07 |
| STM14_RS21785 | tuf | elongation factor Tu | WP_000031748.1 | -0.626794932 | 5.5502E-13 | 9.4174E-12 |
| STM14_RS13920 | hscA | Fe-S protein assembly chaperone HscA | WP_001196669.1 | -0.626911855 | 0.00040656 | 0.00145475 |
| STM14_RS19035 | | 23S rRNA (adenine(2030)-N(6))-methyltransferase RlmJ | WP_000954235.1 | -0.627629617 | 4.4564E-06 | 2.5638E-05 |
| STM14_RS01085 | araA | L-arabinose isomerase | WP_000151691.1 | -0.63035796 | 0.04888218 | 0.09136252 |
| STM14_RS04100 | pgm | phosphoglucomutase (alpha-D-glucose-1%2C6-bisphosphate-depend | WP_000974387.1 | -0.630617548 | 1.2877E-10 | 1.6142E-09 |
| STM14_RS01530 | hrpB | ATP-dependent helicase HrpB | WP_001738008.1 | -0.632888808 | 5.8148E-05 | 0.00026142 |
| STM14_RS15630 | | NAD(P)-dependent oxidoreductase | WP_000206958.1 | -0.633111559 | 6.8554E-05 | 0.00030014 |
| STM14_RS23100 | cysQ | 3'(2')%2C5'-bisphosphate nucleotidase CysQ | WP_000893398.1 | -0.633645131 | 7.1321E-10 | 7.701E-09 |
| STM14_RS22925 | rlmB | 23S rRNA (guanosine(2251)-2'-O)-methyltransferase RlmB | WP_001293274.1 | -0.633848926 | 1.2774E-06 | 8.0398E-06 |
| STM14_RS03735 | dpiA | two-component response regulator DpiA | WP_000138788.1 | -0.63404149 | 0.02465251 | 0.0508378 |
| STM14_RS12765 | elaB | stress response protein ElaB | WP_001522308.1 | -0.634339041 | 7.2868E-13 | 1.1978E-11 |
| STM14_RS15290 | | respiratory chain complex I subunit 1 family protein | WP_000110572.1 | -0.6343465 | 0.02863885 | 0.05787499 |
| STM14_RS17590 | murA | UDP-N-acetylglucosamine 1-carboxyvinyltransferase | WP_000357288.1 | -0.634913672 | 3.7134E-07 | 2.6293E-06 |
| STM14_RS14740 | | AAA family ATPase | WP_001520831.1 | -0.635984176 | 0.00113843 | 0.00359906 |
| STM14_RS01745 | accA | acetyl-CoA carboxylase carboxyl transferase subunit alpha | WP_000055753.1 | -0.637372702 | 1.7141E-08 | 1.5507E-07 |
| STM14_RS18685 | glpE | thiosulfate sulfurtransferase GlpE | WP_000434523.1 | -0.639224956 | 0.00219116 | 0.00641307 |
| STM14_RS04035 | nagE | PTS N-acetyl glucosamine transporter subunit IIABC | WP_001023068.1 | -0.639355309 | 6.7872E-09 | 6.5765E-08 |
| STM14_RS17110 | plsY | glycerol-3-phosphate 1-O-acyltransferase PlsY | WP_001272784.1 | -0.639468243 | 0.00020679 | 0.00079726 |
| STM14_RS19285 | | N-acetyltransferase | WP_000619902.1 | -0.639738374 | 0.00282713 | 0.00802582 |
| STM14_RS05015 | rimK | 30S ribosomal protein S6--L-glutamate ligase | WP_000684361.1 | -0.641712416 | 0.00297084 | 0.00837568 |
| STM14_RS17715 | gltD | glutamate synthase subunit GltD | WP_000081628.1 | -0.642105868 | 0.0003977 | 0.00142583 |
| STM14_RS11370 | | cobalt-precorrin-8 methylmutase | WP_000558988.1 | -0.642330881 | 0.02284909 | 0.04769293 |
| STM14_RS08255 | pntB | Re/Si-specific NAD(P)(+) transhydrogenase subunit beta | WP_000014056.1 | -0.642709908 | 9.5456E-09 | 9.0585E-08 |
| STM14_RS01400 | | D-threonate 4-phosphate dehydrogenase | WP_000448745.1 | -0.643645715 | 2.55E-05 | 0.00012485 |
| STM14_RS08830 | | DMT family transporter | WP_001076571.1 | -0.64417899 | 0.01119662 | 0.02625857 |
| STM14_RS15685 | ispF | 2-C-methyl-D-erythritol 2%2C4-cyclodiphosphate synthase | WP_001219245.1 | -0.645119037 | 0.00075676 | 0.0025239 |
| STM14_RS12955 | yfcF | glutathione transferase | WP_000043010.1 | -0.645571528 | 8.3976E-06 | 4.5205E-05 |
| STM14_RS15905 | fucO | lactaldehyde reductase | WP_000013565.1 | -0.64702732 | 0.00405616 | 0.0109734 |
| STM14_RS13855 | ndk | nucleoside-diphosphate kinase | WP_000963846.1 | -0.648600809 | 1.4317E-10 | 1.769E-09 |
| STM14_RS12995 | | UbiX family flavin prenyltransferase | WP_000825682.1 | -0.649252819 | 0.01390286 | 0.03148237 |
| STM14_RS03770 | lipA | lipoyl synthase | WP_000042640.1 | -0.650804241 | 6.5716E-08 | 5.401E-07 |
| STM14_RS12540 | ada | bifunctional DNA-binding transcriptional regulator/O6-methylguanine-D | WP_000975956.1 | -0.652385123 | 4.1365E-06 | 2.3948E-05 |
| STM14_RS12535 | alkB | DNA oxidative demethylase AlkB | WP_000884990.1 | -0.652540397 | 0.00284929 | 0.0080825 |
| STM14_RS13650 | dapE | succinyl-diaminopimelate desuccinylase | WP_001277825.1 | -0.655394113 | 4.4552E-06 | 2.5638E-05 |
| STM14_RS03370 | fimW | fimbria biosynthesis transcriptional regulator FimW | WP_000944156.1 | -0.656354504 | 2.3084E-05 | 0.00011424 |
| STM14_RS19615 | kbl | glycine C-acetyltransferase | WP_001213790.1 | -0.656386662 | 1.5391E-10 | 1.8827E-09 |
| STM14_RS16540 | | YqgE/AlgH family protein | WP_001053171.1 | -0.656442202 | 0.0001386 | 0.00056079 |
| STM14_RS03115 | | NfeD family protein | WP_000561177.1 | -0.656489687 | 0.00067394 | 0.00227238 |
| STM14_RS04655 | moaE | molybdopterin synthase catalytic subunit MoaE | WP_000545202.1 | -0.657824284 | 0.00428035 | 0.01143947 |
| STM14_RS20275 | rnpA | ribonuclease P protein component | WP_000239725.1 | -0.658956196 | 8.5329E-05 | 0.00036406 |
| STM14_RS13780 | guaB | IMP dehydrogenase | WP_000944174.1 | -0.65901163 | 2.4263E-05 | 0.00011943 |
| STM14_RS20140 | | heme lyase CcmF/NrfE family subunit | WP_000982518.1 | -0.660771939 | 0.04422712 | 0.08398363 |
| STM14_RS10125 | proQ | RNA chaperone ProQ | WP_000431401.1 | -0.660789379 | 5.5221E-10 | 6.1058E-09 |
| STM14_RS04160 | phrB | deoxyribodipyrimidine photo-lyase | WP_001142016.1 | -0.66192539 | 0.00021727 | 0.00083421 |
| STM14_RS19625 | rfaF | ADP-heptose--LPS heptosyltransferase RfaF | WP_000699196.1 | -0.662467942 | 2.6274E-07 | 1.9119E-06 |
| STM14_RS12045 | dld | D-lactate dehydrogenase | WP_000095232.1 | -0.662827754 | 5.0556E-10 | 5.6408E-09 |
| STM14_RS17750 | nanT | sialic acid transporter NanT | WP_000108071.1 | -0.663146474 | 2.583E-07 | 1.8833E-06 |
| STM14_RS13730 | purM | phosphoribosylformylglycinamidine cyclo-ligase | WP_000130477.1 | -0.664502235 | 7.8995E-07 | 5.222E-06 |
| STM14_RS19815 | yicI | alpha-xylosidase | WP_000702987.1 | -0.664530053 | 0.01649803 | 0.03630948 |
| STM14_RS20280 | yidD | membrane protein insertion efficiency factor YidD | WP_001307474.1 | -0.665194855 | 0.00925829 | 0.02232419 |
| STM14_RS00810 | fkpB | FKBP-type peptidyl-prolyl cis-trans isomerase | WP_000047269.1 | -0.666001556 | 1.1459E-05 | 5.9507E-05 |
| STM14_RS01425 | cueO | multicopper oxidase CueO | WP_000946047.1 | -0.66626433 | 1.2369E-06 | 7.8251E-06 |
| STM14_RS17470 | rbfA | 30S ribosome-binding factor RbfA | WP_001040199.1 | -0.666287212 | 2.1599E-05 | 0.00010776 |
| STM14_RS17405 | | N-acetyltransferase | WP_000908562.1 | -0.667225156 | 9.0264E-10 | 9.523E-09 |
| STM14_RS01195 | murE | UDP-N-acetylmuramoyl-L-alanyl-D-glutamate--2%2C6-diaminopimela | WP_000775077.1 | -0.667471576 | 3.9984E-08 | 3.4E-07 |
| STM14_RS04735 | rhlE | ATP-dependent RNA helicase RhlE | WP_000007060.1 | -0.668359068 | 0.00453075 | 0.01197574 |
| STM14_RS13605 | tkt | transketolase | WP_000087323.1 | -0.668862397 | 8.9814E-11 | 1.1482E-09 |
| STM14_RS17260 | | hypothetical protein | WP_000039324.1 | -0.669243398 | 8.066E-05 | 0.00034557 |
| STM14_RS19155 | bcsQ | cellulose biosynthesis protein BcsQ | WP_000996125.1 | -0.669384022 | 5.3486E-05 | 0.00024313 |
| STM14_RS18740 | glgC | glucose-1-phosphate adenylyltransferase | WP_000253995.1 | -0.670782496 | 6.9458E-13 | 1.1468E-11 |
| STM14_RS19140 | bcsZ | cellulose synthase complex periplasmic endoglucanase BcsZ | WP_000988813.1 | -0.671475523 | 6.4672E-05 | 0.00028552 |
| STM14_RS04490 | galM | galactose-1-epimerase | WP_000931428.1 | -0.671749028 | 2.7954E-08 | 2.439E-07 |
| STM14_RS02375 | golT | gold/copper-translocating P-type ATPase GolT | WP_000083318.1 | -0.672129821 | 0.03200223 | 0.06335066 |
| STM14_RS09985 | fadD | long-chain-fatty-acid--CoA ligase FadD | WP_000758418.1 | -0.672579152 | 3.4921E-08 | 3.0112E-07 |
| STM14_RS13510 | eutK | ethanolamine utilization microcompartment protein EutK | WP_000606266.1 | -0.672709488 | 0.00296541 | 0.00837319 |
| STM14_RS21685 | oxyR | DNA-binding transcriptional regulator OxyR | WP_001025919.1 | -0.672747697 | 7.1814E-12 | 1.0619E-10 |
| STM14_RS12825 | nuoG | NADH-quinone oxidoreductase subunit NuoG | WP_001518099.1 | -0.672878349 | 1.6613E-13 | 3.0281E-12 |
| STM14_RS17730 | | cytosine deaminase | WP_001180690.1 | -0.675399585 | 7.5751E-06 | 4.126E-05 |
| STM14_RS04595 | bioB | biotin synthase BioB | WP_000090729.1 | -0.675865414 | 0.00368434 | 0.01007108 |
| STM14_RS17075 | hldE | bifunctional D-glycero-beta-D-manno-heptose-7-phosphate kinase/D-g | WP_000867682.1 | -0.67910968 | 5.2854E-08 | 4.4029E-07 |
| STM14_RS09010 | | S-(hydroxymethyl)glutathione dehydrogenase/class III alcohol dehydrog | WP_000842141.1 | -0.680062419 | 1.438E-07 | 1.1077E-06 |
| STM14_RS02820 | | cytochrome o ubiquinol oxidase subunit IV | WP_000019878.1 | -0.682207717 | 6.4751E-11 | 8.3654E-10 |
| STM14_RS16145 | | acetyl-CoA C-acetyltransferase | WP_000665977.1 | -0.682676544 | 1.2351E-08 | 1.1455E-07 |
| STM14_RS07570 | btuD | vitamin B12 ABC transporter ATP-binding protein BtuD | WP_000080607.1 | -0.683228117 | 0.02950554 | 0.0594259 |
| STM14_RS08955 | sgcA | SgcA family PTS sugar transporter subunit IIA | WP_000606458.1 | -0.684916878 | 0.01169636 | 0.0272225 |
| STM14_RS15345 | sitA | iron/manganese ABC transporter substrate-binding protein SitA | WP_000182930.1 | -0.686376392 | 2.6546E-08 | 2.3272E-07 |
| STM14_RS15785 | queD | 6-carboxytetrahydropterin synthase QueD | WP_000108313.1 | -0.689902586 | 0.02467277 | 0.0508378 |
| STM14_RS14495 | | tRNA/rRNA methyltransferase | WP_000997368.1 | -0.691427267 | 1.5194E-06 | 9.4025E-06 |
| STM14_RS04310 | sucD | succinate--CoA ligase subunit alpha | WP_000114631.1 | -0.691670244 | 1.7653E-14 | 3.7355E-13 |

| | | | | | | |
|---|---|---|---|---|---|---|
| STM14_RS00135 | | hypothetical protein | WP_001266176.1 | -0.692331447 | 0.00032677 | 0.00119873 |
| STM14_RS01095 | araC | arabinose operon transcriptional regulator AraC | WP_000852387.1 | -0.692404084 | 6.1894E-06 | 3.4335E-05 |
| STM14_RS18905 | ftsE | cell division ATP-binding protein FtsE | WP_000617729.1 | -0.692637536 | 1.0019E-06 | 6.4588E-06 |
| STM14_RS04775 | glnQ | glutamine ABC transporter ATP-binding protein GlnQ | WP_000569093.1 | -0.693713087 | 0.0003483 | 0.00126474 |
| STM14_RS13945 | iscR | Fe-S cluster assembly transcriptional regulator IscR | WP_001241346.1 | -0.695119312 | 2.9211E-08 | 2.5367E-07 |
| STM14_RS12735 | menE | o-succinylbenzoate--CoA ligase | WP_000144677.1 | -0.695852812 | 2.6705E-05 | 0.00012989 |
| STM14_RS18840 | ugpB | sn-glycerol-3-phosphate ABC transporter substrate-binding protein Ugp | WP_000624747.1 | -0.696984916 | 9.0262E-12 | 1.3084E-10 |
| STM14_RS04585 | | kinase inhibitor | WP_000767419.1 | -0.699986067 | 0.00080325 | 0.00266207 |
| STM14_RS18680 | glpG | rhomboid family intramembrane serine protease GlpG | WP_000928699.1 | -0.700255878 | 0.00014905 | 0.00059637 |
| STM14_RS03275 | purK | 5-(carboxyamino)imidazole ribonucleotide synthase | WP_000815603.1 | -0.702483957 | 9.502E-06 | 5.0413E-05 |
| STM14_RS14685 | bapA | biofilm-associated protein BapA | WP_001237670.1 | -0.703637389 | 3.738E-06 | 2.1777E-05 |
| STM14_RS03280 | purE | 5-(carboxyamino)imidazole ribonucleotide mutase | WP_000098736.1 | -0.704218561 | 0.00265717 | 0.00760784 |
| STM14_RS02815 | cyoE | heme o synthase | WP_000971352.1 | -0.704990442 | 1.8329E-14 | 3.8564E-13 |
| STM14_RS07575 | | NlpC/P60 family protein | WP_001213256.1 | -0.705334921 | 0.00020126 | 0.00077676 |
| STM14_RS00260 | | DUF2913 family protein | | -0.707516611 | 0.0001611 | 0.00063781 |
| STM14_RS01030 | rsmA | 16S rRNA (adenine(1518)-N(6)/adenine(1519)-N(6))-dimethyltransferas | WP_001065397.1 | -0.708046556 | 1.665E-07 | 1.2614E-06 |
| STM14_RS12915 | | transketolase | WP_001103264.1 | -0.70924267 | 6.0718E-06 | 3.3873E-05 |
| STM14_RS15200 | | PTS glucitol/sorbitol transporter subunit IIB | WP_000199033.1 | -0.709580721 | 0.02191164 | 0.04607303 |
| STM14_RS01020 | apaH | bis(5'-nucleosyl)-tetraphosphatase (symmetrical) ApaH | WP_000257211.1 | -0.709650908 | 1.0199E-08 | 9.6288E-08 |
| STM14_RS02640 | queA | tRNA preQ1(34) S-adenosylmethionine ribosyltransferase-isomerase Q | WP_001266530.1 | -0.709973574 | 0.00858307 | 0.02098464 |
| STM14_RS17735 | nanQ | N-acetylneuraminate anomerase | WP_000979905.1 | -0.711581962 | 3.0104E-10 | 3.4747E-09 |
| STM14_RS20210 | dgoD | galactonate dehydratase | WP_000704735.1 | -0.711846208 | 0.01369064 | 0.03107826 |
| STM14_RS01870 | | MFS transporter | WP_001127538.1 | -0.714933407 | 0.03077646 | 0.06145277 |
| STM14_RS07720 | sufS | cysteine desulfurase SufS | WP_000143859.1 | -0.715440718 | 5.7222E-08 | 4.7346E-07 |
| STM14_RS16385 | | oxidative stress defense protein | WP_000669854.1 | -0.71572806 | 1.4182E-08 | 1.3055E-07 |
| STM14_RS08480 | sad | succinate-semialdehyde dehydrogenase | WP_000178397.1 | -0.718122975 | 9.0694E-08 | 7.2912E-07 |
| STM14_RS08950 | sgcQ | BtpA family protein SgcQ | WP_000118615.1 | -0.71910308 | 0.00192742 | 0.00576037 |
| STM14_RS21290 | rhaD | rhamnulose-1-phosphate aldolase | WP_001179685.1 | -0.725285815 | 0.0005739 | 0.001973 |
| STM14_RS17225 | | phage holin family protein | WP_000841082.1 | -0.726566291 | 1.3987E-12 | 2.1916E-11 |
| STM14_RS23160 | mpl | UDP-N-acetylmuramate:L-alanyl-gamma-D-glutamyl-meso-diaminopin | WP_001219833.1 | -0.726621044 | 7.5585E-08 | 6.1572E-07 |
| STM14_RS15865 | truC | tRNA pseudouridine(65) synthase TruC | WP_000890027.1 | -0.72800919 | 0.01141782 | 0.02672627 |
| STM14_RS19295 | | OmpA family lipoprotein | WP_000747548.1 | -0.728098078 | 0.00076904 | 0.00256253 |
| STM14_RS18735 | glgA | glycogen synthase GlgA | WP_001197669.1 | -0.72872343 | 2.5534E-12 | 3.869E-11 |
| STM14_RS20800 | | DUF484 domain-containing protein | WP_000812765.1 | -0.728818307 | 3.4888E-06 | 2.0488E-05 |
| STM14_RS02430 | | helix-turn-helix domain-containing protein | WP_000074605.1 | -0.728842855 | 0.00134982 | 0.00419411 |
| STM14_RS20670 | wecC | UDP-N-acetyl-D-mannosamine dehydrogenase | WP_000011227.1 | -0.732255085 | 1.9115E-08 | 1.7166E-07 |
| STM14_RS04560 | hutI | imidazolonepropionase | WP_001249472.1 | -0.733609584 | 2.9587E-10 | 3.4258E-09 |
| STM14_RS19135 | bcsC | cellulose synthase complex outer membrane protein BcsC | WP_001225195.1 | -0.735104445 | 5.5171E-10 | 6.1058E-09 |
| STM14_RS03825 | | adenosylcobalamin/alpha-ribazole phosphatase | WP_001241924.1 | -0.736009518 | 0.00092763 | 0.00301725 |
| STM14_RS04185 | pxpA | 5-oxoprolinase subunit PxpA | WP_001017909.1 | -0.73673557 | 6.5075E-05 | 0.00028696 |
| STM14_RS19630 | rfaC | lipopolysaccharide heptosyltransferase RfaC | WP_001264604.1 | -0.737925991 | 6.5766E-07 | 4.4269E-06 |
| STM14_RS11305 | | energy-coupling factor ABC transporter ATP-binding protein | WP_000881552.1 | -0.739267177 | 7.2961E-05 | 0.00031642 |
| STM14_RS15625 | | 3-oxo-tetronate kinase | WP_000912490.1 | -0.739280659 | 7.8997E-07 | 5.222E-06 |
| STM14_RS13525 | eutB | ethanolamine ammonia-lyase subunit alpha | WP_000769990.1 | -0.741514512 | 1.2289E-05 | 6.3371E-05 |
| STM14_RS14970 | glaH | glutarate dioxygenase GlaH | WP_000993096.1 | -0.742956563 | 1.2811E-11 | 1.8354E-10 |
| STM14_RS13255 | mgrA | L-glyceraldehyde 3-phosphate reductase | WP_000640293.1 | -0.743097081 | 2.6437E-07 | 1.92E-06 |
| STM14_RS00130 | ccdB | type II toxin-antitoxin system toxin CcdB | WP_001159863.1 | -0.744459373 | 5.6642E-05 | 0.00025558 |
| STM14_RS19020 | rsmJ | 16S rRNA (guanine(1516)-N(2))-methyltransferase RsmJ | WP_001165134.1 | -0.745044925 | 0.01555544 | 0.03459028 |
| STM14_RS07725 | sufE | cysteine desulfuration protein SufE | WP_000729468.1 | -0.745518318 | 5.4785E-05 | 0.00024751 |
| STM14_RS23505 | | NAD(P)-dependent alcohol dehydrogenase | WP_000152563.1 | -0.746223201 | 1.0197E-07 | 8.0744E-07 |
| STM14_RS19505 | mtlR | mannitol operon repressor MtlR | WP_000193201.1 | -0.746409795 | 0.01037816 | 0.02462137 |
| STM14_RS07160 | | four-helix bundle copper-binding protein | WP_001183699.1 | -0.747135911 | 1.1525E-06 | 7.3455E-06 |
| STM14_RS07490 | pfkB | 6-phosphofructokinase II | WP_000251707.1 | -0.747959598 | 4.4885E-11 | 6.0316E-10 |
| STM14_RS15295 | hycC | formate hydrogenlyase subunit 3 | WP_000097040.1 | -0.751374661 | 0.00119614 | 0.00375783 |
| STM14_RS21665 | argE | acetylornithine deacetylase | WP_000800210.1 | -0.751399126 | 1.1141E-09 | 1.172E-08 |
| STM14_RS18075 | tsaC | L-threonylcarbamoyladenylate synthase type 1 TsaC | WP_001063609.1 | -0.751506558 | 8.368E-06 | 4.5111E-05 |
| STM14_RS06095 | rutR | HTH-type transcriptional regulator RutR | WP_000083126.1 | -0.752042325 | 0.01290537 | 0.02958753 |
| STM14_RS01010 | kefC | glutathione-regulated potassium-efflux system protein KefC | WP_000377155.1 | -0.755122427 | 8.4978E-06 | 4.5677E-05 |
| STM14_RS00585 | thrC | threonine synthase | WP_000781105.1 | -0.755708515 | 4.1877E-12 | 6.2679E-11 |
| STM14_RS02825 | | cytochrome o ubiquinol oxidase subunit III | WP_000179833.1 | -0.75616563 | 5.9409E-13 | 9.943E-12 |
| STM14_RS17390 | yhbO | protein/nucleic acid deglycase | WP_000037599.1 | -0.756411717 | 1.3893E-10 | 1.7224E-09 |
| STM14_RS03595 | entA | 2%2C3-dihydro-2%2C3-dihydroxybenzoate dehydrogenase EntA | WP_000145522.1 | -0.759261627 | 0.02323646 | 0.04836441 |
| STM14_RS07545 | pheS | phenylalanine--tRNA ligase subunit alpha | WP_000018570.1 | -0.760086738 | 2.1516E-10 | 2.5556E-09 |
| STM14_RS13870 | dmsB | dimethylsulfoxide reductase subunit B | WP_000077436.1 | -0.766179562 | 0.02344072 | 0.04870696 |
| STM14_RS02420 | cydB | cytochrome d ubiquinol oxidase subunit II | WP_000151043.1 | -0.769951201 | 1.1632E-05 | 6.024E-05 |
| STM14_RS13890 | sseA | 3-mercaptopyruvate sulfurtransferase | WP_000205241.1 | -0.770319213 | 3.5265E-12 | 5.2999E-11 |
| STM14_RS11335 | | cobalt-precorrin-6A reductase | WP_001001975.1 | -0.771203183 | 0.00217863 | 0.00638671 |
| STM14_RS20690 | rffA | dTDP-4-amino-4%2C6-dideoxygalactose transaminase | WP_000612078.1 | -0.772114504 | 7.6792E-07 | 5.113E-06 |
| STM14_RS17310 | garR | 2-hydroxy-3-oxopropionate reductase | WP_000178106.1 | -0.772672866 | 7.8729E-07 | 5.222E-06 |
| STM14_RS01220 | murG | undecaprenyldiphospho-muramoylpentapeptide beta-N-acetylglucosa | WP_000016613.1 | -0.773375221 | 5.8591E-09 | 5.7224E-08 |
| STM14_RS04785 | glnH | glutamine ABC transporter substrate-binding protein GlnH | WP_000838671.1 | -0.773621277 | 5.8746E-17 | 1.5021E-15 |
| STM14_RS23070 | cycA | D-serine/D-alanine/glycine transporter | WP_000228322.1 | -0.774465685 | 4.796E-11 | 6.3981E-10 |
| STM14_RS10995 | | hypothetical protein | WP_000601353.1 | -0.775924538 | 0.01780951 | 0.03866427 |
| STM14_RS23455 | holC | DNA polymerase III subunit chi | WP_001523689.1 | -0.776621103 | 3.5066E-05 | 0.00016681 |
| STM14_RS21170 | | sugar kinase | WP_001520529.1 | -0.77786984 | 0.00277788 | 0.00789819 |
| STM14_RS07785 | ttrS | tetrathionate respiration histidine kinase TtrS | WP_001214413.1 | -0.779075236 | 3.3847E-12 | 5.1075E-11 |
| STM14_RS00485 | traN | type-F conjugative transfer system mating-pair stabilization protein TraN | WP_000827528.1 | -0.779076847 | 3.3767E-05 | 0.00016084 |
| STM14_RS13550 | | aldehyde dehydrogenase EutE | WP_001075640.1 | -0.780832532 | 0.0035922 | 0.00984847 |
| STM14_RS14940 | | sensor histidine kinase | WP_000872348.1 | -0.780833312 | 5.5372E-06 | 3.1127E-05 |
| STM14_RS23450 | | valine--tRNA ligase | WP_000416329.1 | -0.781966117 | 2.3362E-14 | 4.7788E-13 |
| STM14_RS20895 | | dienelactone hydrolase family protein | WP_000213959.1 | -0.783904834 | 1.1839E-10 | 1.498E-09 |
| STM14_RS19180 | | type I toxin-antitoxin system toxin Ldr family protein | WP_001669334.1 | -0.784518242 | 4.3928E-07 | 3.0517E-06 |

| | | | | | | |
|---|---|---|---|---|---|---|
| STM14_RS21795 | nusG | transcription termination/antitermination protein NusG | WP_001287521.1 | -0.785817812 | 6.0346E-10 | 6.6129E-09 |
| STM14_RS19540 | lldD | quinone-dependent L-lactate dehydrogenase | WP_000586988.1 | -0.787968326 | 1.0179E-05 | 5.3391E-05 |
| STM14_RS16570 | yggU | DUF167 family protein YggU | WP_001277201.1 | -0.787999931 | 0.00210014 | 0.00619657 |
| STM14_RS10835 | | mannosyl-3-phosphoglycerate phosphatase-related protein | WP_000948794.1 | -0.789726228 | 5.2177E-05 | 0.00023806 |
| STM14_RS13435 | cysM | cysteine synthase CysM | WP_001093886.1 | -0.791755248 | 8.9241E-07 | 5.8157E-06 |
| STM14_RS23835 | | YbaK/EbsC family protein | WP_000058276.1 | -0.794177128 | 2.5326E-08 | 2.2309E-07 |
| STM14_RS04950 | | glutathione S-transferase family protein | WP_000631907.1 | -0.796404443 | 1.0425E-08 | 9.7919E-08 |
| STM14_RS11365 | cbiD | cobalt-precorrin-5B (C(1))-methyltransferase CbiD | WP_001292908.1 | -0.796958409 | 1.1127E-05 | 5.795E-05 |
| STM14_RS13630 | aegA | formate-dependent uric acid utilization protein AegA | WP_001078938.1 | -0.797740274 | 5.0823E-05 | 0.00023246 |
| STM14_RS21935 | purH | bifunctional phosphoribosylaminoimidazolecarboxamide formyltransfe | WP_001187509.1 | -0.800560244 | 7.1989E-05 | 0.00031294 |
| STM14_RS15250 | hypF | carbamoyltransferase HypF | WP_000963438.1 | -0.801773198 | 4.6457E-05 | 0.00021516 |
| STM14_RS02415 | | cytochrome ubiquinol oxidase subunit I | WP_000393711.1 | -0.802627164 | 9.0432E-09 | 8.6262E-08 |
| STM14_RS12925 | | PTS sugar transporter subunit IIB | WP_000699818.1 | -0.803102477 | 6.5338E-05 | 0.00028777 |
| STM14_RS00825 | rihC | ribonucleoside hydrolase RihC | WP_000127283.1 | -0.803622259 | 0.00078252 | 0.00259806 |
| STM14_RS20675 | rffG | dTDP-glucose 4%2C6-dehydratase | WP_000822145.1 | -0.806034016 | 1.5585E-08 | 1.4204E-07 |
| STM14_RS13160 | fadI | acetyl-CoA C-acyltransferase FadI | WP_001248125.1 | -0.806141369 | 7.4339E-06 | 4.0671E-05 |
| STM14_RS11490 | pduW | propanediol utilization propionate kinase PduW | WP_000120647.1 | -0.80985368 | 0.00731781 | 0.01830447 |
| STM14_RS23910 | | metal-dependent hydrolase | WP_000119016.1 | -0.81010142 | 0.01006054 | 0.02397542 |
| STM14_RS22865 | tsaE | tRNA (adenosine(37)-N6)-threonylcarbamoyltransferase complex ATPa | WP_000981983.1 | -0.814899678 | 4.06E-07 | 2.8474E-06 |
| STM14_RS16320 | gcvP | aminomethyl-transferring glycine dehydrogenase | WP_000194975.1 | -0.815639454 | 3.0145E-19 | 9.3273E-18 |
| STM14_RS04745 | ybiB | DNA-binding protein YbiB | WP_000386494.1 | -0.81660034 | 7.0747E-08 | 5.7887E-07 |
| STM14_RS23460 | pepA | leucyl aminopeptidase | WP_000397158.1 | -0.818390716 | 4.9286E-14 | 9.6526E-13 |
| STM14_RS19105 | | AsmA family protein | WP_000158675.1 | -0.822421934 | 8.3161E-14 | 1.5543E-12 |
| STM14_RS19900 | | 2-dehydro-3-deoxy-phosphogluconate aldolase | WP_001186146.1 | -0.823091688 | 0.01618068 | 0.03573921 |
| STM14_RS17170 | | NADPH-dependent 2%2C4-dienoyl-CoA reductase | WP_000121523.1 | -0.823234451 | 8.4064E-07 | 5.5272E-06 |
| STM14_RS11585 | hisA | 1-(5-phosphoribosyl)-5-[(5-phosphoribosylamino)methylideneamino]in | WP_000586409.1 | -0.824850101 | 0.00138842 | 0.00429595 |
| STM14_RS13675 | bamC | outer membrane protein assembly factor BamC | WP_000331877.1 | -0.825220462 | 2.6649E-13 | 4.7174E-12 |
| STM14_RS12920 | | PTS ascorbate transporter subunit IIC | WP_001522266.1 | -0.827283405 | 2.8014E-06 | 1.6772E-05 |
| STM14_RS01395 | | D-threonate kinase | WP_000782192.1 | -0.82944272 | 6.0539E-05 | 0.00027051 |
| STM14_RS14115 | pdxJ | pyridoxine 5'-phosphate synthase | WP_000818972.1 | -0.830347693 | 1.4354E-09 | 1.4972E-08 |
| STM14_RS01025 | apaG | Co2+/Mg2+ efflux protein ApaG | WP_000610894.1 | -0.830747015 | 6.7525E-05 | 0.00029634 |
| STM14_RS01725 | lpxA | acyl-ACP--UDP-N-acetylglucosamine O-acyltransferase | WP_000565950.1 | -0.830886893 | 1.8415E-10 | 2.2158E-09 |
| STM14_RS08755 | | NarK family nitrate/nitrite MFS transporter | WP_000198251.1 | -0.832216634 | 0.00048328 | 0.00169954 |
| STM14_RS16090 | | NADP(H)-dependent aldo-keto reductase | WP_000558962.1 | -0.833842654 | 1.4585E-08 | 1.3392E-07 |
| STM14_RS22960 | | glutathionylspermidine synthase family protein | WP_000943932.1 | -0.83500794 | 0.00452478 | 0.01196857 |
| STM14_RS00905 | carB | carbamoyl-phosphate synthase large subunit | WP_001126305.1 | -0.835899001 | 3.6168E-10 | 4.123E-09 |
| STM14_RS03520 | | YbdK family carboxylate-amine ligase | WP_001196903.1 | -0.836466743 | 0.00065418 | 0.00220777 |
| STM14_RS05215 | infA | translation initiation factor IF-1 | WP_001040187.1 | -0.839748961 | 3.1206E-06 | 1.8532E-05 |
| STM14_RS10295 | | carbon-nitrogen hydrolase family protein | WP_000100258.1 | -0.841420984 | 5.7884E-09 | 5.6683E-08 |
| STM14_RS18935 | zntA | Zn(II)/Cd(II)/Pb(II) translocating P-type ATPase ZntA | WP_000106641.1 | -0.844971959 | 4.7625E-09 | 4.6761E-08 |
| STM14_RS11350 | | cobalt-precorrin-4 methyltransferase | WP_000004870.1 | -0.846952289 | 0.00011 | 0.00045508 |
| STM14_RS17685 | mtgA | monofunctional biosynthetic peptidoglycan transglycosylase | WP_000044652.1 | -0.849603183 | 8.0146E-07 | 5.2791E-06 |
| STM14_RS26915 | | sugar phosphate nucleotidyltransferase | WP_223151143.1 | -0.850300605 | 0.0010556 | 0.00335062 |
| STM14_RS13905 | pepB | aminopeptidase PepB | WP_000133541.1 | -0.851922549 | 8.0225E-10 | 8.562E-09 |
| STM14_RS02450 | prpR | propionate catabolism operon regulatory protein PrpR | WP_000205278.1 | -0.852122818 | 2.2846E-07 | 1.6824E-06 |
| STM14_RS08205 | tus | DNA replication terminus site-binding protein | WP_000092490.1 | -0.853188569 | 0.00103954 | 0.0033099 |
| STM14_RS04550 | pgl | 6-phosphogluconolactonase | WP_000815468.1 | -0.855928591 | 1.2862E-12 | 2.0326E-11 |
| STM14_RS12230 | fruK | 1-phosphofructokinase | WP_000091249.1 | -0.856864687 | 0.00120643 | 0.00378048 |
| STM14_RS18830 | ugpE | sn-glycerol-3-phosphate ABC transporter permease UgpE | WP_000572196.1 | -0.858952485 | 0.00236532 | 0.00687923 |
| STM14_RS22435 | | DUF485 domain-containing protein | WP_000999621.1 | -0.859243298 | 4.9071E-06 | 2.8012E-05 |
| STM14_RS11810 | dcd | dCTP deaminase | WP_001234783.1 | -0.859706523 | 1.1787E-07 | 9.2932E-07 |
| STM14_RS16950 | | AraC family transcriptional regulator | WP_000998789.1 | -0.859721979 | 0.00099188 | 0.00318684 |
| STM14_RS04175 | pxpB | 5-oxoprolinase subunit PxpB | WP_001188373.1 | -0.859982147 | 6.2088E-07 | 4.2023E-06 |
| STM14_RS08935 | | M42 family metallopeptidase | WP_000144617.1 | -0.861233863 | 1.456E-06 | 9.0402E-06 |
| STM14_RS15610 | | SDR family oxidoreductase | WP_001165740.1 | -0.862658623 | 8.4288E-05 | 0.00036045 |
| STM14_RS22405 | ghxP | guanine/hypoxanthine transporter GhxP | WP_000106907.1 | -0.863466324 | 1.8658E-07 | 1.3964E-06 |
| STM14_RS20085 | dsdA | D-serine ammonia-lyase | WP_000427997.1 | -0.865156237 | 0.00017965 | 0.00070222 |
| STM14_RS10290 | | DNA polymerase III subunit theta | WP_000856224.1 | -0.868253317 | 0.00163176 | 0.00496132 |
| STM14_RS06075 | agp | bifunctional glucose-1-phosphatase/inositol phosphatase | WP_000749201.1 | -0.869495069 | 1.5415E-18 | 4.5771E-17 |
| STM14_RS23995 | ettA | energy-dependent translational throttle protein EttA | WP_000046770.1 | -0.871240978 | 2.3489E-18 | 6.7567E-17 |
| STM14_RS16770 | yghU | glutathione-dependent disulfide-bond oxidoreductase | WP_000774852.1 | -0.872008883 | 2.7321E-11 | 3.7676E-10 |
| STM14_RS19200 | dppD | dipeptide ABC transporter ATP-binding protein | WP_001196509.1 | -0.873499176 | 5.1516E-08 | 4.3109E-07 |
| STM14_RS04780 | glnP | glutamine ABC transporter permease GlnP | WP_001159076.1 | -0.87828858 | 0.00032687 | 0.00119873 |
| STM14_RS08775 | narI | respiratory nitrate reductase subunit gamma | WP_000617106.1 | -0.879110302 | 0.00335669 | 0.00932075 |
| STM14_RS16450 | speB | agmatinase | WP_000105550.1 | -0.883504278 | 1.673E-10 | 2.0219E-09 |
| STM14_RS04305 | sucC | ADP-forming succinate--CoA ligase subunit beta | WP_001048590.1 | -0.884201433 | 1.6678E-24 | 8.8999E-23 |
| STM14_RS01595 | hemL | glutamate-1-semialdehyde 2%2C1-aminomutase | WP_000045268.1 | -0.884456336 | 1.9858E-11 | 2.7801E-10 |
| STM14_RS19195 | dppF | dipeptide ABC transporter ATP-binding subunit DppF | WP_000103563.1 | -0.885494799 | 4.6753E-06 | 2.6814E-05 |
| STM14_RS13660 | tmcA | tRNA cytosine(34) acetyltransferase TmcA | WP_001279831.1 | -0.886137144 | 8.4244E-09 | 8.0777E-08 |
| STM14_RS18745 | glgX | glycogen debranching protein GlgX | WP_000192491.1 | -0.887927443 | 4.8967E-20 | 1.6541E-18 |
| STM14_RS19795 | | hypothetical protein | WP_000745468.1 | -0.888019243 | 1.4769E-07 | 1.1329E-06 |
| STM14_RS19470 | aldB | aldehyde dehydrogenase AldB | WP_000183985.1 | -0.890229843 | 4.915E-22 | 2.1804E-20 |
| STM14_RS19300 | ghrB | glyoxylate/hydroxypyruvate reductase GhrB | WP_000804674.1 | -0.8906124 | 7.215E-09 | 6.9544E-08 |
| STM14_RS08055 | sodC2 | superoxide dismutase [Cu-Zn] SodC2 | WP_000826819.1 | -0.892586829 | 1.1219E-19 | 3.6557E-18 |
| STM14_RS03315 | ybcJ | ribosome-associated protein YbcJ | WP_000190278.1 | -0.897920117 | 0.0022407 | 0.0065271 |
| STM14_RS15605 | | gluconate:H+ symporter | WP_000202726.1 | -0.897961261 | 1.571E-10 | 1.9154E-09 |
| STM14_RS16340 | ubiH | 2-octaprenyl-6-methoxyphenyl hydroxylase | WP_000111140.1 | -0.901660996 | 3.0393E-11 | 4.1601E-10 |
| STM14_RS19800 | gltS | sodium/glutamate symporter | WP_000462256.1 | -0.902374045 | 1.852E-12 | 2.8532E-11 |
| STM14_RS20970 | fadB | fatty acid oxidation complex subunit alpha FadB | WP_000965998.1 | -0.902999496 | 1.7958E-16 | 4.4675E-15 |
| STM14_RS14000 | | serine hydroxymethyltransferase | WP_000919178.1 | -0.903159574 | 9.9092E-22 | 4.1461E-20 |
| STM14_RS21930 | purD | phosphoribosylamine--glycine ligase | WP_000866767.1 | -0.906303439 | 2.5188E-07 | 1.8446E-06 |
| STM14_RS17240 | | glutathione S-transferase family protein | WP_000531161.1 | -0.906474812 | 9.4111E-11 | 1.1949E-09 |

| | | | | | | |
|---|---|---|---|---|---|---|
| STM14_RS21345 | sodA | superoxide dismutase [Mn] | WP_000122635.1 | -0.908073094 | 1.1417E-24 | 6.1822E-23 |
| STM14_RS07730 | | L%2CD-transpeptidase family protein | WP_000817059.1 | -0.908353755 | 5.633E-15 | 1.2488E-13 |
| STM14_RS19565 | cysE | serine O-acetyltransferase | WP_001112113.1 | -0.912628037 | 7.4512E-12 | 1.0974E-10 |
| STM14_RS10735 | yedE | selenium metabolism membrane protein YedE/FdhT | WP_001535233.1 | -0.913078536 | 1.3914E-05 | 7.1554E-05 |
| STM14_RS16095 | lplT | lysophospholipid transporter LplT | WP_000004679.1 | -0.913251241 | 8.4457E-05 | 0.00036076 |
| STM14_RS08860 | patD | aminobutyraldehyde dehydrogenase | WP_000158107.1 | -0.913995 | 5.1377E-17 | 1.3322E-15 |
| STM14_RS10730 | | lipoprotein YedD | WP_000754695.1 | -0.914242278 | 9.2535E-06 | 4.9236E-05 |
| STM14_RS06895 | | S49 family peptidase | WP_001189493.1 | -0.915808089 | 0.002753 | 0.00783955 |
| STM14_RS04520 | modE | molybdenum-dependent transcriptional regulator | WP_001147410.1 | -0.916558628 | 2.0714E-07 | 1.5456E-06 |
| STM14_RS12975 | hisM | histidine ABC transporter permease HisM | WP_000569771.1 | -0.917536022 | 9.6526E-06 | 5.1125E-05 |
| STM14_RS17980 | prmA | 50S ribosomal protein L11 methyltransferase | WP_001145849.1 | -0.920466182 | 0.00042252 | 0.00150893 |
| STM14_RS19545 | trmL | tRNA (uridine(34)/cytosine(34)/5-carboxymethylaminomethyluridine(34 | WP_000932335.1 | -0.920640577 | 0.00049792 | 0.00174603 |
| STM14_RS11925 | | PfkB family carbohydrate kinase | WP_001012658.1 | -0.921968367 | 0.00891969 | 0.02164951 |
| STM14_RS20590 | ilvD | dihydroxy-acid dehydratase | WP_001127434.1 | -0.92332395 | 7.5416E-10 | 8.0721E-09 |
| STM14_RS14050 | purL | phosphoribosylformylglycinamidine synthase | WP_000970045.1 | -0.925439738 | 3.5747E-13 | 6.2379E-12 |
| STM14_RS02155 | fadE | acyl-CoA dehydrogenase FadE | WP_000973041.1 | -0.928937264 | 7.7847E-16 | 1.8257E-14 |
| STM14_RS19215 | | ABC transporter substrate-binding protein | WP_000028047.1 | -0.931690493 | 3.5048E-21 | 1.3168E-19 |
| STM14_RS22965 | | isovaleryl-CoA dehydrogenase | WP_000118960.1 | -0.93943989 | 1.1342E-21 | 4.5891E-20 |
| STM14_RS00150 | | EAL domain-containing protein | WP_000545754.1 | -0.949061582 | 0.00071367 | 0.00238667 |
| STM14_RS09905 | | D-amino acid dehydrogenase | WP_001266937.1 | -0.960361303 | 1.7915E-21 | 6.9434E-20 |
| STM14_RS08960 | | ribulose-phosphate 3 epimerase family protein | WP_000600616.1 | -0.960849279 | 0.00531794 | 0.01379891 |
| STM14_RS04565 | hutG | formimidoylglutamase | WP_000195682.1 | -0.962681595 | 1.3456E-10 | 1.6739E-09 |
| STM14_RS12805 | nuoK | NADH-quinone oxidoreductase subunit NuoK | WP_000612687.1 | -0.965957227 | 9.6711E-07 | 6.2581E-06 |
| STM14_RS14440 | rseC | SoxR-reducing system protein RseC | WP_000589087.1 | -0.968892288 | 0.00022119 | 0.00084746 |
| STM14_RS08760 | | nitrate reductase subunit alpha | WP_000040398.1 | -0.970807627 | 9.0093E-13 | 1.4613E-11 |
| STM14_RS20965 | fadA | acetyl-CoA C-acyltransferase FadA | WP_000438778.1 | -0.970849976 | 1.1714E-12 | 1.8591E-11 |
| STM14_RS13250 | | alpha-keto acid decarboxylase family protein | WP_001180855.1 | -0.975293606 | 2.1271E-12 | 3.2634E-11 |
| STM14_RS19490 | | glutathione S-transferase | WP_000766684.1 | -0.976070009 | 4.0351E-17 | 1.0612E-15 |
| STM14_RS03350 | sfmF | fimbria assembly protein | WP_000603405.1 | -0.976613194 | 7.915E-06 | 4.292E-05 |
| STM14_RS09460 | acnA | aconitate hydratase AcnA | WP_000099475.1 | -0.981955303 | 2.6803E-31 | 2.2951E-29 |
| STM14_RS14975 | lhgO | L-2-hydroxyglutarate oxidase | WP_000271907.1 | -0.99009518 | 1.5312E-11 | 2.1768E-10 |
| STM14_RS09000 | | LysR substrate-binding domain-containing protein | WP_000414259.1 | -0.995474871 | 4.2287E-13 | 7.2757E-12 |
| STM14_RS16985 | yqiA | esterase YqiA | WP_000105725.1 | -0.995761915 | 2.93E-06 | 1.7457E-05 |
| STM14_RS09900 | dadX | catabolic alanine racemase DadX | WP_000197903.1 | -1.01955399 | 1.9285E-19 | 6.1212E-18 |
| STM14_RS03340 | | fimbrial biogenesis usher protein | WP_000754233.1 | -1.021713777 | 1.0189E-21 | 4.2152E-20 |
| STM14_RS03585 | entE | (2%2C3-dihydroxybenzoyl)adenylate synthase EntE | WP_001222445.1 | -1.022175601 | 0.00047129 | 0.00166057 |
| STM14_RS16115 | lysA | diaminopimelate decarboxylase | WP_000056576.1 | -1.022513069 | 3.0493E-08 | 2.6355E-07 |
| STM14_RS20695 | wzxE | lipid III flippase WzxE | WP_000050295.1 | -1.025287241 | 5.7564E-06 | 3.2309E-05 |
| STM14_RS17600 | mlaB | lipid asymmetry maintenance protein MlaB | WP_000188843.1 | -1.026300663 | 1.7894E-07 | 1.3418E-06 |
| STM14_RS04065 | tcuA | FAD-dependent tricarballylate dehydrogenase TcuA | WP_000228709.1 | -1.031863162 | 0.00013971 | 0.00056343 |
| STM14_RS17385 | | NAD(P)H-binding protein | WP_001521094.1 | -1.032635076 | 3.4866E-07 | 2.4833E-06 |
| STM14_RS17230 | | YqjK family protein | WP_000095496.1 | -1.039266917 | 3.8446E-13 | 6.6459E-12 |
| STM14_RS05035 | potH | putrescine ABC transporter permease PotH | WP_000105452.1 | -1.05047895 | 0.00178453 | 0.00537696 |
| STM14_RS09865 | | Glu/Leu/Phe/Val dehydrogenase | WP_001526439.1 | -1.055712786 | 2.2712E-32 | 2.0396E-30 |
| STM14_RS26800 | | DUF4222 domain-containing protein | | -1.057018423 | 0.00062156 | 0.00211708 |
| STM14_RS14005 | hmpA | NO-inducible flavohemoprotein | WP_000883146.1 | -1.057533327 | 1.4955E-08 | 1.3663E-07 |
| STM14_RS11345 | cbiG | cobalt-precorrin 5A hydrolase | WP_001098914.1 | -1.068545064 | 1.0952E-08 | 1.0234E-07 |
| STM14_RS04180 | | biotin-dependent carboxyltransferase family protein | WP_000932050.1 | -1.074231976 | 5.2365E-11 | 6.9355E-10 |
| STM14_RS01320 | ampE | beta-lactamase regulator AmpE | WP_000172039.1 | -1.097826641 | 1.7553E-07 | 1.3244E-06 |
| STM14_RS08990 | | carboxylesterase/lipase family protein | WP_000933996.1 | -1.10231276 | 5.4562E-14 | 1.0574E-12 |
| STM14_RS14110 | acpS | holo-ACP synthase | WP_000986043.1 | -1.104064948 | 2.8991E-07 | 2.0889E-06 |
| STM14_RS15120 | luxS | S-ribosylhomocysteine lyase | WP_001130194.1 | -1.104298882 | 4.3581E-25 | 2.4055E-23 |
| STM14_RS12990 | argT | lysine/arginine/ornithine ABC transporter substrate-binding protein ArgT | WP_000754423.1 | -1.106259786 | 1.4186E-20 | 5.1716E-19 |
| STM14_RS22430 | actP | cation/acetate symporter ActP | WP_000832536.1 | -1.115625376 | 1.1982E-14 | 2.5503E-13 |
| STM14_RS14080 | murQ | N-acetylmuramic acid 6-phosphate etherase | WP_001048530.1 | -1.120199705 | 4.9633E-06 | 2.8245E-05 |
| STM14_RS22305 | tyrB | aromatic amino acid transaminase | WP_000486903.1 | -1.134265912 | 8.0867E-10 | 8.5807E-09 |
| STM14_RS12910 | | transketolase family protein | WP_000593016.1 | -1.147532166 | 4.1512E-15 | 9.3771E-14 |
| STM14_RS19805 | xanP | xanthine/proton symporter XanP | WP_000115428.1 | -1.155944888 | 0.00029594 | 0.00109402 |
| STM14_RS03045 | recR | recombination mediator RecR | WP_001195023.1 | -1.18968564 | 3.7639E-10 | 4.2774E-09 |
| STM14_RS19210 | dppB | dipeptide ABC transporter permease DppB | WP_000938843.1 | -1.199671086 | 6.1171E-13 | 1.0191E-11 |
| STM14_RS16780 | | ABC transporter substrate-binding protein | WP_001032007.1 | -1.203060389 | 9.4515E-08 | 7.5653E-07 |
| STM14_RS14995 | csiR | DNA-binding transcriptional regulator CsiR | WP_000126152.1 | -1.203093585 | 6.0501E-06 | 3.3804E-05 |
| STM14_RS21400 | | aminoimidazole riboside kinase | WP_000646499.1 | -1.215022551 | 0.00042904 | 0.00152925 |
| STM14_RS01805 | rcsF | Rcs stress response system protein RcsF | WP_001202315.1 | -1.251450944 | 1.0681E-15 | 2.4892E-14 |
| STM14_RS14990 | gabP | GABA permease | WP_000531282.1 | -1.257470946 | 5.5769E-16 | 1.3248E-14 |
| STM14_RS04840 | | DUF1479 domain-containing protein | WP_000210950.1 | -1.283916986 | 1.1497E-06 | 7.3455E-06 |
| STM14_RS07355 | gdhA | NADP-specific glutamate dehydrogenase | WP_000372869.1 | -1.288826665 | 6.121E-11 | 7.9638E-10 |
| STM14_RS23685 | | NAD-dependent succinate-semialdehyde dehydrogenase | WP_000331411.1 | -1.302694766 | 1.7932E-31 | 1.572E-29 |
| STM14_RS18835 | ugpA | sn-glycerol-3-phosphate ABC transporter permease UgpA | WP_000099303.1 | -1.319803818 | 3.66E-06 | 2.1392E-05 |
| STM14_RS03345 | fimH | type 1 fimbria D-mannose specific adhesin FimH | WP_000708653.1 | -1.322290465 | 5.5716E-22 | 2.4422E-20 |
| STM14_RS16545 | ruvX | Holliday junction resolvase RuvX | WP_000017096.1 | -1.32495731 | 0.00017704 | 0.00069422 |
| STM14_RS11405 | pduC | propanediol dehydratase large subunit PduC | WP_001256896.1 | -1.341686826 | 0.00015795 | 0.00062668 |
| STM14_RS11905 | yegS | lipid kinase YegS | WP_001273389.1 | -1.376888115 | 2.6299E-19 | 8.2062E-18 |
| STM14_RS20810 | yigB | 5-amino-6-(5-phospho-D-ribitylamino)uracil phosphatase YigB | WP_001213560.1 | -1.384918606 | 6.0904E-11 | 7.9521E-10 |
| STM14_RS22440 | acs | acetate--CoA ligase | WP_000083882.1 | -1.443030949 | 5.9196E-48 | 1.3622E-45 |
| STM14_RS07390 | astB | N-succinylarginine dihydrolase | WP_000123952.1 | -1.464742804 | 5.8569E-33 | 5.675E-31 |
| STM14_RS07375 | | aspartate aminotransferase family protein | WP_000059493.1 | -1.52596023 | 4.04E-53 | 1.6528E-50 |
| STM14_RS13170 | fadL | long-chain fatty acid transporter FadL | WP_001518618.1 | -1.555252542 | 3.426E-47 | 7.4203E-45 |
| STM14_RS19205 | dppC | dipeptide ABC transporter permease DppC | WP_000084690.1 | -1.633127373 | 2.2001E-14 | 4.5256E-13 |
| STM14_RS07395 | astE | succinylglutamate desuccinylase | WP_000368454.1 | -1.690899338 | 6.8548E-24 | 3.3652E-22 |
| STM14_RS14985 | gabT | 4-aminobutyrate--2-oxoglutarate transaminase | WP_001095556.1 | -1.728891256 | 1.144E-38 | 1.6201E-36 |
| STM14_RS07380 | astA | arginine N-succinyltransferase | WP_001263889.1 | -1.743280593 | 4.5429E-44 | 7.9653E-42 |

| | | | | | | |
|---|---|---|---|---|---|---|
| STM14_RS14980 | gabD | NADP-dependent succinate-semialdehyde dehydrogenase | WP_001176529.1 | -1.799221155 | 1.288E-27 | 8.6224E-26 |
| STM14_RS07385 | astD | succinylglutamate-semialdehyde dehydrogenase | WP_000177268.1 | -1.877598045 | 6.2803E-65 | 3.3034E-62 |
| STM14_RS04575 | hutU | urocanate hydratase | WP_001115209.1 | -1.927429488 | 8.8704E-37 | 1.1665E-34 |
| STM14_RS04580 | hutH | histidine ammonia-lyase | WP_001095227.1 | -2.147460952 | 1.2716E-42 | 2.0356E-40 |
