## Supplementary Table 3 for "Commensal Yeast Promotes *Salmonella* Typhimurium Virulence"

**Table S3: KEGG analysis of pathways differentially regulated in STm co-culture with C. albicans ATCC compared to STm alone.**

| Database | Term | Description | Common | GeneListTotal | PathwayTotal | BackgroundTotal | Log2 Fold Enrichment | PValue | QValue | Column1 |
|---|---|---|---|---|---|---|---|---|---|---|
| seo_pathway.gmt | seo00640 | Propanoate metabolism | 26 | 747 | 30 | 1446 | 2.65 | 6.44E-05 | 2.77E-03 | STM14_0185,STM14_0190,STM14_3586,STM14_5141,STM14_0434,STM14_2529,STM14_0858,STM14_2882,STM14_0857,STM14_0274,STM14_4077,STM14_1099,STM14_4939,STM14_2883,STM14_0433,STM14_3831,STM14_2546,STM14_2914,STM14_4789,STM14_3368,STM14_3924,STM14_3923,STM14_0431,STM14_0432,STM14_4076,STM14_1221 |
| seo_pathway.gmt | seo00620 | Pyruvate metabolism | 41 | 747 | 52 | 1446 | 1.86 | 5.38E-05 | 2.77E-03 | STM14_4396,STM14_0185,STM14_3646,STM14_4052,STM14_1299,STM14_5141,STM14_5173,STM14_2113,STM14_0885,STM14_0183,STM14_3021,STM14_2882,STM14_0274,STM14_1639,STM14_4077,STM14_1772,STM14_1099,STM14_2296,STM14_2883,STM14_2672,STM14_4954,STM14_5221,STM14_2914,STM14_5220,STM14_1733,STM14_5383,STM14_3031,STM14_3024,STM14_1995,STM14_3923,STM14_1991,STM14_1127,STM14_4453,STM14_4438,STM14_1968,STM14_4076,STM14_0184,STM14_1891,STM14_1770,STM14_4863,STM14_4215 |
| seo_pathway.gmt | seo01120 | Microbial metabolism in diverse environments | 143 | 747 | 232 | 1446 | 0.70 | 9.57E-04 | 0.03 | STM14_3557,STM14_0911,STM14_2575,STM14_5075,STM14_4543,STM14_2292,STM14_2886,STM14_3034,STM14_2885,STM14_0815,STM14_4911,STM14_4905,STM14_4396,STM14_2153,STM14_3708,STM14_1610,STM14_2642,STM14_3701,STM14_3033,STM14_5302,STM14_0185,STM14_0190,STM14_3567,STM14_2070,STM14_0849,STM14_3929,STM14_3133,STM14_3646,STM14_3586,STM14_4052,STM14_1299,STM14_5141,STM14_4820,STM14_4929,STM14_1905,STM14_5173,STM14_2113,STM14_5326,STM14_4748,STM14_2721,STM14_2903,STM14_4735,STM14_1964,STM14_0885,STM14_0183,STM14_2888,STM14_3021,STM14_0898,STM14_4459,STM14_0858,STM14_3052,STM14_2882,STM14_3640,STM14_0857,STM14_3046,STM14_3896,STM14_0274,STM14_2956,STM14_1639,STM14_4077,STM14_1772,STM14_1099,STM14_2296,STM14_2883,STM14_0852,STM14_1105,STM14_0003,STM14_2672,STM14_4954,STM14_3638,STM14_0253,STM14_5221,STM14_1577,STM14_1247,STM14_1659,STM14_4773,STM14_3106,STM14_1934,STM14_5430,STM14_2938,STM14_2914,STM14_5220,STM14_0855,STM14_4864,STM14_1565,STM14_0075,STM14_5383,STM14_3031,STM14_3367,STM14_2555,STM14_4789,STM14_2788,STM14_1995,STM14_5289,STM14_0632,STM14_4020,STM14_3366,STM14_0241,STM14_0004,STM14_2707,STM14_3368,STM14_4187,STM14_1252,STM14_3923,STM14_0002,STM14_1906,STM14_0388,STM14_1991,STM14_1250,STM14_0435,STM14_1842,STM14_4620,STM14_0854,STM14_4438,STM14_1968,STM14_3138,STM14_4076,STM14_1221,STM14_3590,STM14_2969,STM14_4021,STM14_0184,STM14_1243,STM14_4260,STM14_5360,STM14_1891,STM14_1770,STM14_0856,STM14_4906,STM14_2683,STM14_1908,STM14_4863,STM14_0851,STM14_4215,STM14_2887,STM14_4245,STM14_4788,STM14_2170,STM14_3365,STM14_1270,STM14_2162,STM14_1856,STM14_2161 |
| seo_pathway.gmt | seo00310 | Lysine degradation | 16 | 747 | 18 | 1446 | 2.93 | 1.36E-03 | 0.03 | STM14_0185,STM14_3646,STM14_1964,STM14_3640,STM14_3896,STM14_1659,STM14_1934,STM14_5430,STM14_3367,STM14_4789,STM14_3366,STM14_3368,STM14_2907,STM14_3138,STM14_0856,STM14_3365 |
| seo_pathway.gmt | seo01200 | Carbon metabolism | 71 | 747 | 107 | 1446 | 0.95 | 1.74E-03 | 0.03 | STM14_3557,STM14_0911,STM14_2575,STM14_5075,STM14_2292,STM14_2886,STM14_3034,STM14_2885,STM14_4911,STM14_4905,STM14_2153,STM14_3708,STM14_1610,STM14_2642,STM14_3701,STM14_3033,STM14_5302,STM14_0185,STM14_0190,STM14_2070,STM14_0849,STM14_3687,STM14_3133,STM14_3646,STM14_3689,STM14_1600,STM14_4052,STM14_5141,STM14_3688,STM14_5173,STM14_0885,STM14_0183,STM14_4459,STM14_0858,STM14_2882,STM14_0857,STM14_0274,STM14_2956,STM14_1639,STM14_4077,STM14_1772,STM14_2296,STM14_2883,STM14_0852,STM14_1105,STM14_4702,STM14_4954,STM14_5221,STM14_2914,STM14_5220,STM14_0855,STM14_3582,STM14_1565,STM14_3031,STM14_4789,STM14_1995,STM14_0632,STM14_2707,STM14_3922,STM14_0854,STM14_3926,STM14_1968,STM14_4076,STM14_0184,STM14_5360,STM14_1770,STM14_0856,STM14_4906,STM14_0851,STM14_4215,STM14_2170 |
| seo_pathway.gmt | seo00071 | Fatty acid degradation | 12 | 747 | 13 | 1446 | 3.51 | 3.49E-03 | 0.03 | STM14_3646,STM14_2113,STM14_2199,STM14_2938,STM14_1005,STM14_4789,STM14_0365,STM14_1968,STM14_1891,STM14_4863,STM14_4788,STM14_3634 |
| seo_pathway.gmt | seo00470 | D-Amino acid metabolism | 15 | 747 | 17 | 1446 | 2.83 | 2.42E-03 | 0.03 | STM14_4748,STM14_2371,STM14_3640,STM14_5105,STM14_4966,STM14_3638,STM14_5417,STM14_0152,STM14_4591,STM14_2907,STM14_2179,STM14_0156,STM14_2178,STM14_1843,STM14_2029 |
| seo_pathway.gmt | seo05132 | Salmonella infection | 24 | 747 | 31 | 1446 | 1.71 | 3.51E-03 | 0.03 | STM14_1565,STM14_2378,STM14_3338,STM14_1702,STM14_3472,STM14_1237,STM14_5562,STM14_3481,STM14_1400,STM14_0572,STM14_3484,STM14_928,STM14_2824,STM14_3164,STM14_3471,STM14_1232,STM14_1193,STM14_2244,STM14_3350,STM14_3477,STM14_3462,STM14_3483,STM14_2428,STM14_1974 |
| seo_pathway.gmt | seo00190 | Oxidative phosphorylation | 32 | 747 | 43 | 1446 | 1.48 | 2.86E-03 | 0.03 | STM14_3066,STM14_0521,STM14_0519,STM14_4664,STM14_0522,STM14_0852,STM14_2866,STM14_4667,STM14_2864,STM14_5221,STM14_2858,STM14_5220,STM14_2863,STM14_2859,STM14_2867,STM14_2865,STM14_0520,STM14_4663,STM14_0854,STM14_2862,STM14_2860,STM14_1385,STM14_2861,STM14_0523,STM14_0851,STM14_2857,STM14_0861,STM14_5301,STM14_0422,STM14_0423,STM14_0862,STM14_0863 |
| seo_pathway.gmt | seo00020 | Citrate cycle (TCA cycle) | 22 | 747 | 29 | 1446 | 1.58 | 8.48E-03 | 0.06 | STM14_0185,STM14_0190,STM14_2070,STM14_0849,STM14_4052,STM14_5173,STM14_0885,STM14_0183,STM14_0858,STM14_0857,STM14_1772,STM14_0852,STM14_5221,STM14_5220,STM14_0855,STM14_1995,STM14_0854,STM14_0184,STM14_1770,STM14_0856,STM14_0851,STM14_4215 |
| seo_pathway.gmt | seo00300 | Lysine biosynthesis | 11 | 747 | 13 | 1446 | 2.38 | 0.02 | 0.12 | STM14_4929,STM14_4748,STM14_0149,STM14_3052,STM14_3046,STM14_3638,STM14_0253,STM14_0075,STM14_0002,STM14_0150,STM14_4260 |
| seo_pathway.gmt | seo00630 | Glyoxylate and dicarboxylate metabolism | 20 | 747 | 27 | 1446 | 1.44 | 0.02 | 0.12 | STM14_4396,STM14_0185,STM14_0190,STM14_2122,STM14_3567,STM14_2070,STM14_0849,STM14_3929,STM14_3687,STM14_3133,STM14_3646,STM14_3689,STM14_3586,STM14_1600,STM14_4052,STM14_3930,STM14_1299,STM14_5141,STM14_3688,STM14_4820 |
| seo_pathway.gmt | seo00010 | Glycolysis / Gluconeogenesis | 28 | 747 | 40 | 1446 | 1.16 | 0.02 | 0.12 | STM14_3557,STM14_5075,STM14_0815,STM14_4911,STM14_3708,STM14_1610,STM14_2642,STM14_5302,STM14_0185,STM14_5141,STM14_2113,STM14_0183,STM14_0898,STM14_2956,STM14_1639,STM14_2296,STM14_1565,STM14_5383,STM14_1995,STM14_4438,STM14_1968,STM14_0184,STM14_1891,STM14_4906,STM14_4863,STM14_4215,STM14_1270,STM14_4554 |
| seo_pathway.gmt | seo00330 | Arginine and proline metabolism | 16 | 747 | 21 | 1446 | 1.60 | 0.03 | 0.12 | STM14_1583,STM14_0379,STM14_3729,STM14_1584,STM14_3896,STM14_3720,STM14_4025,STM14_1586,STM14_1934,STM14_0378,STM14_0456,STM14_1585,STM14_0197,STM14_0818,STM14_1582,ST |
| seo_pathway.gmt | seo03070 | Bacterial secretion system | 23 | 747 | 32 | 1446 | 1.28 | 0.03 | 0.13 | STM14_0481,STM14_3976,STM14_4299,STM14_4637,STM14_4779,STM14_0482,STM14_0736,STM14_4781,STM14_0161,STM14_1711,STM14_1689,STM14_1702,STM14_3490,STM14_3472,STM14_3495,STM14_3785,STM14_1714,STM14_1717,STM14_1715,STM14_1705,STM14_1718,STM14_1710,STM14_0333 |
| seo_pathway.gmt | seo01110 | Biosynthesis of secondary metabolites | 187 | 747 | 329 | 1446 | 0.39 | 0.03 | 0.14 | STM14_3557,STM14_0911,STM14_2575,STM14_5075,STM14_2292,STM14_2886,STM14_0201,STM14_3034,STM14_2885,STM14_0815,STM14_4911,STM14_4905,STM14_4396,STM14_2153,STM14_3708,STM14_1610,STM14_2642,STM14_3701,STM14_3033,STM14_5302,STM14_0185,STM14_0190,STM14_3567,STM14_2070,STM14_0849,STM14_3929,STM14_3687,STM14_3133,STM14_3646,STM14_3689,STM14_1600,STM14_4052,STM14_1299,STM14_5141,STM14_3688,STM14_4929,STM14_5173,STM14_3145,STM14_5358,STM14_0519,STM14_2113,STM14_4720,STM14_4748,STM14_2903,STM14_0459,STM14_4735,STM14_4232,STM14_4199,STM14_3530,STM14_0140,STM14_0885,STM14_0054,STM14_0379,STM14_0183,STM14_5229,STM14_0782,STM14_3729,STM14_3869,STM14_0898,STM14_4459,STM14_0858,STM14_3052,STM14_0857,STM14_1635,STM14_0274,STM14_3076,STM14_2956,STM14_1639,STM14_4077,STM14_1772,STM14_2296,STM14_4700,STM14_0852,STM14_1105,STM14_2053,STM14_0003,STM14_3532,STM14_4702,STM14_3638,STM14_5221,STM14_3480,STM14_2590,STM14_0465,STM14_2844,STM14_5361,STM14_3065,STM14_2938,STM14_3533,STM14_2914,STM14_4793,STM14_5220,STM14_0855,STM14_0462,STM14_0378,STM14_2578,STM14_0059,STM14_2572,STM14_3582,STM14_2848,STM14_1367,STM14_2118,STM14_1005,STM14_1565,STM14_1883,STM14_0075,STM14_5383,STM14_5035,STM14_1369,STM14_4789,STM14_2577,STM14_1995,STM14_5106,STM14_4020,STM14_0241,STM14_0004,STM14_4187,STM14_0624,STM14_0456,STM14_2591,STM14_0002,STM14_4955,STM14_4256,STM14_0792,STM14_2570,STM14_2810,STM14_2172,STM14_3922,STM14_5016,STM14_5017,STM14_0261,STM14_3692,STM14_0576,STM14_3883,STM14_0435,STM14_0854,STM14_2565,STM14_3926,STM14_4438,STM14_4815,STM14_1968,STM14_3138,STM14_4751,STM14_4258,STM14_990,STM14_4076,STM14_2969,STM14_4021,STM14_0184,STM14_0818,STM14_4260,STM14_4706,STM14_0623,STM14_4701,STM14_5241,STM14_5008,STM14_1891,STM14_1770,STM14_3050,STM14_4255,STM14_3098,STM14_3694,STM14_0856,STM14_4906,STM14_2569,STM14_5169,STM14_2845,STM14_3671,STM14_3064,STM14_1531,STM14_0132,STM14_4231,STM14_4863,STM14_0851,STM14_4215,STM14_4245,STM14_1881,STM14_4788,STM14_4918,STM14_2843,STM14_2821,STM14_2819,STM14_4460,STM14_2820,STM14_0696,STM14_0695,STM14_3992,STM14_0694 |
| seo_pathway.gmt | seo01220 | Degradation of aromatic compounds | 8 | 747 | 9 | 1446 | 2.92 | 0.04 | 0.15 | STM14_2113,STM14_2903,STM14_1247,STM14_1252,STM14_1250,STM14_1968,STM14_1891,STM14_4863 |
| seo_pathway.gmt | seo01212 | Fatty acid metabolism | 15 | 747 | 20 | 1446 | 1.51 | 0.04 | 0.16 | STM14_3646,STM14_2199,STM14_0274,STM14_2927,STM14_4077,STM14_2053,STM14_2938,STM14_2914,STM14_1367,STM14_1005,STM14_4789,STM14_0365,STM14_4076,STM14_1366,STM14_4788 |
| seo_pathway.gmt | seo00785 | Lipoic acid metabolism | 9 | 747 | 11 | 1446 | 2.09 | 0.07 | 0.22 | STM14_0185,STM14_3687,STM14_3689,STM14_3688,STM14_0183,STM14_0855,STM14_0184,STM14_0737,STM14_0856 |
| seo_pathway.gmt | seo00030 | Pentose phosphate pathway | 23 | 747 | 34 | 1446 | 0.99 | 0.08 | 0.25 | STM14_0911,STM14_4682,STM14_5487,STM14_2575,STM14_5075,STM14_4543,STM14_2292,STM14_2886,STM14_0201,STM14_3034,STM14_2885,STM14_0815,STM14_4911,STM14_4574,STM14_4905,STM14_4396,STM14_2153,STM14_3708,STM14_1610,STM14_2642,STM14_3701,STM14_3033,STM14_5302 |
| seo_pathway.gmt | seo01100 | Metabolic pathways | 445 | 747 | 829 | 1446 | 0.27 | 0.08 | 0.25 | STM14_3557,STM14_0911,STM14_4682,STM14_5487,STM14_2575,STM14_5075,STM14_4543,STM14_2292,STM14_2886,STM14_0201,STM14_3034,STM14_2885,STM14_0815,STM14_4911,STM14_4574,STM14_4905,STM14_4396,STM14_2153,STM14_3708,STM14_1610,STM14_2642,STM14_3701,STM14_3033,STM14_5302,STM14_0185,STM14_0190,STM14_2122,STM14_3567,STM14_2070,STM14_0849,STM14_3929,STM14_3687,STM14_3133,STM14_3646,STM14_3689,STM14_3586,STM14_1600,STM14_4052,STM14_3930,STM14_1299,STM14_5141,STM14_3688,STM14_4820,STM14_3015,STM14_4929,STM14_2103,STM14_0521,STM14_0917,STM14_1905,STM14_5173,STM14_3145,STM14_4358,STM14_0366,STM14_3978,STM14_5358,STM14_0519,STM14_2113,STM14_2918,STM14_4720,STM14_5326,STM14_0434,STM14_5107,STM14_1200,STM14_5569,STM14_4748,STM14_1583,STM14_2721,STM14_2998,STM14_1590,STM14_4441,STM14_3900,STM14_4276,STM14_2371,STM14_2903,STM14_0459,STM14_4735,STM14_0149,STM14_4232,STM14_1964,STM14_5165,STM14_2407,STM14_4199,STM14_3530,STM14_5193,STM14_4563,STM14_4505,STM14_932,STM14_4975,STM14_4257,STM14_0140,STM14_0885,STM14_0203,STM14_0054,STM14_1749,STM14_4719,STM14_0379,STM14_0183,STM14_5229,STM14_2586,STM14_0782,STM14_1013,STM14_3729,STM14_5352,STM14_2529,STM14_4664,STM14_2888,STM14_2199,STM14_5287,STM14_3869,STM14_0268,STM14_3021,STM14_0898,STM14_4459,STM14_0858,STM14_0831,STM14_4027,STM14_2363,STM14_4561,STM14_3052,STM14_3939,STM14_0914,STM14_0900,STM14_4786,STM14_2520,STM14_2882,STM14_3640,STM14_0078,STM14_0857,STM14_5105,STM14_1635,STM14_4974,STM14_1584,STM14_3046,STM14_3896,STM14_0522,STM14_4469,STM14_0274,STM14_2617,STM14_2927,STM14_1612,STM14_4065,STM14_3076,STM14_2956,STM14_1639,STM14_4077,STM14_4471,STM14_1772,STM14_1099,STM14_4939,STM14_2296,STM14_2830,STM14_2552,STM14_4966,STM14_4700,STM14_2883,STM14_0852,STM14_3876,STM14_1105,STM14_2053,STM14_4327,STM14_3977,STM14_0496,STM14_0003,STM14_2672,STM14_3532,STM14_4702,STM14_2866,STM14_4954,STM14_4667,STM14_3638,STM14_0253,STM14_3739,STM14_2477,STM14_2864,STM14_5221,STM14_5171,STM14_2858,STM14_3402,STM14_3480,STM14_2958,STM14_2590,STM14_3720,STM14_4025,STM14_0465,STM14_0433,STM14_2844,STM14_1577,STM14_3769,STM14_3831,STM14_4473,STM14_5361,STM14_1247,STM14_1659,STM14_2894,STM14_4773,STM14_2546,STM14_3106,STM14_1586,STM14_3065,STM14_4711,STM14_4723,STM14_3151,STM14_1934,STM14_1117,STM14_5430,STM14_4470,STM14_2938,STM14_3533,STM14_2914,STM14_4793,STM14_5220,STM14_0855,STM14_5417,STM14_5202,STM14_1733,STM14_0462,STM14_1787,STM14_0378,STM14_0833,STM14_2578,STM14_0059,STM14_4739,STM14_2572,STM14_3582,STM14_2848,STM14_1367,STM14_3111,STM14_2504,STM14_4864,STM14_2118,STM14_1005,STM14_0899,STM14_1565,STM14_1883,STM14_0075,STM14_5383,STM14_5035,STM14_2514,STM14_3031,STM14_3367,STM14_2863,STM14_3856,STM14_3796,STM14_1369,STM14_2555,STM14_0832,STM14_4789,STM14_1955,STM14_2114,STM14_2517,STM14_3024,STM14_2577,STM14_2788,STM14_4444,STM14_1995,STM14_0247,STM14_2518,STM14_5289,STM14_5106,STM14_1323,STM14_0632,STM14_2859,STM14_4716,STM14_4020,STM14_2507,STM14_3366,STM14_0241,STM14_0004,STM14_2707,STM14_1012,STM14_3368,STM14_3924,STM14_4187,STM14_0624,STM14_1252,STM14_3016,STM14_0456,STM14_2867,STM14_0151,STM14_2591,STM14_0154,STM14_3923,STM14_1090,STM14_0152,STM14_0431,STM14_0002,STM14_1373,STM14_2865,STM14_4955,STM14_4256,STM14_0792,STM14_2570,STM14_0270,STM14_0913,STM14_4482,STM14_1585,STM14_4591,STM14_2810,STM14_2172,STM14_0520,STM14_1906,STM14_3922,STM14_3931,STM14_0388,STM14_1991,STM14_2516,STM14_5016,STM14_3423,STM14_2218,STM14_3158,STM14_5017,STM14_1250,STM14_2813,STM14_0261,STM14_1127,STM14_4663,STM14_2694,STM14_931,STM14_4475,STM14_2521,STM14_5180,STM14_0920,STM14_4453,STM14_2907,STM14_3692,STM14_0576,STM14_3883,STM14_1667,STM14_4718,STM14_0435,STM14_1842,STM14_0150,STM14_3157,STM14_4620,STM14_0854,STM14_4492,STM14_2565,STM14_3926,STM14_934,STM14_0107,STM14_4438,STM14_5505,STM14_4815,STM14_1089,STM14_0432,STM14_0751,STM14_0365,STM14_3994,STM14_2234,STM14_1968,STM14_3138,STM14_2512,STM14_4751,STM14_4258,STM14_990,STM14_2862,STM14_4076,STM14_0290,STM14_2081,STM14_1221,STM14_3590,STM14_0169,STM14_2969,STM14_2860,STM14_4021,STM14_0184,STM14_0197,STM14_0818,STM14_1385,STM14_1243,STM14_1366,STM14_2877,STM14_4260,STM14_0061,STM14_2179,STM14_5360,STM14_2861,STM14_4706,STM14_2812,STM14_2722,STM14_0623,STM14_4487,STM14_2515,STM14_4701,STM14_0155,STM14_0523,STM14_5241,STM14_3075,STM14_0106,STM14_5008,STM14_1891,STM14_1770,STM14_1582,STM14_3050,STM14_4973,STM14_4255,STM14_0737,STM14_3098,STM14_3694,STM14_0856,STM14_4906,STM14_0680,STM14_2683,STM14_0148,STM14_3039,STM14_0156,STM14_2569,STM14_5169,STM14_1332,STM14_1575,STM14_2845,STM14_4472,STM14_3671,STM14_0364,STM14_1908,STM14_0673,STM14_3064,STM14_1531,STM14_0132,STM14_0671,STM14_4014,STM14_4231,STM14_4863,STM14_2178,STM14_0851,STM14_1843,STM14_4215,STM14_2887,STM14_4245,STM14_1881,STM14_0758,STM14_4788,STM14_3418,STM14_3698,STM14_1395,STM14_4918,STM14_5373,STM14_2857,STM14_3747,STM14_1788,STM14_3558,STM14_2170,STM14_3365,STM14_2843,STM14_4003 |
| seo_pathway.gmt | seo02030 | Bacterial chemotaxis | 16 | 747 | 23 | 1446 | 1.11 | 0.09 | 0.28 | STM14_2334,STM14_3894,STM14_3893,STM14_2331,STM14_2332,STM14_1966,STM14_4305,STM14_2338,STM14_2852,STM14_2335,STM14_5446,STM14_2336,STM14_2397,STM14_2337,STM14_4681,STM14_4375 |
| seo_pathway.gmt | seo00280 | Valine, leucine and isoleucine degradation | 8 | 747 | 10 | 1446 | 1.91 | 0.11 | 0.29 | STM14_0185,STM14_3646,STM14_4700,STM14_2938,STM14_1005,STM14_4789,STM14_3368,STM14_4788 |
| seo_pathway.gmt | seo00260 | Glycine, serine and threonine metabolism | 23 | 747 | 35 | 1446 | 0.86 | 0.12 | 0.30 | STM14_4396,STM14_0185,STM14_3567,STM14_3929,STM14_3687,STM14_3133,STM14_3689,STM14_1299,STM14_3688,STM14_4929,STM14_3640,STM14_4469,STM14_1105,STM14_0003,STM14_4702,STM14_4470,STM14_3582,STM14_0004,STM14_0002,STM14_4591,STM14_3922,STM14_3926,STM14_4260 |
| seo_pathway.gmt | seo00860 | Porphyrin metabolism | 23 | 747 | 35 | 1446 | 0.86 | 0.12 | 0.30 | STM14_0519,STM14_4735,STM14_2520,STM14_4793,STM14_2504,STM14_2514,STM14_2517,STM14_2518,STM14_2507,STM14_0241,STM14_4187,STM14_2516,STM14_2521,STM14_0576,STM14_0435,STM14_4815,STM14_0751,STM14_2512,STM14_2081,STM14_2969,STM14_2515,STM14_5008,STM14_4146 |
| seo_pathway.gmt | seo00670 | One carbon pool by folate | 9 | 747 | 12 | 1446 | 1.50 | 0.15 | 0.35 | STM14_2122,STM14_3133,STM14_3689,STM14_3065,STM14_5035,STM14_0632,STM14_5017,STM14_0106,STM14_3698 |
| seo_pathway.gmt | seo00650 | Butanoate metabolism | 16 | 747 | 24 | 1446 | 0.92 | 0.15 | 0.35 | STM14_3646,STM14_2113,STM14_0140,STM14_1099,STM14_0852,STM14_5221,STM14_5430,STM14_5220,STM14_3367,STM14_4789,STM14_1995,STM14_3368,STM14_3923,STM14_1842,STM14_0854,STM14_0851 |
| seo_pathway.gmt | seo02040 | Flagellar assembly | 26 | 747 | 41 | 1446 | 0.72 | 0.15 | 0.35 | STM14_2374,STM14_1342,STM14_2378,STM14_2340,STM14_4008,STM14_2341,STM14_2338,STM14_3338,STM14_1354,STM14_2396,STM14_2397,STM14_2382,STM14_2394,STM14_2395,STM14_1347,STM14_2390,STM14_1355,STM14_2380,STM14_1345,STM14_2392,STM14_2372,STM14_1353,STM14_1341,STM14_2381,STM14_2337,STM14_2373 |
| seo_pathway.gmt | seo00910 | Nitrogen metabolism | 13 | 747 | 19 | 1446 | 1.03 | 0.17 | 0.36 | STM14_4820,STM14_1905,STM14_5569,STM14_0203,STM14_1577,STM14_2788,STM14_4020,STM14_1906,STM14_4021,STM14_5360,STM14_1908,STM14_2170,STM14_1903 |
| seo_pathway.gmt | seo00061 | Fatty acid biosynthesis | 9 | 747 | 13 | 1446 | 1.08 | 0.27 | 0.51 | STM14_2199,STM14_0274,STM14_2927,STM14_4077,STM14_2053,STM14_2914,STM14_1367,STM14_4076,STM14_1366 |
| seo_pathway.gmt | seo01250 | Biosynthesis of nucleotide sugars | 27 | 747 | 45 | 1446 | 0.50 | 0.29 | 0.51 | STM14_5075,STM14_0815,STM14_0366,STM14_4720,STM14_4719,STM14_2586,STM14_0900,STM14_2956,STM14_4471,STM14_3876,STM14_3977,STM14_2590,STM14_0465,STM14_4723,STM14_1117,STM14_2578,STM14_2118,STM14_0899,STM14_2577,STM14_2591,STM14_4256,STM14_4718,STM14_3994,STM14_0290,STM14_4973,STM14_1395,STM14_4003 |
| seo_pathway.gmt | seo00052 | Galactose metabolism | 13 | 747 | 21 | 1446 | 0.61 | 0.39 | 0.65 | STM14_0815,STM14_1610,STM14_4563,STM14_0898,STM14_4561,STM14_3939,STM14_0900,STM14_2956,STM14_5171,STM14_2118,STM14_0899,STM14_1955,STM14_4620 |
| seo_pathway.gmt | seo03430 | Mismatch repair | 14 | 747 | 23 | 1446 | 0.55 | 0.41 | 0.66 | STM14_0273,STM14_5114,STM14_5372,STM14_2982,STM14_1374,STM14_3674,STM14_3627,STM14_2281,STM14_4196,STM14_4752,STM14_4631,STM14_0502,STM14_0309,STM14_2559 |
| seo_pathway.gmt | seo02020 | Two-component system | 89 | 747 | 162 | 1446 | 0.21 | 0.40 | 0.66 | STM14_3646,STM14_4820,STM14_1905,STM14_5193,STM14_4719,STM14_5221,STM14_5220,STM14_1906,STM14_4718,STM14_1908,STM14_1843,STM14_2162,STM14_1856,STM14_2161,STM14_0861,STM14_0422,STM14_0423,STM14_0862,STM14_0863,STM14_2368,STM14_2374,STM14_2136,STM14_1342,STM14_5519,STM14_1501,STM14_0249,STM14_2334,STM14_2256,STM14_3894,STM14_3909,STM14_2378,STM14_4216,STM14_1680,STM14_2946,STM14_2340,STM14_2653,STM14_0771,STM14_3360,STM14_4616,STM14_3044,STM14_4217,STM14_4008,STM14_3893,STM14_3359,STM14_0471,STM14_2800,STM14_2331,STM14_2332,STM14_1966,STM14_4305,STM14_4173,STM14_0254,STM14_0470,STM14_4881,STM14_1686,STM14_4817,STM14_2163,STM14_0773,STM14_1408,STM14_2613,STM14_2341,STM14_2338,STM14_2775,STM14_1409,STM14_1497,STM14_2950,STM14_3363,STM14_3412,STM14_4882,STM14_5509,STM14_2797,STM14_2852,STM14_0641,STM14_0638,STM14_0286,STM14_4351,STM14_2265,STM14_1892,STM14_2335,STM14_1936,STM14_0637,STM14_4652,STM14_5534,STM14_1687,STM14_5163,STM14_0822,STM14_5446,STM14_3338,STM14_2336 |
| seo_pathway.gmt | seo00480 | Glutathione metabolism | 15 | 747 | 25 | 1446 | 0.50 | 0.43 | 0.67 | STM14_2575,STM14_2292,STM14_4276,STM14_1749,STM14_0831,STM14_4327,STM14_3739,STM14_2894,STM14_0833,STM14_3111,STM14_0832,STM14_1012,STM14_0818,STM14_0680,STM14_5373 |
| seo_pathway.gmt | seo00680 | Methane metabolism | 16 | 747 | 27 | 1446 | 0.45 | 0.44 | 0.67 | STM14_3557,STM14_4911,STM14_3708,STM14_1610,STM14_2642,STM14_5302,STM14_3133,STM14_4052,STM14_5141,STM14_2882,STM14_1639,STM14_2883,STM14_1105,STM14_4954,STM14_2707,STM14_1968 |
| seo_pathway.gmt | seo03060 | Protein export | 11 | 747 | 18 | 1446 | 0.56 | 0.48 | 0.68 | STM14_0481,STM14_3976,STM14_4299,STM14_4637,STM14_4779,STM14_0482,STM14_3162,STM14_0736,STM14_4781,STM14_0161,STM14_0056 |
| seo_pathway.gmt | seo00350 | Tyrosine metabolism | 11 | 747 | 18 | 1446 | 0.56 | 0.48 | 0.68 | STM14_2113,STM14_1247,STM14_5430,STM14_3367,STM14_5106,STM14_1252,STM14_1250,STM14_1968,STM14_1891,STM14_2683,STM14_4863 |
| seo_pathway.gmt | seo00340 | Histidine metabolism | 7 | 747 | 11 | 1446 | 0.72 | 0.55 | 0.75 | STM14_0917,STM14_0914,STM14_2572,STM14_2570,STM14_0913,STM14_2565,STM14_2569 |
| seo_pathway.gmt | seo00550 | Peptidoglycan biosynthesis | 15 | 747 | 26 | 1446 | 0.36 | 0.56 | 0.75 | STM14_0149,STM14_1013,STM14_2552,STM14_0151,STM14_0154,STM14_0152,STM14_2218,STM14_0150,STM14_3994,STM14_0155,STM14_4973,STM14_0148,STM14_3039,STM14_0156,STM14_4014 |
| seo_pathway.gmt | seo00900 | Terpenoid backbone biosynthesis | 8 | 747 | 13 | 1446 | 0.59 | 0.58 | 0.77 | STM14_3646,STM14_2903,STM14_3532,STM14_3533,STM14_0059,STM14_0261,STM14_3671,STM14_3992 |
| seo_pathway.gmt | seo00543 | Exopolysaccharide biosynthesis | 3 | 747 | 4 | 1446 | 1.49 | 0.63 | 0.79 | STM14_4459,STM14_4716,STM14_4725 |
| seo_pathway.gmt | seo00051 | Fructose and mannose metabolism | 19 | 747 | 34 | 1446 | 0.25 | 0.73 | 0.90 | STM14_4911,STM14_3708,STM14_1610,STM14_2642,STM14_5302,STM14_2721,STM14_2407,STM14_0465,STM14_2578,STM14_4864,STM14_2577,STM14_4444,STM14_3423,STM14_3590,STM14_2877,STM14_2722,STM14_4906,STM14_0673,STM14_0671 |
| seo_pathway.gmt | seo03018 | RNA degradation | 9 | 747 | 16 | 1446 | 0.27 | 0.80 | 0.94 | STM14_3621,STM14_1357,STM14_3964,STM14_3557,STM14_3962,STM14_951,STM14_5207,STM14_3066,STM14_4761 |
| seo_pathway.gmt | seo00270 | Cysteine and methionine metabolism | 20 | 747 | 37 | 1446 | 0.14 | 0.87 | 0.96 | STM14_4052,STM14_4929,STM14_2998,STM14_2371,STM14_4459,STM14_3640,STM14_4700,STM14_1105,STM14_3739,STM14_3402,STM14_3106,STM14_3582,STM14_5035,STM14_0247,STM14_5106,STM14_0002,STM14_3922,STM14_2234,STM14_0197,STM14_4260 |
| seo_pathway.gmt | seo03440 | Homologous recombination | 15 | 747 | 28 | 1446 | 0.11 | 0.85 | 0.96 | STM14_0273,STM14_5114,STM14_5372,STM14_1374,STM14_3674,STM14_2281,STM14_4631,STM14_0309,STM14_5276,STM14_3417,STM14_5459,STM14_3608,STM14_4509,STM14_2303,STM14_0571 |
| seo_pathway.gmt | seo00521 | Streptomycin biosynthesis | 5 | 747 | 9 | 1446 | 0.23 | 1.00 | 1.00 | STM14_0815,STM14_4720,STM14_2956,STM14_2590,STM14_2591 |
| seo_pathway.gmt | seo00250 | Alanine, aspartate and glutamate metabolism | 17 | 747 | 32 | 1446 | 0.09 | 1.00 | 1.00 | STM14_4820,STM14_5352,STM14_0078,STM14_1577,STM14_5430,STM14_5417,STM14_5202,STM14_3367,STM14_4020,STM14_3368,STM14_0792,STM14_1842,STM14_990,STM14_4021,STM14_0364,STM14_1843,STM14_2170 |
| seo_pathway.gmt | seo00500 | Starch and sucrose metabolism | 18 | 747 | 34 | 1446 | 0.08 | 1.00 | 1.00 | STM14_5075,STM14_0815,STM14_4358,STM14_4232,STM14_4257,STM14_2586,STM14_2956,STM14_0465,STM14_2118,STM14_1883,STM14_4256,STM14_2172,STM14_4258,STM14_4255,STM14_4231,STM14_1881,STM14_4554,STM14_1596 |
| seo_pathway.gmt | seo00130 | Ubiquinone and other terpenoid-quinone biosynt | 10 | 747 | 19 | 1446 | 0.06 | 1.00 | 1.00 | STM14_2903,STM14_0782,STM14_2844,STM14_2848,STM14_2810,STM14_3692,STM14_3694,STM14_2845,STM14_4918,STM14_2843 |
| seo_pathway.gmt | seo01230 | Biosynthesis of amino acids | 62 | 747 | 120 | 1446 | 4.66E-04 | 1.00 | 1.00 | STM14_3557,STM14_2886,STM14_3034,STM14_2885,STM14_4905,STM14_2153,STM14_3708,STM14_1610,STM14_2642,STM14_3701,STM14_3033,STM14_0190,STM14_2070,STM14_0849,STM14_3133,STM14_4820,STM14_4929,STM14_5358,STM14_4748,STM14_2998,STM14_0459,STM14_4199,STM14_0140,STM14_0379,STM14_4459,STM14_3052,STM14_1635,STM14_3046,STM14_2296,STM14_4700,STM14_1105,STM14_0003,STM14_4702,STM14_3638,STM14_0253,STM14_3402,STM14_0378,STM14_2572,STM14_3582,STM14_1565,STM14_0075,STM14_5035,STM14_0247,STM14_5106,STM14_4020,STM14_0004,STM14_0456,STM14_0002,STM14_4955,STM14_0792,STM14_2570,STM14_3922,STM14_2565,STM14_3926,STM14_4021,STM14_4260,STM14_4706,STM14_4701,STM14_4906,STM14_2569,STM14_1531,STM14_0132 |
| seo_pathway.gmt | seo00740 | Riboflavin metabolism | 7 | 747 | 14 | 1446 | -0.10 | 1.00 | 1.00 | STM14_5107,STM14_2903,STM14_5193,STM14_0054,STM14_3869,STM14_4786,STM14_4751 |
| seo_pathway.gmt | seo03030 | DNA replication | 9 | 747 | 18 | 1446 | -0.10 | 1.00 | 1.00 | STM14_0273,STM14_5114,STM14_5372,STM14_2982,STM14_1374,STM14_2281,STM14_4631,STM14_0309,STM14_0308 |
| seo_pathway.gmt | seo00564 | Glycerophospholipid metabolism | 14 | 747 | 28 | 1446 | -0.10 | 1.00 | 1.00 | STM14_3015,STM14_2103,STM14_5229,STM14_2363,STM14_3016,STM14_3883,STM14_3149,STM14_2817,STM14_2821,STM14_4764,STM14_3634,STM14_2819,STM14_4460,STM14_2820 |
| seo_pathway.gmt | seo01210 | 2-Oxocarboxylic acid metabolism | 16 | 747 | 32 | 1446 | -0.10 | 1.00 | 1.00 | STM14_0185,STM14_0190,STM14_2070,STM14_0849,STM14_0140,STM14_0183,STM14_4700,STM14_0855,STM14_1995,STM14_4955,STM14_0184,STM14_4260,STM14_4706,STM14_4701,STM14_0856,STM14_0132 |
| seo_pathway.gmt | seo01501 | beta-Lactam resistance | 9 | 747 | 19 | 1446 | -0.25 | 1.00 | 1.00 | STM14_2218,STM14_0148,STM14_2797,STM14_2108,STM14_2027,STM14_2110,STM14_2106,STM14_2107,STM14_0559 |
| seo_pathway.gmt | seo02060 | Phosphotransferase system (PTS) | 17 | 747 | 36 | 1446 | -0.26 | 1.00 | 1.00 | STM14_2721,STM14_4563,STM14_2888,STM14_4561,STM14_1955,STM14_3423,STM14_2722,STM14_0673,STM14_0671,STM14_2887,STM14_4010,STM14_0670,STM14_5449,STM14_0672,STM14_1596,STM14_2989,STM14_0798 |
| seo_pathway.gmt | seo00220 | Arginine biosynthesis | 8 | 747 | 17 | 1446 | -0.27 | 1.00 | 1.00 | STM14_4820,STM14_5358,STM14_1577,STM14_5361,STM14_4955,STM14_5360,STM14_1843,STM14_2170 |
| seo_pathway.gmt | seo00230 | Purine metabolism | 29 | 747 | 62 | 1446 | -0.29 | 1.00 | 1.00 | STM14_0815,STM14_2153,STM14_3145,STM14_3530,STM14_4505,STM14_5287,STM14_3076,STM14_2477,STM14_3065,STM14_4711,STM14_0462,STM14_4739,STM14_3856,STM14_0624,STM14_5016,STM14_5017,STM14_2813,STM14_0107,STM14_5505,STM14_0169,STM14_1243,STM14_5360,STM14_2812,STM14_0623,STM14_3075,STM14_3050,STM14_3098,STM14_3064,STM14_3747 |
| seo_pathway.gmt | seo02024 | Quorum sensing | 23 | 747 | 50 | 1446 | -0.34 | 1.00 | 1.00 | STM14_2199,STM14_1635,STM14_3402,STM14_2368,STM14_2340,STM14_4173,STM14_2341,STM14_2058,STM14_4897,STM14_0481,STM14_2108,STM14_2027,STM14_1926,STM14_3976,STM14_4904,STM14_4896,STM14_4299,STM14_2110,STM14_2106,STM14_4637,STM14_2107,STM14_4294,STM14_5098 |
| seo_pathway.gmt | seo00520 | Amino sugar and nucleotide sugar metabolism | 22 | 747 | 48 | 1446 | -0.35 | 1.00 | 1.00 | STM14_5075,STM14_0815,STM14_4719,STM14_2586,STM14_4027,STM14_0900,STM14_2956,STM14_3977,STM14_0465,STM14_3151,STM14_2578,STM14_2118,STM14_0899,STM14_2577,STM14_4256,STM14_4718,STM14_3994,STM14_4973,STM14_0673,STM14_0671,STM14_1395,STM14_0798 |
| seo_pathway.gmt | seo00660 | C5-Branched dibasic acid metabolism | 5 | 747 | 11 | 1446 | -0.36 | 1.00 | 1.00 | STM14_0140,STM14_0858,STM14_0857,STM14_3769,STM14_0132 |
| seo_pathway.gmt | seo00561 | Glycerolipid metabolism | 5 | 747 | 11 | 1446 | -0.36 | 1.00 | 1.00 | STM14_3567,STM14_3929,STM14_4939,STM14_5171,STM14_3883 |
| seo_pathway.gmt | seo00541 | O-Antigen nucleotide sugar biosynthesis | 10 | 747 | 22 | 1446 | -0.36 | 1.00 | 1.00 | STM14_4720,STM14_4719,STM14_2586,STM14_2590,STM14_4723,STM14_2578,STM14_2118,STM14_2577,STM14_2591,STM14_4718 |
| seo_pathway.gmt | seo00240 | Pyrimidine metabolism | 20 | 747 | 44 | 1446 | -0.37 | 1.00 | 1.00 | STM14_1200,STM14_3900,STM14_3530,STM14_5352,STM14_5287,STM14_0078,STM14_2617,STM14_4065,STM14_2477,STM14_4025,STM14_0462,STM14_2114,STM14_1373,STM14_2813,STM14_2694,STM14_0061,STM14_2812,STM14_3098,STM14_1332,STM14_3558 |
| seo_pathway.gmt | seo01240 | Biosynthesis of cofactors | 64 | 747 | 142 | 1446 | -0.42 | 1.00 | 1.00 | STM14_0185,STM14_3133,STM14_3978,STM14_0519,STM14_2918,STM14_1200,STM14_1590,STM14_2903,STM14_4735,STM14_932,STM14_4975,STM14_0054,STM14_0782,STM14_5352,STM14_3869,STM14_2520,STM14_0078,STM14_2927,STM14_4700,STM14_1105,STM14_2053,STM14_0496,STM14_3739,STM14_2844,STM14_4793,STM14_2848,STM14_2504,STM14_2118,STM14_2514,STM14_2517,STM14_2518,STM14_0632,STM14_2507,STM14_0241,STM14_4187,STM14_2810,STM14_2516,STM14_3158,STM14_931,STM14_2521,STM14_0920,STM14_3692,STM14_0576,STM14_0435,STM14_4492,STM14_934,STM14_4815,STM14_0751,STM14_2512,STM14_4751,STM14_2081,STM14_2969,STM14_4487,STM14_2515,STM14_0106,STM14_5008,STM14_0737,STM14_3098,STM14_3694,STM14_1332,STM14_2845,STM14_4918,STM14_3558,STM14_2843 |
| seo_pathway.gmt | seo00290 | Valine, leucine and isoleucine biosynthesis | 7 | 747 | 16 | 1446 | -0.46 | 1.00 | 1.00 | STM14_0140,STM14_4700,STM14_4702,STM14_3926,STM14_4706,STM14_4701,STM14_0132 |

| | | | | | | | | | | |
|---|---|---|---|---|---|---|---|---|---|---|
| seo_pathway.gmt | seo03010 | Ribosome | 24 | 747 | 55 | 1446 | -0.48 | 1.00 | 1.00 | STM14_4119,STM14_4121,STM14_5278,STM14_4036,STM14_1622,STM14_4136,STM14_2749,STM14_4120,STM14_0052,STM14_4130,STM14_4117,STM14_4141,STM14_4988,STM14_5275,STM14_4122,STM14_3966,STM14_4151,STM14_3991,STM14_0554,STM14_4138,STM14_4139,STM14_4989,STM14_4135,STM14_1621 |
| seo_pathway.gmt | seo00540 | Lipopolysaccharide biosynthesis | 17 | 747 | 40 | 1446 | -0.55 | 1.00 | 1.00 | STM14_0366,STM14_5165,STM14_0268,STM14_1612,STM14_4471,STM14_3876,STM14_4473,STM14_1117,STM14_1323,STM14_0270,STM14_4482,STM14_4475,STM14_0290,STM14_4472,STM14_4003,STM14_4476,STM14_2773 |
| seo_pathway.gmt | seo02010 | ABC transporters | 58 | 747 | 137 | 1446 | -0.60 | 1.00 | 1.00 | STM14_0771,STM14_0773,STM14_4652,STM14_2108,STM14_2027,STM14_2110,STM14_2106,STM14_2107,STM14_4294,STM14_2372,STM14_0544,STM14_2741,STM14_2898,STM14_4000,STM14_1043,STM14_991,STM14_964,STM14_4282,STM14_4370,STM14_4681,STM14_992,STM14_4372,STM14_4283,STM14_2900,STM14_2740,STM14_2669,STM14_5376,STM14_5232,STM14_2300,STM14_1392,STM14_3459,STM14_3998,STM14_1403,STM14_2508,STM14_4375,STM14_1042,STM14_1029,STM14_2667,STM14_4298,STM14_2999,STM14_5122,STM14_4371,STM14_2901,STM14_3392,STM14_3458,STM14_4280,STM14_962,STM14_4373,STM14_2668,STM14_963,STM14_2897,STM14_5375,STM14_3391,STM14_4297,STM14_4281,STM14_1030,STM14_1393,STM14_3997 |
| seo_pathway.gmt | seo01503 | Cationic antimicrobial peptide (CAMP) resistanc | 13 | 747 | 32 | 1446 | -0.66 | 1.00 | 1.00 | STM14_5165,STM14_0270,STM14_0249,STM14_2946,STM14_4881,STM14_1408,STM14_1409,STM14_4882,STM14_5163,STM14_0559,STM14_1389,STM14_4179,STM14_2842 |
| seo_pathway.gmt | seo00780 | Biotin metabolism | 4 | 747 | 10 | 1446 | -0.68 | 1.00 | 1.00 | STM14_4974,STM14_2927,STM14_2053,STM14_0920 |
| seo_pathway.gmt | seo00970 | Aminoacyl-tRNA biosynthesis | 10 | 747 | 25 | 1446 | -0.69 | 1.00 | 1.00 | STM14_4441,STM14_2969,STM14_4406,STM14_4193,STM14_0055,STM14_1624,STM14_5371,STM14_3093,STM14_2312,STM14_1619 |
| seo_pathway.gmt | seo01232 | Nucleotide metabolism | 20 | 747 | 50 | 1446 | -0.70 | 1.00 | 1.00 | STM14_3900,STM14_3530,STM14_4505,STM14_2617,STM14_4065,STM14_3076,STM14_2477,STM14_4025,STM14_0462,STM14_2114,STM14_1373,STM14_2813,STM14_5505,STM14_0169,STM14_0061,STM14_2812,STM14_3075,STM14_3098,STM14_3747,STM14_3558 |
| seo_pathway.gmt | seo00760 | Nicotinate and nicotinamide metabolism | 9 | 747 | 23 | 1446 | -0.74 | 1.00 | 1.00 | STM14_1590,STM14_3530,STM14_2830,STM14_5430,STM14_1787,STM14_3367,STM14_1842,STM14_3418,STM14_1788 |
| seo_pathway.gmt | seo00770 | Pantothenate and CoA biosynthesis | 9 | 747 | 24 | 1446 | -0.85 | 1.00 | 1.00 | STM14_4975,STM14_0140,STM14_4700,STM14_2694,STM14_3157,STM14_4492,STM14_4706,STM14_4487,STM14_4701 |
| seo_pathway.gmt | seo00450 | Selenocompound metabolism | 5 | 747 | 14 | 1446 | -0.95 | 1.00 | 1.00 | STM14_4441,STM14_5035,STM14_1667,STM14_1575,STM14_1809 |
| seo_pathway.gmt | seo00920 | Sulfur metabolism | 9 | 747 | 26 | 1446 | -1.03 | 1.00 | 1.00 | STM14_4459,STM14_3106,STM14_2555,STM14_5289,STM14_1090,STM14_5180,STM14_1089,STM14_4245,STM14_2999 |
| seo_pathway.gmt | seo00053 | Ascorbate and aldarate metabolism | 3 | 747 | 10 | 1446 | -1.33 | 1.00 | 1.00 | STM14_2888,STM14_3931,STM14_2887 |
| seo_pathway.gmt | seo04122 | Sulfur relay system | 4 | 747 | 15 | 1446 | -1.57 | 1.00 | 1.00 | STM14_932,STM14_3106,STM14_931,STM14_934 |
| seo_pathway.gmt | seo00562 | Inositol phosphate metabolism | 2 | 747 | 8 | 1446 | -1.69 | 1.00 | 1.00 | STM14_5326,STM14_4906 |
| seo_pathway.gmt | seo00400 | Phenylalanine, tyrosine and tryptophan biosynthe | 5 | 747 | 20 | 1446 | -1.70 | 1.00 | 1.00 | STM14_0459,STM14_4199,STM14_1635,STM14_5106,STM14_1531 |
| seo_pathway.gmt | seo03410 | Base excision repair | 3 | 747 | 13 | 1446 | -1.85 | 1.00 | 1.00 | STM14_2982,STM14_3674,STM14_2720 |
| seo_pathway.gmt | seo00040 | Pentose and glucuronate interconversions | 6 | 747 | 26 | 1446 | -1.86 | 1.00 | 1.00 | STM14_4905,STM14_4864,STM14_2118,STM14_3796,STM14_3590,STM14_0758 |
| seo_pathway.gmt | seo00790 | Folate biosynthesis | 5 | 747 | 22 | 1446 | -1.89 | 1.00 | 1.00 | STM14_3978,STM14_932,STM14_931,STM14_934,STM14_0106 |
| seo_pathway.gmt | seo00730 | Thiamine metabolism | 1 | 747 | 10 | 1446 | -3.28 | 1.00 | 1.00 | STM14_0496 |
