## Supplementary Table 4 for "Commensal Yeast Promotes *Salmonella* Typhimurium Virulence"

**Table S4: Real-time PCR analysis of genes encoding for host inflammatory response from the cecum tissue of Uninfected or Uninfected with L-arginine treated mice 24h and 48h p.i. Data is from 2 independent experiments.**

| Gene Name | Time p.i. | Fold difference over Uninfected | | p value using unpaired t-test |
|---|---|---|---|---|
| | | Uninfected | Uninfected + 2% L-arg supplementation | |
| *Il17a* | 24h | 0.82114962454146 | 0.49855069938965 | 0.9447 |
| | | 0.33809862903166 | 0.51811169305562 | |
| | | 6.77887843987691 | 5.47638870259456 | |
| | | 1.86353438481934 | | |
| | | 0.28512745324829 | | |
| | 48h | 4.05046925280069 | 2.927445524 | 0.3804 |
| | | 0.23509156777954 | 1.202939537 | |
| | | 0.47840988784182 | 0.734963206 | |
| | | 2.19511592710664 | 5.316804488 | |
| | | | 4.490032458 | |
| *Tnfa* | 24h | 2.80977745069939 | 3.327630859 | 0.3471 |
| | | 1.12143455554810 | 1.316233089 | |
| | | 6.14857213789811 | 0.45811643 | |
| | | 0.64368334106446 | 15.92119027 | |
| | | 0.03326678857252 | | |
| | | 2.41044220409411 | | |
| | 48h | 1.49697545013517 | 44.44762871 | 0.4925 |
| | | 0.36750018690873 | 1.087713819 | |
| | | 1.16877388773601 | 0.370903523 | |
| | | 1.55523942632196 | 0.331373172 | |
| | | | 0.047360402 | |
| | | | 0.206140101 | |
| *CxCl1* | 24h | 1.52642995018105 | 0.606081085 | 0.4549 |
| | | 0.33375252261578 | 0.915727473 | |
| | | 0.21949612610777 | 0.478942374 | |
| | | 16.01075559514790 | 1.039550798 | |
| | | 0.60686444541057 | | |
| | | 0.92038213960746 | | |
| | 48h | 1.19962103586722 | 33.74905761 | 0.342 |
| | | 1.27496944693282 | 2.510971891 | |
| | | 1.83328051972497 | 1.066151685 | |
| | | 0.35663768006615 | | |
| | | | 1.652378607 | |
| | | | 3.451938312 | |
| *Il22* | 24h | 2.91704584431274 | 2.278875559 | 0.4264 |
| | | 2.07859657938530 | 3.048416359 | |
| | | 1.08596824504133 | 1.47695412 | |
| | | 0.05185394547325 | 1.568799677 | |
| | | 2.10343232572970 | | |
| | | 1.39238411643653 | | |
| | 48h | 2.42094874947106 | 16.9954529 | 0.346 |
| | | 0.11397291139423 | 107.9268361 | |
| | | 2.86808724635132 | 0.132446493 | |
| | | 1.26363138411013 | | |
| | | | 0.222303181 | |
| | | | 2.422240565 | |
| *Ifng* | 24h | 9.51949583617939 | 15.95070635 | 0.9553 |
| | | 7.16201040752615 | 3.393869352 | |
| | | 19.98210835949190 | 0.225477488 | |
| | | 0.09479941230548 | | |
| | | 0.01995308600473 | | |
| | | 0.38805570689336 | | |
| | 48h | 2.88765169405820 | 8.537992326 | 0.5839 |
| | | 0.24027247663182 | 0.310818674 | |
| | | 0.54611020554304 | 0.188864532 | |
| | | 2.63919132269427 | | |
| *Il1b* | 24h | 0.85226738363744 | 0.9406178 | 0.6219 |
| | | 0.60633704948295 | 2.03866865 | |
| | | 0.79383152819254 | 0.437517161 | |
| | | 2.33446416597268 | 0.27534041 | |
| | | 0.52997637394839 | | |
| | | 1.97032542171406 | | |
| | 48h | 0.47893728213948 | 5.698839958 | 0.4604 |
| | | 0.37356076560059 | 0.429999334 | |
| | | 0.32340282157678 | 0.230796559 | |
| | | 17.28288711018700 | 0.896806306 | |
| | | | 0.622651721 | |
| *Il18* | 24h | 5.35702555114206 | 0.830692406 | 0.2697 |
| | | 4.09350675259841 | 0.246563911 | |
| | | 1.52681754951490 | 0.101982526 | |
| | | 0.31840603322364 | | |
| | | 0.35509166826361 | | |
| | | 0.26416292048854 | | |
| | 48h | 1.45818350504268 | 6.079607377 | 0.6671 |
| | | 0.13683194150006 | 0.359446156 | |
| | | 0.32926013784705 | 0.271944229 | |
| | | 15.22163045004800 | 23.13279977 | |
| | | | 9.937555458 | |
| | | | 0.432322613 | |
| *Ifnb* | 24h | 0.61711908421367 | 0.762863368 | 0.4373 |
| | | 0.69712433557998 | 0.487973512 | |
| | | 59.16149752960610 | 0.34190452 | |
| | | 0.35726975639810 | 0.114430219 | |
| | | 0.07519702921079 | | |
| | | 1.46246231383727 | | |
| | 48h | 0.09931074854149 | 1.220334982 | 0.3178 |
| | | 0.99484929220519 | 0.740545806 | |
| | | 12.04557379508250 | 0.296023373 | |
| | | 0.84027019054015 | 3.578531965 | |
| | | | 0.018534634 | |
| | | | 0.015842127 | |

| | | | | |
|---|---|---|---|---|
| *Ido1* | 24h | 0.66599295813364<br>0.65659201418844<br>0.97051295446244<br>1.18521476145245<br>0.54404445576126<br>3.65428141398280 | 1.081219394<br>0.740736176<br>0.702823639<br>1.570276849 | 0.6939 |
| | 48h | 0.89712796972543<br>1.47234617303828<br>0.96946534326029<br>0.78091431300104 | 0.09454107<br>0.46244109<br>0.42144856<br>0.256856733<br>1.24547509<br>1.572511371 | 0.3076 |
| *Ptges* | 24h | 1.64350777147798<br>0.65801790658416<br>1.24546150520367<br>0.75722777169753<br>0.60710916869614<br>1.61497904524025 | 0.779380346<br>0.38592384<br>0.507444439<br>1.210315752 | 0.2309 |
| | 48h | 1.31529178312070<br>0.80188965927771<br>0.76827550581533<br>1.23408846413616 | 0.658193877<br>1.06448569<br>0.883915569<br>1.864290041<br>1.233830705<br>2.724662142 | 0.3831 |
| *Tgfb* | 24h | 1.32344608051042<br>0.41210211883760<br>1.83353371613756 | 0.244022268<br>0.28369578<br>0.530634125 | 0.1205 |
| | 48h | 0.12127861233580<br>0.02935317434010<br>0.04658335156032 | 0.044877043<br>0.027966375<br>0.02432718 | 0.313 |
| *Ifnl2* | 24h | 44.58786461792360<br>99.99639864571520<br>0.16869611026753<br>0.19672805572323<br>0.04140671044183<br>0.16321379722531 | 0.502830166<br>0.269662687<br>0.1523402<br>0.127711026 | 0.2877 |
| | 48h | 32.18993524086030<br>0.17845040790217<br>0.13344965714990<br>1.30450242164940 | 5.152491207<br>0.462445828<br>4.666580737<br>19.60513895<br>0.746364311<br>0.086791764 | 0.661 |
| *Nos2* | 24h | 0.87813822184201<br>0.49774235630734<br>0.28933140788479<br>5.21058662581581<br>0.68432518154441<br>2.21762341189437 | 2.562726089<br>0.966838158<br>1.051433129<br>2.078621083 | 0.973 |
| | 48h | 0.25971856485148<br>3.06894395597814<br>2.53963712630610<br>0.49401072860985 | 0.52555841<br>0.977185343<br>6.919717171<br>9.816324514<br>1.152340214<br>4.109587518 | 0.2809 |
| *Il10* | 24h | 0.88155833826630<br>0.75597483491797<br>0.45865611653627<br>1.09767955636275<br>1.58563670986512<br>1.87964202502003 | 0.243319121<br>0.300408951<br>1.204775102<br>0.475827011 | 0.1254 |
| | 48h | 0.85934103698657<br>1.46465822744944<br>0.51745711237859<br>1.53540797579839 | 0.238450072<br>0.591244311<br>0.321737867<br>1.98703992 | 0.816 |
| *Lcn2* | 24h | 8.77589650094587<br>5.91872455545535<br>9.70508309246834<br>1.30007602252470<br>0.00370829869548<br>0.41146951332894 | 7.586913528<br>3.535607742<br>3.386963129<br>0.38556562 | 0.8083 |
| | 48h | 1.49210710068874<br>0.38544606032151<br>0.37774591823924<br>4.60295375441075 | 1.192445965<br>0.653840683<br>0.542923078<br>5.535308375<br>0.08446477<br>0.233769044 | 0.8028 |
| *Reg3g* | 24h | 3.72954432518003<br>1.20499399888449<br>1.78807789002112<br>7.49786079462886<br>0.29022630166293<br>0.05718719156918 | 3.87505076<br>1.063961706<br>1.808294526<br>0.384453813 | 0.6889 |
| | 48h | 0.41449042905185<br>0.67514222087327<br>1.30572494854129<br>2.73677140166880 | 4.290158272<br>0.260250606<br>0.233246552<br>0.504313436<br>0.193710463<br>0.954050182 | 0.8238 |
| | | 0.93735166280945<br>0.69703394298674 | 0.376245646<br>28.65431375 | |

| | | | | |
|---|---|---|---|---|
| *S100a8* | 24h | 0.77090987149450<br>3.86368946694579<br>0.34694838188680<br>1.48106060955810 | 0.130672385<br>1.242744168 | 0.2959 |
| | 48h | 2.05583991176243<br>0.85347974683837<br>0.87092524339155<br>0.65438993143007 | 0.893687718<br>1.340833219<br>1.15093414<br>24.01572901<br>4.062431248<br>8.227769378 | 0.2641 |
| *S100a9* | 24h | 1.07427713324324<br>0.80608361442806<br>1.16454236142309<br>2.63370212357790<br>0.56395502292472<br>0.66763202564525 | 1.224498384<br>31.25243997<br>0.47810752<br>1.906492583 | 0.2398 |
| | 48h | 2.09293282669222<br>0.53871050081008<br>1.14874604201937<br>0.77208522399127 | 15.78259748<br>1.783896<br>1.215466687<br>7.542794621<br>2.473362557<br>2.639763614 | 0.1937 |
